# Artistoo: build, share, and explore simulations of cells and tissues in the web browser

**DOI:** 10.1101/2020.05.01.072975

**Authors:** Inge M. N. Wortel, Johannes Textor

## Abstract

**Summary:** The Cellular Potts Model (CPM) is a powerful *in silico* method for simulating diverse biological processes at tissue scale. Because of its inherently graphical nature, this model should in theory be accessible to a large audience of biologists – without requiring extensive mathematical expertise. But in practice, CPMs are mostly implemented in specialized frameworks that users need to master before they can run and modify the simulation. We here present Artistoo (Artificial Tissue Toolbox), a JavaScript library for building “explorable” CPM simulations where users can change model parameters and see their effects in real time. Artistoo simulations run directly in the web browser and do not require any third-party software, plugins, or back-end servers. Although implemented in JavaScript, Artistoo does not suffer from a major performance loss compared to frameworks written in C++; it remains sufficiently fast to let users interact with simulations in real time. Artistoo provides an opportunity to unlock CPM models for a broader audience: interactive simulations can be shared through a simple URL in a *zero-install* setting. We discuss how such model sharing may benefit modelling research, science dissemination, open science, and education.

**Availability and Implementation:** Artistoo is an open-source library released under the MIT license, and is freely available on GitHub at https://github.com/ingewortel/artistoo.

## 1 Introduction

A growing community of computational biologists uses simulation models to reason about complex processes in biological systems. The Cellular Potts Model (CPM) is a well-established framework for simulating interacting cells. Originally proposed as a model for cell sorting (Graner and Glazier, 1992), the CPM has since been extended with a plethora of biological processes such as proliferation, apoptosis, cell motion, and chemotaxis – allowing CPM users to model diverse phenomena ranging from slime mould formation to blood vessel development, tumor growth, and cell migration (Marée *et al*., 2007; Szabó *et al*., 2013; Hirashima *et al*., 2017).

Nowadays, several mature professional modelling frameworks with CPM implementations exist, such as CompuCell3D (Swat *et al*., 2012), Morpheus (Starruß *et al*., 2014), the Tissue Simulation Toolkit (Daub and Merks, 2014), and CHASTE (Mirams *et al*., 2013). Although CPMs are relatively efficient models, tissue-scale simulations still require substantial computational resources. For this reason, all of the abovementioned frameworks rely on the C++ programming language for computation steps, which requires them to be built for and installed on the user’s native operating system.

Here, we present “Artistoo” (Artificial Tissue Toolbox), a CPM framework built entirely in JavaScript. Although interpreted languages like JavaScript have classically been deemed too inefficient for running simulations, we found that this no longer holds: investments by major tech companies have tremendously improved JavaScript engines over the past years, to the point that our CPM now has no major performance disadvantage compared to existing C++ frameworks.

The JavaScript implementation of Artistoo opens up new possibilities for rapid and low-barrier sharing of CPM simulations with students, collaborators, and readers or reviewers of a paper. Unlike existing frameworks, Artistoo allows users to view simulations from the web browser without needing to install any software: Artistoo models run on any platform providing a standards-compliant web browser – be it a desktop computer, a tablet, or a mobile phone. These simulations can be published on any web server or saved locally and do not rely on any back-end servers being available. They can be made explorable, enabling visitors to interact with the simulation and see the effect of changing model parameters in real-time.

In this paper, we will first briefly explain the key design principles behind Artistoo. We will then highlight applications in teaching, research, science dissemination, and open science where we envision that the zero-install, web-based architecture of our framework could be particularly useful.

## 2 Implementation

Artistoo is a JavaScript library implemented as an ECMAScript 6 module, which can be loaded into an HTML page or accessed from within a nodejs command line application.

### 2.1 Approachability

The methods currently implemented in the framework allow users to simulate, visualize, and analyze a wide range of CPM models (Figure 1A). Our Github repository contains example code for models of various biological processes (e.g. simulations of tissues, cell migration, and cell interactions). First-time users can download these HTML pages and modify parameters without needing to learn the implementation details of the framework, or to have programmed in JavaScript before. Alternatively, the `Simulation` class provides default methods for setting up and visualizing simulations, allowing users to get started with the library without having to set up this “boilerplate” code themselves. Advanced users can instead build simulations from scratch and customize them using the many available options and methods, or even plug in their own code modules (see section 2.2). An example interactive HTML simulation (Figure 1B) is included in the Supplementary Materials. Full documentation as well as a user manual with step-by-step tutorials are available at https://artistoo.net/.

**Figure 1:**
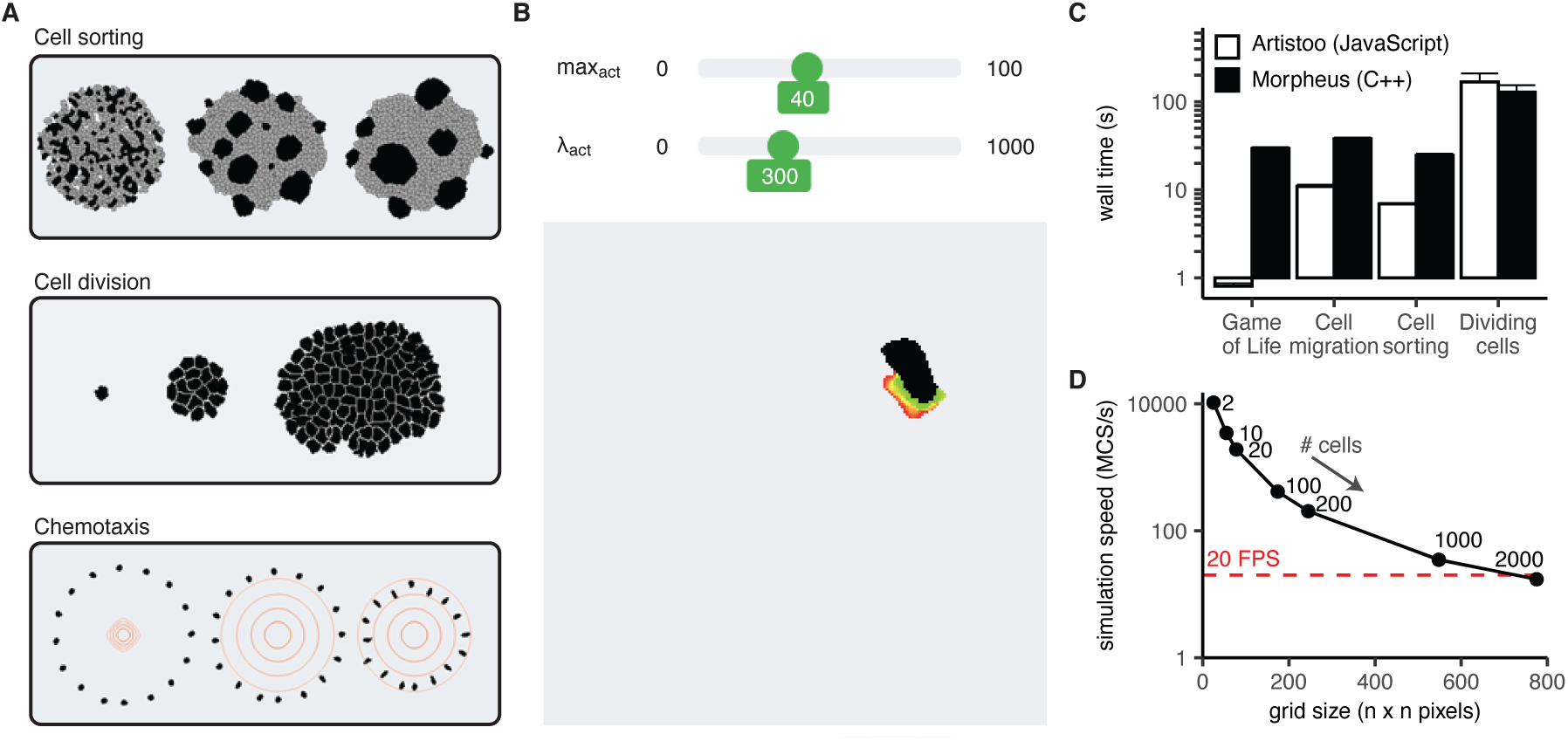
(a) Artistoo supports simulation of diverse biological processes; (b) users can interact with browser-based simulations via sliders, in real-time. (c) Artistoo performance is comparable to that of the Morpheus framework. Data show wall times (mean+SD of 5 runs) for four CPM models implemented in both frameworks (see Supplementary Methods for implementation details, and Supplementary Materials for interactive browser versions). (d) Scalability of the cell sorting simulation; simulation speed in Monte Carlo Steps per second (MCS/s) for different grid sizes (mean ± SD of 5 runs). Red line indicates 20 frames per second, a minimum speed required for a “real-time” simulation for the human visual system.

### 2.2 Modularity and flexibility

A typical CPM simulation consists of different types of components: the grid on which cells are simulated, the energy rules governing cell behavior in the model, separate processes such as cell proliferation or diffusion, and the visualisation and quantification methods used to produce outputs. A key strength of the CPM is that it can be easily extended with custom terms to model specific processes. To facilitate such customization, we have set up the code in a highly modular fashion. These modules can be combined freely to build a custom simulation. In addition, users can supply their own custom modules – containing any of the aforementioned simulation components – to integrate with the framework and to share with other users.

### 2.3 Performance and scalability

Although maximal performance is not a design goal of our framework *per se*, Artistoo should not be much slower than comparable frameworks either: running explorable simulations in real-time is only feasible if computations are reasonably efficient. Indeed, we implemented various simulations both in Artistoo and in Morpheus and found that both frameworks had similar performance (Figure 1C). Even in cases where Artistoo is slower (e.g. in the simulation of cell division), the difference in performance is not so large that real-time browser simulations become infeasible. Simulation speed does decrease for very large systems, but real-time simulations remain feasible for a reasonable range of grid sizes (Figure 1D). This would allow sharing of at least a reasonable prototype of larger-scale models.

## 3 Applications

We here highlight a number of settings where Artistoo might complement other available modelling frameworks, focusing on the unique feature of Artistoo: it allows users to build and share explorable simulations in a *zero-install* setting. We discuss how this opens up novel opportunities of sharing CPM-based research and provide examples from our own work.

### 3.1 Teaching

When organizing practical computer work in the context of classroom teaching, getting software to work on every student’s computer can consume a substantial amount of time and effort. Especially when teaching large classes in limited time, installing an entirely new modelling framework for a single course assignment may not be appropriate. The *zero-install* feature of Artistoo might therefore be attractive for use of CPM modelling in the classroom. We frequently use the framework in teaching and found it feasible to let students run and understand CPM models in a workshop of just a few hours – even when students had no programming experience and were given just a single lecture on the CPM in advance. We provide an introductory assignment on the CPM in the Supplementary Materials, which readers may use and adapt freely for their own courses.

### 3.2 Communication and Open Science

While the move towards open science has prompted many to share their code with publications, understanding and using this code often remains challenging for readers who do not use similar models themselves. We envision that by sharing interactive Artistoo simulations via a simple URL, computational biologists can make their modelling research more accessible for the readers and reviewers of their papers; if readers can interact with model parameters themselves without the barrier of having to install special software, this may greatly improve the transparency of CPM modelling research. This would allow others to evaluate these models more critically, as well as foster the exchange of ideas between scientists from different disciplines.

In addition, interactive simulations can help communicate CPM-based science at conferences or in classrooms. We frequently use the framework `reveal.js` (El Hattab *et al*., 2020) to build slideshows in HTML, in which live, interactive Artistoo demonstrations help explain how models work. Similarly, interactive simulations can be shared on a conference poster via a QR code, which other attendees can explore on their mobile phone. We provide examples of both in the Supplementary Materials.

### 3.3 Research and Collaboration

Although the CPM is extremely flexible in the types of behaviors it can model, it can be difficult to find the parameter ranges where these behaviors occur. We found that an interactive web page with instantaneous feedback where the effect of changing parameters is visible in real-time (Figure 1 and Supplementary Materials) can substantially speed up parameter selection. This visual approach also picks up on unpredicted behaviors and artefacts (e.g. cell breaking) that are difficult to detect from numerical outputs alone. Moreover, we note that sharing these interactive pages allows us to tune parameters in collaboration with experimental biologists, helping us improve our models at an early stage. Thus, building a web-based prototype of a simulation can speed up parameter tuning and help obtain higher quality models.

## 4 Conclusion

Artistoo allows users to build highly customizable simulations, and its highly modular structure makes it easy for future users to extend the framework with custom code. Its performance is comparable to that of existing frameworks. We do not envision Artistoo to *replace* existing modelling software; rather, it can complement existing software directed at computational biologists and developers by letting users build explorable and sharable versions of a simulation. Indeed, to the best of our knowledge, Artistoo is the first simulation framework supporting interactive simulations in the web browser that can be shared via a simple URL. We hope that this will unlock the use of (CPM) simulations to a much larger audience.

## Acknowledgements

The authors thank Nino van Halem and Ankur Ankan for their contributions to the code, and Peter Linders for valuable feedback on an early version of the Artistoo manual.

## Supplementary Methods

This section contains implementation details of the simulations used to assess Artistoo (v1.0.0) performance. All simulations were run in the console mode (using nodejs, which contains the same JavaScript engine as the Chrome web browser).

### Framework comparisons

To compare performance of Artistoo versus that of Morpheus, we performed 4 different simulations in both frameworks. For this, we used the default examples provided with Morpheus, and rebuilt similar simulations in Artistoo. All code is available in the Supplementary Materials, as are interactive HTML versions of each simulation. We refer to the provided code for details of the implementation, but summarise the most important settings here.

#### Game of Life

This is an implementation of the Game of Life, a Cellular Automaton (CA) of John Conway. The simulation was run on a 50 x 50 pixel grid with random initial conditions. The simulation was run for 500 steps, storing a PNG image every 20 steps.

#### Protrusion model

This model of a migrating cell implements an actin-inspired migration model (Niculescu *et al*., 2015). A single cell was seeded in the middle of a 200 x 200 pixel grid. Two obstacles of radius 10 were placed at a distance of 50 pixels to the left and right of the cell, respectively. Simulations were run for 15,000 MCS, logging the cell’s centroid every 10 MCS and saving a PNG every 250 MCS.

#### Cell sorting

This simulation implements the classical CPM model published by Graner and Glazier (Graner and Glazier, 1992). 50 cells each of two cell types were seeded on a 200 x 200 pixel grid within a circle of of radius 67 from the grid midpoint. Simulations were run for 2000 MCS, logging statistics every 10 MCS and saving a PNG every 100 MCS.

#### Cell division

Cell division was simulated on a 500 x 500 pixel grid. The grid was initialised with 20 cells in a circle of radius 35 surrounding the grid midpoint. Simulations were run for 40,000 MCS, logging the number of cells every 100 MCS and saving a PNG every 1000 MCS.

### Scalability of cell sorting

For the scalability simulations, simulations were run without outputting images. This allowed us to investigate the simulation speed separately from the time it takes to draw the entire grid. Note that if the drawing step becomes a limiting factor for running the simulation, it is always possible to speed up the process by drawing only once every few steps, or by choosing a more efficient drawing method (e.g. drawing only cell borders rather than entire cells).

Simulations contained 1, 5, 10, 50, 100, 500, or 1000 cells per cell type. The grid dimensions were adaptively scaled such that 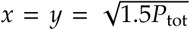, with *P*_tot_ the total number of pixels of all the cells. Cells were seeded within a radius 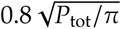 from the grid midpoint. Other settings were the same as in the cell sorting simulation described under *Framework comparisons*.

