## Supplementary material for "Artistoo: build, share, and explore simulations of cells and tissues in the web browser": ref.bib

@article{niculescu_crawling_2015,
	title = {Crawling and {Gliding}: {A} {Computational} {Model} for {Shape}-{Driven} {Cell} {Migration}},
	volume = {11},
	url = {https://doi.org/10.1371/journal.pcbi.1004280},
	doi = {10.1371/journal.pcbi.1004280},
	number = {10},
	journal = {PLOS Computational Biology},
	author = {Niculescu, Ioana and Textor, Johannes and de Boer, Rob J.},
	year = {2015},
	keywords = {2018-cpm, CPM, in silico migration model},
	pages = {1--22}
}
