## Supplementary material for "Artistoo: build, share, and explore simulations of cells and tissues in the web browser": index.html

2020-05-01 Example slides

  

##### Artistoo:

s
  

**Inge Wortel**   
   
 Department of Tumor Immunology, Radboudumc, Nijmegen, the Netherlands

### The Cellular Potts Model (CPM)

Pixels belong to cells, which   
move by copying pixels:

Copy **success** chance (P_copy_) is higher when it helps the cell:

|  |  |  |
| --- | --- | --- |
| Stay together: | Maintain its size: | Maintain its membrane: |
| $\searrow$ | $\downarrow$ | $\swarrow$ |

### Migration: Act model

Cells move if we add **positive feedback**
on protrusive **activity** ($\approx$ actin polymerization)^1^:

|  |  |  |
| --- | --- | --- |
| Parameters: |  |  |
| **λ_act_** | $\approx$ | protrusive **force** |
| **max_act_** | $\approx$ | polymerized actin **lifetime** |

^1^Niculescu et al. PLoS Computational Biology, 2015.

### See also:

https://revealjs.com/#/
