## Supplementary material for "Artistoo: build, share, and explore simulations of cells and tissues in the web browser": TeX-AMS_HTML-full.js

```
/*
 *  /MathJax/config/TeX-AMS_HTML-full.js
 *  
 *  Copyright (c) 2010-2016 The MathJax Consortium
 *
 *  Part of the MathJax library.
 *  See http://www.mathjax.org for details.
 * 
 *  Licensed under the Apache License, Version 2.0;
 *  you may not use this file except in compliance with the License.
 *
 *      http://www.apache.org/licenses/LICENSE-2.0
 */

MathJax.Ajax.Preloading(
  "[MathJax]/jax/input/TeX/config.js",
  "[MathJax]/jax/output/HTML-CSS/config.js",
  "[MathJax]/jax/output/PreviewHTML/config.js",
  "[MathJax]/extensions/tex2jax.js",
  "[MathJax]/extensions/MathEvents.js",
  "[MathJax]/extensions/MathZoom.js",
  "[MathJax]/extensions/MathMenu.js",
  "[MathJax]/jax/element/mml/jax.js",
  "[MathJax]/extensions/toMathML.js",
  "[MathJax]/extensions/TeX/noErrors.js",
  "[MathJax]/extensions/TeX/noUndefined.js",
  "[MathJax]/jax/input/TeX/jax.js",
  "[MathJax]/extensions/TeX/AMSmath.js",
  "[MathJax]/extensions/TeX/AMSsymbols.js",
  "[MathJax]/jax/output/HTML-CSS/jax.js",
  "[MathJax]/jax/output/HTML-CSS/autoload/mtable.js",
  "[MathJax]/jax/output/PreviewHTML/jax.js",
  "[MathJax]/extensions/fast-preview.js",
  "[MathJax]/extensions/AssistiveMML.js"
);

MathJax.Hub.Config({
  extensions: ['[Contrib]/a11y/accessibility-menu.js']
});

MathJax.InputJax.TeX=MathJax.InputJax({id:"TeX",version:"2.7.0",directory:MathJax.InputJax.directory+"/TeX",extensionDir:MathJax.InputJax.extensionDir+"/TeX",config:{TagSide:"right",TagIndent:"0.8em",MultLineWidth:"85%",equationNumbers:{autoNumber:"none",formatNumber:function(a){return a},formatTag:function(a){return"("+a+")"},formatID:function(a){return"mjx-eqn-"+String(a).replace(/[:"'<>&]/g,"")},formatURL:function(b,a){return a+"#"+escape(b)},useLabelIds:true}},resetEquationNumbers:function(){}});MathJax.InputJax.TeX.Register("math/tex");MathJax.InputJax.TeX.loadComplete("config.js");
MathJax.OutputJax["HTML-CSS"]=MathJax.OutputJax({id:"HTML-CSS",version:"2.7.0",directory:MathJax.OutputJax.directory+"/HTML-CSS",extensionDir:MathJax.OutputJax.extensionDir+"/HTML-CSS",autoloadDir:MathJax.OutputJax.directory+"/HTML-CSS/autoload",fontDir:MathJax.OutputJax.directory+"/HTML-CSS/fonts",webfontDir:MathJax.OutputJax.fontDir+"/HTML-CSS",config:{noReflows:true,matchFontHeight:true,scale:100,minScaleAdjust:50,availableFonts:["STIX","TeX"],preferredFont:"TeX",webFont:"TeX",imageFont:"TeX",undefinedFamily:"STIXGeneral,'Arial Unicode MS',serif",mtextFontInherit:false,EqnChunk:(MathJax.Hub.Browser.isMobile?10:50),EqnChunkFactor:1.5,EqnChunkDelay:100,linebreaks:{automatic:false,width:"container"},styles:{".MathJax_Display":{"text-align":"center",margin:"1em 0em"},".MathJax .merror":{"background-color":"#FFFF88",color:"#CC0000",border:"1px solid #CC0000",padding:"1px 3px","font-style":"normal","font-size":"90%"},".MathJax .MJX-monospace":{"font-family":"monospace"},".MathJax .MJX-sans-serif":{"font-family":"sans-serif"},"#MathJax_Tooltip":{"background-color":"InfoBackground",color:"InfoText",border:"1px solid black","box-shadow":"2px 2px 5px #AAAAAA","-webkit-box-shadow":"2px 2px 5px #AAAAAA","-moz-box-shadow":"2px 2px 5px #AAAAAA","-khtml-box-shadow":"2px 2px 5px #AAAAAA",filter:"progid:DXImageTransform.Microsoft.dropshadow(OffX=2, OffY=2, Color='gray', Positive='true')",padding:"3px 4px","z-index":401}}}});if(MathJax.Hub.Browser.isMSIE&&document.documentMode>=9){delete MathJax.OutputJax["HTML-CSS"].config.styles["#MathJax_Tooltip"].filter}if(!MathJax.Hub.config.delayJaxRegistration){MathJax.OutputJax["HTML-CSS"].Register("jax/mml")}MathJax.Hub.Register.StartupHook("End Config",[function(b,c){var a=b.Insert({minBrowserVersion:{Firefox:3,Opera:9.52,MSIE:6,Chrome:0.3,Safari:2,Konqueror:4},inlineMathDelimiters:["$","$"],displayMathDelimiters:["$$","$$"],multilineDisplay:true,minBrowserTranslate:function(f){var e=b.getJaxFor(f),k=["[Math]"],j;var h=document.createElement("span",{className:"MathJax_Preview"});if(e.inputJax==="TeX"){if(e.root.Get("displaystyle")){j=a.displayMathDelimiters;k=[j[0]+e.originalText+j[1]];if(a.multilineDisplay){k=k[0].split(/\n/)}}else{j=a.inlineMathDelimiters;k=[j[0]+e.originalText.replace(/^\s+/,"").replace(/\s+$/,"")+j[1]]}}for(var g=0,d=k.length;g<d;g++){h.appendChild(document.createTextNode(k[g]));if(g<d-1){h.appendChild(document.createElement("br"))}}f.parentNode.insertBefore(h,f)}},(b.config["HTML-CSS"]||{}));if(b.Browser.version!=="0.0"&&!b.Browser.versionAtLeast(a.minBrowserVersion[b.Browser]||0)){c.Translate=a.minBrowserTranslate;b.Config({showProcessingMessages:false});MathJax.Message.Set(["MathJaxNotSupported","Your browser does not support MathJax"],null,4000);b.Startup.signal.Post("MathJax not supported")}},MathJax.Hub,MathJax.OutputJax["HTML-CSS"]]);MathJax.OutputJax["HTML-CSS"].loadComplete("config.js");
MathJax.OutputJax.PreviewHTML=MathJax.OutputJax({id:"PreviewHTML",version:"2.7.0",directory:MathJax.OutputJax.directory+"/PreviewHTML",extensionDir:MathJax.OutputJax.extensionDir+"/PreviewHTML",noFastPreview:true,config:{scale:100,minScaleAdjust:50,mtextFontInherit:false,linebreaks:{automatic:false,width:"container"}}});if(!MathJax.Hub.config.delayJaxRegistration){MathJax.OutputJax.PreviewHTML.Register("jax/mml")}MathJax.OutputJax.PreviewHTML.loadComplete("config.js");
MathJax.Extension.tex2jax={version:"2.7.0",config:{inlineMath:[["\\(","\\)"]],displayMath:[["$$","$$"],["\\[","\\]"]],balanceBraces:true,skipTags:["script","noscript","style","textarea","pre","code","annotation","annotation-xml"],ignoreClass:"tex2jax_ignore",processClass:"tex2jax_process",processEscapes:false,processEnvironments:true,processRefs:true,preview:"TeX"},ignoreTags:{br:(MathJax.Hub.Browser.isMSIE&&document.documentMode<9?"\n":" "),wbr:"","#comment":""},PreProcess:function(a){if(!this.configured){this.config=MathJax.Hub.CombineConfig("tex2jax",this.config);if(this.config.Augment){MathJax.Hub.Insert(this,this.config.Augment)}if(typeof(this.config.previewTeX)!=="undefined"&&!this.config.previewTeX){this.config.preview="none"}this.configured=true}if(typeof(a)==="string"){a=document.getElementById(a)}if(!a){a=document.body}if(this.createPatterns()){this.scanElement(a,a.nextSibling)}},createPatterns:function(){var d=[],e=[],c,a,b=this.config;this.match={};for(c=0,a=b.inlineMath.length;c<a;c++){d.push(this.patternQuote(b.inlineMath[c][0]));this.match[b.inlineMath[c][0]]={mode:"",end:b.inlineMath[c][1],pattern:this.endPattern(b.inlineMath[c][1])}}for(c=0,a=b.displayMath.length;c<a;c++){d.push(this.patternQuote(b.displayMath[c][0]));this.match[b.displayMath[c][0]]={mode:"; mode=display",end:b.displayMath[c][1],pattern:this.endPattern(b.displayMath[c][1])}}if(d.length){e.push(d.sort(this.sortLength).join("|"))}if(b.processEnvironments){e.push("\\\\begin\\{([^}]*)\\}")}if(b.processEscapes){e.push("\\\\*\\\\\\$")}if(b.processRefs){e.push("\\\\(eq)?ref\\{[^}]*\\}")}this.start=new RegExp(e.join("|"),"g");this.skipTags=new RegExp("^("+b.skipTags.join("|")+")$","i");var f=[];if(MathJax.Hub.config.preRemoveClass){f.push(MathJax.Hub.config.preRemoveClass)}if(b.ignoreClass){f.push(b.ignoreClass)}this.ignoreClass=(f.length?new RegExp("(^| )("+f.join("|")+")( |$)"):/^$/);this.processClass=new RegExp("(^| )("+b.processClass+")( |$)");return(e.length>0)},patternQuote:function(a){return a.replace(/([\^$(){}+*?\-|\[\]\:\\])/g,"\\$1")},endPattern:function(a){return new RegExp(this.patternQuote(a)+"|\\\\.|[{}]","g")},sortLength:function(d,c){if(d.length!==c.length){return c.length-d.length}return(d==c?0:(d<c?-1:1))},scanElement:function(c,b,g){var a,e,d,f;while(c&&c!=b){if(c.nodeName.toLowerCase()==="#text"){if(!g){c=this.scanText(c)}}else{a=(typeof(c.className)==="undefined"?"":c.className);e=(typeof(c.tagName)==="undefined"?"":c.tagName);if(typeof(a)!=="string"){a=String(a)}f=this.processClass.exec(a);if(c.firstChild&&!a.match(/(^| )MathJax/)&&(f||!this.skipTags.exec(e))){d=(g||this.ignoreClass.exec(a))&&!f;this.scanElement(c.firstChild,b,d)}}if(c){c=c.nextSibling}}},scanText:function(b){if(b.nodeValue.replace(/\s+/,"")==""){return b}var a,c;this.search={start:true};this.pattern=this.start;while(b){this.pattern.lastIndex=0;while(b&&b.nodeName.toLowerCase()==="#text"&&(a=this.pattern.exec(b.nodeValue))){if(this.search.start){b=this.startMatch(a,b)}else{b=this.endMatch(a,b)}}if(this.search.matched){b=this.encloseMath(b)}if(b){do{c=b;b=b.nextSibling}while(b&&this.ignoreTags[b.nodeName.toLowerCase()]!=null);if(!b||b.nodeName!=="#text"){return(this.search.close?this.prevEndMatch():c)}}}return b},startMatch:function(a,b){var f=this.match[a[0]];if(f!=null){this.search={end:f.end,mode:f.mode,pcount:0,open:b,olen:a[0].length,opos:this.pattern.lastIndex-a[0].length};this.switchPattern(f.pattern)}else{if(a[0].substr(0,6)==="\\begin"){this.search={end:"\\end{"+a[1]+"}",mode:"; mode=display",pcount:0,open:b,olen:0,opos:this.pattern.lastIndex-a[0].length,isBeginEnd:true};this.switchPattern(this.endPattern(this.search.end))}else{if(a[0].substr(0,4)==="\\ref"||a[0].substr(0,6)==="\\eqref"){this.search={mode:"",end:"",open:b,pcount:0,olen:0,opos:this.pattern.lastIndex-a[0].length};return this.endMatch([""],b)}else{var d=a[0].substr(0,a[0].length-1),g,c;if(d.length%2===0){c=[d.replace(/\\\\/g,"\\")];g=1}else{c=[d.substr(1).replace(/\\\\/g,"\\"),"$"];g=0}c=MathJax.HTML.Element("span",null,c);var e=MathJax.HTML.TextNode(b.nodeValue.substr(0,a.index));b.nodeValue=b.nodeValue.substr(a.index+a[0].length-g);b.parentNode.insertBefore(c,b);b.parentNode.insertBefore(e,c);this.pattern.lastIndex=g}}}return b},endMatch:function(a,c){var b=this.search;if(a[0]==b.end){if(!b.close||b.pcount===0){b.close=c;b.cpos=this.pattern.lastIndex;b.clen=(b.isBeginEnd?0:a[0].length)}if(b.pcount===0){b.matched=true;c=this.encloseMath(c);this.switchPattern(this.start)}}else{if(a[0]==="{"){b.pcount++}else{if(a[0]==="}"&&b.pcount){b.pcount--}}}return c},prevEndMatch:function(){this.search.matched=true;var a=this.encloseMath(this.search.close);this.switchPattern(this.start);return a},switchPattern:function(a){a.lastIndex=this.pattern.lastIndex;this.pattern=a;this.search.start=(a===this.start)},encloseMath:function(b){var a=this.search,g=a.close,f,d,c;if(a.cpos===g.length){g=g.nextSibling}else{g=g.splitText(a.cpos)}if(!g){f=g=MathJax.HTML.addText(a.close.parentNode,"")}a.close=g;d=(a.opos?a.open.splitText(a.opos):a.open);while((c=d.nextSibling)&&c!==g){if(c.nodeValue!==null){if(c.nodeName==="#comment"){d.nodeValue+=c.nodeValue.replace(/^\[CDATA\[((.|\n|\r)*)\]\]$/,"$1")}else{d.nodeValue+=c.nodeValue}}else{var h=this.ignoreTags[c.nodeName.toLowerCase()];d.nodeValue+=(h==null?" ":h)}d.parentNode.removeChild(c)}var e=d.nodeValue.substr(a.olen,d.nodeValue.length-a.olen-a.clen);d.parentNode.removeChild(d);if(this.config.preview!=="none"){this.createPreview(a.mode,e)}d=this.createMathTag(a.mode,e);this.search={};this.pattern.lastIndex=0;if(f){f.parentNode.removeChild(f)}return d},insertNode:function(b){var a=this.search;a.close.parentNode.insertBefore(b,a.close)},createPreview:function(d,a){var b=MathJax.Hub.config.preRemoveClass;var c=this.config.preview;if(c==="none"){return}if((this.search.close.previousSibling||{}).className===b){return}if(c==="TeX"){c=[this.filterPreview(a)]}if(c){c=MathJax.HTML.Element("span",{className:b},c);this.insertNode(c)}},createMathTag:function(c,b){var a=document.createElement("script");a.type="math/tex"+c;MathJax.HTML.setScript(a,b);this.insertNode(a);return a},filterPreview:function(a){return a}};MathJax.Hub.Register.PreProcessor(["PreProcess",MathJax.Extension.tex2jax]);MathJax.Ajax.loadComplete("[MathJax]/extensions/tex2jax.js");
(function(d,h,l,g,m,b,j){var p="2.7.0";var i=MathJax.Extension;var c=i.MathEvents={version:p};var k=d.config.menuSettings;var o={hover:500,frame:{x:3.5,y:5,bwidth:1,bcolor:"#A6D",hwidth:"15px",hcolor:"#83A"},button:{x:-6,y:-3,wx:-2},fadeinInc:0.2,fadeoutInc:0.05,fadeDelay:50,fadeoutStart:400,fadeoutDelay:15*1000,styles:{".MathJax_Hover_Frame":{"border-radius":".25em","-webkit-border-radius":".25em","-moz-border-radius":".25em","-khtml-border-radius":".25em","box-shadow":"0px 0px 15px #83A","-webkit-box-shadow":"0px 0px 15px #83A","-moz-box-shadow":"0px 0px 15px #83A","-khtml-box-shadow":"0px 0px 15px #83A",border:"1px solid #A6D ! important",display:"inline-block",position:"absolute"},".MathJax_Menu_Button .MathJax_Hover_Arrow":{position:"absolute",cursor:"pointer",display:"inline-block",border:"2px solid #AAA","border-radius":"4px","-webkit-border-radius":"4px","-moz-border-radius":"4px","-khtml-border-radius":"4px","font-family":"'Courier New',Courier","font-size":"9px",color:"#F0F0F0"},".MathJax_Menu_Button .MathJax_Hover_Arrow span":{display:"block","background-color":"#AAA",border:"1px solid","border-radius":"3px","line-height":0,padding:"4px"},".MathJax_Hover_Arrow:hover":{color:"white!important",border:"2px solid #CCC!important"},".MathJax_Hover_Arrow:hover span":{"background-color":"#CCC!important"}}};var n=c.Event={LEFTBUTTON:0,RIGHTBUTTON:2,MENUKEY:"altKey",KEY:{RETURN:13,ESCAPE:27,SPACE:32,LEFT:37,UP:38,RIGHT:39,DOWN:40},Mousedown:function(q){return n.Handler(q,"Mousedown",this)},Mouseup:function(q){return n.Handler(q,"Mouseup",this)},Mousemove:function(q){return n.Handler(q,"Mousemove",this)},Mouseover:function(q){return n.Handler(q,"Mouseover",this)},Mouseout:function(q){return n.Handler(q,"Mouseout",this)},Click:function(q){return n.Handler(q,"Click",this)},DblClick:function(q){return n.Handler(q,"DblClick",this)},Menu:function(q){return n.Handler(q,"ContextMenu",this)},Handler:function(t,r,s){if(l.loadingMathMenu){return n.False(t)}var q=b[s.jaxID];if(!t){t=window.event}t.isContextMenu=(r==="ContextMenu");if(q[r]){return q[r](t,s)}if(i.MathZoom){return i.MathZoom.HandleEvent(t,r,s)}},False:function(q){if(!q){q=window.event}if(q){if(q.preventDefault){q.preventDefault()}else{q.returnValue=false}if(q.stopPropagation){q.stopPropagation()}q.cancelBubble=true}return false},Keydown:function(r,q){if(!r){r=window.event}if(r.keyCode===n.KEY.SPACE){n.ContextMenu(r,this)}},ContextMenu:function(t,E,w){var B=b[E.jaxID],v=B.getJaxFromMath(E);var F=(B.config.showMathMenu!=null?B:d).config.showMathMenu;if(!F||(k.context!=="MathJax"&&!w)){return}if(c.msieEventBug){t=window.event||t}n.ClearSelection();f.ClearHoverTimer();if(v.hover){if(v.hover.remove){clearTimeout(v.hover.remove);delete v.hover.remove}v.hover.nofade=true}var u=MathJax.Menu;var G,D;if(u){if(u.loadingDomain){return n.False(t)}G=m.loadDomain("MathMenu");if(!G){u.jax=v;var r=u.menu.Find("Show Math As").submenu;r.items[0].name=v.sourceMenuTitle;r.items[0].format=(v.sourceMenuFormat||"MathML");r.items[1].name=j[v.inputJax].sourceMenuTitle;r.items[5].disabled=!j[v.inputJax].annotationEncoding;var A=r.items[2];A.disabled=true;var q=A.submenu.items;annotationList=MathJax.Hub.Config.semanticsAnnotations;for(var z=0,y=q.length;z<y;z++){var s=q[z].name[1];if(v.root&&v.root.getAnnotation(s)!==null){A.disabled=false;q[z].hidden=false}else{q[z].hidden=true}}var x=u.menu.Find("Math Settings","MathPlayer");x.hidden=!(v.outputJax==="NativeMML"&&d.Browser.hasMathPlayer);return u.menu.Post(t)}u.loadingDomain=true;D=function(){delete u.loadingDomain}}else{if(l.loadingMathMenu){return n.False(t)}l.loadingMathMenu=true;G=l.Require("[MathJax]/extensions/MathMenu.js");D=function(){delete l.loadingMathMenu;if(!MathJax.Menu){MathJax.Menu={}}}}var C={pageX:t.pageX,pageY:t.pageY,clientX:t.clientX,clientY:t.clientY};g.Queue(G,D,["ContextMenu",n,C,E,w]);return n.False(t)},AltContextMenu:function(s,r){var t=b[r.jaxID];var q=(t.config.showMathMenu!=null?t:d).config.showMathMenu;if(q){q=(t.config.showMathMenuMSIE!=null?t:d).config.showMathMenuMSIE;if(k.context==="MathJax"&&!k.mpContext&&q){if(!c.noContextMenuBug||s.button!==n.RIGHTBUTTON){return}}else{if(!s[n.MENUKEY]||s.button!==n.LEFTBUTTON){return}}return t.ContextMenu(s,r,true)}},ClearSelection:function(){if(c.safariContextMenuBug){setTimeout("window.getSelection().empty()",0)}if(document.selection){setTimeout("document.selection.empty()",0)}},getBBox:function(s){s.appendChild(c.topImg);var r=c.topImg.offsetTop,t=s.offsetHeight-r,q=s.offsetWidth;s.removeChild(c.topImg);return{w:q,h:r,d:t}}};var f=c.Hover={Mouseover:function(s,r){if(k.discoverable||k.zoom==="Hover"){var u=s.fromElement||s.relatedTarget,t=s.toElement||s.target;if(u&&t&&(d.isMathJaxNode(u)!==d.isMathJaxNode(t)||d.getJaxFor(u)!==d.getJaxFor(t))){var q=this.getJaxFromMath(r);if(q.hover){f.ReHover(q)}else{f.HoverTimer(q,r)}return n.False(s)}}},Mouseout:function(s,r){if(k.discoverable||k.zoom==="Hover"){var u=s.fromElement||s.relatedTarget,t=s.toElement||s.target;if(u&&t&&(d.isMathJaxNode(u)!==d.isMathJaxNode(t)||d.getJaxFor(u)!==d.getJaxFor(t))){var q=this.getJaxFromMath(r);if(q.hover){f.UnHover(q)}else{f.ClearHoverTimer()}return n.False(s)}}},Mousemove:function(s,r){if(k.discoverable||k.zoom==="Hover"){var q=this.getJaxFromMath(r);if(q.hover){return}if(f.lastX==s.clientX&&f.lastY==s.clientY){return}f.lastX=s.clientX;f.lastY=s.clientY;f.HoverTimer(q,r);return n.False(s)}},HoverTimer:function(q,r){this.ClearHoverTimer();this.hoverTimer=setTimeout(g(["Hover",this,q,r]),o.hover)},ClearHoverTimer:function(){if(this.hoverTimer){clearTimeout(this.hoverTimer);delete this.hoverTimer}},Hover:function(q,u){if(i.MathZoom&&i.MathZoom.Hover({},u)){return}var t=b[q.outputJax],v=t.getHoverSpan(q,u),y=t.getHoverBBox(q,v,u),w=(t.config.showMathMenu!=null?t:d).config.showMathMenu;var A=o.frame.x,z=o.frame.y,x=o.frame.bwidth;if(c.msieBorderWidthBug){x=0}q.hover={opacity:0,id:q.inputID+"-Hover"};var r=h.Element("span",{id:q.hover.id,isMathJax:true,style:{display:"inline-block",width:0,height:0,position:"relative"}},[["span",{className:"MathJax_Hover_Frame",isMathJax:true,style:{display:"inline-block",position:"absolute",top:this.Px(-y.h-z-x-(y.y||0)),left:this.Px(-A-x+(y.x||0)),width:this.Px(y.w+2*A),height:this.Px(y.h+y.d+2*z),opacity:0,filter:"alpha(opacity=0)"}}]]);var s=h.Element("span",{isMathJax:true,id:q.hover.id+"Menu",className:"MathJax_Menu_Button",style:{display:"inline-block","z-index":1,width:0,height:0,position:"relative"}},[["span",{className:"MathJax_Hover_Arrow",isMathJax:true,math:u,onclick:this.HoverMenu,jax:t.id,style:{left:this.Px(y.w+A+x+(y.x||0)+o.button.x),top:this.Px(-y.h-z-x-(y.y||0)-o.button.y),opacity:0,filter:"alpha(opacity=0)"}},[["span",{isMathJax:true},"\u25BC"]]]]);if(y.width){r.style.width=s.style.width=y.width;r.style.marginRight=s.style.marginRight="-"+y.width;r.firstChild.style.width=y.width;s.firstChild.style.left="";s.firstChild.style.right=this.Px(o.button.wx)}v.parentNode.insertBefore(r,v);if(w){v.parentNode.insertBefore(s,v)}if(v.style){v.style.position="relative"}this.ReHover(q)},ReHover:function(q){if(q.hover.remove){clearTimeout(q.hover.remove)}q.hover.remove=setTimeout(g(["UnHover",this,q]),o.fadeoutDelay);this.HoverFadeTimer(q,o.fadeinInc)},UnHover:function(q){if(!q.hover.nofade){this.HoverFadeTimer(q,-o.fadeoutInc,o.fadeoutStart)}},HoverFade:function(q){delete q.hover.timer;q.hover.opacity=Math.max(0,Math.min(1,q.hover.opacity+q.hover.inc));q.hover.opacity=Math.floor(1000*q.hover.opacity)/1000;var s=document.getElementById(q.hover.id),r=document.getElementById(q.hover.id+"Menu");s.firstChild.style.opacity=q.hover.opacity;s.firstChild.style.filter="alpha(opacity="+Math.floor(100*q.hover.opacity)+")";if(r){r.firstChild.style.opacity=q.hover.opacity;r.firstChild.style.filter=s.style.filter}if(q.hover.opacity===1){return}if(q.hover.opacity>0){this.HoverFadeTimer(q,q.hover.inc);return}s.parentNode.removeChild(s);if(r){r.parentNode.removeChild(r)}if(q.hover.remove){clearTimeout(q.hover.remove)}delete q.hover},HoverFadeTimer:function(q,s,r){q.hover.inc=s;if(!q.hover.timer){q.hover.timer=setTimeout(g(["HoverFade",this,q]),(r||o.fadeDelay))}},HoverMenu:function(q){if(!q){q=window.event}return b[this.jax].ContextMenu(q,this.math,true)},ClearHover:function(q){if(q.hover.remove){clearTimeout(q.hover.remove)}if(q.hover.timer){clearTimeout(q.hover.timer)}f.ClearHoverTimer();delete q.hover},Px:function(q){if(Math.abs(q)<0.006){return"0px"}return q.toFixed(2).replace(/\.?0+$/,"")+"px"},getImages:function(){if(k.discoverable){var q=new Image();q.src=o.button.src}}};var a=c.Touch={last:0,delay:500,start:function(r){var q=new Date().getTime();var s=(q-a.last<a.delay&&a.up);a.last=q;a.up=false;if(s){a.timeout=setTimeout(a.menu,a.delay,r,this);r.preventDefault()}},end:function(r){var q=new Date().getTime();a.up=(q-a.last<a.delay);if(a.timeout){clearTimeout(a.timeout);delete a.timeout;a.last=0;a.up=false;r.preventDefault();return n.Handler((r.touches[0]||r.touch),"DblClick",this)}},menu:function(r,q){delete a.timeout;a.last=0;a.up=false;return n.Handler((r.touches[0]||r.touch),"ContextMenu",q)}};d.Browser.Select({MSIE:function(q){var s=(document.documentMode||0);var r=q.versionAtLeast("8.0");c.msieBorderWidthBug=(document.compatMode==="BackCompat");c.msieEventBug=q.isIE9;c.msieAlignBug=(!r||s<8);if(s<9){n.LEFTBUTTON=1}},Safari:function(q){c.safariContextMenuBug=true},Opera:function(q){c.operaPositionBug=true},Konqueror:function(q){c.noContextMenuBug=true}});c.topImg=(c.msieAlignBug?h.Element("img",{style:{width:0,height:0,position:"relative"},src:"about:blank"}):h.Element("span",{style:{width:0,height:0,display:"inline-block"}}));if(c.operaPositionBug){c.topImg.style.border="1px solid"}c.config=o=d.CombineConfig("MathEvents",o);var e=function(){var q=o.styles[".MathJax_Hover_Frame"];q.border=o.frame.bwidth+"px solid "+o.frame.bcolor+" ! important";q["box-shadow"]=q["-webkit-box-shadow"]=q["-moz-box-shadow"]=q["-khtml-box-shadow"]="0px 0px "+o.frame.hwidth+" "+o.frame.hcolor};g.Queue(d.Register.StartupHook("End Config",{}),[e],["getImages",f],["Styles",l,o.styles],["Post",d.Startup.signal,"MathEvents Ready"],["loadComplete",l,"[MathJax]/extensions/MathEvents.js"])})(MathJax.Hub,MathJax.HTML,MathJax.Ajax,MathJax.Callback,MathJax.Localization,MathJax.OutputJax,MathJax.InputJax);
(function(a,d,f,c,j){var k="2.7.0";var i=a.CombineConfig("MathZoom",{styles:{"#MathJax_Zoom":{position:"absolute","background-color":"#F0F0F0",overflow:"auto",display:"block","z-index":301,padding:".5em",border:"1px solid black",margin:0,"font-weight":"normal","font-style":"normal","text-align":"left","text-indent":0,"text-transform":"none","line-height":"normal","letter-spacing":"normal","word-spacing":"normal","word-wrap":"normal","white-space":"nowrap","float":"none","-webkit-box-sizing":"content-box","-moz-box-sizing":"content-box","box-sizing":"content-box","box-shadow":"5px 5px 15px #AAAAAA","-webkit-box-shadow":"5px 5px 15px #AAAAAA","-moz-box-shadow":"5px 5px 15px #AAAAAA","-khtml-box-shadow":"5px 5px 15px #AAAAAA",filter:"progid:DXImageTransform.Microsoft.dropshadow(OffX=2, OffY=2, Color='gray', Positive='true')"},"#MathJax_ZoomOverlay":{position:"absolute",left:0,top:0,"z-index":300,display:"inline-block",width:"100%",height:"100%",border:0,padding:0,margin:0,"background-color":"white",opacity:0,filter:"alpha(opacity=0)"},"#MathJax_ZoomFrame":{position:"relative",display:"inline-block",height:0,width:0},"#MathJax_ZoomEventTrap":{position:"absolute",left:0,top:0,"z-index":302,display:"inline-block",border:0,padding:0,margin:0,"background-color":"white",opacity:0,filter:"alpha(opacity=0)"}}});var e,b,g;MathJax.Hub.Register.StartupHook("MathEvents Ready",function(){g=MathJax.Extension.MathEvents.Event;e=MathJax.Extension.MathEvents.Event.False;b=MathJax.Extension.MathEvents.Hover});var h=MathJax.Extension.MathZoom={version:k,settings:a.config.menuSettings,scrollSize:18,HandleEvent:function(n,l,m){if(h.settings.CTRL&&!n.ctrlKey){return true}if(h.settings.ALT&&!n.altKey){return true}if(h.settings.CMD&&!n.metaKey){return true}if(h.settings.Shift&&!n.shiftKey){return true}if(!h[l]){return true}return h[l](n,m)},Click:function(m,l){if(this.settings.zoom==="Click"){return this.Zoom(m,l)}},DblClick:function(m,l){if(this.settings.zoom==="Double-Click"||this.settings.zoom==="DoubleClick"){return this.Zoom(m,l)}},Hover:function(m,l){if(this.settings.zoom==="Hover"){this.Zoom(m,l);return true}return false},Zoom:function(o,u){this.Remove();b.ClearHoverTimer();g.ClearSelection();var s=MathJax.OutputJax[u.jaxID];var p=s.getJaxFromMath(u);if(p.hover){b.UnHover(p)}var q=this.findContainer(u);var l=Math.floor(0.85*q.clientWidth),t=Math.max(document.body.clientHeight,document.documentElement.clientHeight);if(this.getOverflow(q)!=="visible"){t=Math.min(q.clientHeight,t)}t=Math.floor(0.85*t);var n=d.Element("span",{id:"MathJax_ZoomFrame"},[["span",{id:"MathJax_ZoomOverlay",onmousedown:this.Remove}],["span",{id:"MathJax_Zoom",onclick:this.Remove,style:{visibility:"hidden",fontSize:this.settings.zscale}},[["span",{style:{display:"inline-block","white-space":"nowrap"}}]]]]);var z=n.lastChild,w=z.firstChild,r=n.firstChild;u.parentNode.insertBefore(n,u);u.parentNode.insertBefore(u,n);if(w.addEventListener){w.addEventListener("mousedown",this.Remove,true)}var m=z.offsetWidth||z.clientWidth;l-=m;t-=m;z.style.maxWidth=l+"px";z.style.maxHeight=t+"px";if(this.msieTrapEventBug){var y=d.Element("span",{id:"MathJax_ZoomEventTrap",onmousedown:this.Remove});n.insertBefore(y,z)}if(this.msieZIndexBug){var v=d.addElement(document.body,"img",{src:"about:blank",id:"MathJax_ZoomTracker",width:0,height:0,style:{width:0,height:0,position:"relative"}});n.style.position="relative";n.style.zIndex=i.styles["#MathJax_ZoomOverlay"]["z-index"];n=v}var x=s.Zoom(p,w,u,l,t);if(this.msiePositionBug){if(this.msieSizeBug){z.style.height=x.zH+"px";z.style.width=x.zW+"px"}if(z.offsetHeight>t){z.style.height=t+"px";z.style.width=(x.zW+this.scrollSize)+"px"}if(z.offsetWidth>l){z.style.width=l+"px";z.style.height=(x.zH+this.scrollSize)+"px"}}if(this.operaPositionBug){z.style.width=Math.min(l,x.zW)+"px"}if(z.offsetWidth>m&&z.offsetWidth-m<l&&z.offsetHeight-m<t){z.style.overflow="visible"}this.Position(z,x);if(this.msieTrapEventBug){y.style.height=z.clientHeight+"px";y.style.width=z.clientWidth+"px";y.style.left=(parseFloat(z.style.left)+z.clientLeft)+"px";y.style.top=(parseFloat(z.style.top)+z.clientTop)+"px"}z.style.visibility="";if(this.settings.zoom==="Hover"){r.onmouseover=this.Remove}if(window.addEventListener){addEventListener("resize",this.Resize,false)}else{if(window.attachEvent){attachEvent("onresize",this.Resize)}else{this.onresize=window.onresize;window.onresize=this.Resize}}a.signal.Post(["math zoomed",p]);return e(o)},Position:function(p,r){p.style.display="none";var q=this.Resize(),m=q.x,s=q.y,l=r.mW;p.style.display="";var o=-l-Math.floor((p.offsetWidth-l)/2),n=r.Y;p.style.left=Math.max(o,10-m)+"px";p.style.top=Math.max(n,10-s)+"px";if(!h.msiePositionBug){h.SetWH()}},Resize:function(m){if(h.onresize){h.onresize(m)}var q=document.getElementById("MathJax_ZoomFrame"),l=document.getElementById("MathJax_ZoomOverlay");var o=h.getXY(q),n=h.findContainer(q);if(h.getOverflow(n)!=="visible"){l.scroll_parent=n;var p=h.getXY(n);o.x-=p.x;o.y-=p.y;p=h.getBorder(n);o.x-=p.x;o.y-=p.y}l.style.left=(-o.x)+"px";l.style.top=(-o.y)+"px";if(h.msiePositionBug){setTimeout(h.SetWH,0)}else{h.SetWH()}return o},SetWH:function(){var l=document.getElementById("MathJax_ZoomOverlay");if(!l){return}l.style.display="none";var m=l.scroll_parent||document.documentElement||document.body;l.style.width=m.scrollWidth+"px";l.style.height=Math.max(m.clientHeight,m.scrollHeight)+"px";l.style.display=""},findContainer:function(l){l=l.parentNode;while(l.parentNode&&l!==document.body&&h.getOverflow(l)==="visible"){l=l.parentNode}return l},getOverflow:(window.getComputedStyle?function(l){return getComputedStyle(l).overflow}:function(l){return(l.currentStyle||{overflow:"visible"}).overflow}),getBorder:function(o){var m={thin:1,medium:2,thick:3};var n=(window.getComputedStyle?getComputedStyle(o):(o.currentStyle||{borderLeftWidth:0,borderTopWidth:0}));var l=n.borderLeftWidth,p=n.borderTopWidth;if(m[l]){l=m[l]}else{l=parseInt(l)}if(m[p]){p=m[p]}else{p=parseInt(p)}return{x:l,y:p}},getXY:function(o){var l=0,n=0,m;m=o;while(m.offsetParent){l+=m.offsetLeft;m=m.offsetParent}if(h.operaPositionBug){o.style.border="1px solid"}m=o;while(m.offsetParent){n+=m.offsetTop;m=m.offsetParent}if(h.operaPositionBug){o.style.border=""}return{x:l,y:n}},Remove:function(n){var p=document.getElementById("MathJax_ZoomFrame");if(p){var o=MathJax.OutputJax[p.previousSibling.jaxID];var l=o.getJaxFromMath(p.previousSibling);a.signal.Post(["math unzoomed",l]);p.parentNode.removeChild(p);p=document.getElementById("MathJax_ZoomTracker");if(p){p.parentNode.removeChild(p)}if(h.operaRefreshBug){var m=d.addElement(document.body,"div",{style:{position:"fixed",left:0,top:0,width:"100%",height:"100%",backgroundColor:"white",opacity:0},id:"MathJax_OperaDiv"});document.body.removeChild(m)}if(window.removeEventListener){removeEventListener("resize",h.Resize,false)}else{if(window.detachEvent){detachEvent("onresize",h.Resize)}else{window.onresize=h.onresize;delete h.onresize}}}return e(n)}};a.Browser.Select({MSIE:function(l){var n=(document.documentMode||0);var m=(n>=9);h.msiePositionBug=!m;h.msieSizeBug=l.versionAtLeast("7.0")&&(!document.documentMode||n===7||n===8);h.msieZIndexBug=(n<=7);h.msieInlineBlockAlignBug=(n<=7);h.msieTrapEventBug=!window.addEventListener;if(document.compatMode==="BackCompat"){h.scrollSize=52}if(m){delete i.styles["#MathJax_Zoom"].filter}},Opera:function(l){h.operaPositionBug=true;h.operaRefreshBug=true}});h.topImg=(h.msieInlineBlockAlignBug?d.Element("img",{style:{width:0,height:0,position:"relative"},src:"about:blank"}):d.Element("span",{style:{width:0,height:0,display:"inline-block"}}));if(h.operaPositionBug||h.msieTopBug){h.topImg.style.border="1px solid"}MathJax.Callback.Queue(["StartupHook",MathJax.Hub.Register,"Begin Styles",{}],["Styles",f,i.styles],["Post",a.Startup.signal,"MathZoom Ready"],["loadComplete",f,"[MathJax]/extensions/MathZoom.js"])})(MathJax.Hub,MathJax.HTML,MathJax.Ajax,MathJax.OutputJax["HTML-CSS"],MathJax.OutputJax.NativeMML);
(function(f,o,q,e,r){var p="2.7.0";var d=MathJax.Callback.Signal("menu");MathJax.Extension.MathMenu={version:p,signal:d};var t=function(u){return MathJax.Localization._.apply(MathJax.Localization,[["MathMenu",u]].concat([].slice.call(arguments,1)))};var i=MathJax.Object.isArray;var a=f.Browser.isPC,l=f.Browser.isMSIE,m=((document.documentMode||0)>8);var j=(a?null:"5px");var s=f.CombineConfig("MathMenu",{delay:150,showRenderer:true,showMathPlayer:true,showFontMenu:false,showContext:false,showDiscoverable:false,showLocale:true,showLocaleURL:false,semanticsAnnotations:{TeX:["TeX","LaTeX","application/x-tex"],StarMath:["StarMath 5.0"],Maple:["Maple"],ContentMathML:["MathML-Content","application/mathml-content+xml"],OpenMath:["OpenMath"]},windowSettings:{status:"no",toolbar:"no",locationbar:"no",menubar:"no",directories:"no",personalbar:"no",resizable:"yes",scrollbars:"yes",width:400,height:300,left:Math.round((screen.width-400)/2),top:Math.round((screen.height-300)/3)},styles:{"#MathJax_About":{position:"fixed",left:"50%",width:"auto","text-align":"center",border:"3px outset",padding:"1em 2em","background-color":"#DDDDDD",color:"black",cursor:"default","font-family":"message-box","font-size":"120%","font-style":"normal","text-indent":0,"text-transform":"none","line-height":"normal","letter-spacing":"normal","word-spacing":"normal","word-wrap":"normal","white-space":"nowrap","float":"none","z-index":201,"border-radius":"15px","-webkit-border-radius":"15px","-moz-border-radius":"15px","-khtml-border-radius":"15px","box-shadow":"0px 10px 20px #808080","-webkit-box-shadow":"0px 10px 20px #808080","-moz-box-shadow":"0px 10px 20px #808080","-khtml-box-shadow":"0px 10px 20px #808080",filter:"progid:DXImageTransform.Microsoft.dropshadow(OffX=2, OffY=2, Color='gray', Positive='true')"},"#MathJax_About.MathJax_MousePost":{outline:"none"},".MathJax_Menu":{position:"absolute","background-color":"white",color:"black",width:"auto",padding:(a?"2px":"5px 0px"),border:"1px solid #CCCCCC",margin:0,cursor:"default",font:"menu","text-align":"left","text-indent":0,"text-transform":"none","line-height":"normal","letter-spacing":"normal","word-spacing":"normal","word-wrap":"normal","white-space":"nowrap","float":"none","z-index":201,"border-radius":j,"-webkit-border-radius":j,"-moz-border-radius":j,"-khtml-border-radius":j,"box-shadow":"0px 10px 20px #808080","-webkit-box-shadow":"0px 10px 20px #808080","-moz-box-shadow":"0px 10px 20px #808080","-khtml-box-shadow":"0px 10px 20px #808080",filter:"progid:DXImageTransform.Microsoft.dropshadow(OffX=2, OffY=2, Color='gray', Positive='true')"},".MathJax_MenuItem":{padding:(a?"2px 2em":"1px 2em"),background:"transparent"},".MathJax_MenuArrow":{position:"absolute",right:".5em","padding-top":".25em",color:"#666666","font-family":(l?"'Arial unicode MS'":null),"font-size":".75em"},".MathJax_MenuActive .MathJax_MenuArrow":{color:"white"},".MathJax_MenuArrow.RTL":{left:".5em",right:"auto"},".MathJax_MenuCheck":{position:"absolute",left:".7em","font-family":(l?"'Arial unicode MS'":null)},".MathJax_MenuCheck.RTL":{right:".7em",left:"auto"},".MathJax_MenuRadioCheck":{position:"absolute",left:(a?"1em":".7em")},".MathJax_MenuRadioCheck.RTL":{right:(a?"1em":".7em"),left:"auto"},".MathJax_MenuLabel":{padding:(a?"2px 2em 4px 1.33em":"1px 2em 3px 1.33em"),"font-style":"italic"},".MathJax_MenuRule":{"border-top":(a?"1px solid #CCCCCC":"1px solid #DDDDDD"),margin:(a?"4px 1px 0px":"4px 3px")},".MathJax_MenuDisabled":{color:"GrayText"},".MathJax_MenuActive":{"background-color":(a?"Highlight":"#606872"),color:(a?"HighlightText":"white")},".MathJax_MenuDisabled:focus, .MathJax_MenuLabel:focus":{"background-color":"#E8E8E8"},".MathJax_ContextMenu:focus":{outline:"none"},".MathJax_ContextMenu .MathJax_MenuItem:focus":{outline:"none"},"#MathJax_AboutClose":{top:".2em",right:".2em"},".MathJax_Menu .MathJax_MenuClose":{top:"-10px",left:"-10px"},".MathJax_MenuClose":{position:"absolute",cursor:"pointer",display:"inline-block",border:"2px solid #AAA","border-radius":"18px","-webkit-border-radius":"18px","-moz-border-radius":"18px","-khtml-border-radius":"18px","font-family":"'Courier New',Courier","font-size":"24px",color:"#F0F0F0"},".MathJax_MenuClose span":{display:"block","background-color":"#AAA",border:"1.5px solid","border-radius":"18px","-webkit-border-radius":"18px","-moz-border-radius":"18px","-khtml-border-radius":"18px","line-height":0,padding:"8px 0 6px"},".MathJax_MenuClose:hover":{color:"white!important",border:"2px solid #CCC!important"},".MathJax_MenuClose:hover span":{"background-color":"#CCC!important"},".MathJax_MenuClose:hover:focus":{outline:"none"}}});var n,k,b;f.Register.StartupHook("MathEvents Ready",function(){n=MathJax.Extension.MathEvents.Event.False;k=MathJax.Extension.MathEvents.Hover;b=MathJax.Extension.MathEvents.Event.KEY});var h=MathJax.Object.Subclass({Keydown:function(u,v){switch(u.keyCode){case b.ESCAPE:this.Remove(u,v);break;case b.RIGHT:this.Right(u,v);break;case b.LEFT:this.Left(u,v);break;case b.UP:this.Up(u,v);break;case b.DOWN:this.Down(u,v);break;case b.RETURN:case b.SPACE:this.Space(u,v);break;default:return;break}return n(u)},Escape:function(u,v){},Right:function(u,v){},Left:function(u,v){},Up:function(u,v){},Down:function(u,v){},Space:function(u,v){}},{});var g=MathJax.Menu=h.Subclass({version:p,items:[],posted:false,title:null,margin:5,Init:function(u){this.items=[].slice.call(arguments,0)},With:function(u){if(u){f.Insert(this,u)}return this},Post:function(M,E,B){if(!M){M=window.event||{}}var I=document.getElementById("MathJax_MenuFrame");if(!I){I=g.Background(this);delete c.lastItem;delete c.lastMenu;delete g.skipUp;d.Post(["post",g.jax]);g.isRTL=(MathJax.Localization.fontDirection()==="rtl")}var v=o.Element("div",{onmouseup:g.Mouseup,ondblclick:n,ondragstart:n,onselectstart:n,oncontextmenu:n,menuItem:this,className:"MathJax_Menu",onkeydown:g.Keydown,role:"menu"});if(M.type==="contextmenu"||M.type==="mouseover"){v.className+=" MathJax_ContextMenu"}if(!B){MathJax.Localization.setCSS(v)}for(var N=0,K=this.items.length;N<K;N++){this.items[N].Create(v)}if(g.isMobile){o.addElement(v,"span",{className:"MathJax_MenuClose",menu:E,ontouchstart:g.Close,ontouchend:n,onmousedown:g.Close,onmouseup:n},[["span",{},"\u00D7"]])}I.appendChild(v);this.posted=true;if(v.offsetWidth){v.style.width=(v.offsetWidth+2)+"px"}var H=M.pageX,F=M.pageY;var u=document.body.getBoundingClientRect();var C=(window.getComputedStyle?window.getComputedStyle(document.body):{marginLeft:"0px"});var A=u.right-Math.min(0,u.left)+parseFloat(C.marginLeft);if(!H&&!F&&"clientX" in M){H=M.clientX+document.body.scrollLeft+document.documentElement.scrollLeft;F=M.clientY+document.body.scrollTop+document.documentElement.scrollTop}if(!E){var L=g.CurrentNode()||M.target;if((M.type==="keydown"||(!H&&!F))&&L){var P=window.pageXOffset||document.documentElement.scrollLeft;var O=window.pageYOffset||document.documentElement.scrollTop;var w=L.getBoundingClientRect();H=(w.right+w.left)/2+P;F=(w.bottom+w.top)/2+O}if(H+v.offsetWidth>A-this.margin){H=A-v.offsetWidth-this.margin}if(g.isMobile){H=Math.max(5,H-Math.floor(v.offsetWidth/2));F-=20}g.skipUp=M.isContextMenu}else{var z="left",J=E.offsetWidth;H=(g.isMobile?30:J-2);F=0;while(E&&E!==I){H+=E.offsetLeft;F+=E.offsetTop;E=E.parentNode}if(!g.isMobile){if((g.isRTL&&H-J-v.offsetWidth>this.margin)||(!g.isRTL&&H+v.offsetWidth>A-this.margin)){z="right";H=Math.max(this.margin,H-J-v.offsetWidth+6)}}if(!a){v.style["borderRadiusTop"+z]=0;v.style["WebkitBorderRadiusTop"+z]=0;v.style["MozBorderRadiusTop"+z]=0;v.style["KhtmlBorderRadiusTop"+z]=0}}v.style.left=H+"px";v.style.top=F+"px";if(document.selection&&document.selection.empty){document.selection.empty()}var G=window.pageXOffset||document.documentElement.scrollLeft;var D=window.pageYOffset||document.documentElement.scrollTop;g.Focus(v);if(M.type==="keydown"){g.skipMouseoverFromKey=true;setTimeout(function(){delete g.skipMouseoverFromKey},s.delay)}window.scrollTo(G,D);return n(M)},Remove:function(u,v){d.Post(["unpost",g.jax]);var w=document.getElementById("MathJax_MenuFrame");if(w){w.parentNode.removeChild(w);if(this.msieFixedPositionBug){detachEvent("onresize",g.Resize)}}if(g.jax.hover){delete g.jax.hover.nofade;k.UnHover(g.jax)}g.Unfocus(v);if(u.type==="mousedown"){g.CurrentNode().blur()}return n(u)},Find:function(u){return this.FindN(1,u,[].slice.call(arguments,1))},FindId:function(u){return this.FindN(0,u,[].slice.call(arguments,1))},FindN:function(y,v,x){for(var w=0,u=this.items.length;w<u;w++){if(this.items[w].name[y]===v){if(x.length){if(!this.items[w].submenu){return null}return this.items[w].submenu.FindN(y,x[0],x.slice(1))}return this.items[w]}}return null},IndexOf:function(u){return this.IndexOfN(1,u)},IndexOfId:function(u){return this.IndexOfN(0,u)},IndexOfN:function(x,v){for(var w=0,u=this.items.length;w<u;w++){if(this.items[w].name[x]===v){return w}}return null},Right:function(u,v){g.Right(u,v)},Left:function(u,v){g.Left(u,v)},Up:function(v,w){var u=w.lastChild;u.menuItem.Activate(v,u)},Down:function(v,w){var u=w.firstChild;u.menuItem.Activate(v,u)},Space:function(u,v){this.Remove(u,v)}},{config:s,Remove:function(u){return g.Event(u,this,"Remove")},Mouseover:function(u){return g.Event(u,this,"Mouseover")},Mouseout:function(u){return g.Event(u,this,"Mouseout")},Mousedown:function(u){return g.Event(u,this,"Mousedown")},Mouseup:function(u){return g.Event(u,this,"Mouseup")},Keydown:function(u){return g.Event(u,this,"Keydown")},Touchstart:function(u){return g.Event(u,this,"Touchstart")},Touchend:function(u){return g.Event(u,this,"Touchend")},Close:function(u){return g.Event(u,this.menu||this.parentNode,(this.menu?"Touchend":"Remove"))},Event:function(w,y,u,x){if(g.skipMouseover&&u==="Mouseover"&&!x){return n(w)}if(g.skipMouseoverFromKey&&u==="Mouseover"){delete g.skipMouseoverFromKey;return n(w)}if(g.skipUp){if(u.match(/Mouseup|Touchend/)){delete g.skipUp;return n(w)}if(u==="Touchstart"||(u==="Mousedown"&&!g.skipMousedown)){delete g.skipUp}}if(!w){w=window.event}var v=y.menuItem;if(v&&v[u]){return v[u](w,y)}return null},BGSTYLE:{position:"absolute",left:0,top:0,"z-index":200,width:"100%",height:"100%",border:0,padding:0,margin:0},Background:function(v){var w=o.addElement(document.body,"div",{style:this.BGSTYLE,id:"MathJax_MenuFrame"},[["div",{style:this.BGSTYLE,menuItem:v,onmousedown:this.Remove}]]);var u=w.firstChild;if(g.msieBackgroundBug){u.style.backgroundColor="white";u.style.filter="alpha(opacity=0)"}if(g.msieFixedPositionBug){w.width=w.height=0;this.Resize();attachEvent("onresize",this.Resize)}else{u.style.position="fixed"}return w},Resize:function(){setTimeout(g.SetWH,0)},SetWH:function(){var u=document.getElementById("MathJax_MenuFrame");if(u){u=u.firstChild;u.style.width=u.style.height="1px";u.style.width=document.body.scrollWidth+"px";u.style.height=document.body.scrollHeight+"px"}},posted:false,active:null,GetNode:function(u){var v=document.getElementById(u.inputID+"-Frame");return v.isMathJax?v:v.firstChild},CurrentNode:function(){return g.GetNode(g.jax)},AllNodes:function(){var v=MathJax.Hub.getAllJax();var w=[];for(var x=0,u;u=v[x];x++){w.push(g.GetNode(u))}return w},ActiveNode:function(){return g.active},FocusNode:function(u){g.active=u;u.focus()},Focus:function(u){!g.posted?g.Activate(u):g.ActiveNode().tabIndex=-1;u.tabIndex=0;g.FocusNode(u)},Activate:function(u,v){g.UnsetTabIndex();g.posted=true},Unfocus:function(){g.ActiveNode().tabIndex=-1;g.SetTabIndex();g.FocusNode(g.CurrentNode());g.posted=false},MoveHorizontal:function(y,z,w){if(!y.shiftKey){return}var v=g.AllNodes();var u=v.length;if(u===0){return}var x=v[g.Mod(w(g.IndexOf(v,g.CurrentNode())),u)];if(x===g.CurrentNode()){return}g.menu.Remove(y,z);g.jax=MathJax.Hub.getJaxFor(x);g.FocusNode(x);g.menu.Post(null)},Right:function(u,v){g.MoveHorizontal(u,v,function(w){return w+1})},Left:function(u,v){g.MoveHorizontal(u,v,function(w){return w-1})},UnsetTabIndex:function(){var v=g.AllNodes();for(var w=0,u;u=v[w];w++){if(u.tabIndex>0){u.oldTabIndex=u.tabIndex}u.tabIndex=-1}},SetTabIndex:function(){var v=g.AllNodes();for(var w=0,u;u=v[w];w++){if(u.oldTabIndex!==undefined){u.tabIndex=u.oldTabIndex;delete u.oldTabIndex}else{u.tabIndex=f.getTabOrder(u)}}},Mod:function(u,v){return((u%v)+v)%v},IndexOf:(Array.prototype.indexOf?function(u,v,w){return u.indexOf(v,w)}:function(u,x,y){for(var w=(y||0),v=u.length;w<v;w++){if(x===u[w]){return w}}return -1}),saveCookie:function(){o.Cookie.Set("menu",this.cookie)},getCookie:function(){this.cookie=o.Cookie.Get("menu")}});MathJax.Menu.NAV=h;var c=g.ITEM=h.Subclass({name:"",node:null,menu:null,Attributes:function(u){return f.Insert({onmouseup:g.Mouseup,ondragstart:n,onselectstart:n,onselectend:n,ontouchstart:g.Touchstart,ontouchend:g.Touchend,className:"MathJax_MenuItem",role:this.role,menuItem:this},u)},Create:function(w){if(!this.hidden){var v=this.Attributes();var u=this.Label(v,w);o.addElement(w,"div",v,u)}},Name:function(){return t(this.name[0],this.name[1])},Mouseover:function(u,v){if(v.parentNode===g.ActiveNode().parentNode){this.Deactivate(g.ActiveNode())}this.Activate(u,v)},Mouseout:function(u,v){this.Deactivate(v)},Mouseup:function(u,v){return this.Remove(u,v)},DeactivateSubmenus:function(z){var y=document.getElementById("MathJax_MenuFrame").childNodes,v=c.GetMenuNode(z).childNodes;for(var w=0,u=v.length;w<u;w++){var x=v[w].menuItem;if(x&&x.submenu&&x.submenu.posted&&x!==z.menuItem){x.Deactivate(v[w])}}this.RemoveSubmenus(z,y)},RemoveSubmenus:function(w,v){v=v||document.getElementById("MathJax_MenuFrame").childNodes;var u=v.length-1;while(u>=0&&c.GetMenuNode(w).menuItem!==v[u].menuItem){v[u].menuItem.posted=false;v[u].parentNode.removeChild(v[u]);u--}},Touchstart:function(u,v){return this.TouchEvent(u,v,"Mousedown")},Touchend:function(u,v){return this.TouchEvent(u,v,"Mouseup")},TouchEvent:function(v,w,u){if(this!==c.lastItem){if(c.lastMenu){g.Event(v,c.lastMenu,"Mouseout")}g.Event(v,w,"Mouseover",true);c.lastItem=this;c.lastMenu=w}if(this.nativeTouch){return null}g.Event(v,w,u);return false},Remove:function(u,v){v=v.parentNode.menuItem;return v.Remove(u,v)},With:function(u){if(u){f.Insert(this,u)}return this},isRTL:function(){return g.isRTL},rtlClass:function(){return(this.isRTL()?" RTL":"")}},{GetMenuNode:function(u){return u.parentNode}});g.ENTRY=g.ITEM.Subclass({role:"menuitem",Attributes:function(u){u=f.Insert({onmouseover:g.Mouseover,onmouseout:g.Mouseout,onmousedown:g.Mousedown,onkeydown:g.Keydown,"aria-disabled":!!this.disabled},u);u=this.SUPER(arguments).Attributes.call(this,u);if(this.disabled){u.className+=" MathJax_MenuDisabled"}return u},MoveVertical:function(u,E,w){var x=c.GetMenuNode(E);var D=[];for(var z=0,C=x.menuItem.items,y;y=C[z];z++){if(!y.hidden){D.push(y)}}var B=g.IndexOf(D,this);if(B===-1){return}var A=D.length;var v=x.childNodes;do{B=g.Mod(w(B),A)}while(D[B].hidden||!v[B].role||v[B].role==="separator");this.Deactivate(E);D[B].Activate(u,v[B])},Up:function(v,u){this.MoveVertical(v,u,function(w){return w-1})},Down:function(v,u){this.MoveVertical(v,u,function(w){return w+1})},Right:function(v,u){this.MoveHorizontal(v,u,g.Right,!this.isRTL())},Left:function(v,u){this.MoveHorizontal(v,u,g.Left,this.isRTL())},MoveHorizontal:function(A,z,u,B){var x=c.GetMenuNode(z);if(x.menuItem===g.menu&&A.shiftKey){u(A,z)}if(B){return}if(x.menuItem!==g.menu){this.Deactivate(z)}var v=x.previousSibling.childNodes;var y=v.length;while(y--){var w=v[y];if(w.menuItem.submenu&&w.menuItem.submenu===x.menuItem){g.Focus(w);break}}this.RemoveSubmenus(z)},Space:function(u,v){this.Mouseup(u,v)},Activate:function(u,v){this.Deactivate(v);if(!this.disabled){v.className+=" MathJax_MenuActive"}this.DeactivateSubmenus(v);g.Focus(v)},Deactivate:function(u){u.className=u.className.replace(/ MathJax_MenuActive/,"")}});g.ITEM.COMMAND=g.ENTRY.Subclass({action:function(){},Init:function(u,w,v){if(!i(u)){u=[u,u]}this.name=u;this.action=w;this.With(v)},Label:function(u,v){return[this.Name()]},Mouseup:function(u,v){if(!this.disabled){this.Remove(u,v);d.Post(["command",this]);this.action.call(this,u)}return n(u)}});g.ITEM.SUBMENU=g.ENTRY.Subclass({submenu:null,marker:"\u25BA",markerRTL:"\u25C4",Attributes:function(u){u=f.Insert({"aria-haspopup":"true"},u);u=this.SUPER(arguments).Attributes.call(this,u);return u},Init:function(u,w){if(!i(u)){u=[u,u]}this.name=u;var v=1;if(!(w instanceof g.ITEM)){this.With(w),v++}this.submenu=g.apply(g,[].slice.call(arguments,v))},Label:function(u,v){this.submenu.posted=false;return[this.Name()+" ",["span",{className:"MathJax_MenuArrow"+this.rtlClass()},[this.isRTL()?this.markerRTL:this.marker]]]},Timer:function(u,v){this.ClearTimer();u={type:u.type,clientX:u.clientX,clientY:u.clientY};this.timer=setTimeout(e(["Mouseup",this,u,v]),s.delay)},ClearTimer:function(){if(this.timer){clearTimeout(this.timer)}},Touchend:function(v,x){var w=this.submenu.posted;var u=this.SUPER(arguments).Touchend.apply(this,arguments);if(w){this.Deactivate(x);delete c.lastItem;delete c.lastMenu}return u},Mouseout:function(u,v){if(!this.submenu.posted){this.Deactivate(v)}this.ClearTimer()},Mouseover:function(u,v){this.Activate(u,v)},Mouseup:function(u,v){if(!this.disabled){if(!this.submenu.posted){this.ClearTimer();this.submenu.Post(u,v,this.ltr);g.Focus(v)}else{this.DeactivateSubmenus(v)}}return n(u)},Activate:function(u,v){if(!this.disabled){this.Deactivate(v);v.className+=" MathJax_MenuActive"}if(!this.submenu.posted){this.DeactivateSubmenus(v);if(!g.isMobile){this.Timer(u,v)}}g.Focus(v)},MoveVertical:function(w,v,u){this.ClearTimer();this.SUPER(arguments).MoveVertical.apply(this,arguments)},MoveHorizontal:function(w,y,v,x){if(!x){this.SUPER(arguments).MoveHorizontal.apply(this,arguments);return}if(this.disabled){return}if(!this.submenu.posted){this.Activate(w,y);return}var u=c.GetMenuNode(y).nextSibling.childNodes;if(u.length>0){this.submenu.items[0].Activate(w,u[0])}}});g.ITEM.RADIO=g.ENTRY.Subclass({variable:null,marker:(a?"\u25CF":"\u2713"),role:"menuitemradio",Attributes:function(v){var u=s.settings[this.variable]===this.value?"true":"false";v=f.Insert({"aria-checked":u},v);v=this.SUPER(arguments).Attributes.call(this,v);return v},Init:function(v,u,w){if(!i(v)){v=[v,v]}this.name=v;this.variable=u;this.With(w);if(this.value==null){this.value=this.name[0]}},Label:function(v,w){var u={className:"MathJax_MenuRadioCheck"+this.rtlClass()};if(s.settings[this.variable]!==this.value){u={style:{display:"none"}}}return[["span",u,[this.marker]]," "+this.Name()]},Mouseup:function(x,y){if(!this.disabled){var z=y.parentNode.childNodes;for(var v=0,u=z.length;v<u;v++){var w=z[v].menuItem;if(w&&w.variable===this.variable){z[v].firstChild.style.display="none"}}y.firstChild.display="";s.settings[this.variable]=this.value;g.cookie[this.variable]=s.settings[this.variable];g.saveCookie();d.Post(["radio button",this])}this.Remove(x,y);if(this.action&&!this.disabled){this.action.call(g,this)}return n(x)}});g.ITEM.CHECKBOX=g.ENTRY.Subclass({variable:null,marker:"\u2713",role:"menuitemcheckbox",Attributes:function(v){var u=s.settings[this.variable]?"true":"false";v=f.Insert({"aria-checked":u},v);v=this.SUPER(arguments).Attributes.call(this,v);return v},Init:function(v,u,w){if(!i(v)){v=[v,v]}this.name=v;this.variable=u;this.With(w)},Label:function(v,w){var u={className:"MathJax_MenuCheck"+this.rtlClass()};if(!s.settings[this.variable]){u={style:{display:"none"}}}return[["span",u,[this.marker]]," "+this.Name()]},Mouseup:function(u,v){if(!this.disabled){v.firstChild.display=(s.settings[this.variable]?"none":"");s.settings[this.variable]=!s.settings[this.variable];g.cookie[this.variable]=s.settings[this.variable];g.saveCookie();d.Post(["checkbox",this])}this.Remove(u,v);if(this.action&&!this.disabled){this.action.call(g,this)}return n(u)}});g.ITEM.LABEL=g.ENTRY.Subclass({role:"menuitem",Init:function(u,v){if(!i(u)){u=[u,u]}this.name=u;this.With(v)},Label:function(u,v){u.className+=" MathJax_MenuLabel";return[this.Name()]},Activate:function(u,v){this.Deactivate(v);g.Focus(v)},Mouseup:function(u,v){}});g.ITEM.RULE=g.ITEM.Subclass({role:"separator",Attributes:function(u){u=f.Insert({"aria-orientation":"vertical"},u);u=this.SUPER(arguments).Attributes.call(this,u);return u},Label:function(u,v){u.className+=" MathJax_MenuRule";return null}});g.About=function(y){var v=g.About.GetFont();var A=g.About.GetFormat();var u=["MathJax.js v"+MathJax.fileversion,["br"]];u.push(["div",{style:{"border-top":"groove 2px",margin:".25em 0"}}]);g.About.GetJax(u,MathJax.InputJax,["InputJax","%1 Input Jax v%2"]);g.About.GetJax(u,MathJax.OutputJax,["OutputJax","%1 Output Jax v%2"]);g.About.GetJax(u,MathJax.ElementJax,["ElementJax","%1 Element Jax v%2"]);u.push(["div",{style:{"border-top":"groove 2px",margin:".25em 0"}}]);g.About.GetJax(u,MathJax.Extension,["Extension","%1 Extension v%2"],true);u.push(["div",{style:{"border-top":"groove 2px",margin:".25em 0"}}],["center",{},[f.Browser+" v"+f.Browser.version+(A?" \u2014 "+t(A.replace(/ /g,""),A):"")]]);g.About.div=g.Background(g.About);var x=o.addElement(g.About.div,"div",{id:"MathJax_About",tabIndex:0,onkeydown:g.About.Keydown},[["b",{style:{fontSize:"120%"}},["MathJax"]]," v"+MathJax.version,["br"],t(v.replace(/ /g,""),"using "+v),["br"],["br"],["span",{style:{display:"inline-block","text-align":"left","font-size":"80%","max-height":"20em",overflow:"auto","background-color":"#E4E4E4",padding:".4em .6em",border:"1px inset"},tabIndex:0},u],["br"],["br"],["a",{href:"http://www.mathjax.org/"},["www.mathjax.org"]],["span",{className:"MathJax_MenuClose",id:"MathJax_AboutClose",onclick:g.About.Remove,onkeydown:g.About.Keydown,tabIndex:0,role:"button","aria-label":t("CloseAboutDialog","Close about MathJax dialog")},[["span",{},"\u00D7"]]]]);if(y.type==="mouseup"){x.className+=" MathJax_MousePost"}x.focus();MathJax.Localization.setCSS(x);var z=(document.documentElement||{});var w=window.innerHeight||z.clientHeight||z.scrollHeight||0;if(g.prototype.msieAboutBug){x.style.width="20em";x.style.position="absolute";x.style.left=Math.floor((document.documentElement.scrollWidth-x.offsetWidth)/2)+"px";x.style.top=(Math.floor((w-x.offsetHeight)/3)+document.body.scrollTop)+"px"}else{x.style.marginLeft=Math.floor(-x.offsetWidth/2)+"px";x.style.top=Math.floor((w-x.offsetHeight)/3)+"px"}};g.About.Remove=function(u){if(g.About.div){document.body.removeChild(g.About.div);delete g.About.div}};g.About.Keydown=function(u){if(u.keyCode===b.ESCAPE||(this.id==="MathJax_AboutClose"&&(u.keyCode===b.SPACE||u.keyCode===b.RETURN))){g.About.Remove(u);g.CurrentNode().focus();n(u)}},g.About.GetJax=function(v,A,y,x){var z=[];for(var B in A){if(A.hasOwnProperty(B)&&A[B]){if((x&&A[B].version)||(A[B].isa&&A[B].isa(A))){z.push(t(y[0],y[1],(A[B].id||B),A[B].version))}}}z.sort();for(var w=0,u=z.length;w<u;w++){v.push(z[w],["br"])}return v};g.About.GetFont=function(){var u=MathJax.Hub.outputJax["jax/mml"][0]||{};var v={SVG:"web SVG",CommonHTML:"web TeX","HTML-CSS":(u.imgFonts?"image":(u.webFonts?"web":"local")+" "+u.fontInUse)}[u.id]||"generic";return v+" fonts"};g.About.GetFormat=function(){var u=MathJax.Hub.outputJax["jax/mml"][0]||{};if(u.id!=="HTML-CSS"||!u.webFonts||u.imgFonts){return}return u.allowWebFonts.replace(/otf/,"woff or otf")+" fonts"};g.Help=function(u){q.Require("[MathJax]/extensions/HelpDialog.js",function(){MathJax.Extension.Help.Dialog({type:u.type})})};g.ShowSource=function(y){if(!y){y=window.event}var x={screenX:y.screenX,screenY:y.screenY};if(!g.jax){return}if(this.format==="MathML"){var v=MathJax.ElementJax.mml;if(v&&typeof(v.mbase.prototype.toMathML)!=="undefined"){try{g.ShowSource.Text(g.jax.root.toMathML("",g.jax),y)}catch(w){if(!w.restart){throw w}e.After([this,g.ShowSource,x],w.restart)}}else{if(!q.loadingToMathML){q.loadingToMathML=true;g.ShowSource.Window(y);e.Queue(q.Require("[MathJax]/extensions/toMathML.js"),function(){delete q.loadingToMathML;if(!v.mbase.prototype.toMathML){v.mbase.prototype.toMathML=function(){}}},[this,g.ShowSource,x]);return}}}else{if(this.format==="Error"){g.ShowSource.Text(g.jax.errorText,y)}else{if(s.semanticsAnnotations[this.format]){var u=g.jax.root.getAnnotation(this.format);if(u.data[0]){g.ShowSource.Text(u.data[0].toString())}}else{if(g.jax.originalText==null){alert(t("NoOriginalForm","No original form available"));return}g.ShowSource.Text(g.jax.originalText,y)}}}};g.ShowSource.Window=function(v){if(!g.ShowSource.w){var w=[],u=s.windowSettings;for(var x in u){if(u.hasOwnProperty(x)){w.push(x+"="+u[x])}}g.ShowSource.w=window.open("","_blank",w.join(","))}return g.ShowSource.w};g.ShowSource.Text=function(z,x){var u=g.ShowSource.Window(x);delete g.ShowSource.w;z=z.replace(/^\s*/,"").replace(/\s*$/,"");z=z.replace(/&/g,"&amp;").replace(/</g,"&lt;").replace(/>/g,"&gt;");var y=t("EqSource","MathJax Equation Source");if(g.isMobile){u.document.open();u.document.write("<html><head><meta name='viewport' content='width=device-width, initial-scale=1.0' /><title>"+y+"</title></head><body style='font-size:85%'>");u.document.write("<pre>"+z+"</pre>");u.document.write("<hr><input type='button' value='"+t("Close","Close")+"' onclick='window.close()' />");u.document.write("</body></html>");u.document.close()}else{u.document.open();u.document.write("<html><head><title>"+y+"</title></head><body style='font-size:85%'>");u.document.write("<table><tr><td><pre>"+z+"</pre></td></tr></table>");u.document.write("</body></html>");u.document.close();var v=u.document.body.firstChild;setTimeout(function(){var B=(u.outerHeight-u.innerHeight)||30,A=(u.outerWidth-u.innerWidth)||30,w,E;A=Math.max(140,Math.min(Math.floor(0.5*screen.width),v.offsetWidth+A+25));B=Math.max(40,Math.min(Math.floor(0.5*screen.height),v.offsetHeight+B+25));if(g.prototype.msieHeightBug){B+=35}u.resizeTo(A,B);var D;try{D=x.screenX}catch(C){}if(x&&D!=null){w=Math.max(0,Math.min(x.screenX-Math.floor(A/2),screen.width-A-20));E=Math.max(0,Math.min(x.screenY-Math.floor(B/2),screen.height-B-20));u.moveTo(w,E)}},50)}};g.Scale=function(){var z=["CommonHTML","HTML-CSS","SVG","NativeMML","PreviewHTML"],u=z.length,y=100,w,v;for(w=0;w<u;w++){v=r[z[w]];if(v){y=v.config.scale;break}}var x=prompt(t("ScaleMath","Scale all mathematics (compared to surrounding text) by"),y+"%");if(x){if(x.match(/^\s*\d+(\.\d*)?\s*%?\s*$/)){x=parseFloat(x);if(x){if(x!==y){for(w=0;w<u;w++){v=r[z[w]];if(v){v.config.scale=x}}g.cookie.scale=f.config.scale=x;g.saveCookie();f.Queue(["Rerender",f])}}else{alert(t("NonZeroScale","The scale should not be zero"))}}else{alert(t("PercentScale","The scale should be a percentage (e.g., 120%%)"))}}};g.Zoom=function(){if(!MathJax.Extension.MathZoom){q.Require("[MathJax]/extensions/MathZoom.js")}};g.Renderer=function(){var v=f.outputJax["jax/mml"];if(v[0]!==s.settings.renderer){var y=f.Browser,x,u=g.Renderer.Messages,w;switch(s.settings.renderer){case"NativeMML":if(!s.settings.warnedMML){if(y.isChrome&&y.version.substr(0,3)!=="24."){x=u.MML.WebKit}else{if(y.isSafari&&!y.versionAtLeast("5.0")){x=u.MML.WebKit}else{if(y.isMSIE){if(!y.hasMathPlayer){x=u.MML.MSIE}}else{if(y.isEdge){x=u.MML.WebKit}else{x=u.MML[y]}}}}w="warnedMML"}break;case"SVG":if(!s.settings.warnedSVG){if(y.isMSIE&&!m){x=u.SVG.MSIE}}break}if(x){x=t(x[0],x[1]);x+="\n\n";x+=t("SwitchAnyway","Switch the renderer anyway?\n\n(Press OK to switch, CANCEL to continue with the current renderer)");g.cookie.renderer=v[0].id;g.saveCookie();if(!confirm(x)){g.cookie.renderer=s.settings.renderer=o.Cookie.Get("menu").renderer;g.saveCookie();return}if(w){g.cookie.warned=s.settings.warned=true}g.cookie.renderer=s.settings.renderer;g.saveCookie()}f.Queue(["setRenderer",f,s.settings.renderer,"jax/mml"],["Rerender",f])}};g.Renderer.Messages={MML:{WebKit:["WebkitNativeMMLWarning","Your browser doesn't seem to support MathML natively, so switching to MathML output may cause the mathematics on the page to become unreadable."],MSIE:["MSIENativeMMLWarning","Internet Explorer requires the MathPlayer plugin in order to process MathML output."],Opera:["OperaNativeMMLWarning","Opera's support for MathML is limited, so switching to MathML output may cause some expressions to render poorly."],Safari:["SafariNativeMMLWarning","Your browser's native MathML does not implement all the features used by MathJax, so some expressions may not render properly."],Firefox:["FirefoxNativeMMLWarning","Your browser's native MathML does not implement all the features used by MathJax, so some expressions may not render properly."]},SVG:{MSIE:["MSIESVGWarning","SVG is not implemented in Internet Explorer prior to IE9 or when it is emulating IE8 or below. Switching to SVG output will cause the mathematics to not display properly."]}};g.AssistiveMML=function(w,u){var v=MathJax.Extension.AssistiveMML;if(!v){if(!u){q.Require("[MathJax]/extensions/AssistiveMML.js",["AssistiveMML",g,w,true])}return}MathJax.Hub.Queue([(s.settings.assistiveMML?"Add":"Remove")+"AssistiveMathML",v])};g.Font=function(){var u=r["HTML-CSS"];if(!u){return}document.location.reload()};g.Locale=function(){MathJax.Localization.setLocale(s.settings.locale);MathJax.Hub.Queue(["Reprocess",MathJax.Hub])};g.LoadLocale=function(){var u=prompt(t("LoadURL","Load translation data from this URL:"));if(u){if(!u.match(/\.js$/)){alert(t("BadURL","The URL should be for a javascript file that defines MathJax translation data.  Javascript file names should end with '.js'"))}q.Require(u,function(v){if(v!=q.STATUS.OK){alert(t("BadData","Failed to load translation data from %1",u))}})}};g.MPEvents=function(w){var v=s.settings.discoverable,u=g.MPEvents.Messages;if(!m){if(s.settings.mpMouse&&!confirm(t.apply(t,u.IE8warning))){delete g.cookie.mpContext;delete s.settings.mpContext;delete g.cookie.mpMouse;delete s.settings.mpMouse;g.saveCookie();return}s.settings.mpContext=s.settings.mpMouse;g.cookie.mpContext=g.cookie.mpMouse=s.settings.mpMouse;g.saveCookie();MathJax.Hub.Queue(["Rerender",MathJax.Hub])}else{if(!v&&w.name[1]==="Menu Events"&&s.settings.mpContext){alert(t.apply(t,u.IE9warning))}}};g.MPEvents.Messages={IE8warning:["IE8warning","This will disable the MathJax menu and zoom features, but you can Alt-Click on an expression to obtain the MathJax menu instead.\n\nReally change the MathPlayer settings?"],IE9warning:["IE9warning","The MathJax contextual menu will be disabled, but you can Alt-Click on an expression to obtain the MathJax menu instead."]};f.Browser.Select({MSIE:function(u){var v=(document.compatMode==="BackCompat");var w=u.versionAtLeast("8.0")&&document.documentMode>7;g.Augment({margin:20,msieBackgroundBug:((document.documentMode||0)<9),msieFixedPositionBug:(v||!w),msieAboutBug:v,msieHeightBug:((document.documentMode||0)<9)});if(m){delete s.styles["#MathJax_About"].filter;delete s.styles[".MathJax_Menu"].filter}},Firefox:function(u){g.skipMouseover=u.isMobile&&u.versionAtLeast("6.0");g.skipMousedown=u.isMobile}});g.isMobile=f.Browser.isMobile;g.noContextMenu=f.Browser.noContextMenu;g.CreateLocaleMenu=function(){if(!g.menu){return}var z=g.menu.Find("Language").submenu,w=z.items;var v=[],B=MathJax.Localization.strings;for(var A in B){if(B.hasOwnProperty(A)){v.push(A)}}v=v.sort();z.items=[];for(var x=0,u=v.length;x<u;x++){var y=B[v[x]].menuTitle;if(y){y+=" ("+v[x]+")"}else{y=v[x]}z.items.push(c.RADIO([v[x],y],"locale",{action:g.Locale}))}z.items.push(w[w.length-2],w[w.length-1])};g.CreateAnnotationMenu=function(){if(!g.menu){return}var w=g.menu.Find("Show Math As","Annotation").submenu;var v=s.semanticsAnnotations;for(var u in v){if(v.hasOwnProperty(u)){w.items.push(c.COMMAND([u,u],g.ShowSource,{hidden:true,nativeTouch:true,format:u}))}}};f.Register.StartupHook("End Config",function(){s.settings=f.config.menuSettings;if(typeof(s.settings.showRenderer)!=="undefined"){s.showRenderer=s.settings.showRenderer}if(typeof(s.settings.showFontMenu)!=="undefined"){s.showFontMenu=s.settings.showFontMenu}if(typeof(s.settings.showContext)!=="undefined"){s.showContext=s.settings.showContext}g.getCookie();g.menu=g(c.SUBMENU(["Show","Show Math As"],c.COMMAND(["MathMLcode","MathML Code"],g.ShowSource,{nativeTouch:true,format:"MathML"}),c.COMMAND(["Original","Original Form"],g.ShowSource,{nativeTouch:true}),c.SUBMENU(["Annotation","Annotation"],{disabled:true}),c.RULE(),c.CHECKBOX(["texHints","Show TeX hints in MathML"],"texHints"),c.CHECKBOX(["semantics","Add original form as annotation"],"semantics")),c.RULE(),c.SUBMENU(["Settings","Math Settings"],c.SUBMENU(["ZoomTrigger","Zoom Trigger"],c.RADIO(["Hover","Hover"],"zoom",{action:g.Zoom}),c.RADIO(["Click","Click"],"zoom",{action:g.Zoom}),c.RADIO(["DoubleClick","Double-Click"],"zoom",{action:g.Zoom}),c.RADIO(["NoZoom","No Zoom"],"zoom",{value:"None"}),c.RULE(),c.LABEL(["TriggerRequires","Trigger Requires:"]),c.CHECKBOX((f.Browser.isMac?["Option","Option"]:["Alt","Alt"]),"ALT"),c.CHECKBOX(["Command","Command"],"CMD",{hidden:!f.Browser.isMac}),c.CHECKBOX(["Control","Control"],"CTRL",{hidden:f.Browser.isMac}),c.CHECKBOX(["Shift","Shift"],"Shift")),c.SUBMENU(["ZoomFactor","Zoom Factor"],c.RADIO("125%","zscale"),c.RADIO("133%","zscale"),c.RADIO("150%","zscale"),c.RADIO("175%","zscale"),c.RADIO("200%","zscale"),c.RADIO("250%","zscale"),c.RADIO("300%","zscale"),c.RADIO("400%","zscale")),c.RULE(),c.SUBMENU(["Renderer","Math Renderer"],{hidden:!s.showRenderer},c.RADIO(["HTML-CSS","HTML-CSS"],"renderer",{action:g.Renderer}),c.RADIO(["CommonHTML","Common HTML"],"renderer",{action:g.Renderer,value:"CommonHTML"}),c.RADIO(["PreviewHTML","Preview HTML"],"renderer",{action:g.Renderer,value:"PreviewHTML"}),c.RADIO(["MathML","MathML"],"renderer",{action:g.Renderer,value:"NativeMML"}),c.RADIO(["SVG","SVG"],"renderer",{action:g.Renderer}),c.RADIO(["PlainSource","Plain Source"],"renderer",{action:g.Renderer,value:"PlainSource"}),c.RULE(),c.CHECKBOX(["FastPreview","Fast Preview"],"FastPreview")),c.SUBMENU("MathPlayer",{hidden:!f.Browser.isMSIE||!s.showMathPlayer,disabled:!f.Browser.hasMathPlayer},c.LABEL(["MPHandles","Let MathPlayer Handle:"]),c.CHECKBOX(["MenuEvents","Menu Events"],"mpContext",{action:g.MPEvents,hidden:!m}),c.CHECKBOX(["MouseEvents","Mouse Events"],"mpMouse",{action:g.MPEvents,hidden:!m}),c.CHECKBOX(["MenuAndMouse","Mouse and Menu Events"],"mpMouse",{action:g.MPEvents,hidden:m})),c.SUBMENU(["FontPrefs","Font Preference"],{hidden:!s.showFontMenu},c.LABEL(["ForHTMLCSS","For HTML-CSS:"]),c.RADIO(["Auto","Auto"],"font",{action:g.Font}),c.RULE(),c.RADIO(["TeXLocal","TeX (local)"],"font",{action:g.Font}),c.RADIO(["TeXWeb","TeX (web)"],"font",{action:g.Font}),c.RADIO(["TeXImage","TeX (image)"],"font",{action:g.Font}),c.RULE(),c.RADIO(["STIXLocal","STIX (local)"],"font",{action:g.Font}),c.RADIO(["STIXWeb","STIX (web)"],"font",{action:g.Font}),c.RULE(),c.RADIO(["AsanaMathWeb","Asana Math (web)"],"font",{action:g.Font}),c.RADIO(["GyrePagellaWeb","Gyre Pagella (web)"],"font",{action:g.Font}),c.RADIO(["GyreTermesWeb","Gyre Termes (web)"],"font",{action:g.Font}),c.RADIO(["LatinModernWeb","Latin Modern (web)"],"font",{action:g.Font}),c.RADIO(["NeoEulerWeb","Neo Euler (web)"],"font",{action:g.Font})),c.SUBMENU(["ContextMenu","Contextual Menu"],{hidden:!s.showContext},c.RADIO(["MathJax","MathJax"],"context"),c.RADIO(["Browser","Browser"],"context")),c.COMMAND(["Scale","Scale All Math ..."],g.Scale),c.RULE().With({hidden:!s.showDiscoverable,name:["","discover_rule"]}),c.CHECKBOX(["Discoverable","Highlight on Hover"],"discoverable",{hidden:!s.showDiscoverable})),c.SUBMENU(["Accessibility","Accessibility"],c.CHECKBOX(["AssistiveMML","Assistive MathML"],"assistiveMML",{action:g.AssistiveMML}),c.CHECKBOX(["InTabOrder","Include in Tab Order"],"inTabOrder")),c.SUBMENU(["Locale","Language"],{hidden:!s.showLocale,ltr:true},c.RADIO("en","locale",{action:g.Locale}),c.RULE().With({hidden:!s.showLocaleURL,name:["","localURL_rule"]}),c.COMMAND(["LoadLocale","Load from URL ..."],g.LoadLocale,{hidden:!s.showLocaleURL})),c.RULE(),c.COMMAND(["About","About MathJax"],g.About),c.COMMAND(["Help","MathJax Help"],g.Help));if(g.isMobile){(function(){var v=s.settings;var u=g.menu.Find("Math Settings","Zoom Trigger").submenu;u.items[0].disabled=u.items[1].disabled=true;if(v.zoom==="Hover"||v.zoom=="Click"){v.zoom="None"}u.items=u.items.slice(0,4);if(navigator.appVersion.match(/[ (]Android[) ]/)){g.ITEM.SUBMENU.Augment({marker:"\u00BB"})}})()}g.CreateLocaleMenu();g.CreateAnnotationMenu()});g.showRenderer=function(u){g.cookie.showRenderer=s.showRenderer=u;g.saveCookie();g.menu.Find("Math Settings","Math Renderer").hidden=!u};g.showMathPlayer=function(u){g.cookie.showMathPlayer=s.showMathPlayer=u;g.saveCookie();g.menu.Find("Math Settings","MathPlayer").hidden=!u};g.showFontMenu=function(u){g.cookie.showFontMenu=s.showFontMenu=u;g.saveCookie();g.menu.Find("Math Settings","Font Preference").hidden=!u};g.showContext=function(u){g.cookie.showContext=s.showContext=u;g.saveCookie();g.menu.Find("Math Settings","Contextual Menu").hidden=!u};g.showDiscoverable=function(u){g.cookie.showDiscoverable=s.showDiscoverable=u;g.saveCookie();g.menu.Find("Math Settings","Highlight on Hover").hidden=!u;g.menu.Find("Math Settings","discover_rule").hidden=!u};g.showLocale=function(u){g.cookie.showLocale=s.showLocale=u;g.saveCookie();g.menu.Find("Language").hidden=!u};MathJax.Hub.Register.StartupHook("HTML-CSS Jax Ready",function(){if(!MathJax.OutputJax["HTML-CSS"].config.imageFont){g.menu.Find("Math Settings","Font Preference","TeX (image)").disabled=true}});e.Queue(f.Register.StartupHook("End Config",{}),["Styles",q,s.styles],["Post",f.Startup.signal,"MathMenu Ready"],["loadComplete",q,"[MathJax]/extensions/MathMenu.js"])})(MathJax.Hub,MathJax.HTML,MathJax.Ajax,MathJax.CallBack,MathJax.OutputJax);
MathJax.ElementJax.mml=MathJax.ElementJax({mimeType:"jax/mml"},{id:"mml",version:"2.7.0",directory:MathJax.ElementJax.directory+"/mml",extensionDir:MathJax.ElementJax.extensionDir+"/mml",optableDir:MathJax.ElementJax.directory+"/mml/optable"});MathJax.ElementJax.mml.Augment({Init:function(){if(arguments.length===1&&arguments[0].type==="math"){this.root=arguments[0]}else{this.root=MathJax.ElementJax.mml.math.apply(this,arguments)}if(this.root.attr&&this.root.attr.mode){if(!this.root.display&&this.root.attr.mode==="display"){this.root.display="block";this.root.attrNames.push("display")}delete this.root.attr.mode;for(var b=0,a=this.root.attrNames.length;b<a;b++){if(this.root.attrNames[b]==="mode"){this.root.attrNames.splice(b,1);break}}}}},{INHERIT:"_inherit_",AUTO:"_auto_",SIZE:{INFINITY:"infinity",SMALL:"small",NORMAL:"normal",BIG:"big"},COLOR:{TRANSPARENT:"transparent"},VARIANT:{NORMAL:"normal",BOLD:"bold",ITALIC:"italic",BOLDITALIC:"bold-italic",DOUBLESTRUCK:"double-struck",FRAKTUR:"fraktur",BOLDFRAKTUR:"bold-fraktur",SCRIPT:"script",BOLDSCRIPT:"bold-script",SANSSERIF:"sans-serif",BOLDSANSSERIF:"bold-sans-serif",SANSSERIFITALIC:"sans-serif-italic",SANSSERIFBOLDITALIC:"sans-serif-bold-italic",MONOSPACE:"monospace",INITIAL:"inital",TAILED:"tailed",LOOPED:"looped",STRETCHED:"stretched",CALIGRAPHIC:"-tex-caligraphic",OLDSTYLE:"-tex-oldstyle"},FORM:{PREFIX:"prefix",INFIX:"infix",POSTFIX:"postfix"},LINEBREAK:{AUTO:"auto",NEWLINE:"newline",NOBREAK:"nobreak",GOODBREAK:"goodbreak",BADBREAK:"badbreak"},LINEBREAKSTYLE:{BEFORE:"before",AFTER:"after",DUPLICATE:"duplicate",INFIXLINBREAKSTYLE:"infixlinebreakstyle"},INDENTALIGN:{LEFT:"left",CENTER:"center",RIGHT:"right",AUTO:"auto",ID:"id",INDENTALIGN:"indentalign"},INDENTSHIFT:{INDENTSHIFT:"indentshift"},LINETHICKNESS:{THIN:"thin",MEDIUM:"medium",THICK:"thick"},NOTATION:{LONGDIV:"longdiv",ACTUARIAL:"actuarial",RADICAL:"radical",BOX:"box",ROUNDEDBOX:"roundedbox",CIRCLE:"circle",LEFT:"left",RIGHT:"right",TOP:"top",BOTTOM:"bottom",UPDIAGONALSTRIKE:"updiagonalstrike",DOWNDIAGONALSTRIKE:"downdiagonalstrike",UPDIAGONALARROW:"updiagonalarrow",VERTICALSTRIKE:"verticalstrike",HORIZONTALSTRIKE:"horizontalstrike",PHASORANGLE:"phasorangle",MADRUWB:"madruwb"},ALIGN:{TOP:"top",BOTTOM:"bottom",CENTER:"center",BASELINE:"baseline",AXIS:"axis",LEFT:"left",RIGHT:"right"},LINES:{NONE:"none",SOLID:"solid",DASHED:"dashed"},SIDE:{LEFT:"left",RIGHT:"right",LEFTOVERLAP:"leftoverlap",RIGHTOVERLAP:"rightoverlap"},WIDTH:{AUTO:"auto",FIT:"fit"},ACTIONTYPE:{TOGGLE:"toggle",STATUSLINE:"statusline",TOOLTIP:"tooltip",INPUT:"input"},LENGTH:{VERYVERYTHINMATHSPACE:"veryverythinmathspace",VERYTHINMATHSPACE:"verythinmathspace",THINMATHSPACE:"thinmathspace",MEDIUMMATHSPACE:"mediummathspace",THICKMATHSPACE:"thickmathspace",VERYTHICKMATHSPACE:"verythickmathspace",VERYVERYTHICKMATHSPACE:"veryverythickmathspace",NEGATIVEVERYVERYTHINMATHSPACE:"negativeveryverythinmathspace",NEGATIVEVERYTHINMATHSPACE:"negativeverythinmathspace",NEGATIVETHINMATHSPACE:"negativethinmathspace",NEGATIVEMEDIUMMATHSPACE:"negativemediummathspace",NEGATIVETHICKMATHSPACE:"negativethickmathspace",NEGATIVEVERYTHICKMATHSPACE:"negativeverythickmathspace",NEGATIVEVERYVERYTHICKMATHSPACE:"negativeveryverythickmathspace"},OVERFLOW:{LINBREAK:"linebreak",SCROLL:"scroll",ELIDE:"elide",TRUNCATE:"truncate",SCALE:"scale"},UNIT:{EM:"em",EX:"ex",PX:"px",IN:"in",CM:"cm",MM:"mm",PT:"pt",PC:"pc"},TEXCLASS:{ORD:0,OP:1,BIN:2,REL:3,OPEN:4,CLOSE:5,PUNCT:6,INNER:7,VCENTER:8,NONE:-1},TEXCLASSNAMES:["ORD","OP","BIN","REL","OPEN","CLOSE","PUNCT","INNER","VCENTER"],skipAttributes:{texClass:true,useHeight:true,texprimestyle:true},copyAttributes:{displaystyle:1,scriptlevel:1,open:1,close:1,form:1,actiontype:1,fontfamily:true,fontsize:true,fontweight:true,fontstyle:true,color:true,background:true,id:true,"class":1,href:true,style:true},copyAttributeNames:["displaystyle","scriptlevel","open","close","form","actiontype","fontfamily","fontsize","fontweight","fontstyle","color","background","id","class","href","style"],nocopyAttributes:{fontfamily:true,fontsize:true,fontweight:true,fontstyle:true,color:true,background:true,id:true,"class":true,href:true,style:true,xmlns:true},Error:function(d,e){var c=this.merror(d),b=MathJax.Localization.fontDirection(),a=MathJax.Localization.fontFamily();if(e){c=c.With(e)}if(b||a){c=this.mstyle(c);if(b){c.dir=b}if(a){c.style.fontFamily="font-family: "+a}}return c}});(function(a){a.mbase=MathJax.Object.Subclass({type:"base",isToken:false,defaults:{mathbackground:a.INHERIT,mathcolor:a.INHERIT,dir:a.INHERIT},noInherit:{},noInheritAttribute:{texClass:true},getRemoved:{},linebreakContainer:false,Init:function(){this.data=[];if(this.inferRow&&!(arguments.length===1&&arguments[0].inferred)){this.Append(a.mrow().With({inferred:true,notParent:true}))}this.Append.apply(this,arguments)},With:function(e){for(var f in e){if(e.hasOwnProperty(f)){this[f]=e[f]}}return this},Append:function(){if(this.inferRow&&this.data.length){this.data[0].Append.apply(this.data[0],arguments)}else{for(var f=0,e=arguments.length;f<e;f++){this.SetData(this.data.length,arguments[f])}}},SetData:function(e,f){if(f!=null){if(!(f instanceof a.mbase)){f=(this.isToken||this.isChars?a.chars(f):a.mtext(f))}f.parent=this;f.setInherit(this.inheritFromMe?this:this.inherit)}this.data[e]=f},Parent:function(){var e=this.parent;while(e&&e.notParent){e=e.parent}return e},Get:function(f,k,l){if(!l){if(this[f]!=null){return this[f]}if(this.attr&&this.attr[f]!=null){return this.attr[f]}}var g=this.Parent();if(g&&g["adjustChild_"+f]!=null){return(g["adjustChild_"+f])(this.childPosition(),k)}var j=this.inherit;var e=j;while(j){var i=j[f];if(i==null&&j.attr){i=j.attr[f]}if(j.removedStyles&&j.getRemoved[f]&&i==null){i=j.removedStyles[j.getRemoved[f]]}if(i!=null&&j.noInheritAttribute&&!j.noInheritAttribute[f]){var h=j.noInherit[this.type];if(!(h&&h[f])){return i}}e=j;j=j.inherit}if(!k){if(this.defaults[f]===a.AUTO){return this.autoDefault(f)}if(this.defaults[f]!==a.INHERIT&&this.defaults[f]!=null){return this.defaults[f]}if(e){return e.defaults[f]}}return null},hasValue:function(e){return(this.Get(e,true)!=null)},getValues:function(){var f={};for(var g=0,e=arguments.length;g<e;g++){f[arguments[g]]=this.Get(arguments[g])}return f},adjustChild_scriptlevel:function(f,e){return this.Get("scriptlevel",e)},adjustChild_displaystyle:function(f,e){return this.Get("displaystyle",e)},adjustChild_texprimestyle:function(f,e){return this.Get("texprimestyle",e)},childPosition:function(){var h=this,g=h.parent;while(g.notParent){h=g;g=h.parent}for(var f=0,e=g.data.length;f<e;f++){if(g.data[f]===h){return f}}return null},setInherit:function(g){if(g!==this.inherit&&this.inherit==null){this.inherit=g;for(var f=0,e=this.data.length;f<e;f++){if(this.data[f]&&this.data[f].setInherit){this.data[f].setInherit(g)}}}},setTeXclass:function(e){this.getPrevClass(e);return(typeof(this.texClass)!=="undefined"?this:e)},getPrevClass:function(e){if(e){this.prevClass=e.Get("texClass");this.prevLevel=e.Get("scriptlevel")}},updateTeXclass:function(e){if(e){this.prevClass=e.prevClass;delete e.prevClass;this.prevLevel=e.prevLevel;delete e.prevLevel;this.texClass=e.Get("texClass")}},texSpacing:function(){var f=(this.prevClass!=null?this.prevClass:a.TEXCLASS.NONE);var e=(this.Get("texClass")||a.TEXCLASS.ORD);if(f===a.TEXCLASS.NONE||e===a.TEXCLASS.NONE){return""}if(f===a.TEXCLASS.VCENTER){f=a.TEXCLASS.ORD}if(e===a.TEXCLASS.VCENTER){e=a.TEXCLASS.ORD}var g=this.TEXSPACE[f][e];if((this.prevLevel>0||this.Get("scriptlevel")>0)&&g>=0){return""}return this.TEXSPACELENGTH[Math.abs(g)]},TEXSPACELENGTH:["",a.LENGTH.THINMATHSPACE,a.LENGTH.MEDIUMMATHSPACE,a.LENGTH.THICKMATHSPACE],TEXSPACE:[[0,-1,2,3,0,0,0,1],[-1,-1,0,3,0,0,0,1],[2,2,0,0,2,0,0,2],[3,3,0,0,3,0,0,3],[0,0,0,0,0,0,0,0],[0,-1,2,3,0,0,0,1],[1,1,0,1,1,1,1,1],[1,-1,2,3,1,0,1,1]],autoDefault:function(e){return""},isSpacelike:function(){return false},isEmbellished:function(){return false},Core:function(){return this},CoreMO:function(){return this},childIndex:function(g){if(g==null){return}for(var f=0,e=this.data.length;f<e;f++){if(g===this.data[f]){return f}}},CoreIndex:function(){return(this.inferRow?this.data[0]||this:this).childIndex(this.Core())},hasNewline:function(){if(this.isEmbellished()){return this.CoreMO().hasNewline()}if(this.isToken||this.linebreakContainer){return false}for(var f=0,e=this.data.length;f<e;f++){if(this.data[f]&&this.data[f].hasNewline()){return true}}return false},array:function(){if(this.inferred){return this.data}else{return[this]}},toString:function(){return this.type+"("+this.data.join(",")+")"},getAnnotation:function(){return null}},{childrenSpacelike:function(){for(var f=0,e=this.data.length;f<e;f++){if(!this.data[f].isSpacelike()){return false}}return true},childEmbellished:function(){return(this.data[0]&&this.data[0].isEmbellished())},childCore:function(){return(this.inferRow&&this.data[0]?this.data[0].Core():this.data[0])},childCoreMO:function(){return(this.data[0]?this.data[0].CoreMO():null)},setChildTeXclass:function(e){if(this.data[0]){e=this.data[0].setTeXclass(e);this.updateTeXclass(this.data[0])}return e},setBaseTeXclasses:function(g){this.getPrevClass(g);this.texClass=null;if(this.data[0]){if(this.isEmbellished()||this.data[0].isa(a.mi)){g=this.data[0].setTeXclass(g);this.updateTeXclass(this.Core())}else{this.data[0].setTeXclass();g=this}}else{g=this}for(var f=1,e=this.data.length;f<e;f++){if(this.data[f]){this.data[f].setTeXclass()}}return g},setSeparateTeXclasses:function(g){this.getPrevClass(g);for(var f=0,e=this.data.length;f<e;f++){if(this.data[f]){this.data[f].setTeXclass()}}if(this.isEmbellished()){this.updateTeXclass(this.Core())}return this}});a.mi=a.mbase.Subclass({type:"mi",isToken:true,texClass:a.TEXCLASS.ORD,defaults:{mathvariant:a.AUTO,mathsize:a.INHERIT,mathbackground:a.INHERIT,mathcolor:a.INHERIT,dir:a.INHERIT},autoDefault:function(f){if(f==="mathvariant"){var e=(this.data[0]||"").toString();return(e.length===1||(e.length===2&&e.charCodeAt(0)>=55296&&e.charCodeAt(0)<56320)?a.VARIANT.ITALIC:a.VARIANT.NORMAL)}return""},setTeXclass:function(f){this.getPrevClass(f);var e=this.data.join("");if(e.length>1&&e.match(/^[a-z][a-z0-9]*$/i)&&this.texClass===a.TEXCLASS.ORD){this.texClass=a.TEXCLASS.OP;this.autoOP=true}return this}});a.mn=a.mbase.Subclass({type:"mn",isToken:true,texClass:a.TEXCLASS.ORD,defaults:{mathvariant:a.INHERIT,mathsize:a.INHERIT,mathbackground:a.INHERIT,mathcolor:a.INHERIT,dir:a.INHERIT}});a.mo=a.mbase.Subclass({type:"mo",isToken:true,defaults:{mathvariant:a.INHERIT,mathsize:a.INHERIT,mathbackground:a.INHERIT,mathcolor:a.INHERIT,dir:a.INHERIT,form:a.AUTO,fence:a.AUTO,separator:a.AUTO,lspace:a.AUTO,rspace:a.AUTO,stretchy:a.AUTO,symmetric:a.AUTO,maxsize:a.AUTO,minsize:a.AUTO,largeop:a.AUTO,movablelimits:a.AUTO,accent:a.AUTO,linebreak:a.LINEBREAK.AUTO,lineleading:a.INHERIT,linebreakstyle:a.AUTO,linebreakmultchar:a.INHERIT,indentalign:a.INHERIT,indentshift:a.INHERIT,indenttarget:a.INHERIT,indentalignfirst:a.INHERIT,indentshiftfirst:a.INHERIT,indentalignlast:a.INHERIT,indentshiftlast:a.INHERIT,texClass:a.AUTO},defaultDef:{form:a.FORM.INFIX,fence:false,separator:false,lspace:a.LENGTH.THICKMATHSPACE,rspace:a.LENGTH.THICKMATHSPACE,stretchy:false,symmetric:false,maxsize:a.SIZE.INFINITY,minsize:"0em",largeop:false,movablelimits:false,accent:false,linebreak:a.LINEBREAK.AUTO,lineleading:"1ex",linebreakstyle:"before",indentalign:a.INDENTALIGN.AUTO,indentshift:"0",indenttarget:"",indentalignfirst:a.INDENTALIGN.INDENTALIGN,indentshiftfirst:a.INDENTSHIFT.INDENTSHIFT,indentalignlast:a.INDENTALIGN.INDENTALIGN,indentshiftlast:a.INDENTSHIFT.INDENTSHIFT,texClass:a.TEXCLASS.REL},SPACE_ATTR:{lspace:1,rspace:2,form:4},useMMLspacing:7,autoDefault:function(g,n){var l=this.def;if(!l){if(g==="form"){this.useMMLspacing&=~this.SPACE_ATTR.form;return this.getForm()}var k=this.data.join("");var f=[this.Get("form"),a.FORM.INFIX,a.FORM.POSTFIX,a.FORM.PREFIX];for(var h=0,e=f.length;h<e;h++){var j=this.OPTABLE[f[h]][k];if(j){l=this.makeDef(j);break}}if(!l){l=this.CheckRange(k)}if(!l&&n){l={}}else{if(!l){l=MathJax.Hub.Insert({},this.defaultDef)}if(this.parent){this.def=l}else{l=MathJax.Hub.Insert({},l)}l.form=f[0]}}this.useMMLspacing&=~(this.SPACE_ATTR[g]||0);if(l[g]!=null){return l[g]}else{if(!n){return this.defaultDef[g]}}return""},CheckRange:function(j){var k=j.charCodeAt(0);if(k>=55296&&k<56320){k=(((k-55296)<<10)+(j.charCodeAt(1)-56320))+65536}for(var g=0,e=this.RANGES.length;g<e&&this.RANGES[g][0]<=k;g++){if(k<=this.RANGES[g][1]){if(this.RANGES[g][3]){var f=a.optableDir+"/"+this.RANGES[g][3]+".js";this.RANGES[g][3]=null;MathJax.Hub.RestartAfter(MathJax.Ajax.Require(f))}var h=a.TEXCLASSNAMES[this.RANGES[g][2]];h=this.OPTABLE.infix[j]=a.mo.OPTYPES[h==="BIN"?"BIN3":h];return this.makeDef(h)}}return null},makeDef:function(f){if(f[2]==null){f[2]=this.defaultDef.texClass}if(!f[3]){f[3]={}}var e=MathJax.Hub.Insert({},f[3]);e.lspace=this.SPACE[f[0]];e.rspace=this.SPACE[f[1]];e.texClass=f[2];if(e.texClass===a.TEXCLASS.REL&&(this.movablelimits||this.data.join("").match(/^[a-z]+$/i))){e.texClass=a.TEXCLASS.OP}return e},getForm:function(){var e=this,g=this.parent,f=this.Parent();while(f&&f.isEmbellished()){e=g;g=f.parent;f=f.Parent()}if(g&&g.type==="mrow"&&g.NonSpaceLength()!==1){if(g.FirstNonSpace()===e){return a.FORM.PREFIX}if(g.LastNonSpace()===e){return a.FORM.POSTFIX}}return a.FORM.INFIX},isEmbellished:function(){return true},hasNewline:function(){return(this.Get("linebreak")===a.LINEBREAK.NEWLINE)},CoreParent:function(){var e=this;while(e&&e.isEmbellished()&&e.CoreMO()===this&&!e.isa(a.math)){e=e.Parent()}return e},CoreText:function(e){if(!e){return""}if(e.isEmbellished()){return e.CoreMO().data.join("")}while((((e.isa(a.mrow)||e.isa(a.TeXAtom)||e.isa(a.mstyle)||e.isa(a.mphantom))&&e.data.length===1)||e.isa(a.munderover))&&e.data[0]){e=e.data[0]}if(!e.isToken){return""}else{return e.data.join("")}},remapChars:{"*":"\u2217",'"':"\u2033","\u00B0":"\u2218","\u00B2":"2","\u00B3":"3","\u00B4":"\u2032","\u00B9":"1"},remap:function(f,e){f=f.replace(/-/g,"\u2212");if(e){f=f.replace(/'/g,"\u2032").replace(/`/g,"\u2035");if(f.length===1){f=e[f]||f}}return f},setTeXclass:function(f){var e=this.getValues("form","lspace","rspace","fence");if(this.useMMLspacing){this.texClass=a.TEXCLASS.NONE;return this}if(e.fence&&!this.texClass){if(e.form===a.FORM.PREFIX){this.texClass=a.TEXCLASS.OPEN}if(e.form===a.FORM.POSTFIX){this.texClass=a.TEXCLASS.CLOSE}}this.texClass=this.Get("texClass");if(this.data.join("")==="\u2061"){if(f){f.texClass=a.TEXCLASS.OP;f.fnOP=true}this.texClass=this.prevClass=a.TEXCLASS.NONE;return f}return this.adjustTeXclass(f)},adjustTeXclass:function(f){if(this.texClass===a.TEXCLASS.NONE){return f}if(f){if(f.autoOP&&(this.texClass===a.TEXCLASS.BIN||this.texClass===a.TEXCLASS.REL)){f.texClass=a.TEXCLASS.ORD}this.prevClass=f.texClass||a.TEXCLASS.ORD;this.prevLevel=f.Get("scriptlevel")}else{this.prevClass=a.TEXCLASS.NONE}if(this.texClass===a.TEXCLASS.BIN&&(this.prevClass===a.TEXCLASS.NONE||this.prevClass===a.TEXCLASS.BIN||this.prevClass===a.TEXCLASS.OP||this.prevClass===a.TEXCLASS.REL||this.prevClass===a.TEXCLASS.OPEN||this.prevClass===a.TEXCLASS.PUNCT)){this.texClass=a.TEXCLASS.ORD}else{if(this.prevClass===a.TEXCLASS.BIN&&(this.texClass===a.TEXCLASS.REL||this.texClass===a.TEXCLASS.CLOSE||this.texClass===a.TEXCLASS.PUNCT)){f.texClass=this.prevClass=a.TEXCLASS.ORD}else{if(this.texClass===a.TEXCLASS.BIN){var g=this,e=this.parent;while(e&&e.parent&&e.isEmbellished()&&(e.data.length===1||(e.type!=="mrow"&&e.Core()===g))){g=e;e=e.parent}if(e.data[e.data.length-1]===g){this.texClass=a.TEXCLASS.ORD}}}}return this}});a.mtext=a.mbase.Subclass({type:"mtext",isToken:true,isSpacelike:function(){return true},texClass:a.TEXCLASS.ORD,defaults:{mathvariant:a.INHERIT,mathsize:a.INHERIT,mathbackground:a.INHERIT,mathcolor:a.INHERIT,dir:a.INHERIT}});a.mspace=a.mbase.Subclass({type:"mspace",isToken:true,isSpacelike:function(){return true},defaults:{mathbackground:a.INHERIT,mathcolor:a.INHERIT,width:"0em",height:"0ex",depth:"0ex",linebreak:a.LINEBREAK.AUTO},hasDimAttr:function(){return(this.hasValue("width")||this.hasValue("height")||this.hasValue("depth"))},hasNewline:function(){return(!this.hasDimAttr()&&this.Get("linebreak")===a.LINEBREAK.NEWLINE)}});a.ms=a.mbase.Subclass({type:"ms",isToken:true,texClass:a.TEXCLASS.ORD,defaults:{mathvariant:a.INHERIT,mathsize:a.INHERIT,mathbackground:a.INHERIT,mathcolor:a.INHERIT,dir:a.INHERIT,lquote:'"',rquote:'"'}});a.mglyph=a.mbase.Subclass({type:"mglyph",isToken:true,texClass:a.TEXCLASS.ORD,defaults:{mathbackground:a.INHERIT,mathcolor:a.INHERIT,alt:"",src:"",width:a.AUTO,height:a.AUTO,valign:"0em"}});a.mrow=a.mbase.Subclass({type:"mrow",isSpacelike:a.mbase.childrenSpacelike,inferred:false,notParent:false,isEmbellished:function(){var f=false;for(var g=0,e=this.data.length;g<e;g++){if(this.data[g]==null){continue}if(this.data[g].isEmbellished()){if(f){return false}f=true;this.core=g}else{if(!this.data[g].isSpacelike()){return false}}}return f},NonSpaceLength:function(){var g=0;for(var f=0,e=this.data.length;f<e;f++){if(this.data[f]&&!this.data[f].isSpacelike()){g++}}return g},FirstNonSpace:function(){for(var f=0,e=this.data.length;f<e;f++){if(this.data[f]&&!this.data[f].isSpacelike()){return this.data[f]}}return null},LastNonSpace:function(){for(var e=this.data.length-1;e>=0;e--){if(this.data[0]&&!this.data[e].isSpacelike()){return this.data[e]}}return null},Core:function(){if(!(this.isEmbellished())||typeof(this.core)==="undefined"){return this}return this.data[this.core]},CoreMO:function(){if(!(this.isEmbellished())||typeof(this.core)==="undefined"){return this}return this.data[this.core].CoreMO()},toString:function(){if(this.inferred){return"["+this.data.join(",")+"]"}return this.SUPER(arguments).toString.call(this)},setTeXclass:function(g){var f,e=this.data.length;if((this.open||this.close)&&(!g||!g.fnOP)){this.getPrevClass(g);g=null;for(f=0;f<e;f++){if(this.data[f]){g=this.data[f].setTeXclass(g)}}if(!this.hasOwnProperty("texClass")){this.texClass=a.TEXCLASS.INNER}return this}else{for(f=0;f<e;f++){if(this.data[f]){g=this.data[f].setTeXclass(g)}}if(this.data[0]){this.updateTeXclass(this.data[0])}return g}},getAnnotation:function(e){if(this.data.length!=1){return null}return this.data[0].getAnnotation(e)}});a.mfrac=a.mbase.Subclass({type:"mfrac",num:0,den:1,linebreakContainer:true,isEmbellished:a.mbase.childEmbellished,Core:a.mbase.childCore,CoreMO:a.mbase.childCoreMO,defaults:{mathbackground:a.INHERIT,mathcolor:a.INHERIT,linethickness:a.LINETHICKNESS.MEDIUM,numalign:a.ALIGN.CENTER,denomalign:a.ALIGN.CENTER,bevelled:false},adjustChild_displaystyle:function(e){return false},adjustChild_scriptlevel:function(f){var e=this.Get("scriptlevel");if(!this.Get("displaystyle")||e>0){e++}return e},adjustChild_texprimestyle:function(e){if(e==this.den){return true}return this.Get("texprimestyle")},setTeXclass:a.mbase.setSeparateTeXclasses});a.msqrt=a.mbase.Subclass({type:"msqrt",inferRow:true,linebreakContainer:true,texClass:a.TEXCLASS.ORD,setTeXclass:a.mbase.setSeparateTeXclasses,adjustChild_texprimestyle:function(e){return true}});a.mroot=a.mbase.Subclass({type:"mroot",linebreakContainer:true,texClass:a.TEXCLASS.ORD,adjustChild_displaystyle:function(e){if(e===1){return false}return this.Get("displaystyle")},adjustChild_scriptlevel:function(f){var e=this.Get("scriptlevel");if(f===1){e+=2}return e},adjustChild_texprimestyle:function(e){if(e===0){return true}return this.Get("texprimestyle")},setTeXclass:a.mbase.setSeparateTeXclasses});a.mstyle=a.mbase.Subclass({type:"mstyle",isSpacelike:a.mbase.childrenSpacelike,isEmbellished:a.mbase.childEmbellished,Core:a.mbase.childCore,CoreMO:a.mbase.childCoreMO,inferRow:true,defaults:{scriptlevel:a.INHERIT,displaystyle:a.INHERIT,scriptsizemultiplier:Math.sqrt(1/2),scriptminsize:"8pt",mathbackground:a.INHERIT,mathcolor:a.INHERIT,dir:a.INHERIT,infixlinebreakstyle:a.LINEBREAKSTYLE.BEFORE,decimalseparator:"."},adjustChild_scriptlevel:function(g){var f=this.scriptlevel;if(f==null){f=this.Get("scriptlevel")}else{if(String(f).match(/^ *[-+]/)){var e=this.Get("scriptlevel",null,true);f=e+parseInt(f)}}return f},inheritFromMe:true,noInherit:{mpadded:{width:true,height:true,depth:true,lspace:true,voffset:true},mtable:{width:true,height:true,depth:true,align:true}},getRemoved:{fontfamily:"fontFamily",fontweight:"fontWeight",fontstyle:"fontStyle",fontsize:"fontSize"},setTeXclass:a.mbase.setChildTeXclass});a.merror=a.mbase.Subclass({type:"merror",inferRow:true,linebreakContainer:true,texClass:a.TEXCLASS.ORD});a.mpadded=a.mbase.Subclass({type:"mpadded",inferRow:true,isSpacelike:a.mbase.childrenSpacelike,isEmbellished:a.mbase.childEmbellished,Core:a.mbase.childCore,CoreMO:a.mbase.childCoreMO,defaults:{mathbackground:a.INHERIT,mathcolor:a.INHERIT,width:"",height:"",depth:"",lspace:0,voffset:0},setTeXclass:a.mbase.setChildTeXclass});a.mphantom=a.mbase.Subclass({type:"mphantom",texClass:a.TEXCLASS.ORD,inferRow:true,isSpacelike:a.mbase.childrenSpacelike,isEmbellished:a.mbase.childEmbellished,Core:a.mbase.childCore,CoreMO:a.mbase.childCoreMO,setTeXclass:a.mbase.setChildTeXclass});a.mfenced=a.mbase.Subclass({type:"mfenced",defaults:{mathbackground:a.INHERIT,mathcolor:a.INHERIT,open:"(",close:")",separators:","},addFakeNodes:function(){var f=this.getValues("open","close","separators");f.open=f.open.replace(/[ \t\n\r]/g,"");f.close=f.close.replace(/[ \t\n\r]/g,"");f.separators=f.separators.replace(/[ \t\n\r]/g,"");if(f.open!==""){this.SetData("open",a.mo(f.open).With({fence:true,form:a.FORM.PREFIX,texClass:a.TEXCLASS.OPEN}));this.data.open.useMMLspacing=0}if(f.separators!==""){while(f.separators.length<this.data.length){f.separators+=f.separators.charAt(f.separators.length-1)}for(var g=1,e=this.data.length;g<e;g++){if(this.data[g]){this.SetData("sep"+g,a.mo(f.separators.charAt(g-1)).With({separator:true}));this.data["sep"+g].useMMLspacing=0}}}if(f.close!==""){this.SetData("close",a.mo(f.close).With({fence:true,form:a.FORM.POSTFIX,texClass:a.TEXCLASS.CLOSE}));this.data.close.useMMLspacing=0}},texClass:a.TEXCLASS.OPEN,setTeXclass:function(g){this.addFakeNodes();this.getPrevClass(g);if(this.data.open){g=this.data.open.setTeXclass(g)}if(this.data[0]){g=this.data[0].setTeXclass(g)}for(var f=1,e=this.data.length;f<e;f++){if(this.data["sep"+f]){g=this.data["sep"+f].setTeXclass(g)}if(this.data[f]){g=this.data[f].setTeXclass(g)}}if(this.data.close){g=this.data.close.setTeXclass(g)}this.updateTeXclass(this.data.open);this.texClass=a.TEXCLASS.INNER;return g}});a.menclose=a.mbase.Subclass({type:"menclose",inferRow:true,linebreakContainer:true,defaults:{mathbackground:a.INHERIT,mathcolor:a.INHERIT,notation:a.NOTATION.LONGDIV,texClass:a.TEXCLASS.ORD},setTeXclass:a.mbase.setSeparateTeXclasses});a.msubsup=a.mbase.Subclass({type:"msubsup",base:0,sub:1,sup:2,isEmbellished:a.mbase.childEmbellished,Core:a.mbase.childCore,CoreMO:a.mbase.childCoreMO,defaults:{mathbackground:a.INHERIT,mathcolor:a.INHERIT,subscriptshift:"",superscriptshift:"",texClass:a.AUTO},autoDefault:function(e){if(e==="texClass"){return(this.isEmbellished()?this.CoreMO().Get(e):a.TEXCLASS.ORD)}return 0},adjustChild_displaystyle:function(e){if(e>0){return false}return this.Get("displaystyle")},adjustChild_scriptlevel:function(f){var e=this.Get("scriptlevel");if(f>0){e++}return e},adjustChild_texprimestyle:function(e){if(e===this.sub){return true}return this.Get("texprimestyle")},setTeXclass:a.mbase.setBaseTeXclasses});a.msub=a.msubsup.Subclass({type:"msub"});a.msup=a.msubsup.Subclass({type:"msup",sub:2,sup:1});a.mmultiscripts=a.msubsup.Subclass({type:"mmultiscripts",adjustChild_texprimestyle:function(e){if(e%2===1){return true}return this.Get("texprimestyle")}});a.mprescripts=a.mbase.Subclass({type:"mprescripts"});a.none=a.mbase.Subclass({type:"none"});a.munderover=a.mbase.Subclass({type:"munderover",base:0,under:1,over:2,sub:1,sup:2,ACCENTS:["","accentunder","accent"],linebreakContainer:true,isEmbellished:a.mbase.childEmbellished,Core:a.mbase.childCore,CoreMO:a.mbase.childCoreMO,defaults:{mathbackground:a.INHERIT,mathcolor:a.INHERIT,accent:a.AUTO,accentunder:a.AUTO,align:a.ALIGN.CENTER,texClass:a.AUTO,subscriptshift:"",superscriptshift:""},autoDefault:function(e){if(e==="texClass"){return(this.isEmbellished()?this.CoreMO().Get(e):a.TEXCLASS.ORD)}if(e==="accent"&&this.data[this.over]){return this.data[this.over].CoreMO().Get("accent")}if(e==="accentunder"&&this.data[this.under]){return this.data[this.under].CoreMO().Get("accent")}return false},adjustChild_displaystyle:function(e){if(e>0){return false}return this.Get("displaystyle")},adjustChild_scriptlevel:function(g){var f=this.Get("scriptlevel");var e=(this.data[this.base]&&!this.Get("displaystyle")&&this.data[this.base].CoreMO().Get("movablelimits"));if(g==this.under&&(e||!this.Get("accentunder"))){f++}if(g==this.over&&(e||!this.Get("accent"))){f++}return f},adjustChild_texprimestyle:function(e){if(e===this.base&&this.data[this.over]){return true}return this.Get("texprimestyle")},setTeXclass:a.mbase.setBaseTeXclasses});a.munder=a.munderover.Subclass({type:"munder"});a.mover=a.munderover.Subclass({type:"mover",over:1,under:2,sup:1,sub:2,ACCENTS:["","accent","accentunder"]});a.mtable=a.mbase.Subclass({type:"mtable",defaults:{mathbackground:a.INHERIT,mathcolor:a.INHERIT,align:a.ALIGN.AXIS,rowalign:a.ALIGN.BASELINE,columnalign:a.ALIGN.CENTER,groupalign:"{left}",alignmentscope:true,columnwidth:a.WIDTH.AUTO,width:a.WIDTH.AUTO,rowspacing:"1ex",columnspacing:".8em",rowlines:a.LINES.NONE,columnlines:a.LINES.NONE,frame:a.LINES.NONE,framespacing:"0.4em 0.5ex",equalrows:false,equalcolumns:false,displaystyle:false,side:a.SIDE.RIGHT,minlabelspacing:"0.8em",texClass:a.TEXCLASS.ORD,useHeight:1},adjustChild_displaystyle:function(){return(this.displaystyle!=null?this.displaystyle:this.defaults.displaystyle)},inheritFromMe:true,noInherit:{mover:{align:true},munder:{align:true},munderover:{align:true},mtable:{align:true,rowalign:true,columnalign:true,groupalign:true,alignmentscope:true,columnwidth:true,width:true,rowspacing:true,columnspacing:true,rowlines:true,columnlines:true,frame:true,framespacing:true,equalrows:true,equalcolumns:true,displaystyle:true,side:true,minlabelspacing:true,texClass:true,useHeight:1}},linebreakContainer:true,Append:function(){for(var f=0,e=arguments.length;f<e;f++){if(!((arguments[f] instanceof a.mtr)||(arguments[f] instanceof a.mlabeledtr))){arguments[f]=a.mtr(arguments[f])}}this.SUPER(arguments).Append.apply(this,arguments)},setTeXclass:a.mbase.setSeparateTeXclasses});a.mtr=a.mbase.Subclass({type:"mtr",defaults:{mathbackground:a.INHERIT,mathcolor:a.INHERIT,rowalign:a.INHERIT,columnalign:a.INHERIT,groupalign:a.INHERIT},inheritFromMe:true,noInherit:{mrow:{rowalign:true,columnalign:true,groupalign:true},mtable:{rowalign:true,columnalign:true,groupalign:true}},linebreakContainer:true,Append:function(){for(var f=0,e=arguments.length;f<e;f++){if(!(arguments[f] instanceof a.mtd)){arguments[f]=a.mtd(arguments[f])}}this.SUPER(arguments).Append.apply(this,arguments)},setTeXclass:a.mbase.setSeparateTeXclasses});a.mtd=a.mbase.Subclass({type:"mtd",inferRow:true,linebreakContainer:true,isEmbellished:a.mbase.childEmbellished,Core:a.mbase.childCore,CoreMO:a.mbase.childCoreMO,defaults:{mathbackground:a.INHERIT,mathcolor:a.INHERIT,rowspan:1,columnspan:1,rowalign:a.INHERIT,columnalign:a.INHERIT,groupalign:a.INHERIT},setTeXclass:a.mbase.setSeparateTeXclasses});a.maligngroup=a.mbase.Subclass({type:"maligngroup",isSpacelike:function(){return true},defaults:{mathbackground:a.INHERIT,mathcolor:a.INHERIT,groupalign:a.INHERIT},inheritFromMe:true,noInherit:{mrow:{groupalign:true},mtable:{groupalign:true}}});a.malignmark=a.mbase.Subclass({type:"malignmark",defaults:{mathbackground:a.INHERIT,mathcolor:a.INHERIT,edge:a.SIDE.LEFT},isSpacelike:function(){return true}});a.mlabeledtr=a.mtr.Subclass({type:"mlabeledtr"});a.maction=a.mbase.Subclass({type:"maction",defaults:{mathbackground:a.INHERIT,mathcolor:a.INHERIT,actiontype:a.ACTIONTYPE.TOGGLE,selection:1},selected:function(){return this.data[this.Get("selection")-1]||a.NULL},isEmbellished:function(){return this.selected().isEmbellished()},isSpacelike:function(){return this.selected().isSpacelike()},Core:function(){return this.selected().Core()},CoreMO:function(){return this.selected().CoreMO()},setTeXclass:function(f){if(this.Get("actiontype")===a.ACTIONTYPE.TOOLTIP&&this.data[1]){this.data[1].setTeXclass()}var e=this.selected();f=e.setTeXclass(f);this.updateTeXclass(e);return f}});a.semantics=a.mbase.Subclass({type:"semantics",notParent:true,isEmbellished:a.mbase.childEmbellished,Core:a.mbase.childCore,CoreMO:a.mbase.childCoreMO,defaults:{definitionURL:null,encoding:null},setTeXclass:a.mbase.setChildTeXclass,getAnnotation:function(g){var l=MathJax.Hub.config.MathMenu.semanticsAnnotations[g];if(l){for(var h=0,e=this.data.length;h<e;h++){var k=this.data[h].Get("encoding");if(k){for(var f=0,o=l.length;f<o;f++){if(l[f]===k){return this.data[h]}}}}}return null}});a.annotation=a.mbase.Subclass({type:"annotation",isChars:true,linebreakContainer:true,defaults:{definitionURL:null,encoding:null,cd:"mathmlkeys",name:"",src:null}});a["annotation-xml"]=a.mbase.Subclass({type:"annotation-xml",linebreakContainer:true,defaults:{definitionURL:null,encoding:null,cd:"mathmlkeys",name:"",src:null}});a.math=a.mstyle.Subclass({type:"math",defaults:{mathvariant:a.VARIANT.NORMAL,mathsize:a.SIZE.NORMAL,mathcolor:"",mathbackground:a.COLOR.TRANSPARENT,dir:"ltr",scriptlevel:0,displaystyle:a.AUTO,display:"inline",maxwidth:"",overflow:a.OVERFLOW.LINEBREAK,altimg:"","altimg-width":"","altimg-height":"","altimg-valign":"",alttext:"",cdgroup:"",scriptsizemultiplier:Math.sqrt(1/2),scriptminsize:"8px",infixlinebreakstyle:a.LINEBREAKSTYLE.BEFORE,lineleading:"1ex",indentshift:"auto",indentalign:a.INDENTALIGN.AUTO,indentalignfirst:a.INDENTALIGN.INDENTALIGN,indentshiftfirst:a.INDENTSHIFT.INDENTSHIFT,indentalignlast:a.INDENTALIGN.INDENTALIGN,indentshiftlast:a.INDENTSHIFT.INDENTSHIFT,decimalseparator:".",texprimestyle:false},autoDefault:function(e){if(e==="displaystyle"){return this.Get("display")==="block"}return""},linebreakContainer:true,setTeXclass:a.mbase.setChildTeXclass,getAnnotation:function(e){if(this.data.length!=1){return null}return this.data[0].getAnnotation(e)}});a.chars=a.mbase.Subclass({type:"chars",Append:function(){this.data.push.apply(this.data,arguments)},value:function(){return this.data.join("")},toString:function(){return this.data.join("")}});a.entity=a.mbase.Subclass({type:"entity",Append:function(){this.data.push.apply(this.data,arguments)},value:function(){if(this.data[0].substr(0,2)==="#x"){return parseInt(this.data[0].substr(2),16)}else{if(this.data[0].substr(0,1)==="#"){return parseInt(this.data[0].substr(1))}else{return 0}}},toString:function(){var e=this.value();if(e<=65535){return String.fromCharCode(e)}e-=65536;return String.fromCharCode((e>>10)+55296)+String.fromCharCode((e&1023)+56320)}});a.xml=a.mbase.Subclass({type:"xml",Init:function(){this.div=document.createElement("div");return this.SUPER(arguments).Init.apply(this,arguments)},Append:function(){for(var f=0,e=arguments.length;f<e;f++){var g=this.Import(arguments[f]);this.data.push(g);this.div.appendChild(g)}},Import:function(j){if(document.importNode){return document.importNode(j,true)}var f,g,e;if(j.nodeType===1){f=document.createElement(j.nodeName);for(g=0,e=j.attributes.length;g<e;g++){var h=j.attributes[g];if(h.specified&&h.nodeValue!=null&&h.nodeValue!=""){f.setAttribute(h.nodeName,h.nodeValue)}if(h.nodeName==="style"){f.style.cssText=h.nodeValue}}if(j.className){f.className=j.className}}else{if(j.nodeType===3||j.nodeType===4){f=document.createTextNode(j.nodeValue)}else{if(j.nodeType===8){f=document.createComment(j.nodeValue)}else{return document.createTextNode("")}}}for(g=0,e=j.childNodes.length;g<e;g++){f.appendChild(this.Import(j.childNodes[g]))}return f},value:function(){return this.div},toString:function(){return this.div.innerHTML}});a.TeXAtom=a.mbase.Subclass({type:"texatom",linebreakContainer:true,inferRow:true,notParent:true,texClass:a.TEXCLASS.ORD,Core:a.mbase.childCore,CoreMO:a.mbase.childCoreMO,isEmbellished:a.mbase.childEmbellished,setTeXclass:function(e){this.data[0].setTeXclass();return this.adjustTeXclass(e)},adjustTeXclass:a.mo.prototype.adjustTeXclass});a.NULL=a.mbase().With({type:"null"});var b=a.TEXCLASS;var d={ORD:[0,0,b.ORD],ORD11:[1,1,b.ORD],ORD21:[2,1,b.ORD],ORD02:[0,2,b.ORD],ORD55:[5,5,b.ORD],OP:[1,2,b.OP,{largeop:true,movablelimits:true,symmetric:true}],OPFIXED:[1,2,b.OP,{largeop:true,movablelimits:true}],INTEGRAL:[0,1,b.OP,{largeop:true,symmetric:true}],INTEGRAL2:[1,2,b.OP,{largeop:true,symmetric:true}],BIN3:[3,3,b.BIN],BIN4:[4,4,b.BIN],BIN01:[0,1,b.BIN],BIN5:[5,5,b.BIN],TALLBIN:[4,4,b.BIN,{stretchy:true}],BINOP:[4,4,b.BIN,{largeop:true,movablelimits:true}],REL:[5,5,b.REL],REL1:[1,1,b.REL,{stretchy:true}],REL4:[4,4,b.REL],RELSTRETCH:[5,5,b.REL,{stretchy:true}],RELACCENT:[5,5,b.REL,{accent:true}],WIDEREL:[5,5,b.REL,{accent:true,stretchy:true}],OPEN:[0,0,b.OPEN,{fence:true,stretchy:true,symmetric:true}],CLOSE:[0,0,b.CLOSE,{fence:true,stretchy:true,symmetric:true}],INNER:[0,0,b.INNER],PUNCT:[0,3,b.PUNCT],ACCENT:[0,0,b.ORD,{accent:true}],WIDEACCENT:[0,0,b.ORD,{accent:true,stretchy:true}]};a.mo.Augment({SPACE:["0em","0.1111em","0.1667em","0.2222em","0.2667em","0.3333em"],RANGES:[[32,127,b.REL,"BasicLatin"],[160,255,b.ORD,"Latin1Supplement"],[256,383,b.ORD],[384,591,b.ORD],[688,767,b.ORD,"SpacingModLetters"],[768,879,b.ORD,"CombDiacritMarks"],[880,1023,b.ORD,"GreekAndCoptic"],[7680,7935,b.ORD],[8192,8303,b.PUNCT,"GeneralPunctuation"],[8304,8351,b.ORD],[8352,8399,b.ORD],[8400,8447,b.ORD,"CombDiactForSymbols"],[8448,8527,b.ORD,"LetterlikeSymbols"],[8528,8591,b.ORD],[8592,8703,b.REL,"Arrows"],[8704,8959,b.BIN,"MathOperators"],[8960,9215,b.ORD,"MiscTechnical"],[9312,9471,b.ORD],[9472,9631,b.ORD],[9632,9727,b.ORD,"GeometricShapes"],[9984,10175,b.ORD,"Dingbats"],[10176,10223,b.ORD,"MiscMathSymbolsA"],[10224,10239,b.REL,"SupplementalArrowsA"],[10496,10623,b.REL,"SupplementalArrowsB"],[10624,10751,b.ORD,"MiscMathSymbolsB"],[10752,11007,b.BIN,"SuppMathOperators"],[11008,11263,b.ORD,"MiscSymbolsAndArrows"],[119808,120831,b.ORD]],OPTABLE:{prefix:{"\u2200":d.ORD21,"\u2202":d.ORD21,"\u2203":d.ORD21,"\u2207":d.ORD21,"\u220F":d.OP,"\u2210":d.OP,"\u2211":d.OP,"\u2212":d.BIN01,"\u2213":d.BIN01,"\u221A":[1,1,b.ORD,{stretchy:true}],"\u2220":d.ORD,"\u222B":d.INTEGRAL,"\u222E":d.INTEGRAL,"\u22C0":d.OP,"\u22C1":d.OP,"\u22C2":d.OP,"\u22C3":d.OP,"\u2308":d.OPEN,"\u230A":d.OPEN,"\u27E8":d.OPEN,"\u27EE":d.OPEN,"\u2A00":d.OP,"\u2A01":d.OP,"\u2A02":d.OP,"\u2A04":d.OP,"\u2A06":d.OP,"\u00AC":d.ORD21,"\u00B1":d.BIN01,"(":d.OPEN,"+":d.BIN01,"-":d.BIN01,"[":d.OPEN,"{":d.OPEN,"|":d.OPEN},postfix:{"!":[1,0,b.CLOSE],"&":d.ORD,"\u2032":d.ORD02,"\u203E":d.WIDEACCENT,"\u2309":d.CLOSE,"\u230B":d.CLOSE,"\u23DE":d.WIDEACCENT,"\u23DF":d.WIDEACCENT,"\u266D":d.ORD02,"\u266E":d.ORD02,"\u266F":d.ORD02,"\u27E9":d.CLOSE,"\u27EF":d.CLOSE,"\u02C6":d.WIDEACCENT,"\u02C7":d.WIDEACCENT,"\u02C9":d.WIDEACCENT,"\u02CA":d.ACCENT,"\u02CB":d.ACCENT,"\u02D8":d.ACCENT,"\u02D9":d.ACCENT,"\u02DC":d.WIDEACCENT,"\u0302":d.WIDEACCENT,"\u00A8":d.ACCENT,"\u00AF":d.WIDEACCENT,")":d.CLOSE,"]":d.CLOSE,"^":d.WIDEACCENT,_:d.WIDEACCENT,"`":d.ACCENT,"|":d.CLOSE,"}":d.CLOSE,"~":d.WIDEACCENT},infix:{"":d.ORD,"%":[3,3,b.ORD],"\u2022":d.BIN4,"\u2026":d.INNER,"\u2044":d.TALLBIN,"\u2061":d.ORD,"\u2062":d.ORD,"\u2063":[0,0,b.ORD,{linebreakstyle:"after",separator:true}],"\u2064":d.ORD,"\u2190":d.WIDEREL,"\u2191":d.RELSTRETCH,"\u2192":d.WIDEREL,"\u2193":d.RELSTRETCH,"\u2194":d.WIDEREL,"\u2195":d.RELSTRETCH,"\u2196":d.RELSTRETCH,"\u2197":d.RELSTRETCH,"\u2198":d.RELSTRETCH,"\u2199":d.RELSTRETCH,"\u21A6":d.WIDEREL,"\u21A9":d.WIDEREL,"\u21AA":d.WIDEREL,"\u21BC":d.WIDEREL,"\u21BD":d.WIDEREL,"\u21C0":d.WIDEREL,"\u21C1":d.WIDEREL,"\u21CC":d.WIDEREL,"\u21D0":d.WIDEREL,"\u21D1":d.RELSTRETCH,"\u21D2":d.WIDEREL,"\u21D3":d.RELSTRETCH,"\u21D4":d.WIDEREL,"\u21D5":d.RELSTRETCH,"\u2208":d.REL,"\u2209":d.REL,"\u220B":d.REL,"\u2212":d.BIN4,"\u2213":d.BIN4,"\u2215":d.TALLBIN,"\u2216":d.BIN4,"\u2217":d.BIN4,"\u2218":d.BIN4,"\u2219":d.BIN4,"\u221D":d.REL,"\u2223":d.REL,"\u2225":d.REL,"\u2227":d.BIN4,"\u2228":d.BIN4,"\u2229":d.BIN4,"\u222A":d.BIN4,"\u223C":d.REL,"\u2240":d.BIN4,"\u2243":d.REL,"\u2245":d.REL,"\u2248":d.REL,"\u224D":d.REL,"\u2250":d.REL,"\u2260":d.REL,"\u2261":d.REL,"\u2264":d.REL,"\u2265":d.REL,"\u226A":d.REL,"\u226B":d.REL,"\u227A":d.REL,"\u227B":d.REL,"\u2282":d.REL,"\u2283":d.REL,"\u2286":d.REL,"\u2287":d.REL,"\u228E":d.BIN4,"\u2291":d.REL,"\u2292":d.REL,"\u2293":d.BIN4,"\u2294":d.BIN4,"\u2295":d.BIN4,"\u2296":d.BIN4,"\u2297":d.BIN4,"\u2298":d.BIN4,"\u2299":d.BIN4,"\u22A2":d.REL,"\u22A3":d.REL,"\u22A4":d.ORD55,"\u22A5":d.REL,"\u22A8":d.REL,"\u22C4":d.BIN4,"\u22C5":d.BIN4,"\u22C6":d.BIN4,"\u22C8":d.REL,"\u22EE":d.ORD55,"\u22EF":d.INNER,"\u22F1":[5,5,b.INNER],"\u25B3":d.BIN4,"\u25B5":d.BIN4,"\u25B9":d.BIN4,"\u25BD":d.BIN4,"\u25BF":d.BIN4,"\u25C3":d.BIN4,"\u2758":d.REL,"\u27F5":d.WIDEREL,"\u27F6":d.WIDEREL,"\u27F7":d.WIDEREL,"\u27F8":d.WIDEREL,"\u27F9":d.WIDEREL,"\u27FA":d.WIDEREL,"\u27FC":d.WIDEREL,"\u2A2F":d.BIN4,"\u2A3F":d.BIN4,"\u2AAF":d.REL,"\u2AB0":d.REL,"\u00B1":d.BIN4,"\u00B7":d.BIN4,"\u00D7":d.BIN4,"\u00F7":d.BIN4,"*":d.BIN3,"+":d.BIN4,",":[0,3,b.PUNCT,{linebreakstyle:"after",separator:true}],"-":d.BIN4,".":[3,3,b.ORD],"/":d.ORD11,":":[1,2,b.REL],";":[0,3,b.PUNCT,{linebreakstyle:"after",separator:true}],"<":d.REL,"=":d.REL,">":d.REL,"?":[1,1,b.CLOSE],"\\":d.ORD,"^":d.ORD11,_:d.ORD11,"|":[2,2,b.ORD,{fence:true,stretchy:true,symmetric:true}],"#":d.ORD,"$":d.ORD,"\u002E":[0,3,b.PUNCT,{separator:true}],"\u02B9":d.ORD,"\u0300":d.ACCENT,"\u0301":d.ACCENT,"\u0303":d.WIDEACCENT,"\u0304":d.ACCENT,"\u0306":d.ACCENT,"\u0307":d.ACCENT,"\u0308":d.ACCENT,"\u030C":d.ACCENT,"\u0332":d.WIDEACCENT,"\u0338":d.REL4,"\u2015":[0,0,b.ORD,{stretchy:true}],"\u2017":[0,0,b.ORD,{stretchy:true}],"\u2020":d.BIN3,"\u2021":d.BIN3,"\u20D7":d.ACCENT,"\u2111":d.ORD,"\u2113":d.ORD,"\u2118":d.ORD,"\u211C":d.ORD,"\u2205":d.ORD,"\u221E":d.ORD,"\u2305":d.BIN3,"\u2306":d.BIN3,"\u2322":d.REL4,"\u2323":d.REL4,"\u2329":d.OPEN,"\u232A":d.CLOSE,"\u23AA":d.ORD,"\u23AF":[0,0,b.ORD,{stretchy:true}],"\u23B0":d.OPEN,"\u23B1":d.CLOSE,"\u2500":d.ORD,"\u25EF":d.BIN3,"\u2660":d.ORD,"\u2661":d.ORD,"\u2662":d.ORD,"\u2663":d.ORD,"\u3008":d.OPEN,"\u3009":d.CLOSE,"\uFE37":d.WIDEACCENT,"\uFE38":d.WIDEACCENT}}},{OPTYPES:d});var c=a.mo.prototype.OPTABLE;c.infix["^"]=d.WIDEREL;c.infix._=d.WIDEREL;c.prefix["\u2223"]=d.OPEN;c.prefix["\u2225"]=d.OPEN;c.postfix["\u2223"]=d.CLOSE;c.postfix["\u2225"]=d.CLOSE})(MathJax.ElementJax.mml);MathJax.ElementJax.mml.loadComplete("jax.js");
MathJax.Hub.Register.LoadHook("[MathJax]/jax/element/mml/jax.js",function(){var c="2.7.0";var a=MathJax.ElementJax.mml,b=MathJax.Hub.config.menuSettings;a.mbase.Augment({toMathML:function(l){var h=(this.inferred&&this.parent.inferRow);if(l==null){l=""}var f=this.type,e=this.toMathMLattributes();if(f==="mspace"){return l+"<"+f+e+" />"}var k=[],j=(this.isToken?"":l+(h?"":"  "));for(var g=0,d=this.data.length;g<d;g++){if(this.data[g]){k.push(this.data[g].toMathML(j))}else{if(!this.isToken&&!this.isChars){k.push(j+"<mrow />")}}}if(this.isToken||this.isChars){return l+"<"+f+e+">"+k.join("")+"</"+f+">"}if(h){return k.join("\n")}if(k.length===0||(k.length===1&&k[0]==="")){return l+"<"+f+e+" />"}return l+"<"+f+e+">\n"+k.join("\n")+"\n"+l+"</"+f+">"},toMathMLattributes:function(){var j=(this.type==="mstyle"?a.math.prototype.defaults:this.defaults);var h=(this.attrNames||a.copyAttributeNames),g=a.skipAttributes,l=a.copyAttributes;var e=[];if(this.type==="math"&&(!this.attr||!this.attr.xmlns)){e.push('xmlns="http://www.w3.org/1998/Math/MathML"')}if(!this.attrNames){for(var k in j){if(!g[k]&&!l[k]&&j.hasOwnProperty(k)){if(this[k]!=null&&this[k]!==j[k]){if(this.Get(k,null,1)!==this[k]){e.push(k+'="'+this.toMathMLattribute(this[k])+'"')}}}}}for(var f=0,d=h.length;f<d;f++){if(l[h[f]]===1&&!j.hasOwnProperty(h[f])){continue}value=(this.attr||{})[h[f]];if(value==null){value=this[h[f]]}if(value!=null){e.push(h[f]+'="'+this.toMathMLquote(value)+'"')}}this.toMathMLclass(e);if(e.length){return" "+e.join(" ")}else{return""}},toMathMLclass:function(d){var f=[];if(this["class"]){f.push(this["class"])}if(this.isa(a.TeXAtom)&&b.texHints){var e=["ORD","OP","BIN","REL","OPEN","CLOSE","PUNCT","INNER","VCENTER"][this.texClass];if(e){f.push("MJX-TeXAtom-"+e);if(e==="OP"&&!this.movablelimits){f.push("MJX-fixedlimits")}}}if(this.mathvariant&&this.toMathMLvariants[this.mathvariant]){f.push("MJX"+this.mathvariant)}if(this.variantForm){f.push("MJX-variant")}if(f.length){d.unshift('class="'+f.join(" ")+'"')}},toMathMLattribute:function(d){if(typeof(d)==="string"&&d.replace(/ /g,"").match(/^(([-+])?(\d+(\.\d*)?|\.\d+))mu$/)){return(RegExp.$2||"")+((1/18)*RegExp.$3).toFixed(3).replace(/\.?0+$/,"")+"em"}else{if(this.toMathMLvariants[d]){return this.toMathMLvariants[d]}}return this.toMathMLquote(d)},toMathMLvariants:{"-tex-caligraphic":a.VARIANT.SCRIPT,"-tex-caligraphic-bold":a.VARIANT.BOLDSCRIPT,"-tex-oldstyle":a.VARIANT.NORMAL,"-tex-oldstyle-bold":a.VARIANT.BOLD,"-tex-mathit":a.VARIANT.ITALIC},toMathMLquote:function(f){f=String(f).split("");for(var g=0,d=f.length;g<d;g++){var k=f[g].charCodeAt(0);if(k<=55295||57344<=k){if(k>126||(k<32&&k!==10&&k!==13&&k!==9)){f[g]="&#x"+k.toString(16).toUpperCase()+";"}else{var j={"&":"&amp;","<":"&lt;",">":"&gt;",'"':"&quot;"}[f[g]];if(j){f[g]=j}}}else{if(g+1<d){var h=f[g+1].charCodeAt(0);var e=(((k-55296)<<10)+(h-56320)+65536);f[g]="&#x"+e.toString(16).toUpperCase()+";";f[g+1]="";g++}else{f[g]=""}}}return f.join("")}});a.math.Augment({toMathML:function(d,e){var g;if(d==null){d=""}if(e&&e.originalText&&b.semantics){g=MathJax.InputJax[e.inputJax].annotationEncoding}var n=(this.data[0]&&this.data[0].data.length>1);var p=this.type,k=this.toMathMLattributes();var j=[],o=d+(g?"  "+(n?"  ":""):"")+"  ";for(var h=0,f=this.data.length;h<f;h++){if(this.data[h]){j.push(this.data[h].toMathML(o))}else{j.push(o+"<mrow />")}}if(j.length===0||(j.length===1&&j[0]==="")){if(!g){return"<"+p+k+" />"}j.push(o+"<mrow />")}if(g){if(n){j.unshift(d+"    <mrow>");j.push(d+"    </mrow>")}j.unshift(d+"  <semantics>");var l=e.originalText.replace(/[&<>]/g,function(i){return{">":"&gt;","<":"&lt;","&":"&amp;"}[i]});j.push(d+'    <annotation encoding="'+g+'">'+l+"</annotation>");j.push(d+"  </semantics>")}return d+"<"+p+k+">\n"+j.join("\n")+"\n"+d+"</"+p+">"}});a.msubsup.Augment({toMathML:function(j){var f=this.type;if(this.data[this.sup]==null){f="msub"}if(this.data[this.sub]==null){f="msup"}var e=this.toMathMLattributes();delete this.data[0].inferred;var h=[];for(var g=0,d=this.data.length;g<d;g++){if(this.data[g]){h.push(this.data[g].toMathML(j+"  "))}}return j+"<"+f+e+">\n"+h.join("\n")+"\n"+j+"</"+f+">"}});a.munderover.Augment({toMathML:function(k){var f=this.type;var j=this.data[this.base];if(j&&j.isa(a.TeXAtom)&&j.movablelimits&&!j.Get("displaystyle")){type="msubsup";if(this.data[this.under]==null){f="msup"}if(this.data[this.over]==null){f="msub"}}else{if(this.data[this.under]==null){f="mover"}if(this.data[this.over]==null){f="munder"}}var e=this.toMathMLattributes();delete this.data[0].inferred;var h=[];for(var g=0,d=this.data.length;g<d;g++){if(this.data[g]){h.push(this.data[g].toMathML(k+"  "))}}return k+"<"+f+e+">\n"+h.join("\n")+"\n"+k+"</"+f+">"}});a.TeXAtom.Augment({toMathML:function(e){var d=this.toMathMLattributes();if(!d&&this.data[0].data.length===1){return e.substr(2)+this.data[0].toMathML(e)}return e+"<mrow"+d+">\n"+this.data[0].toMathML(e+"  ")+"\n"+e+"</mrow>"}});a.chars.Augment({toMathML:function(d){return(d||"")+this.toMathMLquote(this.toString())}});a.entity.Augment({toMathML:function(d){return(d||"")+"&"+this.data[0]+";<!-- "+this.toString()+" -->"}});a.xml.Augment({toMathML:function(d){return(d||"")+this.toString()}});MathJax.Hub.Register.StartupHook("TeX mathchoice Ready",function(){a.TeXmathchoice.Augment({toMathML:function(d){return this.Core().toMathML(d)}})});MathJax.Hub.Startup.signal.Post("toMathML Ready")});MathJax.Ajax.loadComplete("[MathJax]/extensions/toMathML.js");
(function(b,e){var d="2.7.0";var a=b.CombineConfig("TeX.noErrors",{disabled:false,multiLine:true,inlineDelimiters:["",""],style:{"font-size":"90%","text-align":"left",color:"black",padding:"1px 3px",border:"1px solid"}});var c="\u00A0";MathJax.Extension["TeX/noErrors"]={version:d,config:a};b.Register.StartupHook("TeX Jax Ready",function(){var f=MathJax.InputJax.TeX.formatError;MathJax.InputJax.TeX.Augment({formatError:function(j,i,k,g){if(a.disabled){return f.apply(this,arguments)}var h=j.message.replace(/\n.*/,"");b.signal.Post(["TeX Jax - parse error",h,i,k,g]);var m=a.inlineDelimiters;var l=(k||a.multiLine);if(!k){i=m[0]+i+m[1]}if(l){i=i.replace(/ /g,c)}else{i=i.replace(/\n/g," ")}return MathJax.ElementJax.mml.merror(i).With({isError:true,multiLine:l})}})});b.Register.StartupHook("HTML-CSS Jax Config",function(){b.Config({"HTML-CSS":{styles:{".MathJax .noError":b.Insert({"vertical-align":(b.Browser.isMSIE&&a.multiLine?"-2px":"")},a.style)}}})});b.Register.StartupHook("HTML-CSS Jax Ready",function(){var g=MathJax.ElementJax.mml;var h=MathJax.OutputJax["HTML-CSS"];var f=g.math.prototype.toHTML,i=g.merror.prototype.toHTML;g.math.Augment({toHTML:function(j,k){var l=this.data[0];if(l&&l.data[0]&&l.data[0].isError){j.style.fontSize="";j=this.HTMLcreateSpan(j);j.bbox=l.data[0].toHTML(j).bbox}else{j=f.apply(this,arguments)}return j}});g.merror.Augment({toHTML:function(p){if(!this.isError){return i.apply(this,arguments)}p=this.HTMLcreateSpan(p);p.className="noError";if(this.multiLine){p.style.display="inline-block"}var r=this.data[0].data[0].data.join("").split(/\n/);for(var o=0,l=r.length;o<l;o++){h.addText(p,r[o]);if(o!==l-1){h.addElement(p,"br",{isMathJax:true})}}var q=h.getHD(p.parentNode),k=h.getW(p.parentNode);if(l>1){var n=(q.h+q.d)/2,j=h.TeX.x_height/2;p.parentNode.style.verticalAlign=h.Em(q.d+(j-n));q.h=j+n;q.d=n-j}p.bbox={h:q.h,d:q.d,w:k,lw:0,rw:k};return p}})});b.Register.StartupHook("SVG Jax Config",function(){b.Config({SVG:{styles:{".MathJax_SVG .noError":b.Insert({"vertical-align":(b.Browser.isMSIE&&a.multiLine?"-2px":"")},a.style)}}})});b.Register.StartupHook("SVG Jax Ready",function(){var g=MathJax.ElementJax.mml;var f=g.math.prototype.toSVG,h=g.merror.prototype.toSVG;g.math.Augment({toSVG:function(i,j){var k=this.data[0];if(k&&k.data[0]&&k.data[0].isError){i=k.data[0].toSVG(i)}else{i=f.apply(this,arguments)}return i}});g.merror.Augment({toSVG:function(n){if(!this.isError||this.Parent().type!=="math"){return h.apply(this,arguments)}n=e.addElement(n,"span",{className:"noError",isMathJax:true});if(this.multiLine){n.style.display="inline-block"}var o=this.data[0].data[0].data.join("").split(/\n/);for(var l=0,j=o.length;l<j;l++){e.addText(n,o[l]);if(l!==j-1){e.addElement(n,"br",{isMathJax:true})}}if(j>1){var k=n.offsetHeight/2;n.style.verticalAlign=(-k+(k/j))+"px"}return n}})});b.Register.StartupHook("NativeMML Jax Ready",function(){var h=MathJax.ElementJax.mml;var g=MathJax.Extension["TeX/noErrors"].config;var f=h.math.prototype.toNativeMML,i=h.merror.prototype.toNativeMML;h.math.Augment({toNativeMML:function(j){var k=this.data[0];if(k&&k.data[0]&&k.data[0].isError){j=k.data[0].toNativeMML(j)}else{j=f.apply(this,arguments)}return j}});h.merror.Augment({toNativeMML:function(n){if(!this.isError){return i.apply(this,arguments)}n=n.appendChild(document.createElement("span"));var o=this.data[0].data[0].data.join("").split(/\n/);for(var l=0,k=o.length;l<k;l++){n.appendChild(document.createTextNode(o[l]));if(l!==k-1){n.appendChild(document.createElement("br"))}}if(this.multiLine){n.style.display="inline-block";if(k>1){n.style.verticalAlign="middle"}}for(var p in g.style){if(g.style.hasOwnProperty(p)){var j=p.replace(/-./g,function(m){return m.charAt(1).toUpperCase()});n.style[j]=g.style[p]}}return n}})});b.Register.StartupHook("PreviewHTML Jax Config",function(){b.Config({PreviewHTML:{styles:{".MathJax_PHTML .noError":b.Insert({"vertical-align":(b.Browser.isMSIE&&a.multiLine?"-2px":"")},a.style)}}})});b.Register.StartupHook("PreviewHTML Jax Ready",function(){var f=MathJax.ElementJax.mml;var h=MathJax.HTML;var g=f.merror.prototype.toPreviewHTML;f.merror.Augment({toPreviewHTML:function(l){if(!this.isError){return g.apply(this,arguments)}l=this.PHTMLcreateSpan(l);l.className="noError";if(this.multiLine){l.style.display="inline-block"}var n=this.data[0].data[0].data.join("").split(/\n/);for(var k=0,j=n.length;k<j;k++){h.addText(l,n[k]);if(k!==j-1){h.addElement(l,"br",{isMathJax:true})}}return l}})});b.Register.StartupHook("CommonHTML Jax Config",function(){b.Config({CommonHTML:{styles:{".mjx-chtml .mjx-noError":b.Insert({"line-height":1.2,"vertical-align":(b.Browser.isMSIE&&a.multiLine?"-2px":"")},a.style)}}})});b.Register.StartupHook("CommonHTML Jax Ready",function(){var f=MathJax.ElementJax.mml;var g=MathJax.OutputJax.CommonHTML;var i=MathJax.HTML;var h=f.merror.prototype.toCommonHTML;f.merror.Augment({toCommonHTML:function(n){if(!this.isError){return h.apply(this,arguments)}n=g.addElement(n,"mjx-noError");var p=this.data[0].data[0].data.join("").split(/\n/);for(var k=0,j=p.length;k<j;k++){i.addText(n,p[k]);if(k!==j-1){g.addElement(n,"br",{isMathJax:true})}}var o=this.CHTML=g.BBOX.zero();o.w=(n.offsetWidth)/g.em;if(j>1){var l=1.2*j/2;o.h=l+0.25;o.d=l-0.25;n.style.verticalAlign=g.Em(0.45-l)}else{o.h=1;o.d=0.2+2/g.em}return n}})});b.Startup.signal.Post("TeX noErrors Ready")})(MathJax.Hub,MathJax.HTML);MathJax.Ajax.loadComplete("[MathJax]/extensions/TeX/noErrors.js");
MathJax.Extension["TeX/noUndefined"]={version:"2.7.0",config:MathJax.Hub.CombineConfig("TeX.noUndefined",{disabled:false,attributes:{mathcolor:"red"}})};MathJax.Hub.Register.StartupHook("TeX Jax Ready",function(){var b=MathJax.Extension["TeX/noUndefined"].config;var a=MathJax.ElementJax.mml;var c=MathJax.InputJax.TeX.Parse.prototype.csUndefined;MathJax.InputJax.TeX.Parse.Augment({csUndefined:function(d){if(b.disabled){return c.apply(this,arguments)}MathJax.Hub.signal.Post(["TeX Jax - undefined control sequence",d]);this.Push(a.mtext(d).With(b.attributes))}});MathJax.Hub.Startup.signal.Post("TeX noUndefined Ready")});MathJax.Ajax.loadComplete("[MathJax]/extensions/TeX/noUndefined.js");
(function(d,c,j){var i,h="\u00A0";var k=function(m){return MathJax.Localization._.apply(MathJax.Localization,[["TeX",m]].concat([].slice.call(arguments,1)))};var f=MathJax.Object.isArray;var e=MathJax.Object.Subclass({Init:function(n,m){this.global={isInner:m};this.data=[b.start(this.global)];if(n){this.data[0].env=n}this.env=this.data[0].env},Push:function(){var o,n,p,q;for(o=0,n=arguments.length;o<n;o++){p=arguments[o];if(!p){continue}if(p instanceof i.mbase){p=b.mml(p)}p.global=this.global;q=(this.data.length?this.Top().checkItem(p):true);if(q instanceof Array){this.Pop();this.Push.apply(this,q)}else{if(q instanceof b){this.Pop();this.Push(q)}else{if(q){this.data.push(p);if(p.env){if(p.copyEnv!==false){for(var r in this.env){if(this.env.hasOwnProperty(r)){p.env[r]=this.env[r]}}}this.env=p.env}else{p.env=this.env}}}}}},Pop:function(){var m=this.data.pop();if(!m.isOpen){delete m.env}this.env=(this.data.length?this.Top().env:{});return m},Top:function(m){if(m==null){m=1}if(this.data.length<m){return null}return this.data[this.data.length-m]},Prev:function(m){var n=this.Top();if(m){return n.data[n.data.length-1]}else{return n.Pop()}},toString:function(){return"stack[\n  "+this.data.join("\n  ")+"\n]"}});var b=e.Item=MathJax.Object.Subclass({type:"base",endError:["ExtraOpenMissingClose","Extra open brace or missing close brace"],closeError:["ExtraCloseMissingOpen","Extra close brace or missing open brace"],rightError:["MissingLeftExtraRight","Missing \\left or extra \\right"],Init:function(){if(this.isOpen){this.env={}}this.data=[];this.Push.apply(this,arguments)},Push:function(){this.data.push.apply(this.data,arguments)},Pop:function(){return this.data.pop()},mmlData:function(m,n){if(m==null){m=true}if(this.data.length===1&&!n){return this.data[0]}return i.mrow.apply(i,this.data).With((m?{inferred:true}:{}))},checkItem:function(m){if(m.type==="over"&&this.isOpen){m.num=this.mmlData(false);this.data=[]}if(m.type==="cell"&&this.isOpen){if(m.linebreak){return false}d.Error(["Misplaced","Misplaced %1",m.name])}if(m.isClose&&this[m.type+"Error"]){d.Error(this[m.type+"Error"])}if(!m.isNotStack){return true}this.Push(m.data[0]);return false},With:function(m){for(var n in m){if(m.hasOwnProperty(n)){this[n]=m[n]}}return this},toString:function(){return this.type+"["+this.data.join("; ")+"]"}});b.start=b.Subclass({type:"start",isOpen:true,Init:function(m){this.SUPER(arguments).Init.call(this);this.global=m},checkItem:function(m){if(m.type==="stop"){return b.mml(this.mmlData())}return this.SUPER(arguments).checkItem.call(this,m)}});b.stop=b.Subclass({type:"stop",isClose:true});b.open=b.Subclass({type:"open",isOpen:true,stopError:["ExtraOpenMissingClose","Extra open brace or missing close brace"],checkItem:function(n){if(n.type==="close"){var m=this.mmlData();return b.mml(i.TeXAtom(m))}return this.SUPER(arguments).checkItem.call(this,n)}});b.close=b.Subclass({type:"close",isClose:true});b.prime=b.Subclass({type:"prime",checkItem:function(m){if(this.data[0].type!=="msubsup"){return[i.msup(this.data[0],this.data[1]),m]}this.data[0].SetData(this.data[0].sup,this.data[1]);return[this.data[0],m]}});b.subsup=b.Subclass({type:"subsup",stopError:["MissingScript","Missing superscript or subscript argument"],supError:["MissingOpenForSup","Missing open brace for superscript"],subError:["MissingOpenForSub","Missing open brace for subscript"],checkItem:function(m){if(m.type==="open"||m.type==="left"){return true}if(m.type==="mml"){if(this.primes){if(this.position!==2){this.data[0].SetData(2,this.primes)}else{m.data[0]=i.mrow(this.primes.With({variantForm:true}),m.data[0])}}this.data[0].SetData(this.position,m.data[0]);if(this.movesupsub!=null){this.data[0].movesupsub=this.movesupsub}return b.mml(this.data[0])}if(this.SUPER(arguments).checkItem.call(this,m)){d.Error(this[["","subError","supError"][this.position]])}},Pop:function(){}});b.over=b.Subclass({type:"over",isClose:true,name:"\\over",checkItem:function(o,m){if(o.type==="over"){d.Error(["AmbiguousUseOf","Ambiguous use of %1",o.name])}if(o.isClose){var n=i.mfrac(this.num,this.mmlData(false));if(this.thickness!=null){n.linethickness=this.thickness}if(this.open||this.close){n.texWithDelims=true;n=d.fixedFence(this.open,n,this.close)}return[b.mml(n),o]}return this.SUPER(arguments).checkItem.call(this,o)},toString:function(){return"over["+this.num+" / "+this.data.join("; ")+"]"}});b.left=b.Subclass({type:"left",isOpen:true,delim:"(",stopError:["ExtraLeftMissingRight","Extra \\left or missing \\right"],checkItem:function(m){if(m.type==="right"){return b.mml(d.fenced(this.delim,this.mmlData(),m.delim))}return this.SUPER(arguments).checkItem.call(this,m)}});b.right=b.Subclass({type:"right",isClose:true,delim:")"});b.begin=b.Subclass({type:"begin",isOpen:true,checkItem:function(m){if(m.type==="end"){if(m.name!==this.name){d.Error(["EnvBadEnd","\\begin{%1} ended with \\end{%2}",this.name,m.name])}if(!this.end){return b.mml(this.mmlData())}return this.parse[this.end].call(this.parse,this,this.data)}if(m.type==="stop"){d.Error(["EnvMissingEnd","Missing \\end{%1}",this.name])}return this.SUPER(arguments).checkItem.call(this,m)}});b.end=b.Subclass({type:"end",isClose:true});b.style=b.Subclass({type:"style",checkItem:function(n){if(!n.isClose){return this.SUPER(arguments).checkItem.call(this,n)}var m=i.mstyle.apply(i,this.data).With(this.styles);return[b.mml(m),n]}});b.position=b.Subclass({type:"position",checkItem:function(n){if(n.isClose){d.Error(["MissingBoxFor","Missing box for %1",this.name])}if(n.isNotStack){var m=n.mmlData();switch(this.move){case"vertical":m=i.mpadded(m).With({height:this.dh,depth:this.dd,voffset:this.dh});return[b.mml(m)];case"horizontal":return[b.mml(this.left),n,b.mml(this.right)]}}return this.SUPER(arguments).checkItem.call(this,n)}});b.array=b.Subclass({type:"array",isOpen:true,copyEnv:false,arraydef:{},Init:function(){this.table=[];this.row=[];this.frame=[];this.hfill=[];this.SUPER(arguments).Init.apply(this,arguments)},checkItem:function(n){if(n.isClose&&n.type!=="over"){if(n.isEntry){this.EndEntry();this.clearEnv();return false}if(n.isCR){this.EndEntry();this.EndRow();this.clearEnv();return false}this.EndTable();this.clearEnv();var o=this.arraydef.scriptlevel;delete this.arraydef.scriptlevel;var m=i.mtable.apply(i,this.table).With(this.arraydef);if(this.frame.length===4){m.frame=(this.frame.dashed?"dashed":"solid")}else{if(this.frame.length){m.hasFrame=true;if(this.arraydef.rowlines){this.arraydef.rowlines=this.arraydef.rowlines.replace(/none( none)+$/,"none")}m=i.menclose(m).With({notation:this.frame.join(" "),isFrame:true});if((this.arraydef.columnlines||"none")!="none"||(this.arraydef.rowlines||"none")!="none"){m.padding=0}}}if(o){m=i.mstyle(m).With({scriptlevel:o})}if(this.open||this.close){m=d.fenced(this.open,m,this.close)}m=b.mml(m);if(this.requireClose){if(n.type==="close"){return m}d.Error(["MissingCloseBrace","Missing close brace"])}return[m,n]}return this.SUPER(arguments).checkItem.call(this,n)},EndEntry:function(){var m=i.mtd.apply(i,this.data);if(this.hfill.length){if(this.hfill[0]===0){m.columnalign="right"}if(this.hfill[this.hfill.length-1]===this.data.length){m.columnalign=(m.columnalign?"center":"left")}}this.row.push(m);this.data=[];this.hfill=[]},EndRow:function(){var m=i.mtr;if(this.isNumbered&&this.row.length===3){this.row.unshift(this.row.pop());m=i.mlabeledtr}this.table.push(m.apply(i,this.row));this.row=[]},EndTable:function(){if(this.data.length||this.row.length){this.EndEntry();this.EndRow()}this.checkLines()},checkLines:function(){if(this.arraydef.rowlines){var m=this.arraydef.rowlines.split(/ /);if(m.length===this.table.length){this.frame.push("bottom");m.pop();this.arraydef.rowlines=m.join(" ")}else{if(m.length<this.table.length-1){this.arraydef.rowlines+=" none"}}}if(this.rowspacing){var n=this.arraydef.rowspacing.split(/ /);while(n.length<this.table.length){n.push(this.rowspacing+"em")}this.arraydef.rowspacing=n.join(" ")}},clearEnv:function(){for(var m in this.env){if(this.env.hasOwnProperty(m)){delete this.env[m]}}}});b.cell=b.Subclass({type:"cell",isClose:true});b.mml=b.Subclass({type:"mml",isNotStack:true,Add:function(){this.data.push.apply(this.data,arguments);return this}});b.fn=b.Subclass({type:"fn",checkItem:function(n){if(this.data[0]){if(n.isOpen){return true}if(n.type!=="fn"){if(n.type!=="mml"||!n.data[0]){return[this.data[0],n]}if(n.data[0].isa(i.mspace)){return[this.data[0],n]}var m=n.data[0];if(m.isEmbellished()){m=m.CoreMO()}if([0,0,1,1,0,1,1,0,0,0][m.Get("texClass")]){return[this.data[0],n]}}return[this.data[0],i.mo(i.entity("#x2061")).With({texClass:i.TEXCLASS.NONE}),n]}return this.SUPER(arguments).checkItem.apply(this,arguments)}});b.not=b.Subclass({type:"not",checkItem:function(n){var m,o;if(n.type==="open"||n.type==="left"){return true}if(n.type==="mml"&&n.data[0].type.match(/^(mo|mi|mtext)$/)){m=n.data[0],o=m.data.join("");if(o.length===1&&!m.movesupsub){o=b.not.remap[o.charCodeAt(0)];if(o){m.SetData(0,i.chars(String.fromCharCode(o)))}else{m.Append(i.chars("\u0338"))}return n}}m=i.mpadded(i.mtext("\u29F8")).With({width:0});m=i.TeXAtom(m).With({texClass:i.TEXCLASS.REL});return[m,n]}});b.not.remap={8592:8602,8594:8603,8596:8622,8656:8653,8658:8655,8660:8654,8712:8713,8715:8716,8739:8740,8741:8742,8764:8769,126:8769,8771:8772,8773:8775,8776:8777,8781:8813,61:8800,8801:8802,60:8814,62:8815,8804:8816,8805:8817,8818:8820,8819:8821,8822:8824,8823:8825,8826:8832,8827:8833,8834:8836,8835:8837,8838:8840,8839:8841,8866:8876,8872:8877,8873:8878,8875:8879,8828:8928,8829:8929,8849:8930,8850:8931,8882:8938,8883:8939,8884:8940,8885:8941,8707:8708};b.dots=b.Subclass({type:"dots",checkItem:function(n){if(n.type==="open"||n.type==="left"){return true}var o=this.ldots;if(n.type==="mml"&&n.data[0].isEmbellished()){var m=n.data[0].CoreMO().Get("texClass");if(m===i.TEXCLASS.BIN||m===i.TEXCLASS.REL){o=this.cdots}}return[o,n]}});var g={Add:function(m,p,o){if(!p){p=this}for(var n in m){if(m.hasOwnProperty(n)){if(typeof m[n]==="object"&&!f(m[n])&&(typeof p[n]==="object"||typeof p[n]==="function")){this.Add(m[n],p[n],m[n],o)}else{if(!p[n]||!p[n].isUser||!o){p[n]=m[n]}}}}return p}};var l=function(){i=MathJax.ElementJax.mml;c.Insert(g,{letter:/[a-z]/i,digit:/[0-9.]/,number:/^(?:[0-9]+(?:\{,\}[0-9]{3})*(?:\.[0-9]*)*|\.[0-9]+)/,special:{"\\":"ControlSequence","{":"Open","}":"Close","~":"Tilde","^":"Superscript",_:"Subscript"," ":"Space","\t":"Space","\r":"Space","\n":"Space","'":"Prime","%":"Comment","&":"Entry","#":"Hash","\u00A0":"Space","\u2019":"Prime"},remap:{"-":"2212","*":"2217","`":"2018"},mathchar0mi:{alpha:"03B1",beta:"03B2",gamma:"03B3",delta:"03B4",epsilon:"03F5",zeta:"03B6",eta:"03B7",theta:"03B8",iota:"03B9",kappa:"03BA",lambda:"03BB",mu:"03BC",nu:"03BD",xi:"03BE",omicron:"03BF",pi:"03C0",rho:"03C1",sigma:"03C3",tau:"03C4",upsilon:"03C5",phi:"03D5",chi:"03C7",psi:"03C8",omega:"03C9",varepsilon:"03B5",vartheta:"03D1",varpi:"03D6",varrho:"03F1",varsigma:"03C2",varphi:"03C6",S:["00A7",{mathvariant:i.VARIANT.NORMAL}],aleph:["2135",{mathvariant:i.VARIANT.NORMAL}],hbar:["210F",{variantForm:true}],imath:"0131",jmath:"0237",ell:"2113",wp:["2118",{mathvariant:i.VARIANT.NORMAL}],Re:["211C",{mathvariant:i.VARIANT.NORMAL}],Im:["2111",{mathvariant:i.VARIANT.NORMAL}],partial:["2202",{mathvariant:i.VARIANT.NORMAL}],infty:["221E",{mathvariant:i.VARIANT.NORMAL}],prime:["2032",{mathvariant:i.VARIANT.NORMAL,variantForm:true}],emptyset:["2205",{mathvariant:i.VARIANT.NORMAL}],nabla:["2207",{mathvariant:i.VARIANT.NORMAL}],top:["22A4",{mathvariant:i.VARIANT.NORMAL}],bot:["22A5",{mathvariant:i.VARIANT.NORMAL}],angle:["2220",{mathvariant:i.VARIANT.NORMAL}],triangle:["25B3",{mathvariant:i.VARIANT.NORMAL}],backslash:["2216",{mathvariant:i.VARIANT.NORMAL,variantForm:true}],forall:["2200",{mathvariant:i.VARIANT.NORMAL}],exists:["2203",{mathvariant:i.VARIANT.NORMAL}],neg:["00AC",{mathvariant:i.VARIANT.NORMAL}],lnot:["00AC",{mathvariant:i.VARIANT.NORMAL}],flat:["266D",{mathvariant:i.VARIANT.NORMAL}],natural:["266E",{mathvariant:i.VARIANT.NORMAL}],sharp:["266F",{mathvariant:i.VARIANT.NORMAL}],clubsuit:["2663",{mathvariant:i.VARIANT.NORMAL}],diamondsuit:["2662",{mathvariant:i.VARIANT.NORMAL}],heartsuit:["2661",{mathvariant:i.VARIANT.NORMAL}],spadesuit:["2660",{mathvariant:i.VARIANT.NORMAL}]},mathchar0mo:{surd:"221A",coprod:["2210",{texClass:i.TEXCLASS.OP,movesupsub:true}],bigvee:["22C1",{texClass:i.TEXCLASS.OP,movesupsub:true}],bigwedge:["22C0",{texClass:i.TEXCLASS.OP,movesupsub:true}],biguplus:["2A04",{texClass:i.TEXCLASS.OP,movesupsub:true}],bigcap:["22C2",{texClass:i.TEXCLASS.OP,movesupsub:true}],bigcup:["22C3",{texClass:i.TEXCLASS.OP,movesupsub:true}],"int":["222B",{texClass:i.TEXCLASS.OP}],intop:["222B",{texClass:i.TEXCLASS.OP,movesupsub:true,movablelimits:true}],iint:["222C",{texClass:i.TEXCLASS.OP}],iiint:["222D",{texClass:i.TEXCLASS.OP}],prod:["220F",{texClass:i.TEXCLASS.OP,movesupsub:true}],sum:["2211",{texClass:i.TEXCLASS.OP,movesupsub:true}],bigotimes:["2A02",{texClass:i.TEXCLASS.OP,movesupsub:true}],bigoplus:["2A01",{texClass:i.TEXCLASS.OP,movesupsub:true}],bigodot:["2A00",{texClass:i.TEXCLASS.OP,movesupsub:true}],oint:["222E",{texClass:i.TEXCLASS.OP}],bigsqcup:["2A06",{texClass:i.TEXCLASS.OP,movesupsub:true}],smallint:["222B",{largeop:false}],triangleleft:"25C3",triangleright:"25B9",bigtriangleup:"25B3",bigtriangledown:"25BD",wedge:"2227",land:"2227",vee:"2228",lor:"2228",cap:"2229",cup:"222A",ddagger:"2021",dagger:"2020",sqcap:"2293",sqcup:"2294",uplus:"228E",amalg:"2A3F",diamond:"22C4",bullet:"2219",wr:"2240",div:"00F7",odot:["2299",{largeop:false}],oslash:["2298",{largeop:false}],otimes:["2297",{largeop:false}],ominus:["2296",{largeop:false}],oplus:["2295",{largeop:false}],mp:"2213",pm:"00B1",circ:"2218",bigcirc:"25EF",setminus:["2216",{variantForm:true}],cdot:"22C5",ast:"2217",times:"00D7",star:"22C6",propto:"221D",sqsubseteq:"2291",sqsupseteq:"2292",parallel:"2225",mid:"2223",dashv:"22A3",vdash:"22A2",leq:"2264",le:"2264",geq:"2265",ge:"2265",lt:"003C",gt:"003E",succ:"227B",prec:"227A",approx:"2248",succeq:"2AB0",preceq:"2AAF",supset:"2283",subset:"2282",supseteq:"2287",subseteq:"2286","in":"2208",ni:"220B",notin:"2209",owns:"220B",gg:"226B",ll:"226A",sim:"223C",simeq:"2243",perp:"22A5",equiv:"2261",asymp:"224D",smile:"2323",frown:"2322",ne:"2260",neq:"2260",cong:"2245",doteq:"2250",bowtie:"22C8",models:"22A8",notChar:"29F8",Leftrightarrow:"21D4",Leftarrow:"21D0",Rightarrow:"21D2",leftrightarrow:"2194",leftarrow:"2190",gets:"2190",rightarrow:"2192",to:"2192",mapsto:"21A6",leftharpoonup:"21BC",leftharpoondown:"21BD",rightharpoonup:"21C0",rightharpoondown:"21C1",nearrow:"2197",searrow:"2198",nwarrow:"2196",swarrow:"2199",rightleftharpoons:"21CC",hookrightarrow:"21AA",hookleftarrow:"21A9",longleftarrow:"27F5",Longleftarrow:"27F8",longrightarrow:"27F6",Longrightarrow:"27F9",Longleftrightarrow:"27FA",longleftrightarrow:"27F7",longmapsto:"27FC",ldots:"2026",cdots:"22EF",vdots:"22EE",ddots:"22F1",dotsc:"2026",dotsb:"22EF",dotsm:"22EF",dotsi:"22EF",dotso:"2026",ldotp:["002E",{texClass:i.TEXCLASS.PUNCT}],cdotp:["22C5",{texClass:i.TEXCLASS.PUNCT}],colon:["003A",{texClass:i.TEXCLASS.PUNCT}]},mathchar7:{Gamma:"0393",Delta:"0394",Theta:"0398",Lambda:"039B",Xi:"039E",Pi:"03A0",Sigma:"03A3",Upsilon:"03A5",Phi:"03A6",Psi:"03A8",Omega:"03A9",_:"005F","#":"0023","$":"0024","%":"0025","&":"0026",And:"0026"},delimiter:{"(":"(",")":")","[":"[","]":"]","<":"27E8",">":"27E9","\\lt":"27E8","\\gt":"27E9","/":"/","|":["|",{texClass:i.TEXCLASS.ORD}],".":"","\\\\":"\\","\\lmoustache":"23B0","\\rmoustache":"23B1","\\lgroup":"27EE","\\rgroup":"27EF","\\arrowvert":"23D0","\\Arrowvert":"2016","\\bracevert":"23AA","\\Vert":["2016",{texClass:i.TEXCLASS.ORD}],"\\|":["2016",{texClass:i.TEXCLASS.ORD}],"\\vert":["|",{texClass:i.TEXCLASS.ORD}],"\\uparrow":"2191","\\downarrow":"2193","\\updownarrow":"2195","\\Uparrow":"21D1","\\Downarrow":"21D3","\\Updownarrow":"21D5","\\backslash":"\\","\\rangle":"27E9","\\langle":"27E8","\\rbrace":"}","\\lbrace":"{","\\}":"}","\\{":"{","\\rceil":"2309","\\lceil":"2308","\\rfloor":"230B","\\lfloor":"230A","\\lbrack":"[","\\rbrack":"]"},macros:{displaystyle:["SetStyle","D",true,0],textstyle:["SetStyle","T",false,0],scriptstyle:["SetStyle","S",false,1],scriptscriptstyle:["SetStyle","SS",false,2],rm:["SetFont",i.VARIANT.NORMAL],mit:["SetFont",i.VARIANT.ITALIC],oldstyle:["SetFont",i.VARIANT.OLDSTYLE],cal:["SetFont",i.VARIANT.CALIGRAPHIC],it:["SetFont","-tex-mathit"],bf:["SetFont",i.VARIANT.BOLD],bbFont:["SetFont",i.VARIANT.DOUBLESTRUCK],scr:["SetFont",i.VARIANT.SCRIPT],frak:["SetFont",i.VARIANT.FRAKTUR],sf:["SetFont",i.VARIANT.SANSSERIF],tt:["SetFont",i.VARIANT.MONOSPACE],tiny:["SetSize",0.5],Tiny:["SetSize",0.6],scriptsize:["SetSize",0.7],small:["SetSize",0.85],normalsize:["SetSize",1],large:["SetSize",1.2],Large:["SetSize",1.44],LARGE:["SetSize",1.73],huge:["SetSize",2.07],Huge:["SetSize",2.49],arcsin:["NamedFn"],arccos:["NamedFn"],arctan:["NamedFn"],arg:["NamedFn"],cos:["NamedFn"],cosh:["NamedFn"],cot:["NamedFn"],coth:["NamedFn"],csc:["NamedFn"],deg:["NamedFn"],det:"NamedOp",dim:["NamedFn"],exp:["NamedFn"],gcd:"NamedOp",hom:["NamedFn"],inf:"NamedOp",ker:["NamedFn"],lg:["NamedFn"],lim:"NamedOp",liminf:["NamedOp","lim&thinsp;inf"],limsup:["NamedOp","lim&thinsp;sup"],ln:["NamedFn"],log:["NamedFn"],max:"NamedOp",min:"NamedOp",Pr:"NamedOp",sec:["NamedFn"],sin:["NamedFn"],sinh:["NamedFn"],sup:"NamedOp",tan:["NamedFn"],tanh:["NamedFn"],limits:["Limits",1],nolimits:["Limits",0],overline:["UnderOver","00AF",null,1],underline:["UnderOver","005F"],overbrace:["UnderOver","23DE",1],underbrace:["UnderOver","23DF",1],overparen:["UnderOver","23DC"],underparen:["UnderOver","23DD"],overrightarrow:["UnderOver","2192"],underrightarrow:["UnderOver","2192"],overleftarrow:["UnderOver","2190"],underleftarrow:["UnderOver","2190"],overleftrightarrow:["UnderOver","2194"],underleftrightarrow:["UnderOver","2194"],overset:"Overset",underset:"Underset",stackrel:["Macro","\\mathrel{\\mathop{#2}\\limits^{#1}}",2],over:"Over",overwithdelims:"Over",atop:"Over",atopwithdelims:"Over",above:"Over",abovewithdelims:"Over",brace:["Over","{","}"],brack:["Over","[","]"],choose:["Over","(",")"],frac:"Frac",sqrt:"Sqrt",root:"Root",uproot:["MoveRoot","upRoot"],leftroot:["MoveRoot","leftRoot"],left:"LeftRight",right:"LeftRight",middle:"Middle",llap:"Lap",rlap:"Lap",raise:"RaiseLower",lower:"RaiseLower",moveleft:"MoveLeftRight",moveright:"MoveLeftRight",",":["Spacer",i.LENGTH.THINMATHSPACE],":":["Spacer",i.LENGTH.MEDIUMMATHSPACE],">":["Spacer",i.LENGTH.MEDIUMMATHSPACE],";":["Spacer",i.LENGTH.THICKMATHSPACE],"!":["Spacer",i.LENGTH.NEGATIVETHINMATHSPACE],enspace:["Spacer",".5em"],quad:["Spacer","1em"],qquad:["Spacer","2em"],thinspace:["Spacer",i.LENGTH.THINMATHSPACE],negthinspace:["Spacer",i.LENGTH.NEGATIVETHINMATHSPACE],hskip:"Hskip",hspace:"Hskip",kern:"Hskip",mskip:"Hskip",mspace:"Hskip",mkern:"Hskip",Rule:["Rule"],Space:["Rule","blank"],big:["MakeBig",i.TEXCLASS.ORD,0.85],Big:["MakeBig",i.TEXCLASS.ORD,1.15],bigg:["MakeBig",i.TEXCLASS.ORD,1.45],Bigg:["MakeBig",i.TEXCLASS.ORD,1.75],bigl:["MakeBig",i.TEXCLASS.OPEN,0.85],Bigl:["MakeBig",i.TEXCLASS.OPEN,1.15],biggl:["MakeBig",i.TEXCLASS.OPEN,1.45],Biggl:["MakeBig",i.TEXCLASS.OPEN,1.75],bigr:["MakeBig",i.TEXCLASS.CLOSE,0.85],Bigr:["MakeBig",i.TEXCLASS.CLOSE,1.15],biggr:["MakeBig",i.TEXCLASS.CLOSE,1.45],Biggr:["MakeBig",i.TEXCLASS.CLOSE,1.75],bigm:["MakeBig",i.TEXCLASS.REL,0.85],Bigm:["MakeBig",i.TEXCLASS.REL,1.15],biggm:["MakeBig",i.TEXCLASS.REL,1.45],Biggm:["MakeBig",i.TEXCLASS.REL,1.75],mathord:["TeXAtom",i.TEXCLASS.ORD],mathop:["TeXAtom",i.TEXCLASS.OP],mathopen:["TeXAtom",i.TEXCLASS.OPEN],mathclose:["TeXAtom",i.TEXCLASS.CLOSE],mathbin:["TeXAtom",i.TEXCLASS.BIN],mathrel:["TeXAtom",i.TEXCLASS.REL],mathpunct:["TeXAtom",i.TEXCLASS.PUNCT],mathinner:["TeXAtom",i.TEXCLASS.INNER],vcenter:["TeXAtom",i.TEXCLASS.VCENTER],mathchoice:["Extension","mathchoice"],buildrel:"BuildRel",hbox:["HBox",0],text:"HBox",mbox:["HBox",0],fbox:"FBox",strut:"Strut",mathstrut:["Macro","\\vphantom{(}"],phantom:"Phantom",vphantom:["Phantom",1,0],hphantom:["Phantom",0,1],smash:"Smash",acute:["Accent","00B4"],grave:["Accent","0060"],ddot:["Accent","00A8"],tilde:["Accent","007E"],bar:["Accent","00AF"],breve:["Accent","02D8"],check:["Accent","02C7"],hat:["Accent","005E"],vec:["Accent","2192"],dot:["Accent","02D9"],widetilde:["Accent","007E",1],widehat:["Accent","005E",1],matrix:"Matrix",array:"Matrix",pmatrix:["Matrix","(",")"],cases:["Matrix","{","","left left",null,".1em",null,true],eqalign:["Matrix",null,null,"right left",i.LENGTH.THICKMATHSPACE,".5em","D"],displaylines:["Matrix",null,null,"center",null,".5em","D"],cr:"Cr","\\":"CrLaTeX",newline:"Cr",hline:["HLine","solid"],hdashline:["HLine","dashed"],eqalignno:["Matrix",null,null,"right left",i.LENGTH.THICKMATHSPACE,".5em","D",null,"right"],leqalignno:["Matrix",null,null,"right left",i.LENGTH.THICKMATHSPACE,".5em","D",null,"left"],hfill:"HFill",hfil:"HFill",hfilll:"HFill",bmod:["Macro",'\\mmlToken{mo}[lspace="thickmathspace" rspace="thickmathspace"]{mod}'],pmod:["Macro","\\pod{\\mmlToken{mi}{mod}\\kern 6mu #1}",1],mod:["Macro","\\mathchoice{\\kern18mu}{\\kern12mu}{\\kern12mu}{\\kern12mu}\\mmlToken{mi}{mod}\\,\\,#1",1],pod:["Macro","\\mathchoice{\\kern18mu}{\\kern8mu}{\\kern8mu}{\\kern8mu}(#1)",1],iff:["Macro","\\;\\Longleftrightarrow\\;"],skew:["Macro","{{#2{#3\\mkern#1mu}\\mkern-#1mu}{}}",3],mathcal:["Macro","{\\cal #1}",1],mathscr:["Macro","{\\scr #1}",1],mathrm:["Macro","{\\rm #1}",1],mathbf:["Macro","{\\bf #1}",1],mathbb:["Macro","{\\bbFont #1}",1],Bbb:["Macro","{\\bbFont #1}",1],mathit:["Macro","{\\it #1}",1],mathfrak:["Macro","{\\frak #1}",1],mathsf:["Macro","{\\sf #1}",1],mathtt:["Macro","{\\tt #1}",1],textrm:["Macro","\\mathord{\\rm\\text{#1}}",1],textit:["Macro","\\mathord{\\it\\text{#1}}",1],textbf:["Macro","\\mathord{\\bf\\text{#1}}",1],textsf:["Macro","\\mathord{\\sf\\text{#1}}",1],texttt:["Macro","\\mathord{\\tt\\text{#1}}",1],pmb:["Macro","\\rlap{#1}\\kern1px{#1}",1],TeX:["Macro","T\\kern-.14em\\lower.5ex{E}\\kern-.115em X"],LaTeX:["Macro","L\\kern-.325em\\raise.21em{\\scriptstyle{A}}\\kern-.17em\\TeX"]," ":["Macro","\\text{ }"],not:"Not",dots:"Dots",space:"Tilde","\u00A0":"Tilde",begin:"BeginEnd",end:"BeginEnd",newcommand:["Extension","newcommand"],renewcommand:["Extension","newcommand"],newenvironment:["Extension","newcommand"],renewenvironment:["Extension","newcommand"],def:["Extension","newcommand"],let:["Extension","newcommand"],verb:["Extension","verb"],boldsymbol:["Extension","boldsymbol"],tag:["Extension","AMSmath"],notag:["Extension","AMSmath"],label:["Extension","AMSmath"],ref:["Extension","AMSmath"],eqref:["Extension","AMSmath"],nonumber:["Macro","\\notag"],unicode:["Extension","unicode"],color:"Color",href:["Extension","HTML"],"class":["Extension","HTML"],style:["Extension","HTML"],cssId:["Extension","HTML"],bbox:["Extension","bbox"],mmlToken:"MmlToken",require:"Require"},environment:{array:["AlignedArray"],matrix:["Array",null,null,null,"c"],pmatrix:["Array",null,"(",")","c"],bmatrix:["Array",null,"[","]","c"],Bmatrix:["Array",null,"\\{","\\}","c"],vmatrix:["Array",null,"\\vert","\\vert","c"],Vmatrix:["Array",null,"\\Vert","\\Vert","c"],cases:["Array",null,"\\{",".","ll",null,".2em","T"],equation:[null,"Equation"],"equation*":[null,"Equation"],eqnarray:["ExtensionEnv",null,"AMSmath"],"eqnarray*":["ExtensionEnv",null,"AMSmath"],align:["ExtensionEnv",null,"AMSmath"],"align*":["ExtensionEnv",null,"AMSmath"],aligned:["ExtensionEnv",null,"AMSmath"],multline:["ExtensionEnv",null,"AMSmath"],"multline*":["ExtensionEnv",null,"AMSmath"],split:["ExtensionEnv",null,"AMSmath"],gather:["ExtensionEnv",null,"AMSmath"],"gather*":["ExtensionEnv",null,"AMSmath"],gathered:["ExtensionEnv",null,"AMSmath"],alignat:["ExtensionEnv",null,"AMSmath"],"alignat*":["ExtensionEnv",null,"AMSmath"],alignedat:["ExtensionEnv",null,"AMSmath"]},p_height:1.2/0.85});if(this.config.Macros){var m=this.config.Macros;for(var n in m){if(m.hasOwnProperty(n)){if(typeof(m[n])==="string"){g.macros[n]=["Macro",m[n]]}else{g.macros[n]=["Macro"].concat(m[n])}g.macros[n].isUser=true}}}};var a=MathJax.Object.Subclass({Init:function(n,o){this.string=n;this.i=0;this.macroCount=0;var m;if(o){m={};for(var p in o){if(o.hasOwnProperty(p)){m[p]=o[p]}}}this.stack=d.Stack(m,!!o);this.Parse();this.Push(b.stop())},Parse:function(){var o,m;while(this.i<this.string.length){o=this.string.charAt(this.i++);m=o.charCodeAt(0);if(m>=55296&&m<56320){o+=this.string.charAt(this.i++)}if(g.special[o]){this[g.special[o]](o)}else{if(g.letter.test(o)){this.Variable(o)}else{if(g.digit.test(o)){this.Number(o)}else{this.Other(o)}}}}},Push:function(){this.stack.Push.apply(this.stack,arguments)},mml:function(){if(this.stack.Top().type!=="mml"){return null}return this.stack.Top().data[0]},mmlToken:function(m){return m},ControlSequence:function(p){var m=this.GetCS(),o=this.csFindMacro(m);if(o){if(!f(o)){o=[o]}var n=o[0];if(!(n instanceof Function)){n=this[n]}n.apply(this,[p+m].concat(o.slice(1)))}else{if(g.mathchar0mi[m]){this.csMathchar0mi(m,g.mathchar0mi[m])}else{if(g.mathchar0mo[m]){this.csMathchar0mo(m,g.mathchar0mo[m])}else{if(g.mathchar7[m]){this.csMathchar7(m,g.mathchar7[m])}else{if(g.delimiter["\\"+m]!=null){this.csDelimiter(m,g.delimiter["\\"+m])}else{this.csUndefined(p+m)}}}}}},csFindMacro:function(m){return g.macros[m]},csMathchar0mi:function(m,o){var n={mathvariant:i.VARIANT.ITALIC};if(f(o)){n=o[1];o=o[0]}this.Push(this.mmlToken(i.mi(i.entity("#x"+o)).With(n)))},csMathchar0mo:function(m,o){var n={stretchy:false};if(f(o)){n=o[1];n.stretchy=false;o=o[0]}this.Push(this.mmlToken(i.mo(i.entity("#x"+o)).With(n)))},csMathchar7:function(m,o){var n={mathvariant:i.VARIANT.NORMAL};if(f(o)){n=o[1];o=o[0]}if(this.stack.env.font){n.mathvariant=this.stack.env.font}this.Push(this.mmlToken(i.mi(i.entity("#x"+o)).With(n)))},csDelimiter:function(m,o){var n={};if(f(o)){n=o[1];o=o[0]}if(o.length===4){o=i.entity("#x"+o)}else{o=i.chars(o)}this.Push(this.mmlToken(i.mo(o).With({fence:false,stretchy:false}).With(n)))},csUndefined:function(m){d.Error(["UndefinedControlSequence","Undefined control sequence %1",m])},Variable:function(n){var m={};if(this.stack.env.font){m.mathvariant=this.stack.env.font}this.Push(this.mmlToken(i.mi(i.chars(n)).With(m)))},Number:function(p){var m,o=this.string.slice(this.i-1).match(g.number);if(o){m=i.mn(o[0].replace(/[{}]/g,""));this.i+=o[0].length-1}else{m=i.mo(i.chars(p))}if(this.stack.env.font){m.mathvariant=this.stack.env.font}this.Push(this.mmlToken(m))},Open:function(m){this.Push(b.open())},Close:function(m){this.Push(b.close())},Tilde:function(m){this.Push(i.mtext(i.chars(h)))},Space:function(m){},Superscript:function(r){if(this.GetNext().match(/\d/)){this.string=this.string.substr(0,this.i+1)+" "+this.string.substr(this.i+1)}var q,o,p=this.stack.Top();if(p.type==="prime"){o=p.data[0];q=p.data[1];this.stack.Pop()}else{o=this.stack.Prev();if(!o){o=i.mi("")}}if(o.isEmbellishedWrapper){o=o.data[0].data[0]}var n=o.movesupsub,m=o.sup;if((o.type==="msubsup"&&o.data[o.sup])||(o.type==="munderover"&&o.data[o.over]&&!o.subsupOK)){d.Error(["DoubleExponent","Double exponent: use braces to clarify"])}if(o.type!=="msubsup"){if(n){if(o.type!=="munderover"||o.data[o.over]){if(o.movablelimits&&o.isa(i.mi)){o=this.mi2mo(o)}o=i.munderover(o,null,null).With({movesupsub:true})}m=o.over}else{o=i.msubsup(o,null,null);m=o.sup}}this.Push(b.subsup(o).With({position:m,primes:q,movesupsub:n}))},Subscript:function(r){if(this.GetNext().match(/\d/)){this.string=this.string.substr(0,this.i+1)+" "+this.string.substr(this.i+1)}var q,o,p=this.stack.Top();if(p.type==="prime"){o=p.data[0];q=p.data[1];this.stack.Pop()}else{o=this.stack.Prev();if(!o){o=i.mi("")}}if(o.isEmbellishedWrapper){o=o.data[0].data[0]}var n=o.movesupsub,m=o.sub;if((o.type==="msubsup"&&o.data[o.sub])||(o.type==="munderover"&&o.data[o.under]&&!o.subsupOK)){d.Error(["DoubleSubscripts","Double subscripts: use braces to clarify"])}if(o.type!=="msubsup"){if(n){if(o.type!=="munderover"||o.data[o.under]){if(o.movablelimits&&o.isa(i.mi)){o=this.mi2mo(o)}o=i.munderover(o,null,null).With({movesupsub:true})}m=o.under}else{o=i.msubsup(o,null,null);m=o.sub}}this.Push(b.subsup(o).With({position:m,primes:q,movesupsub:n}))},PRIME:"\u2032",SMARTQUOTE:"\u2019",Prime:function(o){var n=this.stack.Prev();if(!n){n=i.mi()}if(n.type==="msubsup"&&n.data[n.sup]){d.Error(["DoubleExponentPrime","Prime causes double exponent: use braces to clarify"])}var m="";this.i--;do{m+=this.PRIME;this.i++,o=this.GetNext()}while(o==="'"||o===this.SMARTQUOTE);m=["","\u2032","\u2033","\u2034","\u2057"][m.length]||m;this.Push(b.prime(n,this.mmlToken(i.mo(m))))},mi2mo:function(m){var n=i.mo();n.Append.apply(n,m.data);var o;for(o in n.defaults){if(n.defaults.hasOwnProperty(o)&&m[o]!=null){n[o]=m[o]}}for(o in i.copyAttributes){if(i.copyAttributes.hasOwnProperty(o)&&m[o]!=null){n[o]=m[o]}}n.lspace=n.rspace="0";n.useMMLspacing&=~(n.SPACE_ATTR.lspace|n.SPACE_ATTR.rspace);return n},Comment:function(m){while(this.i<this.string.length&&this.string.charAt(this.i)!="\n"){this.i++}},Hash:function(m){d.Error(["CantUseHash1","You can't use 'macro parameter character #' in math mode"])},Other:function(o){var n,m;if(this.stack.env.font){n={mathvariant:this.stack.env.font}}if(g.remap[o]){o=g.remap[o];if(f(o)){n=o[1];o=o[0]}m=i.mo(i.entity("#x"+o)).With(n)}else{m=i.mo(o).With(n)}if(m.autoDefault("stretchy",true)){m.stretchy=false}if(m.autoDefault("texClass",true)==""){m=i.TeXAtom(m)}this.Push(this.mmlToken(m))},SetFont:function(n,m){this.stack.env.font=m},SetStyle:function(n,m,o,p){this.stack.env.style=m;this.stack.env.level=p;this.Push(b.style().With({styles:{displaystyle:o,scriptlevel:p}}))},SetSize:function(m,n){this.stack.env.size=n;this.Push(b.style().With({styles:{mathsize:n+"em"}}))},Color:function(o){var n=this.GetArgument(o);var m=this.stack.env.color;this.stack.env.color=n;var p=this.ParseArg(o);if(m){this.stack.env.color}else{delete this.stack.env.color}this.Push(i.mstyle(p).With({mathcolor:n}))},Spacer:function(m,n){this.Push(i.mspace().With({width:n,mathsize:i.SIZE.NORMAL,scriptlevel:0}))},LeftRight:function(m){this.Push(b[m.substr(1)]().With({delim:this.GetDelimiter(m)}))},Middle:function(m){var n=this.GetDelimiter(m);this.Push(i.TeXAtom().With({texClass:i.TEXCLASS.CLOSE}));if(this.stack.Top().type!=="left"){d.Error(["MisplacedMiddle","%1 must be within \\left and \\right",m])}this.Push(i.mo(n).With({stretchy:true}));this.Push(i.TeXAtom().With({texClass:i.TEXCLASS.OPEN}))},NamedFn:function(n,o){if(!o){o=n.substr(1)}var m=i.mi(o).With({texClass:i.TEXCLASS.OP});this.Push(b.fn(this.mmlToken(m)))},NamedOp:function(n,o){if(!o){o=n.substr(1)}o=o.replace(/&thinsp;/,"\u2006");var m=i.mo(o).With({movablelimits:true,movesupsub:true,form:i.FORM.PREFIX,texClass:i.TEXCLASS.OP});m.useMMLspacing&=~m.SPACE_ATTR.form;this.Push(this.mmlToken(m))},Limits:function(n,m){var p=this.stack.Prev("nopop");if(!p||(p.Get("texClass")!==i.TEXCLASS.OP&&p.movesupsub==null)){d.Error(["MisplacedLimits","%1 is allowed only on operators",n])}var o=this.stack.Top();if(p.type==="munderover"&&!m){p=o.data[o.data.length-1]=i.msubsup.apply(i.subsup,p.data)}else{if(p.type==="msubsup"&&m){p=o.data[o.data.length-1]=i.munderover.apply(i.underover,p.data)}}p.movesupsub=(m?true:false);p.Core().movablelimits=false;if(p.movablelimits){p.movablelimits=false}},Over:function(o,n,p){var m=b.over().With({name:o});if(n||p){m.open=n;m.close=p}else{if(o.match(/withdelims$/)){m.open=this.GetDelimiter(o);m.close=this.GetDelimiter(o)}}if(o.match(/^\\above/)){m.thickness=this.GetDimen(o)}else{if(o.match(/^\\atop/)||n||p){m.thickness=0}}this.Push(m)},Frac:function(n){var m=this.ParseArg(n);var o=this.ParseArg(n);this.Push(i.mfrac(m,o))},Sqrt:function(p){var q=this.GetBrackets(p),m=this.GetArgument(p);if(m==="\\frac"){m+="{"+this.GetArgument(m)+"}{"+this.GetArgument(m)+"}"}var o=d.Parse(m,this.stack.env).mml();if(!q){o=i.msqrt.apply(i,o.array())}else{o=i.mroot(o,this.parseRoot(q))}this.Push(o)},Root:function(o){var p=this.GetUpTo(o,"\\of");var m=this.ParseArg(o);this.Push(i.mroot(m,this.parseRoot(p)))},parseRoot:function(r){var o=this.stack.env,m=o.inRoot;o.inRoot=true;var q=d.Parse(r,o);r=q.mml();var p=q.stack.global;if(p.leftRoot||p.upRoot){r=i.mpadded(r);if(p.leftRoot){r.width=p.leftRoot}if(p.upRoot){r.voffset=p.upRoot;r.height=p.upRoot}}o.inRoot=m;return r},MoveRoot:function(m,p){if(!this.stack.env.inRoot){d.Error(["MisplacedMoveRoot","%1 can appear only within a root",m])}if(this.stack.global[p]){d.Error(["MultipleMoveRoot","Multiple use of %1",m])}var o=this.GetArgument(m);if(!o.match(/-?[0-9]+/)){d.Error(["IntegerArg","The argument to %1 must be an integer",m])}o=(o/15)+"em";if(o.substr(0,1)!=="-"){o="+"+o}this.stack.global[p]=o},Accent:function(o,m,s){var r=this.ParseArg(o);var q={accent:true};if(this.stack.env.font){q.mathvariant=this.stack.env.font}var n=this.mmlToken(i.mo(i.entity("#x"+m)).With(q));n.stretchy=(s?true:false);var p=(r.isEmbellished()?r.CoreMO():r);if(p.isa(i.mo)){p.movablelimits=false}this.Push(i.TeXAtom(i.munderover(r,null,n).With({accent:true})))},UnderOver:function(o,s,m,q){var r={o:"over",u:"under"}[o.charAt(1)];var p=this.ParseArg(o);if(p.Get("movablelimits")){p.movablelimits=false}if(p.isa(i.munderover)&&p.isEmbellished()){p.Core().With({lspace:0,rspace:0});p=i.mrow(i.mo().With({rspace:0}),p)}var n=i.munderover(p,null,null);n.SetData(n[r],this.mmlToken(i.mo(i.entity("#x"+s)).With({stretchy:true,accent:!q})));if(m){n=i.TeXAtom(n).With({texClass:i.TEXCLASS.OP,movesupsub:true})}this.Push(n.With({subsupOK:true}))},Overset:function(m){var o=this.ParseArg(m),n=this.ParseArg(m);if(n.movablelimits){n.movablelimits=false}this.Push(i.mover(n,o))},Underset:function(m){var o=this.ParseArg(m),n=this.ParseArg(m);if(n.movablelimits){n.movablelimits=false}this.Push(i.munder(n,o))},TeXAtom:function(p,r){var q={texClass:r},o;if(r==i.TEXCLASS.OP){q.movesupsub=q.movablelimits=true;var m=this.GetArgument(p);var n=m.match(/^\s*\\rm\s+([a-zA-Z0-9 ]+)$/);if(n){q.mathvariant=i.VARIANT.NORMAL;o=b.fn(this.mmlToken(i.mi(n[1]).With(q)))}else{o=b.fn(i.TeXAtom(d.Parse(m,this.stack.env).mml()).With(q))}}else{o=i.TeXAtom(this.ParseArg(p)).With(q)}this.Push(o)},MmlToken:function(o){var p=this.GetArgument(o),m=this.GetBrackets(o,"").replace(/^\s+/,""),s=this.GetArgument(o),r={attrNames:[]},n;if(!i[p]||!i[p].prototype.isToken){d.Error(["NotMathMLToken","%1 is not a token element",p])}while(m!==""){n=m.match(/^([a-z]+)\s*=\s*('[^']*'|"[^"]*"|[^ ,]*)\s*,?\s*/i);if(!n){d.Error(["InvalidMathMLAttr","Invalid MathML attribute: %1",m])}if(i[p].prototype.defaults[n[1]]==null&&!this.MmlTokenAllow[n[1]]){d.Error(["UnknownAttrForElement","%1 is not a recognized attribute for %2",n[1],p])}var q=this.MmlFilterAttribute(n[1],n[2].replace(/^(['"])(.*)\1$/,"$2"));if(q){if(q.toLowerCase()==="true"){q=true}else{if(q.toLowerCase()==="false"){q=false}}r[n[1]]=q;r.attrNames.push(n[1])}m=m.substr(n[0].length)}this.Push(this.mmlToken(i[p](s).With(r)))},MmlFilterAttribute:function(m,n){return n},MmlTokenAllow:{fontfamily:1,fontsize:1,fontweight:1,fontstyle:1,color:1,background:1,id:1,"class":1,href:1,style:1},Strut:function(m){this.Push(i.mpadded(i.mrow()).With({height:"8.6pt",depth:"3pt",width:0}))},Phantom:function(n,m,o){var p=i.mphantom(this.ParseArg(n));if(m||o){p=i.mpadded(p);if(o){p.height=p.depth=0}if(m){p.width=0}}this.Push(i.TeXAtom(p))},Smash:function(o){var n=this.trimSpaces(this.GetBrackets(o,""));var m=i.mpadded(this.ParseArg(o));switch(n){case"b":m.depth=0;break;case"t":m.height=0;break;default:m.height=m.depth=0}this.Push(i.TeXAtom(m))},Lap:function(n){var m=i.mpadded(this.ParseArg(n)).With({width:0});if(n==="\\llap"){m.lspace="-1width"}this.Push(i.TeXAtom(m))},RaiseLower:function(m){var n=this.GetDimen(m);var o=b.position().With({name:m,move:"vertical"});if(n.charAt(0)==="-"){n=n.slice(1);m={raise:"\\lower",lower:"\\raise"}[m.substr(1)]}if(m==="\\lower"){o.dh="-"+n;o.dd="+"+n}else{o.dh="+"+n;o.dd="-"+n}this.Push(o)},MoveLeftRight:function(m){var p=this.GetDimen(m);var o=(p.charAt(0)==="-"?p.slice(1):"-"+p);if(m==="\\moveleft"){var n=p;p=o;o=n}this.Push(b.position().With({name:m,move:"horizontal",left:i.mspace().With({width:p,mathsize:i.SIZE.NORMAL}),right:i.mspace().With({width:o,mathsize:i.SIZE.NORMAL})}))},Hskip:function(m){this.Push(i.mspace().With({width:this.GetDimen(m),mathsize:i.SIZE.NORMAL}))},Rule:function(o,q){var m=this.GetDimen(o),p=this.GetDimen(o),s=this.GetDimen(o);var n,r={width:m,height:p,depth:s};if(q!=="blank"){if(parseFloat(m)&&parseFloat(p)+parseFloat(s)){r.mathbackground=(this.stack.env.color||"black")}n=i.mpadded(i.mrow()).With(r)}else{n=i.mspace().With(r)}this.Push(n)},MakeBig:function(m,p,n){n*=g.p_height;n=String(n).replace(/(\.\d\d\d).+/,"$1")+"em";var o=this.GetDelimiter(m,true);this.Push(i.TeXAtom(i.mo(o).With({minsize:n,maxsize:n,fence:true,stretchy:true,symmetric:true})).With({texClass:p}))},BuildRel:function(m){var n=this.ParseUpTo(m,"\\over");var o=this.ParseArg(m);this.Push(i.TeXAtom(i.munderover(o,null,n)).With({texClass:i.TEXCLASS.REL}))},HBox:function(m,n){this.Push.apply(this,this.InternalMath(this.GetArgument(m),n))},FBox:function(m){this.Push(i.menclose.apply(i,this.InternalMath(this.GetArgument(m))).With({notation:"box"}))},Not:function(m){this.Push(b.not())},Dots:function(m){this.Push(b.dots().With({ldots:this.mmlToken(i.mo(i.entity("#x2026")).With({stretchy:false})),cdots:this.mmlToken(i.mo(i.entity("#x22EF")).With({stretchy:false}))}))},Require:function(m){var n=this.GetArgument(m).replace(/.*\//,"").replace(/[^a-z0-9_.-]/ig,"");this.Extension(null,n)},Extension:function(m,n,o){if(m&&!typeof(m)==="string"){m=m.name}n=d.extensionDir+"/"+n;if(!n.match(/\.js$/)){n+=".js"}if(!j.loaded[j.fileURL(n)]){if(m!=null){delete g[o||"macros"][m.replace(/^\\/,"")]}c.RestartAfter(j.Require(n))}},Macro:function(o,r,q,s){if(q){var n=[];if(s!=null){var m=this.GetBrackets(o);n.push(m==null?s:m)}for(var p=n.length;p<q;p++){n.push(this.GetArgument(o))}r=this.SubstituteArgs(n,r)}this.string=this.AddArgs(r,this.string.slice(this.i));this.i=0;if(++this.macroCount>d.config.MAXMACROS){d.Error(["MaxMacroSub1","MathJax maximum macro substitution count exceeded; is there a recursive macro call?"])}},Matrix:function(n,p,v,r,u,o,m,w,t){var s=this.GetNext();if(s===""){d.Error(["MissingArgFor","Missing argument for %1",n])}if(s==="{"){this.i++}else{this.string=s+"}"+this.string.slice(this.i+1);this.i=0}var q=b.array().With({requireClose:true,arraydef:{rowspacing:(o||"4pt"),columnspacing:(u||"1em")}});if(w){q.isCases=true}if(t){q.isNumbered=true;q.arraydef.side=t}if(p||v){q.open=p;q.close=v}if(m==="D"){q.arraydef.displaystyle=true}if(r!=null){q.arraydef.columnalign=r}this.Push(q)},Entry:function(p){this.Push(b.cell().With({isEntry:true,name:p}));if(this.stack.Top().isCases){var o=this.string;var s=0,q=this.i,n=o.length;while(q<n){var t=o.charAt(q);if(t==="{"){s++;q++}else{if(t==="}"){if(s===0){n=0}else{s--;q++}}else{if(t==="&"&&s===0){d.Error(["ExtraAlignTab","Extra alignment tab in \\cases text"])}else{if(t==="\\"){if(o.substr(q).match(/^((\\cr)[^a-zA-Z]|\\\\)/)){n=0}else{q+=2}}else{q++}}}}}var r=o.substr(this.i,q-this.i);if(!r.match(/^\s*\\text[^a-zA-Z]/)){this.Push.apply(this,this.InternalMath(r,0));this.i=q}}},Cr:function(m){this.Push(b.cell().With({isCR:true,name:m}))},CrLaTeX:function(m){var q;if(this.string.charAt(this.i)==="["){q=this.GetBrackets(m,"").replace(/ /g,"").replace(/,/,".");if(q&&!this.matchDimen(q)){d.Error(["BracketMustBeDimension","Bracket argument to %1 must be a dimension",m])}}this.Push(b.cell().With({isCR:true,name:m,linebreak:true}));var p=this.stack.Top();if(p.isa(b.array)){if(q&&p.arraydef.rowspacing){var o=p.arraydef.rowspacing.split(/ /);if(!p.rowspacing){p.rowspacing=this.dimen2em(o[0])}while(o.length<p.table.length){o.push(this.Em(p.rowspacing))}o[p.table.length-1]=this.Em(Math.max(0,p.rowspacing+this.dimen2em(q)));p.arraydef.rowspacing=o.join(" ")}}else{if(q){this.Push(i.mspace().With({depth:q}))}this.Push(i.mspace().With({linebreak:i.LINEBREAK.NEWLINE}))}},emPerInch:7.2,pxPerInch:72,matchDimen:function(m){return m.match(/^(-?(?:\.\d+|\d+(?:\.\d*)?))(px|pt|em|ex|mu|pc|in|mm|cm)$/)},dimen2em:function(q){var o=this.matchDimen(q);var n=parseFloat(o[1]||"1"),p=o[2];if(p==="em"){return n}if(p==="ex"){return n*0.43}if(p==="pt"){return n/10}if(p==="pc"){return n*1.2}if(p==="px"){return n*this.emPerInch/this.pxPerInch}if(p==="in"){return n*this.emPerInch}if(p==="cm"){return n*this.emPerInch/2.54}if(p==="mm"){return n*this.emPerInch/25.4}if(p==="mu"){return n/18}return 0},Em:function(n){if(Math.abs(n)<0.0006){return"0em"}return n.toFixed(3).replace(/\.?0+$/,"")+"em"},HLine:function(n,o){if(o==null){o="solid"}var p=this.stack.Top();if(!p.isa(b.array)||p.data.length){d.Error(["Misplaced","Misplaced %1",n])}if(p.table.length==0){p.frame.push("top")}else{var m=(p.arraydef.rowlines?p.arraydef.rowlines.split(/ /):[]);while(m.length<p.table.length){m.push("none")}m[p.table.length-1]=o;p.arraydef.rowlines=m.join(" ")}},HFill:function(m){var n=this.stack.Top();if(n.isa(b.array)){n.hfill.push(n.data.length)}else{d.Error(["UnsupportedHFill","Unsupported use of %1",m])}},BeginEnd:function(o){var p=this.GetArgument(o),r=false;if(p.match(/^\\end\\/)){r=true;p=p.substr(5)}if(p.match(/\\/i)){d.Error(["InvalidEnv","Invalid environment name '%1'",p])}var q=this.envFindName(p);if(!q){d.Error(["UnknownEnv","Unknown environment '%1'",p])}if(!f(q)){q=[q]}var m=(f(q[1])?q[1][0]:q[1]);var n=b.begin().With({name:p,end:m,parse:this});if(o==="\\end"){if(!r&&f(q[1])&&this[q[1][1]]){n=this[q[1][1]].apply(this,[n].concat(q.slice(2)))}else{n=b.end().With({name:p})}}else{if(++this.macroCount>d.config.MAXMACROS){d.Error(["MaxMacroSub2","MathJax maximum substitution count exceeded; is there a recursive latex environment?"])}if(q[0]&&this[q[0]]){n=this[q[0]].apply(this,[n].concat(q.slice(2)))}}this.Push(n)},envFindName:function(m){return g.environment[m]},Equation:function(m,n){return n},ExtensionEnv:function(n,m){this.Extension(n.name,m,"environment")},Array:function(n,p,u,s,t,o,m,q){if(!s){s=this.GetArgument("\\begin{"+n.name+"}")}var v=("c"+s).replace(/[^clr|:]/g,"").replace(/[^|:]([|:])+/g,"$1");s=s.replace(/[^clr]/g,"").split("").join(" ");s=s.replace(/l/g,"left").replace(/r/g,"right").replace(/c/g,"center");var r=b.array().With({arraydef:{columnalign:s,columnspacing:(t||"1em"),rowspacing:(o||"4pt")}});if(v.match(/[|:]/)){if(v.charAt(0).match(/[|:]/)){r.frame.push("left");r.frame.dashed=v.charAt(0)===":"}if(v.charAt(v.length-1).match(/[|:]/)){r.frame.push("right")}v=v.substr(1,v.length-2);r.arraydef.columnlines=v.split("").join(" ").replace(/[^|: ]/g,"none").replace(/\|/g,"solid").replace(/:/g,"dashed")}if(p){r.open=this.convertDelimiter(p)}if(u){r.close=this.convertDelimiter(u)}if(m==="D"){r.arraydef.displaystyle=true}else{if(m){r.arraydef.displaystyle=false}}if(m==="S"){r.arraydef.scriptlevel=1}if(q){r.arraydef.useHeight=false}this.Push(n);return r},AlignedArray:function(m){var n=this.GetBrackets("\\begin{"+m.name+"}");return this.setArrayAlign(this.Array.apply(this,arguments),n)},setArrayAlign:function(n,m){m=this.trimSpaces(m||"");if(m==="t"){n.arraydef.align="baseline 1"}else{if(m==="b"){n.arraydef.align="baseline -1"}else{if(m==="c"){n.arraydef.align="center"}else{if(m){n.arraydef.align=m}}}}return n},convertDelimiter:function(m){if(m){m=g.delimiter[m]}if(m==null){return null}if(f(m)){m=m[0]}if(m.length===4){m=String.fromCharCode(parseInt(m,16))}return m},trimSpaces:function(n){if(typeof(n)!="string"){return n}var m=n.replace(/^\s+|\s+$/g,"");if(m.match(/\\$/)&&n.match(/ $/)){m+=" "}return m},nextIsSpace:function(){return this.string.charAt(this.i).match(/\s/)},GetNext:function(){while(this.nextIsSpace()){this.i++}return this.string.charAt(this.i)},GetCS:function(){var m=this.string.slice(this.i).match(/^([a-z]+|.) ?/i);if(m){this.i+=m[1].length;return m[1]}else{this.i++;return" "}},GetArgument:function(n,o){switch(this.GetNext()){case"":if(!o){d.Error(["MissingArgFor","Missing argument for %1",n])}return null;case"}":if(!o){d.Error(["ExtraCloseMissingOpen","Extra close brace or missing open brace"])}return null;case"\\":this.i++;return"\\"+this.GetCS();case"{":var m=++this.i,p=1;while(this.i<this.string.length){switch(this.string.charAt(this.i++)){case"\\":this.i++;break;case"{":p++;break;case"}":if(--p==0){return this.string.slice(m,this.i-1)}break}}d.Error(["MissingCloseBrace","Missing close brace"]);break}return this.string.charAt(this.i++)},GetBrackets:function(n,p){if(this.GetNext()!="["){return p}var m=++this.i,o=0;while(this.i<this.string.length){switch(this.string.charAt(this.i++)){case"{":o++;break;case"\\":this.i++;break;case"}":if(o--<=0){d.Error(["ExtraCloseLooking","Extra close brace while looking for %1","']'"])}break;case"]":if(o==0){return this.string.slice(m,this.i-1)}break}}d.Error(["MissingCloseBracket","Couldn't find closing ']' for argument to %1",n])},GetDelimiter:function(m,n){while(this.nextIsSpace()){this.i++}var o=this.string.charAt(this.i);this.i++;if(this.i<=this.string.length){if(o=="\\"){o+=this.GetCS(m)}else{if(o==="{"&&n){this.i--;o=this.GetArgument(m)}}if(g.delimiter[o]!=null){return this.convertDelimiter(o)}}d.Error(["MissingOrUnrecognizedDelim","Missing or unrecognized delimiter for %1",m])},GetDimen:function(n){var o;if(this.nextIsSpace()){this.i++}if(this.string.charAt(this.i)=="{"){o=this.GetArgument(n);if(o.match(/^\s*([-+]?([.,]\d+|\d+([.,]\d*)?))\s*(pt|em|ex|mu|px|mm|cm|in|pc)\s*$/)){return o.replace(/ /g,"").replace(/,/,".")}}else{o=this.string.slice(this.i);var m=o.match(/^\s*(([-+]?([.,]\d+|\d+([.,]\d*)?))\s*(pt|em|ex|mu|px|mm|cm|in|pc)) ?/);if(m){this.i+=m[0].length;return m[1].replace(/ /g,"").replace(/,/,".")}}d.Error(["MissingDimOrUnits","Missing dimension or its units for %1",n])},GetUpTo:function(o,p){while(this.nextIsSpace()){this.i++}var n=this.i,m,r,q=0;while(this.i<this.string.length){m=this.i;r=this.string.charAt(this.i++);switch(r){case"\\":r+=this.GetCS();break;case"{":q++;break;case"}":if(q==0){d.Error(["ExtraCloseLooking","Extra close brace while looking for %1",p])}q--;break}if(q==0&&r==p){return this.string.slice(n,m)}}d.Error(["TokenNotFoundForCommand","Couldn't find %1 for %2",p,o])},ParseArg:function(m){return d.Parse(this.GetArgument(m),this.stack.env).mml()},ParseUpTo:function(m,n){return d.Parse(this.GetUpTo(m,n),this.stack.env).mml()},InternalMath:function(v,m){var o=(this.stack.env.font?{mathvariant:this.stack.env.font}:{});var n=[],r=0,q=0,u,s="",p=0;if(v.match(/\\?[${}\\]|\\\(|\\(eq)?ref\s*\{/)){while(r<v.length){u=v.charAt(r++);if(u==="$"){if(s==="$"&&p===0){n.push(i.TeXAtom(d.Parse(v.slice(q,r-1),{}).mml()));s="";q=r}else{if(s===""){if(q<r-1){n.push(this.InternalText(v.slice(q,r-1),o))}s="$";q=r}}}else{if(u==="{"&&s!==""){p++}else{if(u==="}"){if(s==="}"&&p===0){n.push(i.TeXAtom(d.Parse(v.slice(q,r),{}).mml().With(o)));s="";q=r}else{if(s!==""){if(p){p--}}}}else{if(u==="\\"){if(s===""&&v.substr(r).match(/^(eq)?ref\s*\{/)){var t=RegExp["$&"].length;if(q<r-1){n.push(this.InternalText(v.slice(q,r-1),o))}s="}";q=r-1;r+=t}else{u=v.charAt(r++);if(u==="("&&s===""){if(q<r-2){n.push(this.InternalText(v.slice(q,r-2),o))}s=")";q=r}else{if(u===")"&&s===")"&&p===0){n.push(i.TeXAtom(d.Parse(v.slice(q,r-2),{}).mml()));s="";q=r}else{if(u.match(/[${}\\]/)&&s===""){r--;v=v.substr(0,r-1)+v.substr(r)}}}}}}}}}if(s!==""){d.Error(["MathNotTerminated","Math not terminated in text box"])}}if(q<v.length){n.push(this.InternalText(v.slice(q),o))}if(m!=null){n=[i.mstyle.apply(i,n).With({displaystyle:false,scriptlevel:m})]}else{if(n.length>1){n=[i.mrow.apply(i,n)]}}return n},InternalText:function(n,m){n=n.replace(/^\s+/,h).replace(/\s+$/,h);return i.mtext(i.chars(n)).With(m)},SubstituteArgs:function(n,m){var q="";var p="";var r;var o=0;while(o<m.length){r=m.charAt(o++);if(r==="\\"){q+=r+m.charAt(o++)}else{if(r==="#"){r=m.charAt(o++);if(r==="#"){q+=r}else{if(!r.match(/[1-9]/)||r>n.length){d.Error(["IllegalMacroParam","Illegal macro parameter reference"])}p=this.AddArgs(this.AddArgs(p,q),n[r-1]);q=""}}else{q+=r}}}return this.AddArgs(p,q)},AddArgs:function(n,m){if(m.match(/^[a-z]/i)&&n.match(/(^|[^\\])(\\\\)*\\[a-z]+$/i)){n+=" "}if(n.length+m.length>d.config.MAXBUFFER){d.Error(["MaxBufferSize","MathJax internal buffer size exceeded; is there a recursive macro call?"])}return n+m}});d.Augment({Stack:e,Parse:a,Definitions:g,Startup:l,config:{MAXMACROS:10000,MAXBUFFER:5*1024},sourceMenuTitle:["TeXCommands","TeX Commands"],annotationEncoding:"application/x-tex",prefilterHooks:MathJax.Callback.Hooks(true),postfilterHooks:MathJax.Callback.Hooks(true),Config:function(){this.SUPER(arguments).Config.apply(this,arguments);if(this.config.equationNumbers.autoNumber!=="none"){if(!this.config.extensions){this.config.extensions=[]}this.config.extensions.push("AMSmath.js")}},Translate:function(m){var n,o=false,q=MathJax.HTML.getScript(m);var s=(m.type.replace(/\n/g," ").match(/(;|\s|\n)mode\s*=\s*display(;|\s|\n|$)/)!=null);var r={math:q,display:s,script:m};var t=this.prefilterHooks.Execute(r);if(t){return t}q=r.math;try{n=d.Parse(q).mml()}catch(p){if(!p.texError){throw p}n=this.formatError(p,q,s,m);o=true}if(n.isa(i.mtable)&&n.displaystyle==="inherit"){n.displaystyle=s}if(n.inferred){n=i.apply(MathJax.ElementJax,n.data)}else{n=i(n)}if(s){n.root.display="block"}if(o){n.texError=true}r.math=n;return this.postfilterHooks.Execute(r)||r.math},prefilterMath:function(n,o,m){return n},postfilterMath:function(n,o,m){this.combineRelations(n.root);return n},formatError:function(p,o,q,m){var n=p.message.replace(/\n.*/,"");c.signal.Post(["TeX Jax - parse error",n,o,q,m]);return i.Error(n)},Error:function(m){if(f(m)){m=k.apply(k,m)}throw c.Insert(Error(m),{texError:true})},Macro:function(m,n,o){g.macros[m]=["Macro"].concat([].slice.call(arguments,1));g.macros[m].isUser=true},fenced:function(o,n,p){var m=i.mrow().With({open:o,close:p,texClass:i.TEXCLASS.INNER});m.Append(i.mo(o).With({fence:true,stretchy:true,symmetric:true,texClass:i.TEXCLASS.OPEN}),n,i.mo(p).With({fence:true,stretchy:true,symmetric:true,texClass:i.TEXCLASS.CLOSE}));return m},fixedFence:function(o,n,p){var m=i.mrow().With({open:o,close:p,texClass:i.TEXCLASS.ORD});if(o){m.Append(this.mathPalette(o,"l"))}if(n.type==="mrow"){m.Append.apply(m,n.data)}else{m.Append(n)}if(p){m.Append(this.mathPalette(p,"r"))}return m},mathPalette:function(p,n){if(p==="{"||p==="}"){p="\\"+p}var o="{\\bigg"+n+" "+p+"}",m="{\\big"+n+" "+p+"}";return d.Parse("\\mathchoice"+o+m+m+m,{}).mml()},combineRelations:function(q){var r,n,p,o;for(r=0,n=q.data.length;r<n;r++){if(q.data[r]){if(q.isa(i.mrow)){while(r+1<n&&(p=q.data[r])&&(o=q.data[r+1])&&p.isa(i.mo)&&o.isa(i.mo)&&p.Get("texClass")===i.TEXCLASS.REL&&o.Get("texClass")===i.TEXCLASS.REL){if(p.variantForm==o.variantForm&&p.Get("mathvariant")==o.Get("mathvariant")&&p.style==o.style&&p["class"]==o["class"]&&!p.id&&!o.id){p.Append.apply(p,o.data);q.data.splice(r+1,1);n--}else{p.rspace=o.lspace="0pt";r++}}}if(!q.data[r].isToken){this.combineRelations(q.data[r])}}}}});d.prefilterHooks.Add(function(m){m.math=d.prefilterMath(m.math,m.display,m.script)});d.postfilterHooks.Add(function(m){m.math=d.postfilterMath(m.math,m.display,m.script)});d.loadComplete("jax.js")})(MathJax.InputJax.TeX,MathJax.Hub,MathJax.Ajax);
MathJax.Extension["TeX/AMSmath"]={version:"2.7.0",number:0,startNumber:0,IDs:{},eqIDs:{},labels:{},eqlabels:{},refs:[]};MathJax.Hub.Register.StartupHook("TeX Jax Ready",function(){var b=MathJax.ElementJax.mml,h=MathJax.InputJax.TeX,g=MathJax.Extension["TeX/AMSmath"];var d=h.Definitions,f=h.Stack.Item,a=h.config.equationNumbers;var c=function(k){var n=[];for(var l=0,j=k.length;l<j;l++){n[l]=h.Parse.prototype.Em(k[l])}return n.join(" ")};var e=(document.getElementsByTagName("base").length===0)?"":String(document.location).replace(/#.*$/,"");d.Add({mathchar0mo:{iiiint:["2A0C",{texClass:b.TEXCLASS.OP}]},macros:{mathring:["Accent","2DA"],nobreakspace:"Tilde",negmedspace:["Spacer",b.LENGTH.NEGATIVEMEDIUMMATHSPACE],negthickspace:["Spacer",b.LENGTH.NEGATIVETHICKMATHSPACE],idotsint:["MultiIntegral","\\int\\cdots\\int"],dddot:["Accent","20DB"],ddddot:["Accent","20DC"],sideset:["Macro","\\mathop{\\mathop{\\rlap{\\phantom{#3}}}\\nolimits#1\\!\\mathop{#3}\\nolimits#2}",3],boxed:["Macro","\\fbox{$\\displaystyle{#1}$}",1],tag:"HandleTag",notag:"HandleNoTag",label:"HandleLabel",ref:"HandleRef",eqref:["HandleRef",true],substack:["Macro","\\begin{subarray}{c}#1\\end{subarray}",1],injlim:["NamedOp","inj&thinsp;lim"],projlim:["NamedOp","proj&thinsp;lim"],varliminf:["Macro","\\mathop{\\underline{\\mmlToken{mi}{lim}}}"],varlimsup:["Macro","\\mathop{\\overline{\\mmlToken{mi}{lim}}}"],varinjlim:["Macro","\\mathop{\\underrightarrow{\\mmlToken{mi}{lim}}}"],varprojlim:["Macro","\\mathop{\\underleftarrow{\\mmlToken{mi}{lim}}}"],DeclareMathOperator:"HandleDeclareOp",operatorname:"HandleOperatorName",SkipLimits:"SkipLimits",genfrac:"Genfrac",frac:["Genfrac","","","",""],tfrac:["Genfrac","","","",1],dfrac:["Genfrac","","","",0],binom:["Genfrac","(",")","0",""],tbinom:["Genfrac","(",")","0",1],dbinom:["Genfrac","(",")","0",0],cfrac:"CFrac",shoveleft:["HandleShove",b.ALIGN.LEFT],shoveright:["HandleShove",b.ALIGN.RIGHT],xrightarrow:["xArrow",8594,5,6],xleftarrow:["xArrow",8592,7,3]},environment:{align:["AMSarray",null,true,true,"rlrlrlrlrlrl",c([0,2,0,2,0,2,0,2,0,2,0])],"align*":["AMSarray",null,false,true,"rlrlrlrlrlrl",c([0,2,0,2,0,2,0,2,0,2,0])],multline:["Multline",null,true],"multline*":["Multline",null,false],split:["AMSarray",null,false,false,"rl",c([0])],gather:["AMSarray",null,true,true,"c"],"gather*":["AMSarray",null,false,true,"c"],alignat:["AlignAt",null,true,true],"alignat*":["AlignAt",null,false,true],alignedat:["AlignAt",null,false,false],aligned:["AlignedAMSArray",null,null,null,"rlrlrlrlrlrl",c([0,2,0,2,0,2,0,2,0,2,0]),".5em","D"],gathered:["AlignedAMSArray",null,null,null,"c",null,".5em","D"],subarray:["Array",null,null,null,null,c([0]),"0.1em","S",1],smallmatrix:["Array",null,null,null,"c",c([1/3]),".2em","S",1],equation:["EquationBegin","Equation",true],"equation*":["EquationBegin","EquationStar",false],eqnarray:["AMSarray",null,true,true,"rcl","0 "+b.LENGTH.THICKMATHSPACE,".5em"],"eqnarray*":["AMSarray",null,false,true,"rcl","0 "+b.LENGTH.THICKMATHSPACE,".5em"]},delimiter:{"\\lvert":["007C",{texClass:b.TEXCLASS.OPEN}],"\\rvert":["007C",{texClass:b.TEXCLASS.CLOSE}],"\\lVert":["2016",{texClass:b.TEXCLASS.OPEN}],"\\rVert":["2016",{texClass:b.TEXCLASS.CLOSE}]}},null,true);h.Parse.Augment({HandleTag:function(k){var m=this.GetStar();var j=this.trimSpaces(this.GetArgument(k)),i=j;if(!m){j=a.formatTag(j)}var l=this.stack.global;l.tagID=i;if(l.notags){h.Error(["CommandNotAllowedInEnv","%1 not allowed in %2 environment",k,l.notags])}if(l.tag){h.Error(["MultipleCommand","Multiple %1",k])}l.tag=b.mtd.apply(b,this.InternalMath(j)).With({id:a.formatID(i)})},HandleNoTag:function(i){if(this.stack.global.tag){delete this.stack.global.tag}this.stack.global.notag=true},HandleLabel:function(j){var k=this.stack.global,i=this.GetArgument(j);if(i===""){return}if(!g.refUpdate){if(k.label){h.Error(["MultipleCommand","Multiple %1",j])}k.label=i;if(g.labels[i]||g.eqlabels[i]){h.Error(["MultipleLabel","Label '%1' multiply defined",i])}g.eqlabels[i]={tag:"???",id:""}}},HandleRef:function(k,m){var j=this.GetArgument(k);var l=g.labels[j]||g.eqlabels[j];if(!l){l={tag:"???",id:""};g.badref=!g.refUpdate}var i=l.tag;if(m){i=a.formatTag(i)}this.Push(b.mrow.apply(b,this.InternalMath(i)).With({href:a.formatURL(l.id,e),"class":"MathJax_ref"}))},HandleDeclareOp:function(j){var i=(this.GetStar()?"":"\\nolimits\\SkipLimits");var k=this.trimSpaces(this.GetArgument(j));if(k.charAt(0)=="\\"){k=k.substr(1)}var l=this.GetArgument(j);l=l.replace(/\*/g,"\\text{*}").replace(/-/g,"\\text{-}");h.Definitions.macros[k]=["Macro","\\mathop{\\rm "+l+"}"+i]},HandleOperatorName:function(j){var i=(this.GetStar()?"":"\\nolimits\\SkipLimits");var k=this.trimSpaces(this.GetArgument(j));k=k.replace(/\*/g,"\\text{*}").replace(/-/g,"\\text{-}");this.string="\\mathop{\\rm "+k+"}"+i+" "+this.string.slice(this.i);this.i=0},SkipLimits:function(j){var l=this.GetNext(),k=this.i;if(l==="\\"&&++this.i&&this.GetCS()!=="limits"){this.i=k}},HandleShove:function(j,i){var k=this.stack.Top();if(k.type!=="multline"||k.data.length){h.Error(["CommandAtTheBeginingOfLine","%1 must come at the beginning of the line",j])}k.data.shove=i},CFrac:function(l){var i=this.trimSpaces(this.GetBrackets(l,"")),k=this.GetArgument(l),m=this.GetArgument(l);var j=b.mfrac(h.Parse("\\strut\\textstyle{"+k+"}",this.stack.env).mml(),h.Parse("\\strut\\textstyle{"+m+"}",this.stack.env).mml());i=({l:b.ALIGN.LEFT,r:b.ALIGN.RIGHT,"":""})[i];if(i==null){h.Error(["IllegalAlign","Illegal alignment specified in %1",l])}if(i){j.numalign=j.denomalign=i}this.Push(j)},Genfrac:function(j,l,q,n,i){if(l==null){l=this.GetDelimiterArg(j)}if(q==null){q=this.GetDelimiterArg(j)}if(n==null){n=this.GetArgument(j)}if(i==null){i=this.trimSpaces(this.GetArgument(j))}var m=this.ParseArg(j);var p=this.ParseArg(j);var k=b.mfrac(m,p);if(n!==""){k.linethickness=n}if(l||q){k=h.fixedFence(l,k.With({texWithDelims:true}),q)}if(i!==""){var o=(["D","T","S","SS"])[i];if(o==null){h.Error(["BadMathStyleFor","Bad math style for %1",j])}k=b.mstyle(k);if(o==="D"){k.displaystyle=true;k.scriptlevel=0}else{k.displaystyle=false;k.scriptlevel=i-1}}this.Push(k)},Multline:function(j,i){this.Push(j);this.checkEqnEnv();return f.multline(i,this.stack).With({arraydef:{displaystyle:true,rowspacing:".5em",width:h.config.MultLineWidth,columnwidth:"100%",side:h.config.TagSide,minlabelspacing:h.config.TagIndent}})},AMSarray:function(k,j,i,m,l){this.Push(k);if(i){this.checkEqnEnv()}m=m.replace(/[^clr]/g,"").split("").join(" ");m=m.replace(/l/g,"left").replace(/r/g,"right").replace(/c/g,"center");return f.AMSarray(k.name,j,i,this.stack).With({arraydef:{displaystyle:true,rowspacing:".5em",columnalign:m,columnspacing:(l||"1em"),rowspacing:"3pt",side:h.config.TagSide,minlabelspacing:h.config.TagIndent}})},AlignedAMSArray:function(i){var j=this.GetBrackets("\\begin{"+i.name+"}");return this.setArrayAlign(this.AMSarray.apply(this,arguments),j)},AlignAt:function(l,j,i){var q,k,p="",o=[];if(!i){k=this.GetBrackets("\\begin{"+l.name+"}")}q=this.GetArgument("\\begin{"+l.name+"}");if(q.match(/[^0-9]/)){h.Error(["PositiveIntegerArg","Argument to %1 must me a positive integer","\\begin{"+l.name+"}"])}while(q>0){p+="rl";o.push("0em 0em");q--}o=o.join(" ");if(i){return this.AMSarray(l,j,i,p,o)}var m=this.AMSarray(l,j,i,p,o);return this.setArrayAlign(m,k)},EquationBegin:function(i,j){this.checkEqnEnv();this.stack.global.forcetag=(j&&a.autoNumber!=="none");return i},EquationStar:function(i,j){this.stack.global.tagged=true;return j},checkEqnEnv:function(){if(this.stack.global.eqnenv){h.Error(["ErroneousNestingEq","Erroneous nesting of equation structures"])}this.stack.global.eqnenv=true},MultiIntegral:function(j,m){var l=this.GetNext();if(l==="\\"){var k=this.i;l=this.GetArgument(j);this.i=k;if(l==="\\limits"){if(j==="\\idotsint"){m="\\!\\!\\mathop{\\,\\,"+m+"}"}else{m="\\!\\!\\!\\mathop{\\,\\,\\,"+m+"}"}}}this.string=m+" "+this.string.slice(this.i);this.i=0},xArrow:function(k,o,n,i){var m={width:"+"+(n+i)+"mu",lspace:n+"mu"};var p=this.GetBrackets(k),q=this.ParseArg(k);var s=b.mo(b.chars(String.fromCharCode(o))).With({stretchy:true,texClass:b.TEXCLASS.REL});var j=b.munderover(s);j.SetData(j.over,b.mpadded(q).With(m).With({voffset:".15em"}));if(p){p=h.Parse(p,this.stack.env).mml();j.SetData(j.under,b.mpadded(p).With(m).With({voffset:"-.24em"}))}this.Push(j.With({subsupOK:true}))},GetDelimiterArg:function(i){var j=this.trimSpaces(this.GetArgument(i));if(j==""){return null}if(j in d.delimiter){return j}h.Error(["MissingOrUnrecognizedDelim","Missing or unrecognized delimiter for %1",i])},GetStar:function(){var i=(this.GetNext()==="*");if(i){this.i++}return i}});f.Augment({autoTag:function(){var j=this.global;if(!j.notag){g.number++;j.tagID=a.formatNumber(g.number.toString());var i=h.Parse("\\text{"+a.formatTag(j.tagID)+"}",{}).mml();j.tag=b.mtd(i).With({id:a.formatID(j.tagID)})}},getTag:function(){var m=this.global,k=m.tag;m.tagged=true;if(m.label){if(a.useLabelIds){k.id=a.formatID(m.label)}g.eqlabels[m.label]={tag:m.tagID,id:k.id}}if(document.getElementById(k.id)||g.IDs[k.id]||g.eqIDs[k.id]){var l=0,j;do{l++;j=k.id+"_"+l}while(document.getElementById(j)||g.IDs[j]||g.eqIDs[j]);k.id=j;if(m.label){g.eqlabels[m.label].id=j}}g.eqIDs[k.id]=1;this.clearTag();return k},clearTag:function(){var i=this.global;delete i.tag;delete i.tagID;delete i.label},fixInitialMO:function(l){for(var k=0,j=l.length;k<j;k++){if(l[k]&&(l[k].type!=="mspace"&&(l[k].type!=="texatom"||(l[k].data[0]&&l[k].data[0].data.length)))){if(l[k].isEmbellished()){l.unshift(b.mi())}break}}}});f.multline=f.array.Subclass({type:"multline",Init:function(j,i){this.SUPER(arguments).Init.apply(this);this.numbered=(j&&a.autoNumber!=="none");this.save={notag:i.global.notag};i.global.tagged=!j&&!i.global.forcetag},EndEntry:function(){if(this.table.length){this.fixInitialMO(this.data)}var i=b.mtd.apply(b,this.data);if(this.data.shove){i.columnalign=this.data.shove}this.row.push(i);this.data=[]},EndRow:function(){if(this.row.length!=1){h.Error(["MultlineRowsOneCol","The rows within the %1 environment must have exactly one column","multline"])}this.table.push(this.row);this.row=[]},EndTable:function(){this.SUPER(arguments).EndTable.call(this);if(this.table.length){var k=this.table.length-1,n,l=-1;if(!this.table[0][0].columnalign){this.table[0][0].columnalign=b.ALIGN.LEFT}if(!this.table[k][0].columnalign){this.table[k][0].columnalign=b.ALIGN.RIGHT}if(!this.global.tag&&this.numbered){this.autoTag()}if(this.global.tag&&!this.global.notags){l=(this.arraydef.side==="left"?0:this.table.length-1);this.table[l]=[this.getTag()].concat(this.table[l])}for(n=0,k=this.table.length;n<k;n++){var j=(n===l?b.mlabeledtr:b.mtr);this.table[n]=j.apply(b,this.table[n])}}this.global.notag=this.save.notag}});f.AMSarray=f.array.Subclass({type:"AMSarray",Init:function(l,k,j,i){this.SUPER(arguments).Init.apply(this);this.numbered=(k&&a.autoNumber!=="none");this.save={notags:i.global.notags,notag:i.global.notag};i.global.notags=(j?null:l);i.global.tagged=!k&&!i.global.forcetag},EndEntry:function(){if(this.row.length){this.fixInitialMO(this.data)}this.row.push(b.mtd.apply(b,this.data));this.data=[]},EndRow:function(){var i=b.mtr;if(!this.global.tag&&this.numbered){this.autoTag()}if(this.global.tag&&!this.global.notags){this.row=[this.getTag()].concat(this.row);i=b.mlabeledtr}else{this.clearTag()}if(this.numbered){delete this.global.notag}this.table.push(i.apply(b,this.row));this.row=[]},EndTable:function(){this.SUPER(arguments).EndTable.call(this);this.global.notags=this.save.notags;this.global.notag=this.save.notag}});f.start.Augment({oldCheckItem:f.start.prototype.checkItem,checkItem:function(k){if(k.type==="stop"){var i=this.mmlData(),j=this.global;if(g.display&&!j.tag&&!j.tagged&&!j.isInner&&(a.autoNumber==="all"||j.forcetag)){this.autoTag()}if(j.tag){var m=[this.getTag(),b.mtd(i)];var l={side:h.config.TagSide,minlabelspacing:h.config.TagIndent,displaystyle:"inherit"};i=b.mtable(b.mlabeledtr.apply(b,m)).With(l)}return f.mml(i)}return this.oldCheckItem.call(this,k)}});h.prefilterHooks.Add(function(i){g.display=i.display;g.number=g.startNumber;g.eqlabels={};g.eqIDs={};g.badref=false;if(g.refUpdate){g.number=i.script.MathJax.startNumber}});h.postfilterHooks.Add(function(i){i.script.MathJax.startNumber=g.startNumber;g.startNumber=g.number;MathJax.Hub.Insert(g.IDs,g.eqIDs);MathJax.Hub.Insert(g.labels,g.eqlabels);if(g.badref&&!i.math.texError){g.refs.push(i.script)}},100);MathJax.Hub.Register.MessageHook("Begin Math Input",function(){g.refs=[];g.refUpdate=false});MathJax.Hub.Register.MessageHook("End Math Input",function(l){if(g.refs.length){g.refUpdate=true;for(var k=0,j=g.refs.length;k<j;k++){g.refs[k].MathJax.state=MathJax.ElementJax.STATE.UPDATE}return MathJax.Hub.processInput({scripts:g.refs,start:new Date().getTime(),i:0,j:0,jax:{},jaxIDs:[]})}return null});h.resetEquationNumbers=function(j,i){g.startNumber=(j||0);if(!i){g.labels={};g.IDs={}}};MathJax.Hub.Startup.signal.Post("TeX AMSmath Ready")});MathJax.Ajax.loadComplete("[MathJax]/extensions/TeX/AMSmath.js");
MathJax.Extension["TeX/AMSsymbols"]={version:"2.7.0"};MathJax.Hub.Register.StartupHook("TeX Jax Ready",function(){var a=MathJax.ElementJax.mml,b=MathJax.InputJax.TeX.Definitions;b.Add({mathchar0mi:{digamma:"03DD",varkappa:"03F0",varGamma:["0393",{mathvariant:a.VARIANT.ITALIC}],varDelta:["0394",{mathvariant:a.VARIANT.ITALIC}],varTheta:["0398",{mathvariant:a.VARIANT.ITALIC}],varLambda:["039B",{mathvariant:a.VARIANT.ITALIC}],varXi:["039E",{mathvariant:a.VARIANT.ITALIC}],varPi:["03A0",{mathvariant:a.VARIANT.ITALIC}],varSigma:["03A3",{mathvariant:a.VARIANT.ITALIC}],varUpsilon:["03A5",{mathvariant:a.VARIANT.ITALIC}],varPhi:["03A6",{mathvariant:a.VARIANT.ITALIC}],varPsi:["03A8",{mathvariant:a.VARIANT.ITALIC}],varOmega:["03A9",{mathvariant:a.VARIANT.ITALIC}],beth:"2136",gimel:"2137",daleth:"2138",backprime:["2035",{variantForm:true}],hslash:"210F",varnothing:["2205",{variantForm:true}],blacktriangle:"25B4",triangledown:["25BD",{variantForm:true}],blacktriangledown:"25BE",square:"25FB",Box:"25FB",blacksquare:"25FC",lozenge:"25CA",Diamond:"25CA",blacklozenge:"29EB",circledS:["24C8",{mathvariant:a.VARIANT.NORMAL}],bigstar:"2605",sphericalangle:"2222",measuredangle:"2221",nexists:"2204",complement:"2201",mho:"2127",eth:["00F0",{mathvariant:a.VARIANT.NORMAL}],Finv:"2132",diagup:"2571",Game:"2141",diagdown:"2572",Bbbk:["006B",{mathvariant:a.VARIANT.DOUBLESTRUCK}],yen:"00A5",circledR:"00AE",checkmark:"2713",maltese:"2720"},mathchar0mo:{dotplus:"2214",ltimes:"22C9",smallsetminus:"2216",rtimes:"22CA",Cap:"22D2",doublecap:"22D2",leftthreetimes:"22CB",Cup:"22D3",doublecup:"22D3",rightthreetimes:"22CC",barwedge:"22BC",curlywedge:"22CF",veebar:"22BB",curlyvee:"22CE",doublebarwedge:"2A5E",boxminus:"229F",circleddash:"229D",boxtimes:"22A0",circledast:"229B",boxdot:"22A1",circledcirc:"229A",boxplus:"229E",centerdot:["22C5",{variantForm:true}],divideontimes:"22C7",intercal:"22BA",leqq:"2266",geqq:"2267",leqslant:"2A7D",geqslant:"2A7E",eqslantless:"2A95",eqslantgtr:"2A96",lesssim:"2272",gtrsim:"2273",lessapprox:"2A85",gtrapprox:"2A86",approxeq:"224A",lessdot:"22D6",gtrdot:"22D7",lll:"22D8",llless:"22D8",ggg:"22D9",gggtr:"22D9",lessgtr:"2276",gtrless:"2277",lesseqgtr:"22DA",gtreqless:"22DB",lesseqqgtr:"2A8B",gtreqqless:"2A8C",doteqdot:"2251",Doteq:"2251",eqcirc:"2256",risingdotseq:"2253",circeq:"2257",fallingdotseq:"2252",triangleq:"225C",backsim:"223D",thicksim:["223C",{variantForm:true}],backsimeq:"22CD",thickapprox:["2248",{variantForm:true}],subseteqq:"2AC5",supseteqq:"2AC6",Subset:"22D0",Supset:"22D1",sqsubset:"228F",sqsupset:"2290",preccurlyeq:"227C",succcurlyeq:"227D",curlyeqprec:"22DE",curlyeqsucc:"22DF",precsim:"227E",succsim:"227F",precapprox:"2AB7",succapprox:"2AB8",vartriangleleft:"22B2",lhd:"22B2",vartriangleright:"22B3",rhd:"22B3",trianglelefteq:"22B4",unlhd:"22B4",trianglerighteq:"22B5",unrhd:"22B5",vDash:"22A8",Vdash:"22A9",Vvdash:"22AA",smallsmile:["2323",{variantForm:true}],shortmid:["2223",{variantForm:true}],smallfrown:["2322",{variantForm:true}],shortparallel:["2225",{variantForm:true}],bumpeq:"224F",between:"226C",Bumpeq:"224E",pitchfork:"22D4",varpropto:"221D",backepsilon:"220D",blacktriangleleft:"25C2",blacktriangleright:"25B8",therefore:"2234",because:"2235",eqsim:"2242",vartriangle:["25B3",{variantForm:true}],Join:"22C8",nless:"226E",ngtr:"226F",nleq:"2270",ngeq:"2271",nleqslant:["2A87",{variantForm:true}],ngeqslant:["2A88",{variantForm:true}],nleqq:["2270",{variantForm:true}],ngeqq:["2271",{variantForm:true}],lneq:"2A87",gneq:"2A88",lneqq:"2268",gneqq:"2269",lvertneqq:["2268",{variantForm:true}],gvertneqq:["2269",{variantForm:true}],lnsim:"22E6",gnsim:"22E7",lnapprox:"2A89",gnapprox:"2A8A",nprec:"2280",nsucc:"2281",npreceq:["22E0",{variantForm:true}],nsucceq:["22E1",{variantForm:true}],precneqq:"2AB5",succneqq:"2AB6",precnsim:"22E8",succnsim:"22E9",precnapprox:"2AB9",succnapprox:"2ABA",nsim:"2241",ncong:"2246",nshortmid:["2224",{variantForm:true}],nshortparallel:["2226",{variantForm:true}],nmid:"2224",nparallel:"2226",nvdash:"22AC",nvDash:"22AD",nVdash:"22AE",nVDash:"22AF",ntriangleleft:"22EA",ntriangleright:"22EB",ntrianglelefteq:"22EC",ntrianglerighteq:"22ED",nsubseteq:"2288",nsupseteq:"2289",nsubseteqq:["2288",{variantForm:true}],nsupseteqq:["2289",{variantForm:true}],subsetneq:"228A",supsetneq:"228B",varsubsetneq:["228A",{variantForm:true}],varsupsetneq:["228B",{variantForm:true}],subsetneqq:"2ACB",supsetneqq:"2ACC",varsubsetneqq:["2ACB",{variantForm:true}],varsupsetneqq:["2ACC",{variantForm:true}],leftleftarrows:"21C7",rightrightarrows:"21C9",leftrightarrows:"21C6",rightleftarrows:"21C4",Lleftarrow:"21DA",Rrightarrow:"21DB",twoheadleftarrow:"219E",twoheadrightarrow:"21A0",leftarrowtail:"21A2",rightarrowtail:"21A3",looparrowleft:"21AB",looparrowright:"21AC",leftrightharpoons:"21CB",rightleftharpoons:["21CC",{variantForm:true}],curvearrowleft:"21B6",curvearrowright:"21B7",circlearrowleft:"21BA",circlearrowright:"21BB",Lsh:"21B0",Rsh:"21B1",upuparrows:"21C8",downdownarrows:"21CA",upharpoonleft:"21BF",upharpoonright:"21BE",downharpoonleft:"21C3",restriction:"21BE",multimap:"22B8",downharpoonright:"21C2",leftrightsquigarrow:"21AD",rightsquigarrow:"21DD",leadsto:"21DD",dashrightarrow:"21E2",dashleftarrow:"21E0",nleftarrow:"219A",nrightarrow:"219B",nLeftarrow:"21CD",nRightarrow:"21CF",nleftrightarrow:"21AE",nLeftrightarrow:"21CE"},delimiter:{"\\ulcorner":"231C","\\urcorner":"231D","\\llcorner":"231E","\\lrcorner":"231F"},macros:{implies:["Macro","\\;\\Longrightarrow\\;"],impliedby:["Macro","\\;\\Longleftarrow\\;"]}},null,true);var c=a.mo.OPTYPES.REL;MathJax.Hub.Insert(a.mo.prototype,{OPTABLE:{infix:{"\u2322":c,"\u2323":c,"\u25B3":c,"\uE006":c,"\uE007":c,"\uE00C":c,"\uE00D":c,"\uE00E":c,"\uE00F":c,"\uE010":c,"\uE011":c,"\uE016":c,"\uE017":c,"\uE018":c,"\uE019":c,"\uE01A":c,"\uE01B":c,"\uE04B":c,"\uE04F":c}}});MathJax.Hub.Startup.signal.Post("TeX AMSsymbols Ready")});MathJax.Ajax.loadComplete("[MathJax]/extensions/TeX/AMSsymbols.js");
(function(j,b,d){var i,k=b.Browser.isMobile;var h=MathJax.Object.isArray;var e=function(){var m=[].slice.call(arguments,0);m[0][0]=["HTML-CSS",m[0][0]];return MathJax.Message.Set.apply(MathJax.Message,m)};var f=MathJax.Object.Subclass({timeout:(k?15:8)*1000,comparisonFont:["sans-serif","monospace","script","Times","Courier","Arial","Helvetica"],testSize:["40px","50px","60px","30px","20px"],FedoraSTIXcheck:{family:"STIXSizeOneSym",testString:"abcABC",noStyleChar:true},Init:function(){this.div=MathJax.HTML.addElement(document.body,"div",{style:{position:"absolute",width:0,height:0,overflow:"hidden",padding:0,border:0,margin:0}},[["div",{id:"MathJax_Font_Test",style:{position:"absolute",visibility:"hidden",top:0,left:0,width:"auto",padding:0,border:0,margin:0,whiteSpace:"nowrap",textAlign:"left",textIndent:0,textTransform:"none",lineHeight:"normal",letterSpacing:"normal",wordSpacing:"normal",fontSize:this.testSize[0],fontWeight:"normal",fontStyle:"normal",fontSizeAdjust:"none"}},[""]]]).firstChild;this.text=this.div.firstChild},findFont:function(r,o){var q=null;if(o&&this.testCollection(o)){q=o}else{for(var p=0,n=r.length;p<n;p++){if(r[p]===o){continue}if(this.testCollection(r[p])){q=r[p];break}}}if(q==="STIX"&&this.testFont(this.FedoraSTIXcheck)){q=null}return q},testCollection:function(n){var m={testString:"() {} []"};m.family={TeX:"MathJax_Size1",STIX:"STIXSizeOneSym"}[n]||n.replace(/-(Math)?/,"")+"MathJax_Size1";if(n==="STIX"){m.noStyleChar=true}return this.testFont(m)},testFont:function(p){if(p.isWebFont&&d.FontFaceBug){this.div.style.fontWeight=this.div.style.fontStyle="normal"}else{this.div.style.fontWeight=(p.weight||"normal");this.div.style.fontStyle=(p.style||"normal")}var r=p.familyFixed||p.family;if(!p.isWebFont&&!r.match(/^(STIX|MathJax)|'/)){r=r.replace(/_/g," ").replace(/([a-z])([A-Z])/g,"$1 $2").replace(/ Jax/,"Jax")+"','"+r+"','"+r+"-";if(p.weight){r+="Bold"}if(p.style){r+="Italic"}if(!p.weight&&!p.style){r+="Regular"}p.familyFixed=r="'"+r+"'"}var o=this.getComparisonWidths(p.testString,p.noStyleChar);var s=null;if(o){this.div.style.fontFamily=r+","+this.comparisonFont[0];if(this.div.offsetWidth==o[0]){this.div.style.fontFamily=r+","+this.comparisonFont[o[2]];if(this.div.offsetWidth==o[1]){s=false}}if(s===null&&(this.div.offsetWidth!=o[3]||this.div.offsetHeight!=o[4])){if(!p.noStyleChar&&d.FONTDATA&&d.FONTDATA.hasStyleChar){for(var q=0,n=this.testSize.length;q<n;q++){if(this.testStyleChar(p,this.testSize[q])){s=true;n=0}}}else{s=true}}}if(d.safariTextNodeBug){this.div.innerHTML=""}else{this.text.nodeValue=""}return s},styleChar:"\uEFFD",versionChar:"\uEFFE",compChar:"\uEFFF",testStyleChar:function(p,r){var u=3+(p.weight?2:0)+(p.style?4:0);var o="",q=0;var t=this.div.style.fontSize;this.div.style.fontSize=r;if(d.msieItalicWidthBug&&p.style==="italic"){this.text.nodeValue=o=this.compChar;q=this.div.offsetWidth}if(d.safariTextNodeBug){this.div.innerHTML=this.compChar+o}else{this.text.nodeValue=this.compChar+o}var m=this.div.offsetWidth-q;if(d.safariTextNodeBug){this.div.innerHTML=this.styleChar+o}else{this.text.nodeValue=this.styleChar+o}var s=Math.floor((this.div.offsetWidth-q)/m+0.5);if(s===u){if(d.safariTextNodeBug){this.div.innerHTML=this.versionChar+o}else{this.text.nodeValue=this.versionChar+o}p.version=Math.floor((this.div.offsetWidth-q)/m+1.5)/2}this.div.style.fontSize=t;return(s===u)},getComparisonWidths:function(r,p){if(d.FONTDATA&&d.FONTDATA.hasStyleChar&&!p){r+=this.styleChar+" "+this.compChar}if(d.safariTextNodeBug){this.div.innerHTML=r}else{this.text.nodeValue=r}this.div.style.fontFamily=this.comparisonFont[0];var o=this.div.offsetWidth;this.div.style.fontFamily=d.webFontDefault;var t=this.div.offsetWidth,q=this.div.offsetHeight;for(var s=1,n=this.comparisonFont.length;s<n;s++){this.div.style.fontFamily=this.comparisonFont[s];if(this.div.offsetWidth!=o){return[o,this.div.offsetWidth,s,t,q]}}return null},loadWebFont:function(o){b.Startup.signal.Post("HTML-CSS Jax - Web-Font "+d.fontInUse+"/"+o.directory);var q=e(["LoadWebFont","Loading web-font %1",d.fontInUse+"/"+o.directory]);var m=MathJax.Callback({});var p=MathJax.Callback(["loadComplete",this,o,q,m]);j.timer.start(j,[this.checkWebFont,o,p],0,this.timeout);return m},loadComplete:function(p,r,o,m){MathJax.Message.Clear(r);if(m===j.STATUS.OK){this.webFontLoaded=true;o();return}this.loadError(p);if(b.Browser.isFirefox&&d.allowWebFonts){var q=document.location.protocol+"//"+document.location.hostname;if(document.location.port!=""){q+=":"+document.location.port}q+="/";if(j.fileURL(d.webfontDir).substr(0,q.length)!==q){this.firefoxFontError(p)}}if(!this.webFontLoaded){d.loadWebFontError(p,o)}else{o()}},loadError:function(m){e(["CantLoadWebFont","Can't load web font %1",d.fontInUse+"/"+m.directory],null,2000);b.Startup.signal.Post(["HTML-CSS Jax - web font error",d.fontInUse+"/"+m.directory,m])},firefoxFontError:function(m){e(["FirefoxCantLoadWebFont","Firefox can't load web fonts from a remote host"],null,3000);b.Startup.signal.Post("HTML-CSS Jax - Firefox web fonts on remote host error")},checkWebFont:function(m,n,o){if(m.time(o)){return}if(d.Font.testFont(n)){o(m.STATUS.OK)}else{setTimeout(m,m.delay)}},fontFace:function(q){var r=d.allowWebFonts;var t=d.FONTDATA.FONTS[q];if(d.msieFontCSSBug&&!t.family.match(/-Web$/)){t.family+="-Web"}if(t.isWebFont){delete t.familyFixed}var m=d.webfontDir+"/"+r;var p=j.fileURL(m);var o=q.replace(/-b/,"-B").replace(/-i/,"-I").replace(/-Bold-/,"-Bold");if(!o.match(/-/)){o+="-Regular"}if(r==="svg"){o+=".svg#"+o}else{o+="."+r}var n=j.fileRev(m+"/"+o.replace(/#.*/,""));var s={"font-family":t.family,src:"url('"+p+"/"+o+n+"')"};if(r==="otf"){o=o.replace(/otf$/,"woff");n=j.fileRev(m+"/"+o);s.src+=" format('opentype')";p=j.fileURL(d.webfontDir+"/woff");s.src="url('"+p+"/"+o+n+"') format('woff'), "+s.src}else{if(r!=="eot"){s.src+=" format('"+r+"')"}}if(!(d.FontFaceBug&&t.isWebFont)){if(q.match(/-bold/)){s["font-weight"]="bold"}if(q.match(/-italic/)){s["font-style"]="italic"}}return s}});var l,a,c;var g=MathJax.Hub.Browser.isMSIE&&(document.documentMode||0)<8;d.Augment({config:{styles:{".MathJax":{display:"inline","font-style":"normal","font-weight":"normal","line-height":"normal","font-size":"100%","font-size-adjust":"none","text-indent":0,"text-align":"left","text-transform":"none","letter-spacing":"normal","word-spacing":"normal","word-wrap":"normal","white-space":"nowrap","float":"none",direction:"ltr","max-width":"none","max-height":"none","min-width":0,"min-height":0,border:0,padding:0,margin:0},".MathJax:focus, body :focus .MathJax":{display:"inline-table"},".MathJax_Display":{position:"relative",display:"block!important","text-indent":0,"max-width":"none","max-height":"none","min-width":0,"min-height":0,width:"100%"},".MathJax.MathJax_FullWidth":{"text-align":"center",display:(g?"block":"table-cell")+"!important",width:(g?"100%":"10000em")+"!important"},".MathJax img, .MathJax nobr, .MathJax a":{border:0,padding:0,margin:0,"max-width":"none","max-height":"none","min-width":0,"min-height":0,"vertical-align":0,"line-height":"normal","text-decoration":"none"},"img.MathJax_strut":{border:"0!important",padding:"0!important",margin:"0!important","vertical-align":"0!important"},".MathJax span":{display:"inline",position:"static",border:0,padding:0,margin:0,"vertical-align":0,"line-height":"normal","text-decoration":"none"},".MathJax nobr":{"white-space":"nowrap!important"},".MathJax img":{display:"inline!important","float":"none!important"},".MathJax *":{transition:"none","-webkit-transition":"none","-moz-transition":"none","-ms-transition":"none","-o-transition":"none"},".MathJax_Processing":{visibility:"hidden",position:"fixed",width:0,height:0,overflow:"hidden"},".MathJax_Processed":{display:"none!important"},".MathJax_ExBox":{display:"block!important",overflow:"hidden",width:"1px",height:"60ex","min-height":0,"max-height":"none"},".MathJax .MathJax_EmBox":{display:"block!important",overflow:"hidden",width:"1px",height:"60em","min-height":0,"max-height":"none"},".MathJax_LineBox":{display:(g?"block":"table")+"!important"},".MathJax_LineBox span":{display:(g?"block":"table-cell")+"!important",width:(g?"100%":"10000em")+"!important","min-width":0,"max-width":"none",padding:0,border:0,margin:0},".MathJax .MathJax_HitBox":{cursor:"text",background:"white",opacity:0,filter:"alpha(opacity=0)"},".MathJax .MathJax_HitBox *":{filter:"none",opacity:1,background:"transparent"},"#MathJax_Tooltip":{position:"absolute",left:0,top:0,width:"auto",height:"auto",display:"none"},"#MathJax_Tooltip *":{filter:"none",opacity:1,background:"transparent"},"@font-face":{"font-family":"MathJax_Blank",src:"url('about:blank')"}}},settings:b.config.menuSettings,Font:null,webFontDefault:"MathJax_Blank",allowWebFonts:"otf",maxStretchyParts:1000,fontName:{TeXLocal:"TeX",TeXWeb:["","TeX"],TeXImage:["",""],STIXLocal:["STIX","STIX-Web"],STIXWeb:"STIX-Web",AsanaMathWeb:"Asana-Math",GyrePagellaWeb:"Gyre-Pagella",GyreTermesWeb:"Gyre-Termes",LatinModernWeb:"Latin-Modern",NeoEulerWeb:"Neo-Euler"},fontInUse:"generic",FONTDATA:{TeX_factor:1,baselineskip:1.2,lineH:0.8,lineD:0.2,ffLineH:0.8,FONTS:{},VARIANT:{normal:{fonts:[]},"-generic-variant":{},"-largeOp":{},"-smallOp":{}},RANGES:[],DELIMITERS:{},RULECHAR:45,REMAP:{}},Config:function(){if(!this.require){this.require=[]}this.Font=f();this.SUPER(arguments).Config.call(this);var o=this.settings,n=this.config,m=o.font;if(this.adjustAvailableFonts){this.adjustAvailableFonts(n.availableFonts)}if(o.scale){n.scale=o.scale}if(m&&m!=="Auto"&&this.fontName[m]){n.availableFonts=[];delete n.fonts;if(h(this.fontName[m])){n.preferredFont=this.fontName[m][0];n.webFont=this.fontName[m][1]}else{n.preferredFont=n.webFont=this.fontName[m]}if(n.preferredFont){n.availableFonts[0]=n.preferredFont}}if(n.fonts){n.availableFonts=n.fonts;n.preferredFont=n.webFont=n.fonts[0];if(n.webFont==="STIX"){n.webFont+="-Web"}}m=this.Font.findFont(n.availableFonts,n.preferredFont);if(!m&&this.allowWebFonts){m=n.webFont;if(m){this.webFonts=true}}if(!m&&this.config.imageFont){m=n.imageFont;this.imgFonts=true}if(m){this.fontInUse=m;this.fontDir+="/"+m;this.webfontDir+="/"+m;this.require.push(this.fontDir+"/fontdata.js");if(this.imgFonts){this.require.push(this.directory+"/imageFonts.js");b.Startup.signal.Post("HTML-CSS Jax - using image fonts")}}else{e(["CantFindFontUsing","Can't find a valid font using %1","["+this.config.availableFonts.join(", ")+"]"],null,3000);b.Startup.signal.Post("HTML-CSS Jax - no valid font")}this.require.push(MathJax.OutputJax.extensionDir+"/MathEvents.js")},Startup:function(){l=MathJax.Extension.MathEvents.Event;a=MathJax.Extension.MathEvents.Touch;c=MathJax.Extension.MathEvents.Hover;this.ContextMenu=l.ContextMenu;this.Mousedown=l.AltContextMenu;this.Mouseover=c.Mouseover;this.Mouseout=c.Mouseout;this.Mousemove=c.Mousemove;this.hiddenDiv=this.Element("div",{style:{visibility:"hidden",overflow:"hidden",position:"absolute",top:0,height:"1px",width:"auto",padding:0,border:0,margin:0,textAlign:"left",textIndent:0,textTransform:"none",lineHeight:"normal",letterSpacing:"normal",wordSpacing:"normal"}});if(!document.body.firstChild){document.body.appendChild(this.hiddenDiv)}else{document.body.insertBefore(this.hiddenDiv,document.body.firstChild)}this.hiddenDiv=this.addElement(this.hiddenDiv,"div",{id:"MathJax_Hidden"});var n=this.addElement(this.hiddenDiv,"div",{style:{width:"5in"}});this.pxPerInch=n.offsetWidth/5;this.hiddenDiv.removeChild(n);this.startMarker=this.createStrut(this.Element("span"),10,true);this.endMarker=this.addText(this.Element("span"),"x").parentNode;this.HDspan=this.Element("span");if(this.operaHeightBug){this.createStrut(this.HDspan,0)}if(this.msieInlineBlockAlignBug){this.HDimg=this.addElement(this.HDspan,"img",{style:{height:"0px",width:"1px"}});try{this.HDimg.src="about:blank"}catch(m){}}else{this.HDimg=this.createStrut(this.HDspan,0)}this.EmExSpan=this.Element("span",{style:{position:"absolute","font-size-adjust":"none"}},[["span",{className:"MathJax_ExBox"}],["span",{className:"MathJax"},[["span",{className:"MathJax_EmBox"}]]]]);this.linebreakSpan=MathJax.HTML.Element("span",{className:"MathJax_LineBox"},[["span"]]);return j.Styles(this.config.styles,["InitializeHTML",this])},removeSTIXfonts:function(p){for(var o=0,n=p.length;o<n;o++){if(p[o]==="STIX"){p.splice(o,1);n--;o--}}if(this.config.preferredFont==="STIX"){this.config.preferredFont=p[0]}},PreloadWebFonts:function(){if(!d.allowWebFonts||!d.config.preloadWebFonts){return}for(var o=0,n=d.config.preloadWebFonts.length;o<n;o++){var p=d.FONTDATA.FONTS[d.config.preloadWebFonts[o]];if(!p.available){d.Font.testFont(p)}}},InitializeHTML:function(){this.PreloadWebFonts();this.getDefaultExEm();if(this.defaultEm){return}var m=MathJax.Callback();j.timer.start(j,function(n){if(n.time(m)){b.signal.Post(["HTML-CSS Jax - no default em size"]);return}d.getDefaultExEm();if(d.defaultEm){m()}else{setTimeout(n,n.delay)}},this.defaultEmDelay,this.defaultEmTimeout);return m},defaultEmDelay:100,defaultEmTimeout:1000,getDefaultExEm:function(){document.body.appendChild(this.EmExSpan);document.body.appendChild(this.linebreakSpan);this.defaultEx=this.EmExSpan.firstChild.offsetHeight/60;this.defaultEm=this.EmExSpan.lastChild.firstChild.offsetHeight/60;this.defaultWidth=this.linebreakSpan.firstChild.offsetWidth;document.body.removeChild(this.linebreakSpan);document.body.removeChild(this.EmExSpan)},preTranslate:function(s){var r=s.jax[this.id],D,z=r.length,y,G,w,C,u,E,p,F,o,H,v,t=false,A,q=this.config.linebreaks.automatic,x=this.config.linebreaks.width;if(q){t=(x.match(/^\s*(\d+(\.\d*)?%\s*)?container\s*$/)!=null);if(t){x=x.replace(/\s*container\s*/,"")}else{v=this.defaultWidth}if(x===""){x="100%"}}else{v=100000}for(D=0;D<z;D++){G=r[D];if(!G.parentNode){continue}w=G.previousSibling;if(w&&String(w.className).match(/^MathJax(_Display)?( MathJax_Process(ing|ed))?$/)){w.parentNode.removeChild(w)}if(G.MathJax.preview){G.MathJax.preview.style.display="none"}p=G.MathJax.elementJax;if(!p){continue}p.HTMLCSS={display:(p.root.Get("display")==="block")};C=u=this.Element("span",{className:"MathJax",id:p.inputID+"-Frame",isMathJax:true,jaxID:this.id,oncontextmenu:l.Menu,onmousedown:l.Mousedown,onmouseover:l.Mouseover,onmouseout:l.Mouseout,onmousemove:l.Mousemove,onclick:l.Click,ondblclick:l.DblClick,onkeydown:l.Keydown,tabIndex:b.getTabOrder(p)});if(b.Browser.noContextMenu){C.ontouchstart=a.start;C.ontouchend=a.end}if(p.HTMLCSS.display){u=this.Element("div",{className:"MathJax_Display"});u.appendChild(C)}else{if(this.msieDisappearingBug){C.style.display="inline-block"}}u.className+=" MathJax_Processing";G.parentNode.insertBefore(u,G);p.HTMLCSS.span=C;p.HTMLCSS.div=u;G.parentNode.insertBefore(this.EmExSpan.cloneNode(true),G);u.parentNode.insertBefore(this.linebreakSpan.cloneNode(true),u)}var B=[];for(D=0;D<z;D++){G=r[D];if(!G.parentNode){continue}E=G.previousSibling;u=E.previousSibling;p=G.MathJax.elementJax;if(!p){continue}F=E.firstChild.offsetHeight/60;o=E.lastChild.firstChild.offsetHeight/60;A=Math.max(0,u.previousSibling.firstChild.offsetWidth-2);if(F===0||F==="NaN"){B.push(u);p.HTMLCSS.isHidden=true;F=this.defaultEx;o=this.defaultEm;A=this.defaultWidth}if(t){v=A}H=(this.config.matchFontHeight?F/this.TeX.x_height/o:1);H=Math.floor(Math.max(this.config.minScaleAdjust/100,H)*this.config.scale);p.HTMLCSS.scale=H/100;p.HTMLCSS.fontSize=H+"%";p.HTMLCSS.em=p.HTMLCSS.outerEm=o;this.em=o*H/100;p.HTMLCSS.ex=F;p.HTMLCSS.cwidth=A/this.em;p.HTMLCSS.lineWidth=(q?this.length2em(x,1,v/this.em):1000000)}for(D=0,y=B.length;D<y;D++){this.hiddenDiv.appendChild(B[D]);this.addElement(this.hiddenDiv,"br")}for(D=0;D<z;D++){G=r[D];if(!G.parentNode){continue}E=r[D].previousSibling;p=r[D].MathJax.elementJax;if(!p){continue}C=E.previousSibling;if(!p.HTMLCSS.isHidden){C=C.previousSibling}C.parentNode.removeChild(C);E.parentNode.removeChild(E);if(G.MathJax.preview){G.MathJax.preview.style.display=""}}s.HTMLCSSeqn=s.HTMLCSSlast=0;s.HTMLCSSi=-1;s.HTMLCSSchunk=this.config.EqnChunk;s.HTMLCSSdelay=false},PHASE:{I:1,II:2,III:3},Translate:function(n,r){if(!n.parentNode){return}if(r.HTMLCSSdelay){r.HTMLCSSdelay=false;b.RestartAfter(MathJax.Callback.Delay(this.config.EqnChunkDelay))}var m=n.MathJax.elementJax,q=m.root,s=m.HTMLCSS.div,o=m.HTMLCSS.span;if(!document.getElementById(o.id)){return}this.getMetrics(m);if(this.scale!==1){o.style.fontSize=m.HTMLCSS.fontSize}this.initImg(o);this.initHTML(q,o);this.savePreview(n);try{q.setTeXclass();q.toHTML(o,s,this.PHASE.I)}catch(p){if(p.restart){while(o.firstChild){o.removeChild(o.firstChild)}}this.restorePreview(n);throw p}this.restorePreview(n);s.className=s.className.split(/ /)[0]+" MathJax_Processed";b.signal.Post(["New Math Pending",m.inputID]);r.HTMLCSSeqn+=(r.i-r.HTMLCSSi);r.HTMLCSSi=r.i;if(r.HTMLCSSeqn>=r.HTMLCSSlast+r.HTMLCSSchunk){this.postTranslate(r,true);r.HTMLCSSchunk=Math.floor(r.HTMLCSSchunk*this.config.EqnChunkFactor);r.HTMLCSSdelay=true}return false},savePreview:function(m){var n=m.MathJax.preview;if(n){m.MathJax.tmpPreview=document.createElement("span");n.parentNode.replaceChild(m.MathJax.tmpPreview,n)}},restorePreview:function(m){var n=m.MathJax.tmpPreview;if(n){n.parentNode.replaceChild(m.MathJax.preview,n);delete m.MathJax.tmpPreview}},getMetrics:function(m){var n=m.HTMLCSS;this.em=i.mbase.prototype.em=n.em*n.scale;this.outerEm=n.em;this.scale=n.scale;this.cwidth=n.cwidth;this.linebreakWidth=n.lineWidth},postTranslate:function(o,u){var r=o.jax[this.id],v,p,s,q;for(s=o.HTMLCSSlast,q=o.HTMLCSSeqn;s<q;s++){v=r[s];if(v&&v.parentNode&&v.MathJax.elementJax){var n=(v.MathJax.elementJax.HTMLCSS||{}).div;if(n){n.className=n.className.split(/ /)[0]}if(v.MathJax.preview){v.MathJax.preview.innerHTML="";v.MathJax.preview.style.display="none"}}}for(s=o.HTMLCSSlast,q=o.HTMLCSSeqn;s<q;s++){v=r[s];if(v&&v.parentNode&&v.MathJax.elementJax){p=v.MathJax.elementJax;this.getMetrics(p);if(p.HTMLCSS.span&&p.HTMLCSS.div){p.root.toHTML(p.HTMLCSS.span,p.HTMLCSS.div,this.PHASE.II)}}}for(s=o.HTMLCSSlast,q=o.HTMLCSSeqn;s<q;s++){v=r[s];if(v&&v.parentNode&&v.MathJax.elementJax){p=v.MathJax.elementJax;this.getMetrics(p);if(p.HTMLCSS.span&&p.HTMLCSS.div){p.root.toHTML(p.HTMLCSS.span,p.HTMLCSS.div,this.PHASE.III);if(p.HTMLCSS.isHidden){v.parentNode.insertBefore(p.HTMLCSS.div,v)}delete p.HTMLCSS.span;delete p.HTMLCSS.div;v.MathJax.state=p.STATE.PROCESSED;b.signal.Post(["New Math",v.MathJax.elementJax.inputID])}}}if(this.forceReflow){var t=(document.styleSheets||[])[0]||{};t.disabled=true;t.disabled=false}o.HTMLCSSlast=o.HTMLCSSeqn},getJaxFromMath:function(m){if(m.parentNode.className.match(/MathJax_Display/)){m=m.parentNode}do{m=m.nextSibling}while(m&&m.nodeName.toLowerCase()!=="script");return b.getJaxFor(m)},getHoverSpan:function(m,n){return m.root.HTMLspanElement()},getHoverBBox:function(m,p,q){var r=p.bbox,o=m.HTMLCSS.outerEm;var n={w:r.w*o,h:r.h*o,d:r.d*o};if(r.width){n.width=r.width}return n},Zoom:function(n,y,x,m,v){y.className="MathJax";y.style.fontSize=n.HTMLCSS.fontSize;var B=y.appendChild(this.EmExSpan.cloneNode(true));var q=B.lastChild.firstChild.offsetHeight/60;this.em=i.mbase.prototype.em=q;this.outerEm=q/n.HTMLCSS.scale;B.parentNode.removeChild(B);this.scale=n.HTMLCSS.scale;this.linebreakWidth=n.HTMLCSS.lineWidth;this.cwidth=n.HTMLCSS.cwidth;this.zoomScale=parseInt(b.config.menuSettings.zscale)/100;this.idPostfix="-zoom";n.root.toHTML(y,y);this.idPostfix="";this.zoomScale=1;var z=n.root.HTMLspanElement().bbox,p=z.width;if(p){if(z.tw){m=z.tw*q}if(z.w*q<m){m=z.w*q}y.style.width=Math.floor(m-1.5*d.em)+"px";y.style.display="inline-block";var o=(n.root.id||"MathJax-Span-"+n.root.spanID)+"-zoom";var r=document.getElementById(o).firstChild;while(r&&r.style.width!==p){r=r.nextSibling}if(r){var u=r.offsetWidth;r.style.width="100%";if(u>m){y.style.width=(u+100)+"px"}}}r=y.firstChild.firstChild.style;if(z.H!=null&&z.H>z.h){r.marginTop=d.Em(z.H-Math.max(z.h,d.FONTDATA.lineH))}if(z.D!=null&&z.D>z.d){r.marginBottom=d.Em(z.D-Math.max(z.d,d.FONTDATA.lineD))}if(z.lw<0){r.paddingLeft=d.Em(-z.lw)}if(z.rw>z.w){r.marginRight=d.Em(z.rw-z.w)}y.style.position="absolute";if(!p){x.style.position="absolute"}var w=y.offsetWidth,t=y.offsetHeight,A=x.offsetHeight,s=x.offsetWidth;y.style.position=x.style.position="";return{Y:-l.getBBox(y).h,mW:s,mH:A,zW:w,zH:t}},initImg:function(m){},initHTML:function(n,m){},initFont:function(m){var o=d.FONTDATA.FONTS,n=d.config.availableFonts;if(n&&n.length&&d.Font.testFont(o[m])){o[m].available=true;if(o[m].familyFixed){o[m].family=o[m].familyFixed;delete o[m].familyFixed}return null}if(!this.allowWebFonts){return null}o[m].isWebFont=true;if(d.FontFaceBug){o[m].family=m;if(d.msieFontCSSBug){o[m].family+="-Web"}}return j.Styles({"@font-face":this.Font.fontFace(m)})},Remove:function(m){var n=document.getElementById(m.inputID+"-Frame");if(n){if(m.HTMLCSS.display){n=n.parentNode}n.parentNode.removeChild(n)}delete m.HTMLCSS},getHD:function(n,o){if(n.bbox&&this.config.noReflows&&!o){return{h:n.bbox.h,d:n.bbox.d}}var m=n.style.position;n.style.position="absolute";this.HDimg.style.height="0px";n.appendChild(this.HDspan);var p={h:n.offsetHeight};this.HDimg.style.height=p.h+"px";p.d=n.offsetHeight-p.h;p.h-=p.d;p.h/=this.em;p.d/=this.em;n.removeChild(this.HDspan);n.style.position=m;return p},getW:function(q){var n,p,o=(q.bbox||{}).w,r=q;if(q.bbox&&this.config.noReflows&&q.bbox.exactW!==false){if(!q.bbox.exactW){if(q.style.paddingLeft){o+=this.unEm(q.style.paddingLeft)*(q.scale||1)}if(q.style.paddingRight){o+=this.unEm(q.style.paddingRight)*(q.scale||1)}}return o}if(q.bbox&&q.bbox.exactW){return o}if((q.bbox&&o>=0&&!this.initialSkipBug&&!this.msieItalicWidthBug)||this.negativeBBoxes||!q.firstChild){n=q.offsetWidth;p=q.parentNode.offsetHeight}else{if(q.bbox&&o<0&&this.msieNegativeBBoxBug){n=-q.offsetWidth,p=q.parentNode.offsetHeight}else{var m=q.style.position;q.style.position="absolute";r=this.startMarker;q.insertBefore(r,q.firstChild);q.appendChild(this.endMarker);n=this.endMarker.offsetLeft-r.offsetLeft;q.removeChild(this.endMarker);q.removeChild(r);q.style.position=m}}if(p!=null){q.parentNode.HH=p/this.em}return n/this.em},Measured:function(o,n){var p=o.bbox;if(p.width==null&&p.w&&!p.isMultiline){var m=this.getW(o);p.rw+=m-p.w;p.w=m;p.exactW=true}if(!n){n=o.parentNode}if(!n.bbox){n.bbox=p}return o},Remeasured:function(n,m){m.bbox=this.Measured(n,m).bbox},MeasureSpans:function(q){var t=[],v,s,p,w,n,r,o,u;for(s=0,p=q.length;s<p;s++){v=q[s];if(!v){continue}w=v.bbox;u=this.parentNode(v);if(w.exactW||w.width||w.w===0||w.isMultiline||(this.config.noReflows&&w.exactW!==false)){if(!u.bbox){u.bbox=w}continue}if(this.negativeBBoxes||!v.firstChild||(w.w>=0&&!this.initialSkipBug)||(w.w<0&&this.msieNegativeBBoxBug)){t.push([v])}else{if(this.initialSkipBug){n=this.startMarker.cloneNode(true);r=this.endMarker.cloneNode(true);v.insertBefore(n,v.firstChild);v.appendChild(r);t.push([v,n,r,v.style.position]);v.style.position="absolute"}else{r=this.endMarker.cloneNode(true);v.appendChild(r);t.push([v,null,r])}}}for(s=0,p=t.length;s<p;s++){v=t[s][0];w=v.bbox;u=this.parentNode(v);if((w.w>=0&&!this.initialSkipBug)||this.negativeBBoxes||!v.firstChild){o=v.offsetWidth;u.HH=u.offsetHeight/this.em}else{if(w.w<0&&this.msieNegativeBBoxBug){o=-v.offsetWidth,u.HH=u.offsetHeight/this.em}else{o=t[s][2].offsetLeft-((t[s][1]||{}).offsetLeft||0)}}o/=this.em;w.rw+=o-w.w;w.w=o;w.exactW=true;if(!u.bbox){u.bbox=w}}for(s=0,p=t.length;s<p;s++){v=t[s];if(v[1]){v[1].parentNode.removeChild(v[1]),v[0].style.position=v[3]}if(v[2]){v[2].parentNode.removeChild(v[2])}}},Em:function(n){if(Math.abs(n)<0.0006){return"0em"}return n.toFixed(3).replace(/\.?0+$/,"")+"em"},EmRounded:function(n){if(Math.abs(n)<0.0006){return"0em"}n=(Math.round(n*d.em)+0.05)/d.em;return n.toFixed(3).replace(/\.?0+$/,"")+"em"},unEm:function(n){return parseFloat(n)},Px:function(n){n*=this.em;var o=(n<0?"-":"");return o+Math.abs(n).toFixed(1).replace(/\.?0+$/,"")+"px"},unPx:function(n){return parseFloat(n)/this.em},Percent:function(n){return(100*n).toFixed(1).replace(/\.?0+$/,"")+"%"},length2em:function(t,o,r){if(typeof(t)!=="string"){t=t.toString()}if(t===""){return""}if(t===i.SIZE.NORMAL){return 1}if(t===i.SIZE.BIG){return 2}if(t===i.SIZE.SMALL){return 0.71}if(t==="infinity"){return d.BIGDIMEN}var q=this.FONTDATA.TeX_factor,u=(d.zoomScale||1)/d.em;if(t.match(/mathspace$/)){return d.MATHSPACE[t]*q}var p=t.match(/^\s*([-+]?(?:\.\d+|\d+(?:\.\d*)?))?(pt|em|ex|mu|px|pc|in|mm|cm|%)?/);var n=parseFloat(p[1]||"1"),s=p[2];if(r==null){r=1}if(o==null){o=1}if(s==="em"){return n*q}if(s==="ex"){return n*d.TeX.x_height*q}if(s==="%"){return n/100*r}if(s==="px"){return n*u}if(s==="pt"){return n/10*q}if(s==="pc"){return n*1.2*q}if(s==="in"){return n*this.pxPerInch*u}if(s==="cm"){return n*this.pxPerInch*u/2.54}if(s==="mm"){return n*this.pxPerInch*u/25.4}if(s==="mu"){return n/18*q*o}return n*r},thickness2em:function(n,m){var o=d.TeX.rule_thickness;if(n===i.LINETHICKNESS.MEDIUM){return o}if(n===i.LINETHICKNESS.THIN){return 0.67*o}if(n===i.LINETHICKNESS.THICK){return 1.67*o}return this.length2em(n,m,o)},getPadding:function(n){var p={top:0,right:0,bottom:0,left:0},m=false;for(var q in p){if(p.hasOwnProperty(q)){var o=n.style["padding"+q.charAt(0).toUpperCase()+q.substr(1)];if(o){p[q]=this.length2em(o);m=true}}}return(m?p:false)},getBorders:function(r){var o={top:0,right:0,bottom:0,left:0},p={},n=false;for(var s in o){if(o.hasOwnProperty(s)){var m="border"+s.charAt(0).toUpperCase()+s.substr(1);var q=r.style[m+"Style"];if(q){n=true;o[s]=this.length2em(r.style[m+"Width"]);p[m]=[r.style[m+"Width"],r.style[m+"Style"],r.style[m+"Color"]].join(" ")}}}o.css=p;return(n?o:false)},setBorders:function(m,n){if(n){for(var o in n.css){if(n.css.hasOwnProperty(o)){m.style[o]=n.css[o]}}}},createStrut:function(o,n,p){var m=this.Element("span",{isMathJax:true,style:{display:"inline-block",overflow:"hidden",height:n+"px",width:"1px",marginRight:"-1px"}});if(p){o.insertBefore(m,o.firstChild)}else{o.appendChild(m)}return m},createBlank:function(n,m,o){var p=this.Element("span",{isMathJax:true,style:{display:"inline-block",overflow:"hidden",height:"1px",width:this.Em(m)}});if(m<0){p.style.marginRight=p.style.width;p.style.width=0}if(o){n.insertBefore(p,n.firstChild)}else{n.appendChild(p)}return p},createShift:function(n,m,p){var o=this.Element("span",{style:{marginLeft:this.Em(m)},isMathJax:true});if(p){n.insertBefore(o,n.firstChild)}else{n.appendChild(o)}return o},createSpace:function(r,p,q,s,o,u){if(p<-q){q=-p}var t=this.Em(p+q),m=this.Em(-q);if(this.msieInlineBlockAlignBug){m=this.Em(d.getHD(r.parentNode,true).d-q)}if(r.isBox||u){var n=(r.scale==null?1:r.scale);r.bbox={exactW:true,h:p*n,d:q*n,w:s*n,rw:s*n,lw:0};r.style.height=t;r.style.verticalAlign=m;r.HH=(p+q)*n}else{r=this.addElement(r,"span",{style:{height:t,verticalAlign:m},isMathJax:true})}if(s>=0){r.style.width=this.Em(s);r.style.display="inline-block";r.style.overflow="hidden"}else{if(this.msieNegativeSpaceBug){r.style.height=""}r.style.marginLeft=this.Em(s);if(d.safariNegativeSpaceBug&&r.parentNode.firstChild==r){this.createBlank(r,0,true)}}if(o&&o!==i.COLOR.TRANSPARENT){r.style.backgroundColor=o;r.style.position="relative"}return r},createRule:function(t,p,r,u,n){if(p<-r){r=-p}var o=d.TeX.min_rule_thickness,q=1;if(u>0&&u*this.em<o){u=o/this.em}if(p+r>0&&(p+r)*this.em<o){q=1/(p+r)*(o/this.em);p*=q;r*=q}if(!n){n="solid"}else{n="solid "+n}var m={display:"inline-block",overflow:"hidden",verticalAlign:this.Em(-r)};if(u>p+r){m.borderTop=this.Px(p+r)+" "+n;m.width=this.Em(u);m.height=(this.msieRuleBug&&p+r>0?this.Em(p+r):0)}else{m.borderLeft=this.Px(u)+" "+n;m.width=(this.msieRuleBug&&u>0?this.Em(u):0);m.height=this.Em(p+r)}var s=this.addElement(t,"span",{style:m,noAdjust:true,HH:p+r,isMathJax:true,bbox:{h:p,d:r,w:u,rw:u,lw:0,exactW:true}});if(t.isBox||t.className=="mspace"){t.bbox=s.bbox,t.HH=p+r}return s},createFrame:function(v,s,u,x,z,n){if(s<-u){u=-s}var r=2*z;if(this.msieFrameSizeBug){if(x<r){x=r}if(s+u<r){s=r-u}}if(this.msieBorderWidthBug){r=0}var y=this.Em(s+u-r),m=this.Em(-u-z),q=this.Em(x-r);var o=this.Px(z)+" "+n;var p=this.addElement(v,"span",{style:{border:o,display:"inline-block",overflow:"hidden",width:q,height:y},bbox:{h:s,d:u,w:x,rw:x,lw:0,exactW:true},noAdjust:true,HH:s+u,isMathJax:true});if(m){p.style.verticalAlign=m}return p},parentNode:function(n){var m=n.parentNode;if(m.nodeName.toLowerCase()==="a"){m=m.parentNode}return m},createStack:function(o,q,n){if(this.msiePaddingWidthBug){this.createStrut(o,0)}var p=String(n).match(/%$/);var m=(!p&&n!=null?n:0);o=this.addElement(o,"span",{noAdjust:true,HH:0,isMathJax:true,style:{display:"inline-block",position:"relative",width:(p?"100%":this.Em(m)),height:0}});if(!q){o.parentNode.bbox=o.bbox={exactW:true,h:-this.BIGDIMEN,d:-this.BIGDIMEN,w:m,lw:this.BIGDIMEN,rw:(!p&&n!=null?n:-this.BIGDIMEN)};if(p){o.bbox.width=n}}return o},createBox:function(n,m){var o=this.addElement(n,"span",{style:{position:"absolute"},isBox:true,isMathJax:true});if(m!=null){o.style.width=m}return o},addBox:function(m,n){n.style.position="absolute";n.isBox=n.isMathJax=true;return m.appendChild(n)},placeBox:function(v,u,s,p){v.isMathJax=true;var w=d.parentNode(v),E=v.bbox,A=w.bbox;if(this.msiePlaceBoxBug){this.addText(v,this.NBSP)}if(this.imgSpaceBug){this.addText(v,this.imgSpace)}var z,G=0;if(v.HH!=null){z=v.HH}else{if(E){z=Math.max(3,E.h+E.d)}else{z=v.offsetHeight/this.em}}if(!v.noAdjust){z+=1;z=Math.round(z*this.em)/this.em;if(this.msieInlineBlockAlignBug){this.addElement(v,"img",{className:"MathJax_strut",border:0,src:"about:blank",isMathJax:true,style:{width:0,height:this.Em(z)}})}else{this.addElement(v,"span",{isMathJax:true,style:{display:"inline-block",width:0,height:this.Em(z)}});if(d.chromeHeightBug){z-=(v.lastChild.offsetHeight-Math.round(z*this.em))/this.em}}}if(E){if(this.initialSkipBug){if(E.lw<0){G=E.lw;d.createBlank(v,-G,true)}if(E.rw>E.w){d.createBlank(v,E.rw-E.w+0.1)}}if(!this.msieClipRectBug&&!E.noclip&&!p){var C=3/this.em;var B=(E.H==null?E.h:E.H),n=(E.D==null?E.d:E.D);var F=z-B-C,q=z+n+C,o=-1000,m=E.rw+1000;v.style.clip="rect("+this.Em(F)+" "+this.Em(m)+" "+this.Em(q)+" "+this.Em(o)+")"}}v.style.top=this.Em(-s-z);v.style.left=this.Em(u+G);if(E&&A){if(E.H!=null&&(A.H==null||E.H+s>A.H)){A.H=E.H+s}if(E.D!=null&&(A.D==null||E.D-s>A.D)){A.D=E.D-s}if(E.h+s>A.h){A.h=E.h+s}if(E.d-s>A.d){A.d=E.d-s}if(A.H!=null&&A.H<=A.h){delete A.H}if(A.D!=null&&A.D<=A.d){delete A.D}if(E.w+u>A.w){A.w=E.w+u;if(A.width==null){w.style.width=this.Em(A.w)}}if(E.rw+u>A.rw){A.rw=E.rw+u}if(E.lw+u<A.lw){A.lw=E.lw+u}if(E.width!=null&&!E.isFixed){if(A.width==null){w.style.width=A.width="100%";if(E.minWidth){w.style.minWidth=A.minWidth=E.minWidth}}v.style.width=E.width}if(E.tw){A.tw=E.tw}}},alignBox:function(t,p,s,x,w){if(x==null){x=0}this.placeBox(t,x,s);if(this.msiePlaceBoxBug){var n=t.lastChild;while(n&&n.nodeName!=="#text"){n=n.previousSibling}if(n){t.removeChild(n)}}var v=t.bbox;if(v.isMultiline&&!w){return}var u=v.width!=null&&!v.isFixed;var m=0,q=x-v.w/2,o="50%";if(this.initialSkipBug){m=v.w-v.rw-0.1;q+=v.lw}if(this.msieMarginScaleBug){q=(q*this.em)+"px"}else{q=this.Em(q)}if(u){q=(x===0?"":this.Em(x));o=(50-parseFloat(v.width)/2)+"%"}b.Insert(t.style,({right:{left:"",right:this.Em(m-x)},center:{left:o,marginLeft:q}})[p])},setStackWidth:function(n,m){if(typeof(m)==="number"){n.style.width=this.Em(Math.max(0,m));var o=n.bbox;if(o){o.w=m;o.exactW=true}o=n.parentNode.bbox;if(o){o.w=m;o.exactW=true}}else{n.style.width=n.parentNode.style.width="100%";if(n.bbox){n.bbox.width=m}if(n.parentNode.bbox){n.parentNode.bbox.width=m}}},createDelimiter:function(w,n,p,s,q){if(!n){w.bbox={h:0,d:0,w:this.TeX.nulldelimiterspace,lw:0};w.bbox.rw=w.bbox.w;this.createSpace(w,w.bbox.h,w.bbox.d,w.bbox.w);return}if(!s){s=1}if(!(p instanceof Array)){p=[p,p]}var v=p[1];p=p[0];var o={alias:n};while(o.alias){n=o.alias;o=this.FONTDATA.DELIMITERS[n];if(!o){o={HW:[0,this.FONTDATA.VARIANT[i.VARIANT.NORMAL]]}}}if(o.load){b.RestartAfter(j.Require(this.fontDir+"/fontdata-"+o.load+".js"))}for(var u=0,r=o.HW.length;u<r;u++){if(o.HW[u][0]*s>=p-0.01||(u==r-1&&!o.stretch)){if(o.HW[u][2]){s*=o.HW[u][2]}if(o.HW[u][3]){n=o.HW[u][3]}var t=this.addElement(w,"span");this.createChar(t,[n,o.HW[u][1]],s,q);w.bbox=t.bbox;w.offset=0.65*w.bbox.w;w.scale=s;return}}if(o.stretch){this["extendDelimiter"+o.dir](w,v,o.stretch,s,q)}},extendDelimiterV:function(B,u,F,G,x){var p=this.createStack(B,true);var w=this.createBox(p),v=this.createBox(p);this.createChar(w,(F.top||F.ext),G,x);this.createChar(v,(F.bot||F.ext),G,x);var o={bbox:{w:0,lw:0,rw:0}},E=o,q;var C=w.bbox.h+w.bbox.d+v.bbox.h+v.bbox.d;var s=-w.bbox.h;this.placeBox(w,0,s,true);s-=w.bbox.d;if(F.mid){E=this.createBox(p);this.createChar(E,F.mid,G,x);C+=E.bbox.h+E.bbox.d}if(F.min&&u<C*F.min){u=C*F.min}if(u>C){o=this.Element("span");this.createChar(o,F.ext,G,x);var D=o.bbox.h+o.bbox.d,m=D-0.05,z,r,A=(F.mid?2:1);r=z=Math.min(Math.ceil((u-C)/(A*m)),this.maxStretchyParts);if(!F.fullExtenders){m=(u-C)/(A*z)}var t=(z/(z+1))*(D-m);m=D-t;s+=t+m-o.bbox.h;while(A-->0){while(z-->0){if(!this.msieCloneNodeBug){q=o.cloneNode(true)}else{q=this.Element("span");this.createChar(q,F.ext,G,x)}q.bbox=o.bbox;s-=m;this.placeBox(this.addBox(p,q),0,s,true)}s+=t-o.bbox.d;if(F.mid&&A){this.placeBox(E,0,s-E.bbox.h,true);z=r;s+=-(E.bbox.h+E.bbox.d)+t+m-o.bbox.h}}}else{s+=(C-u)/2;if(F.mid){this.placeBox(E,0,s-E.bbox.h,true);s+=-(E.bbox.h+E.bbox.d)}s+=(C-u)/2}this.placeBox(v,0,s-v.bbox.h,true);s-=v.bbox.h+v.bbox.d;B.bbox={w:Math.max(w.bbox.w,o.bbox.w,v.bbox.w,E.bbox.w),lw:Math.min(w.bbox.lw,o.bbox.lw,v.bbox.lw,E.bbox.lw),rw:Math.max(w.bbox.rw,o.bbox.rw,v.bbox.rw,E.bbox.rw),h:0,d:-s,exactW:true};B.scale=G;B.offset=0.55*B.bbox.w;B.isMultiChar=true;this.setStackWidth(p,B.bbox.w)},extendDelimiterH:function(C,p,F,H,z){var s=this.createStack(C,true);var q=this.createBox(s),D=this.createBox(s);this.createChar(q,(F.left||F.rep),H,z);this.createChar(D,(F.right||F.rep),H,z);var m=this.Element("span");this.createChar(m,F.rep,H,z);var E={bbox:{h:-this.BIGDIMEN,d:-this.BIGDIMEN}},o;this.placeBox(q,-q.bbox.lw,0,true);var v=(q.bbox.rw-q.bbox.lw)+(D.bbox.rw-D.bbox.lw)-0.05,u=q.bbox.rw-q.bbox.lw-0.025,y;if(F.mid){E=this.createBox(s);this.createChar(E,F.mid,H,z);v+=E.bbox.w}if(F.min&&p<v*F.min){p=v*F.min}if(p>v){var G=m.bbox.rw-m.bbox.lw,r=G-0.05,A,t,B=(F.mid?2:1);t=A=Math.min(Math.ceil((p-v)/(B*r)),this.maxStretchyParts);if(!F.fillExtenders){r=(p-v)/(B*A)}y=(A/(A+1))*(G-r);r=G-y;u-=m.bbox.lw+y;while(B-->0){while(A-->0){if(!this.cloneNodeBug){o=m.cloneNode(true)}else{o=this.Element("span");this.createChar(o,F.rep,H,z)}o.bbox=m.bbox;this.placeBox(this.addBox(s,o),u,0,true);u+=r}if(F.mid&&B){this.placeBox(E,u,0,true);u+=E.bbox.w-y;A=t}}}else{u-=(v-p)/2;if(F.mid){this.placeBox(E,u,0,true);u+=E.bbox.w}u-=(v-p)/2}this.placeBox(D,u,0,true);C.bbox={w:u+D.bbox.rw,lw:0,rw:u+D.bbox.rw,h:Math.max(q.bbox.h,m.bbox.h,D.bbox.h,E.bbox.h),d:Math.max(q.bbox.d,m.bbox.d,D.bbox.d,E.bbox.d),exactW:true};C.scale=H;C.isMultiChar=true;this.setStackWidth(s,C.bbox.w)},createChar:function(u,r,p,n){u.isMathJax=true;var t=u,v="",q={fonts:[r[1]],noRemap:true};if(n&&n===i.VARIANT.BOLD){q.fonts=[r[1]+"-bold",r[1]]}if(typeof(r[1])!=="string"){q=r[1]}if(r[0] instanceof Array){for(var s=0,o=r[0].length;s<o;s++){v+=String.fromCharCode(r[0][s])}}else{v=String.fromCharCode(r[0])}if(r[4]){p*=r[4]}if(p!==1||r[3]){t=this.addElement(u,"span",{style:{fontSize:this.Percent(p)},scale:p,isMathJax:true});this.handleVariant(t,q,v);u.bbox=t.bbox}else{this.handleVariant(u,q,v)}if(r[2]){u.style.marginLeft=this.Em(r[2])}if(r[3]){u.firstChild.style.verticalAlign=this.Em(r[3]);u.bbox.h+=r[3];if(u.bbox.h<0){u.bbox.h=0}}if(r[5]){u.bbox.h+=r[5]}if(r[6]){u.bbox.d+=r[6]}if(this.AccentBug&&u.bbox.w===0){t.firstChild.nodeValue+=this.NBSP}},positionDelimiter:function(n,m){m-=n.bbox.h;n.bbox.d-=m;n.bbox.h+=m;if(m){if(this.safariVerticalAlignBug||this.konquerorVerticalAlignBug||(this.operaVerticalAlignBug&&n.isMultiChar)){if(n.firstChild.style.display===""&&n.style.top!==""){n=n.firstChild;m-=d.unEm(n.style.top)}n.style.position="relative";n.style.top=this.Em(-m)}else{n.style.verticalAlign=this.Em(m);if(d.ffVerticalAlignBug){d.createRule(n.parentNode,n.bbox.h,0,0);delete n.parentNode.bbox}}}},handleVariant:function(B,q,t){var A="",y,D,u,E,o=B,p=!!B.style.fontFamily;if(t.length===0){return}if(!B.bbox){B.bbox={w:0,h:-this.BIGDIMEN,d:-this.BIGDIMEN,rw:-this.BIGDIMEN,lw:this.BIGDIMEN}}if(!q){q=this.FONTDATA.VARIANT[i.VARIANT.NORMAL]}E=q;for(var C=0,z=t.length;C<z;C++){q=E;y=t.charCodeAt(C);D=t.charAt(C);if(y>=55296&&y<56319){C++;y=(((y-55296)<<10)+(t.charCodeAt(C)-56320))+65536;if(this.FONTDATA.RemapPlane1){var F=this.FONTDATA.RemapPlane1(y,q);y=F.n;q=F.variant}}else{var v,s,w=this.FONTDATA.RANGES;for(v=0,s=w.length;v<s;v++){if(w[v].name==="alpha"&&q.noLowerCase){continue}var r=q["offset"+w[v].offset];if(r&&y>=w[v].low&&y<=w[v].high){if(w[v].remap&&w[v].remap[y]){y=r+w[v].remap[y]}else{y=y-w[v].low+r;if(w[v].add){y+=w[v].add}}if(q["variant"+w[v].offset]){q=this.FONTDATA.VARIANT[q["variant"+w[v].offset]]}break}}}if(q.remap&&q.remap[y]){y=q.remap[y];if(q.remap.variant){q=this.FONTDATA.VARIANT[q.remap.variant]}}else{if(this.FONTDATA.REMAP[y]&&!q.noRemap){y=this.FONTDATA.REMAP[y]}}if(h(y)){q=this.FONTDATA.VARIANT[y[1]];y=y[0]}if(typeof(y)==="string"){t=y+t.substr(C+1);z=t.length;C=-1;continue}u=this.lookupChar(q,y);D=u[y];if(p||(!this.checkFont(u,o.style)&&!D[5].img)){if(A.length){this.addText(o,A);A=""}var x=!!o.style.fontFamily||!!B.style.fontStyle||!!B.style.fontWeight||!u.directory||p;p=false;if(o!==B){x=!this.checkFont(u,B.style);o=B}if(x){o=this.addElement(B,"span",{isMathJax:true,subSpan:true})}this.handleFont(o,u,o!==B)}A=this.handleChar(o,u,D,y,A);if(!(D[5]||{}).space){if(D[0]/1000>B.bbox.h){B.bbox.h=D[0]/1000}if(D[1]/1000>B.bbox.d){B.bbox.d=D[1]/1000}}if(B.bbox.w+D[3]/1000<B.bbox.lw){B.bbox.lw=B.bbox.w+D[3]/1000}if(B.bbox.w+D[4]/1000>B.bbox.rw){B.bbox.rw=B.bbox.w+D[4]/1000}B.bbox.w+=D[2]/1000;if((D[5]||{}).isUnknown){B.bbox.exactW=false}}if(A.length){this.addText(o,A)}if(B.scale&&B.scale!==1){B.bbox.h*=B.scale;B.bbox.d*=B.scale;B.bbox.w*=B.scale;B.bbox.lw*=B.scale;B.bbox.rw*=B.scale}if(t.length==1&&u.skew&&u.skew[y]){B.bbox.skew=u.skew[y]}},checkFont:function(m,n){var o=(n.fontWeight||"normal");if(o.match(/^\d+$/)){o=(parseInt(o)>=600?"bold":"normal")}return(m.family.replace(/'/g,"")===n.fontFamily.replace(/'/g,"")&&(m.style||"normal")===(n.fontStyle||"normal")&&(m.weight||"normal")===o)},handleFont:function(o,m,q){o.style.fontFamily=m.family;if(!m.directory){o.style.fontSize=Math.floor(d.config.scale/d.scale+0.5)+"%"}if(!(d.FontFaceBug&&m.isWebFont)){var n=m.style||"normal",p=m.weight||"normal";if(n!=="normal"||q){o.style.fontStyle=n}if(p!=="normal"||q){o.style.fontWeight=p}}},handleChar:function(o,m,u,t,s){var r=u[5];if(r.space){if(s.length){this.addText(o,s)}d.createShift(o,u[2]/1000);return""}if(r.img){return this.handleImg(o,m,u,t,s)}if(r.isUnknown&&this.FONTDATA.DELIMITERS[t]){if(s.length){this.addText(o,s)}var q=o.scale;d.createDelimiter(o,t,0,1,m);if(this.FONTDATA.DELIMITERS[t].dir==="V"){o.style.verticalAlign=this.Em(o.bbox.d);o.bbox.h+=o.bbox.d;o.bbox.d=0}o.scale=q;u[0]=o.bbox.h*1000;u[1]=o.bbox.d*1000;u[2]=o.bbox.w*1000;u[3]=o.bbox.lw*1000;u[4]=o.bbox.rw*1000;return""}if(r.c==null){if(t<=65535){r.c=String.fromCharCode(t)}else{var p=t-65536;r.c=String.fromCharCode((p>>10)+55296)+String.fromCharCode((p&1023)+56320)}}if(d.ffFontOptimizationBug&&u[4]-u[2]>125){o.style.textRendering="optimizeLegibility"}if(r.rfix){this.addText(o,s+r.c);d.createShift(o,r.rfix/1000);return""}if(u[2]||!this.msieAccentBug||s.length){return s+r.c}d.createShift(o,u[3]/1000);d.createShift(o,(u[4]-u[3])/1000);this.addText(o,r.c);d.createShift(o,-u[4]/1000);return""},handleImg:function(o,m,r,q,p){return p},lookupChar:function(r,u){var q,o;if(!r.FONTS){var t=this.FONTDATA.FONTS;var s=(r.fonts||this.FONTDATA.VARIANT.normal.fonts);if(!(s instanceof Array)){s=[s]}if(r.fonts!=s){r.fonts=s}r.FONTS=[];for(q=0,o=s.length;q<o;q++){if(t[s[q]]){r.FONTS.push(t[s[q]]);t[s[q]].name=s[q]}}}for(q=0,o=r.FONTS.length;q<o;q++){var p=r.FONTS[q];if(typeof(p)==="string"){delete r.FONTS;this.loadFont(p)}if(p[u]){if(p[u].length===5){p[u][5]={}}if(d.allowWebFonts&&!p.available){this.loadWebFont(p)}else{return p}}else{this.findBlock(p,u)}}return this.unknownChar(r,u)},unknownChar:function(m,p){var o=(m.defaultFont||{family:d.config.undefinedFamily});if(m.bold){o.weight="bold"}if(m.italic){o.style="italic"}if(!o[p]){o[p]=[800,200,500,0,500,{isUnknown:true}]}b.signal.Post(["HTML-CSS Jax - unknown char",p,m]);return o},findBlock:function(o,s){if(o.Ranges){for(var r=0,n=o.Ranges.length;r<n;r++){if(s<o.Ranges[r][0]){return}if(s<=o.Ranges[r][1]){var q=o.Ranges[r][2];for(var p=o.Ranges.length-1;p>=0;p--){if(o.Ranges[p][2]==q){o.Ranges.splice(p,1)}}this.loadFont(o.directory+"/"+q+".js")}}}},loadFont:function(n){var m=MathJax.Callback.Queue();m.Push(["Require",j,this.fontDir+"/"+n]);if(this.imgFonts){if(!MathJax.isPacked){n=n.replace(/\/([^\/]*)$/,d.imgPacked+"/$1")}m.Push(["Require",j,this.webfontDir+"/png/"+n])}b.RestartAfter(m.Push({}))},loadWebFont:function(m){m.available=m.isWebFont=true;if(d.FontFaceBug){m.family=m.name;if(d.msieFontCSSBug){m.family+="-Web"}}b.RestartAfter(this.Font.loadWebFont(m))},loadWebFontError:function(n,m){b.Startup.signal.Post("HTML-CSS Jax - disable web fonts");n.isWebFont=false;if(this.config.imageFont&&this.config.imageFont===this.fontInUse){this.imgFonts=true;b.Startup.signal.Post("HTML-CSS Jax - switch to image fonts");b.Startup.signal.Post("HTML-CSS Jax - using image fonts");e(["WebFontNotAvailable","Web-Fonts not available -- using image fonts instead"],null,3000);j.Require(this.directory+"/imageFonts.js",m)}else{this.allowWebFonts=false;m()}},Element:MathJax.HTML.Element,addElement:MathJax.HTML.addElement,TextNode:MathJax.HTML.TextNode,addText:MathJax.HTML.addText,ucMatch:MathJax.HTML.ucMatch,BIGDIMEN:10000000,ID:0,idPostfix:"",GetID:function(){this.ID++;return this.ID},MATHSPACE:{veryverythinmathspace:1/18,verythinmathspace:2/18,thinmathspace:3/18,mediummathspace:4/18,thickmathspace:5/18,verythickmathspace:6/18,veryverythickmathspace:7/18,negativeveryverythinmathspace:-1/18,negativeverythinmathspace:-2/18,negativethinmathspace:-3/18,negativemediummathspace:-4/18,negativethickmathspace:-5/18,negativeverythickmathspace:-6/18,negativeveryverythickmathspace:-7/18},TeX:{x_height:0.430554,quad:1,num1:0.676508,num2:0.393732,num3:0.44373,denom1:0.685951,denom2:0.344841,sup1:0.412892,sup2:0.362892,sup3:0.288888,sub1:0.15,sub2:0.247217,sup_drop:0.386108,sub_drop:0.05,delim1:2.39,delim2:1,axis_height:0.25,rule_thickness:0.06,big_op_spacing1:0.111111,big_op_spacing2:0.166666,big_op_spacing3:0.2,big_op_spacing4:0.6,big_op_spacing5:0.1,scriptspace:0.1,nulldelimiterspace:0.12,delimiterfactor:901,delimitershortfall:0.3,min_rule_thickness:1.25},NBSP:"\u00A0",rfuzz:0});MathJax.Hub.Register.StartupHook("mml Jax Ready",function(){i=MathJax.ElementJax.mml;i.mbase.Augment({toHTML:function(q){q=this.HTMLcreateSpan(q);if(this.type!="mrow"){q=this.HTMLhandleSize(q)}for(var o=0,n=this.data.length;o<n;o++){if(this.data[o]){this.data[o].toHTML(q)}}var u=this.HTMLcomputeBBox(q);var p=q.bbox.h,t=q.bbox.d,r=false,s;for(o=0,n=u.length;o<n;o++){s=u[o].HTMLspanElement().bbox;if(u[o].forceStretch||s.h!==p||s.d!==t){u[o].HTMLstretchV(q,p,t);r=true}else{if(u[o].needsBBox){r=true}}}if(r){this.HTMLcomputeBBox(q,true)}if(this.HTMLlineBreaks(q)){q=this.HTMLmultiline(q)}this.HTMLhandleSpace(q);this.HTMLhandleColor(q);if(this.data.length===1&&this.data[0]){s=this.data[0].HTMLspanElement().bbox;if(s.skew){q.bbox.skew=s.skew}}return q},HTMLlineBreaks:function(){return false},HTMLmultiline:function(){i.mbase.HTMLautoloadFile("multiline")},HTMLcomputeBBox:function(s,r,q,n){if(q==null){q=0}if(n==null){n=this.data.length}var p=s.bbox={exactW:true},t=[];while(q<n){var o=this.data[q];if(!o){continue}if(!r&&o.HTMLcanStretch("Vertical")){t.push(o);o=(o.CoreMO()||o);t[t.length-1].needsBBox=(o!==this.data[q])}this.HTMLcombineBBoxes(o,p);q++}this.HTMLcleanBBox(p);return t},HTMLcombineBBoxes:function(m,n){if(n.w==null){this.HTMLemptyBBox(n)}var p=(m.bbox?m:m.HTMLspanElement());if(!p||!p.bbox){return}var o=p.bbox;if(o.d>n.d){n.d=o.d}if(o.h>n.h){n.h=o.h}if(o.D!=null&&o.D>n.D){n.D=o.D}if(o.H!=null&&o.H>n.H){n.H=o.H}if(p.style.paddingLeft){n.w+=d.unEm(p.style.paddingLeft)*(p.scale||1)}if(n.w+o.lw<n.lw){n.lw=n.w+o.lw}if(n.w+o.rw>n.rw){n.rw=n.w+o.rw}n.w+=o.w;if(p.style.paddingRight){n.w+=d.unEm(p.style.paddingRight)*(p.scale||1)}if(o.width){n.width=o.width;n.minWidth=o.minWidth}if(o.tw){n.tw=o.tw}if(o.ic){n.ic=o.ic}else{delete n.ic}if(n.exactW&&!o.exactW){n.exactW=o.exactW}},HTMLemptyBBox:function(m){m.h=m.d=m.H=m.D=m.rw=-d.BIGDIMEN;m.w=0;m.lw=d.BIGDIMEN;return m},HTMLcleanBBox:function(m){if(m.h===this.BIGDIMEN){m.h=m.d=m.H=m.D=m.w=m.rw=m.lw=0}if(m.D<=m.d){delete m.D}if(m.H<=m.h){delete m.H}},HTMLzeroBBox:function(){return{h:0,d:0,w:0,lw:0,rw:0}},HTMLcanStretch:function(n){if(this.isEmbellished()){var m=this.Core();if(m&&m!==this){return m.HTMLcanStretch(n)}}return false},HTMLstretchH:function(n,m){return this.HTMLspanElement()},HTMLstretchV:function(n,m,o){return this.HTMLspanElement()},HTMLnotEmpty:function(m){while(m){if((m.type!=="mrow"&&m.type!=="texatom")||m.data.length>1){return true}m=m.data[0]}return false},HTMLmeasureChild:function(o,m){if(this.data[o]){d.Measured(this.data[o].toHTML(m),m)}else{m.bbox=this.HTMLzeroBBox()}},HTMLboxChild:function(o,m){if(!this.data[o]){this.SetData(o,i.mrow())}return this.data[o].toHTML(m)},HTMLcreateSpan:function(m){if(this.spanID){var n=this.HTMLspanElement();if(n&&(n.parentNode===m||(n.parentNode||{}).parentNode===m)){while(n.firstChild){n.removeChild(n.firstChild)}n.bbox=this.HTMLzeroBBox();n.scale=1;n.isMultChar=n.HH=null;n.style.cssText="";return n}}if(this.href){m=d.addElement(m,"a",{href:this.href,isMathJax:true})}m=d.addElement(m,"span",{className:this.type,isMathJax:true});if(d.imgHeightBug){m.style.display="inline-block"}if(this["class"]){m.className+=" "+this["class"]}if(!this.spanID){this.spanID=d.GetID()}m.id=(this.id||"MathJax-Span-"+this.spanID)+d.idPostfix;m.bbox=this.HTMLzeroBBox();this.styles={};if(this.style){m.style.cssText=this.style;if(m.style.fontSize){this.mathsize=m.style.fontSize;m.style.fontSize=""}this.styles={border:d.getBorders(m),padding:d.getPadding(m)};if(this.styles.border){m.style.border=""}if(this.styles.padding){m.style.padding=""}}if(this.href){m.parentNode.bbox=m.bbox}this.HTMLaddAttributes(m);return m},HTMLaddAttributes:function(p){if(this.attrNames){var u=this.attrNames,q=i.nocopyAttributes,t=b.config.ignoreMMLattributes;var r=(this.type==="mstyle"?i.math.prototype.defaults:this.defaults);for(var o=0,n=u.length;o<n;o++){var s=u[o];if(t[s]==false||(!q[s]&&!t[s]&&r[s]==null&&typeof(p[s])==="undefined")){p.setAttribute(s,this.attr[s])}}}},HTMLspanElement:function(){if(!this.spanID){return null}return document.getElementById((this.id||"MathJax-Span-"+this.spanID)+d.idPostfix)},HTMLhandleVariant:function(n,m,o){d.handleVariant(n,m,o)},HTMLhandleSize:function(m){if(!m.scale){m.scale=this.HTMLgetScale();if(m.scale!==1){m.style.fontSize=d.Percent(m.scale)}}return m},HTMLhandleDir:function(n){var m=this.Get("dir",true);if(m){n.dir=m}return n},HTMLhandleColor:function(y){var A=this.getValues("mathcolor","color");if(this.mathbackground){A.mathbackground=this.mathbackground}if(this.background){A.background=this.background}if(this.style&&y.style.backgroundColor){A.mathbackground=y.style.backgroundColor;y.style.backgroundColor="transparent"}var v=(this.styles||{}).border,x=(this.styles||{}).padding;if(A.color&&!this.mathcolor){A.mathcolor=A.color}if(A.background&&!this.mathbackground){A.mathbackground=A.background}if(A.mathcolor){y.style.color=A.mathcolor}if((A.mathbackground&&A.mathbackground!==i.COLOR.TRANSPARENT)||v||x){var C=y.bbox,B=(C.exact?0:1/d.em),w=0,u=0,o=y.style.paddingLeft,s=y.style.paddingRight;if(this.isToken){w=C.lw;u=C.rw-C.w}if(o!==""){w+=d.unEm(o)*(y.scale||1)}if(s!==""){u-=d.unEm(s)*(y.scale||1)}var n=(d.PaddingWidthBug||C.keepPadding||C.exactW?0:u-w);var q=Math.max(0,d.getW(y)+n);var z=C.h+C.d,m=-C.d,t=0,r=0;if(q>0){q+=2*B;w-=B}if(z>0){z+=2*B;m-=B}u=-q-w;if(v){u-=v.right;m-=v.bottom;t+=v.left;r+=v.right;C.h+=v.top;C.d+=v.bottom;C.w+=v.left+v.right;C.lw-=v.left;C.rw+=v.right}if(x){z+=x.top+x.bottom;q+=x.left+x.right;u-=x.right;m-=x.bottom;t+=x.left;r+=x.right;C.h+=x.top;C.d+=x.bottom;C.w+=x.left+x.right;C.lw-=x.left;C.rw+=x.right}if(r){y.style.paddingRight=d.Em(r)}var p=d.Element("span",{id:"MathJax-Color-"+this.spanID+d.idPostfix,isMathJax:true,style:{display:"inline-block",backgroundColor:A.mathbackground,width:d.Em(q),height:d.Em(z),verticalAlign:d.Em(m),marginLeft:d.Em(w),marginRight:d.Em(u)}});d.setBorders(p,v);if(C.width){p.style.width=C.width;p.style.marginRight="-"+C.width}if(d.msieInlineBlockAlignBug){p.style.position="relative";p.style.width=p.style.height=0;p.style.verticalAlign=p.style.marginLeft=p.style.marginRight="";p.style.border=p.style.padding="";if(v&&d.msieBorderWidthBug){z+=v.top+v.bottom;q+=v.left+v.right}p.style.width=d.Em(t+B);d.placeBox(d.addElement(p,"span",{noAdjust:true,isMathJax:true,style:{display:"inline-block",position:"absolute",overflow:"hidden",background:(A.mathbackground||"transparent"),width:d.Em(q),height:d.Em(z)}}),w,C.h+B);d.setBorders(p.firstChild,v)}y.parentNode.insertBefore(p,y);if(d.msieColorPositionBug){y.style.position="relative"}return p}return null},HTMLremoveColor:function(){var m=document.getElementById("MathJax-Color-"+this.spanID+d.idPostfix);if(m){m.parentNode.removeChild(m)}},HTMLhandleSpace:function(q){if(this.useMMLspacing){if(this.type!=="mo"){return}var o=this.getValues("scriptlevel","lspace","rspace");if(o.scriptlevel<=0||this.hasValue("lspace")||this.hasValue("rspace")){var n=this.HTMLgetMu(q);o.lspace=Math.max(0,d.length2em(o.lspace,n));o.rspace=Math.max(0,d.length2em(o.rspace,n));var m=this,p=this.Parent();while(p&&p.isEmbellished()&&p.Core()===m){m=p;p=p.Parent();q=m.HTMLspanElement()}if(o.lspace){q.style.paddingLeft=d.Em(o.lspace)}if(o.rspace){q.style.paddingRight=d.Em(o.rspace)}}}else{var r=this.texSpacing();if(r!==""){this.HTMLgetScale();r=d.length2em(r,this.scale)/(q.scale||1)*this.mscale;if(q.style.paddingLeft){r+=d.unEm(q.style.paddingLeft)}q.style.paddingLeft=d.Em(r)}}},HTMLgetScale:function(){if(this.scale){return this.scale*this.mscale}var o=1,m=this.getValues("scriptlevel","fontsize");m.mathsize=(this.isToken?this:this.Parent()).Get("mathsize");if(this.style){var n=this.HTMLspanElement();if(n.style.fontSize!=""){m.fontsize=n.style.fontSize}}if(m.fontsize&&!this.mathsize){m.mathsize=m.fontsize}if(m.scriptlevel!==0){if(m.scriptlevel>2){m.scriptlevel=2}o=Math.pow(this.Get("scriptsizemultiplier"),m.scriptlevel);m.scriptminsize=d.length2em(this.Get("scriptminsize"));if(o<m.scriptminsize){o=m.scriptminsize}}this.scale=o;this.mscale=d.length2em(m.mathsize);return o*this.mscale},HTMLgetMu:function(o){var m=1,n=this.getValues("scriptlevel","scriptsizemultiplier");if(o.scale&&o.scale!==1){m=1/o.scale}if(n.scriptlevel!==0){if(n.scriptlevel>2){n.scriptlevel=2}m=Math.sqrt(Math.pow(n.scriptsizemultiplier,n.scriptlevel))}return m},HTMLgetVariant:function(){var m=this.getValues("mathvariant","fontfamily","fontweight","fontstyle");m.hasVariant=this.Get("mathvariant",true);if(!m.hasVariant){m.family=m.fontfamily;m.weight=m.fontweight;m.style=m.fontstyle}if(this.style){var o=this.HTMLspanElement();if(!m.family&&o.style.fontFamily){m.family=o.style.fontFamily}if(!m.weight&&o.style.fontWeight){m.weight=o.style.fontWeight}if(!m.style&&o.style.fontStyle){m.style=o.style.fontStyle}}if(m.weight&&m.weight.match(/^\d+$/)){m.weight=(parseInt(m.weight)>600?"bold":"normal")}var n=m.mathvariant;if(this.variantForm){n="-"+d.fontInUse+"-variant"}if(m.family&&!m.hasVariant){if(!m.weight&&m.mathvariant.match(/bold/)){m.weight="bold"}if(!m.style&&m.mathvariant.match(/italic/)){m.style="italic"}return{FONTS:[],fonts:[],noRemap:true,defaultFont:{family:m.family,style:m.style,weight:m.weight}}}if(m.weight==="bold"){n={normal:i.VARIANT.BOLD,italic:i.VARIANT.BOLDITALIC,fraktur:i.VARIANT.BOLDFRAKTUR,script:i.VARIANT.BOLDSCRIPT,"sans-serif":i.VARIANT.BOLDSANSSERIF,"sans-serif-italic":i.VARIANT.SANSSERIFBOLDITALIC}[n]||n}else{if(m.weight==="normal"){n={bold:i.VARIANT.normal,"bold-italic":i.VARIANT.ITALIC,"bold-fraktur":i.VARIANT.FRAKTUR,"bold-script":i.VARIANT.SCRIPT,"bold-sans-serif":i.VARIANT.SANSSERIF,"sans-serif-bold-italic":i.VARIANT.SANSSERIFITALIC}[n]||n}}if(m.style==="italic"){n={normal:i.VARIANT.ITALIC,bold:i.VARIANT.BOLDITALIC,"sans-serif":i.VARIANT.SANSSERIFITALIC,"bold-sans-serif":i.VARIANT.SANSSERIFBOLDITALIC}[n]||n}else{if(m.style==="normal"){n={italic:i.VARIANT.NORMAL,"bold-italic":i.VARIANT.BOLD,"sans-serif-italic":i.VARIANT.SANSSERIF,"sans-serif-bold-italic":i.VARIANT.BOLDSANSSERIF}[n]||n}}if(!(n in d.FONTDATA.VARIANT)){n="normal"}return d.FONTDATA.VARIANT[n]},HTMLdrawBBox:function(m){var o=m.bbox;var n=d.Element("span",{style:{"font-size":m.style.fontSize,display:"inline-block",opacity:0.25,"margin-left":d.Em(-o.w)}},[["span",{style:{height:d.Em(o.h),width:d.Em(o.w),"background-color":"red",display:"inline-block"}}],["span",{style:{height:d.Em(o.d),width:d.Em(o.w),"margin-left":d.Em(-o.w),"vertical-align":d.Em(-o.d),"background-color":"green",display:"inline-block"}}]]);if(m.nextSibling){m.parentNode.insertBefore(n,m.nextSibling)}else{m.parentNode.appendChild(n)}}},{HTMLautoload:function(){var m=d.autoloadDir+"/"+this.type+".js";b.RestartAfter(j.Require(m))},HTMLautoloadFile:function(m){var n=d.autoloadDir+"/"+m+".js";b.RestartAfter(j.Require(n))},HTMLstretchH:function(n,m){this.HTMLremoveColor();return this.toHTML(n,m)},HTMLstretchV:function(n,m,o){this.HTMLremoveColor();return this.toHTML(n,m,o)}});i.chars.Augment({toHTML:function(p,o,n,q){var t=this.data.join("").replace(/[\u2061-\u2064]/g,"");if(n){t=n(t,q)}if(o.fontInherit){var s=Math.floor(d.config.scale/d.scale+0.5)+"%";d.addElement(p,"span",{style:{"font-size":s}},[t]);if(o.bold){p.lastChild.style.fontWeight="bold"}if(o.italic){p.lastChild.style.fontStyle="italic"}p.bbox=null;var r=d.getHD(p),m=d.getW(p);p.bbox={h:r.h,d:r.d,w:m,lw:0,rw:m,exactW:true}}else{this.HTMLhandleVariant(p,o,t)}}});i.entity.Augment({toHTML:function(p,o,n,q){var t=this.toString().replace(/[\u2061-\u2064]/g,"");if(n){t=n(t,q)}if(o.fontInherit){var s=Math.floor(d.config.scale/d.scale+0.5)+"%";d.addElement(p,"span",{style:{"font-size":s}},[t]);if(o.bold){p.lastChild.style.fontWeight="bold"}if(o.italic){p.lastChild.style.fontStyle="italic"}delete p.bbox;var r=d.getHD(p),m=d.getW(p);p.bbox={h:r.h,d:r.d,w:m,lw:0,rw:m,exactW:true}}else{this.HTMLhandleVariant(p,o,t)}}});i.mi.Augment({toHTML:function(q){q=this.HTMLhandleSize(this.HTMLcreateSpan(q));q.bbox=null;var p=this.HTMLgetVariant();for(var o=0,n=this.data.length;o<n;o++){if(this.data[o]){this.data[o].toHTML(q,p)}}if(!q.bbox){q.bbox=this.HTMLzeroBBox()}var s=this.data.join(""),r=q.bbox;if(r.skew&&s.length!==1){delete r.skew}if(r.rw>r.w&&s.length===1&&!p.noIC){r.ic=r.rw-r.w;d.createBlank(q,r.ic/this.mscale);r.w=r.rw}this.HTMLhandleSpace(q);this.HTMLhandleColor(q);this.HTMLhandleDir(q);return q}});i.mn.Augment({HTMLremapMinus:function(m){return m.replace(/^-/,"\u2212")},toHTML:function(r){r=this.HTMLhandleSize(this.HTMLcreateSpan(r));r.bbox=null;var q=this.HTMLgetVariant();var p=this.HTMLremapMinus;for(var o=0,n=this.data.length;o<n;o++){if(this.data[o]){this.data[o].toHTML(r,q,p);p=null}}if(!r.bbox){r.bbox=this.HTMLzeroBBox()}if(this.data.join("").length!==1){delete r.bbox.skew}this.HTMLhandleSpace(r);this.HTMLhandleColor(r);this.HTMLhandleDir(r);return r}});i.mo.Augment({toHTML:function(x){x=this.HTMLhandleSize(this.HTMLcreateSpan(x));if(this.data.length==0){return x}else{x.bbox=null}var A=this.data.join("");var s=this.HTMLgetVariant();var z=this.getValues("largeop","displaystyle");if(z.largeop){s=d.FONTDATA.VARIANT[z.displaystyle?"-largeOp":"-smallOp"]}var y=this.CoreParent(),r=(y&&y.isa(i.msubsup)&&this!==y.data[y.base]),o=(r?this.remapChars:null);if(A.length===1&&y&&y.isa(i.munderover)&&this.CoreText(y.data[y.base]).length===1){var u=y.data[y.over],w=y.data[y.under];if(u&&this===u.CoreMO()&&y.Get("accent")){o=d.FONTDATA.REMAPACCENT}else{if(w&&this===w.CoreMO()&&y.Get("accentunder")){o=d.FONTDATA.REMAPACCENTUNDER}}}if(r&&A.match(/['`"\u00B4\u2032-\u2037\u2057]/)){s=d.FONTDATA.VARIANT["-"+d.fontInUse+"-variant"]}for(var t=0,q=this.data.length;t<q;t++){if(this.data[t]){this.data[t].toHTML(x,s,this.remap,o)}}if(!x.bbox){x.bbox=this.HTMLzeroBBox()}if(A.length!==1){delete x.bbox.skew}if(d.AccentBug&&x.bbox.w===0&&A.length===1&&x.firstChild){x.firstChild.nodeValue+=d.NBSP;d.createSpace(x,0,0,-x.offsetWidth/d.em)}if(z.largeop){var v=d.TeX.axis_height*this.scale*this.mscale;var n=(x.bbox.h-x.bbox.d)/2-v;if(d.safariVerticalAlignBug&&x.lastChild.nodeName==="IMG"){x.lastChild.style.verticalAlign=d.Em(d.unEm(x.lastChild.style.verticalAlign||0)/d.em-n/x.scale)}else{if(d.konquerorVerticalAlignBug&&x.lastChild.nodeName==="IMG"){x.style.position="relative";x.lastChild.style.position="relative";x.lastChild.style.top=d.Em(n/x.scale)}else{x.style.verticalAlign=d.Em(-n/x.scale)}}x.bbox.h-=n;x.bbox.d+=n;if(x.bbox.rw>x.bbox.w){x.bbox.ic=x.bbox.rw-x.bbox.w;d.createBlank(x,x.bbox.ic/this.mscale);x.bbox.w=x.bbox.rw}}this.HTMLhandleSpace(x);this.HTMLhandleColor(x);this.HTMLhandleDir(x);return x},HTMLcanStretch:function(q){if(!this.Get("stretchy")){return false}var r=this.data.join("");if(r.length>1){return false}var o=this.CoreParent();if(o&&o.isa(i.munderover)&&this.CoreText(o.data[o.base]).length===1){var p=o.data[o.over],n=o.data[o.under];if(p&&this===p.CoreMO()&&o.Get("accent")){r=d.FONTDATA.REMAPACCENT[r]||r}else{if(n&&this===n.CoreMO()&&o.Get("accentunder")){r=d.FONTDATA.REMAPACCENTUNDER[r]||r}}}r=d.FONTDATA.DELIMITERS[r.charCodeAt(0)];var m=(r&&r.dir===q.substr(0,1));this.forceStretch=(m&&(this.Get("minsize",true)||this.Get("maxsize",true)));return m},HTMLstretchV:function(o,p,q){this.HTMLremoveColor();var t=this.getValues("symmetric","maxsize","minsize");var r=this.HTMLspanElement(),u=this.HTMLgetMu(r),s;var n=this.HTMLgetScale(),m=d.TeX.axis_height*n;if(t.symmetric){s=2*Math.max(p-m,q+m)}else{s=p+q}t.maxsize=d.length2em(t.maxsize,u,r.bbox.h+r.bbox.d);t.minsize=d.length2em(t.minsize,u,r.bbox.h+r.bbox.d);s=Math.max(t.minsize,Math.min(t.maxsize,s));if(s!=t.minsize){s=[Math.max(s*d.TeX.delimiterfactor/1000,s-d.TeX.delimitershortfall),s]}r=this.HTMLcreateSpan(o);d.createDelimiter(r,this.data.join("").charCodeAt(0),s,n);if(t.symmetric){s=(r.bbox.h+r.bbox.d)/2+m}else{s=(r.bbox.h+r.bbox.d)*p/(p+q)}d.positionDelimiter(r,s);this.HTMLhandleSpace(r);this.HTMLhandleColor(r);return r},HTMLstretchH:function(q,m){this.HTMLremoveColor();var o=this.getValues("maxsize","minsize","mathvariant","fontweight");if((o.fontweight==="bold"||parseInt(o.fontweight)>=600)&&!this.Get("mathvariant",true)){o.mathvariant=i.VARIANT.BOLD}var p=this.HTMLspanElement(),n=this.HTMLgetMu(p),r=p.scale;o.maxsize=d.length2em(o.maxsize,n,p.bbox.w);o.minsize=d.length2em(o.minsize,n,p.bbox.w);m=Math.max(o.minsize,Math.min(o.maxsize,m));p=this.HTMLcreateSpan(q);d.createDelimiter(p,this.data.join("").charCodeAt(0),m,r,o.mathvariant);this.HTMLhandleSpace(p);this.HTMLhandleColor(p);return p}});i.mtext.Augment({toHTML:function(q){q=this.HTMLhandleSize(this.HTMLcreateSpan(q));var p=this.HTMLgetVariant();if(d.config.mtextFontInherit||this.Parent().type==="merror"){var r=this.Get("mathvariant");if(r==="monospace"){q.className+=" MJX-monospace"}else{if(r.match(/sans-serif/)){q.className+=" MJX-sans-serif"}}p={bold:p.bold,italic:p.italic,fontInherit:true}}for(var o=0,n=this.data.length;o<n;o++){if(this.data[o]){this.data[o].toHTML(q,p)}}if(!q.bbox){q.bbox=this.HTMLzeroBBox()}if(this.data.join("").length!==1){delete q.bbox.skew}this.HTMLhandleSpace(q);this.HTMLhandleColor(q);this.HTMLhandleDir(q);return q}});i.merror.Augment({toHTML:function(n){var p=MathJax.HTML.addElement(n,"span",{style:{display:"inline-block"}});n=this.SUPER(arguments).toHTML.call(this,p);var o=d.getHD(p),m=d.getW(p);p.bbox={h:o.h,d:o.d,w:m,lw:0,rw:m,exactW:true};p.id=n.id;n.id=null;return p}});i.ms.Augment({toHTML:i.mbase.HTMLautoload});i.mglyph.Augment({toHTML:i.mbase.HTMLautoload});i.mspace.Augment({toHTML:function(q){q=this.HTMLcreateSpan(q);var o=this.getValues("height","depth","width");var n=this.HTMLgetMu(q);this.HTMLgetScale();o.mathbackground=this.mathbackground;if(this.background&&!this.mathbackground){o.mathbackground=this.background}var p=d.length2em(o.height,n)*this.mscale,r=d.length2em(o.depth,n)*this.mscale,m=d.length2em(o.width,n)*this.mscale;d.createSpace(q,p,r,m,o.mathbackground,true);return q}});i.mphantom.Augment({toHTML:function(q,o,s){q=this.HTMLcreateSpan(q);if(this.data[0]!=null){var r=this.data[0].toHTML(q);if(s!=null){d.Remeasured(this.data[0].HTMLstretchV(q,o,s),q)}else{if(o!=null){d.Remeasured(this.data[0].HTMLstretchH(q,o),q)}else{r=d.Measured(r,q)}}q.bbox={w:r.bbox.w,h:r.bbox.h,d:r.bbox.d,lw:0,rw:0,exactW:true};for(var p=0,n=q.childNodes.length;p<n;p++){q.childNodes[p].style.visibility="hidden"}}this.HTMLhandleSpace(q);this.HTMLhandleColor(q);return q},HTMLstretchH:i.mbase.HTMLstretchH,HTMLstretchV:i.mbase.HTMLstretchV});i.mpadded.Augment({toHTML:function(u,o,m){u=this.HTMLcreateSpan(u);if(this.data[0]!=null){var s=d.createStack(u,true);var p=d.createBox(s);var n=this.data[0].toHTML(p);if(m!=null){d.Remeasured(this.data[0].HTMLstretchV(p,o,m),p)}else{if(o!=null){d.Remeasured(this.data[0].HTMLstretchH(p,o),p)}else{d.Measured(n,p)}}var v=this.getValues("height","depth","width","lspace","voffset"),t=0,r=0,w=this.HTMLgetMu(u);this.HTMLgetScale();if(v.lspace){t=this.HTMLlength2em(p,v.lspace,w)}if(v.voffset){r=this.HTMLlength2em(p,v.voffset,w)}d.placeBox(p,t,r);t/=this.mscale;r/=this.mscale;u.bbox={h:p.bbox.h,d:p.bbox.d,w:p.bbox.w,exactW:true,lw:p.bbox.lw+t,rw:p.bbox.rw+t,H:Math.max((p.bbox.H==null?-d.BIGDIMEN:p.bbox.H+r),p.bbox.h+r),D:Math.max((p.bbox.D==null?-d.BIGDIMEN:p.bbox.D-r),p.bbox.d-r)};if(v.height!==""){u.bbox.h=this.HTMLlength2em(p,v.height,w,"h",0)}if(v.depth!==""){u.bbox.d=this.HTMLlength2em(p,v.depth,w,"d",0)}if(v.width!==""){u.bbox.w=this.HTMLlength2em(p,v.width,w,"w",0)}if(u.bbox.H<=u.bbox.h){delete u.bbox.H}if(u.bbox.D<=u.bbox.d){delete u.bbox.D}var q=/^\s*(\d+(\.\d*)?|\.\d+)\s*(pt|em|ex|mu|px|pc|in|mm|cm)\s*$/;u.bbox.exact=!!((this.data[0]&&this.data[0].data.length==0)||q.exec(v.height)||q.exec(v.width)||q.exec(v.depth));d.setStackWidth(s,u.bbox.w)}this.HTMLhandleSpace(u);this.HTMLhandleColor(u);return u},HTMLlength2em:function(s,t,o,u,n){if(n==null){n=-d.BIGDIMEN}var q=String(t).match(/width|height|depth/);var r=(q?s.bbox[q[0].charAt(0)]:(u?s.bbox[u]:0));var p=d.length2em(t,o,r/this.mscale)*this.mscale;if(u&&String(t).match(/^\s*[-+]/)){return Math.max(n,s.bbox[u]+p)}else{return p}},HTMLstretchH:i.mbase.HTMLstretchH,HTMLstretchV:i.mbase.HTMLstretchV});i.mrow.Augment({HTMLlineBreaks:function(m){if(!this.parent.linebreakContainer){return false}return(d.config.linebreaks.automatic&&m.bbox.w>d.linebreakWidth)||this.hasNewline()},HTMLstretchH:function(o,m){this.HTMLremoveColor();var n=this.HTMLspanElement();this.data[this.core].HTMLstretchH(n,m);this.HTMLcomputeBBox(n,true);this.HTMLhandleColor(n);return n},HTMLstretchV:function(o,n,p){this.HTMLremoveColor();var m=this.HTMLspanElement();this.data[this.core].HTMLstretchV(m,n,p);this.HTMLcomputeBBox(m,true);this.HTMLhandleColor(m);return m}});i.mstyle.Augment({toHTML:function(n,m,o){n=this.HTMLcreateSpan(n);if(this.data[0]!=null){var p=this.data[0].toHTML(n);if(o!=null){this.data[0].HTMLstretchV(n,m,o)}else{if(m!=null){this.data[0].HTMLstretchH(n,m)}}n.bbox=p.bbox}this.HTMLhandleSpace(n);this.HTMLhandleColor(n);return n},HTMLstretchH:i.mbase.HTMLstretchH,HTMLstretchV:i.mbase.HTMLstretchV});i.mfrac.Augment({toHTML:function(F){F=this.HTMLcreateSpan(F);var o=d.createStack(F);var w=d.createBox(o),s=d.createBox(o);d.MeasureSpans([this.HTMLboxChild(0,w),this.HTMLboxChild(1,s)]);var m=this.getValues("displaystyle","linethickness","numalign","denomalign","bevelled");var K=this.HTMLgetScale(),E=m.displaystyle;var J=d.TeX.axis_height*K;if(m.bevelled){var I=(E?0.4:0.15);var x=Math.max(w.bbox.h+w.bbox.d,s.bbox.h+s.bbox.d)+2*I;var G=d.createBox(o);d.createDelimiter(G,47,x);d.placeBox(w,0,(w.bbox.d-w.bbox.h)/2+J+I);d.placeBox(G,w.bbox.w-I/2,(G.bbox.d-G.bbox.h)/2+J);d.placeBox(s,w.bbox.w+G.bbox.w-I,(s.bbox.d-s.bbox.h)/2+J-I)}else{var n=Math.max(w.bbox.w,s.bbox.w);var A=d.thickness2em(m.linethickness,this.scale)*this.mscale,C,B,z,y;var D=d.TeX.min_rule_thickness/this.em;if(E){z=d.TeX.num1;y=d.TeX.denom1}else{z=(A===0?d.TeX.num3:d.TeX.num2);y=d.TeX.denom2}z*=K;y*=K;if(A===0){C=Math.max((E?7:3)*d.TeX.rule_thickness,2*D);B=(z-w.bbox.d)-(s.bbox.h-y);if(B<C){z+=(C-B)/2;y+=(C-B)/2}}else{C=Math.max((E?2:0)*D+A,A/2+1.5*D);B=(z-w.bbox.d)-(J+A/2);if(B<C){z+=C-B}B=(J-A/2)-(s.bbox.h-y);if(B<C){y+=C-B}var r=d.createBox(o);d.createRule(r,A,0,n+2*A);d.placeBox(r,0,J-A/2)}d.alignBox(w,m.numalign,z,0,true);d.alignBox(s,m.denomalign,-y,0,true)}this.HTMLhandleSpace(F);this.HTMLhandleColor(F);return F},HTMLcanStretch:function(m){return false},HTMLhandleSpace:function(n){if(!this.texWithDelims&&!this.useMMLspacing){var o=d.TeX.nulldelimiterspace*this.mscale;var m=n.childNodes[d.msiePaddingWidthBug?1:0].style;m.marginLeft=m.marginRight=d.Em(o);n.bbox.w+=2*o;n.bbox.rw+=2*o}this.SUPER(arguments).HTMLhandleSpace.call(this,n)}});i.msqrt.Augment({toHTML:function(z){z=this.HTMLcreateSpan(z);var B=d.createStack(z);var r=d.createBox(B),w=d.createBox(B),v=d.createBox(B);var u=this.HTMLgetScale();var C=d.TeX.rule_thickness*u,o,n,A,s;if(this.Get("displaystyle")){o=d.TeX.x_height*u}else{o=C}n=Math.max(C+o/4,1.5*d.TeX.min_rule_thickness/this.em);var m=this.HTMLboxChild(0,r);A=m.bbox.h+m.bbox.d+n+C;d.createDelimiter(v,8730,A,u);d.MeasureSpans([m,v]);s=m.bbox.w;var y=0;if(v.isMultiChar||(d.AdjustSurd&&d.imgFonts)){v.bbox.w*=0.95}if(v.bbox.h+v.bbox.d>A){n=((v.bbox.h+v.bbox.d)-(A-C))/2}var D=d.FONTDATA.DELIMITERS[d.FONTDATA.RULECHAR];if(!D||s<(D.HW[0]||[0])[0]*u||u<0.75){d.createRule(w,0,C,s);w.bbox.h=-C}else{d.createDelimiter(w,d.FONTDATA.RULECHAR,s,u)}A=m.bbox.h+n+C;n=A*d.rfuzz;if(v.isMultiChar){n=d.rfuzz}y=this.HTMLaddRoot(B,v,y,v.bbox.h+v.bbox.d-A,u);d.placeBox(v,y,A-v.bbox.h);d.placeBox(w,y+v.bbox.w,A-w.bbox.h+n);d.placeBox(r,y+v.bbox.w,0);this.HTMLhandleSpace(z);this.HTMLhandleColor(z);return z},HTMLaddRoot:function(o,n,m,q,p){return m}});i.mroot.Augment({toHTML:i.msqrt.prototype.toHTML,HTMLaddRoot:function(u,n,s,q,m){var o=d.createBox(u);if(this.data[1]){var r=this.data[1].toHTML(o);r.style.paddingRight=r.style.paddingLeft="";d.Measured(r,o)}else{o.bbox=this.HTMLzeroBBox()}var p=this.HTMLrootHeight(n.bbox.h+n.bbox.d,m,o)-q;var t=Math.min(o.bbox.w,o.bbox.rw);s=Math.max(t,n.offset);d.placeBox(o,s-t,p);return s-n.offset},HTMLrootHeight:function(o,n,m){return 0.45*(o-0.9*n)+0.6*n+Math.max(0,m.bbox.d-0.075)}});i.mfenced.Augment({toHTML:function(q){q=this.HTMLcreateSpan(q);if(this.data.open){this.data.open.toHTML(q)}if(this.data[0]!=null){this.data[0].toHTML(q)}for(var o=1,n=this.data.length;o<n;o++){if(this.data[o]){if(this.data["sep"+o]){this.data["sep"+o].toHTML(q)}this.data[o].toHTML(q)}}if(this.data.close){this.data.close.toHTML(q)}var s=this.HTMLcomputeBBox(q);var p=q.bbox.h,r=q.bbox.d;for(o=0,n=s.length;o<n;o++){s[o].HTMLstretchV(q,p,r)}if(s.length){this.HTMLcomputeBBox(q,true)}this.HTMLhandleSpace(q);this.HTMLhandleColor(q);return q},HTMLcomputeBBox:function(r,q){var o=r.bbox={},s=[];this.HTMLcheckStretchy(this.data.open,o,s,q);this.HTMLcheckStretchy(this.data[0],o,s,q);for(var p=1,n=this.data.length;p<n;p++){if(this.data[p]){this.HTMLcheckStretchy(this.data["sep"+p],o,s,q);this.HTMLcheckStretchy(this.data[p],o,s,q)}}this.HTMLcheckStretchy(this.data.close,o,s,q);this.HTMLcleanBBox(o);return s},HTMLcheckStretchy:function(m,n,p,o){if(m){if(!o&&m.HTMLcanStretch("Vertical")){p.push(m);m=(m.CoreMO()||m)}this.HTMLcombineBBoxes(m,n)}}});i.menclose.Augment({toHTML:i.mbase.HTMLautoload});i.maction.Augment({toHTML:i.mbase.HTMLautoload});i.semantics.Augment({toHTML:function(n,m,o){n=this.HTMLcreateSpan(n);if(this.data[0]!=null){var p=this.data[0].toHTML(n);if(o!=null){this.data[0].HTMLstretchV(n,m,o)}else{if(m!=null){this.data[0].HTMLstretchH(n,m)}}n.bbox=p.bbox}this.HTMLhandleSpace(n);return n},HTMLstretchH:i.mbase.HTMLstretchH,HTMLstretchV:i.mbase.HTMLstretchV});i.munderover.Augment({toHTML:function(M,I,G){var n=this.getValues("displaystyle","accent","accentunder","align");var q=this.data[this.base];if(!n.displaystyle&&q!=null&&(q.movablelimits||q.CoreMO().Get("movablelimits"))){return i.msubsup.prototype.toHTML.call(this,M)}M=this.HTMLcreateSpan(M);var Q=this.HTMLgetScale();var r=d.createStack(M);var s=[],p=[],O=[],z,N,J;for(N=0,J=this.data.length;N<J;N++){if(this.data[N]!=null){z=s[N]=d.createBox(r);p[N]=this.data[N].toHTML(z);if(N==this.base){if(G!=null){this.data[this.base].HTMLstretchV(z,I,G)}else{if(I!=null){this.data[this.base].HTMLstretchH(z,I)}}O[N]=(G==null&&I!=null?false:this.data[N].HTMLcanStretch("Horizontal"))}else{O[N]=this.data[N].HTMLcanStretch("Horizontal");p[N].style.paddingLeft=p[N].style.paddingRight=""}}}d.MeasureSpans(p);var o=-d.BIGDIMEN,L=o;for(N=0,J=this.data.length;N<J;N++){if(this.data[N]){if(s[N].bbox.w>L){L=s[N].bbox.w}if(!O[N]&&L>o){o=L}}}if(G==null&&I!=null){o=I}else{if(o==-d.BIGDIMEN){o=L}}for(N=L=0,J=this.data.length;N<J;N++){if(this.data[N]){z=s[N];if(O[N]){z.bbox=this.data[N].HTMLstretchH(z,o).bbox;if(N!==this.base){p[N].style.paddingLeft=p[N].style.paddingRight=""}}if(z.bbox.w>L){L=z.bbox.w}}}var F=d.TeX.rule_thickness*this.mscale,H=d.FONTDATA.TeX_factor;var w,u,B,A,v,E,K,P=0;q=s[this.base]||{bbox:this.HTMLzeroBBox()};if(q.bbox.ic){P=1.3*q.bbox.ic+0.05}for(N=0,J=this.data.length;N<J;N++){if(this.data[N]!=null){z=s[N];v=d.TeX.big_op_spacing5*Q;var C=(N!=this.base&&n[this.ACCENTS[N]]);if(C&&z.bbox.w<=1/d.em+0.0001){z.bbox.w=z.bbox.rw-z.bbox.lw;z.bbox.noclip=true;if(z.bbox.lw){z.insertBefore(d.createSpace(z.parentNode,0,0,-z.bbox.lw),z.firstChild)}d.createBlank(z,0,0,z.bbox.rw+0.1)}E={left:0,center:(L-z.bbox.w)/2,right:L-z.bbox.w}[n.align];w=E;u=0;if(N==this.over){if(C){K=Math.max(F*Q*H,2.5/this.em);v=0;if(q.bbox.skew){w+=q.bbox.skew;M.bbox.skew=q.bbox.skew;if(w+z.bbox.w>L){M.bbox.skew+=(L-z.bbox.w-w)/2}}}else{B=d.TeX.big_op_spacing1*Q*H;A=d.TeX.big_op_spacing3*Q*H;K=Math.max(B,A-Math.max(0,z.bbox.d))}K=Math.max(K,1.5/this.em);w+=P/2;u=q.bbox.h+z.bbox.d+K;z.bbox.h+=v}else{if(N==this.under){if(C){K=3*F*Q*H;v=0}else{B=d.TeX.big_op_spacing2*Q*H;A=d.TeX.big_op_spacing4*Q*H;K=Math.max(B,A-z.bbox.h)}K=Math.max(K,1.5/this.em);w-=P/2;u=-(q.bbox.d+z.bbox.h+K);z.bbox.d+=v}}d.placeBox(z,w,u)}}this.HTMLhandleSpace(M);this.HTMLhandleColor(M);return M},HTMLstretchH:i.mbase.HTMLstretchH,HTMLstretchV:i.mbase.HTMLstretchV});i.msubsup.Augment({toHTML:function(M,K,F){M=this.HTMLcreateSpan(M);var P=this.HTMLgetScale(),J=this.HTMLgetMu(M);var y=d.createStack(M),n,w=[];var x=d.createBox(y);if(this.data[this.base]){w.push(this.data[this.base].toHTML(x));if(F!=null){this.data[this.base].HTMLstretchV(x,K,F)}else{if(K!=null){this.data[this.base].HTMLstretchH(x,K)}}}else{x.bbox=this.HTMLzeroBBox()}var N=d.TeX.x_height*P,E=d.TeX.scriptspace*P*0.75;var m,z;if(this.HTMLnotEmpty(this.data[this.sup])){m=d.createBox(y);w.push(this.data[this.sup].toHTML(m))}if(this.HTMLnotEmpty(this.data[this.sub])){z=d.createBox(y);w.push(this.data[this.sub].toHTML(z))}d.MeasureSpans(w);if(m){m.bbox.w+=E;m.bbox.rw=Math.max(m.bbox.w,m.bbox.rw)}if(z){z.bbox.w+=E;z.bbox.rw=Math.max(z.bbox.w,z.bbox.rw)}d.placeBox(x,0,0);var o=P;if(m){o=this.data[this.sup].HTMLgetScale()}else{if(z){o=this.data[this.sub].HTMLgetScale()}}var H=d.TeX.sup_drop*o,G=d.TeX.sub_drop*o;var B=x.bbox.h-H,A=x.bbox.d+G,O=0,I;if(x.bbox.ic){x.bbox.w-=x.bbox.ic;O=1.3*x.bbox.ic+0.05}if(this.data[this.base]&&K==null&&F==null&&(this.data[this.base].type==="mi"||this.data[this.base].type==="mo")){if(this.data[this.base].data.join("").length===1&&w[0].scale===1&&!this.data[this.base].Get("largeop")){B=A=0}}var L=this.getValues("subscriptshift","superscriptshift");L.subscriptshift=(L.subscriptshift===""?0:d.length2em(L.subscriptshift,J));L.superscriptshift=(L.superscriptshift===""?0:d.length2em(L.superscriptshift,J));if(!m){if(z){A=Math.max(A,d.TeX.sub1*P,z.bbox.h-(4/5)*N,L.subscriptshift);d.placeBox(z,x.bbox.w,-A,z.bbox)}}else{if(!z){n=this.getValues("displaystyle","texprimestyle");I=d.TeX[(n.displaystyle?"sup1":(n.texprimestyle?"sup3":"sup2"))];B=Math.max(B,I*P,m.bbox.d+(1/4)*N,L.superscriptshift);d.placeBox(m,x.bbox.w+O,B,m.bbox)}else{A=Math.max(A,d.TeX.sub2*P);var C=d.TeX.rule_thickness*P;if((B-m.bbox.d)-(z.bbox.h-A)<3*C){A=3*C-B+m.bbox.d+z.bbox.h;H=(4/5)*N-(B-m.bbox.d);if(H>0){B+=H;A-=H}}d.placeBox(m,x.bbox.w+O,Math.max(B,L.superscriptshift));d.placeBox(z,x.bbox.w,-Math.max(A,L.subscriptshift))}}this.HTMLhandleSpace(M);this.HTMLhandleColor(M);return M},HTMLstretchH:i.mbase.HTMLstretchH,HTMLstretchV:i.mbase.HTMLstretchV});i.mmultiscripts.Augment({toHTML:i.mbase.HTMLautoload});i.mtable.Augment({toHTML:i.mbase.HTMLautoload});i["annotation-xml"].Augment({toHTML:i.mbase.HTMLautoload});i.annotation.Augment({toHTML:function(m){return this.HTMLcreateSpan(m)}});i.math.Augment({toHTML:function(E,B,t){var u,w,x,r,m=E;if(!t||t===d.PHASE.I){var C=d.addElement(E,"nobr",{isMathJax:true});E=this.HTMLcreateSpan(C);var n=this.Get("alttext");if(n&&!E.getAttribute("aria-label")){E.setAttribute("aria-label",n)}u=d.createStack(E);w=d.createBox(u);u.style.fontSize=C.parentNode.style.fontSize;C.parentNode.style.fontSize="";if(this.data[0]!=null){i.mbase.prototype.displayAlign=b.config.displayAlign;i.mbase.prototype.displayIndent=b.config.displayIndent;if(String(b.config.displayIndent).match(/^0($|[a-z%])/i)){i.mbase.prototype.displayIndent="0"}x=this.data[0].toHTML(w);x.bbox.exactW=false}}else{E=E.firstChild.firstChild;if(this.href){E=E.firstChild}u=E.firstChild;if(u.style.position!=="relative"){u=u.nextSibling}w=u.firstChild;x=w.firstChild}r=((!t||t===d.PHASE.II)?d.Measured(x,w):x);if(!t||t===d.PHASE.III){d.placeBox(w,0,0);var q=r.bbox.w;q=Math.abs(q)<0.006?0:Math.max(0,Math.round(q*this.em)+0.25);E.style.width=d.EmRounded(q/d.outerEm);E.style.display="inline-block";var A=1/d.em,G=d.em/d.outerEm;d.em/=G;E.bbox.h*=G;E.bbox.d*=G;E.bbox.w*=G;E.bbox.lw*=G;E.bbox.rw*=G;if(E.bbox.H){E.bbox.H*=G}if(E.bbox.D){E.bbox.D*=G}if(r&&r.bbox.width!=null){E.style.minWidth=(r.bbox.minWidth||E.style.width);E.style.width=r.bbox.width;w.style.width=u.style.width="100%";m.className+=" MathJax_FullWidth"}var D=this.HTMLhandleColor(E);if(r){d.createRule(E,(r.bbox.h+A)*G,(r.bbox.d+A)*G,0)}if(!this.isMultiline&&this.Get("display")==="block"&&E.bbox.width==null){var o=this.getValues("indentalignfirst","indentshiftfirst","indentalign","indentshift");if(o.indentalignfirst!==i.INDENTALIGN.INDENTALIGN){o.indentalign=o.indentalignfirst}if(o.indentalign===i.INDENTALIGN.AUTO){o.indentalign=this.displayAlign}if(o.indentshiftfirst!==i.INDENTSHIFT.INDENTSHIFT){o.indentshift=o.indentshiftfirst}if(o.indentshift==="auto"){o.indentshift="0"}var F=d.length2em(o.indentshift,1,d.scale*d.cwidth);if(this.displayIndent!=="0"){var y=d.length2em(this.displayIndent,1,d.scale*d.cwidth);F+=(o.indentalign===i.INDENTALIGN.RIGHT?-y:y)}m.style.textAlign=B.style.textAlign=o.indentalign;if(F){b.Insert(E.style,({left:{marginLeft:d.Em(F)},right:{marginRight:d.Em(-F)},center:{marginLeft:d.Em(F),marginRight:d.Em(-F)}})[o.indentalign]);if(D){var v=parseFloat(D.style.marginLeft||"0")+F,s=parseFloat(D.style.marginRight||"0")-F;D.style.marginLeft=d.Em(v);D.style.marginRight=d.Em(s+(o.indentalign==="right"?E.bbox.w+F-E.bbox.w:0));if(d.msieColorBug&&o.indentalign==="right"){if(parseFloat(D.style.marginLeft)>0){var z=MathJax.HTML.addElement(D.parentNode,"span");z.style.marginLeft=d.Em(s+Math.min(0,E.bbox.w+F));D.nextSibling.style.marginRight="0em"}D.nextSibling.style.marginLeft="0em";D.style.marginRight=D.style.marginLeft="0em"}}}}}return E},HTMLspanElement:i.mbase.prototype.HTMLspanElement});i.TeXAtom.Augment({toHTML:function(q,o,s){q=this.HTMLcreateSpan(q);if(this.data[0]!=null){if(this.texClass===i.TEXCLASS.VCENTER){var m=d.createStack(q);var r=d.createBox(m);var t=this.data[0].toHTML(r);if(s!=null){d.Remeasured(this.data[0].HTMLstretchV(r,o,s),r)}else{if(o!=null){d.Remeasured(this.data[0].HTMLstretchH(r,o),r)}else{d.Measured(t,r)}}var n=d.TeX.axis_height*this.HTMLgetScale();d.placeBox(r,0,n-(r.bbox.h+r.bbox.d)/2+r.bbox.d)}else{var p=this.data[0].toHTML(q,o,s);if(s!=null){p=this.data[0].HTMLstretchV(r,o,s)}else{if(o!=null){p=this.data[0].HTMLstretchH(r,o)}}q.bbox=p.bbox}}this.HTMLhandleSpace(q);this.HTMLhandleColor(q);return q},HTMLstretchH:i.mbase.HTMLstretchH,HTMLstretchV:i.mbase.HTMLstretchV});b.Register.StartupHook("onLoad",function(){setTimeout(MathJax.Callback(["loadComplete",d,"jax.js"]),0)})});b.Register.StartupHook("End Config",function(){b.Browser.Select({MSIE:function(m){var q=(document.documentMode||0);var p=m.versionAtLeast("7.0");var o=m.versionAtLeast("8.0")&&q>7;var n=(document.compatMode==="BackCompat");if(q<9){d.config.styles[".MathJax .MathJax_HitBox"]["background-color"]="white";d.config.styles[".MathJax .MathJax_HitBox"].opacity=0;d.config.styles[".MathJax .MathJax_HitBox"].filter="alpha(opacity=0)"}d.Augment({PaddingWidthBug:true,msieAccentBug:true,msieColorBug:(q<8),msieColorPositionBug:true,msieRelativeWidthBug:n,msieDisappearingBug:(q>=8),msieMarginScaleBug:(q<8),msiePaddingWidthBug:true,msieBorderWidthBug:n,msieFrameSizeBug:(q<=8),msieInlineBlockAlignBug:(!o||n),msiePlaceBoxBug:(o&&!n),msieClipRectBug:!o,msieNegativeSpaceBug:n,msieRuleBug:(q<7),cloneNodeBug:(o&&m.version==="8.0"),msieItalicWidthBug:true,initialSkipBug:(q<8),msieNegativeBBoxBug:(q>=8),msieIE6:!p,msieItalicWidthBug:true,FontFaceBug:(q<9),msieFontCSSBug:m.isIE9,allowWebFonts:(q>=9?"woff":"eot")})},Firefox:function(n){var o=false;if(n.versionAtLeast("3.5")){var m=String(document.location).replace(/[^\/]*$/,"");if(document.location.protocol!=="file:"||b.config.root.match(/^https?:\/\//)||(b.config.root+"/").substr(0,m.length)===m){o="otf"}}d.Augment({ffVerticalAlignBug:!n.versionAtLeast("20.0"),AccentBug:true,allowWebFonts:o,ffFontOptimizationBug:true})},Safari:function(r){var p=r.versionAtLeast("3.0");var o=r.versionAtLeast("3.1");var m=navigator.appVersion.match(/ Safari\/\d/)&&navigator.appVersion.match(/ Version\/\d/)&&navigator.vendor.match(/Apple/);var n=(navigator.appVersion.match(/ Android (\d+)\.(\d+)/));var s=(o&&r.isMobile&&((navigator.platform.match(/iPad|iPod|iPhone/)&&!r.versionAtLeast("5.0"))||(n!=null&&(n[1]<2||(n[1]==2&&n[2]<2)))));d.Augment({config:{styles:{".MathJax img, .MathJax nobr, .MathJax a":{"max-width":"5000em","max-height":"5000em"}}},Em:((r.webkit||0)>=538?d.EmRounded:d.Em),rfuzz:0.011,AccentBug:true,AdjustSurd:true,negativeBBoxes:true,safariNegativeSpaceBug:true,safariVerticalAlignBug:!o,safariTextNodeBug:!p,forceReflow:true,FontFaceBug:true,allowWebFonts:(o&&!s?"otf":false)});if(m){d.Augment({webFontDefault:(r.isMobile?"sans-serif":"serif")})}if(r.isPC){d.Augment({adjustAvailableFonts:d.removeSTIXfonts,checkWebFontsTwice:true})}if(s){var q=b.config["HTML-CSS"];if(q){q.availableFonts=[];q.preferredFont=null}else{b.config["HTML-CSS"]={availableFonts:[],preferredFont:null}}}},Chrome:function(m){d.Augment({Em:d.EmRounded,cloneNodeBug:true,rfuzz:-0.02,AccentBug:true,AdjustSurd:true,FontFaceBug:m.versionAtLeast("32.0"),negativeBBoxes:true,safariNegativeSpaceBug:true,safariWebFontSerif:[""],forceReflow:true,allowWebFonts:(m.versionAtLeast("4.0")?"otf":"svg")})},Opera:function(m){m.isMini=(navigator.appVersion.match("Opera Mini")!=null);d.config.styles[".MathJax .merror"]["vertical-align"]=null;d.config.styles[".MathJax span"]["z-index"]=0;d.Augment({operaHeightBug:true,operaVerticalAlignBug:true,operaFontSizeBug:m.versionAtLeast("10.61"),initialSkipBug:true,FontFaceBug:true,PaddingWidthBug:true,allowWebFonts:(m.versionAtLeast("10.0")&&!m.isMini?"otf":false),adjustAvailableFonts:d.removeSTIXfonts})},Konqueror:function(m){d.Augment({konquerorVerticalAlignBug:true})}})});MathJax.Hub.Register.StartupHook("End Cookie",function(){if(b.config.menuSettings.zoom!=="None"){j.Require("[MathJax]/extensions/MathZoom.js")}})})(MathJax.Ajax,MathJax.Hub,MathJax.OutputJax["HTML-CSS"]);
MathJax.Hub.Register.StartupHook("HTML-CSS Jax Ready",function(){var c="2.7.0";var a=MathJax.ElementJax.mml,b=MathJax.OutputJax["HTML-CSS"];a.mtable.Augment({toHTML:function(r){r=this.HTMLcreateSpan(r);if(this.data.length===0){return r}var I=this.getValues("columnalign","rowalign","columnspacing","rowspacing","columnwidth","equalcolumns","equalrows","columnlines","rowlines","frame","framespacing","align","useHeight","width","side","minlabelspacing");var aM=I.width.match(/%$/);var ay=b.createStack(r);var aJ=this.HTMLgetScale(),aB=this.HTMLgetMu(r),aC=-1;var aq=[],au=[],aj=[],aw=[],av=[],ae,ad,ap=-1,ac,ao,X,aH,Q,aE,aR=[],aW;var G=b.FONTDATA.lineH*aJ*I.useHeight,N=b.FONTDATA.lineD*aJ*I.useHeight;for(ae=0,ac=this.data.length;ae<ac;ae++){aH=this.data[ae];X=(aH.type==="mlabeledtr"?aC:0);aw[ae]=[];aq[ae]=G;au[ae]=N;for(ad=X,ao=aH.data.length+X;ad<ao;ad++){if(aj[ad]==null){if(ad>ap){ap=ad}av[ad]=b.createStack(b.createBox(ay));aj[ad]=-b.BIGDIMEN}aw[ae][ad]=b.createBox(av[ad]);aR.push(aH.data[ad-X].toHTML(aw[ae][ad]))}}b.MeasureSpans(aR);for(ae=0,ac=this.data.length;ae<ac;ae++){aH=this.data[ae];X=(aH.type==="mlabeledtr"?aC:0);for(ad=X,ao=aH.data.length+X;ad<ao;ad++){Q=aH.data[ad-X];if(Q.isMultiline){aw[ae][ad].style.width="100%"}if(Q.isEmbellished()){aE=Q.CoreMO();var aV=aE.Get("minsize",true);if(aV){var aO=aE.HTMLspanElement().bbox;if(aE.HTMLcanStretch("Vertical")){aW=aO.h+aO.d;if(aW){aV=b.length2em(aV,aB,aW);if(aV*aO.h/aW>aq[ae]){aq[ae]=aV*aO.h/aW}if(aV*aO.d/aW>au[ae]){au[ae]=aV*aO.d/aW}}}else{if(aE.HTMLcanStretch("Horizontal")){aV=b.length2em(aV,aB,aO.w);if(aV>aj[ad]){aj[ad]=aV}}}}}if(aw[ae][ad].bbox.h>aq[ae]){aq[ae]=aw[ae][ad].bbox.h}if(aw[ae][ad].bbox.d>au[ae]){au[ae]=aw[ae][ad].bbox.d}if(aw[ae][ad].bbox.w>aj[ad]){aj[ad]=aw[ae][ad].bbox.w}}}var aG=MathJax.Hub.SplitList;var aA=aG(I.columnspacing),aT=aG(I.rowspacing),e=aG(I.columnalign),B=aG(I.rowalign),d=aG(I.columnlines),w=aG(I.rowlines),aP=aG(I.columnwidth),U=[];for(ae=0,ac=aA.length;ae<ac;ae++){aA[ae]=b.length2em(aA[ae],aB)}for(ae=0,ac=aT.length;ae<ac;ae++){aT[ae]=b.length2em(aT[ae],aB)}while(aA.length<ap){aA.push(aA[aA.length-1])}while(e.length<=ap){e.push(e[e.length-1])}while(d.length<ap){d.push(d[d.length-1])}while(aP.length<=ap){aP.push(aP[aP.length-1])}while(aT.length<aw.length){aT.push(aT[aT.length-1])}while(B.length<=aw.length){B.push(B[B.length-1])}while(w.length<aw.length){w.push(w[w.length-1])}if(av[aC]){e[aC]=(I.side.substr(0,1)==="l"?"left":"right");aA[aC]=-aj[aC]}for(ae=0,ac=aw.length;ae<ac;ae++){aH=this.data[ae];U[ae]=[];if(aH.rowalign){B[ae]=aH.rowalign}if(aH.columnalign){U[ae]=aG(aH.columnalign);while(U[ae].length<=ap){U[ae].push(U[ae][U[ae].length-1])}}}if(I.equalrows){var aF=Math.max.apply(Math,aq),V=Math.max.apply(Math,au);for(ae=0,ac=aw.length;ae<ac;ae++){X=((aF+V)-(aq[ae]+au[ae]))/2;aq[ae]+=X;au[ae]+=X}}aW=aq[0]+au[aw.length-1];for(ae=0,ac=aw.length-1;ae<ac;ae++){aW+=Math.max(0,au[ae]+aq[ae+1]+aT[ae])}var aL=0,aK=0,aZ,g=aW;if(I.frame!=="none"||(I.columnlines+I.rowlines).match(/solid|dashed/)){var v=aG(I.framespacing);if(v.length!=2){v=aG(this.defaults.framespacing)}aL=b.length2em(v[0],aB);aK=b.length2em(v[1],aB);g=aW+2*aK}var ai,aY,aa="";if(typeof(I.align)!=="string"){I.align=String(I.align)}if(I.align.match(/(top|bottom|center|baseline|axis)( +(-?\d+))?/)){aa=RegExp.$3||"";I.align=RegExp.$1}else{I.align=this.defaults.align}if(aa!==""){aa=parseInt(aa);if(aa<0){aa=aw.length+1+aa}if(aa<1){aa=1}else{if(aa>aw.length){aa=aw.length}}ai=0;aY=-(aW+aK)+aq[0];for(ae=0,ac=aa-1;ae<ac;ae++){var L=Math.max(0,au[ae]+aq[ae+1]+aT[ae]);ai+=L;aY+=L}}else{ai=({top:-(aq[0]+aK),bottom:aW+aK-aq[0],center:aW/2-aq[0],baseline:aW/2-aq[0],axis:aW/2+b.TeX.axis_height*aJ-aq[0]})[I.align];aY=({top:-(aW+2*aK),bottom:0,center:-(aW/2+aK),baseline:-(aW/2+aK),axis:b.TeX.axis_height*aJ-aW/2-aK})[I.align]}var ab,af=0,z=0,K=0,Z=0,ag=0,am=[],at=[],R=1;if(I.equalcolumns&&I.width!=="auto"){if(aM){ab=(100/(ap+1)).toFixed(2).replace(/\.?0+$/,"")+"%";for(ae=0,ac=Math.min(ap+1,aP.length);ae<ac;ae++){aP[ae]=ab}ab=0;af=1;ag=ap+1;for(ae=0,ac=Math.min(ap+1,aA.length);ae<ac;ae++){ab+=aA[ae]}}else{ab=b.length2em(I.width,aB);for(ae=0,ac=Math.min(ap,aA.length);ae<ac;ae++){ab-=aA[ae]}ab/=ap;for(ae=0,ac=Math.min(ap+1,aP.length);ae<ac;ae++){aj[ae]=ab}}}else{for(ae=0,ac=Math.min(ap+1,aP.length);ae<ac;ae++){if(aP[ae]==="auto"){z+=aj[ae]}else{if(aP[ae]==="fit"){at[ag]=ae;ag++;z+=aj[ae]}else{if(aP[ae].match(/%$/)){am[Z]=ae;Z++;K+=aj[ae];af+=b.length2em(aP[ae],aB,1)}else{aj[ae]=b.length2em(aP[ae],aB);z+=aj[ae]}}}}if(aM){ab=0;for(ae=0,ac=Math.min(ap,aA.length);ae<ac;ae++){ab+=aA[ae]}if(af>0.98){R=0.98/af;af=0.98}}else{if(I.width==="auto"){if(af>0.98){R=K/(z+K);ab=z+K}else{ab=z/(1-af)}}else{ab=b.length2em(I.width,aB);for(ae=0,ac=Math.min(ap,aA.length);ae<ac;ae++){ab-=aA[ae]}}for(ae=0,ac=am.length;ae<ac;ae++){aj[am[ae]]=b.length2em(aP[am[ae]],aB,ab*R);z+=aj[am[ae]]}if(Math.abs(ab-z)>0.01){if(ag&&ab>z){ab=(ab-z)/ag;for(ae=0,ac=at.length;ae<ac;ae++){aj[at[ae]]+=ab}}else{ab=ab/z;for(ad=0;ad<=ap;ad++){aj[ad]*=ab}}}if(I.equalcolumns){var O=Math.max.apply(Math,aj);for(ad=0;ad<=ap;ad++){aj[ad]=O}}}}var S=ai,o,q,aU;X=(av[aC]?aC:0);for(ad=X;ad<=ap;ad++){for(ae=0,ac=aw.length;ae<ac;ae++){if(aw[ae][ad]){X=(this.data[ae].type==="mlabeledtr"?aC:0);Q=this.data[ae].data[ad-X];if(Q.HTMLcanStretch("Horizontal")){aw[ae][ad].bbox=Q.HTMLstretchH(av[ad],aj[ad]).bbox}else{if(Q.HTMLcanStretch("Vertical")){aE=Q.CoreMO();var aN=aE.symmetric;aE.symmetric=false;aw[ae][ad].bbox=Q.HTMLstretchV(av[ad],aq[ae],au[ae]).bbox;aw[ae][ad].HH=null;if(aw[ae][ad].bbox.h>aq[ae]){aw[ae][ad].bbox.H=aw[ae][ad].bbox.h;aw[ae][ad].bbox.h=aq[ae]}if(aw[ae][ad].bbox.d>au[ae]){aw[ae][ad].bbox.D=aw[ae][ad].bbox.d;aw[ae][ad].bbox.d=au[ae]}aE.symmetric=aN}}aU=Q.rowalign||this.data[ae].rowalign||B[ae];o=({top:aq[ae]-aw[ae][ad].bbox.h,bottom:aw[ae][ad].bbox.d-au[ae],center:((aq[ae]-au[ae])-(aw[ae][ad].bbox.h-aw[ae][ad].bbox.d))/2,baseline:0,axis:0})[aU]||0;aU=(Q.columnalign||U[ae][ad]||e[ad]);b.alignBox(aw[ae][ad],aU,S+o)}if(ae<aw.length-1){S-=Math.max(0,au[ae]+aq[ae+1]+aT[ae])}}S=ai}if(aM){var E=b.createBox(ay);E.style.left=E.style.top=0;E.style.right=b.Em(ab+2*aL);E.style.display="inline-block";E.style.height="0px";if(b.msieRelativeWidthBug){E=b.createBox(E);E.style.position="relative";E.style.height="1em";E.style.width="100%";E.bbox=ay.bbox}var aS=0,a0=aL,k,l;if(ag){k=100*(1-af)/ag,l=z/ag}else{k=100*(1-af)/(ap+1);l=z/(ap+1)}for(ad=0;ad<=ap;ad++){b.placeBox(av[ad].parentNode,0,0);av[ad].style.position="relative";av[ad].style.left=b.Em(a0);av[ad].style.width="100%";av[ad].parentNode.parentNode.removeChild(av[ad].parentNode);var al=b.createBox(E);b.addBox(al,av[ad]);av[ad]=al;var h=al.style;h.display="inline-block";h.left=aS+"%";if(aP[ad].match(/%$/)){var t=parseFloat(aP[ad])*R;if(ag===0){h.width=(k+t)+"%";aS+=k+t;al=b.createBox(al);b.addBox(al,av[ad].firstChild);al.style.left=0;al.style.right=b.Em(l);a0-=l}else{h.width=t+"%";aS+=t}}else{if(aP[ad]==="fit"||ag===0){h.width=k+"%";al=b.createBox(al);b.addBox(al,av[ad].firstChild);al.style.left=0;al.style.right=b.Em(l-aj[ad]);a0+=aj[ad]-l;aS+=k}else{h.width=b.Em(aj[ad]);a0+=aj[ad]}}if(b.msieRelativeWidthBug){b.addText(al.firstChild,b.NBSP);al.firstChild.style.position="relative"}a0+=aA[ad];if(d[ad]!=="none"&&ad<ap&&ad!==aC){q=b.createBox(E);q.style.left=aS+"%";q=b.createRule(q,g,0,1.25/b.em);q.style.position="absolute";q.bbox={h:g,d:0,w:0,rw:1.25/b.em,lw:0};q.parentNode.bbox=ay.bbox;b.placeBox(q,a0-aA[ad]/2,aY,true);q.style.borderStyle=d[ad]}}}else{var T=aL;for(ad=0;ad<=ap;ad++){if(!av[ad].bbox.width){b.setStackWidth(av[ad],aj[ad])}if(aP[ad]!=="auto"&&aP[ad]!=="fit"){av[ad].bbox.width=aj[ad];av[ad].bbox.isFixed=true}b.placeBox(av[ad].parentNode,T,0);T+=aj[ad]+aA[ad];if(d[ad]!=="none"&&ad<ap&&ad!==aC){q=b.createRule(ay,g,0,1.25/b.em);b.addBox(ay,q);q.bbox={h:g,d:0,w:0,rw:1.25/b.em,lw:0};b.placeBox(q,T-aA[ad]/2,aY,true);q.style.borderStyle=d[ad]}}}ay.bbox.d=-aY;ay.bbox.h=g+aY;b.setStackWidth(ay,ay.bbox.w+aL);aZ=ay.bbox.w;var ah;if(I.frame!=="none"){ah=b.createFrame(ay,g,0,aZ,1.25/b.em,I.frame);b.addBox(ay,ah);b.placeBox(ah,0,aY,true);if(aM){ah.style.width="100%"}}S=ai;for(ae=0,ac=aw.length-1;ae<ac;ae++){o=Math.max(0,au[ae]+aq[ae+1]+aT[ae]);if(w[ae]!==a.LINES.NONE&&w[ae]!==""){q=b.createRule(ay,1.25/b.em,0,aZ);b.addBox(ay,q);q.bbox={h:1.25/b.em,d:0,w:aZ,rw:aZ,lw:0};b.placeBox(q,0,S-au[ae]-(o-au[ae]-aq[ae+1])/2,true);if(w[ae]===a.LINES.DASHED){q.style.borderTopStyle="dashed"}if(aM){q.style.width="100%"}}S-=o}if(aM){r.bbox.width=I.width;ay.style.width="100%"}if(av[aC]){var ax=ay.bbox.w;var ar=this.getValues("indentalignfirst","indentshiftfirst","indentalign","indentshift");if(ar.indentalignfirst!==a.INDENTALIGN.INDENTALIGN){ar.indentalign=ar.indentalignfirst}if(ar.indentalign===a.INDENTALIGN.AUTO){ar.indentalign=this.displayAlign}if(ar.indentshiftfirst!==a.INDENTSHIFT.INDENTSHIFT){ar.indentshift=ar.indentshiftfirst}if(ar.indentshift==="auto"){ar.indentshift="0"}var an=b.length2em(ar.indentshift,aB,b.cwidth);var aD=b.length2em(I.minlabelspacing,aB,b.cwidth);var aX=aD+av[aC].bbox.w,az=0,ak=ax;var aI=b.length2em(this.displayIndent,aB,b.cwidth);X=(e[aC]===a.INDENTALIGN.RIGHT?-1:1);if(ar.indentalign===a.INDENTALIGN.CENTER){ak+=2*(aX-X*(an+aI));an+=aI}else{if(e[aC]===ar.indentalign){if(aI<0){az=X*aI;aI=0}an+=X*aI;if(aX>X*an){an=X*aX}an+=az;ak+=X*an}else{ak+=aX-X*an+aI;an-=X*aI}}var aQ=b.createStack(r,false,"100%");b.addBox(aQ,ay);b.alignBox(ay,ar.indentalign,0,an);av[aC].parentNode.parentNode.removeChild(av[aC].parentNode);b.addBox(aQ,av[aC]);b.alignBox(av[aC],e[aC],0);if(b.msieRelativeWidthBug){ay.style.top=av[aC].style.top=""}if(aM){ay.style.width=I.width;r.bbox.width="100%"}av[aC].style[X===1?"marginLeft":"marginRight"]=b.Em(X*az);r.bbox.tw=ak;r.style.minWidth=r.bbox.minWidth=b.Em(ak);aQ.style.minWidth=aQ.bbox.minWidth=b.Em(ak/aJ)}if(!aM){this.HTMLhandleSpace(r)}var u=this.HTMLhandleColor(r);if(u&&aM){if(!ah){ah=b.createFrame(ay,g,0,aZ,0,"none");b.addBox(ay,ah);b.placeBox(ah,0,aY,true);ah.style.width="100%"}ah.style.backgroundColor=u.style.backgroundColor;ah.parentNode.insertBefore(ah,ah.parentNode.firstChild);u.parentNode.removeChild(u)}return r},HTMLhandleSpace:function(d){d.bbox.keepPadding=true;d.bbox.exact=true;if(!this.hasFrame&&d.bbox.width==null){d.style.paddingLeft=d.style.paddingRight=b.Em(1/6)}this.SUPER(arguments).HTMLhandleSpace.call(this,d)}});a.mtd.Augment({toHTML:function(e,d,g){e=this.HTMLcreateSpan(e);if(this.data[0]){var f=this.data[0].toHTML(e);if(g!=null){f=this.data[0].HTMLstretchV(e,d,g)}else{if(d!=null){f=this.data[0].HTMLstretchH(e,d)}}e.bbox=f.bbox}this.HTMLhandleSpace(e);this.HTMLhandleColor(e);return e},HTMLstretchH:a.mbase.HTMLstretchH,HTMLstretchV:a.mbase.HTMLstretchV});MathJax.Hub.Startup.signal.Post("HTML-CSS mtable Ready");MathJax.Ajax.loadComplete(b.autoloadDir+"/mtable.js")});
(function(i,b,e,g){var h;var j,a,d;var f="'Times New Roman',Times,STIXGeneral,serif";var m={".MJXp-script":{"font-size":".8em"},".MJXp-right":{"-webkit-transform-origin":"right","-moz-transform-origin":"right","-ms-transform-origin":"right","-o-transform-origin":"right","transform-origin":"right"},".MJXp-bold":{"font-weight":"bold"},".MJXp-italic":{"font-style":"italic"},".MJXp-scr":{"font-family":"MathJax_Script,"+f},".MJXp-frak":{"font-family":"MathJax_Fraktur,"+f},".MJXp-sf":{"font-family":"MathJax_SansSerif,"+f},".MJXp-cal":{"font-family":"MathJax_Caligraphic,"+f},".MJXp-mono":{"font-family":"MathJax_Typewriter,"+f},".MJXp-largeop":{"font-size":"150%"},".MJXp-largeop.MJXp-int":{"vertical-align":"-.2em"},".MJXp-math":{display:"inline-block","line-height":"1.2","text-indent":"0","font-family":f,"white-space":"nowrap","border-collapse":"collapse"},".MJXp-display":{display:"block","text-align":"center",margin:"1em 0"},".MJXp-math span":{display:"inline-block"},".MJXp-box":{display:"block!important","text-align":"center"},".MJXp-box:after":{content:'" "'},".MJXp-rule":{display:"block!important","margin-top":".1em"},".MJXp-char":{display:"block!important"},".MJXp-mo":{margin:"0 .15em"},".MJXp-mfrac":{margin:"0 .125em","vertical-align":".25em"},".MJXp-denom":{display:"inline-table!important",width:"100%"},".MJXp-denom > *":{display:"table-row!important"},".MJXp-surd":{"vertical-align":"top"},".MJXp-surd > *":{display:"block!important"},".MJXp-script-box > * ":{display:"table!important",height:"50%"},".MJXp-script-box > * > *":{display:"table-cell!important","vertical-align":"top"},".MJXp-script-box > *:last-child > *":{"vertical-align":"bottom"},".MJXp-script-box > * > * > *":{display:"block!important"},".MJXp-mphantom":{visibility:"hidden"},".MJXp-munderover":{display:"inline-table!important"},".MJXp-over":{display:"inline-block!important","text-align":"center"},".MJXp-over > *":{display:"block!important"},".MJXp-munderover > *":{display:"table-row!important"},".MJXp-mtable":{"vertical-align":".25em",margin:"0 .125em"},".MJXp-mtable > *":{display:"inline-table!important","vertical-align":"middle"},".MJXp-mtr":{display:"table-row!important"},".MJXp-mtd":{display:"table-cell!important","text-align":"center",padding:".5em 0 0 .5em"},".MJXp-mtr > .MJXp-mtd:first-child":{"padding-left":0},".MJXp-mtr:first-child > .MJXp-mtd":{"padding-top":0},".MJXp-mlabeledtr":{display:"table-row!important"},".MJXp-mlabeledtr > .MJXp-mtd:first-child":{"padding-left":0},".MJXp-mlabeledtr:first-child > .MJXp-mtd":{"padding-top":0},".MJXp-merror":{"background-color":"#FFFF88",color:"#CC0000",border:"1px solid #CC0000",padding:"1px 3px","font-style":"normal","font-size":"90%"}};(function(){for(var n=0;n<10;n++){var o="scaleX(."+n+")";m[".MJXp-scale"+n]={"-webkit-transform":o,"-moz-transform":o,"-ms-transform":o,"-o-transform":o,transform:o}}})();var k=1000000;var c="V",l="H";g.Augment({settings:b.config.menuSettings,config:{styles:m},hideProcessedMath:false,maxStretchyParts:1000,Config:function(){if(!this.require){this.require=[]}this.SUPER(arguments).Config.call(this);var n=this.settings;if(n.scale){this.config.scale=n.scale}this.require.push(MathJax.OutputJax.extensionDir+"/MathEvents.js")},Startup:function(){j=MathJax.Extension.MathEvents.Event;a=MathJax.Extension.MathEvents.Touch;d=MathJax.Extension.MathEvents.Hover;this.ContextMenu=j.ContextMenu;this.Mousedown=j.AltContextMenu;this.Mouseover=d.Mouseover;this.Mouseout=d.Mouseout;this.Mousemove=d.Mousemove;var n=e.addElement(document.body,"div",{style:{width:"5in"}});this.pxPerInch=n.offsetWidth/5;n.parentNode.removeChild(n);return i.Styles(this.config.styles,["InitializePHTML",this])},InitializePHTML:function(){},preTranslate:function(p){var s=p.jax[this.id],t,q=s.length,u,r,v,o,n;for(t=0;t<q;t++){u=s[t];if(!u.parentNode){continue}r=u.previousSibling;if(r&&String(r.className).match(/^MathJax(_PHTML)?(_Display)?( MathJax_Process(ing|ed))?$/)){r.parentNode.removeChild(r)}n=u.MathJax.elementJax;if(!n){continue}n.PHTML={display:(n.root.Get("display")==="block")};v=o=e.Element("span",{className:"MathJax_PHTML",id:n.inputID+"-Frame",isMathJax:true,jaxID:this.id,oncontextmenu:j.Menu,onmousedown:j.Mousedown,onmouseover:j.Mouseover,onmouseout:j.Mouseout,onmousemove:j.Mousemove,onclick:j.Click,ondblclick:j.DblClick,onkeydown:j.Keydown,tabIndex:b.getTabOrder(n)});if(b.Browser.noContextMenu){v.ontouchstart=a.start;v.ontouchend=a.end}if(n.PHTML.display){o=e.Element("div",{className:"MathJax_PHTML_Display"});o.appendChild(v)}o.className+=" MathJax_Processing";u.parentNode.insertBefore(o,u)}},Translate:function(o,s){if(!o.parentNode){return}var n=o.MathJax.elementJax,r=n.root,p=document.getElementById(n.inputID+"-Frame"),t=(n.PHTML.display?p.parentNode:p);this.initPHTML(r,p);try{r.toPreviewHTML(p)}catch(q){if(q.restart){while(p.firstChild){p.removeChild(p.firstChild)}}throw q}t.className=t.className.split(/ /)[0];if(this.hideProcessedMath){t.className+=" MathJax_Processed";if(o.MathJax.preview){n.PHTML.preview=o.MathJax.preview;delete o.MathJax.preview}}},postTranslate:function(s){var o=s.jax[this.id];if(!this.hideProcessedMath){return}for(var q=0,n=o.length;q<n;q++){var p=o[q];if(p&&p.MathJax.elementJax){p.previousSibling.className=p.previousSibling.className.split(/ /)[0];var r=p.MathJax.elementJax.PHTML;if(r.preview){r.preview.innerHTML="";r.preview.style.display="none";p.MathJax.preview=r.preview;delete r.preview}}}},getJaxFromMath:function(n){if(n.parentNode.className.match(/MathJax_PHTML_Display/)){n=n.parentNode}do{n=n.nextSibling}while(n&&n.nodeName.toLowerCase()!=="script");return b.getJaxFor(n)},getHoverSpan:function(n,o){return n.root.PHTMLspanElement()},getHoverBBox:function(n,q,r){var s=n.root.PHTML,p=n.PHTML.outerEm;var o={w:s.w*p,h:s.h*p,d:s.d*p};if(s.width){o.width=s.width}return o},Zoom:function(o,u,s,n,r){u.className="MathJax";this.idPostfix="-zoom";o.root.toPHTML(u,u);this.idPostfix="";u.style.position="absolute";if(!width){s.style.position="absolute"}var t=u.offsetWidth,q=u.offsetHeight,v=s.offsetHeight,p=s.offsetWidth;if(p===0){p=s.parentNode.offsetWidth}u.style.position=s.style.position="";return{Y:-j.getBBox(u).h,mW:p,mH:v,zW:t,zH:q}},initPHTML:function(o,n){},Remove:function(n){var o=document.getElementById(n.inputID+"-Frame");if(o){if(n.PHTML.display){o=o.parentNode}o.parentNode.removeChild(o)}delete n.PHTML},ID:0,idPostfix:"",GetID:function(){this.ID++;return this.ID},VARIANT:{bold:"MJXp-bold",italic:"MJXp-italic","bold-italic":"MJXp-bold MJXp-italic",script:"MJXp-scr","bold-script":"MJXp-scr MJXp-bold",fraktur:"MJXp-frak","bold-fraktur":"MJXp-frak MJXp-bold",monospace:"MJXp-mono","sans-serif":"MJXp-sf","-tex-caligraphic":"MJXp-cal"},MATHSPACE:{veryverythinmathspace:1/18,verythinmathspace:2/18,thinmathspace:3/18,mediummathspace:4/18,thickmathspace:5/18,verythickmathspace:6/18,veryverythickmathspace:7/18,negativeveryverythinmathspace:-1/18,negativeverythinmathspace:-2/18,negativethinmathspace:-3/18,negativemediummathspace:-4/18,negativethickmathspace:-5/18,negativeverythickmathspace:-6/18,negativeveryverythickmathspace:-7/18,thin:0.08,medium:0.1,thick:0.15,infinity:k},TeX:{x_height:0.430554},pxPerInch:72,em:16,DELIMITERS:{"(":{dir:c},"{":{dir:c,w:0.58},"[":{dir:c},"|":{dir:c,w:0.275},")":{dir:c},"}":{dir:c,w:0.58},"]":{dir:c},"/":{dir:c},"\\":{dir:c},"\u2223":{dir:c,w:0.275},"\u2225":{dir:c,w:0.55},"\u230A":{dir:c,w:0.5},"\u230B":{dir:c,w:0.5},"\u2308":{dir:c,w:0.5},"\u2309":{dir:c,w:0.5},"\u27E8":{dir:c,w:0.5},"\u27E9":{dir:c,w:0.5},"\u2191":{dir:c,w:0.65},"\u2193":{dir:c,w:0.65},"\u21D1":{dir:c,w:0.75},"\u21D3":{dir:c,w:0.75},"\u2195":{dir:c,w:0.65},"\u21D5":{dir:c,w:0.75},"\u27EE":{dir:c,w:0.275},"\u27EF":{dir:c,w:0.275},"\u23B0":{dir:c,w:0.6},"\u23B1":{dir:c,w:0.6}},REMAPACCENT:{"\u20D7":"\u2192","'":"\u02CB","`":"\u02CA",".":"\u02D9","^":"\u02C6","-":"\u02C9","~":"\u02DC","\u00AF":"\u02C9","\u00B0":"\u02DA","\u00B4":"\u02CA","\u0300":"\u02CB","\u0301":"\u02CA","\u0302":"\u02C6","\u0303":"\u02DC","\u0304":"\u02C9","\u0305":"\u02C9","\u0306":"\u02D8","\u0307":"\u02D9","\u0308":"\u00A8","\u030C":"\u02C7"},REMAPACCENTUNDER:{},length2em:function(r,p){if(typeof(r)!=="string"){r=r.toString()}if(r===""){return""}if(r===h.SIZE.NORMAL){return 1}if(r===h.SIZE.BIG){return 2}if(r===h.SIZE.SMALL){return 0.71}if(this.MATHSPACE[r]){return this.MATHSPACE[r]}var o=r.match(/^\s*([-+]?(?:\.\d+|\d+(?:\.\d*)?))?(pt|em|ex|mu|px|pc|in|mm|cm|%)?/);var n=parseFloat(o[1]||"1"),q=o[2];if(p==null){p=1}if(q==="em"){return n}if(q==="ex"){return n*this.TeX.x_height}if(q==="%"){return n/100*p}if(q==="px"){return n/this.em}if(q==="pt"){return n/10}if(q==="pc"){return n*1.2}if(q==="in"){return n*this.pxPerInch/this.em}if(q==="cm"){return n*this.pxPerInch/this.em/2.54}if(q==="mm"){return n*this.pxPerInch/this.em/25.4}if(q==="mu"){return n/18}return n*p},Em:function(n){if(Math.abs(n)<0.001){return"0em"}return(n.toFixed(3).replace(/\.?0+$/,""))+"em"},arrayEntry:function(n,o){return n[Math.max(0,Math.min(o,n.length-1))]}});MathJax.Hub.Register.StartupHook("mml Jax Ready",function(){h=MathJax.ElementJax.mml;h.mbase.Augment({toPreviewHTML:function(o,n){return this.PHTMLdefaultSpan(o,n)},PHTMLdefaultSpan:function(q,o){if(!o){o={}}q=this.PHTMLcreateSpan(q);this.PHTMLhandleStyle(q);this.PHTMLhandleColor(q);if(this.isToken){this.PHTMLhandleToken(q)}for(var p=0,n=this.data.length;p<n;p++){this.PHTMLaddChild(q,p,o)}return q},PHTMLaddChild:function(p,o,n){var q=this.data[o];if(q){if(n.childSpans){p=e.addElement(p,"span",{className:n.className})}q.toPreviewHTML(p);if(!n.noBBox){this.PHTML.w+=q.PHTML.w+q.PHTML.l+q.PHTML.r;if(q.PHTML.h>this.PHTML.h){this.PHTML.h=q.PHTML.h}if(q.PHTML.d>this.PHTML.d){this.PHTML.d=q.PHTML.d}if(q.PHTML.t>this.PHTML.t){this.PHTML.t=q.PHTML.t}if(q.PHTML.b>this.PHTML.b){this.PHTML.b=q.PHTML.b}}}else{if(n.forceChild){e.addElement(p,"span")}}},PHTMLstretchChild:function(q,p,s){var r=this.data[q];if(r&&r.PHTMLcanStretch("Vertical",p,s)){var t=this.PHTML,o=r.PHTML,n=o.w;r.PHTMLstretchV(p,s);t.w+=o.w-n;if(o.h>t.h){t.h=o.h}if(o.d>t.d){t.d=o.d}}},PHTMLcreateSpan:function(n){if(!this.PHTML){this.PHTML={}}this.PHTML={w:0,h:0,d:0,l:0,r:0,t:0,b:0};if(this.inferred){return n}if(this.type==="mo"&&this.data.join("")==="\u222B"){g.lastIsInt=true}else{if(this.type!=="mspace"||this.width!=="negativethinmathspace"){g.lastIsInt=false}}if(!this.PHTMLspanID){this.PHTMLspanID=g.GetID()}var o=(this.id||"MJXp-Span-"+this.PHTMLspanID);return e.addElement(n,"span",{className:"MJXp-"+this.type,id:o})},PHTMLspanElement:function(){if(!this.PHTMLspanID){return null}return document.getElementById(this.id||"MJXp-Span-"+this.PHTMLspanID)},PHTMLhandleToken:function(o){var n=this.getValues("mathvariant");if(n.mathvariant!==h.VARIANT.NORMAL){o.className+=" "+g.VARIANT[n.mathvariant]}},PHTMLhandleStyle:function(n){if(this.style){n.style.cssText=this.style}},PHTMLhandleColor:function(n){if(this.mathcolor){n.style.color=this.mathcolor}if(this.mathbackground){n.style.backgroundColor=this.mathbackground}},PHTMLhandleScriptlevel:function(n){var o=this.Get("scriptlevel");if(o){n.className+=" MJXp-script"}},PHTMLhandleText:function(y,A){var v,p;var z=0,o=0,q=0;for(var s=0,r=A.length;s<r;s++){p=A.charCodeAt(s);v=A.charAt(s);if(p>=55296&&p<56319){s++;p=(((p-55296)<<10)+(A.charCodeAt(s)-56320))+65536}var t=0.7,u=0.22,x=0.5;if(p<127){if(v.match(/[A-Za-ehik-or-xz0-9]/)){u=0}if(v.match(/[A-HK-Z]/)){x=0.67}else{if(v.match(/[IJ]/)){x=0.36}}if(v.match(/[acegm-su-z]/)){t=0.45}else{if(v.match(/[ij]/)){t=0.75}}if(v.match(/[ijlt]/)){x=0.28}}if(g.DELIMITERS[v]){x=g.DELIMITERS[v].w||0.4}if(t>z){z=t}if(u>o){o=u}q+=x}if(!this.CHML){this.PHTML={}}this.PHTML={h:0.9,d:0.3,w:q,l:0,r:0,t:z,b:o};e.addText(y,A)},PHTMLbboxFor:function(o){if(this.data[o]&&this.data[o].PHTML){return this.data[o].PHTML}return{w:0,h:0,d:0,l:0,r:0,t:0,b:0}},PHTMLcanStretch:function(q,o,p){if(this.isEmbellished()){var n=this.Core();if(n&&n!==this){return n.PHTMLcanStretch(q,o,p)}}return false},PHTMLstretchV:function(n,o){},PHTMLstretchH:function(n){},CoreParent:function(){var n=this;while(n&&n.isEmbellished()&&n.CoreMO()===this&&!n.isa(h.math)){n=n.Parent()}return n},CoreText:function(n){if(!n){return""}if(n.isEmbellished()){return n.CoreMO().data.join("")}while((n.isa(h.mrow)||n.isa(h.TeXAtom)||n.isa(h.mstyle)||n.isa(h.mphantom))&&n.data.length===1&&n.data[0]){n=n.data[0]}if(!n.isToken){return""}else{return n.data.join("")}}});h.chars.Augment({toPreviewHTML:function(n){var o=this.toString().replace(/[\u2061-\u2064]/g,"");this.PHTMLhandleText(n,o)}});h.entity.Augment({toPreviewHTML:function(n){var o=this.toString().replace(/[\u2061-\u2064]/g,"");this.PHTMLhandleText(n,o)}});h.math.Augment({toPreviewHTML:function(n){n=this.PHTMLdefaultSpan(n);if(this.Get("display")==="block"){n.className+=" MJXp-display"}return n}});h.mo.Augment({toPreviewHTML:function(o){o=this.PHTMLdefaultSpan(o);this.PHTMLadjustAccent(o);var n=this.getValues("lspace","rspace","scriptlevel","displaystyle","largeop");if(n.scriptlevel===0){this.PHTML.l=g.length2em(n.lspace);this.PHTML.r=g.length2em(n.rspace);o.style.marginLeft=g.Em(this.PHTML.l);o.style.marginRight=g.Em(this.PHTML.r)}else{this.PHTML.l=0.15;this.PHTML.r=0.1}if(n.displaystyle&&n.largeop){var p=e.Element("span",{className:"MJXp-largeop"});p.appendChild(o.firstChild);o.appendChild(p);this.PHTML.h*=1.2;this.PHTML.d*=1.2;if(this.data.join("")==="\u222B"){p.className+=" MJXp-int"}}return o},PHTMLadjustAccent:function(p){var o=this.CoreParent();if(o&&o.isa(h.munderover)&&this.CoreText(o.data[o.base]).length===1){var q=o.data[o.over],n=o.data[o.under];var s=this.data.join(""),r;if(q&&this===q.CoreMO()&&o.Get("accent")){r=g.REMAPACCENT[s]}else{if(n&&this===n.CoreMO()&&o.Get("accentunder")){r=g.REMAPACCENTUNDER[s]}}if(r){s=p.innerHTML=r}if(s.match(/[\u02C6-\u02DC\u00A8]/)){this.PHTML.acc=-0.52}else{if(s==="\u2192"){this.PHTML.acc=-0.15;this.PHTML.vec=true}}}},PHTMLcanStretch:function(q,o,p){if(!this.Get("stretchy")){return false}var r=this.data.join("");if(r.length>1){return false}r=g.DELIMITERS[r];var n=(r&&r.dir===q.substr(0,1));if(n){n=(this.PHTML.h!==o||this.PHTML.d!==p||(this.Get("minsize",true)||this.Get("maxsize",true)))}return n},PHTMLstretchV:function(p,u){var o=this.PHTMLspanElement(),t=this.PHTML;var n=this.getValues("symmetric","maxsize","minsize");if(n.symmetric){l=2*Math.max(p-0.25,u+0.25)}else{l=p+u}n.maxsize=g.length2em(n.maxsize,t.h+t.d);n.minsize=g.length2em(n.minsize,t.h+t.d);l=Math.max(n.minsize,Math.min(n.maxsize,l));var s=l/(t.h+t.d-0.3);var q=e.Element("span",{style:{"font-size":g.Em(s)}});if(s>1.25){var r=Math.ceil(1.25/s*10);q.className="MJXp-right MJXp-scale"+r;q.style.marginLeft=g.Em(t.w*(r/10-1)+0.07);t.w*=s*r/10}q.appendChild(o.firstChild);o.appendChild(q);if(n.symmetric){o.style.verticalAlign=g.Em(0.25*(1-s))}}});h.mspace.Augment({toPreviewHTML:function(q){q=this.PHTMLdefaultSpan(q);var o=this.getValues("height","depth","width");var n=g.length2em(o.width),p=g.length2em(o.height),s=g.length2em(o.depth);var r=this.PHTML;r.w=n;r.h=p;r.d=s;if(n<0){if(!g.lastIsInt){q.style.marginLeft=g.Em(n)}n=0}q.style.width=g.Em(n);q.style.height=g.Em(p+s);if(s){q.style.verticalAlign=g.Em(-s)}return q}});h.mpadded.Augment({toPreviewHTML:function(u){u=this.PHTMLdefaultSpan(u,{childSpans:true,className:"MJXp-box",forceChild:true});var o=u.firstChild;var v=this.getValues("width","height","depth","lspace","voffset");var s=this.PHTMLdimen(v.lspace);var q=0,n=0,t=s.len,r=-s.len,p=0;if(v.width!==""){s=this.PHTMLdimen(v.width,"w",0);if(s.pm){r+=s.len}else{u.style.width=g.Em(s.len)}}if(v.height!==""){s=this.PHTMLdimen(v.height,"h",0);if(!s.pm){q+=-this.PHTMLbboxFor(0).h}q+=s.len}if(v.depth!==""){s=this.PHTMLdimen(v.depth,"d",0);if(!s.pm){n+=-this.PHTMLbboxFor(0).d;p+=-s.len}n+=s.len}if(v.voffset!==""){s=this.PHTMLdimen(v.voffset);q-=s.len;n+=s.len;p+=s.len}if(q){o.style.marginTop=g.Em(q)}if(n){o.style.marginBottom=g.Em(n)}if(t){o.style.marginLeft=g.Em(t)}if(r){o.style.marginRight=g.Em(r)}if(p){u.style.verticalAlign=g.Em(p)}return u},PHTMLdimen:function(q,r,n){if(n==null){n=-k}q=String(q);var o=q.match(/width|height|depth/);var p=(o?this.PHTML[o[0].charAt(0)]:(r?this.PHTML[r]:0));return{len:g.length2em(q,p)||0,pm:!!q.match(/^[-+]/)}}});h.munderover.Augment({toPreviewHTML:function(r){var t=this.getValues("displaystyle","accent","accentunder","align");var n=this.data[this.base];if(!t.displaystyle&&n!=null&&(n.movablelimits||n.CoreMO().Get("movablelimits"))){r=h.msubsup.prototype.toPreviewHTML.call(this,r);r.className=r.className.replace(/munderover/,"msubsup");return r}r=this.PHTMLdefaultSpan(r,{childSpans:true,className:"",noBBox:true});var p=this.PHTMLbboxFor(this.over),v=this.PHTMLbboxFor(this.under),u=this.PHTMLbboxFor(this.base),s=this.PHTML,o=p.acc;if(this.data[this.over]){if(r.lastChild.firstChild){r.lastChild.firstChild.style.marginLeft=p.l=r.lastChild.firstChild.style.marginRight=p.r=0}var q=e.Element("span",{},[["span",{className:"MJXp-over"}]]);q.firstChild.appendChild(r.lastChild);if(r.childNodes.length>(this.data[this.under]?1:0)){q.firstChild.appendChild(r.firstChild)}this.data[this.over].PHTMLhandleScriptlevel(q.firstChild.firstChild);if(o!=null){if(p.vec){q.firstChild.firstChild.firstChild.style.fontSize="60%";p.h*=0.6;p.d*=0.6;p.w*=0.6}o=o-p.d+0.1;if(u.t!=null){o+=u.t-u.h}q.firstChild.firstChild.style.marginBottom=g.Em(o)}if(r.firstChild){r.insertBefore(q,r.firstChild)}else{r.appendChild(q)}}if(this.data[this.under]){if(r.lastChild.firstChild){r.lastChild.firstChild.style.marginLeft=v.l=r.lastChild.firstChild.marginRight=v.r=0}this.data[this.under].PHTMLhandleScriptlevel(r.lastChild)}s.w=Math.max(0.8*p.w,0.8*v.w,u.w);s.h=0.8*(p.h+p.d+(o||0))+u.h;s.d=u.d+0.8*(v.h+v.d);return r}});h.msubsup.Augment({toPreviewHTML:function(q){q=this.PHTMLdefaultSpan(q,{noBBox:true});if(!this.data[this.base]){if(q.firstChild){q.insertBefore(e.Element("span"),q.firstChild)}else{q.appendChild(e.Element("span"))}}var s=this.data[this.base],p=this.data[this.sub],n=this.data[this.sup];if(!s){s={bbox:{h:0.8,d:0.2}}}q.firstChild.style.marginRight=".05em";var o=Math.max(0.4,s.PHTML.h-0.4),u=Math.max(0.2,s.PHTML.d+0.1);var t=this.PHTML;if(n&&p){var r=e.Element("span",{className:"MJXp-script-box",style:{height:g.Em(o+n.PHTML.h*0.8+u+p.PHTML.d*0.8),"vertical-align":g.Em(-u-p.PHTML.d*0.8)}},[["span",{},[["span",{},[["span",{style:{"margin-bottom":g.Em(-(n.PHTML.d-0.05))}}]]]]],["span",{},[["span",{},[["span",{style:{"margin-top":g.Em(-(n.PHTML.h-0.05))}}]]]]]]);p.PHTMLhandleScriptlevel(r.firstChild);n.PHTMLhandleScriptlevel(r.lastChild);r.firstChild.firstChild.firstChild.appendChild(q.lastChild);r.lastChild.firstChild.firstChild.appendChild(q.lastChild);q.appendChild(r);t.h=Math.max(s.PHTML.h,n.PHTML.h*0.8+o);t.d=Math.max(s.PHTML.d,p.PHTML.d*0.8+u);t.w=s.PHTML.w+Math.max(n.PHTML.w,p.PHTML.w)+0.07}else{if(n){q.lastChild.style.verticalAlign=g.Em(o);n.PHTMLhandleScriptlevel(q.lastChild);t.h=Math.max(s.PHTML.h,n.PHTML.h*0.8+o);t.d=Math.max(s.PHTML.d,n.PHTML.d*0.8-o);t.w=s.PHTML.w+n.PHTML.w+0.07}else{if(p){q.lastChild.style.verticalAlign=g.Em(-u);p.PHTMLhandleScriptlevel(q.lastChild);t.h=Math.max(s.PHTML.h,p.PHTML.h*0.8-u);t.d=Math.max(s.PHTML.d,p.PHTML.d*0.8+u);t.w=s.PHTML.w+p.PHTML.w+0.07}}}return q}});h.mfrac.Augment({toPreviewHTML:function(r){r=this.PHTMLdefaultSpan(r,{childSpans:true,className:"MJXp-box",forceChild:true,noBBox:true});var o=this.getValues("linethickness","displaystyle");if(!o.displaystyle){if(this.data[0]){this.data[0].PHTMLhandleScriptlevel(r.firstChild)}if(this.data[1]){this.data[1].PHTMLhandleScriptlevel(r.lastChild)}}var n=e.Element("span",{className:"MJXp-box"},[["span",{className:"MJXp-denom"},[["span",{},[["span",{className:"MJXp-rule",style:{height:"1em"}}]]],["span"]]]]);n.firstChild.lastChild.appendChild(r.lastChild);r.appendChild(n);var s=this.PHTMLbboxFor(0),p=this.PHTMLbboxFor(1),v=this.PHTML;v.w=Math.max(s.w,p.w)*0.8;v.h=s.h+s.d+0.1+0.25;v.d=p.h+p.d-0.25;v.l=v.r=0.125;o.linethickness=Math.max(0,g.length2em(o.linethickness||"0",0));if(o.linethickness){var u=n.firstChild.firstChild.firstChild;var q=g.Em(o.linethickness);u.style.borderTop="none";u.style.borderBottom=(o.linethickness<0.15?"1px":q)+" solid";u.style.margin=q+" 0";q=o.linethickness;n.style.marginTop=g.Em(3*q-1.2);r.style.verticalAlign=g.Em(1.5*q+0.1);v.h+=1.5*q-0.1;v.d+=1.5*q}else{n.style.marginTop="-.7em"}return r}});h.msqrt.Augment({toPreviewHTML:function(n){n=this.PHTMLdefaultSpan(n,{childSpans:true,className:"MJXp-box",forceChild:true,noBBox:true});this.PHTMLlayoutRoot(n,n.firstChild);return n},PHTMLlayoutRoot:function(u,n){var v=this.PHTMLbboxFor(0);var q=Math.ceil((v.h+v.d+0.14)*100),w=g.Em(14/q);var r=e.Element("span",{className:"MJXp-surd"},[["span",{style:{"font-size":q+"%","margin-top":w}},["\u221A"]]]);var s=e.Element("span",{className:"MJXp-root"},[["span",{className:"MJXp-rule",style:{"border-top":".08em solid"}}]]);var p=(1.2/2.2)*q/100;if(q>150){var o=Math.ceil(150/q*10);r.firstChild.className="MJXp-right MJXp-scale"+o;r.firstChild.style.marginLeft=g.Em(p*(o/10-1)/q*100);p=p*o/10;s.firstChild.style.borderTopWidth=g.Em(0.08/Math.sqrt(o/10))}s.appendChild(n);u.appendChild(r);u.appendChild(s);this.PHTML.h=v.h+0.18;this.PHTML.d=v.d;this.PHTML.w=v.w+p;return u}});h.mroot.Augment({toPreviewHTML:function(q){q=this.PHTMLdefaultSpan(q,{childSpans:true,className:"MJXp-box",forceChild:true,noBBox:true});var p=this.PHTMLbboxFor(1),n=q.removeChild(q.lastChild);var t=this.PHTMLlayoutRoot(e.Element("span"),q.firstChild);n.className="MJXp-script";var u=parseInt(t.firstChild.firstChild.style.fontSize);var o=0.55*(u/120)+p.d*0.8,s=-0.6*(u/120);if(u>150){s*=0.95*Math.ceil(150/u*10)/10}n.style.marginRight=g.Em(s);n.style.verticalAlign=g.Em(o);if(-s>p.w*0.8){n.style.marginLeft=g.Em(-s-p.w*0.8)}q.appendChild(n);q.appendChild(t);this.PHTML.w+=Math.max(0,p.w*0.8+s);this.PHTML.h=Math.max(this.PHTML.h,p.h*0.8+o);return q},PHTMLlayoutRoot:h.msqrt.prototype.PHTMLlayoutRoot});h.mfenced.Augment({toPreviewHTML:function(q){q=this.PHTMLcreateSpan(q);this.PHTMLhandleStyle(q);this.PHTMLhandleColor(q);this.addFakeNodes();this.PHTMLaddChild(q,"open",{});for(var p=0,n=this.data.length;p<n;p++){this.PHTMLaddChild(q,"sep"+p,{});this.PHTMLaddChild(q,p,{})}this.PHTMLaddChild(q,"close",{});var o=this.PHTML.h,r=this.PHTML.d;this.PHTMLstretchChild("open",o,r);for(p=0,n=this.data.length;p<n;p++){this.PHTMLstretchChild("sep"+p,o,r);this.PHTMLstretchChild(p,o,r)}this.PHTMLstretchChild("close",o,r);return q}});h.mrow.Augment({toPreviewHTML:function(q){q=this.PHTMLdefaultSpan(q);var p=this.PHTML.h,r=this.PHTML.d;for(var o=0,n=this.data.length;o<n;o++){this.PHTMLstretchChild(o,p,r)}return q}});h.mstyle.Augment({toPreviewHTML:function(n){n=this.PHTMLdefaultSpan(n);this.PHTMLhandleScriptlevel(n);return n}});h.TeXAtom.Augment({toPreviewHTML:function(n){n=this.PHTMLdefaultSpan(n);n.className="MJXp-mrow";return n}});h.mtable.Augment({toPreviewHTML:function(E){E=this.PHTMLdefaultSpan(E,{noBBox:true});var r=this.getValues("columnalign","rowalign","columnspacing","rowspacing","columnwidth","equalcolumns","equalrows","columnlines","rowlines","frame","framespacing","align","width");var u=MathJax.Hub.SplitList,F,A,D,z;var N=u(r.columnspacing),w=u(r.rowspacing),L=u(r.columnalign),t=u(r.rowalign);for(F=0,A=N.length;F<A;F++){N[F]=g.length2em(N[F])}for(F=0,A=w.length;F<A;F++){w[F]=g.length2em(w[F])}var K=e.Element("span");while(E.firstChild){K.appendChild(E.firstChild)}E.appendChild(K);var y=0,s=0;for(F=0,A=this.data.length;F<A;F++){var v=this.data[F];if(v){var J=g.arrayEntry(w,F-1),C=g.arrayEntry(t,F);var x=v.PHTML,q=v.PHTMLspanElement();q.style.verticalAlign=C;var B=(v.type==="mlabeledtr"?1:0);for(D=0,z=v.data.length;D<z-B;D++){var p=v.data[D+B];if(p){var M=g.arrayEntry(N,D-1),G=g.arrayEntry(L,D);var I=p.PHTMLspanElement();if(D){x.w+=M;I.style.paddingLeft=g.Em(M)}if(F){I.style.paddingTop=g.Em(J)}I.style.textAlign=G}}y+=x.h+x.d;if(F){y+=J}if(x.w>s){s=x.w}}}var o=this.PHTML;o.w=s;o.h=y/2+0.25;o.d=y/2-0.25;o.l=o.r=0.125;return E}});h.mlabeledtr.Augment({PHTMLdefaultSpan:function(q,o){if(!o){o={}}q=this.PHTMLcreateSpan(q);this.PHTMLhandleStyle(q);this.PHTMLhandleColor(q);if(this.isToken){this.PHTMLhandleToken(q)}for(var p=1,n=this.data.length;p<n;p++){this.PHTMLaddChild(q,p,o)}return q}});h.semantics.Augment({toPreviewHTML:function(n){n=this.PHTMLcreateSpan(n);if(this.data[0]){this.data[0].toPreviewHTML(n);MathJax.Hub.Insert(this.data[0].PHTML||{},this.PHTML)}return n}});h.annotation.Augment({toPreviewHTML:function(n){}});h["annotation-xml"].Augment({toPreviewHTML:function(n){}});MathJax.Hub.Register.StartupHook("onLoad",function(){setTimeout(MathJax.Callback(["loadComplete",g,"jax.js"]),0)})});MathJax.Hub.Register.StartupHook("End Cookie",function(){if(b.config.menuSettings.zoom!=="None"){i.Require("[MathJax]/extensions/MathZoom.js")}})})(MathJax.Ajax,MathJax.Hub,MathJax.HTML,MathJax.OutputJax.PreviewHTML);
(function(b,g,f){var c=b.config.menuSettings;var e=MathJax.OutputJax;var a=f.isMSIE&&(document.documentMode||0)<8;var d=MathJax.Extension["fast-preview"]={version:"2.7.0",enabled:true,config:b.CombineConfig("fast-preview",{Chunks:{EqnChunk:10000,EqnChunkFactor:1,EqnChunkDelay:0},color:"inherit!important",updateTime:30,updateDelay:6,messageStyle:"none",disabled:f.isMSIE&&!f.versionAtLeast("8.0")}),Config:function(){if(b.config["CHTML-preview"]){MathJax.Hub.Config({"fast-preview":b.config["CHTML-preview"]})}var m,j,k,h,l;var i=this.config;if(!i.disabled&&c.FastPreview==null){b.Config({menuSettings:{FastPreview:true}})}if(c.FastPreview){MathJax.Ajax.Styles({".MathJax_Preview .MJXf-math":{color:i.color}});b.Config({"HTML-CSS":i.Chunks,CommonHTML:i.Chunks,SVG:i.Chunks})}b.Register.MessageHook("Begin Math Output",function(){if(!h&&d.Active()){m=b.processUpdateTime;j=b.processUpdateDelay;k=b.config.messageStyle;b.processUpdateTime=i.updateTime;b.processUpdateDelay=i.updateDelay;b.Config({messageStyle:i.messageStyle});MathJax.Message.Clear(0,0);l=true}});b.Register.MessageHook("End Math Output",function(){if(!h&&l){b.processUpdateTime=m;b.processUpdateDelay=j;b.Config({messageStyle:k});h=true}})},Disable:function(){this.enabled=false},Enable:function(){this.enabled=true},Active:function(){return c.FastPreview&&this.enabled&&!(e[c.renderer]||{}).noFastPreview},Preview:function(h){if(!this.Active()||!h.script.parentNode){return}var i=h.script.MathJax.preview||h.script.previousSibling;if(!i||i.className!==MathJax.Hub.config.preRemoveClass){i=g.Element("span",{className:MathJax.Hub.config.preRemoveClass});h.script.parentNode.insertBefore(i,h.script);h.script.MathJax.preview=i}i.innerHTML="";i.style.color=(a?"black":"inherit");return this.postFilter(i,h)},postFilter:function(j,i){if(!i.math.root.toPreviewHTML){var h=MathJax.Callback.Queue();h.Push(["Require",MathJax.Ajax,"[MathJax]/jax/output/PreviewHTML/config.js"],["Require",MathJax.Ajax,"[MathJax]/jax/output/PreviewHTML/jax.js"]);b.RestartAfter(h.Push({}))}i.math.root.toPreviewHTML(j)},Register:function(h){b.Register.StartupHook(h+" Jax Require",function(){var i=MathJax.InputJax[h];i.postfilterHooks.Add(["Preview",MathJax.Extension["fast-preview"]],50)})}};d.Register("TeX");d.Register("MathML");d.Register("AsciiMath");b.Register.StartupHook("End Config",["Config",d]);b.Startup.signal.Post("fast-preview Ready")})(MathJax.Hub,MathJax.HTML,MathJax.Hub.Browser);MathJax.Ajax.loadComplete("[MathJax]/extensions/fast-preview.js");
(function(a,e,b,f){var c=b.config.menuSettings;var d=MathJax.Extension.AssistiveMML={version:"2.7.0",config:b.CombineConfig("AssistiveMML",{disabled:false,styles:{".MJX_Assistive_MathML":{position:"absolute!important",top:0,left:0,clip:(b.Browser.isMSIE&&(document.documentMode||0)<8?"rect(1px 1px 1px 1px)":"rect(1px, 1px, 1px, 1px)"),padding:"1px 0 0 0!important",border:"0!important",height:"1px!important",width:"1px!important",overflow:"hidden!important",display:"block!important","-webkit-touch-callout":"none","-webkit-user-select":"none","-khtml-user-select":"none","-moz-user-select":"none","-ms-user-select":"none","user-select":"none"},".MJX_Assistive_MathML.MJX_Assistive_MathML_Block":{width:"100%!important"}}}),Config:function(){if(!this.config.disabled&&c.assistiveMML==null){b.Config({menuSettings:{assistiveMML:true}})}a.Styles(this.config.styles);b.Register.MessageHook("End Math",function(g){if(c.assistiveMML){return d.AddAssistiveMathML(g[1])}})},AddAssistiveMathML:function(g){var h={jax:b.getAllJax(g),i:0,callback:MathJax.Callback({})};this.HandleMML(h);return h.callback},RemoveAssistiveMathML:function(k){var h=b.getAllJax(k),l;for(var j=0,g=h.length;j<g;j++){l=document.getElementById(h[j].inputID+"-Frame");if(l&&l.getAttribute("data-mathml")){l.removeAttribute("data-mathml");if(l.lastChild&&l.lastChild.className.match(/MJX_Assistive_MathML/)){l.removeChild(l.lastChild)}}}},HandleMML:function(l){var g=l.jax.length,h,i,n,j;while(l.i<g){h=l.jax[l.i];n=document.getElementById(h.inputID+"-Frame");if(h.outputJax!=="NativeMML"&&h.outputJax!=="PlainSource"&&n&&!n.getAttribute("data-mathml")){try{i=h.root.toMathML("").replace(/\n */g,"").replace(/<!--.*?-->/g,"")}catch(k){if(!k.restart){throw k}return MathJax.Callback.After(["HandleMML",this,l],k.restart)}n.setAttribute("data-mathml",i);j=f.addElement(n,"span",{isMathJax:true,unselectable:"on",className:"MJX_Assistive_MathML"+(h.root.Get("display")==="block"?" MJX_Assistive_MathML_Block":"")});try{j.innerHTML=i}catch(k){}n.style.position="relative";n.setAttribute("role","presentation");n.firstChild.setAttribute("aria-hidden","true");j.setAttribute("role","presentation")}l.i++}l.callback()}};b.Startup.signal.Post("AssistiveMML Ready")})(MathJax.Ajax,MathJax.Callback,MathJax.Hub,MathJax.HTML);MathJax.Callback.Queue(["Require",MathJax.Ajax,"[MathJax]/extensions/toMathML.js"],["loadComplete",MathJax.Ajax,"[MathJax]/extensions/AssistiveMML.js"],function(){MathJax.Hub.Register.StartupHook("End Config",["Config",MathJax.Extension.AssistiveMML])});
MathJax.Ajax.loadComplete("[MathJax]/config/TeX-AMS_HTML-full.js");
```
