## Supplementary material for "Artistoo: build, share, and explore simulations of cells and tissues in the web browser": fontdata.js

```
/*
 *  /MathJax/jax/output/HTML-CSS/fonts/STIX/fontdata.js
 *
 *  Copyright (c) 2009-2016 The MathJax Consortium
 *
 *  Licensed under the Apache License, Version 2.0 (the "License");
 *  you may not use this file except in compliance with the License.
 *  You may obtain a copy of the License at
 *
 *      http://www.apache.org/licenses/LICENSE-2.0
 *
 *  Unless required by applicable law or agreed to in writing, software
 *  distributed under the License is distributed on an "AS IS" BASIS,
 *  WITHOUT WARRANTIES OR CONDITIONS OF ANY KIND, either express or implied.
 *  See the License for the specific language governing permissions and
 *  limitations under the License.
 */

(function(j,b,s){var t="2.7.0";j.allowWebFonts=false;var m="STIXGeneral",u="STIXGeneral-bold",q="STIXGeneral-italic",v="STIXGeneral-bold-italic",r="STIXNonUnicode",p="STIXNonUnicode-italic",i="STIXSizeOneSym",h="STIXSizeTwoSym",g="STIXSizeThreeSym",d="STIXSizeFourSym",c="STIXSizeFiveSym";var l="H",a="V",n={load:"extra",dir:l},f={load:"extra",dir:a};var k=[8722,m,0,0,0,-0.26,-0.26];j.Augment({FONTDATA:{version:t,STIXversion:"1.1",TeX_factor:1.125,baselineskip:1.2,lineH:0.8,lineD:0.2,FONTS:{STIXGeneral:"General/Regular/Main.js","STIXGeneral-italic":"General/Italic/Main.js","STIXGeneral-bold":"General/Bold/Main.js","STIXGeneral-bold-italic":"General/BoldItalic/Main.js",STIXNonUnicode:"NonUnicode/Regular/Main.js","STIXNonUnicode-italic":"NonUnicode/Italic/Main.js","STIXNonUnicode-bold":"NonUnicode/Bold/Main.js","STIXNonUnicode-bold-italic":"NonUnicode/BoldItalic/Main.js",STIXVariants:"Variants/Regular/All.js",STIXSizeOneSym:"SizeOneSym/Regular/All.js",STIXSizeTwoSym:"SizeTwoSym/Regular/All.js",STIXSizeThreeSym:"SizeThreeSym/Regular/All.js",STIXSizeFourSym:"SizeFourSym/Regular/All.js",STIXSizeFiveSym:"SizeFiveSym/Regular/All.js",STIXIntegralsD:"IntegralsD/Regular/All.js"},VARIANT:{normal:{fonts:[m,r,i],remap:{8709:[8709,"-STIX-variant"],124:[124,"-STIX-variant"]}},bold:{fonts:[u,"STIXNonUnicode-bold","STIXSizeOneSym-bold"],bold:true},italic:{fonts:[q,p,m,r,i],italic:true},"bold-italic":{fonts:[v,"STIXNonUnicode-bold-italic"],bold:true,italic:true},"double-struck":{offsetA:120120,offsetN:120792,remap:{120122:8450,120127:8461,120133:8469,120135:8473,120136:8474,120137:8477,120145:8484}},fraktur:{offsetA:120068,remap:{120070:8493,120075:8460,120076:8465,120085:8476,120093:8488}},"bold-fraktur":{fonts:[u],offsetA:120172,bold:true},script:{fonts:[q],offsetA:119964,italic:true,remap:{119965:8492,119968:8496,119969:8497,119971:8459,119972:8464,119975:8466,119976:8499,119981:8475,119994:8495,119996:8458,120004:8500}},"bold-script":{fonts:[v],offsetA:120016,bold:true,italic:true},"sans-serif":{offsetA:120224,offsetN:120802,offsetG:57725,offsetE:57725},"bold-sans-serif":{offsetA:120276,offsetG:120662,offsetN:120812,bold:true},"sans-serif-italic":{fonts:[q,p],offsetA:120328,offsetN:57780,offsetG:57791,offsetE:57791,italic:true},"sans-serif-bold-italic":{fonts:[v,"STIXNonUnicode-bold-italic"],offsetA:120380,offsetN:57846,offsetG:120720,bold:true,italic:true},monospace:{offsetA:120432,offsetN:120822,remap:{32:[32,"-STIX-variant"]}},"-STIX-variant":{fonts:["STIXVariants",r,m],remap:{10887:57360,10888:57359,8816:57361,8817:57358,8928:57419,8929:57423,8840:57366,8841:57368,9651:9653,9661:9663,8709:[8709,b.VARIANT.NORMAL],124:[124,b.VARIANT.NORMAL]}},"-tex-caligraphic":{fonts:[q,p,r,i],offsetA:57901,noLowerCase:1},"-tex-oldstyle":{offsetN:57953,remap:{57954:57957,57955:57961,57956:57965,57957:57969,57958:57973,57959:57977,57960:57981,57961:57985,57962:57989}},"-tex-mathit":{fonts:[q,p,m,r,i],italic:true,noIC:true},"-largeOp":{fonts:[i,"STIXIntegralsD",r,m]},"-smallOp":{},"-tex-caligraphic-bold":{fonts:["STIXGeneral-bold-italic","STIXNonUnicode-bold-italic","STIXNonUnicode","STIXGeneral","STIXSizeOneSym"],bold:true,offsetA:57927,noLowerCase:1},"-tex-oldstyle-bold":{fonts:["STIXGeneral-bold","STIXNonUnicode-bold","STIXGeneral","STIXSizeOneSym"],bold:true,offsetN:57955,remap:{57956:57959,57957:57963,57958:57967,57959:57971,57960:57975,57961:57979,57962:57983,57963:57987,57964:57991}}},RANGES:[{name:"alpha",low:97,high:122,offset:"A",add:26},{name:"Alpha",low:65,high:90,offset:"A"},{name:"number",low:48,high:57,offset:"N"},{name:"greek-non-unicode",low:945,high:969,offset:"E",add:25},{name:"greek",low:945,high:969,offset:"G",add:26},{name:"Greek",low:913,high:1014,offset:"G",remap:{1013:53,977:54,1008:55,981:56,1009:57,982:58,1012:17}}],RULECHAR:8254,REMAP:{755:730,756:714,65079:9182,65080:9183,12296:10216,12297:10217,10072:8739},REMAPACCENT:{"\u007E":"\u0303","\u2192":"\u20D7","\u2190":"\u20D6","\u0060":"\u0300","\u005E":"\u0302","\u00B4":"\u0301","\u2032":"\u0301","\u2035":"\u0300"},REMAPACCENTUNDER:{},DELIMITERS:{40:{dir:a,HW:[[0.844,m],[1.23,i],[1.353,i,1.1],[1.845,h],[2.048,h,1.11],[2.46,g],[2.472,g,1.005],[3.075,d]],stretch:{top:[9115,i],ext:[9116,i],bot:[9117,i]}},41:{dir:a,HW:[[0.844,m],[1.23,i],[1.353,i,1.1],[1.845,h],[2.048,h,1.11],[2.46,g],[2.472,g,1.005],[3.075,d]],stretch:{top:[9118,i],ext:[9119,i],bot:[9120,i]}},47:{dir:a,HW:[[0.69,m],[1.23,i],[1.353,i,1.1],[1.845,h],[2.048,h,1.11],[2.46,g],[2.472,g,1.005],[3.075,d]]},91:{dir:a,HW:[[0.818,m],[1.23,i],[1.353,i,1.1],[1.845,h],[2.048,h,1.11],[2.46,g],[2.472,g,1.005],[3.075,d]],stretch:{top:[9121,i],ext:[9122,i],bot:[9123,i]}},92:{dir:a,HW:[[0.69,m],[1.23,i],[1.353,i,1.1],[1.845,h],[2.048,h,1.11],[2.46,g],[2.472,g,1.005],[3.075,d]]},93:{dir:a,HW:[[0.818,m],[1.23,i],[1.353,i,1.1],[1.845,h],[2.048,h,1.11],[2.46,g],[2.472,g,1.005],[3.075,d]],stretch:{top:[9124,i],ext:[9125,i],bot:[9126,i]}},123:{dir:a,HW:[[0.861,m],[1.23,i],[1.353,i,1.1],[1.845,h],[2.048,h,1.11],[2.46,g],[2.472,g,1.005],[3.075,d]],stretch:{top:[9127,i],mid:[9128,i],bot:[9129,i],ext:[9130,i]}},124:{dir:a,HW:[[0.69,m]],stretch:{ext:[8739,m]}},125:{dir:a,HW:[[0.861,m],[1.23,i],[1.353,i,1.1],[1.845,h],[2.048,h,1.11],[2.46,g],[2.472,g,1.005],[3.075,d]],stretch:{top:[9131,i],mid:[9132,i],bot:[9133,i],ext:[9130,i]}},710:{dir:l,HW:[[0.333,m],[0.56,i],[0.979,h],[1.46,g],[1.886,d],[2.328,c]]},711:{dir:l,HW:[[0.333,m],[0.56,i],[0.979,h],[1.46,g],[1.886,d],[2.328,c]]},732:{dir:l,HW:[[0.333,m],[0.558,i],[0.978,h],[1.458,g],[1.886,d],[2.328,c]]},8214:{dir:a,HW:[[0.879,m]],stretch:{ext:[8214,m]}},8254:{dir:l,HW:[[0.5,m]],stretch:{rep:[8254,m]}},8592:{dir:l,HW:[[0.926,m]],stretch:{left:[8592,m],rep:k}},8593:{dir:a,HW:[[0.818,m]],stretch:{top:[8593,m],ext:[9168,m]}},8594:{dir:l,HW:[[0.926,m]],stretch:{rep:k,right:[8594,m]}},8595:{dir:a,HW:[[0.818,m]],stretch:{ext:[9168,m],bot:[8595,m]}},8596:{dir:l,HW:[[0.926,m]],stretch:{left:[8592,m],rep:k,right:[8594,m]}},8597:{dir:a,HW:[[0.818,m]],stretch:{top:[8593,m],ext:[9168,m],bot:[8595,m]}},8656:{dir:l,HW:[[0.926,m]],stretch:{left:[8656,m],rep:[61,m]}},8657:{dir:a,HW:[[0.818,m]],stretch:{top:[8657,m],ext:[8741,m,0.082]}},8658:{dir:l,HW:[[0.926,m]],stretch:{rep:[61,m],right:[8658,m]}},8659:{dir:a,HW:[[0.818,m]],stretch:{ext:[8741,m,0.082],bot:[8659,m]}},8660:{dir:l,HW:[[0.926,m]],stretch:{left:[8656,m],rep:[61,m],right:[8658,m]}},8661:{dir:a,HW:[[0.818,m]],stretch:{top:[8657,m],ext:[8741,m,0.082],bot:[8659,m]}},8730:{dir:a,HW:[[0.954,"STIXVariants"],[1.232,m],[1.847,i],[2.46,h],[3.075,g]],stretch:{top:[57345,r],ext:[57344,r],bot:[9143,i],fullExtenders:true}},8739:{dir:a,HW:[[0.879,m]],stretch:{ext:[8739,m]}},8741:{dir:a,HW:[[0.879,m]],stretch:{ext:[8741,m]}},8968:{dir:a,HW:[[0.926,m],[1.23,i],[1.353,i,1.1],[1.845,h],[2.048,h,1.11],[2.46,g],[2.472,g,1.005],[3.075,d]],stretch:{top:[9121,i],ext:[9122,i]}},8969:{dir:a,HW:[[0.926,m],[1.23,i],[1.353,i,1.1],[1.845,h],[2.048,h,1.11],[2.46,g],[2.472,g,1.005],[3.075,d]],streth:{top:[9124,i],ext:[9125,i]}},8970:{dir:a,HW:[[0.926,m],[1.23,i],[1.353,i,1.1],[1.845,h],[2.048,h,1.11],[2.46,g],[2.472,g,1.005],[3.075,d]],stretch:{ext:[9122,i],bot:[9123,i]}},8971:{dir:a,HW:[[0.926,m],[1.23,i],[1.353,i,1.1],[1.845,h],[2.048,h,1.11],[2.46,g],[2.472,g,1.005],[3.075,d]],stretch:{ext:[9125,i],bot:[9126,i]}},9130:{dir:a,HW:[[1.01,i]],stretch:{top:[9130,i],ext:[9130,i],bot:[9130,i]}},9135:{dir:l,HW:[[0.315,m]],stretch:{rep:[9135,m]}},9136:{dir:a,HW:[[1,i]],stretch:{top:[9127,i],ext:[9130,i],bot:[9133,i]}},9137:{dir:a,HW:[[1,i]],stretch:{top:[9131,i],ext:[9130,i],bot:[9129,i]}},9168:{dir:a,HW:[[0.304,m],[0.69,m,null,124],[0.879,m,null,8739]],stretch:{ext:[8739,m]}},9182:{dir:l,HW:[[0.926,i],[1,m],[1.46,h],[1.886,g],[2.328,d],[3.238,c]],stretch:{left:[57659,r],mid:[57664,r],right:[57660,r],rep:[57674,r]}},9183:{dir:l,HW:[[0.926,i],[1,m],[1.46,h],[1.886,g],[2.328,d],[3.238,c]],stretch:{left:[57661,r],mid:[57665,r],right:[57662,r],rep:[57675,r]}},10216:{dir:a,HW:[[0.926,m],[1.23,i],[1.353,i,1.1],[1.845,h],[2.048,h,1.11],[2.46,g],[2.472,g,1.005],[3.075,d]]},10217:{dir:a,HW:[[0.926,m],[1.23,i],[1.353,i,1.1],[1.845,h],[2.048,h,1.11],[2.46,g],[2.472,g,1.005],[3.075,d]]},10222:{dir:a,HW:[[0.853,m]],stretch:{top:[9127,i],ext:[9130,i],bot:[9129,i]}},10223:{dir:a,HW:[[0.853,m]],stretch:{top:[9131,i],ext:[9130,i],bot:[9133,i]}},45:{alias:9135,dir:l},94:{alias:710,dir:l},95:{alias:9135,dir:l},126:{alias:732,dir:l},175:{alias:9135,dir:l},713:{alias:9135,dir:l},770:{alias:710,dir:l},771:{alias:732,dir:l},780:{alias:711,dir:l},818:{alias:9135,dir:l},8213:{alias:9135,dir:l},8215:{alias:9135,dir:l},8407:{alias:8594,dir:l},8722:{alias:9135,dir:l},8725:{alias:47,dir:a},9001:{alias:10216,dir:a},9002:{alias:10217,dir:a},9472:{alias:8722,dir:l},10072:{alias:8739,dir:a},12296:{alias:10216,dir:a},12297:{alias:10217,dir:a},65079:{alias:9182,dir:l},65080:{alias:9183,dir:l},61:n,8606:n,8608:n,8612:n,8613:f,8614:n,8615:f,8624:f,8625:f,8636:n,8637:n,8638:f,8639:f,8640:n,8641:n,8642:f,8643:f,8666:n,8667:n,8747:f,9140:n,9141:n,9180:n,9181:n,9184:n,9185:n,10502:n,10503:n,10574:n,10575:f,10576:n,10577:f,10586:n,10587:n,10588:f,10589:f,10590:n,10591:n,10592:f,10593:f,8978:{alias:9180,dir:l},8994:{alias:9180,dir:l},8995:{alias:9181,dir:l},10229:{alias:8592,dir:l},10230:{alias:8594,dir:l},10231:{alias:8596,dir:l},10232:{alias:8656,dir:l},10233:{alias:8658,dir:l},10234:{alias:8660,dir:l},10235:{alias:8612,dir:l},10236:{alias:8614,dir:l},10237:{alias:10502,dir:l},10238:{alias:10503,dir:l},711:n,717:n,759:n,8607:f,8609:f,8616:f,8617:n,8618:n,8626:f,8627:f,8628:n,8629:f,8651:n,8652:n,8672:n,8673:f,8674:n,8675:f,8676:n,8677:n,8701:n,8702:n,8703:n,10214:f,10215:f,10218:f,10219:f,10506:f,10507:f,10514:f,10515:f,10578:n,10579:n,10580:f,10581:f,10582:n,10583:n,10584:f,10585:f,10624:f,10647:f,10648:f}}});j.FONTDATA.FONTS.STIXGeneral={directory:"General/Regular",family:"STIXGeneral",Ranges:[[160,255,"Latin1Supplement"],[256,383,"LatinExtendedA"],[384,591,"LatinExtendedB"],[592,687,"IPAExtensions"],[688,767,"SpacingModLetters"],[768,879,"CombDiacritMarks"],[880,1023,"GreekAndCoptic"],[1024,1279,"Cyrillic"],[7424,7615,"PhoneticExtensions"],[7680,7935,"LatinExtendedAdditional"],[8192,8303,"GeneralPunctuation"],[8304,8351,"SuperAndSubscripts"],[8352,8399,"CurrencySymbols"],[8400,8447,"CombDiactForSymbols"],[8448,8527,"LetterlikeSymbols"],[8528,8591,"NumberForms"],[8592,8703,"Arrows"],[8704,8959,"MathOperators"],[8960,9215,"MiscTechnical"],[9216,9279,"ControlPictures"],[9312,9471,"EnclosedAlphanum"],[9472,9599,"BoxDrawing"],[9600,9631,"BlockElements"],[9632,9727,"GeometricShapes"],[9728,9983,"MiscSymbols"],[9984,10175,"Dingbats"],[10176,10223,"MiscMathSymbolsA"],[10224,10239,"SupplementalArrowsA"],[10496,10623,"SupplementalArrowsB"],[10624,10751,"MiscMathSymbolsB"],[10752,11007,"SuppMathOperators"],[11008,11263,"MiscSymbolsAndArrows"],[12288,12351,"CJK"],[12352,12447,"Hiragana"],[42784,43007,"LatinExtendedD"],[64256,64335,"AlphaPresentForms"],[65520,65535,"Specials"],[119808,119859,"MathBold"],[119860,119911,"MathItalic"],[119912,119963,"MathBoldItalic"],[119964,120015,"MathScript"],[120016,120067,"MathBoldScript"],[120068,120119,"Fraktur"],[120120,120171,"BBBold"],[120172,120223,"BoldFraktur"],[120224,120275,"MathSS"],[120276,120327,"MathSSBold"],[120328,120379,"MathSSItalic"],[120380,120431,"MathSSItalicBold"],[120432,120483,"MathTT"],[120484,120485,"ij"],[120488,120545,"GreekBold"],[120546,120603,"GreekItalic"],[120604,120661,"GreekBoldItalic"],[120662,120719,"GreekSSBold"],[120720,120777,"GreekSSBoldItalic"],[120782,120791,"MathBold"],[120792,120801,"BBBold"],[120802,120811,"MathSS"],[120812,120822,"MathSSBold"],[120822,120831,"MathTT"]],32:[0,0,250,0,0],33:[676,9,333,130,236],34:[676,-431,408,77,331],35:[662,0,500,6,495],36:[727,87,500,44,458],37:[706,19,747,61,686],38:[676,13,778,42,750],39:[676,-431,180,48,133],40:[676,177,333,48,304],41:[676,177,333,29,285],42:[676,-265,500,68,433],43:[547,41,685,48,636],44:[102,141,250,55,195],45:[257,-194,333,39,285],46:[100,11,250,70,181],47:[676,14,278,-9,287],48:[676,14,500,24,476],49:[676,0,500,111,394],50:[676,0,500,29,474],51:[676,14,500,41,431],52:[676,0,500,12,473],53:[688,14,500,31,438],54:[684,14,500,34,468],55:[662,8,500,20,449],56:[676,14,500,56,445],57:[676,22,500,30,459],58:[459,11,278,81,192],59:[459,141,278,80,219],60:[534,24,685,56,621],61:[386,-120,685,48,637],62:[534,24,685,56,621],63:[676,8,444,68,414],64:[676,14,921,116,809],65:[674,0,722,15,707],66:[662,0,667,17,593],67:[676,14,667,28,633],68:[662,0,722,16,685],69:[662,0,611,12,597],70:[662,0,556,11,546],71:[676,14,722,32,709],72:[662,0,722,18,703],73:[662,0,333,18,315],74:[662,14,373,-6,354],75:[662,0,722,33,723],76:[662,0,611,12,598],77:[662,0,889,12,864],78:[662,11,722,12,707],79:[676,14,722,34,688],80:[662,0,557,16,542],81:[676,177,722,34,701],82:[662,0,667,17,660],83:[676,14,556,43,491],84:[662,0,611,17,593],85:[662,14,722,14,705],86:[662,11,722,16,697],87:[662,11,944,5,932],88:[662,0,722,10,704],89:[662,0,722,22,703],90:[662,0,612,10,598],91:[662,156,333,88,299],92:[676,14,278,-9,287],93:[662,156,333,34,245],94:[662,-297,469,24,446],95:[-75,125,500,0,500],96:[678,-507,333,18,242],97:[460,10,444,37,442],98:[683,10,500,3,468],99:[460,10,444,25,412],100:[683,10,500,27,491],101:[460,10,444,25,424],102:[683,0,333,20,383],103:[460,218,500,28,470],104:[683,0,500,9,487],105:[683,0,278,16,253],106:[683,218,278,-70,194],107:[683,0,500,7,505],108:[683,0,278,19,257],109:[460,0,778,16,775],110:[460,0,500,16,485],111:[460,10,500,29,470],112:[460,217,500,5,470],113:[460,217,500,24,488],114:[460,0,333,5,335],115:[459,10,389,51,348],116:[579,10,278,13,279],117:[450,10,500,9,480],118:[450,14,500,19,477],119:[450,14,722,21,694],120:[450,0,500,17,479],121:[450,218,500,14,475],122:[450,0,444,27,418],123:[680,181,480,100,350],124:[676,14,200,67,133],125:[680,181,480,130,380],126:[325,-183,541,40,502],160:[0,0,250,0,0],168:[622,-523,333,18,316],172:[393,-115,600,48,552],175:[601,-547,333,11,322],177:[502,87,685,48,637],183:[310,-199,250,70,181],215:[529,25,640,43,597],247:[516,10,564,30,534],305:[460,0,278,16,253],567:[460,218,278,-70,193],710:[674,-507,333,11,322],711:[674,-507,333,11,322],713:[601,-547,334,11,322],714:[679,-509,333,93,320],715:[679,-509,333,22,249],728:[664,-507,335,27,308],729:[622,-523,333,118,217],732:[638,-532,333,1,331],768:[678,-507,0,-371,-147],769:[678,-507,0,-371,-147],770:[674,-507,0,-386,-75],771:[638,-532,0,-395,-65],772:[601,-547,0,-385,-74],774:[664,-507,0,-373,-92],775:[622,-523,0,-280,-181],776:[622,-523,0,-379,-81],778:[711,-512,0,-329,-130],779:[678,-507,0,-401,-22],780:[674,-507,0,-385,-74],824:[662,156,0,-380,31],915:[662,0,587,11,577],916:[674,0,722,48,675],920:[676,14,722,34,688],923:[674,0,702,15,687],926:[662,0,643,29,614],928:[662,0,722,18,703],931:[662,0,624,30,600],933:[674,0,722,29,703],934:[662,0,763,35,728],936:[690,0,746,22,724],937:[676,0,744,29,715],8224:[676,149,500,59,442],8225:[676,153,500,58,442],8230:[100,11,1000,111,888],8242:[678,-402,289,75,214],8254:[820,-770,500,0,500],8407:[760,-548,0,-453,-17],8465:[695,34,762,45,711],8472:[547,217,826,52,799],8476:[704,22,874,50,829],8501:[677,13,682,43,634],8592:[449,-58,926,71,857],8593:[662,156,511,60,451],8594:[448,-57,926,70,856],8595:[662,156,511,60,451],8596:[449,-57,926,38,888],8597:[730,224,511,60,451],8598:[662,156,926,70,856],8599:[662,156,926,70,856],8600:[662,156,926,70,856],8601:[662,156,926,70,856],8614:[450,-57,926,70,857],8617:[553,-57,926,70,856],8618:[553,-57,926,70,856],8636:[494,-220,955,54,901],8637:[286,-12,955,54,901],8640:[494,-220,955,54,901],8641:[286,-12,955,54,901],8652:[539,33,926,70,856],8656:[551,45,926,60,866],8657:[662,156,685,45,641],8658:[551,45,926,60,866],8659:[662,156,685,45,641],8660:[517,10,926,20,906],8661:[730,224,685,45,641],8704:[662,0,560,2,558],8706:[668,11,471,40,471],8707:[662,0,560,73,487],8709:[583,79,762,50,712],8711:[662,12,731,63,667],8712:[531,27,685,60,625],8713:[662,157,685,60,625],8715:[531,27,685,60,625],8719:[763,259,1000,52,948],8720:[763,259,1000,52,948],8721:[763,259,914,58,856],8722:[286,-220,685,64,621],8723:[502,87,685,48,637],8725:[710,222,523,46,478],8726:[411,-93,428,25,403],8727:[471,-33,523,67,457],8728:[387,-117,350,40,310],8729:[387,-117,350,40,310],8730:[973,259,928,112,963],8733:[430,0,685,41,643],8734:[430,0,926,70,854],8736:[547,0,685,23,643],8739:[690,189,266,100,166],8741:[690,189,523,129,394],8743:[536,29,620,31,589],8744:[536,29,620,31,589],8745:[536,31,620,48,572],8746:[536,31,620,48,572],8747:[824,320,459,32,639],8764:[362,-148,685,48,637],8768:[547,42,286,35,249],8771:[445,-55,685,48,637],8773:[532,27,685,48,637],8776:[475,-25,685,48,637],8781:[498,-8,685,48,637],8784:[611,-120,685,48,637],8800:[662,156,685,48,637],8801:[478,-28,685,48,637],8804:[609,103,685,64,629],8805:[609,103,685,64,629],8810:[532,26,933,25,908],8811:[532,26,933,25,908],8826:[532,26,685,64,621],8827:[532,26,685,64,621],8828:[628,120,685,64,621],8829:[629,119,685,64,621],8834:[531,25,685,64,621],8835:[531,25,685,64,621],8838:[607,103,685,64,621],8839:[607,103,685,64,621],8846:[536,31,620,48,572],8849:[607,103,685,64,621],8850:[607,103,685,64,621],8851:[536,31,620,48,572],8852:[536,31,620,48,572],8853:[623,119,842,50,792],8854:[623,119,842,50,792],8855:[623,119,842,50,792],8856:[623,119,842,50,792],8857:[583,79,762,50,712],8866:[662,0,685,64,621],8867:[662,0,685,64,621],8868:[662,0,685,48,637],8869:[662,0,685,48,637],8872:[662,0,685,64,621],8896:[763,259,924,54,870],8897:[763,259,924,54,870],8898:[778,254,924,94,830],8899:[768,264,924,94,830],8900:[488,-16,523,26,497],8901:[313,-193,286,83,203],8902:[597,13,700,35,665],8904:[582,80,810,54,756],8942:[606,104,511,192,319],8943:[316,-189,926,108,818],8945:[520,18,926,194,732],8968:[713,213,469,188,447],8969:[713,213,469,27,286],8970:[713,213,469,188,447],8971:[713,213,469,27,286],8994:[360,-147,1019,54,965],8995:[360,-147,1019,54,965],9135:[286,-220,315,0,315],9168:[405,-101,511,222,288],9651:[811,127,1145,35,1110],9657:[555,50,660,80,605],9661:[811,127,1145,35,1110],9667:[554,51,660,55,580],9711:[785,282,1207,70,1137],9824:[609,99,685,34,651],9825:[603,105,685,34,651],9826:[609,105,685,41,643],9827:[603,99,685,34,651],9837:[768,10,426,57,346],9838:[768,181,426,75,350],9839:[768,181,426,41,386],10216:[713,213,400,77,335],10217:[713,213,400,65,323],10222:[676,177,233,56,211],10223:[676,177,233,22,177],10229:[449,-58,1574,55,1519],10230:[449,-57,1574,55,1519],10231:[449,-57,1574,55,1519],10232:[551,45,1574,55,1519],10233:[551,45,1574,55,1519],10234:[517,10,1574,55,1519],10235:[450,-57,1574,55,1519],10236:[450,-57,1574,55,1519],10741:[710,222,523,46,478],10752:[763,259,1126,53,1073],10753:[763,259,1126,53,1073],10754:[763,259,1126,53,1073],10755:[768,264,924,94,830],10756:[768,264,924,94,830],10757:[763,259,924,94,830],10758:[763,259,924,94,830],10815:[662,0,694,30,664],10927:[609,103,685,64,621],10928:[609,103,685,64,621]};j.FONTDATA.FONTS["STIXGeneral-bold"]={directory:"General/Bold",family:"STIXGeneral",weight:"bold",Ranges:[[160,255,"Latin1Supplement"],[256,383,"LatinExtendedA"],[384,591,"LatinExtendedB"],[592,687,"IPAExtensions"],[688,767,"SpacingModLetters"],[768,879,"CombDiacritMarks"],[880,1023,"GreekAndCoptic"],[1024,1279,"Cyrillic"],[7424,7615,"PhoneticExtensions"],[7680,7935,"LatinExtendedAdditional"],[8192,8303,"GeneralPunctuation"],[8304,8351,"SuperAndSubscripts"],[8352,8399,"CurrencySymbols"],[8400,8447,"CombDiactForSymbols"],[8448,8527,"LetterlikeSymbols"],[8528,8591,"NumberForms"],[8592,8703,"Arrows"],[8704,8959,"MathOperators"],[8960,9215,"MiscTechnical"],[9216,9279,"ControlPictures"],[9312,9471,"EnclosedAlphanum"],[9472,9599,"BoxDrawing"],[9632,9727,"GeometricShapes"],[9728,9983,"MiscSymbols"],[10176,10223,"MiscMathSymbolsA"],[10624,10751,"MiscMathSymbolsB"],[10752,11007,"SuppMathOperators"],[42784,43007,"LatinExtendedD"],[64256,64335,"AlphaPresentForms"],[119808,119859,"MathBold"],[120120,120171,"BBBold"],[120172,120223,"BoldFraktur"],[120276,120327,"MathSSBold"],[120488,120545,"GreekBold"],[120662,120719,"GreekSSBold"],[120782,120791,"MathBold"],[120812,120822,"MathSSBold"]],32:[0,0,250,0,0],33:[691,13,333,81,251],34:[691,-404,555,83,472],35:[700,0,500,5,495],36:[750,99,500,29,472],37:[706,29,749,61,688],38:[691,16,833,62,789],39:[691,-404,278,75,204],40:[694,168,333,46,306],41:[694,168,333,27,287],42:[691,-255,500,56,448],43:[563,57,750,65,685],44:[155,180,250,39,223],45:[287,-171,333,44,287],46:[156,13,250,41,210],47:[691,19,278,-24,302],48:[688,13,500,24,476],49:[688,0,500,65,441],50:[688,0,500,17,478],51:[688,14,500,16,468],52:[688,0,500,19,476],53:[676,8,500,22,470],54:[688,13,500,28,475],55:[676,0,500,17,477],56:[688,13,500,28,472],57:[688,13,500,26,473],58:[472,13,333,82,251],59:[472,180,333,82,266],60:[534,24,750,80,670],61:[399,-107,750,68,682],62:[534,24,750,80,670],63:[689,13,500,57,445],64:[691,19,930,108,822],65:[690,0,722,9,689],66:[676,0,667,16,619],67:[691,19,722,49,687],68:[676,0,722,14,690],69:[676,0,667,16,641],70:[676,0,611,16,583],71:[691,19,778,37,755],72:[676,0,778,21,759],73:[676,0,389,20,370],74:[676,96,500,3,478],75:[676,0,778,30,769],76:[677,0,667,19,638],77:[676,0,944,14,921],78:[676,18,722,16,701],79:[691,19,778,35,743],80:[676,0,611,16,600],81:[691,176,778,35,743],82:[676,0,722,26,716],83:[692,19,556,35,513],84:[676,0,667,31,636],85:[676,19,722,16,701],86:[676,18,722,16,701],87:[676,15,1000,19,981],88:[676,0,722,16,699],89:[676,0,722,15,699],90:[676,0,667,28,634],91:[678,149,333,67,301],92:[691,19,278,-25,303],93:[678,149,333,32,266],94:[676,-311,581,73,509],95:[-75,125,500,0,500],96:[713,-528,333,8,246],97:[473,14,500,25,488],98:[676,14,556,17,521],99:[473,14,444,25,430],100:[676,14,556,25,534],101:[473,14,444,25,427],102:[691,0,333,14,389],103:[473,206,500,28,483],104:[676,0,556,15,534],105:[691,0,278,15,256],106:[691,203,333,-57,263],107:[676,0,556,22,543],108:[676,0,278,15,256],109:[473,0,833,15,814],110:[473,0,556,21,539],111:[473,14,500,25,476],112:[473,205,556,19,524],113:[473,205,556,34,536],114:[473,0,444,28,434],115:[473,14,389,25,361],116:[630,12,333,19,332],117:[461,14,556,16,538],118:[461,14,500,21,485],119:[461,14,722,23,707],120:[461,0,500,12,484],121:[461,205,500,16,482],122:[461,0,444,21,420],123:[698,175,394,22,340],124:[691,19,220,66,154],125:[698,175,394,54,372],126:[333,-173,520,29,491],915:[676,0,620,16,593],916:[690,0,722,33,673],920:[692,18,778,35,743],923:[690,0,707,9,674],926:[676,0,647,40,607],928:[676,0,778,21,759],931:[676,0,671,28,641],933:[692,0,703,7,693],934:[676,0,836,18,818],936:[692,0,808,15,797],937:[692,0,768,28,740]};j.FONTDATA.FONTS["STIXGeneral-italic"]={directory:"General/Italic",family:"STIXGeneral",style:"italic",Ranges:[[160,255,"Latin1Supplement"],[256,383,"LatinExtendedA"],[384,591,"LatinExtendedB"],[592,687,"IPAExtensions"],[688,767,"SpacingModLetters"],[880,1023,"GreekAndCoptic"],[1024,1279,"Cyrillic"],[7680,7935,"LatinExtendedAdditional"],[8192,8303,"GeneralPunctuation"],[8352,8399,"CurrencySymbols"],[8400,8447,"CombDiactForSymbols"],[8448,8527,"LetterlikeSymbols"],[8704,8959,"MathOperators"],[9216,9279,"ControlPictures"],[9312,9471,"EnclosedAlphanum"],[9472,9599,"BoxDrawing"],[64256,64335,"AlphaPresentForms"],[119860,119911,"MathItalic"],[119964,120015,"MathScript"],[120328,120379,"MathSSItalic"],[120484,120485,"ij"],[120546,120603,"GreekItalic"]],32:[0,0,250,0,0],33:[667,11,333,39,304],34:[666,-421,420,144,432],35:[676,0,501,2,540],36:[731,89,500,32,497],37:[706,19,755,80,705],38:[666,18,778,76,723],39:[666,-421,214,132,241],40:[669,181,333,42,315],41:[669,180,333,16,289],42:[666,-255,500,128,492],43:[506,0,675,86,590],44:[101,129,250,-5,135],45:[255,-192,333,49,282],46:[100,11,250,27,138],47:[666,18,278,-65,386],48:[676,7,500,32,497],49:[676,0,500,50,409],50:[676,0,500,12,452],51:[676,7,500,16,465],52:[676,0,500,1,479],53:[666,7,500,15,491],54:[686,7,500,30,521],55:[666,8,500,75,537],56:[676,7,500,30,493],57:[676,17,500,23,492],58:[441,11,333,50,261],59:[441,129,333,26,261],60:[516,10,675,84,592],61:[386,-120,675,86,590],62:[516,10,675,84,592],63:[664,12,500,132,472],64:[666,18,920,118,806],65:[668,0,611,-51,564],66:[653,0,611,-8,588],67:[666,18,667,66,689],68:[653,0,722,-8,700],69:[653,0,611,-1,634],70:[653,0,611,8,645],71:[666,18,722,52,722],72:[653,0,722,-8,769],73:[653,0,333,-8,384],74:[653,18,444,-6,491],75:[653,0,667,7,722],76:[653,0,556,-8,559],77:[653,0,833,-18,872],78:[653,15,667,-20,727],79:[667,18,722,60,699],80:[653,0,611,0,605],81:[666,182,722,59,699],82:[653,0,611,-13,588],83:[667,18,500,17,508],84:[653,0,556,59,633],85:[653,18,722,102,765],86:[653,18,611,76,688],87:[653,18,833,71,906],88:[653,0,611,-29,655],89:[653,0,556,78,633],90:[653,0,556,-6,606],91:[663,153,389,21,391],92:[666,18,278,-41,319],93:[663,153,389,12,382],94:[666,-301,422,0,422],95:[-75,125,500,0,500],96:[664,-492,333,120,311],97:[441,11,501,17,476],98:[683,11,500,23,473],99:[441,11,444,30,425],100:[683,13,500,15,527],101:[441,11,444,31,412],102:[678,207,278,-147,424],103:[441,206,500,8,471],104:[683,9,500,19,478],105:[654,11,278,49,264],106:[652,207,278,-124,279],107:[683,11,444,14,461],108:[683,11,278,41,279],109:[441,9,722,12,704],110:[441,9,500,14,474],111:[441,11,500,27,468],112:[441,205,504,-75,472],113:[441,209,500,25,484],114:[441,0,389,45,412],115:[442,13,389,16,366],116:[546,11,278,38,296],117:[441,11,500,42,475],118:[441,18,444,20,426],119:[441,18,667,15,648],120:[441,11,444,-27,447],121:[441,206,444,-24,426],122:[428,81,389,-2,380],123:[687,177,400,51,407],124:[666,18,275,105,171],125:[687,177,400,-7,349],126:[323,-183,541,40,502],305:[441,11,278,47,235],567:[441,207,278,-124,246],915:[653,0,611,8,645],916:[668,0,611,-32,526],920:[667,18,722,60,699],923:[668,0,611,-51,564],926:[653,0,651,-6,680],928:[653,0,722,-8,769],931:[653,0,620,-6,659],933:[668,0,556,78,648],934:[653,0,741,50,731],936:[667,0,675,77,778],937:[666,0,762,-6,739],945:[441,11,552,27,549],946:[678,205,506,-40,514],947:[435,206,410,19,438],948:[668,11,460,24,460],949:[441,11,444,30,425],950:[683,185,454,30,475],951:[441,205,474,14,442],952:[678,11,480,27,494],953:[441,11,278,49,235],954:[441,13,444,14,465],955:[678,16,458,-12,431],956:[428,205,526,-33,483],957:[441,18,470,20,459],958:[683,185,454,30,446],959:[441,11,500,27,468],960:[428,18,504,19,536],961:[441,205,504,-40,471],962:[441,185,454,30,453],963:[428,11,498,27,531],964:[428,11,410,12,426],965:[441,10,478,19,446],966:[441,205,622,27,590],967:[441,207,457,-108,498],968:[441,205,584,15,668],969:[439,11,686,27,654],977:[678,10,556,19,526],981:[683,205,627,27,595],982:[428,11,792,17,832],1009:[441,205,516,27,484],1013:[441,11,444,30,420],8467:[687,11,579,48,571]};j.FONTDATA.FONTS.STIXIntegralsD={directory:"IntegralsD/Regular",family:"STIXIntegralsD",Ranges:[[32,32,"All"],[160,160,"All"],[8747,8755,"All"],[10763,10780,"All"]],8747:[2000,269,585,56,1035],8750:[2000,269,635,56,1035]};j.FONTDATA.FONTS.STIXNonUnicode={directory:"NonUnicode/Regular",family:"STIXNonUnicode",Ranges:[[32,32,"All"],[160,160,"All"],[57344,63743,"PrivateUse"]],57344:[610,25,1184,829,895],57345:[667,-41,1184,829,1211],57656:[634,-584,480,-10,490],57657:[-127,177,480,-10,490],57659:[955,-512,897,-25,908],57660:[955,-512,897,-11,922],57661:[182,261,897,-25,908],57662:[182,261,897,-11,922],57664:[1218,-820,1844,-10,1854],57665:[-126,524,1844,-10,1854],57674:[955,-820,633,-1,634],57675:[-126,261,633,-1,634],57953:[422,10,523,41,481],57957:[421,0,523,127,405],57961:[421,0,523,68,455],57965:[424,198,523,47,463],57969:[420,198,523,58,480],57973:[421,198,523,66,457],57977:[612,8,523,37,486],57981:[421,198,523,25,490],57985:[606,12,523,47,477],57989:[421,200,523,41,483],57999:[135,0,325,-1,326],58000:[135,0,633,-1,634]};j.FONTDATA.FONTS["STIXNonUnicode-bold"]={directory:"NonUnicode/Bold",family:"STIXNonUnicode",weight:"bold",Ranges:[[32,32,"All"],[160,160,"All"],[57344,63743,"PrivateUse"]]};j.FONTDATA.FONTS["STIXNonUnicode-italic"]={directory:"NonUnicode/Italic",family:"STIXNonUnicode",style:"italic",Ranges:[[32,32,"All"],[160,160,"All"],[57344,63743,"PrivateUse"]],57901:[677,45,852,43,812],57902:[670,3,724,35,709],57903:[671,11,569,43,586],57904:[662,0,801,34,788],57905:[670,4,553,40,599],57906:[662,0,652,43,710],57907:[671,131,580,40,580],57908:[664,21,831,41,845],57909:[662,0,575,38,591],57910:[662,120,632,31,785],57911:[670,13,809,30,783],57912:[670,7,693,30,653],57913:[671,45,1166,40,1128],57914:[795,37,957,40,1064],57915:[669,10,737,38,729],57916:[662,0,667,38,709],57917:[671,131,744,43,704],57918:[662,3,854,38,816],57919:[671,0,634,38,671],57920:[721,0,509,41,730],57921:[672,13,817,37,950],57922:[677,33,638,33,680],57923:[685,32,956,33,998],57924:[672,13,692,38,739],57925:[675,131,719,34,763],57926:[664,94,752,38,714]};j.FONTDATA.FONTS.STIXSizeOneSym={directory:"SizeOneSym/Regular",family:"STIXSizeOneSym",Ranges:[[688,767,"All"],[768,824,"All"],[8254,8254,"All"],[8400,8431,"All"],[8512,8512,"All"],[8730,8732,"All"],[8992,8993,"All"],[9115,9145,"All"],[9180,9185,"All"],[10098,10099,"All"],[10214,10219,"All"],[10627,10630,"All"],[10744,10745,"All"],[10752,10762,"All"],[11004,11007,"All"]],32:[0,0,250,0,0],40:[1066,164,468,139,382],41:[1066,164,468,86,329],47:[1066,164,579,25,552],91:[1066,164,383,180,363],92:[1066,164,579,27,552],93:[1066,164,383,20,203],95:[-127,177,1000,0,1000],123:[1066,164,575,114,466],125:[1066,164,575,109,461],160:[0,0,250,0,0],770:[767,-554,0,-720,-160],771:[750,-598,0,-722,-162],8719:[1500,-49,1355,50,1305],8720:[1500,-49,1355,50,1305],8721:[1499,-49,1292,90,1202],8730:[1552,295,1057,112,1089],8896:[1500,-49,1265,60,1205],8897:[1500,-49,1265,60,1205],8898:[1510,-49,1265,118,1147],8899:[1500,-39,1265,118,1147],8968:[1066,164,453,180,426],8969:[1066,164,453,25,273],8970:[1066,164,453,180,428],8971:[1066,164,453,27,273],9115:[700,305,450,50,400],9116:[705,305,450,50,174],9117:[705,300,450,50,400],9118:[700,305,450,50,400],9119:[705,305,450,276,400],9120:[705,300,450,50,400],9121:[682,323,450,50,415],9122:[687,323,450,50,150],9123:[687,318,450,50,415],9124:[682,323,450,35,400],9125:[687,323,450,300,400],9126:[687,318,450,35,400],9127:[700,305,640,260,600],9128:[705,305,640,40,380],9129:[705,300,640,260,600],9130:[705,305,640,260,380],9131:[700,305,640,40,380],9132:[705,305,640,260,600],9133:[705,300,640,40,380],9134:[610,25,688,294,394],9136:[700,301,600,35,566],9137:[700,301,600,35,566],9143:[1510,345,1184,112,895],9144:[1566,289,721,0,66],9145:[1566,289,721,655,721],9182:[136,89,926,0,925],9183:[789,-564,926,0,925],10216:[1066,164,578,116,462],10217:[1066,164,578,116,462],10752:[1500,-49,1555,52,1503],10753:[1500,-49,1555,52,1503],10754:[1500,-49,1555,52,1503],10756:[1500,-39,1265,118,1147],10757:[1500,-49,1153,82,1071],10758:[1500,-49,1153,82,1071]};j.FONTDATA.FONTS.STIXSizeTwoSym={directory:"SizeTwoSym/Regular",family:"STIXSizeTwoSym",Ranges:[[688,767,"All"],[768,824,"All"],[8254,8254,"All"],[8400,8431,"All"],[8730,8732,"All"],[9115,9145,"All"],[9180,9185,"All"],[10098,10099,"All"],[10214,10219,"All"],[10627,10630,"All"],[11004,11007,"All"]],32:[0,0,250,0,0],40:[1566,279,589,139,503],41:[1566,279,608,114,478],47:[1566,279,806,25,781],91:[1566,279,459,190,422],92:[1566,279,806,25,781],93:[1566,279,459,37,269],95:[-127,177,1500,0,1500],123:[1566,279,717,124,531],125:[1566,279,717,186,593],160:[0,0,250,0,0],770:[777,-564,0,-1150,-171],771:[760,-608,0,-1152,-173],8730:[2056,404,1124,110,1157],8968:[1566,279,524,190,479],8969:[1566,279,526,47,336],8970:[1566,279,524,190,479],8971:[1566,279,526,47,336],9182:[143,81,1460,0,1460],9183:[797,-573,1460,0,1460],10216:[1566,279,622,95,531],10217:[1566,279,622,91,527]};j.FONTDATA.FONTS.STIXSizeThreeSym={directory:"SizeThreeSym/Regular",family:"STIXSizeThreeSym",Ranges:[[688,767,"All"],[768,824,"All"],[8254,8254,"All"],[8400,8431,"All"],[8730,8732,"All"],[9115,9145,"All"],[9180,9185,"All"],[10098,10099,"All"],[10214,10219,"All"],[10627,10630,"All"]],32:[0,0,250,0,0],40:[2066,394,750,182,667],41:[2066,394,750,83,568],47:[2066,394,1101,30,1071],91:[2066,394,508,225,491],92:[2066,394,1101,30,1071],93:[2066,394,508,17,283],95:[-127,177,2000,0,2000],123:[2066,394,906,143,717],125:[2066,394,906,189,763],160:[0,0,250,0,0],770:[777,-564,0,-1610,-150],771:[774,-608,0,-1612,-152],8730:[2565,510,1076,112,1110],8968:[2066,394,565,225,550],8969:[2066,394,565,15,340],8970:[2066,394,565,225,550],8971:[2066,394,565,15,340],9182:[157,86,1886,0,1886],9183:[815,-572,1886,0,1886],10216:[2066,394,765,96,670],10217:[2066,394,765,95,669]};j.FONTDATA.FONTS.STIXSizeFourSym={directory:"SizeFourSym/Regular",family:"STIXSizeFourSym",Ranges:[[688,767,"All"],[768,824,"All"],[8254,8254,"All"],[8400,8431,"All"],[8730,8732,"All"],[9115,9145,"All"],[9180,9185,"All"],[10098,10099,"All"],[10214,10219,"All"],[10627,10630,"All"]],32:[0,0,250,0,0],40:[2566,509,808,124,732],41:[2566,509,808,76,684],47:[2566,509,1309,16,1293],91:[2566,509,661,295,634],92:[2566,509,1309,16,1293],93:[2566,509,661,27,366],95:[-127,177,2500,0,2500],123:[2566,509,1076,173,882],125:[2566,509,1076,194,903],160:[0,0,250,0,0],770:[796,-573,0,-2040,-154],771:[771,-608,0,-2040,-154],8730:[1510,345,1184,112,895],8968:[2566,509,682,295,655],8969:[2566,509,682,27,387],8970:[2566,509,682,295,655],8971:[2566,509,682,27,387],9182:[175,90,2328,0,2328],9183:[837,-572,2328,0,2328],10216:[2566,509,908,113,796],10217:[2566,509,908,112,795]};j.FONTDATA.FONTS.STIXSizeFiveSym={directory:"SizeFiveSym/Regular",family:"STIXSizeFiveSym",Ranges:[[688,767,"All"],[768,824,"All"],[8254,8254,"All"],[8400,8431,"All"],[9115,9145,"All"],[9180,9185,"All"]],32:[0,0,250,0,0],95:[-127,177,3000,0,3000],160:[0,0,250,0,0],770:[816,-572,0,-2485,-157],771:[780,-617,0,-2485,-157],9182:[181,90,3238,0,3238],9183:[844,-573,3238,0,3238]};j.FONTDATA.FONTS.STIXVariants={directory:"Variants/Regular",family:"STIXVariants",Ranges:[[32,32,"All"],[119,124,"All"],[160,160,"All"],[411,411,"All"],[612,612,"All"],[8242,8279,"All"],[8512,8512,"All"],[8592,8595,"All"],[8657,8674,"All"],[8709,8941,"All"],[8994,8995,"All"],[9251,9251,"All"],[9641,9641,"All"],[10812,10990,"All"]],8242:[565,-28,340,44,295],8463:[683,10,579,47,547],8709:[729,74,523,28,502],8726:[710,222,523,46,478],8730:[943,11,737,67,767]};j.FONTDATA.FONTS.STIXGeneral[8722][0]=j.FONTDATA.FONTS.STIXGeneral[43][0];j.FONTDATA.FONTS.STIXGeneral[8722][1]=j.FONTDATA.FONTS.STIXGeneral[43][1];j.FONTDATA.FONTS.STIXGeneral[8942][0]+=400;j.FONTDATA.FONTS.STIXGeneral[8945][0]+=500;j.FONTDATA.FONTS.STIXGeneral[8722][1]+=100;j.FONTDATA.FONTS.STIXGeneral[61][1]+=100;j.FONTDATA.FONTS.STIXNonUnicode[57674][0]+=200;j.FONTDATA.FONTS.STIXNonUnicode[57674][1]+=200;j.FONTDATA.FONTS.STIXNonUnicode[57675][0]+=200;j.FONTDATA.FONTS.STIXNonUnicode[57675][1]+=200;MathJax.Hub.Register.LoadHook(j.fontDir+"/Variants/Regular/All.js",function(){j.FONTDATA.FONTS.STIXVariants[32][2]+=275;j.FONTDATA.FONTS.STIXVariants[32][5]={rfix:275}});MathJax.Hub.Insert(j.FONTDATA.FONTS.STIXGeneral,{8192:[0,0,500,0,0,{space:1}],8193:[0,0,1000,0,0,{space:1}],8194:[0,0,500,0,0,{space:1}],8195:[0,0,1000,0,0,{space:1}],8196:[0,0,333,0,0,{space:1}],8197:[0,0,250,0,0,{space:1}],8198:[0,0,167,0,0,{space:1}],8201:[0,0,167,0,0,{space:1}],8202:[0,0,83,0,0,{space:1}],8203:[0,0,0,0,0,{space:1}]});MathJax.Hub.Browser.Select({MSIE:function(w){if(!w.versionAtLeast("8.0")||document.documentMode<8){var x=j.FONTDATA;x.REMAP[713]=175;x.REMAP[714]=180;x.REMAP[715]=96;x.REMAP[730]=176;x.RANGES[5]=x.RANGES[4];x.RANGES[4]=x.RANGES[3];x.RANGES[3]={name:"greek",low:945,high:1014,offset:"GG",remap:{1013:26,977:27,1008:28,981:29,1009:30,982:31}};x.VARIANT.bold.offsetGG=120514;x.VARIANT.bold.offsetG=120488;x.VARIANT.italic.offsetGG=120572;x.VARIANT.italic.offsetG=120546;x.VARIANT["bold-italic"].offsetGG=120630;x.VARIANT["bold-italic"].offsetG=120604}},Safari:function(w){w.STIXfontBug=w.versionAtLeast("5.1")&&w.isMac},Chrome:function(x){if(x.isMac){var w=navigator.appVersion.match(/AppleWebKit\/(\d+)/);if(w&&parseInt(w[1])>534){x.STIXfontBug=true}}}});if(MathJax.Hub.Browser.STIXfontBug){j.FONTDATA.FONTS.STIXGeneral.family="STIXGeneral-Regular";j.FONTDATA.FONTS["STIXGeneral-italic"].family="STIXGeneral-Italic";delete j.FONTDATA.FONTS["STIXGeneral-italic"].style;j.FONTDATA.FONTS.STIXNonUnicode.family="STIXNonUnicode-Regular";j.FONTDATA.FONTS["STIXNonUnicode-italic"].family="STIXNonUnicode-Italic";delete j.FONTDATA.FONTS["STIXNonUnicode-italic"].style}var e=[];var o=j.Font.div;s.addElement(o,"span",{style:{display:"inline-block","font-family":"STIXNonUnicode","font-weight":"bold"}},["\uE38C\uE38C\uE38C\uE38C\uE38C"]);s.addElement(o,"span",{style:{display:"inline-block","font-family":"STIXNonUnicode","font-weight":"bold"}},["\uE39A\uE39A\uE39A\uE39A\uE39A"]);if(o.lastChild.previousSibling.offsetWidth<o.lastChild.offsetWidth){e.push(["Require",MathJax.Ajax,j.fontDir+"/fontdata-1.0.js"])}o.removeChild(o.lastChild);o.removeChild(o.lastChild);if(!j.Font.testFont({family:"STIXSizeOneSym",testString:"\u02C6"})){e.push(["Require",MathJax.Ajax,j.fontDir+"/fontdata-beta.js"])}e.push(["loadComplete",MathJax.Ajax,j.fontDir+"/fontdata.js"]);MathJax.Callback.Queue.apply(MathJax.Callback,e)})(MathJax.OutputJax["HTML-CSS"],MathJax.ElementJax.mml,MathJax.HTML);
```
