## Supplementary material for "Artistoo: build, share, and explore simulations of cells and tissues in the web browser": All.js

```
/*
 *  /MathJax/jax/output/HTML-CSS/fonts/STIX/Variants/Regular/All.js
 *
 *  Copyright (c) 2009-2016 The MathJax Consortium
 *
 *  Licensed under the Apache License, Version 2.0 (the "License");
 *  you may not use this file except in compliance with the License.
 *  You may obtain a copy of the License at
 *
 *      http://www.apache.org/licenses/LICENSE-2.0
 *
 *  Unless required by applicable law or agreed to in writing, software
 *  distributed under the License is distributed on an "AS IS" BASIS,
 *  WITHOUT WARRANTIES OR CONDITIONS OF ANY KIND, either express or implied.
 *  See the License for the specific language governing permissions and
 *  limitations under the License.
 */

MathJax.Hub.Insert(MathJax.OutputJax["HTML-CSS"].FONTDATA.FONTS.STIXVariants,{32:[0,0,250,0,0],119:[80,244,515,22,493],124:[690,189,320,127,193],160:[0,0,250,0,0],411:[668,0,520,55,516],612:[450,10,460,18,441],8243:[565,-28,605,43,561],8244:[565,-28,873,43,829],8245:[565,-28,340,45,296],8246:[565,-28,605,44,561],8247:[565,-28,873,43,829],8279:[565,-28,1139,43,1096],8512:[662,0,718,50,668],8592:[449,-57,415,55,415],8593:[600,15,598,82,518],8594:[449,-57,415,0,360],8595:[600,15,598,80,516],8657:[600,15,794,63,729],8659:[600,15,794,65,731],8672:[449,-58,463,70,393],8674:[449,-58,463,70,393],8712:[516,13,402,64,338],8713:[662,156,685,60,625],8715:[516,13,402,64,338],8716:[662,156,685,60,625],8719:[662,0,694,30,664],8720:[662,0,694,30,664],8721:[662,0,694,38,656],8731:[946,-55,737,72,767],8732:[943,-55,737,72,767],8733:[428,0,685,41,646],8739:[451,19,266,100,166],8740:[451,19,404,23,381],8741:[451,11,446,90,356],8742:[451,19,609,23,586],8745:[602,31,620,10,610],8746:[602,31,620,10,610],8764:[362,-148,685,48,637],8766:[344,-130,1086,55,1031],8767:[461,-43,520,0,586],8769:[462,-48,685,48,637],8772:[529,35,685,48,637],8775:[606,117,685,48,637],8776:[475,-25,685,48,637],8777:[549,49,685,48,637],8780:[586,82,685,48,637],8800:[662,156,685,48,637],8802:[662,156,685,48,637],8808:[718,275,685,56,621],8809:[718,275,685,56,621],8813:[572,66,685,48,637],8814:[662,156,685,47,612],8815:[662,156,685,73,638],8816:[695,189,685,56,621],8817:[695,189,685,64,629],8818:[673,103,685,10,632],8819:[673,144,685,58,624],8820:[730,227,685,48,637],8821:[730,227,685,48,650],8824:[818,311,685,56,621],8825:[818,311,685,55,620],8832:[662,156,685,64,621],8833:[662,156,685,64,621],8836:[662,156,685,55,620],8837:[662,156,685,65,630],8840:[707,203,695,65,630],8841:[707,203,695,65,630],8842:[607,229,685,51,616],8843:[607,229,685,69,634],8851:[536,31,620,10,610],8852:[536,31,620,10,610],8853:[623,119,842,50,792],8855:[623,119,842,50,792],8860:[623,119,842,50,792],8922:[768,262,685,60,625],8923:[768,262,685,60,625],8928:[803,212,685,60,625],8929:[803,212,685,60,625],8940:[695,189,685,54,611],8941:[695,189,685,74,631],8994:[386,-120,685,48,637],8995:[386,-120,685,48,637],9251:[22,119,500,48,453],9641:[662,158,910,45,865],10812:[633,129,463,51,411],10813:[633,129,463,51,411],10909:[647,166,685,49,627],10910:[615,166,685,54,676],10924:[625,137,685,60,625],10925:[625,137,685,60,625],10955:[718,321,685,64,629],10956:[718,321,685,74,639],10990:[451,19,404,23,381]});MathJax.Ajax.loadComplete(MathJax.OutputJax["HTML-CSS"].fontDir+"/Variants/Regular/All.js");
```
