## Supplementary material for "Artistoo: build, share, and explore simulations of cells and tissues in the web browser": MathJax.js

```
/*
 *  /MathJax.js
 *
 *  Copyright (c) 2009-2016 The MathJax Consortium
 *
 *  Licensed under the Apache License, Version 2.0 (the "License");
 *  you may not use this file except in compliance with the License.
 *  You may obtain a copy of the License at
 *
 *      http://www.apache.org/licenses/LICENSE-2.0
 *
 *  Unless required by applicable law or agreed to in writing, software
 *  distributed under the License is distributed on an "AS IS" BASIS,
 *  WITHOUT WARRANTIES OR CONDITIONS OF ANY KIND, either express or implied.
 *  See the License for the specific language governing permissions and
 *  limitations under the License.
 */

if(document.getElementById&&document.childNodes&&document.createElement){if(!(window.MathJax&&MathJax.Hub)){if(window.MathJax){window.MathJax={AuthorConfig:window.MathJax}}else{window.MathJax={}}MathJax.isPacked=true;MathJax.version="2.7.0";MathJax.fileversion="2.7.0";MathJax.cdnVersion="2.7.0";MathJax.cdnFileVersions={};(function(d){var b=window[d];if(!b){b=window[d]={}}var e=[];var c=function(f){var g=f.constructor;if(!g){g=function(){}}for(var h in f){if(h!=="constructor"&&f.hasOwnProperty(h)){g[h]=f[h]}}return g};var a=function(){return function(){return arguments.callee.Init.call(this,arguments)}};b.Object=c({constructor:a(),Subclass:function(f,h){var g=a();g.SUPER=this;g.Init=this.Init;g.Subclass=this.Subclass;g.Augment=this.Augment;g.protoFunction=this.protoFunction;g.can=this.can;g.has=this.has;g.isa=this.isa;g.prototype=new this(e);g.prototype.constructor=g;g.Augment(f,h);return g},Init:function(f){var g=this;if(f.length===1&&f[0]===e){return g}if(!(g instanceof f.callee)){g=new f.callee(e)}return g.Init.apply(g,f)||g},Augment:function(f,g){var h;if(f!=null){for(h in f){if(f.hasOwnProperty(h)){this.protoFunction(h,f[h])}}if(f.toString!==this.prototype.toString&&f.toString!=={}.toString){this.protoFunction("toString",f.toString)}}if(g!=null){for(h in g){if(g.hasOwnProperty(h)){this[h]=g[h]}}}return this},protoFunction:function(g,f){this.prototype[g]=f;if(typeof f==="function"){f.SUPER=this.SUPER.prototype}},prototype:{Init:function(){},SUPER:function(f){return f.callee.SUPER},can:function(f){return typeof(this[f])==="function"},has:function(f){return typeof(this[f])!=="undefined"},isa:function(f){return(f instanceof Object)&&(this instanceof f)}},can:function(f){return this.prototype.can.call(this,f)},has:function(f){return this.prototype.has.call(this,f)},isa:function(g){var f=this;while(f){if(f===g){return true}else{f=f.SUPER}}return false},SimpleSUPER:c({constructor:function(f){return this.SimpleSUPER.define(f)},define:function(f){var h={};if(f!=null){for(var g in f){if(f.hasOwnProperty(g)){h[g]=this.wrap(g,f[g])}}if(f.toString!==this.prototype.toString&&f.toString!=={}.toString){h.toString=this.wrap("toString",f.toString)}}return h},wrap:function(i,h){if(typeof(h)!=="function"||!h.toString().match(/\.\s*SUPER\s*\(/)){return h}var g=function(){this.SUPER=g.SUPER[i];try{var f=h.apply(this,arguments)}catch(j){delete this.SUPER;throw j}delete this.SUPER;return f};g.toString=function(){return h.toString.apply(h,arguments)};return g}})});b.Object.isArray=Array.isArray||function(f){return Object.prototype.toString.call(f)==="[object Array]"};b.Object.Array=Array})("MathJax");(function(BASENAME){var BASE=window[BASENAME];if(!BASE){BASE=window[BASENAME]={}}var isArray=BASE.Object.isArray;var CALLBACK=function(data){var cb=function(){return arguments.callee.execute.apply(arguments.callee,arguments)};for(var id in CALLBACK.prototype){if(CALLBACK.prototype.hasOwnProperty(id)){if(typeof(data[id])!=="undefined"){cb[id]=data[id]}else{cb[id]=CALLBACK.prototype[id]}}}cb.toString=CALLBACK.prototype.toString;return cb};CALLBACK.prototype={isCallback:true,hook:function(){},data:[],object:window,execute:function(){if(!this.called||this.autoReset){this.called=!this.autoReset;return this.hook.apply(this.object,this.data.concat([].slice.call(arguments,0)))}},reset:function(){delete this.called},toString:function(){return this.hook.toString.apply(this.hook,arguments)}};var ISCALLBACK=function(f){return(typeof(f)==="function"&&f.isCallback)};var EVAL=function(code){return eval.call(window,code)};var TESTEVAL=function(){EVAL("var __TeSt_VaR__ = 1");if(window.__TeSt_VaR__){try{delete window.__TeSt_VaR__}catch(error){window.__TeSt_VaR__=null}}else{if(window.execScript){EVAL=function(code){BASE.__code=code;code="try {"+BASENAME+".__result = eval("+BASENAME+".__code)} catch(err) {"+BASENAME+".__result = err}";window.execScript(code);var result=BASE.__result;delete BASE.__result;delete BASE.__code;if(result instanceof Error){throw result}return result}}else{EVAL=function(code){BASE.__code=code;code="try {"+BASENAME+".__result = eval("+BASENAME+".__code)} catch(err) {"+BASENAME+".__result = err}";var head=(document.getElementsByTagName("head"))[0];if(!head){head=document.body}var script=document.createElement("script");script.appendChild(document.createTextNode(code));head.appendChild(script);head.removeChild(script);var result=BASE.__result;delete BASE.__result;delete BASE.__code;if(result instanceof Error){throw result}return result}}}TESTEVAL=null};var USING=function(args,i){if(arguments.length>1){if(arguments.length===2&&!(typeof arguments[0]==="function")&&arguments[0] instanceof Object&&typeof arguments[1]==="number"){args=[].slice.call(args,i)}else{args=[].slice.call(arguments,0)}}if(isArray(args)&&args.length===1){args=args[0]}if(typeof args==="function"){if(args.execute===CALLBACK.prototype.execute){return args}return CALLBACK({hook:args})}else{if(isArray(args)){if(typeof(args[0])==="string"&&args[1] instanceof Object&&typeof args[1][args[0]]==="function"){return CALLBACK({hook:args[1][args[0]],object:args[1],data:args.slice(2)})}else{if(typeof args[0]==="function"){return CALLBACK({hook:args[0],data:args.slice(1)})}else{if(typeof args[1]==="function"){return CALLBACK({hook:args[1],object:args[0],data:args.slice(2)})}}}}else{if(typeof(args)==="string"){if(TESTEVAL){TESTEVAL()}return CALLBACK({hook:EVAL,data:[args]})}else{if(args instanceof Object){return CALLBACK(args)}else{if(typeof(args)==="undefined"){return CALLBACK({})}}}}}throw Error("Can't make callback from given data")};var DELAY=function(time,callback){callback=USING(callback);callback.timeout=setTimeout(callback,time);return callback};var WAITFOR=function(callback,signal){callback=USING(callback);if(!callback.called){WAITSIGNAL(callback,signal);signal.pending++}};var WAITEXECUTE=function(){var signals=this.signal;delete this.signal;this.execute=this.oldExecute;delete this.oldExecute;var result=this.execute.apply(this,arguments);if(ISCALLBACK(result)&&!result.called){WAITSIGNAL(result,signals)}else{for(var i=0,m=signals.length;i<m;i++){signals[i].pending--;if(signals[i].pending<=0){signals[i].call()}}}};var WAITSIGNAL=function(callback,signals){if(!isArray(signals)){signals=[signals]}if(!callback.signal){callback.oldExecute=callback.execute;callback.execute=WAITEXECUTE;callback.signal=signals}else{if(signals.length===1){callback.signal.push(signals[0])}else{callback.signal=callback.signal.concat(signals)}}};var AFTER=function(callback){callback=USING(callback);callback.pending=0;for(var i=1,m=arguments.length;i<m;i++){if(arguments[i]){WAITFOR(arguments[i],callback)}}if(callback.pending===0){var result=callback();if(ISCALLBACK(result)){callback=result}}return callback};var HOOKS=MathJax.Object.Subclass({Init:function(reset){this.hooks=[];this.remove=[];this.reset=reset;this.running=false},Add:function(hook,priority){if(priority==null){priority=10}if(!ISCALLBACK(hook)){hook=USING(hook)}hook.priority=priority;var i=this.hooks.length;while(i>0&&priority<this.hooks[i-1].priority){i--}this.hooks.splice(i,0,hook);return hook},Remove:function(hook){for(var i=0,m=this.hooks.length;i<m;i++){if(this.hooks[i]===hook){if(this.running){this.remove.push(i)}else{this.hooks.splice(i,1)}return}}},Execute:function(){var callbacks=[{}];this.running=true;for(var i=0,m=this.hooks.length;i<m;i++){if(this.reset){this.hooks[i].reset()}var result=this.hooks[i].apply(window,arguments);if(ISCALLBACK(result)&&!result.called){callbacks.push(result)}}this.running=false;if(this.remove.length){this.RemovePending()}if(callbacks.length===1){return null}if(callbacks.length===2){return callbacks[1]}return AFTER.apply({},callbacks)},RemovePending:function(){this.remove=this.remove.sort();for(var i=this.remove.length-1;i>=0;i--){this.hooks.splice(i,1)}this.remove=[]}});var EXECUTEHOOKS=function(hooks,data,reset){if(!hooks){return null}if(!isArray(hooks)){hooks=[hooks]}if(!isArray(data)){data=(data==null?[]:[data])}var handler=HOOKS(reset);for(var i=0,m=hooks.length;i<m;i++){handler.Add(hooks[i])}return handler.Execute.apply(handler,data)};var QUEUE=BASE.Object.Subclass({Init:function(){this.pending=this.running=0;this.queue=[];this.Push.apply(this,arguments)},Push:function(){var callback;for(var i=0,m=arguments.length;i<m;i++){callback=USING(arguments[i]);if(callback===arguments[i]&&!callback.called){callback=USING(["wait",this,callback])}this.queue.push(callback)}if(!this.running&&!this.pending){this.Process()}return callback},Process:function(queue){while(!this.running&&!this.pending&&this.queue.length){var callback=this.queue[0];queue=this.queue.slice(1);this.queue=[];this.Suspend();var result=callback();this.Resume();if(queue.length){this.queue=queue.concat(this.queue)}if(ISCALLBACK(result)&&!result.called){WAITFOR(result,this)}}},Suspend:function(){this.running++},Resume:function(){if(this.running){this.running--}},call:function(){this.Process.apply(this,arguments)},wait:function(callback){return callback}});var SIGNAL=QUEUE.Subclass({Init:function(name){QUEUE.prototype.Init.call(this);this.name=name;this.posted=[];this.listeners=HOOKS(true);this.posting=false;this.callback=null},Post:function(message,callback,forget){callback=USING(callback);if(this.posting||this.pending){this.Push(["Post",this,message,callback,forget])}else{this.callback=callback;callback.reset();if(!forget){this.posted.push(message)}this.Suspend();this.posting=true;var result=this.listeners.Execute(message);if(ISCALLBACK(result)&&!result.called){WAITFOR(result,this)}this.Resume();this.posting=false;if(!this.pending){this.call()}}return callback},Clear:function(callback){callback=USING(callback);if(this.posting||this.pending){callback=this.Push(["Clear",this,callback])}else{this.posted=[];callback()}return callback},call:function(){this.callback(this);this.Process()},Interest:function(callback,ignorePast,priority){callback=USING(callback);this.listeners.Add(callback,priority);if(!ignorePast){for(var i=0,m=this.posted.length;i<m;i++){callback.reset();var result=callback(this.posted[i]);if(ISCALLBACK(result)&&i===this.posted.length-1){WAITFOR(result,this)}}}return callback},NoInterest:function(callback){this.listeners.Remove(callback)},MessageHook:function(msg,callback,priority){callback=USING(callback);if(!this.hooks){this.hooks={};this.Interest(["ExecuteHooks",this])}if(!this.hooks[msg]){this.hooks[msg]=HOOKS(true)}this.hooks[msg].Add(callback,priority);for(var i=0,m=this.posted.length;i<m;i++){if(this.posted[i]==msg){callback.reset();callback(this.posted[i])}}callback.msg=msg;return callback},ExecuteHooks:function(msg){var type=(isArray(msg)?msg[0]:msg);if(!this.hooks[type]){return null}return this.hooks[type].Execute(msg)},RemoveHook:function(hook){this.hooks[hook.msg].Remove(hook)}},{signals:{},find:function(name){if(!SIGNAL.signals[name]){SIGNAL.signals[name]=new SIGNAL(name)}return SIGNAL.signals[name]}});BASE.Callback=BASE.CallBack=USING;BASE.Callback.Delay=DELAY;BASE.Callback.After=AFTER;BASE.Callback.Queue=QUEUE;BASE.Callback.Signal=SIGNAL.find;BASE.Callback.Hooks=HOOKS;BASE.Callback.ExecuteHooks=EXECUTEHOOKS})("MathJax");(function(e){var a=window[e];if(!a){a=window[e]={}}var d=(navigator.vendor==="Apple Computer, Inc."&&typeof navigator.vendorSub==="undefined");var g=0;var h=function(i){if(document.styleSheets&&document.styleSheets.length>g){g=document.styleSheets.length}if(!i){i=document.head||((document.getElementsByTagName("head"))[0]);if(!i){i=document.body}}return i};var f=[];var c=function(){for(var k=0,j=f.length;k<j;k++){a.Ajax.head.removeChild(f[k])}f=[]};var b={};b[e]="";b.Contrib="https://cdn.mathjax.org/mathjax/contrib";a.Ajax={loaded:{},loading:{},loadHooks:{},timeout:15*1000,styleDelay:1,config:{root:"",path:b},params:{},STATUS:{OK:1,ERROR:-1},fileURL:function(j){var i=j.match(/^\[([-._a-z0-9]+)\]/i);if(i&&i[1] in b){j=(b[i[1]]||this.config.root)+j.substr(i[1].length+2)}return j},fileName:function(j){var i=this.config.root;if(j.substr(0,i.length)===i){j="["+e+"]"+j.substr(i.length)}else{for(var k in b){if(b.hasOwnProperty(k)&&b[k]){if(j.substr(0,b[k].length)===b[k]){j="["+k+"]"+j.substr(b[k].length);break}}}}return j},fileRev:function(j){var i=a.cdnFileVersions[j]||a.cdnVersion||"";if(i){i="?V="+i}return i},urlRev:function(i){return this.fileURL(i)+this.fileRev(i)},Require:function(k,n){n=a.Callback(n);var l;if(k instanceof Object){for(var j in k){if(k.hasOwnProperty(j)){l=j.toUpperCase();k=k[j]}}}else{l=k.split(/\./).pop().toUpperCase()}if(this.params.noContrib&&k.substr(0,9)==="[Contrib]"){n(this.STATUS.ERROR)}else{k=this.fileURL(k);if(this.loaded[k]){n(this.loaded[k])}else{var m={};m[l]=k;this.Load(m,n)}}return n},Load:function(k,m){m=a.Callback(m);var l;if(k instanceof Object){for(var j in k){if(k.hasOwnProperty(j)){l=j.toUpperCase();k=k[j]}}}else{l=k.split(/\./).pop().toUpperCase()}k=this.fileURL(k);if(this.loading[k]){this.addHook(k,m)}else{this.head=h(this.head);if(this.loader[l]){this.loader[l].call(this,k,m)}else{throw Error("Can't load files of type "+l)}}return m},LoadHook:function(l,m,k){m=a.Callback(m);if(l instanceof Object){for(var j in l){if(l.hasOwnProperty(j)){l=l[j]}}}l=this.fileURL(l);if(this.loaded[l]){m(this.loaded[l])}else{this.addHook(l,m,k)}return m},addHook:function(j,k,i){if(!this.loadHooks[j]){this.loadHooks[j]=MathJax.Callback.Hooks()}this.loadHooks[j].Add(k,i);k.file=j},removeHook:function(i){if(this.loadHooks[i.file]){this.loadHooks[i.file].Remove(i);if(!this.loadHooks[i.file].hooks.length){delete this.loadHooks[i.file]}}},Preloading:function(){for(var l=0,j=arguments.length;l<j;l++){var k=this.fileURL(arguments[l]);if(!this.loading[k]){this.loading[k]={preloaded:true}}}},loader:{JS:function(k,m){var j=this.fileName(k);var i=document.createElement("script");var l=a.Callback(["loadTimeout",this,k]);this.loading[k]={callback:m,timeout:setTimeout(l,this.timeout),status:this.STATUS.OK,script:i};this.loading[k].message=a.Message.File(j);i.onerror=l;i.type="text/javascript";i.src=k+this.fileRev(j);this.head.appendChild(i)},CSS:function(j,l){var i=this.fileName(j);var k=document.createElement("link");k.rel="stylesheet";k.type="text/css";k.href=j+this.fileRev(i);this.loading[j]={callback:l,message:a.Message.File(i),status:this.STATUS.OK};this.head.appendChild(k);this.timer.create.call(this,[this.timer.file,j],k)}},timer:{create:function(j,i){j=a.Callback(j);if(i.nodeName==="STYLE"&&i.styleSheet&&typeof(i.styleSheet.cssText)!=="undefined"){j(this.STATUS.OK)}else{if(window.chrome&&i.nodeName==="LINK"){j(this.STATUS.OK)}else{if(d){this.timer.start(this,[this.timer.checkSafari2,g++,j],this.styleDelay)}else{this.timer.start(this,[this.timer.checkLength,i,j],this.styleDelay)}}}return j},start:function(j,i,k,l){i=a.Callback(i);i.execute=this.execute;i.time=this.time;i.STATUS=j.STATUS;i.timeout=l||j.timeout;i.delay=i.total=k||0;if(k){setTimeout(i,k)}else{i()}},time:function(i){this.total+=this.delay;this.delay=Math.floor(this.delay*1.05+5);if(this.total>=this.timeout){i(this.STATUS.ERROR);return 1}return 0},file:function(j,i){if(i<0){a.Ajax.loadTimeout(j)}else{a.Ajax.loadComplete(j)}},execute:function(){this.hook.call(this.object,this,this.data[0],this.data[1])},checkSafari2:function(i,j,k){if(i.time(k)){return}if(document.styleSheets.length>j&&document.styleSheets[j].cssRules&&document.styleSheets[j].cssRules.length){k(i.STATUS.OK)}else{setTimeout(i,i.delay)}},checkLength:function(i,l,n){if(i.time(n)){return}var m=0;var j=(l.sheet||l.styleSheet);try{if((j.cssRules||j.rules||[]).length>0){m=1}}catch(k){if(k.message.match(/protected variable|restricted URI/)){m=1}else{if(k.message.match(/Security error/)){m=1}}}if(m){setTimeout(a.Callback([n,i.STATUS.OK]),0)}else{setTimeout(i,i.delay)}}},loadComplete:function(i){i=this.fileURL(i);var j=this.loading[i];if(j&&!j.preloaded){a.Message.Clear(j.message);clearTimeout(j.timeout);if(j.script){if(f.length===0){setTimeout(c,0)}f.push(j.script)}this.loaded[i]=j.status;delete this.loading[i];this.addHook(i,j.callback)}else{if(j){delete this.loading[i]}this.loaded[i]=this.STATUS.OK;j={status:this.STATUS.OK}}if(!this.loadHooks[i]){return null}return this.loadHooks[i].Execute(j.status)},loadTimeout:function(i){if(this.loading[i].timeout){clearTimeout(this.loading[i].timeout)}this.loading[i].status=this.STATUS.ERROR;this.loadError(i);this.loadComplete(i)},loadError:function(i){a.Message.Set(["LoadFailed","File failed to load: %1",i],null,2000);a.Hub.signal.Post(["file load error",i])},Styles:function(k,l){var i=this.StyleString(k);if(i===""){l=a.Callback(l);l()}else{var j=document.createElement("style");j.type="text/css";this.head=h(this.head);this.head.appendChild(j);if(j.styleSheet&&typeof(j.styleSheet.cssText)!=="undefined"){j.styleSheet.cssText=i}else{j.appendChild(document.createTextNode(i))}l=this.timer.create.call(this,l,j)}return l},StyleString:function(n){if(typeof(n)==="string"){return n}var k="",o,m;for(o in n){if(n.hasOwnProperty(o)){if(typeof n[o]==="string"){k+=o+" {"+n[o]+"}\n"}else{if(a.Object.isArray(n[o])){for(var l=0;l<n[o].length;l++){m={};m[o]=n[o][l];k+=this.StyleString(m)}}else{if(o.substr(0,6)==="@media"){k+=o+" {"+this.StyleString(n[o])+"}\n"}else{if(n[o]!=null){m=[];for(var j in n[o]){if(n[o].hasOwnProperty(j)){if(n[o][j]!=null){m[m.length]=j+": "+n[o][j]}}}k+=o+" {"+m.join("; ")+"}\n"}}}}}}return k}}})("MathJax");MathJax.HTML={Element:function(d,f,e){var g=document.createElement(d),h;if(f){if(f.hasOwnProperty("style")){var c=f.style;f.style={};for(h in c){if(c.hasOwnProperty(h)){f.style[h.replace(/-([a-z])/g,this.ucMatch)]=c[h]}}}MathJax.Hub.Insert(g,f);for(h in f){if(h==="role"||h.substr(0,5)==="aria-"){g.setAttribute(h,f[h])}}}if(e){if(!MathJax.Object.isArray(e)){e=[e]}for(var b=0,a=e.length;b<a;b++){if(MathJax.Object.isArray(e[b])){g.appendChild(this.Element(e[b][0],e[b][1],e[b][2]))}else{if(d==="script"){this.setScript(g,e[b])}else{g.appendChild(document.createTextNode(e[b]))}}}}return g},ucMatch:function(a,b){return b.toUpperCase()},addElement:function(b,a,d,c){return b.appendChild(this.Element(a,d,c))},TextNode:function(a){return document.createTextNode(a)},addText:function(a,b){return a.appendChild(this.TextNode(b))},setScript:function(a,b){if(this.setScriptBug){a.text=b}else{while(a.firstChild){a.removeChild(a.firstChild)}this.addText(a,b)}},getScript:function(a){var b=(a.text===""?a.innerHTML:a.text);return b.replace(/^\s+/,"").replace(/\s+$/,"")},Cookie:{prefix:"mjx",expires:365,Set:function(a,e){var d=[];if(e){for(var g in e){if(e.hasOwnProperty(g)){d.push(g+":"+e[g].toString().replace(/&/g,"&&"))}}}var b=this.prefix+"."+a+"="+escape(d.join("&;"));if(this.expires){var f=new Date();f.setDate(f.getDate()+this.expires);b+="; expires="+f.toGMTString()}try{document.cookie=b+"; path=/"}catch(c){}},Get:function(a,d){if(!d){d={}}var g=new RegExp("(?:^|;\\s*)"+this.prefix+"\\."+a+"=([^;]*)(?:;|$)");var f;try{f=g.exec(document.cookie)}catch(c){}if(f&&f[1]!==""){var j=unescape(f[1]).split("&;");for(var e=0,b=j.length;e<b;e++){f=j[e].match(/([^:]+):(.*)/);var h=f[2].replace(/&&/g,"&");if(h==="true"){h=true}else{if(h==="false"){h=false}else{if(h.match(/^-?(\d+(\.\d+)?|\.\d+)$/)){h=parseFloat(h)}}}d[f[1]]=h}}return d}}};MathJax.Localization={locale:"en",directory:"[MathJax]/localization",strings:{ast:{menuTitle:"asturianu"},bg:{menuTitle:"\u0431\u044A\u043B\u0433\u0430\u0440\u0441\u043A\u0438"},bcc:{menuTitle:"\u0628\u0644\u0648\u0686\u06CC"},br:{menuTitle:"brezhoneg"},ca:{menuTitle:"catal\u00E0"},cdo:{menuTitle:"M\u00ECng-d\u0115\u0324ng-ng\u1E73\u0304"},cs:{menuTitle:"\u010De\u0161tina"},da:{menuTitle:"dansk"},de:{menuTitle:"Deutsch"},diq:{menuTitle:"Zazaki"},en:{menuTitle:"English",isLoaded:true},eo:{menuTitle:"Esperanto"},es:{menuTitle:"espa\u00F1ol"},fa:{menuTitle:"\u0641\u0627\u0631\u0633\u06CC"},fi:{menuTitle:"suomi"},fr:{menuTitle:"fran\u00E7ais"},gl:{menuTitle:"galego"},he:{menuTitle:"\u05E2\u05D1\u05E8\u05D9\u05EA"},ia:{menuTitle:"interlingua"},it:{menuTitle:"italiano"},ja:{menuTitle:"\u65E5\u672C\u8A9E"},kn:{menuTitle:"\u0C95\u0CA8\u0CCD\u0CA8\u0CA1"},ko:{menuTitle:"\uD55C\uAD6D\uC5B4"},lb:{menuTitle:"L\u00EBtzebuergesch"},lki:{menuTitle:"\u0644\u06D5\u06A9\u06CC"},lt:{menuTitle:"lietuvi\u0173"},mk:{menuTitle:"\u043C\u0430\u043A\u0435\u0434\u043E\u043D\u0441\u043A\u0438"},nl:{menuTitle:"Nederlands"},oc:{menuTitle:"occitan"},pl:{menuTitle:"polski"},pt:{menuTitle:"portugus\u00EA"},"pt-br":{menuTitle:"portugu\u00EAs do Brasil"},ru:{menuTitle:"\u0440\u0443\u0441\u0441\u043A\u0438\u0439"},sco:{menuTitle:"Scots"},scn:{menuTitle:"sicilianu"},sl:{menuTitle:"sloven\u0161\u010Dina"},sv:{menuTitle:"svenska"},tr:{menuTitle:"T\u00FCrk\u00E7e"},uk:{menuTitle:"\u0443\u043A\u0440\u0430\u0457\u043D\u0441\u044C\u043A\u0430"},vi:{menuTitle:"Ti\u1EBFng Vi\u1EC7t"},"zh-hans":{menuTitle:"\u4E2D\u6587\uFF08\u7B80\u4F53\uFF09"}},pattern:/%(\d+|\{\d+\}|\{[a-z]+:\%\d+(?:\|(?:%\{\d+\}|%.|[^\}])*)+\}|.)/g,SPLIT:("axb".split(/(x)/).length===3?function(a,b){return a.split(b)}:function(c,e){var a=[],b,d=0;e.lastIndex=0;while((b=e.exec(c))){a.push(c.substr(d,b.index-d));a.push.apply(a,b.slice(1));d=b.index+b[0].length}a.push(c.substr(d));return a}),_:function(b,a){if(MathJax.Object.isArray(a)){return this.processSnippet(b,a)}return this.processString(this.lookupPhrase(b,a),[].slice.call(arguments,2))},processString:function(l,p,g){var j,e,o=MathJax.Object.isArray;for(j=0,e=p.length;j<e;j++){if(g&&o(p[j])){p[j]=this.processSnippet(g,p[j])}}var f=this.SPLIT(l,this.pattern);for(j=1,e=f.length;j<e;j+=2){var q=f[j].charAt(0);if(q>="0"&&q<="9"){f[j]=p[f[j]-1];if(typeof f[j]==="number"){f[j]=this.number(f[j])}}else{if(q==="{"){q=f[j].substr(1);if(q>="0"&&q<="9"){f[j]=p[f[j].substr(1,f[j].length-2)-1];if(typeof f[j]==="number"){f[j]=this.number(f[j])}}else{var k=f[j].match(/^\{([a-z]+):%(\d+)\|(.*)\}$/);if(k){if(k[1]==="plural"){var d=p[k[2]-1];if(typeof d==="undefined"){f[j]="???"}else{d=this.plural(d)-1;var h=k[3].replace(/(^|[^%])(%%)*%\|/g,"$1$2%\uEFEF").split(/\|/);if(d>=0&&d<h.length){f[j]=this.processString(h[d].replace(/\uEFEF/g,"|"),p,g)}else{f[j]="???"}}}else{f[j]="%"+f[j]}}}}}if(f[j]==null){f[j]="???"}}if(!g){return f.join("")}var a=[],b="";for(j=0;j<e;j++){b+=f[j];j++;if(j<e){if(o(f[j])){a.push(b);a=a.concat(f[j]);b=""}else{b+=f[j]}}}if(b!==""){a.push(b)}return a},processSnippet:function(g,e){var c=[];for(var d=0,b=e.length;d<b;d++){if(MathJax.Object.isArray(e[d])){var f=e[d];if(typeof f[1]==="string"){var h=f[0];if(!MathJax.Object.isArray(h)){h=[g,h]}var a=this.lookupPhrase(h,f[1]);c=c.concat(this.processMarkdown(a,f.slice(2),g))}else{if(MathJax.Object.isArray(f[1])){c=c.concat(this.processSnippet.apply(this,f))}else{if(f.length>=3){c.push([f[0],f[1],this.processSnippet(g,f[2])])}else{c.push(e[d])}}}}else{c.push(e[d])}}return c},markdownPattern:/(%.)|(\*{1,3})((?:%.|.)+?)\2|(`+)((?:%.|.)+?)\4|\[((?:%.|.)+?)\]\(([^\s\)]+)\)/,processMarkdown:function(b,h,d){var j=[],e;var c=b.split(this.markdownPattern);var g=c[0];for(var f=1,a=c.length;f<a;f+=8){if(c[f+1]){e=this.processString(c[f+2],h,d);if(!MathJax.Object.isArray(e)){e=[e]}e=[["b","i","i"][c[f+1].length-1],{},e];if(c[f+1].length===3){e=["b",{},e]}}else{if(c[f+3]){e=this.processString(c[f+4].replace(/^\s/,"").replace(/\s$/,""),h,d);if(!MathJax.Object.isArray(e)){e=[e]}e=["code",{},e]}else{if(c[f+5]){e=this.processString(c[f+5],h,d);if(!MathJax.Object.isArray(e)){e=[e]}e=["a",{href:this.processString(c[f+6],h),target:"_blank"},e]}else{g+=c[f];e=null}}}if(e){j=this.concatString(j,g,h,d);j.push(e);g=""}if(c[f+7]!==""){g+=c[f+7]}}j=this.concatString(j,g,h,d);return j},concatString:function(a,c,b,d){if(c!=""){c=this.processString(c,b,d);if(!MathJax.Object.isArray(c)){c=[c]}a=a.concat(c)}return a},lookupPhrase:function(f,a,d){if(!d){d="_"}if(MathJax.Object.isArray(f)){d=(f[0]||"_");f=(f[1]||"")}var c=this.loadDomain(d);if(c){MathJax.Hub.RestartAfter(c)}var b=this.strings[this.locale];if(b){if(b.domains&&d in b.domains){var e=b.domains[d];if(e.strings&&f in e.strings){a=e.strings[f]}}}return a},loadFile:function(b,d,e){e=MathJax.Callback(e);b=(d.file||b);if(!b.match(/\.js$/)){b+=".js"}if(!b.match(/^([a-z]+:|\[MathJax\])/)){var a=(this.strings[this.locale].directory||this.directory+"/"+this.locale||"[MathJax]/localization/"+this.locale);b=a+"/"+b}var c=MathJax.Ajax.Require(b,function(){d.isLoaded=true;return e()});return(c.called?null:c)},loadDomain:function(c,e){var b,a=this.strings[this.locale];if(a){if(!a.isLoaded){b=this.loadFile(this.locale,a);if(b){return MathJax.Callback.Queue(b,["loadDomain",this,c]).Push(e||{})}}if(a.domains&&c in a.domains){var d=a.domains[c];if(!d.isLoaded){b=this.loadFile(c,d);if(b){return MathJax.Callback.Queue(b).Push(e)}}}}return MathJax.Callback(e)()},Try:function(a){a=MathJax.Callback(a);a.autoReset=true;try{a()}catch(b){if(!b.restart){throw b}MathJax.Callback.After(["Try",this,a],b.restart)}},resetLocale:function(a){if(!a){return}a=a.toLowerCase();while(!this.strings[a]){var c=a.lastIndexOf("-");if(c===-1){return}a=a.substring(0,c)}var b=this.strings[a].remap;this.locale=b?b:a},setLocale:function(a){this.resetLocale(a);if(MathJax.Menu){this.loadDomain("MathMenu")}},addTranslation:function(b,e,c){var d=this.strings[b],a=false;if(!d){d=this.strings[b]={};a=true}if(!d.domains){d.domains={}}if(e){if(!d.domains[e]){d.domains[e]={}}d=d.domains[e]}MathJax.Hub.Insert(d,c);if(a&&MathJax.Menu.menu){MathJax.Menu.CreateLocaleMenu()}},setCSS:function(b){var a=this.strings[this.locale];if(a){if(a.fontFamily){b.style.fontFamily=a.fontFamily}if(a.fontDirection){b.style.direction=a.fontDirection;if(a.fontDirection==="rtl"){b.style.textAlign="right"}}}return b},fontFamily:function(){var a=this.strings[this.locale];return(a?a.fontFamily:null)},fontDirection:function(){var a=this.strings[this.locale];return(a?a.fontDirection:null)},plural:function(b){var a=this.strings[this.locale];if(a&&a.plural){return a.plural(b)}if(b==1){return 1}return 2},number:function(b){var a=this.strings[this.locale];if(a&&a.number){return a.number(b)}return b}};MathJax.Message={ready:false,log:[{}],current:null,textNodeBug:(navigator.vendor==="Apple Computer, Inc."&&typeof navigator.vendorSub==="undefined")||(window.hasOwnProperty&&window.hasOwnProperty("konqueror")),styles:{"#MathJax_Message":{position:"fixed",left:"1px",bottom:"2px","background-color":"#E6E6E6",border:"1px solid #959595",margin:"0px",padding:"2px 8px","z-index":"102",color:"black","font-size":"80%",width:"auto","white-space":"nowrap"},"#MathJax_MSIE_Frame":{position:"absolute",top:0,left:0,width:"0px","z-index":101,border:"0px",margin:"0px",padding:"0px"}},browsers:{MSIE:function(a){MathJax.Message.msieFixedPositionBug=((document.documentMode||0)<7);if(MathJax.Message.msieFixedPositionBug){MathJax.Hub.config.styles["#MathJax_Message"].position="absolute"}MathJax.Message.quirks=(document.compatMode==="BackCompat")},Chrome:function(a){MathJax.Hub.config.styles["#MathJax_Message"].bottom="1.5em";MathJax.Hub.config.styles["#MathJax_Message"].left="1em"}},Init:function(a){if(a){this.ready=true}if(!document.body||!this.ready){return false}if(this.div&&this.div.parentNode==null){this.div=document.getElementById("MathJax_Message");if(this.div){this.text=this.div.firstChild}}if(!this.div){var b=document.body;if(this.msieFixedPositionBug&&window.attachEvent){b=this.frame=this.addDiv(document.body);b.removeAttribute("id");b.style.position="absolute";b.style.border=b.style.margin=b.style.padding="0px";b.style.zIndex="101";b.style.height="0px";b=this.addDiv(b);b.id="MathJax_MSIE_Frame";window.attachEvent("onscroll",this.MoveFrame);window.attachEvent("onresize",this.MoveFrame);this.MoveFrame()}this.div=this.addDiv(b);this.div.style.display="none";this.text=this.div.appendChild(document.createTextNode(""))}return true},addDiv:function(a){var b=document.createElement("div");b.id="MathJax_Message";if(a.firstChild){a.insertBefore(b,a.firstChild)}else{a.appendChild(b)}return b},MoveFrame:function(){var a=(MathJax.Message.quirks?document.body:document.documentElement);var b=MathJax.Message.frame;b.style.left=a.scrollLeft+"px";b.style.top=a.scrollTop+"px";b.style.width=a.clientWidth+"px";b=b.firstChild;b.style.height=a.clientHeight+"px"},localize:function(a){return MathJax.Localization._(a,a)},filterText:function(a,c,b){if(MathJax.Hub.config.messageStyle==="simple"){if(b==="LoadFile"){if(!this.loading){this.loading=this.localize("Loading")+" "}a=this.loading;this.loading+="."}else{if(b==="ProcessMath"){if(!this.processing){this.processing=this.localize("Processing")+" "}a=this.processing;this.processing+="."}else{if(b==="TypesetMath"){if(!this.typesetting){this.typesetting=this.localize("Typesetting")+" "}a=this.typesetting;this.typesetting+="."}}}}return a},clearCounts:function(){delete this.loading;delete this.processing;delete this.typesetting},Set:function(c,e,b){if(e==null){e=this.log.length;this.log[e]={}}var d="";if(MathJax.Object.isArray(c)){d=c[0];if(MathJax.Object.isArray(d)){d=d[1]}try{c=MathJax.Localization._.apply(MathJax.Localization,c)}catch(a){if(!a.restart){throw a}if(!a.restart.called){if(this.log[e].restarted==null){this.log[e].restarted=0}this.log[e].restarted++;delete this.log[e].cleared;MathJax.Callback.After(["Set",this,c,e,b],a.restart);return e}}}if(this.timer){clearTimeout(this.timer);delete this.timer}this.log[e].text=c;this.log[e].filteredText=c=this.filterText(c,e,d);if(typeof(this.log[e].next)==="undefined"){this.log[e].next=this.current;if(this.current!=null){this.log[this.current].prev=e}this.current=e}if(this.current===e&&MathJax.Hub.config.messageStyle!=="none"){if(this.Init()){if(this.textNodeBug){this.div.innerHTML=c}else{this.text.nodeValue=c}this.div.style.display="";if(this.status){window.status="";delete this.status}}else{window.status=c;this.status=true}}if(this.log[e].restarted){if(this.log[e].cleared){b=0}if(--this.log[e].restarted===0){delete this.log[e].cleared}}if(b){setTimeout(MathJax.Callback(["Clear",this,e]),b)}else{if(b==0){this.Clear(e,0)}}return e},Clear:function(b,a){if(this.log[b].prev!=null){this.log[this.log[b].prev].next=this.log[b].next}if(this.log[b].next!=null){this.log[this.log[b].next].prev=this.log[b].prev}if(this.current===b){this.current=this.log[b].next;if(this.text){if(this.div.parentNode==null){this.Init()}if(this.current==null){if(this.timer){clearTimeout(this.timer);delete this.timer}if(a==null){a=600}if(a===0){this.Remove()}else{this.timer=setTimeout(MathJax.Callback(["Remove",this]),a)}}else{if(MathJax.Hub.config.messageStyle!=="none"){if(this.textNodeBug){this.div.innerHTML=this.log[this.current].filteredText}else{this.text.nodeValue=this.log[this.current].filteredText}}}if(this.status){window.status="";delete this.status}}else{if(this.status){window.status=(this.current==null?"":this.log[this.current].text)}}}delete this.log[b].next;delete this.log[b].prev;delete this.log[b].filteredText;if(this.log[b].restarted){this.log[b].cleared=true}},Remove:function(){this.text.nodeValue="";this.div.style.display="none"},File:function(a){return this.Set(["LoadFile","Loading %1",a],null,null)},Log:function(){var b=[];for(var c=1,a=this.log.length;c<a;c++){b[c]=this.log[c].text}return b.join("\n")}};MathJax.Hub={config:{root:"",config:[],styleSheets:[],styles:{".MathJax_Preview":{color:"#888"}},jax:[],extensions:[],preJax:null,postJax:null,displayAlign:"center",displayIndent:"0",preRemoveClass:"MathJax_Preview",showProcessingMessages:true,messageStyle:"normal",delayStartupUntil:"none",skipStartupTypeset:false,elements:[],positionToHash:true,showMathMenu:true,showMathMenuMSIE:true,menuSettings:{zoom:"None",CTRL:false,ALT:false,CMD:false,Shift:false,discoverable:false,zscale:"200%",renderer:null,font:"Auto",context:"MathJax",locale:null,mpContext:false,mpMouse:false,texHints:true,FastPreview:null,assistiveMML:null,inTabOrder:true,semantics:false},errorSettings:{message:["[",["MathProcessingError","Math Processing Error"],"]"],style:{color:"#CC0000","font-style":"italic"}},ignoreMMLattributes:{}},preProcessors:MathJax.Callback.Hooks(true),inputJax:{},outputJax:{order:{}},processSectionDelay:50,processUpdateTime:250,processUpdateDelay:10,signal:MathJax.Callback.Signal("Hub"),Config:function(a){this.Insert(this.config,a);if(this.config.Augment){this.Augment(this.config.Augment)}},CombineConfig:function(c,f){var b=this.config,g,e;c=c.split(/\./);for(var d=0,a=c.length;d<a;d++){g=c[d];if(!b[g]){b[g]={}}e=b;b=b[g]}e[g]=b=this.Insert(f,b);return b},Register:{PreProcessor:function(){return MathJax.Hub.preProcessors.Add.apply(MathJax.Hub.preProcessors,arguments)},MessageHook:function(){return MathJax.Hub.signal.MessageHook.apply(MathJax.Hub.signal,arguments)},StartupHook:function(){return MathJax.Hub.Startup.signal.MessageHook.apply(MathJax.Hub.Startup.signal,arguments)},LoadHook:function(){return MathJax.Ajax.LoadHook.apply(MathJax.Ajax,arguments)}},UnRegister:{PreProcessor:function(a){MathJax.Hub.preProcessors.Remove(a)},MessageHook:function(a){MathJax.Hub.signal.RemoveHook(a)},StartupHook:function(a){MathJax.Hub.Startup.signal.RemoveHook(a)},LoadHook:function(a){MathJax.Ajax.removeHook(a)}},getAllJax:function(e){var c=[],b=this.elementScripts(e);for(var d=0,a=b.length;d<a;d++){if(b[d].MathJax&&b[d].MathJax.elementJax){c.push(b[d].MathJax.elementJax)}}return c},getJaxByType:function(f,e){var c=[],b=this.elementScripts(e);for(var d=0,a=b.length;d<a;d++){if(b[d].MathJax&&b[d].MathJax.elementJax&&b[d].MathJax.elementJax.mimeType===f){c.push(b[d].MathJax.elementJax)}}return c},getJaxByInputType:function(f,e){var c=[],b=this.elementScripts(e);for(var d=0,a=b.length;d<a;d++){if(b[d].MathJax&&b[d].MathJax.elementJax&&b[d].type&&b[d].type.replace(/ *;(.|\s)*/,"")===f){c.push(b[d].MathJax.elementJax)}}return c},getJaxFor:function(a){if(typeof(a)==="string"){a=document.getElementById(a)}if(a&&a.MathJax){return a.MathJax.elementJax}if(this.isMathJaxNode(a)){if(!a.isMathJax){a=a.firstChild}while(a&&!a.jaxID){a=a.parentNode}if(a){return MathJax.OutputJax[a.jaxID].getJaxFromMath(a)}}return null},isJax:function(a){if(typeof(a)==="string"){a=document.getElementById(a)}if(this.isMathJaxNode(a)){return 1}if(a&&(a.tagName||"").toLowerCase()==="script"){if(a.MathJax){return(a.MathJax.state===MathJax.ElementJax.STATE.PROCESSED?1:-1)}if(a.type&&this.inputJax[a.type.replace(/ *;(.|\s)*/,"")]){return -1}}return 0},isMathJaxNode:function(a){return !!a&&(a.isMathJax||(a.className||"")==="MathJax_MathML")},setRenderer:function(d,c){if(!d){return}if(!MathJax.OutputJax[d]){this.config.menuSettings.renderer="";var b="[MathJax]/jax/output/"+d+"/config.js";return MathJax.Ajax.Require(b,["setRenderer",this,d,c])}else{this.config.menuSettings.renderer=d;if(c==null){c="jax/mml"}var a=this.outputJax;if(a[c]&&a[c].length){if(d!==a[c][0].id){a[c].unshift(MathJax.OutputJax[d]);return this.signal.Post(["Renderer Selected",d])}}return null}},Queue:function(){return this.queue.Push.apply(this.queue,arguments)},Typeset:function(c,d){if(!MathJax.isReady){return null}var b=this.elementCallback(c,d);if(b.count){var a=MathJax.Callback.Queue(["PreProcess",this,b.elements],["Process",this,b.elements])}return a.Push(b.callback)},PreProcess:function(e,g){var c=this.elementCallback(e,g);var b=MathJax.Callback.Queue();if(c.count){var f=(c.count===1?[c.elements]:c.elements);b.Push(["Post",this.signal,["Begin PreProcess",c.elements]]);for(var d=0,a=f.length;d<a;d++){if(f[d]){b.Push(["Execute",this.preProcessors,f[d]])}}b.Push(["Post",this.signal,["End PreProcess",c.elements]])}return b.Push(c.callback)},Process:function(a,b){return this.takeAction("Process",a,b)},Update:function(a,b){return this.takeAction("Update",a,b)},Reprocess:function(a,b){return this.takeAction("Reprocess",a,b)},Rerender:function(a,b){return this.takeAction("Rerender",a,b)},takeAction:function(g,d,h){var c=this.elementCallback(d,h);var f=c.elements;var a=MathJax.Callback.Queue(["Clear",this.signal]);var e={scripts:[],start:new Date().getTime(),i:0,j:0,jax:{},jaxIDs:[]};if(c.count){var b=["Delay",MathJax.Callback,this.processSectionDelay];if(!b[2]){b={}}a.Push(["clearCounts",MathJax.Message],["Post",this.signal,["Begin "+g,f]],["Post",this.signal,["Begin Math",f,g]],["prepareScripts",this,g,f,e],["Post",this.signal,["Begin Math Input",f,g]],["processInput",this,e],["Post",this.signal,["End Math Input",f,g]],b,["prepareOutput",this,e,"preProcess"],b,["Post",this.signal,["Begin Math Output",f,g]],["processOutput",this,e],["Post",this.signal,["End Math Output",f,g]],b,["prepareOutput",this,e,"postProcess"],b,["Post",this.signal,["End Math",f,g]],["Post",this.signal,["End "+g,f]],["clearCounts",MathJax.Message])}return a.Push(c.callback)},scriptAction:{Process:function(a){},Update:function(b){var a=b.MathJax.elementJax;if(a&&a.needsUpdate()){a.Remove(true);b.MathJax.state=a.STATE.UPDATE}else{b.MathJax.state=a.STATE.PROCESSED}},Reprocess:function(b){var a=b.MathJax.elementJax;if(a){a.Remove(true);b.MathJax.state=a.STATE.UPDATE}},Rerender:function(b){var a=b.MathJax.elementJax;if(a){a.Remove(true);b.MathJax.state=a.STATE.OUTPUT}}},prepareScripts:function(h,e,g){if(arguments.callee.disabled){return}var b=this.elementScripts(e);var f=MathJax.ElementJax.STATE;for(var d=0,a=b.length;d<a;d++){var c=b[d];if(c.type&&this.inputJax[c.type.replace(/ *;(.|\n)*/,"")]){if(c.MathJax){if(c.MathJax.elementJax&&c.MathJax.elementJax.hover){MathJax.Extension.MathEvents.Hover.ClearHover(c.MathJax.elementJax)}if(c.MathJax.state!==f.PENDING){this.scriptAction[h](c)}}if(!c.MathJax){c.MathJax={state:f.PENDING}}if(c.MathJax.error){delete c.MathJax.error}if(c.MathJax.state!==f.PROCESSED){g.scripts.push(c)}}}},checkScriptSiblings:function(a){if(a.MathJax.checked){return}var b=this.config,f=a.previousSibling;if(f&&f.nodeName==="#text"){var d,e,c=a.nextSibling;if(c&&c.nodeName!=="#text"){c=null}if(b.preJax){if(typeof(b.preJax)==="string"){b.preJax=new RegExp(b.preJax+"$")}d=f.nodeValue.match(b.preJax)}if(b.postJax&&c){if(typeof(b.postJax)==="string"){b.postJax=new RegExp("^"+b.postJax)}e=c.nodeValue.match(b.postJax)}if(d&&(!b.postJax||e)){f.nodeValue=f.nodeValue.replace(b.preJax,(d.length>1?d[1]:""));f=null}if(e&&(!b.preJax||d)){c.nodeValue=c.nodeValue.replace(b.postJax,(e.length>1?e[1]:""))}if(f&&!f.nodeValue.match(/\S/)){f=f.previousSibling}}if(b.preRemoveClass&&f&&f.className===b.preRemoveClass){a.MathJax.preview=f}a.MathJax.checked=1},processInput:function(a){var b,i=MathJax.ElementJax.STATE;var h,e,d=a.scripts.length;try{while(a.i<d){h=a.scripts[a.i];if(!h){a.i++;continue}e=h.previousSibling;if(e&&e.className==="MathJax_Error"){e.parentNode.removeChild(e)}if(!h.parentNode||!h.MathJax||h.MathJax.state===i.PROCESSED){a.i++;continue}if(!h.MathJax.elementJax||h.MathJax.state===i.UPDATE){this.checkScriptSiblings(h);var g=h.type.replace(/ *;(.|\s)*/,"");var j=this.inputJax[g];b=j.Process(h,a);if(typeof b==="function"){if(b.called){continue}this.RestartAfter(b)}b=b.Attach(h,j.id);this.saveScript(b,a,h,i);this.postInputHooks.Execute(b,j.id,h)}else{if(h.MathJax.state===i.OUTPUT){this.saveScript(h.MathJax.elementJax,a,h,i)}}a.i++;var c=new Date().getTime();if(c-a.start>this.processUpdateTime&&a.i<a.scripts.length){a.start=c;this.RestartAfter(MathJax.Callback.Delay(1))}}}catch(f){return this.processError(f,a,"Input")}if(a.scripts.length&&this.config.showProcessingMessages){MathJax.Message.Set(["ProcessMath","Processing math: %1%%",100],0)}a.start=new Date().getTime();a.i=a.j=0;return null},postInputHooks:MathJax.Callback.Hooks(true),saveScript:function(a,d,b,c){if(!this.outputJax[a.mimeType]){b.MathJax.state=c.UPDATE;throw Error("No output jax registered for "+a.mimeType)}a.outputJax=this.outputJax[a.mimeType][0].id;if(!d.jax[a.outputJax]){if(d.jaxIDs.length===0){d.jax[a.outputJax]=d.scripts}else{if(d.jaxIDs.length===1){d.jax[d.jaxIDs[0]]=d.scripts.slice(0,d.i)}d.jax[a.outputJax]=[]}d.jaxIDs.push(a.outputJax)}if(d.jaxIDs.length>1){d.jax[a.outputJax].push(b)}b.MathJax.state=c.OUTPUT},prepareOutput:function(c,f){while(c.j<c.jaxIDs.length){var e=c.jaxIDs[c.j],d=MathJax.OutputJax[e];if(d[f]){try{var a=d[f](c);if(typeof a==="function"){if(a.called){continue}this.RestartAfter(a)}}catch(b){if(!b.restart){MathJax.Message.Set(["PrepError","Error preparing %1 output (%2)",e,f],null,600);MathJax.Hub.lastPrepError=b;c.j++}return MathJax.Callback.After(["prepareOutput",this,c,f],b.restart)}}c.j++}return null},processOutput:function(h){var b,g=MathJax.ElementJax.STATE,d,a=h.scripts.length;try{while(h.i<a){d=h.scripts[h.i];if(!d||!d.parentNode||!d.MathJax||d.MathJax.error){h.i++;continue}var c=d.MathJax.elementJax;if(!c){h.i++;continue}b=MathJax.OutputJax[c.outputJax].Process(d,h);if(b!==false){d.MathJax.state=g.PROCESSED;if(d.MathJax.preview){d.MathJax.preview.innerHTML="";d.MathJax.preview.style.display="none"}this.signal.Post(["New Math",c.inputID])}h.i++;var e=new Date().getTime();if(e-h.start>this.processUpdateTime&&h.i<h.scripts.length){h.start=e;this.RestartAfter(MathJax.Callback.Delay(this.processUpdateDelay))}}}catch(f){return this.processError(f,h,"Output")}if(h.scripts.length&&this.config.showProcessingMessages){MathJax.Message.Set(["TypesetMath","Typesetting math: %1%%",100],0);MathJax.Message.Clear(0)}h.i=h.j=0;return null},processMessage:function(d,b){var a=Math.floor(d.i/(d.scripts.length)*100);var c=(b==="Output"?["TypesetMath","Typesetting math: %1%%"]:["ProcessMath","Processing math: %1%%"]);if(this.config.showProcessingMessages){MathJax.Message.Set(c.concat(a),0)}},processError:function(b,c,a){if(!b.restart){if(!this.config.errorSettings.message){throw b}this.formatError(c.scripts[c.i],b);c.i++}this.processMessage(c,a);return MathJax.Callback.After(["process"+a,this,c],b.restart)},formatError:function(b,f){var h=function(l,k,j,i){return MathJax.Localization._(l,k,j,i)};var e=h("ErrorMessage","Error: %1",f.message)+"\n";if(f.sourceURL||f.fileName){e+="\n"+h("ErrorFile","file: %1",f.sourceURL||f.fileName)}if(f.line||f.lineNumber){e+="\n"+h("ErrorLine","line: %1",f.line||f.lineNumber)}e+="\n\n"+h("ErrorTips","Debugging tips: use %1, inspect %2 in the browser console","'unpacked/MathJax.js'","'MathJax.Hub.lastError'");b.MathJax.error=MathJax.OutputJax.Error.Jax(e,b);if(b.MathJax.elementJax){b.MathJax.error.inputID=b.MathJax.elementJax.inputID}var g=this.config.errorSettings;var a=h(g.messageId,g.message);var c=MathJax.HTML.Element("span",{className:"MathJax_Error",jaxID:"Error",isMathJax:true,id:b.MathJax.error.inputID+"-Frame"},[["span",null,a]]);MathJax.Ajax.Require("[MathJax]/extensions/MathEvents.js",function(){var j=MathJax.Extension.MathEvents.Event,i=MathJax.Hub;c.oncontextmenu=j.Menu;c.onmousedown=j.Mousedown;c.onkeydown=j.Keydown;c.tabIndex=i.getTabOrder(i.getJaxFor(b))});var d=document.getElementById(c.id);if(d){d.parentNode.removeChild(d)}if(b.parentNode){b.parentNode.insertBefore(c,b)}if(b.MathJax.preview){b.MathJax.preview.innerHTML="";b.MathJax.preview.style.display="none"}this.lastError=f;this.signal.Post(["Math Processing Error",b,f])},RestartAfter:function(a){throw this.Insert(Error("restart"),{restart:MathJax.Callback(a)})},elementCallback:function(c,f){if(f==null&&(MathJax.Object.isArray(c)||typeof c==="function")){try{MathJax.Callback(c);f=c;c=null}catch(d){}}if(c==null){c=this.config.elements||[]}if(this.isHTMLCollection(c)){c=this.HTMLCollection2Array(c)}if(!MathJax.Object.isArray(c)){c=[c]}c=[].concat(c);for(var b=0,a=c.length;b<a;b++){if(typeof(c[b])==="string"){c[b]=document.getElementById(c[b])}}if(!document.body){document.body=document.getElementsByTagName("body")[0]}if(c.length==0){c.push(document.body)}if(!f){f={}}return{count:c.length,elements:(c.length===1?c[0]:c),callback:f}},elementScripts:function(e){var b=[];if(MathJax.Object.isArray(e)||this.isHTMLCollection(e)){for(var d=0,a=e.length;d<a;d++){var f=0;for(var c=0;c<d&&!f;c++){f=e[c].contains(e[d])}if(!f){b.push.apply(b,this.elementScripts(e[d]))}}return b}if(typeof(e)==="string"){e=document.getElementById(e)}if(!document.body){document.body=document.getElementsByTagName("body")[0]}if(e==null){e=document.body}if(e.tagName!=null&&e.tagName.toLowerCase()==="script"){return[e]}b=e.getElementsByTagName("script");if(this.msieHTMLCollectionBug){b=this.HTMLCollection2Array(b)}return b},isHTMLCollection:function(a){return("HTMLCollection" in window&&typeof(a)==="object"&&a instanceof HTMLCollection)},HTMLCollection2Array:function(c){if(!this.msieHTMLCollectionBug){return[].slice.call(c)}var b=[];for(var d=0,a=c.length;d<a;d++){b[d]=c[d]}return b},Insert:function(c,a){for(var b in a){if(a.hasOwnProperty(b)){if(typeof a[b]==="object"&&!(MathJax.Object.isArray(a[b]))&&(typeof c[b]==="object"||typeof c[b]==="function")){this.Insert(c[b],a[b])}else{c[b]=a[b]}}}return c},getTabOrder:function(a){return this.config.menuSettings.inTabOrder?0:-1},SplitList:("trim" in String.prototype?function(a){return a.trim().split(/\s+/)}:function(a){return a.replace(/^\s+/,"").replace(/\s+$/,"").split(/\s+/)})};MathJax.Hub.Insert(MathJax.Hub.config.styles,MathJax.Message.styles);MathJax.Hub.Insert(MathJax.Hub.config.styles,{".MathJax_Error":MathJax.Hub.config.errorSettings.style});MathJax.Extension={};MathJax.Hub.Configured=MathJax.Callback({});MathJax.Hub.Startup={script:"",queue:MathJax.Callback.Queue(),signal:MathJax.Callback.Signal("Startup"),params:{},Config:function(){this.queue.Push(["Post",this.signal,"Begin Config"]);if(MathJax.AuthorConfig&&MathJax.AuthorConfig.root){MathJax.Ajax.config.root=MathJax.AuthorConfig.root}if(this.params.locale){MathJax.Localization.resetLocale(this.params.locale);MathJax.Hub.config.menuSettings.locale=this.params.locale}if(this.params.config){var c=this.params.config.split(/,/);for(var b=0,a=c.length;b<a;b++){if(!c[b].match(/\.js$/)){c[b]+=".js"}this.queue.Push(["Require",MathJax.Ajax,this.URL("config",c[b])])}}this.queue.Push(["Config",MathJax.Hub,MathJax.AuthorConfig]);if(this.script.match(/\S/)){this.queue.Push(this.script+";\n1;")}this.queue.Push(["ConfigDelay",this],["ConfigBlocks",this],[function(d){return d.loadArray(MathJax.Hub.config.config,"config",null,true)},this],["Post",this.signal,"End Config"])},ConfigDelay:function(){var a=this.params.delayStartupUntil||MathJax.Hub.config.delayStartupUntil;if(a==="onload"){return this.onload}if(a==="configured"){return MathJax.Hub.Configured}return a},ConfigBlocks:function(){var c=document.getElementsByTagName("script");var b=MathJax.Callback.Queue();for(var d=0,a=c.length;d<a;d++){var e=String(c[d].type).replace(/ /g,"");if(e.match(/^text\/x-mathjax-config(;.*)?$/)&&!e.match(/;executed=true/)){c[d].type+=";executed=true";b.Push(c[d].innerHTML+";\n1;")}}return b.Push(function(){MathJax.Ajax.config.root=MathJax.Hub.config.root})},Cookie:function(){return this.queue.Push(["Post",this.signal,"Begin Cookie"],["Get",MathJax.HTML.Cookie,"menu",MathJax.Hub.config.menuSettings],[function(e){var d=e.menuSettings;if(d.locale){MathJax.Localization.resetLocale(d.locale)}var g=e.menuSettings.renderer,b=e.jax;if(g){var c="output/"+g;b.sort();for(var f=0,a=b.length;f<a;f++){if(b[f].substr(0,7)==="output/"){break}}if(f==a-1){b.pop()}else{while(f<a){if(b[f]===c){b.splice(f,1);break}f++}}b.unshift(c)}if(d.CHTMLpreview!=null){if(d.FastPreview==null){d.FastPreview=d.CHTMLpreview}delete d.CHTMLpreview}if(d.FastPreview&&!MathJax.Extension["fast-preview"]){MathJax.Hub.config.extensions.push("fast-preview.js")}if(e.menuSettings.assistiveMML&&!MathJax.Extension.AssistiveMML){MathJax.Hub.config.extensions.push("AssistiveMML.js")}},MathJax.Hub.config],["Post",this.signal,"End Cookie"])},Styles:function(){return this.queue.Push(["Post",this.signal,"Begin Styles"],["loadArray",this,MathJax.Hub.config.styleSheets,"config"],["Styles",MathJax.Ajax,MathJax.Hub.config.styles],["Post",this.signal,"End Styles"])},Jax:function(){var f=MathJax.Hub.config,c=MathJax.Hub.outputJax;for(var g=0,b=f.jax.length,d=0;g<b;g++){var e=f.jax[g].substr(7);if(f.jax[g].substr(0,7)==="output/"&&c.order[e]==null){c.order[e]=d;d++}}var a=MathJax.Callback.Queue();return a.Push(["Post",this.signal,"Begin Jax"],["loadArray",this,f.jax,"jax","config.js"],["Post",this.signal,"End Jax"])},Extensions:function(){var a=MathJax.Callback.Queue();return a.Push(["Post",this.signal,"Begin Extensions"],["loadArray",this,MathJax.Hub.config.extensions,"extensions"],["Post",this.signal,"End Extensions"])},Message:function(){MathJax.Message.Init(true)},Menu:function(){var b=MathJax.Hub.config.menuSettings,a=MathJax.Hub.outputJax,d;for(var c in a){if(a.hasOwnProperty(c)){if(a[c].length){d=a[c];break}}}if(d&&d.length){if(b.renderer&&b.renderer!==d[0].id){d.unshift(MathJax.OutputJax[b.renderer])}b.renderer=d[0].id}},Hash:function(){if(MathJax.Hub.config.positionToHash&&document.location.hash&&document.body&&document.body.scrollIntoView){var d=document.location.hash.substr(1);var f=document.getElementById(d);if(!f){var c=document.getElementsByTagName("a");for(var e=0,b=c.length;e<b;e++){if(c[e].name===d){f=c[e];break}}}if(f){while(!f.scrollIntoView){f=f.parentNode}f=this.HashCheck(f);if(f&&f.scrollIntoView){setTimeout(function(){f.scrollIntoView(true)},1)}}}},HashCheck:function(b){var a=MathJax.Hub.getJaxFor(b);if(a&&MathJax.OutputJax[a.outputJax].hashCheck){b=MathJax.OutputJax[a.outputJax].hashCheck(b)}return b},MenuZoom:function(){if(MathJax.Hub.config.showMathMenu){if(!MathJax.Extension.MathMenu){setTimeout(function(){MathJax.Callback.Queue(["Require",MathJax.Ajax,"[MathJax]/extensions/MathMenu.js",{}],["loadDomain",MathJax.Localization,"MathMenu"])},1000)}else{setTimeout(MathJax.Callback(["loadDomain",MathJax.Localization,"MathMenu"]),1000)}if(!MathJax.Extension.MathZoom){setTimeout(MathJax.Callback(["Require",MathJax.Ajax,"[MathJax]/extensions/MathZoom.js",{}]),2000)}}},onLoad:function(){var a=this.onload=MathJax.Callback(function(){MathJax.Hub.Startup.signal.Post("onLoad")});if(document.body&&document.readyState){if(MathJax.Hub.Browser.isMSIE){if(document.readyState==="complete"){return[a]}}else{if(document.readyState!=="loading"){return[a]}}}if(window.addEventListener){window.addEventListener("load",a,false);if(!this.params.noDOMContentEvent){window.addEventListener("DOMContentLoaded",a,false)}}else{if(window.attachEvent){window.attachEvent("onload",a)}else{window.onload=a}}return a},Typeset:function(a,b){if(MathJax.Hub.config.skipStartupTypeset){return function(){}}return this.queue.Push(["Post",this.signal,"Begin Typeset"],["Typeset",MathJax.Hub,a,b],["Post",this.signal,"End Typeset"])},URL:function(b,a){if(!a.match(/^([a-z]+:\/\/|\[|\/)/)){a="[MathJax]/"+b+"/"+a}return a},loadArray:function(b,f,c,a){if(b){if(!MathJax.Object.isArray(b)){b=[b]}if(b.length){var h=MathJax.Callback.Queue(),j={},e;for(var g=0,d=b.length;g<d;g++){e=this.URL(f,b[g]);if(c){e+="/"+c}if(a){h.Push(["Require",MathJax.Ajax,e,j])}else{h.Push(MathJax.Ajax.Require(e,j))}}return h.Push({})}}return null}};(function(d){var b=window[d],e="["+d+"]";var c=b.Hub,a=b.Ajax,f=b.Callback;var g=MathJax.Object.Subclass({JAXFILE:"jax.js",require:null,config:{},Init:function(i,h){if(arguments.length===0){return this}return(this.constructor.Subclass(i,h))()},Augment:function(k,j){var i=this.constructor,h={};if(k!=null){for(var l in k){if(k.hasOwnProperty(l)){if(typeof k[l]==="function"){i.protoFunction(l,k[l])}else{h[l]=k[l]}}}if(k.toString!==i.prototype.toString&&k.toString!=={}.toString){i.protoFunction("toString",k.toString)}}c.Insert(i.prototype,h);i.Augment(null,j);return this},Translate:function(h,i){throw Error(this.directory+"/"+this.JAXFILE+" failed to define the Translate() method")},Register:function(h){},Config:function(){this.config=c.CombineConfig(this.id,this.config);if(this.config.Augment){this.Augment(this.config.Augment)}},Startup:function(){},loadComplete:function(i){if(i==="config.js"){return a.loadComplete(this.directory+"/"+i)}else{var h=f.Queue();h.Push(c.Register.StartupHook("End Config",{}),["Post",c.Startup.signal,this.id+" Jax Config"],["Config",this],["Post",c.Startup.signal,this.id+" Jax Require"],[function(j){return MathJax.Hub.Startup.loadArray(j.require,this.directory)},this],[function(j,k){return MathJax.Hub.Startup.loadArray(j.extensions,"extensions/"+k)},this.config||{},this.id],["Post",c.Startup.signal,this.id+" Jax Startup"],["Startup",this],["Post",c.Startup.signal,this.id+" Jax Ready"]);if(this.copyTranslate){h.Push([function(j){j.preProcess=j.preTranslate;j.Process=j.Translate;j.postProcess=j.postTranslate},this.constructor.prototype])}return h.Push(["loadComplete",a,this.directory+"/"+i])}}},{id:"Jax",version:"2.7.0",directory:e+"/jax",extensionDir:e+"/extensions"});b.InputJax=g.Subclass({elementJax:"mml",sourceMenuTitle:["Original","Original Form"],copyTranslate:true,Process:function(l,q){var j=f.Queue(),o;var k=this.elementJax;if(!b.Object.isArray(k)){k=[k]}for(var n=0,h=k.length;n<h;n++){o=b.ElementJax.directory+"/"+k[n]+"/"+this.JAXFILE;if(!this.require){this.require=[]}else{if(!b.Object.isArray(this.require)){this.require=[this.require]}}this.require.push(o);j.Push(a.Require(o))}o=this.directory+"/"+this.JAXFILE;var p=j.Push(a.Require(o));if(!p.called){this.constructor.prototype.Process=function(){if(!p.called){return p}throw Error(o+" failed to load properly")}}k=c.outputJax["jax/"+k[0]];if(k){j.Push(a.Require(k[0].directory+"/"+this.JAXFILE))}return j.Push({})},needsUpdate:function(h){var i=h.SourceElement();return(h.originalText!==b.HTML.getScript(i))},Register:function(h){if(!c.inputJax){c.inputJax={}}c.inputJax[h]=this}},{id:"InputJax",version:"2.7.0",directory:g.directory+"/input",extensionDir:g.extensionDir});b.OutputJax=g.Subclass({copyTranslate:true,preProcess:function(j){var i,h=this.directory+"/"+this.JAXFILE;this.constructor.prototype.preProcess=function(k){if(!i.called){return i}throw Error(h+" failed to load properly")};i=a.Require(h);return i},Register:function(i){var h=c.outputJax;if(!h[i]){h[i]=[]}if(h[i].length&&(this.id===c.config.menuSettings.renderer||(h.order[this.id]||0)<(h.order[h[i][0].id]||0))){h[i].unshift(this)}else{h[i].push(this)}if(!this.require){this.require=[]}else{if(!b.Object.isArray(this.require)){this.require=[this.require]}}this.require.push(b.ElementJax.directory+"/"+(i.split(/\//)[1])+"/"+this.JAXFILE)},Remove:function(h){}},{id:"OutputJax",version:"2.7.0",directory:g.directory+"/output",extensionDir:g.extensionDir,fontDir:e+(b.isPacked?"":"/..")+"/fonts",imageDir:e+(b.isPacked?"":"/..")+"/images"});b.ElementJax=g.Subclass({Init:function(i,h){return this.constructor.Subclass(i,h)},inputJax:null,outputJax:null,inputID:null,originalText:"",mimeType:"",sourceMenuTitle:["MathMLcode","MathML Code"],Text:function(i,j){var h=this.SourceElement();b.HTML.setScript(h,i);h.MathJax.state=this.STATE.UPDATE;return c.Update(h,j)},Reprocess:function(i){var h=this.SourceElement();h.MathJax.state=this.STATE.UPDATE;return c.Reprocess(h,i)},Update:function(h){return this.Rerender(h)},Rerender:function(i){var h=this.SourceElement();h.MathJax.state=this.STATE.OUTPUT;return c.Process(h,i)},Remove:function(h){if(this.hover){this.hover.clear(this)}b.OutputJax[this.outputJax].Remove(this);if(!h){c.signal.Post(["Remove Math",this.inputID]);this.Detach()}},needsUpdate:function(){return b.InputJax[this.inputJax].needsUpdate(this)},SourceElement:function(){return document.getElementById(this.inputID)},Attach:function(i,j){var h=i.MathJax.elementJax;if(i.MathJax.state===this.STATE.UPDATE){h.Clone(this)}else{h=i.MathJax.elementJax=this;if(i.id){this.inputID=i.id}else{i.id=this.inputID=b.ElementJax.GetID();this.newID=1}}h.originalText=b.HTML.getScript(i);h.inputJax=j;if(h.root){h.root.inputID=h.inputID}return h},Detach:function(){var h=this.SourceElement();if(!h){return}try{delete h.MathJax}catch(i){h.MathJax=null}if(this.newID){h.id=""}},Clone:function(h){var i;for(i in this){if(!this.hasOwnProperty(i)){continue}if(typeof(h[i])==="undefined"&&i!=="newID"){delete this[i]}}for(i in h){if(!h.hasOwnProperty(i)){continue}if(typeof(this[i])==="undefined"||(this[i]!==h[i]&&i!=="inputID")){this[i]=h[i]}}}},{id:"ElementJax",version:"2.7.0",directory:g.directory+"/element",extensionDir:g.extensionDir,ID:0,STATE:{PENDING:1,PROCESSED:2,UPDATE:3,OUTPUT:4},GetID:function(){this.ID++;return"MathJax-Element-"+this.ID},Subclass:function(){var h=g.Subclass.apply(this,arguments);h.loadComplete=this.prototype.loadComplete;return h}});b.ElementJax.prototype.STATE=b.ElementJax.STATE;b.OutputJax.Error={id:"Error",version:"2.7.0",config:{},errors:0,ContextMenu:function(){return b.Extension.MathEvents.Event.ContextMenu.apply(b.Extension.MathEvents.Event,arguments)},Mousedown:function(){return b.Extension.MathEvents.Event.AltContextMenu.apply(b.Extension.MathEvents.Event,arguments)},getJaxFromMath:function(h){return(h.nextSibling.MathJax||{}).error},Jax:function(j,i){var h=MathJax.Hub.inputJax[i.type.replace(/ *;(.|\s)*/,"")];this.errors++;return{inputJax:(h||{id:"Error"}).id,outputJax:"Error",inputID:"MathJax-Error-"+this.errors,sourceMenuTitle:["ErrorMessage","Error Message"],sourceMenuFormat:"Error",originalText:MathJax.HTML.getScript(i),errorText:j}}};b.InputJax.Error={id:"Error",version:"2.7.0",config:{},sourceMenuTitle:["Original","Original Form"]}})("MathJax");(function(o){var h=window[o];if(!h){h=window[o]={}}var d=h.Hub;var s=d.Startup;var w=d.config;var g=document.head||(document.getElementsByTagName("head")[0]);if(!g){g=document.childNodes[0]}var b=(document.documentElement||document).getElementsByTagName("script");if(b.length===0&&g.namespaceURI){b=document.getElementsByTagNameNS(g.namespaceURI,"script")}var f=new RegExp("(^|/)"+o+"\\.js(\\?.*)?$");for(var q=b.length-1;q>=0;q--){if((b[q].src||"").match(f)){s.script=b[q].innerHTML;if(RegExp.$2){var t=RegExp.$2.substr(1).split(/\&/);for(var p=0,l=t.length;p<l;p++){var n=t[p].match(/(.*)=(.*)/);if(n){s.params[unescape(n[1])]=unescape(n[2])}else{s.params[t[p]]=true}}}w.root=b[q].src.replace(/(^|\/)[^\/]*(\?.*)?$/,"").replace(/^(https?:\/\/cdn.mathjax.org\/mathjax\/)(latest)/,"$1"+h.version.split(/\./).slice(0,2).join(".")+"-$2");h.Ajax.config.root=w.root;h.Ajax.params=s.params;break}}var k=navigator.userAgent;var a={isMac:(navigator.platform.substr(0,3)==="Mac"),isPC:(navigator.platform.substr(0,3)==="Win"),isMSIE:("ActiveXObject" in window&&"clipboardData" in window),isEdge:("MSGestureEvent" in window&&"chrome" in window&&window.chrome.loadTimes==null),isFirefox:(!!k.match(/Gecko\//)&&!k.match(/like Gecko/)),isSafari:(!!k.match(/ (Apple)?WebKit\//)&&!k.match(/ like iPhone /)&&(!window.chrome||window.chrome.app==null)),isChrome:("chrome" in window&&window.chrome.loadTimes!=null),isOpera:("opera" in window&&window.opera.version!=null),isKonqueror:("konqueror" in window&&navigator.vendor=="KDE"),versionAtLeast:function(y){var x=(this.version).split(".");y=(new String(y)).split(".");for(var z=0,j=y.length;z<j;z++){if(x[z]!=y[z]){return parseInt(x[z]||"0")>=parseInt(y[z])}}return true},Select:function(j){var i=j[d.Browser];if(i){return i(d.Browser)}return null}};var e=k.replace(/^Mozilla\/(\d+\.)+\d+ /,"").replace(/[a-z][-a-z0-9._: ]+\/\d+[^ ]*-[^ ]*\.([a-z][a-z])?\d+ /i,"").replace(/Gentoo |Ubuntu\/(\d+\.)*\d+ (\([^)]*\) )?/,"");d.Browser=d.Insert(d.Insert(new String("Unknown"),{version:"0.0"}),a);for(var v in a){if(a.hasOwnProperty(v)){if(a[v]&&v.substr(0,2)==="is"){v=v.slice(2);if(v==="Mac"||v==="PC"){continue}d.Browser=d.Insert(new String(v),a);var r=new RegExp(".*(Version/| Trident/.*; rv:)((?:\\d+\\.)+\\d+)|.*("+v+")"+(v=="MSIE"?" ":"/")+"((?:\\d+\\.)*\\d+)|(?:^|\\(| )([a-z][-a-z0-9._: ]+|(?:Apple)?WebKit)/((?:\\d+\\.)+\\d+)");var u=r.exec(e)||["","","","unknown","0.0"];d.Browser.name=(u[1]!=""?v:(u[3]||u[5]));d.Browser.version=u[2]||u[4]||u[6];break}}}try{d.Browser.Select({Safari:function(j){var i=parseInt((String(j.version).split("."))[0]);if(i>85){j.webkit=j.version}if(i>=538){j.version="8.0"}else{if(i>=537){j.version="7.0"}else{if(i>=536){j.version="6.0"}else{if(i>=534){j.version="5.1"}else{if(i>=533){j.version="5.0"}else{if(i>=526){j.version="4.0"}else{if(i>=525){j.version="3.1"}else{if(i>500){j.version="3.0"}else{if(i>400){j.version="2.0"}else{if(i>85){j.version="1.0"}}}}}}}}}}j.webkit=(navigator.appVersion.match(/WebKit\/(\d+)\./))[1];j.isMobile=(navigator.appVersion.match(/Mobile/i)!=null);j.noContextMenu=j.isMobile},Firefox:function(j){if((j.version==="0.0"||k.match(/Firefox/)==null)&&navigator.product==="Gecko"){var m=k.match(/[\/ ]rv:(\d+\.\d.*?)[\) ]/);if(m){j.version=m[1]}else{var i=(navigator.buildID||navigator.productSub||"0").substr(0,8);if(i>="20111220"){j.version="9.0"}else{if(i>="20111120"){j.version="8.0"}else{if(i>="20110927"){j.version="7.0"}else{if(i>="20110816"){j.version="6.0"}else{if(i>="20110621"){j.version="5.0"}else{if(i>="20110320"){j.version="4.0"}else{if(i>="20100121"){j.version="3.6"}else{if(i>="20090630"){j.version="3.5"}else{if(i>="20080617"){j.version="3.0"}else{if(i>="20061024"){j.version="2.0"}}}}}}}}}}}}j.isMobile=(navigator.appVersion.match(/Android/i)!=null||k.match(/ Fennec\//)!=null||k.match(/Mobile/)!=null)},Chrome:function(i){i.noContextMenu=i.isMobile=!!navigator.userAgent.match(/ Mobile[ \/]/)},Opera:function(i){i.version=opera.version()},Edge:function(i){i.isMobile=!!navigator.userAgent.match(/ Phone/)},MSIE:function(j){j.isMobile=!!navigator.userAgent.match(/ Phone/);j.isIE9=!!(document.documentMode&&(window.performance||window.msPerformance));MathJax.HTML.setScriptBug=!j.isIE9||document.documentMode<9;MathJax.Hub.msieHTMLCollectionBug=(document.documentMode<9);if(document.documentMode<10&&!s.params.NoMathPlayer){try{new ActiveXObject("MathPlayer.Factory.1");j.hasMathPlayer=true}catch(m){}try{if(j.hasMathPlayer){var i=document.createElement("object");i.id="mathplayer";i.classid="clsid:32F66A20-7614-11D4-BD11-00104BD3F987";g.appendChild(i);document.namespaces.add("m","http://www.w3.org/1998/Math/MathML");j.mpNamespace=true;if(document.readyState&&(document.readyState==="loading"||document.readyState==="interactive")){document.write('<?import namespace="m" implementation="#MathPlayer">');j.mpImported=true}}else{document.namespaces.add("mjx_IE_fix","http://www.w3.org/1999/xlink")}}catch(m){}}}})}catch(c){console.error(c.message)}d.Browser.Select(MathJax.Message.browsers);if(h.AuthorConfig&&typeof h.AuthorConfig.AuthorInit==="function"){h.AuthorConfig.AuthorInit()}d.queue=h.Callback.Queue();d.queue.Push(["Post",s.signal,"Begin"],["Config",s],["Cookie",s],["Styles",s],["Message",s],function(){var i=h.Callback.Queue(s.Jax(),s.Extensions());return i.Push({})},["Menu",s],s.onLoad(),function(){MathJax.isReady=true},["Typeset",s],["Hash",s],["MenuZoom",s],["Post",s.signal,"End"])})("MathJax")}};
```
