## Supplementary material for "Artistoo: build, share, and explore simulations of cells and tissues in the web browser": reveal.js

```
/*!
 * reveal.js
 * http://revealjs.com
 * MIT licensed
 *
 * Copyright (C) 2017 Hakim El Hattab, http://hakim.se
 */
(function( root, factory ) {
	if( typeof define === 'function' && define.amd ) {
		// AMD. Register as an anonymous module.
		define( function() {
			root.Reveal = factory();
			return root.Reveal;
		} );
	} else if( typeof exports === 'object' ) {
		// Node. Does not work with strict CommonJS.
		module.exports = factory();
	} else {
		// Browser globals.
		root.Reveal = factory();
	}
}( this, function() {

	'use strict';
	var Reveal;

	// The reveal.js version
	var VERSION = '3.6.0';

	var SLIDES_SELECTOR = '.slides section',
		HORIZONTAL_SLIDES_SELECTOR = '.slides>section',
		VERTICAL_SLIDES_SELECTOR = '.slides>section.present>section',
		HOME_SLIDE_SELECTOR = '.slides>section:first-of-type',
		UA = navigator.userAgent,

		// Configuration defaults, can be overridden at initialization time
		config = {

			// The "normal" size of the presentation, aspect ratio will be preserved
			// when the presentation is scaled to fit different resolutions
			width: 960,
			height: 700,

			// Factor of the display size that should remain empty around the content
			margin: 0.04,

			// Bounds for smallest/largest possible scale to apply to content
			minScale: 0.2,
			maxScale: 2.0,

			// Display presentation control arrows
			controls: true,

			// Help the user learn the controls by providing hints, for example by
			// bouncing the down arrow when they first encounter a vertical slide
			controlsTutorial: true,

			// Determines where controls appear, "edges" or "bottom-right"
			controlsLayout: 'bottom-right',

			// Visibility rule for backwards navigation arrows; "faded", "hidden"
			// or "visible"
			controlsBackArrows: 'faded',

			// Display a presentation progress bar
			progress: true,

			// Display the page number of the current slide
			slideNumber: false,

			// Determine which displays to show the slide number on
			showSlideNumber: 'all',

			// Push each slide change to the browser history
			history: false,

			// Enable keyboard shortcuts for navigation
			keyboard: true,

			// Optional function that blocks keyboard events when retuning false
			keyboardCondition: null,

			// Enable the slide overview mode
			overview: true,

			// Vertical centering of slides
			center: true,

			// Enables touch navigation on devices with touch input
			touch: true,

			// Loop the presentation
			loop: false,

			// Change the presentation direction to be RTL
			rtl: false,

			// Randomizes the order of slides each time the presentation loads
			shuffle: false,

			// Turns fragments on and off globally
			fragments: true,

			// Flags if the presentation is running in an embedded mode,
			// i.e. contained within a limited portion of the screen
			embedded: false,

			// Flags if we should show a help overlay when the question-mark
			// key is pressed
			help: true,

			// Flags if it should be possible to pause the presentation (blackout)
			pause: true,

			// Flags if speaker notes should be visible to all viewers
			showNotes: false,

			// Global override for autolaying embedded media (video/audio/iframe)
			// - null:   Media will only autoplay if data-autoplay is present
			// - true:   All media will autoplay, regardless of individual setting
			// - false:  No media will autoplay, regardless of individual setting
			autoPlayMedia: null,

			// Controls automatic progression to the next slide
			// - 0:      Auto-sliding only happens if the data-autoslide HTML attribute
			//           is present on the current slide or fragment
			// - 1+:     All slides will progress automatically at the given interval
			// - false:  No auto-sliding, even if data-autoslide is present
			autoSlide: 0,

			// Stop auto-sliding after user input
			autoSlideStoppable: true,

			// Use this method for navigation when auto-sliding (defaults to navigateNext)
			autoSlideMethod: null,

			// Enable slide navigation via mouse wheel
			mouseWheel: false,

			// Apply a 3D roll to links on hover
			rollingLinks: false,

			// Hides the address bar on mobile devices
			hideAddressBar: true,

			// Opens links in an iframe preview overlay
			// Add `data-preview-link` and `data-preview-link="false"` to customise each link
			// individually
			previewLinks: false,

			// Exposes the reveal.js API through window.postMessage
			postMessage: true,

			// Dispatches all reveal.js events to the parent window through postMessage
			postMessageEvents: false,

			// Focuses body when page changes visibility to ensure keyboard shortcuts work
			focusBodyOnPageVisibilityChange: true,

			// Transition style
			transition: 'slide', // none/fade/slide/convex/concave/zoom

			// Transition speed
			transitionSpeed: 'default', // default/fast/slow

			// Transition style for full page slide backgrounds
			backgroundTransition: 'fade', // none/fade/slide/convex/concave/zoom

			// Parallax background image
			parallaxBackgroundImage: '', // CSS syntax, e.g. "a.jpg"

			// Parallax background size
			parallaxBackgroundSize: '', // CSS syntax, e.g. "3000px 2000px"

			// Amount of pixels to move the parallax background per slide step
			parallaxBackgroundHorizontal: null,
			parallaxBackgroundVertical: null,

			// The maximum number of pages a single slide can expand onto when printing
			// to PDF, unlimited by default
			pdfMaxPagesPerSlide: Number.POSITIVE_INFINITY,

			// Offset used to reduce the height of content within exported PDF pages.
			// This exists to account for environment differences based on how you
			// print to PDF. CLI printing options, like phantomjs and wkpdf, can end
			// on precisely the total height of the document whereas in-browser
			// printing has to end one pixel before.
			pdfPageHeightOffset: -1,

			// Number of slides away from the current that are visible
			viewDistance: 3,

			// The display mode that will be used to show slides
			display: 'block',

			// Script dependencies to load
			dependencies: []

		},

		// Flags if Reveal.initialize() has been called
		initialized = false,

		// Flags if reveal.js is loaded (has dispatched the 'ready' event)
		loaded = false,

		// Flags if the overview mode is currently active
		overview = false,

		// Holds the dimensions of our overview slides, including margins
		overviewSlideWidth = null,
		overviewSlideHeight = null,

		// The horizontal and vertical index of the currently active slide
		indexh,
		indexv,

		// The previous and current slide HTML elements
		previousSlide,
		currentSlide,

		previousBackground,

		// Remember which directions that the user has navigated towards
		hasNavigatedRight = false,
		hasNavigatedDown = false,

		// Slides may hold a data-state attribute which we pick up and apply
		// as a class to the body. This list contains the combined state of
		// all current slides.
		state = [],

		// The current scale of the presentation (see width/height config)
		scale = 1,

		// CSS transform that is currently applied to the slides container,
		// split into two groups
		slidesTransform = { layout: '', overview: '' },

		// Cached references to DOM elements
		dom = {},

		// Features supported by the browser, see #checkCapabilities()
		features = {},

		// Client is a mobile device, see #checkCapabilities()
		isMobileDevice,

		// Client is a desktop Chrome, see #checkCapabilities()
		isChrome,

		// Throttles mouse wheel navigation
		lastMouseWheelStep = 0,

		// Delays updates to the URL due to a Chrome thumbnailer bug
		writeURLTimeout = 0,

		// Flags if the interaction event listeners are bound
		eventsAreBound = false,

		// The current auto-slide duration
		autoSlide = 0,

		// Auto slide properties
		autoSlidePlayer,
		autoSlideTimeout = 0,
		autoSlideStartTime = -1,
		autoSlidePaused = false,

		// Holds information about the currently ongoing touch input
		touch = {
			startX: 0,
			startY: 0,
			startSpan: 0,
			startCount: 0,
			captured: false,
			threshold: 40
		},

		// Holds information about the keyboard shortcuts
		keyboardShortcuts = {
			'N  ,  SPACE':			'Next slide',
			'P':					'Previous slide',
			'&#8592;  ,  H':		'Navigate left',
			'&#8594;  ,  L':		'Navigate right',
			'&#8593;  ,  K':		'Navigate up',
			'&#8595;  ,  J':		'Navigate down',
			'Home':					'First slide',
			'End':					'Last slide',
			'B  ,  .':				'Pause',
			'F':					'Fullscreen',
			'ESC, O':				'Slide overview'
		};

	/**
	 * Starts up the presentation if the client is capable.
	 */
	function initialize( options ) {

		// Make sure we only initialize once
		if( initialized === true ) return;

		initialized = true;

		checkCapabilities();

		if( !features.transforms2d && !features.transforms3d ) {
			document.body.setAttribute( 'class', 'no-transforms' );

			// Since JS won't be running any further, we load all lazy
			// loading elements upfront
			var images = toArray( document.getElementsByTagName( 'img' ) ),
				iframes = toArray( document.getElementsByTagName( 'iframe' ) );

			var lazyLoadable = images.concat( iframes );

			for( var i = 0, len = lazyLoadable.length; i < len; i++ ) {
				var element = lazyLoadable[i];
				if( element.getAttribute( 'data-src' ) ) {
					element.setAttribute( 'src', element.getAttribute( 'data-src' ) );
					element.removeAttribute( 'data-src' );
				}
			}

			// If the browser doesn't support core features we won't be
			// using JavaScript to control the presentation
			return;
		}

		// Cache references to key DOM elements
		dom.wrapper = document.querySelector( '.reveal' );
		dom.slides = document.querySelector( '.reveal .slides' );

		// Force a layout when the whole page, incl fonts, has loaded
		window.addEventListener( 'load', layout, false );

		var query = Reveal.getQueryHash();

		// Do not accept new dependencies via query config to avoid
		// the potential of malicious script injection
		if( typeof query['dependencies'] !== 'undefined' ) delete query['dependencies'];

		// Copy options over to our config object
		extend( config, options );
		extend( config, query );

		// Hide the address bar in mobile browsers
		hideAddressBar();

		// Loads the dependencies and continues to #start() once done
		load();

	}

	/**
	 * Inspect the client to see what it's capable of, this
	 * should only happens once per runtime.
	 */
	function checkCapabilities() {

		isMobileDevice = /(iphone|ipod|ipad|android)/gi.test( UA );
		isChrome = /chrome/i.test( UA ) && !/edge/i.test( UA );

		var testElement = document.createElement( 'div' );

		features.transforms3d = 'WebkitPerspective' in testElement.style ||
								'MozPerspective' in testElement.style ||
								'msPerspective' in testElement.style ||
								'OPerspective' in testElement.style ||
								'perspective' in testElement.style;

		features.transforms2d = 'WebkitTransform' in testElement.style ||
								'MozTransform' in testElement.style ||
								'msTransform' in testElement.style ||
								'OTransform' in testElement.style ||
								'transform' in testElement.style;

		features.requestAnimationFrameMethod = window.requestAnimationFrame || window.webkitRequestAnimationFrame || window.mozRequestAnimationFrame;
		features.requestAnimationFrame = typeof features.requestAnimationFrameMethod === 'function';

		features.canvas = !!document.createElement( 'canvas' ).getContext;

		// Transitions in the overview are disabled in desktop and
		// Safari due to lag
		features.overviewTransitions = !/Version\/[\d\.]+.*Safari/.test( UA );

		// Flags if we should use zoom instead of transform to scale
		// up slides. Zoom produces crisper results but has a lot of
		// xbrowser quirks so we only use it in whitelsited browsers.
		features.zoom = 'zoom' in testElement.style && !isMobileDevice &&
						( isChrome || /Version\/[\d\.]+.*Safari/.test( UA ) );

	}

    /**
     * Loads the dependencies of reveal.js. Dependencies are
     * defined via the configuration option 'dependencies'
     * and will be loaded prior to starting/binding reveal.js.
     * Some dependencies may have an 'async' flag, if so they
     * will load after reveal.js has been started up.
     */
	function load() {

		var scripts = [],
			scriptsAsync = [],
			scriptsToPreload = 0;

		// Called once synchronous scripts finish loading
		function proceed() {
			if( scriptsAsync.length ) {
				// Load asynchronous scripts
				head.js.apply( null, scriptsAsync );
			}

			start();
		}

		function loadScript( s ) {
			head.ready( s.src.match( /([\w\d_\-]*)\.?js$|[^\\\/]*$/i )[0], function() {
				// Extension may contain callback functions
				if( typeof s.callback === 'function' ) {
					s.callback.apply( this );
				}

				if( --scriptsToPreload === 0 ) {
					proceed();
				}
			});
		}

		for( var i = 0, len = config.dependencies.length; i < len; i++ ) {
			var s = config.dependencies[i];

			// Load if there's no condition or the condition is truthy
			if( !s.condition || s.condition() ) {
				if( s.async ) {
					scriptsAsync.push( s.src );
				}
				else {
					scripts.push( s.src );
				}

				loadScript( s );
			}
		}

		if( scripts.length ) {
			scriptsToPreload = scripts.length;

			// Load synchronous scripts
			head.js.apply( null, scripts );
		}
		else {
			proceed();
		}

	}

	/**
	 * Starts up reveal.js by binding input events and navigating
	 * to the current URL deeplink if there is one.
	 */
	function start() {

		loaded = true;

		// Make sure we've got all the DOM elements we need
		setupDOM();

		// Listen to messages posted to this window
		setupPostMessage();

		// Prevent the slides from being scrolled out of view
		setupScrollPrevention();

		// Resets all vertical slides so that only the first is visible
		resetVerticalSlides();

		// Updates the presentation to match the current configuration values
		configure();

		// Read the initial hash
		readURL();

		// Update all backgrounds
		updateBackground( true );

		// Notify listeners that the presentation is ready but use a 1ms
		// timeout to ensure it's not fired synchronously after #initialize()
		setTimeout( function() {
			// Enable transitions now that we're loaded
			dom.slides.classList.remove( 'no-transition' );

			dom.wrapper.classList.add( 'ready' );

			dispatchEvent( 'ready', {
				'indexh': indexh,
				'indexv': indexv,
				'currentSlide': currentSlide
			} );
		}, 1 );

		// Special setup and config is required when printing to PDF
		if( isPrintingPDF() ) {
			removeEventListeners();

			// The document needs to have loaded for the PDF layout
			// measurements to be accurate
			if( document.readyState === 'complete' ) {
				setupPDF();
			}
			else {
				window.addEventListener( 'load', setupPDF );
			}
		}

	}

	/**
	 * Finds and stores references to DOM elements which are
	 * required by the presentation. If a required element is
	 * not found, it is created.
	 */
	function setupDOM() {

		// Prevent transitions while we're loading
		dom.slides.classList.add( 'no-transition' );

		if( isMobileDevice ) {
			dom.wrapper.classList.add( 'no-hover' );
		}
		else {
			dom.wrapper.classList.remove( 'no-hover' );
		}

		if( /iphone/gi.test( UA ) ) {
			dom.wrapper.classList.add( 'ua-iphone' );
		}
		else {
			dom.wrapper.classList.remove( 'ua-iphone' );
		}

		// Background element
		dom.background = createSingletonNode( dom.wrapper, 'div', 'backgrounds', null );

		// Progress bar
		dom.progress = createSingletonNode( dom.wrapper, 'div', 'progress', '<span></span>' );
		dom.progressbar = dom.progress.querySelector( 'span' );

		// Arrow controls
		dom.controls = createSingletonNode( dom.wrapper, 'aside', 'controls',
			'<button class="navigate-left" aria-label="previous slide"><div class="controls-arrow"></div></button>' +
			'<button class="navigate-right" aria-label="next slide"><div class="controls-arrow"></div></button>' +
			'<button class="navigate-up" aria-label="above slide"><div class="controls-arrow"></div></button>' +
			'<button class="navigate-down" aria-label="below slide"><div class="controls-arrow"></div></button>' );

		// Slide number
		dom.slideNumber = createSingletonNode( dom.wrapper, 'div', 'slide-number', '' );

		// Element containing notes that are visible to the audience
		dom.speakerNotes = createSingletonNode( dom.wrapper, 'div', 'speaker-notes', null );
		dom.speakerNotes.setAttribute( 'data-prevent-swipe', '' );
		dom.speakerNotes.setAttribute( 'tabindex', '0' );

		// Overlay graphic which is displayed during the paused mode
		createSingletonNode( dom.wrapper, 'div', 'pause-overlay', null );

		dom.wrapper.setAttribute( 'role', 'application' );

		// There can be multiple instances of controls throughout the page
		dom.controlsLeft = toArray( document.querySelectorAll( '.navigate-left' ) );
		dom.controlsRight = toArray( document.querySelectorAll( '.navigate-right' ) );
		dom.controlsUp = toArray( document.querySelectorAll( '.navigate-up' ) );
		dom.controlsDown = toArray( document.querySelectorAll( '.navigate-down' ) );
		dom.controlsPrev = toArray( document.querySelectorAll( '.navigate-prev' ) );
		dom.controlsNext = toArray( document.querySelectorAll( '.navigate-next' ) );

		// The right and down arrows in the standard reveal.js controls
		dom.controlsRightArrow = dom.controls.querySelector( '.navigate-right' );
		dom.controlsDownArrow = dom.controls.querySelector( '.navigate-down' );

		dom.statusDiv = createStatusDiv();
	}

	/**
	 * Creates a hidden div with role aria-live to announce the
	 * current slide content. Hide the div off-screen to make it
	 * available only to Assistive Technologies.
	 *
	 * @return {HTMLElement}
	 */
	function createStatusDiv() {

		var statusDiv = document.getElementById( 'aria-status-div' );
		if( !statusDiv ) {
			statusDiv = document.createElement( 'div' );
			statusDiv.style.position = 'absolute';
			statusDiv.style.height = '1px';
			statusDiv.style.width = '1px';
			statusDiv.style.overflow = 'hidden';
			statusDiv.style.clip = 'rect( 1px, 1px, 1px, 1px )';
			statusDiv.setAttribute( 'id', 'aria-status-div' );
			statusDiv.setAttribute( 'aria-live', 'polite' );
			statusDiv.setAttribute( 'aria-atomic','true' );
			dom.wrapper.appendChild( statusDiv );
		}
		return statusDiv;

	}

	/**
	 * Converts the given HTML element into a string of text
	 * that can be announced to a screen reader. Hidden
	 * elements are excluded.
	 */
	function getStatusText( node ) {

		var text = '';

		// Text node
		if( node.nodeType === 3 ) {
			text += node.textContent;
		}
		// Element node
		else if( node.nodeType === 1 ) {

			var isAriaHidden = node.getAttribute( 'aria-hidden' );
			var isDisplayHidden = window.getComputedStyle( node )['display'] === 'none';
			if( isAriaHidden !== 'true' && !isDisplayHidden ) {

				toArray( node.childNodes ).forEach( function( child ) {
					text += getStatusText( child );
				} );

			}

		}

		return text;

	}

	/**
	 * Configures the presentation for printing to a static
	 * PDF.
	 */
	function setupPDF() {

		var slideSize = getComputedSlideSize( window.innerWidth, window.innerHeight );

		// Dimensions of the PDF pages
		var pageWidth = Math.floor( slideSize.width * ( 1 + config.margin ) ),
			pageHeight = Math.floor( slideSize.height * ( 1 + config.margin ) );

		// Dimensions of slides within the pages
		var slideWidth = slideSize.width,
			slideHeight = slideSize.height;

		// Let the browser know what page size we want to print
		injectStyleSheet( '@page{size:'+ pageWidth +'px '+ pageHeight +'px; margin: 0px;}' );

		// Limit the size of certain elements to the dimensions of the slide
		injectStyleSheet( '.reveal section>img, .reveal section>video, .reveal section>iframe{max-width: '+ slideWidth +'px; max-height:'+ slideHeight +'px}' );

		document.body.classList.add( 'print-pdf' );
		document.body.style.width = pageWidth + 'px';
		document.body.style.height = pageHeight + 'px';

		// Make sure stretch elements fit on slide
		layoutSlideContents( slideWidth, slideHeight );

		// Add each slide's index as attributes on itself, we need these
		// indices to generate slide numbers below
		toArray( dom.wrapper.querySelectorAll( HORIZONTAL_SLIDES_SELECTOR ) ).forEach( function( hslide, h ) {
			hslide.setAttribute( 'data-index-h', h );

			if( hslide.classList.contains( 'stack' ) ) {
				toArray( hslide.querySelectorAll( 'section' ) ).forEach( function( vslide, v ) {
					vslide.setAttribute( 'data-index-h', h );
					vslide.setAttribute( 'data-index-v', v );
				} );
			}
		} );

		// Slide and slide background layout
		toArray( dom.wrapper.querySelectorAll( SLIDES_SELECTOR ) ).forEach( function( slide ) {

			// Vertical stacks are not centred since their section
			// children will be
			if( slide.classList.contains( 'stack' ) === false ) {
				// Center the slide inside of the page, giving the slide some margin
				var left = ( pageWidth - slideWidth ) / 2,
					top = ( pageHeight - slideHeight ) / 2;

				var contentHeight = slide.scrollHeight;
				var numberOfPages = Math.max( Math.ceil( contentHeight / pageHeight ), 1 );

				// Adhere to configured pages per slide limit
				numberOfPages = Math.min( numberOfPages, config.pdfMaxPagesPerSlide );

				// Center slides vertically
				if( numberOfPages === 1 && config.center || slide.classList.contains( 'center' ) ) {
					top = Math.max( ( pageHeight - contentHeight ) / 2, 0 );
				}

				// Wrap the slide in a page element and hide its overflow
				// so that no page ever flows onto another
				var page = document.createElement( 'div' );
				page.className = 'pdf-page';
				page.style.height = ( ( pageHeight + config.pdfPageHeightOffset ) * numberOfPages ) + 'px';
				slide.parentNode.insertBefore( page, slide );
				page.appendChild( slide );

				// Position the slide inside of the page
				slide.style.left = left + 'px';
				slide.style.top = top + 'px';
				slide.style.width = slideWidth + 'px';

				if( slide.slideBackgroundElement ) {
					page.insertBefore( slide.slideBackgroundElement, slide );
				}

				// Inject notes if `showNotes` is enabled
				if( config.showNotes ) {

					// Are there notes for this slide?
					var notes = getSlideNotes( slide );
					if( notes ) {

						var notesSpacing = 8;
						var notesLayout = typeof config.showNotes === 'string' ? config.showNotes : 'inline';
						var notesElement = document.createElement( 'div' );
						notesElement.classList.add( 'speaker-notes' );
						notesElement.classList.add( 'speaker-notes-pdf' );
						notesElement.setAttribute( 'data-layout', notesLayout );
						notesElement.innerHTML = notes;

						if( notesLayout === 'separate-page' ) {
							page.parentNode.insertBefore( notesElement, page.nextSibling );
						}
						else {
							notesElement.style.left = notesSpacing + 'px';
							notesElement.style.bottom = notesSpacing + 'px';
							notesElement.style.width = ( pageWidth - notesSpacing*2 ) + 'px';
							page.appendChild( notesElement );
						}

					}

				}

				// Inject slide numbers if `slideNumbers` are enabled
				if( config.slideNumber && /all|print/i.test( config.showSlideNumber ) ) {
					var slideNumberH = parseInt( slide.getAttribute( 'data-index-h' ), 10 ) + 1,
						slideNumberV = parseInt( slide.getAttribute( 'data-index-v' ), 10 ) + 1;

					var numberElement = document.createElement( 'div' );
					numberElement.classList.add( 'slide-number' );
					numberElement.classList.add( 'slide-number-pdf' );
					numberElement.innerHTML = formatSlideNumber( slideNumberH, '.', slideNumberV );
					page.appendChild( numberElement );
				}
			}

		} );

		// Show all fragments
		toArray( dom.wrapper.querySelectorAll( SLIDES_SELECTOR + ' .fragment' ) ).forEach( function( fragment ) {
			fragment.classList.add( 'visible' );
		} );

		// Notify subscribers that the PDF layout is good to go
		dispatchEvent( 'pdf-ready' );

	}

	/**
	 * This is an unfortunate necessity. Some actions – such as
	 * an input field being focused in an iframe or using the
	 * keyboard to expand text selection beyond the bounds of
	 * a slide – can trigger our content to be pushed out of view.
	 * This scrolling can not be prevented by hiding overflow in
	 * CSS (we already do) so we have to resort to repeatedly
	 * checking if the slides have been offset :(
	 */
	function setupScrollPrevention() {

		setInterval( function() {
			if( dom.wrapper.scrollTop !== 0 || dom.wrapper.scrollLeft !== 0 ) {
				dom.wrapper.scrollTop = 0;
				dom.wrapper.scrollLeft = 0;
			}
		}, 1000 );

	}

	/**
	 * Creates an HTML element and returns a reference to it.
	 * If the element already exists the existing instance will
	 * be returned.
	 *
	 * @param {HTMLElement} container
	 * @param {string} tagname
	 * @param {string} classname
	 * @param {string} innerHTML
	 *
	 * @return {HTMLElement}
	 */
	function createSingletonNode( container, tagname, classname, innerHTML ) {

		// Find all nodes matching the description
		var nodes = container.querySelectorAll( '.' + classname );

		// Check all matches to find one which is a direct child of
		// the specified container
		for( var i = 0; i < nodes.length; i++ ) {
			var testNode = nodes[i];
			if( testNode.parentNode === container ) {
				return testNode;
			}
		}

		// If no node was found, create it now
		var node = document.createElement( tagname );
		node.className = classname;
		if( typeof innerHTML === 'string' ) {
			node.innerHTML = innerHTML;
		}
		container.appendChild( node );

		return node;

	}

	/**
	 * Creates the slide background elements and appends them
	 * to the background container. One element is created per
	 * slide no matter if the given slide has visible background.
	 */
	function createBackgrounds() {

		var printMode = isPrintingPDF();

		// Clear prior backgrounds
		dom.background.innerHTML = '';
		dom.background.classList.add( 'no-transition' );

		// Iterate over all horizontal slides
		toArray( dom.wrapper.querySelectorAll( HORIZONTAL_SLIDES_SELECTOR ) ).forEach( function( slideh ) {

			var backgroundStack = createBackground( slideh, dom.background );

			// Iterate over all vertical slides
			toArray( slideh.querySelectorAll( 'section' ) ).forEach( function( slidev ) {

				createBackground( slidev, backgroundStack );

				backgroundStack.classList.add( 'stack' );

			} );

		} );

		// Add parallax background if specified
		if( config.parallaxBackgroundImage ) {

			dom.background.style.backgroundImage = 'url("' + config.parallaxBackgroundImage + '")';
			dom.background.style.backgroundSize = config.parallaxBackgroundSize;

			// Make sure the below properties are set on the element - these properties are
			// needed for proper transitions to be set on the element via CSS. To remove
			// annoying background slide-in effect when the presentation starts, apply
			// these properties after short time delay
			setTimeout( function() {
				dom.wrapper.classList.add( 'has-parallax-background' );
			}, 1 );

		}
		else {

			dom.background.style.backgroundImage = '';
			dom.wrapper.classList.remove( 'has-parallax-background' );

		}

	}

	/**
	 * Creates a background for the given slide.
	 *
	 * @param {HTMLElement} slide
	 * @param {HTMLElement} container The element that the background
	 * should be appended to
	 * @return {HTMLElement} New background div
	 */
	function createBackground( slide, container ) {

		var data = {
			background: slide.getAttribute( 'data-background' ),
			backgroundSize: slide.getAttribute( 'data-background-size' ),
			backgroundImage: slide.getAttribute( 'data-background-image' ),
			backgroundVideo: slide.getAttribute( 'data-background-video' ),
			backgroundIframe: slide.getAttribute( 'data-background-iframe' ),
			backgroundColor: slide.getAttribute( 'data-background-color' ),
			backgroundRepeat: slide.getAttribute( 'data-background-repeat' ),
			backgroundPosition: slide.getAttribute( 'data-background-position' ),
			backgroundTransition: slide.getAttribute( 'data-background-transition' )
		};

		var element = document.createElement( 'div' );

		// Carry over custom classes from the slide to the background
		element.className = 'slide-background ' + slide.className.replace( /present|past|future/, '' );

		if( data.background ) {
			// Auto-wrap image urls in url(...)
			if( /^(http|file|\/\/)/gi.test( data.background ) || /\.(svg|png|jpg|jpeg|gif|bmp)([?#]|$)/gi.test( data.background ) ) {
				slide.setAttribute( 'data-background-image', data.background );
			}
			else {
				element.style.background = data.background;
			}
		}

		// Create a hash for this combination of background settings.
		// This is used to determine when two slide backgrounds are
		// the same.
		if( data.background || data.backgroundColor || data.backgroundImage || data.backgroundVideo || data.backgroundIframe ) {
			element.setAttribute( 'data-background-hash', data.background +
															data.backgroundSize +
															data.backgroundImage +
															data.backgroundVideo +
															data.backgroundIframe +
															data.backgroundColor +
															data.backgroundRepeat +
															data.backgroundPosition +
															data.backgroundTransition );
		}

		// Additional and optional background properties
		if( data.backgroundSize ) element.style.backgroundSize = data.backgroundSize;
		if( data.backgroundSize ) element.setAttribute( 'data-background-size', data.backgroundSize );
		if( data.backgroundColor ) element.style.backgroundColor = data.backgroundColor;
		if( data.backgroundRepeat ) element.style.backgroundRepeat = data.backgroundRepeat;
		if( data.backgroundPosition ) element.style.backgroundPosition = data.backgroundPosition;
		if( data.backgroundTransition ) element.setAttribute( 'data-background-transition', data.backgroundTransition );

		container.appendChild( element );

		// If backgrounds are being recreated, clear old classes
		slide.classList.remove( 'has-dark-background' );
		slide.classList.remove( 'has-light-background' );

		slide.slideBackgroundElement = element;

		// If this slide has a background color, add a class that
		// signals if it is light or dark. If the slide has no background
		// color, no class will be set
		var computedBackgroundStyle = window.getComputedStyle( element );
		if( computedBackgroundStyle && computedBackgroundStyle.backgroundColor ) {
			var rgb = colorToRgb( computedBackgroundStyle.backgroundColor );

			// Ignore fully transparent backgrounds. Some browsers return
			// rgba(0,0,0,0) when reading the computed background color of
			// an element with no background
			if( rgb && rgb.a !== 0 ) {
				if( colorBrightness( computedBackgroundStyle.backgroundColor ) < 128 ) {
					slide.classList.add( 'has-dark-background' );
				}
				else {
					slide.classList.add( 'has-light-background' );
				}
			}
		}

		return element;

	}

	/**
	 * Registers a listener to postMessage events, this makes it
	 * possible to call all reveal.js API methods from another
	 * window. For example:
	 *
	 * revealWindow.postMessage( JSON.stringify({
	 *   method: 'slide',
	 *   args: [ 2 ]
	 * }), '*' );
	 */
	function setupPostMessage() {

		if( config.postMessage ) {
			window.addEventListener( 'message', function ( event ) {
				var data = event.data;

				// Make sure we're dealing with JSON
				if( typeof data === 'string' && data.charAt( 0 ) === '{' && data.charAt( data.length - 1 ) === '}' ) {
					data = JSON.parse( data );

					// Check if the requested method can be found
					if( data.method && typeof Reveal[data.method] === 'function' ) {
						Reveal[data.method].apply( Reveal, data.args );
					}
				}
			}, false );
		}

	}

	/**
	 * Applies the configuration settings from the config
	 * object. May be called multiple times.
	 *
	 * @param {object} options
	 */
	function configure( options ) {

		var oldTransition = config.transition;

		// New config options may be passed when this method
		// is invoked through the API after initialization
		if( typeof options === 'object' ) extend( config, options );

		// Abort if reveal.js hasn't finished loading, config
		// changes will be applied automatically once loading
		// finishes
		if( loaded === false ) return;

		var numberOfSlides = dom.wrapper.querySelectorAll( SLIDES_SELECTOR ).length;

		// Remove the previously configured transition class
		dom.wrapper.classList.remove( oldTransition );

		// Force linear transition based on browser capabilities
		if( features.transforms3d === false ) config.transition = 'linear';

		dom.wrapper.classList.add( config.transition );

		dom.wrapper.setAttribute( 'data-transition-speed', config.transitionSpeed );
		dom.wrapper.setAttribute( 'data-background-transition', config.backgroundTransition );

		dom.controls.style.display = config.controls ? 'block' : 'none';
		dom.progress.style.display = config.progress ? 'block' : 'none';

		dom.controls.setAttribute( 'data-controls-layout', config.controlsLayout );
		dom.controls.setAttribute( 'data-controls-back-arrows', config.controlsBackArrows );

		if( config.shuffle ) {
			shuffle();
		}

		if( config.rtl ) {
			dom.wrapper.classList.add( 'rtl' );
		}
		else {
			dom.wrapper.classList.remove( 'rtl' );
		}

		if( config.center ) {
			dom.wrapper.classList.add( 'center' );
		}
		else {
			dom.wrapper.classList.remove( 'center' );
		}

		// Exit the paused mode if it was configured off
		if( config.pause === false ) {
			resume();
		}

		if( config.showNotes ) {
			dom.speakerNotes.setAttribute( 'data-layout', typeof config.showNotes === 'string' ? config.showNotes : 'inline' );
		}

		if( config.mouseWheel ) {
			document.addEventListener( 'DOMMouseScroll', onDocumentMouseScroll, false ); // FF
			document.addEventListener( 'mousewheel', onDocumentMouseScroll, false );
		}
		else {
			document.removeEventListener( 'DOMMouseScroll', onDocumentMouseScroll, false ); // FF
			document.removeEventListener( 'mousewheel', onDocumentMouseScroll, false );
		}

		// Rolling 3D links
		if( config.rollingLinks ) {
			enableRollingLinks();
		}
		else {
			disableRollingLinks();
		}

		// Iframe link previews
		if( config.previewLinks ) {
			enablePreviewLinks();
			disablePreviewLinks( '[data-preview-link=false]' );
		}
		else {
			disablePreviewLinks();
			enablePreviewLinks( '[data-preview-link]:not([data-preview-link=false])' );
		}

		// Remove existing auto-slide controls
		if( autoSlidePlayer ) {
			autoSlidePlayer.destroy();
			autoSlidePlayer = null;
		}

		// Generate auto-slide controls if needed
		if( numberOfSlides > 1 && config.autoSlide && config.autoSlideStoppable && features.canvas && features.requestAnimationFrame ) {
			autoSlidePlayer = new Playback( dom.wrapper, function() {
				return Math.min( Math.max( ( Date.now() - autoSlideStartTime ) / autoSlide, 0 ), 1 );
			} );

			autoSlidePlayer.on( 'click', onAutoSlidePlayerClick );
			autoSlidePaused = false;
		}

		// When fragments are turned off they should be visible
		if( config.fragments === false ) {
			toArray( dom.slides.querySelectorAll( '.fragment' ) ).forEach( function( element ) {
				element.classList.add( 'visible' );
				element.classList.remove( 'current-fragment' );
			} );
		}

		// Slide numbers
		var slideNumberDisplay = 'none';
		if( config.slideNumber && !isPrintingPDF() ) {
			if( config.showSlideNumber === 'all' ) {
				slideNumberDisplay = 'block';
			}
			else if( config.showSlideNumber === 'speaker' && isSpeakerNotes() ) {
				slideNumberDisplay = 'block';
			}
		}

		dom.slideNumber.style.display = slideNumberDisplay;

		sync();

	}

	/**
	 * Binds all event listeners.
	 */
	function addEventListeners() {

		eventsAreBound = true;

		window.addEventListener( 'hashchange', onWindowHashChange, false );
		window.addEventListener( 'resize', onWindowResize, false );

		if( config.touch ) {
			dom.wrapper.addEventListener( 'touchstart', onTouchStart, false );
			dom.wrapper.addEventListener( 'touchmove', onTouchMove, false );
			dom.wrapper.addEventListener( 'touchend', onTouchEnd, false );

			// Support pointer-style touch interaction as well
			if( window.navigator.pointerEnabled ) {
				// IE 11 uses un-prefixed version of pointer events
				dom.wrapper.addEventListener( 'pointerdown', onPointerDown, false );
				dom.wrapper.addEventListener( 'pointermove', onPointerMove, false );
				dom.wrapper.addEventListener( 'pointerup', onPointerUp, false );
			}
			else if( window.navigator.msPointerEnabled ) {
				// IE 10 uses prefixed version of pointer events
				dom.wrapper.addEventListener( 'MSPointerDown', onPointerDown, false );
				dom.wrapper.addEventListener( 'MSPointerMove', onPointerMove, false );
				dom.wrapper.addEventListener( 'MSPointerUp', onPointerUp, false );
			}
		}

		if( config.keyboard ) {
			document.addEventListener( 'keydown', onDocumentKeyDown, false );
			document.addEventListener( 'keypress', onDocumentKeyPress, false );
		}

		if( config.progress && dom.progress ) {
			dom.progress.addEventListener( 'click', onProgressClicked, false );
		}

		if( config.focusBodyOnPageVisibilityChange ) {
			var visibilityChange;

			if( 'hidden' in document ) {
				visibilityChange = 'visibilitychange';
			}
			else if( 'msHidden' in document ) {
				visibilityChange = 'msvisibilitychange';
			}
			else if( 'webkitHidden' in document ) {
				visibilityChange = 'webkitvisibilitychange';
			}

			if( visibilityChange ) {
				document.addEventListener( visibilityChange, onPageVisibilityChange, false );
			}
		}

		// Listen to both touch and click events, in case the device
		// supports both
		var pointerEvents = [ 'touchstart', 'click' ];

		// Only support touch for Android, fixes double navigations in
		// stock browser
		if( UA.match( /android/gi ) ) {
			pointerEvents = [ 'touchstart' ];
		}

		pointerEvents.forEach( function( eventName ) {
			dom.controlsLeft.forEach( function( el ) { el.addEventListener( eventName, onNavigateLeftClicked, false ); } );
			dom.controlsRight.forEach( function( el ) { el.addEventListener( eventName, onNavigateRightClicked, false ); } );
			dom.controlsUp.forEach( function( el ) { el.addEventListener( eventName, onNavigateUpClicked, false ); } );
			dom.controlsDown.forEach( function( el ) { el.addEventListener( eventName, onNavigateDownClicked, false ); } );
			dom.controlsPrev.forEach( function( el ) { el.addEventListener( eventName, onNavigatePrevClicked, false ); } );
			dom.controlsNext.forEach( function( el ) { el.addEventListener( eventName, onNavigateNextClicked, false ); } );
		} );

	}

	/**
	 * Unbinds all event listeners.
	 */
	function removeEventListeners() {

		eventsAreBound = false;

		document.removeEventListener( 'keydown', onDocumentKeyDown, false );
		document.removeEventListener( 'keypress', onDocumentKeyPress, false );
		window.removeEventListener( 'hashchange', onWindowHashChange, false );
		window.removeEventListener( 'resize', onWindowResize, false );

		dom.wrapper.removeEventListener( 'touchstart', onTouchStart, false );
		dom.wrapper.removeEventListener( 'touchmove', onTouchMove, false );
		dom.wrapper.removeEventListener( 'touchend', onTouchEnd, false );

		// IE11
		if( window.navigator.pointerEnabled ) {
			dom.wrapper.removeEventListener( 'pointerdown', onPointerDown, false );
			dom.wrapper.removeEventListener( 'pointermove', onPointerMove, false );
			dom.wrapper.removeEventListener( 'pointerup', onPointerUp, false );
		}
		// IE10
		else if( window.navigator.msPointerEnabled ) {
			dom.wrapper.removeEventListener( 'MSPointerDown', onPointerDown, false );
			dom.wrapper.removeEventListener( 'MSPointerMove', onPointerMove, false );
			dom.wrapper.removeEventListener( 'MSPointerUp', onPointerUp, false );
		}

		if ( config.progress && dom.progress ) {
			dom.progress.removeEventListener( 'click', onProgressClicked, false );
		}

		[ 'touchstart', 'click' ].forEach( function( eventName ) {
			dom.controlsLeft.forEach( function( el ) { el.removeEventListener( eventName, onNavigateLeftClicked, false ); } );
			dom.controlsRight.forEach( function( el ) { el.removeEventListener( eventName, onNavigateRightClicked, false ); } );
			dom.controlsUp.forEach( function( el ) { el.removeEventListener( eventName, onNavigateUpClicked, false ); } );
			dom.controlsDown.forEach( function( el ) { el.removeEventListener( eventName, onNavigateDownClicked, false ); } );
			dom.controlsPrev.forEach( function( el ) { el.removeEventListener( eventName, onNavigatePrevClicked, false ); } );
			dom.controlsNext.forEach( function( el ) { el.removeEventListener( eventName, onNavigateNextClicked, false ); } );
		} );

	}

	/**
	 * Extend object a with the properties of object b.
	 * If there's a conflict, object b takes precedence.
	 *
	 * @param {object} a
	 * @param {object} b
	 */
	function extend( a, b ) {

		for( var i in b ) {
			a[ i ] = b[ i ];
		}

		return a;

	}

	/**
	 * Converts the target object to an array.
	 *
	 * @param {object} o
	 * @return {object[]}
	 */
	function toArray( o ) {

		return Array.prototype.slice.call( o );

	}

	/**
	 * Utility for deserializing a value.
	 *
	 * @param {*} value
	 * @return {*}
	 */
	function deserialize( value ) {

		if( typeof value === 'string' ) {
			if( value === 'null' ) return null;
			else if( value === 'true' ) return true;
			else if( value === 'false' ) return false;
			else if( value.match( /^-?[\d\.]+$/ ) ) return parseFloat( value );
		}

		return value;

	}

	/**
	 * Measures the distance in pixels between point a
	 * and point b.
	 *
	 * @param {object} a point with x/y properties
	 * @param {object} b point with x/y properties
	 *
	 * @return {number}
	 */
	function distanceBetween( a, b ) {

		var dx = a.x - b.x,
			dy = a.y - b.y;

		return Math.sqrt( dx*dx + dy*dy );

	}

	/**
	 * Applies a CSS transform to the target element.
	 *
	 * @param {HTMLElement} element
	 * @param {string} transform
	 */
	function transformElement( element, transform ) {

		element.style.WebkitTransform = transform;
		element.style.MozTransform = transform;
		element.style.msTransform = transform;
		element.style.transform = transform;

	}

	/**
	 * Applies CSS transforms to the slides container. The container
	 * is transformed from two separate sources: layout and the overview
	 * mode.
	 *
	 * @param {object} transforms
	 */
	function transformSlides( transforms ) {

		// Pick up new transforms from arguments
		if( typeof transforms.layout === 'string' ) slidesTransform.layout = transforms.layout;
		if( typeof transforms.overview === 'string' ) slidesTransform.overview = transforms.overview;

		// Apply the transforms to the slides container
		if( slidesTransform.layout ) {
			transformElement( dom.slides, slidesTransform.layout + ' ' + slidesTransform.overview );
		}
		else {
			transformElement( dom.slides, slidesTransform.overview );
		}

	}

	/**
	 * Injects the given CSS styles into the DOM.
	 *
	 * @param {string} value
	 */
	function injectStyleSheet( value ) {

		var tag = document.createElement( 'style' );
		tag.type = 'text/css';
		if( tag.styleSheet ) {
			tag.styleSheet.cssText = value;
		}
		else {
			tag.appendChild( document.createTextNode( value ) );
		}
		document.getElementsByTagName( 'head' )[0].appendChild( tag );

	}

	/**
	 * Find the closest parent that matches the given
	 * selector.
	 *
	 * @param {HTMLElement} target The child element
	 * @param {String} selector The CSS selector to match
	 * the parents against
	 *
	 * @return {HTMLElement} The matched parent or null
	 * if no matching parent was found
	 */
	function closestParent( target, selector ) {

		var parent = target.parentNode;

		while( parent ) {

			// There's some overhead doing this each time, we don't
			// want to rewrite the element prototype but should still
			// be enough to feature detect once at startup...
			var matchesMethod = parent.matches || parent.matchesSelector || parent.msMatchesSelector;

			// If we find a match, we're all set
			if( matchesMethod && matchesMethod.call( parent, selector ) ) {
				return parent;
			}

			// Keep searching
			parent = parent.parentNode;

		}

		return null;

	}

	/**
	 * Converts various color input formats to an {r:0,g:0,b:0} object.
	 *
	 * @param {string} color The string representation of a color
	 * @example
	 * colorToRgb('#000');
	 * @example
	 * colorToRgb('#000000');
	 * @example
	 * colorToRgb('rgb(0,0,0)');
	 * @example
	 * colorToRgb('rgba(0,0,0)');
	 *
	 * @return {{r: number, g: number, b: number, [a]: number}|null}
	 */
	function colorToRgb( color ) {

		var hex3 = color.match( /^#([0-9a-f]{3})$/i );
		if( hex3 && hex3[1] ) {
			hex3 = hex3[1];
			return {
				r: parseInt( hex3.charAt( 0 ), 16 ) * 0x11,
				g: parseInt( hex3.charAt( 1 ), 16 ) * 0x11,
				b: parseInt( hex3.charAt( 2 ), 16 ) * 0x11
			};
		}

		var hex6 = color.match( /^#([0-9a-f]{6})$/i );
		if( hex6 && hex6[1] ) {
			hex6 = hex6[1];
			return {
				r: parseInt( hex6.substr( 0, 2 ), 16 ),
				g: parseInt( hex6.substr( 2, 2 ), 16 ),
				b: parseInt( hex6.substr( 4, 2 ), 16 )
			};
		}

		var rgb = color.match( /^rgb\s*\(\s*(\d+)\s*,\s*(\d+)\s*,\s*(\d+)\s*\)$/i );
		if( rgb ) {
			return {
				r: parseInt( rgb[1], 10 ),
				g: parseInt( rgb[2], 10 ),
				b: parseInt( rgb[3], 10 )
			};
		}

		var rgba = color.match( /^rgba\s*\(\s*(\d+)\s*,\s*(\d+)\s*,\s*(\d+)\s*\,\s*([\d]+|[\d]*.[\d]+)\s*\)$/i );
		if( rgba ) {
			return {
				r: parseInt( rgba[1], 10 ),
				g: parseInt( rgba[2], 10 ),
				b: parseInt( rgba[3], 10 ),
				a: parseFloat( rgba[4] )
			};
		}

		return null;

	}

	/**
	 * Calculates brightness on a scale of 0-255.
	 *
	 * @param {string} color See colorToRgb for supported formats.
	 * @see {@link colorToRgb}
	 */
	function colorBrightness( color ) {

		if( typeof color === 'string' ) color = colorToRgb( color );

		if( color ) {
			return ( color.r * 299 + color.g * 587 + color.b * 114 ) / 1000;
		}

		return null;

	}

	/**
	 * Returns the remaining height within the parent of the
	 * target element.
	 *
	 * remaining height = [ configured parent height ] - [ current parent height ]
	 *
	 * @param {HTMLElement} element
	 * @param {number} [height]
	 */
	function getRemainingHeight( element, height ) {

		height = height || 0;

		if( element ) {
			var newHeight, oldHeight = element.style.height;

			// Change the .stretch element height to 0 in order find the height of all
			// the other elements
			element.style.height = '0px';
			newHeight = height - element.parentNode.offsetHeight;

			// Restore the old height, just in case
			element.style.height = oldHeight + 'px';

			return newHeight;
		}

		return height;

	}

	/**
	 * Checks if this instance is being used to print a PDF.
	 */
	function isPrintingPDF() {

		return ( /print-pdf/gi ).test( window.location.search );

	}

	/**
	 * Hides the address bar if we're on a mobile device.
	 */
	function hideAddressBar() {

		if( config.hideAddressBar && isMobileDevice ) {
			// Events that should trigger the address bar to hide
			window.addEventListener( 'load', removeAddressBar, false );
			window.addEventListener( 'orientationchange', removeAddressBar, false );
		}

	}

	/**
	 * Causes the address bar to hide on mobile devices,
	 * more vertical space ftw.
	 */
	function removeAddressBar() {

		setTimeout( function() {
			window.scrollTo( 0, 1 );
		}, 10 );

	}

	/**
	 * Dispatches an event of the specified type from the
	 * reveal DOM element.
	 */
	function dispatchEvent( type, args ) {

		var event = document.createEvent( 'HTMLEvents', 1, 2 );
		event.initEvent( type, true, true );
		extend( event, args );
		dom.wrapper.dispatchEvent( event );

		// If we're in an iframe, post each reveal.js event to the
		// parent window. Used by the notes plugin
		if( config.postMessageEvents && window.parent !== window.self ) {
			window.parent.postMessage( JSON.stringify({ namespace: 'reveal', eventName: type, state: getState() }), '*' );
		}

	}

	/**
	 * Wrap all links in 3D goodness.
	 */
	function enableRollingLinks() {

		if( features.transforms3d && !( 'msPerspective' in document.body.style ) ) {
			var anchors = dom.wrapper.querySelectorAll( SLIDES_SELECTOR + ' a' );

			for( var i = 0, len = anchors.length; i < len; i++ ) {
				var anchor = anchors[i];

				if( anchor.textContent && !anchor.querySelector( '*' ) && ( !anchor.className || !anchor.classList.contains( anchor, 'roll' ) ) ) {
					var span = document.createElement('span');
					span.setAttribute('data-title', anchor.text);
					span.innerHTML = anchor.innerHTML;

					anchor.classList.add( 'roll' );
					anchor.innerHTML = '';
					anchor.appendChild(span);
				}
			}
		}

	}

	/**
	 * Unwrap all 3D links.
	 */
	function disableRollingLinks() {

		var anchors = dom.wrapper.querySelectorAll( SLIDES_SELECTOR + ' a.roll' );

		for( var i = 0, len = anchors.length; i < len; i++ ) {
			var anchor = anchors[i];
			var span = anchor.querySelector( 'span' );

			if( span ) {
				anchor.classList.remove( 'roll' );
				anchor.innerHTML = span.innerHTML;
			}
		}

	}

	/**
	 * Bind preview frame links.
	 *
	 * @param {string} [selector=a] - selector for anchors
	 */
	function enablePreviewLinks( selector ) {

		var anchors = toArray( document.querySelectorAll( selector ? selector : 'a' ) );

		anchors.forEach( function( element ) {
			if( /^(http|www)/gi.test( element.getAttribute( 'href' ) ) ) {
				element.addEventListener( 'click', onPreviewLinkClicked, false );
			}
		} );

	}

	/**
	 * Unbind preview frame links.
	 */
	function disablePreviewLinks( selector ) {

		var anchors = toArray( document.querySelectorAll( selector ? selector : 'a' ) );

		anchors.forEach( function( element ) {
			if( /^(http|www)/gi.test( element.getAttribute( 'href' ) ) ) {
				element.removeEventListener( 'click', onPreviewLinkClicked, false );
			}
		} );

	}

	/**
	 * Opens a preview window for the target URL.
	 *
	 * @param {string} url - url for preview iframe src
	 */
	function showPreview( url ) {

		closeOverlay();

		dom.overlay = document.createElement( 'div' );
		dom.overlay.classList.add( 'overlay' );
		dom.overlay.classList.add( 'overlay-preview' );
		dom.wrapper.appendChild( dom.overlay );

		dom.overlay.innerHTML = [
			'<header>',
				'<a class="close" href="#"><span class="icon"></span></a>',
				'<a class="external" href="'+ url +'" target="_blank"><span class="icon"></span></a>',
			'</header>',
			'<div class="spinner"></div>',
			'<div class="viewport">',
				'<iframe src="'+ url +'"></iframe>',
				'<small class="viewport-inner">',
					'<span class="x-frame-error">Unable to load iframe. This is likely due to the site\'s policy (x-frame-options).</span>',
				'</small>',
			'</div>'
		].join('');

		dom.overlay.querySelector( 'iframe' ).addEventListener( 'load', function( event ) {
			dom.overlay.classList.add( 'loaded' );
		}, false );

		dom.overlay.querySelector( '.close' ).addEventListener( 'click', function( event ) {
			closeOverlay();
			event.preventDefault();
		}, false );

		dom.overlay.querySelector( '.external' ).addEventListener( 'click', function( event ) {
			closeOverlay();
		}, false );

		setTimeout( function() {
			dom.overlay.classList.add( 'visible' );
		}, 1 );

	}

	/**
	 * Open or close help overlay window.
	 *
	 * @param {Boolean} [override] Flag which overrides the
	 * toggle logic and forcibly sets the desired state. True means
	 * help is open, false means it's closed.
	 */
	function toggleHelp( override ){

		if( typeof override === 'boolean' ) {
			override ? showHelp() : closeOverlay();
		}
		else {
			if( dom.overlay ) {
				closeOverlay();
			}
			else {
				showHelp();
			}
		}
	}

	/**
	 * Opens an overlay window with help material.
	 */
	function showHelp() {

		if( config.help ) {

			closeOverlay();

			dom.overlay = document.createElement( 'div' );
			dom.overlay.classList.add( 'overlay' );
			dom.overlay.classList.add( 'overlay-help' );
			dom.wrapper.appendChild( dom.overlay );

			var html = '<p class="title">Keyboard Shortcuts</p><br/>';

			html += '<table><th>KEY</th><th>ACTION</th>';
			for( var key in keyboardShortcuts ) {
				html += '<tr><td>' + key + '</td><td>' + keyboardShortcuts[ key ] + '</td></tr>';
			}

			html += '</table>';

			dom.overlay.innerHTML = [
				'<header>',
					'<a class="close" href="#"><span class="icon"></span></a>',
				'</header>',
				'<div class="viewport">',
					'<div class="viewport-inner">'+ html +'</div>',
				'</div>'
			].join('');

			dom.overlay.querySelector( '.close' ).addEventListener( 'click', function( event ) {
				closeOverlay();
				event.preventDefault();
			}, false );

			setTimeout( function() {
				dom.overlay.classList.add( 'visible' );
			}, 1 );

		}

	}

	/**
	 * Closes any currently open overlay.
	 */
	function closeOverlay() {

		if( dom.overlay ) {
			dom.overlay.parentNode.removeChild( dom.overlay );
			dom.overlay = null;
		}

	}

	/**
	 * Applies JavaScript-controlled layout rules to the
	 * presentation.
	 */
	function layout() {

		if( dom.wrapper && !isPrintingPDF() ) {

			var size = getComputedSlideSize();

			// Layout the contents of the slides
			layoutSlideContents( config.width, config.height );

			dom.slides.style.width = size.width + 'px';
			dom.slides.style.height = size.height + 'px';

			// Determine scale of content to fit within available space
			scale = Math.min( size.presentationWidth / size.width, size.presentationHeight / size.height );

			// Respect max/min scale settings
			scale = Math.max( scale, config.minScale );
			scale = Math.min( scale, config.maxScale );

			// Don't apply any scaling styles if scale is 1
			if( scale === 1 ) {
				dom.slides.style.zoom = '';
				dom.slides.style.left = '';
				dom.slides.style.top = '';
				dom.slides.style.bottom = '';
				dom.slides.style.right = '';
				transformSlides( { layout: '' } );
			}
			else {
				// Prefer zoom for scaling up so that content remains crisp.
				// Don't use zoom to scale down since that can lead to shifts
				// in text layout/line breaks.
				if( scale > 1 && features.zoom ) {
					dom.slides.style.zoom = scale;
					dom.slides.style.left = '';
					dom.slides.style.top = '';
					dom.slides.style.bottom = '';
					dom.slides.style.right = '';
					transformSlides( { layout: '' } );
				}
				// Apply scale transform as a fallback
				else {
					dom.slides.style.zoom = '';
					dom.slides.style.left = '50%';
					dom.slides.style.top = '50%';
					dom.slides.style.bottom = 'auto';
					dom.slides.style.right = 'auto';
					transformSlides( { layout: 'translate(-50%, -50%) scale('+ scale +')' } );
				}
			}

			// Select all slides, vertical and horizontal
			var slides = toArray( dom.wrapper.querySelectorAll( SLIDES_SELECTOR ) );

			for( var i = 0, len = slides.length; i < len; i++ ) {
				var slide = slides[ i ];

				// Don't bother updating invisible slides
				if( slide.style.display === 'none' ) {
					continue;
				}

				if( config.center || slide.classList.contains( 'center' ) ) {
					// Vertical stacks are not centred since their section
					// children will be
					if( slide.classList.contains( 'stack' ) ) {
						slide.style.top = 0;
					}
					else {
						slide.style.top = Math.max( ( size.height - slide.scrollHeight ) / 2, 0 ) + 'px';
					}
				}
				else {
					slide.style.top = '';
				}

			}

			updateProgress();
			updateParallax();

			if( isOverview() ) {
				updateOverview();
			}

		}

	}

	/**
	 * Applies layout logic to the contents of all slides in
	 * the presentation.
	 *
	 * @param {string|number} width
	 * @param {string|number} height
	 */
	function layoutSlideContents( width, height ) {

		// Handle sizing of elements with the 'stretch' class
		toArray( dom.slides.querySelectorAll( 'section > .stretch' ) ).forEach( function( element ) {

			// Determine how much vertical space we can use
			var remainingHeight = getRemainingHeight( element, height );

			// Consider the aspect ratio of media elements
			if( /(img|video)/gi.test( element.nodeName ) ) {
				var nw = element.naturalWidth || element.videoWidth,
					nh = element.naturalHeight || element.videoHeight;

				var es = Math.min( width / nw, remainingHeight / nh );

				element.style.width = ( nw * es ) + 'px';
				element.style.height = ( nh * es ) + 'px';

			}
			else {
				element.style.width = width + 'px';
				element.style.height = remainingHeight + 'px';
			}

		} );

	}

	/**
	 * Calculates the computed pixel size of our slides. These
	 * values are based on the width and height configuration
	 * options.
	 *
	 * @param {number} [presentationWidth=dom.wrapper.offsetWidth]
	 * @param {number} [presentationHeight=dom.wrapper.offsetHeight]
	 */
	function getComputedSlideSize( presentationWidth, presentationHeight ) {

		var size = {
			// Slide size
			width: config.width,
			height: config.height,

			// Presentation size
			presentationWidth: presentationWidth || dom.wrapper.offsetWidth,
			presentationHeight: presentationHeight || dom.wrapper.offsetHeight
		};

		// Reduce available space by margin
		size.presentationWidth -= ( size.presentationWidth * config.margin );
		size.presentationHeight -= ( size.presentationHeight * config.margin );

		// Slide width may be a percentage of available width
		if( typeof size.width === 'string' && /%$/.test( size.width ) ) {
			size.width = parseInt( size.width, 10 ) / 100 * size.presentationWidth;
		}

		// Slide height may be a percentage of available height
		if( typeof size.height === 'string' && /%$/.test( size.height ) ) {
			size.height = parseInt( size.height, 10 ) / 100 * size.presentationHeight;
		}

		return size;

	}

	/**
	 * Stores the vertical index of a stack so that the same
	 * vertical slide can be selected when navigating to and
	 * from the stack.
	 *
	 * @param {HTMLElement} stack The vertical stack element
	 * @param {string|number} [v=0] Index to memorize
	 */
	function setPreviousVerticalIndex( stack, v ) {

		if( typeof stack === 'object' && typeof stack.setAttribute === 'function' ) {
			stack.setAttribute( 'data-previous-indexv', v || 0 );
		}

	}

	/**
	 * Retrieves the vertical index which was stored using
	 * #setPreviousVerticalIndex() or 0 if no previous index
	 * exists.
	 *
	 * @param {HTMLElement} stack The vertical stack element
	 */
	function getPreviousVerticalIndex( stack ) {

		if( typeof stack === 'object' && typeof stack.setAttribute === 'function' && stack.classList.contains( 'stack' ) ) {
			// Prefer manually defined start-indexv
			var attributeName = stack.hasAttribute( 'data-start-indexv' ) ? 'data-start-indexv' : 'data-previous-indexv';

			return parseInt( stack.getAttribute( attributeName ) || 0, 10 );
		}

		return 0;

	}

	/**
	 * Displays the overview of slides (quick nav) by scaling
	 * down and arranging all slide elements.
	 */
	function activateOverview() {

		// Only proceed if enabled in config
		if( config.overview && !isOverview() ) {

			overview = true;

			dom.wrapper.classList.add( 'overview' );
			dom.wrapper.classList.remove( 'overview-deactivating' );

			if( features.overviewTransitions ) {
				setTimeout( function() {
					dom.wrapper.classList.add( 'overview-animated' );
				}, 1 );
			}

			// Don't auto-slide while in overview mode
			cancelAutoSlide();

			// Move the backgrounds element into the slide container to
			// that the same scaling is applied
			dom.slides.appendChild( dom.background );

			// Clicking on an overview slide navigates to it
			toArray( dom.wrapper.querySelectorAll( SLIDES_SELECTOR ) ).forEach( function( slide ) {
				if( !slide.classList.contains( 'stack' ) ) {
					slide.addEventListener( 'click', onOverviewSlideClicked, true );
				}
			} );

			// Calculate slide sizes
			var margin = 70;
			var slideSize = getComputedSlideSize();
			overviewSlideWidth = slideSize.width + margin;
			overviewSlideHeight = slideSize.height + margin;

			// Reverse in RTL mode
			if( config.rtl ) {
				overviewSlideWidth = -overviewSlideWidth;
			}

			updateSlidesVisibility();
			layoutOverview();
			updateOverview();

			layout();

			// Notify observers of the overview showing
			dispatchEvent( 'overviewshown', {
				'indexh': indexh,
				'indexv': indexv,
				'currentSlide': currentSlide
			} );

		}

	}

	/**
	 * Uses CSS transforms to position all slides in a grid for
	 * display inside of the overview mode.
	 */
	function layoutOverview() {

		// Layout slides
		toArray( dom.wrapper.querySelectorAll( HORIZONTAL_SLIDES_SELECTOR ) ).forEach( function( hslide, h ) {
			hslide.setAttribute( 'data-index-h', h );
			transformElement( hslide, 'translate3d(' + ( h * overviewSlideWidth ) + 'px, 0, 0)' );

			if( hslide.classList.contains( 'stack' ) ) {

				toArray( hslide.querySelectorAll( 'section' ) ).forEach( function( vslide, v ) {
					vslide.setAttribute( 'data-index-h', h );
					vslide.setAttribute( 'data-index-v', v );

					transformElement( vslide, 'translate3d(0, ' + ( v * overviewSlideHeight ) + 'px, 0)' );
				} );

			}
		} );

		// Layout slide backgrounds
		toArray( dom.background.childNodes ).forEach( function( hbackground, h ) {
			transformElement( hbackground, 'translate3d(' + ( h * overviewSlideWidth ) + 'px, 0, 0)' );

			toArray( hbackground.querySelectorAll( '.slide-background' ) ).forEach( function( vbackground, v ) {
				transformElement( vbackground, 'translate3d(0, ' + ( v * overviewSlideHeight ) + 'px, 0)' );
			} );
		} );

	}

	/**
	 * Moves the overview viewport to the current slides.
	 * Called each time the current slide changes.
	 */
	function updateOverview() {

		var vmin = Math.min( window.innerWidth, window.innerHeight );
		var scale = Math.max( vmin / 5, 150 ) / vmin;

		transformSlides( {
			overview: [
				'scale('+ scale +')',
				'translateX('+ ( -indexh * overviewSlideWidth ) +'px)',
				'translateY('+ ( -indexv * overviewSlideHeight ) +'px)'
			].join( ' ' )
		} );

	}

	/**
	 * Exits the slide overview and enters the currently
	 * active slide.
	 */
	function deactivateOverview() {

		// Only proceed if enabled in config
		if( config.overview ) {

			overview = false;

			dom.wrapper.classList.remove( 'overview' );
			dom.wrapper.classList.remove( 'overview-animated' );

			// Temporarily add a class so that transitions can do different things
			// depending on whether they are exiting/entering overview, or just
			// moving from slide to slide
			dom.wrapper.classList.add( 'overview-deactivating' );

			setTimeout( function () {
				dom.wrapper.classList.remove( 'overview-deactivating' );
			}, 1 );

			// Move the background element back out
			dom.wrapper.appendChild( dom.background );

			// Clean up changes made to slides
			toArray( dom.wrapper.querySelectorAll( SLIDES_SELECTOR ) ).forEach( function( slide ) {
				transformElement( slide, '' );

				slide.removeEventListener( 'click', onOverviewSlideClicked, true );
			} );

			// Clean up changes made to backgrounds
			toArray( dom.background.querySelectorAll( '.slide-background' ) ).forEach( function( background ) {
				transformElement( background, '' );
			} );

			transformSlides( { overview: '' } );

			slide( indexh, indexv );

			layout();

			cueAutoSlide();

			// Notify observers of the overview hiding
			dispatchEvent( 'overviewhidden', {
				'indexh': indexh,
				'indexv': indexv,
				'currentSlide': currentSlide
			} );

		}
	}

	/**
	 * Toggles the slide overview mode on and off.
	 *
	 * @param {Boolean} [override] Flag which overrides the
	 * toggle logic and forcibly sets the desired state. True means
	 * overview is open, false means it's closed.
	 */
	function toggleOverview( override ) {

		if( typeof override === 'boolean' ) {
			override ? activateOverview() : deactivateOverview();
		}
		else {
			isOverview() ? deactivateOverview() : activateOverview();
		}

	}

	/**
	 * Checks if the overview is currently active.
	 *
	 * @return {Boolean} true if the overview is active,
	 * false otherwise
	 */
	function isOverview() {

		return overview;

	}

	/**
	 * Checks if the current or specified slide is vertical
	 * (nested within another slide).
	 *
	 * @param {HTMLElement} [slide=currentSlide] The slide to check
	 * orientation of
	 * @return {Boolean}
	 */
	function isVerticalSlide( slide ) {

		// Prefer slide argument, otherwise use current slide
		slide = slide ? slide : currentSlide;

		return slide && slide.parentNode && !!slide.parentNode.nodeName.match( /section/i );

	}

	/**
	 * Handling the fullscreen functionality via the fullscreen API
	 *
	 * @see http://fullscreen.spec.whatwg.org/
	 * @see https://developer.mozilla.org/en-US/docs/DOM/Using_fullscreen_mode
	 */
	function enterFullscreen() {

		var element = document.documentElement;

		// Check which implementation is available
		var requestMethod = element.requestFullscreen ||
							element.webkitRequestFullscreen ||
							element.webkitRequestFullScreen ||
							element.mozRequestFullScreen ||
							element.msRequestFullscreen;

		if( requestMethod ) {
			requestMethod.apply( element );
		}

	}

	/**
	 * Enters the paused mode which fades everything on screen to
	 * black.
	 */
	function pause() {

		if( config.pause ) {
			var wasPaused = dom.wrapper.classList.contains( 'paused' );

			cancelAutoSlide();
			dom.wrapper.classList.add( 'paused' );

			if( wasPaused === false ) {
				dispatchEvent( 'paused' );
			}
		}

	}

	/**
	 * Exits from the paused mode.
	 */
	function resume() {

		var wasPaused = dom.wrapper.classList.contains( 'paused' );
		dom.wrapper.classList.remove( 'paused' );

		cueAutoSlide();

		if( wasPaused ) {
			dispatchEvent( 'resumed' );
		}

	}

	/**
	 * Toggles the paused mode on and off.
	 */
	function togglePause( override ) {

		if( typeof override === 'boolean' ) {
			override ? pause() : resume();
		}
		else {
			isPaused() ? resume() : pause();
		}

	}

	/**
	 * Checks if we are currently in the paused mode.
	 *
	 * @return {Boolean}
	 */
	function isPaused() {

		return dom.wrapper.classList.contains( 'paused' );

	}

	/**
	 * Toggles the auto slide mode on and off.
	 *
	 * @param {Boolean} [override] Flag which sets the desired state.
	 * True means autoplay starts, false means it stops.
	 */

	function toggleAutoSlide( override ) {

		if( typeof override === 'boolean' ) {
			override ? resumeAutoSlide() : pauseAutoSlide();
		}

		else {
			autoSlidePaused ? resumeAutoSlide() : pauseAutoSlide();
		}

	}

	/**
	 * Checks if the auto slide mode is currently on.
	 *
	 * @return {Boolean}
	 */
	function isAutoSliding() {

		return !!( autoSlide && !autoSlidePaused );

	}

	/**
	 * Steps from the current point in the presentation to the
	 * slide which matches the specified horizontal and vertical
	 * indices.
	 *
	 * @param {number} [h=indexh] Horizontal index of the target slide
	 * @param {number} [v=indexv] Vertical index of the target slide
	 * @param {number} [f] Index of a fragment within the
	 * target slide to activate
	 * @param {number} [o] Origin for use in multimaster environments
	 */
	function slide( h, v, f, o ) {

		// Remember where we were at before
		previousSlide = currentSlide;

		// Query all horizontal slides in the deck
		var horizontalSlides = dom.wrapper.querySelectorAll( HORIZONTAL_SLIDES_SELECTOR );

		// Abort if there are no slides
		if( horizontalSlides.length === 0 ) return;

		// If no vertical index is specified and the upcoming slide is a
		// stack, resume at its previous vertical index
		if( v === undefined && !isOverview() ) {
			v = getPreviousVerticalIndex( horizontalSlides[ h ] );
		}

		// If we were on a vertical stack, remember what vertical index
		// it was on so we can resume at the same position when returning
		if( previousSlide && previousSlide.parentNode && previousSlide.parentNode.classList.contains( 'stack' ) ) {
			setPreviousVerticalIndex( previousSlide.parentNode, indexv );
		}

		// Remember the state before this slide
		var stateBefore = state.concat();

		// Reset the state array
		state.length = 0;

		var indexhBefore = indexh || 0,
			indexvBefore = indexv || 0;

		// Activate and transition to the new slide
		indexh = updateSlides( HORIZONTAL_SLIDES_SELECTOR, h === undefined ? indexh : h );
		indexv = updateSlides( VERTICAL_SLIDES_SELECTOR, v === undefined ? indexv : v );

		// Update the visibility of slides now that the indices have changed
		updateSlidesVisibility();

		layout();

		// Apply the new state
		stateLoop: for( var i = 0, len = state.length; i < len; i++ ) {
			// Check if this state existed on the previous slide. If it
			// did, we will avoid adding it repeatedly
			for( var j = 0; j < stateBefore.length; j++ ) {
				if( stateBefore[j] === state[i] ) {
					stateBefore.splice( j, 1 );
					continue stateLoop;
				}
			}

			document.documentElement.classList.add( state[i] );

			// Dispatch custom event matching the state's name
			dispatchEvent( state[i] );
		}

		// Clean up the remains of the previous state
		while( stateBefore.length ) {
			document.documentElement.classList.remove( stateBefore.pop() );
		}

		// Update the overview if it's currently active
		if( isOverview() ) {
			updateOverview();
		}

		// Find the current horizontal slide and any possible vertical slides
		// within it
		var currentHorizontalSlide = horizontalSlides[ indexh ],
			currentVerticalSlides = currentHorizontalSlide.querySelectorAll( 'section' );

		// Store references to the previous and current slides
		currentSlide = currentVerticalSlides[ indexv ] || currentHorizontalSlide;

		// Show fragment, if specified
		if( typeof f !== 'undefined' ) {
			navigateFragment( f );
		}

		// Dispatch an event if the slide changed
		var slideChanged = ( indexh !== indexhBefore || indexv !== indexvBefore );
		if( slideChanged ) {
			dispatchEvent( 'slidechanged', {
				'indexh': indexh,
				'indexv': indexv,
				'previousSlide': previousSlide,
				'currentSlide': currentSlide,
				'origin': o
			} );
		}
		else {
			// Ensure that the previous slide is never the same as the current
			previousSlide = null;
		}

		// Solves an edge case where the previous slide maintains the
		// 'present' class when navigating between adjacent vertical
		// stacks
		if( previousSlide ) {
			previousSlide.classList.remove( 'present' );
			previousSlide.setAttribute( 'aria-hidden', 'true' );

			// Reset all slides upon navigate to home
			// Issue: #285
			if ( dom.wrapper.querySelector( HOME_SLIDE_SELECTOR ).classList.contains( 'present' ) ) {
				// Launch async task
				setTimeout( function () {
					var slides = toArray( dom.wrapper.querySelectorAll( HORIZONTAL_SLIDES_SELECTOR + '.stack') ), i;
					for( i in slides ) {
						if( slides[i] ) {
							// Reset stack
							setPreviousVerticalIndex( slides[i], 0 );
						}
					}
				}, 0 );
			}
		}

		// Handle embedded content
		if( slideChanged || !previousSlide ) {
			stopEmbeddedContent( previousSlide );
			startEmbeddedContent( currentSlide );
		}

		// Announce the current slide contents, for screen readers
		dom.statusDiv.textContent = getStatusText( currentSlide );

		updateControls();
		updateProgress();
		updateBackground();
		updateParallax();
		updateSlideNumber();
		updateNotes();

		// Update the URL hash
		writeURL();

		cueAutoSlide();

	}

	/**
	 * Syncs the presentation with the current DOM. Useful
	 * when new slides or control elements are added or when
	 * the configuration has changed.
	 */
	function sync() {

		// Subscribe to input
		removeEventListeners();
		addEventListeners();

		// Force a layout to make sure the current config is accounted for
		layout();

		// Reflect the current autoSlide value
		autoSlide = config.autoSlide;

		// Start auto-sliding if it's enabled
		cueAutoSlide();

		// Re-create the slide backgrounds
		createBackgrounds();

		// Write the current hash to the URL
		writeURL();

		sortAllFragments();

		updateControls();
		updateProgress();
		updateSlideNumber();
		updateSlidesVisibility();
		updateBackground( true );
		updateNotesVisibility();
		updateNotes();

		formatEmbeddedContent();

		// Start or stop embedded content depending on global config
		if( config.autoPlayMedia === false ) {
			stopEmbeddedContent( currentSlide, { unloadIframes: false } );
		}
		else {
			startEmbeddedContent( currentSlide );
		}

		if( isOverview() ) {
			layoutOverview();
		}

	}

	/**
	 * Resets all vertical slides so that only the first
	 * is visible.
	 */
	function resetVerticalSlides() {

		var horizontalSlides = toArray( dom.wrapper.querySelectorAll( HORIZONTAL_SLIDES_SELECTOR ) );
		horizontalSlides.forEach( function( horizontalSlide ) {

			var verticalSlides = toArray( horizontalSlide.querySelectorAll( 'section' ) );
			verticalSlides.forEach( function( verticalSlide, y ) {

				if( y > 0 ) {
					verticalSlide.classList.remove( 'present' );
					verticalSlide.classList.remove( 'past' );
					verticalSlide.classList.add( 'future' );
					verticalSlide.setAttribute( 'aria-hidden', 'true' );
				}

			} );

		} );

	}

	/**
	 * Sorts and formats all of fragments in the
	 * presentation.
	 */
	function sortAllFragments() {

		var horizontalSlides = toArray( dom.wrapper.querySelectorAll( HORIZONTAL_SLIDES_SELECTOR ) );
		horizontalSlides.forEach( function( horizontalSlide ) {

			var verticalSlides = toArray( horizontalSlide.querySelectorAll( 'section' ) );
			verticalSlides.forEach( function( verticalSlide, y ) {

				sortFragments( verticalSlide.querySelectorAll( '.fragment' ) );

			} );

			if( verticalSlides.length === 0 ) sortFragments( horizontalSlide.querySelectorAll( '.fragment' ) );

		} );

	}

	/**
	 * Randomly shuffles all slides in the deck.
	 */
	function shuffle() {

		var slides = toArray( dom.wrapper.querySelectorAll( HORIZONTAL_SLIDES_SELECTOR ) );

		slides.forEach( function( slide ) {

			// Insert this slide next to another random slide. This may
			// cause the slide to insert before itself but that's fine.
			dom.slides.insertBefore( slide, slides[ Math.floor( Math.random() * slides.length ) ] );

		} );

	}

	/**
	 * Updates one dimension of slides by showing the slide
	 * with the specified index.
	 *
	 * @param {string} selector A CSS selector that will fetch
	 * the group of slides we are working with
	 * @param {number} index The index of the slide that should be
	 * shown
	 *
	 * @return {number} The index of the slide that is now shown,
	 * might differ from the passed in index if it was out of
	 * bounds.
	 */
	function updateSlides( selector, index ) {

		// Select all slides and convert the NodeList result to
		// an array
		var slides = toArray( dom.wrapper.querySelectorAll( selector ) ),
			slidesLength = slides.length;

		var printMode = isPrintingPDF();

		if( slidesLength ) {

			// Should the index loop?
			if( config.loop ) {
				index %= slidesLength;

				if( index < 0 ) {
					index = slidesLength + index;
				}
			}

			// Enforce max and minimum index bounds
			index = Math.max( Math.min( index, slidesLength - 1 ), 0 );

			for( var i = 0; i < slidesLength; i++ ) {
				var element = slides[i];

				var reverse = config.rtl && !isVerticalSlide( element );

				element.classList.remove( 'past' );
				element.classList.remove( 'present' );
				element.classList.remove( 'future' );

				// http://www.w3.org/html/wg/drafts/html/master/editing.html#the-hidden-attribute
				element.setAttribute( 'hidden', '' );
				element.setAttribute( 'aria-hidden', 'true' );

				// If this element contains vertical slides
				if( element.querySelector( 'section' ) ) {
					element.classList.add( 'stack' );
				}

				// If we're printing static slides, all slides are "present"
				if( printMode ) {
					element.classList.add( 'present' );
					continue;
				}

				if( i < index ) {
					// Any element previous to index is given the 'past' class
					element.classList.add( reverse ? 'future' : 'past' );

					if( config.fragments ) {
						var pastFragments = toArray( element.querySelectorAll( '.fragment' ) );

						// Show all fragments on prior slides
						while( pastFragments.length ) {
							var pastFragment = pastFragments.pop();
							pastFragment.classList.add( 'visible' );
							pastFragment.classList.remove( 'current-fragment' );
						}
					}
				}
				else if( i > index ) {
					// Any element subsequent to index is given the 'future' class
					element.classList.add( reverse ? 'past' : 'future' );

					if( config.fragments ) {
						var futureFragments = toArray( element.querySelectorAll( '.fragment.visible' ) );

						// No fragments in future slides should be visible ahead of time
						while( futureFragments.length ) {
							var futureFragment = futureFragments.pop();
							futureFragment.classList.remove( 'visible' );
							futureFragment.classList.remove( 'current-fragment' );
						}
					}
				}
			}

			// Mark the current slide as present
			slides[index].classList.add( 'present' );
			slides[index].removeAttribute( 'hidden' );
			slides[index].removeAttribute( 'aria-hidden' );

			// If this slide has a state associated with it, add it
			// onto the current state of the deck
			var slideState = slides[index].getAttribute( 'data-state' );
			if( slideState ) {
				state = state.concat( slideState.split( ' ' ) );
			}

		}
		else {
			// Since there are no slides we can't be anywhere beyond the
			// zeroth index
			index = 0;
		}

		return index;

	}

	/**
	 * Optimization method; hide all slides that are far away
	 * from the present slide.
	 */
	function updateSlidesVisibility() {

		// Select all slides and convert the NodeList result to
		// an array
		var horizontalSlides = toArray( dom.wrapper.querySelectorAll( HORIZONTAL_SLIDES_SELECTOR ) ),
			horizontalSlidesLength = horizontalSlides.length,
			distanceX,
			distanceY;

		if( horizontalSlidesLength && typeof indexh !== 'undefined' ) {

			// The number of steps away from the present slide that will
			// be visible
			var viewDistance = isOverview() ? 10 : config.viewDistance;

			// Limit view distance on weaker devices
			if( isMobileDevice ) {
				viewDistance = isOverview() ? 6 : 2;
			}

			// All slides need to be visible when exporting to PDF
			if( isPrintingPDF() ) {
				viewDistance = Number.MAX_VALUE;
			}

			for( var x = 0; x < horizontalSlidesLength; x++ ) {
				var horizontalSlide = horizontalSlides[x];

				var verticalSlides = toArray( horizontalSlide.querySelectorAll( 'section' ) ),
					verticalSlidesLength = verticalSlides.length;

				// Determine how far away this slide is from the present
				distanceX = Math.abs( ( indexh || 0 ) - x ) || 0;

				// If the presentation is looped, distance should measure
				// 1 between the first and last slides
				if( config.loop ) {
					distanceX = Math.abs( ( ( indexh || 0 ) - x ) % ( horizontalSlidesLength - viewDistance ) ) || 0;
				}

				// Show the horizontal slide if it's within the view distance
				if( distanceX < viewDistance ) {
					loadSlide( horizontalSlide );
				}
				else {
					unloadSlide( horizontalSlide );
				}

				if( verticalSlidesLength ) {

					var oy = getPreviousVerticalIndex( horizontalSlide );

					for( var y = 0; y < verticalSlidesLength; y++ ) {
						var verticalSlide = verticalSlides[y];

						distanceY = x === ( indexh || 0 ) ? Math.abs( ( indexv || 0 ) - y ) : Math.abs( y - oy );

						if( distanceX + distanceY < viewDistance ) {
							loadSlide( verticalSlide );
						}
						else {
							unloadSlide( verticalSlide );
						}
					}

				}
			}

			// Flag if there are ANY vertical slides, anywhere in the deck
			if( dom.wrapper.querySelectorAll( '.slides>section>section' ).length ) {
				dom.wrapper.classList.add( 'has-vertical-slides' );
			}
			else {
				dom.wrapper.classList.remove( 'has-vertical-slides' );
			}

			// Flag if there are ANY horizontal slides, anywhere in the deck
			if( dom.wrapper.querySelectorAll( '.slides>section' ).length > 1 ) {
				dom.wrapper.classList.add( 'has-horizontal-slides' );
			}
			else {
				dom.wrapper.classList.remove( 'has-horizontal-slides' );
			}

		}

	}

	/**
	 * Pick up notes from the current slide and display them
	 * to the viewer.
	 *
	 * @see {@link config.showNotes}
	 */
	function updateNotes() {

		if( config.showNotes && dom.speakerNotes && currentSlide && !isPrintingPDF() ) {

			dom.speakerNotes.innerHTML = getSlideNotes() || '<span class="notes-placeholder">No notes on this slide.</span>';

		}

	}

	/**
	 * Updates the visibility of the speaker notes sidebar that
	 * is used to share annotated slides. The notes sidebar is
	 * only visible if showNotes is true and there are notes on
	 * one or more slides in the deck.
	 */
	function updateNotesVisibility() {

		if( config.showNotes && hasNotes() ) {
			dom.wrapper.classList.add( 'show-notes' );
		}
		else {
			dom.wrapper.classList.remove( 'show-notes' );
		}

	}

	/**
	 * Checks if there are speaker notes for ANY slide in the
	 * presentation.
	 */
	function hasNotes() {

		return dom.slides.querySelectorAll( '[data-notes], aside.notes' ).length > 0;

	}

	/**
	 * Updates the progress bar to reflect the current slide.
	 */
	function updateProgress() {

		// Update progress if enabled
		if( config.progress && dom.progressbar ) {

			dom.progressbar.style.width = getProgress() * dom.wrapper.offsetWidth + 'px';

		}

	}

	/**
	 * Updates the slide number div to reflect the current slide.
	 *
	 * The following slide number formats are available:
	 *  "h.v":	horizontal . vertical slide number (default)
	 *  "h/v":	horizontal / vertical slide number
	 *    "c":	flattened slide number
	 *  "c/t":	flattened slide number / total slides
	 */
	function updateSlideNumber() {

		// Update slide number if enabled
		if( config.slideNumber && dom.slideNumber ) {

			var value = [];
			var format = 'h.v';

			// Check if a custom number format is available
			if( typeof config.slideNumber === 'string' ) {
				format = config.slideNumber;
			}

			switch( format ) {
				case 'c':
					value.push( getSlidePastCount() + 1 );
					break;
				case 'c/t':
					value.push( getSlidePastCount() + 1, '/', getTotalSlides() );
					break;
				case 'h/v':
					value.push( indexh + 1 );
					if( isVerticalSlide() ) value.push( '/', indexv + 1 );
					break;
				default:
					value.push( indexh + 1 );
					if( isVerticalSlide() ) value.push( '.', indexv + 1 );
			}

			dom.slideNumber.innerHTML = formatSlideNumber( value[0], value[1], value[2] );
		}

	}

	/**
	 * Applies HTML formatting to a slide number before it's
	 * written to the DOM.
	 *
	 * @param {number} a Current slide
	 * @param {string} delimiter Character to separate slide numbers
	 * @param {(number|*)} b Total slides
	 * @return {string} HTML string fragment
	 */
	function formatSlideNumber( a, delimiter, b ) {

		if( typeof b === 'number' && !isNaN( b ) ) {
			return  '<span class="slide-number-a">'+ a +'</span>' +
					'<span class="slide-number-delimiter">'+ delimiter +'</span>' +
					'<span class="slide-number-b">'+ b +'</span>';
		}
		else {
			return '<span class="slide-number-a">'+ a +'</span>';
		}

	}

	/**
	 * Updates the state of all control/navigation arrows.
	 */
	function updateControls() {

		var routes = availableRoutes();
		var fragments = availableFragments();

		// Remove the 'enabled' class from all directions
		dom.controlsLeft.concat( dom.controlsRight )
						.concat( dom.controlsUp )
						.concat( dom.controlsDown )
						.concat( dom.controlsPrev )
						.concat( dom.controlsNext ).forEach( function( node ) {
			node.classList.remove( 'enabled' );
			node.classList.remove( 'fragmented' );

			// Set 'disabled' attribute on all directions
			node.setAttribute( 'disabled', 'disabled' );
		} );

		// Add the 'enabled' class to the available routes; remove 'disabled' attribute to enable buttons
		if( routes.left ) dom.controlsLeft.forEach( function( el ) { el.classList.add( 'enabled' ); el.removeAttribute( 'disabled' ); } );
		if( routes.right ) dom.controlsRight.forEach( function( el ) { el.classList.add( 'enabled' ); el.removeAttribute( 'disabled' ); } );
		if( routes.up ) dom.controlsUp.forEach( function( el ) { el.classList.add( 'enabled' ); el.removeAttribute( 'disabled' ); } );
		if( routes.down ) dom.controlsDown.forEach( function( el ) { el.classList.add( 'enabled' ); el.removeAttribute( 'disabled' ); } );

		// Prev/next buttons
		if( routes.left || routes.up ) dom.controlsPrev.forEach( function( el ) { el.classList.add( 'enabled' ); el.removeAttribute( 'disabled' ); } );
		if( routes.right || routes.down ) dom.controlsNext.forEach( function( el ) { el.classList.add( 'enabled' ); el.removeAttribute( 'disabled' ); } );

		// Highlight fragment directions
		if( currentSlide ) {

			// Always apply fragment decorator to prev/next buttons
			if( fragments.prev ) dom.controlsPrev.forEach( function( el ) { el.classList.add( 'fragmented', 'enabled' ); el.removeAttribute( 'disabled' ); } );
			if( fragments.next ) dom.controlsNext.forEach( function( el ) { el.classList.add( 'fragmented', 'enabled' ); el.removeAttribute( 'disabled' ); } );

			// Apply fragment decorators to directional buttons based on
			// what slide axis they are in
			if( isVerticalSlide( currentSlide ) ) {
				if( fragments.prev ) dom.controlsUp.forEach( function( el ) { el.classList.add( 'fragmented', 'enabled' ); el.removeAttribute( 'disabled' ); } );
				if( fragments.next ) dom.controlsDown.forEach( function( el ) { el.classList.add( 'fragmented', 'enabled' ); el.removeAttribute( 'disabled' ); } );
			}
			else {
				if( fragments.prev ) dom.controlsLeft.forEach( function( el ) { el.classList.add( 'fragmented', 'enabled' ); el.removeAttribute( 'disabled' ); } );
				if( fragments.next ) dom.controlsRight.forEach( function( el ) { el.classList.add( 'fragmented', 'enabled' ); el.removeAttribute( 'disabled' ); } );
			}

		}

		if( config.controlsTutorial ) {

			// Highlight control arrows with an animation to ensure
			// that the viewer knows how to navigate
			if( !hasNavigatedDown && routes.down ) {
				dom.controlsDownArrow.classList.add( 'highlight' );
			}
			else {
				dom.controlsDownArrow.classList.remove( 'highlight' );

				if( !hasNavigatedRight && routes.right && indexv === 0 ) {
					dom.controlsRightArrow.classList.add( 'highlight' );
				}
				else {
					dom.controlsRightArrow.classList.remove( 'highlight' );
				}
			}

		}

	}

	/**
	 * Updates the background elements to reflect the current
	 * slide.
	 *
	 * @param {boolean} includeAll If true, the backgrounds of
	 * all vertical slides (not just the present) will be updated.
	 */
	function updateBackground( includeAll ) {

		var currentBackground = null;

		// Reverse past/future classes when in RTL mode
		var horizontalPast = config.rtl ? 'future' : 'past',
			horizontalFuture = config.rtl ? 'past' : 'future';

		// Update the classes of all backgrounds to match the
		// states of their slides (past/present/future)
		toArray( dom.background.childNodes ).forEach( function( backgroundh, h ) {

			backgroundh.classList.remove( 'past' );
			backgroundh.classList.remove( 'present' );
			backgroundh.classList.remove( 'future' );

			if( h < indexh ) {
				backgroundh.classList.add( horizontalPast );
			}
			else if ( h > indexh ) {
				backgroundh.classList.add( horizontalFuture );
			}
			else {
				backgroundh.classList.add( 'present' );

				// Store a reference to the current background element
				currentBackground = backgroundh;
			}

			if( includeAll || h === indexh ) {
				toArray( backgroundh.querySelectorAll( '.slide-background' ) ).forEach( function( backgroundv, v ) {

					backgroundv.classList.remove( 'past' );
					backgroundv.classList.remove( 'present' );
					backgroundv.classList.remove( 'future' );

					if( v < indexv ) {
						backgroundv.classList.add( 'past' );
					}
					else if ( v > indexv ) {
						backgroundv.classList.add( 'future' );
					}
					else {
						backgroundv.classList.add( 'present' );

						// Only if this is the present horizontal and vertical slide
						if( h === indexh ) currentBackground = backgroundv;
					}

				} );
			}

		} );

		// Stop content inside of previous backgrounds
		if( previousBackground ) {

			stopEmbeddedContent( previousBackground );

		}

		// Start content in the current background
		if( currentBackground ) {

			startEmbeddedContent( currentBackground );

			var backgroundImageURL = currentBackground.style.backgroundImage || '';

			// Restart GIFs (doesn't work in Firefox)
			if( /\.gif/i.test( backgroundImageURL ) ) {
				currentBackground.style.backgroundImage = '';
				window.getComputedStyle( currentBackground ).opacity;
				currentBackground.style.backgroundImage = backgroundImageURL;
			}

			// Don't transition between identical backgrounds. This
			// prevents unwanted flicker.
			var previousBackgroundHash = previousBackground ? previousBackground.getAttribute( 'data-background-hash' ) : null;
			var currentBackgroundHash = currentBackground.getAttribute( 'data-background-hash' );
			if( currentBackgroundHash && currentBackgroundHash === previousBackgroundHash && currentBackground !== previousBackground ) {
				dom.background.classList.add( 'no-transition' );
			}

			previousBackground = currentBackground;

		}

		// If there's a background brightness flag for this slide,
		// bubble it to the .reveal container
		if( currentSlide ) {
			[ 'has-light-background', 'has-dark-background' ].forEach( function( classToBubble ) {
				if( currentSlide.classList.contains( classToBubble ) ) {
					dom.wrapper.classList.add( classToBubble );
				}
				else {
					dom.wrapper.classList.remove( classToBubble );
				}
			} );
		}

		// Allow the first background to apply without transition
		setTimeout( function() {
			dom.background.classList.remove( 'no-transition' );
		}, 1 );

	}

	/**
	 * Updates the position of the parallax background based
	 * on the current slide index.
	 */
	function updateParallax() {

		if( config.parallaxBackgroundImage ) {

			var horizontalSlides = dom.wrapper.querySelectorAll( HORIZONTAL_SLIDES_SELECTOR ),
				verticalSlides = dom.wrapper.querySelectorAll( VERTICAL_SLIDES_SELECTOR );

			var backgroundSize = dom.background.style.backgroundSize.split( ' ' ),
				backgroundWidth, backgroundHeight;

			if( backgroundSize.length === 1 ) {
				backgroundWidth = backgroundHeight = parseInt( backgroundSize[0], 10 );
			}
			else {
				backgroundWidth = parseInt( backgroundSize[0], 10 );
				backgroundHeight = parseInt( backgroundSize[1], 10 );
			}

			var slideWidth = dom.background.offsetWidth,
				horizontalSlideCount = horizontalSlides.length,
				horizontalOffsetMultiplier,
				horizontalOffset;

			if( typeof config.parallaxBackgroundHorizontal === 'number' ) {
				horizontalOffsetMultiplier = config.parallaxBackgroundHorizontal;
			}
			else {
				horizontalOffsetMultiplier = horizontalSlideCount > 1 ? ( backgroundWidth - slideWidth ) / ( horizontalSlideCount-1 ) : 0;
			}

			horizontalOffset = horizontalOffsetMultiplier * indexh * -1;

			var slideHeight = dom.background.offsetHeight,
				verticalSlideCount = verticalSlides.length,
				verticalOffsetMultiplier,
				verticalOffset;

			if( typeof config.parallaxBackgroundVertical === 'number' ) {
				verticalOffsetMultiplier = config.parallaxBackgroundVertical;
			}
			else {
				verticalOffsetMultiplier = ( backgroundHeight - slideHeight ) / ( verticalSlideCount-1 );
			}

			verticalOffset = verticalSlideCount > 0 ?  verticalOffsetMultiplier * indexv : 0;

			dom.background.style.backgroundPosition = horizontalOffset + 'px ' + -verticalOffset + 'px';

		}

	}

	/**
	 * Called when the given slide is within the configured view
	 * distance. Shows the slide element and loads any content
	 * that is set to load lazily (data-src).
	 *
	 * @param {HTMLElement} slide Slide to show
	 */
	function loadSlide( slide, options ) {

		options = options || {};

		// Show the slide element
		slide.style.display = config.display;

		// Media elements with data-src attributes
		toArray( slide.querySelectorAll( 'img[data-src], video[data-src], audio[data-src], iframe[data-src]' ) ).forEach( function( element ) {
			element.setAttribute( 'src', element.getAttribute( 'data-src' ) );
			element.setAttribute( 'data-lazy-loaded', '' );
			element.removeAttribute( 'data-src' );
		} );

		// Media elements with <source> children
		toArray( slide.querySelectorAll( 'video, audio' ) ).forEach( function( media ) {
			var sources = 0;

			toArray( media.querySelectorAll( 'source[data-src]' ) ).forEach( function( source ) {
				source.setAttribute( 'src', source.getAttribute( 'data-src' ) );
				source.removeAttribute( 'data-src' );
				source.setAttribute( 'data-lazy-loaded', '' );
				sources += 1;
			} );

			// If we rewrote sources for this video/audio element, we need
			// to manually tell it to load from its new origin
			if( sources > 0 ) {
				media.load();
			}
		} );


		// Show the corresponding background element
		var indices = getIndices( slide );
		var background = getSlideBackground( indices.h, indices.v );
		if( background ) {
			background.style.display = 'block';

			// If the background contains media, load it
			if( background.hasAttribute( 'data-loaded' ) === false ) {
				background.setAttribute( 'data-loaded', 'true' );

				var backgroundImage = slide.getAttribute( 'data-background-image' ),
					backgroundVideo = slide.getAttribute( 'data-background-video' ),
					backgroundVideoLoop = slide.hasAttribute( 'data-background-video-loop' ),
					backgroundVideoMuted = slide.hasAttribute( 'data-background-video-muted' ),
					backgroundIframe = slide.getAttribute( 'data-background-iframe' );

				// Images
				if( backgroundImage ) {
					background.style.backgroundImage = 'url('+ backgroundImage +')';
				}
				// Videos
				else if ( backgroundVideo && !isSpeakerNotes() ) {
					var video = document.createElement( 'video' );

					if( backgroundVideoLoop ) {
						video.setAttribute( 'loop', '' );
					}

					if( backgroundVideoMuted ) {
						video.muted = true;
					}

					// Inline video playback works (at least in Mobile Safari) as
					// long as the video is muted and the `playsinline` attribute is
					// present
					if( isMobileDevice ) {
						video.muted = true;
						video.autoplay = true;
						video.setAttribute( 'playsinline', '' );
					}

					// Support comma separated lists of video sources
					backgroundVideo.split( ',' ).forEach( function( source ) {
						video.innerHTML += '<source src="'+ source +'">';
					} );

					background.appendChild( video );
				}
				// Iframes
				else if( backgroundIframe && options.excludeIframes !== true ) {
					var iframe = document.createElement( 'iframe' );
					iframe.setAttribute( 'allowfullscreen', '' );
					iframe.setAttribute( 'mozallowfullscreen', '' );
					iframe.setAttribute( 'webkitallowfullscreen', '' );

					// Only load autoplaying content when the slide is shown to
					// avoid having it play in the background
					if( /autoplay=(1|true|yes)/gi.test( backgroundIframe ) ) {
						iframe.setAttribute( 'data-src', backgroundIframe );
					}
					else {
						iframe.setAttribute( 'src', backgroundIframe );
					}

					iframe.style.width  = '100%';
					iframe.style.height = '100%';
					iframe.style.maxHeight = '100%';
					iframe.style.maxWidth = '100%';

					background.appendChild( iframe );
				}
			}

		}

	}

	/**
	 * Unloads and hides the given slide. This is called when the
	 * slide is moved outside of the configured view distance.
	 *
	 * @param {HTMLElement} slide
	 */
	function unloadSlide( slide ) {

		// Hide the slide element
		slide.style.display = 'none';

		// Hide the corresponding background element
		var indices = getIndices( slide );
		var background = getSlideBackground( indices.h, indices.v );
		if( background ) {
			background.style.display = 'none';
		}

		// Reset lazy-loaded media elements with src attributes
		toArray( slide.querySelectorAll( 'video[data-lazy-loaded][src], audio[data-lazy-loaded][src]' ) ).forEach( function( element ) {
			element.setAttribute( 'data-src', element.getAttribute( 'src' ) );
			element.removeAttribute( 'src' );
		} );

		// Reset lazy-loaded media elements with <source> children
		toArray( slide.querySelectorAll( 'video[data-lazy-loaded] source[src], audio source[src]' ) ).forEach( function( source ) {
			source.setAttribute( 'data-src', source.getAttribute( 'src' ) );
			source.removeAttribute( 'src' );
		} );

	}

	/**
	 * Determine what available routes there are for navigation.
	 *
	 * @return {{left: boolean, right: boolean, up: boolean, down: boolean}}
	 */
	function availableRoutes() {

		var horizontalSlides = dom.wrapper.querySelectorAll( HORIZONTAL_SLIDES_SELECTOR ),
			verticalSlides = dom.wrapper.querySelectorAll( VERTICAL_SLIDES_SELECTOR );

		var routes = {
			left: indexh > 0 || config.loop,
			right: indexh < horizontalSlides.length - 1 || config.loop,
			up: indexv > 0,
			down: indexv < verticalSlides.length - 1
		};

		// reverse horizontal controls for rtl
		if( config.rtl ) {
			var left = routes.left;
			routes.left = routes.right;
			routes.right = left;
		}

		return routes;

	}

	/**
	 * Returns an object describing the available fragment
	 * directions.
	 *
	 * @return {{prev: boolean, next: boolean}}
	 */
	function availableFragments() {

		if( currentSlide && config.fragments ) {
			var fragments = currentSlide.querySelectorAll( '.fragment' );
			var hiddenFragments = currentSlide.querySelectorAll( '.fragment:not(.visible)' );

			return {
				prev: fragments.length - hiddenFragments.length > 0,
				next: !!hiddenFragments.length
			};
		}
		else {
			return { prev: false, next: false };
		}

	}

	/**
	 * Enforces origin-specific format rules for embedded media.
	 */
	function formatEmbeddedContent() {

		var _appendParamToIframeSource = function( sourceAttribute, sourceURL, param ) {
			toArray( dom.slides.querySelectorAll( 'iframe['+ sourceAttribute +'*="'+ sourceURL +'"]' ) ).forEach( function( el ) {
				var src = el.getAttribute( sourceAttribute );
				if( src && src.indexOf( param ) === -1 ) {
					el.setAttribute( sourceAttribute, src + ( !/\?/.test( src ) ? '?' : '&' ) + param );
				}
			});
		};

		// YouTube frames must include "?enablejsapi=1"
		_appendParamToIframeSource( 'src', 'youtube.com/embed/', 'enablejsapi=1' );
		_appendParamToIframeSource( 'data-src', 'youtube.com/embed/', 'enablejsapi=1' );

		// Vimeo frames must include "?api=1"
		_appendParamToIframeSource( 'src', 'player.vimeo.com/', 'api=1' );
		_appendParamToIframeSource( 'data-src', 'player.vimeo.com/', 'api=1' );

		// Always show media controls on mobile devices
		if( isMobileDevice ) {
			toArray( dom.slides.querySelectorAll( 'video, audio' ) ).forEach( function( el ) {
				el.controls = true;
			} );
		}

	}

	/**
	 * Start playback of any embedded content inside of
	 * the given element.
	 *
	 * @param {HTMLElement} element
	 */
	function startEmbeddedContent( element ) {

		if( element && !isSpeakerNotes() ) {

			// Restart GIFs
			toArray( element.querySelectorAll( 'img[src$=".gif"]' ) ).forEach( function( el ) {
				// Setting the same unchanged source like this was confirmed
				// to work in Chrome, FF & Safari
				el.setAttribute( 'src', el.getAttribute( 'src' ) );
			} );

			// HTML5 media elements
			toArray( element.querySelectorAll( 'video, audio' ) ).forEach( function( el ) {
				if( closestParent( el, '.fragment' ) && !closestParent( el, '.fragment.visible' ) ) {
					return;
				}

				// Prefer an explicit global autoplay setting
				var autoplay = config.autoPlayMedia;

				// If no global setting is available, fall back on the element's
				// own autoplay setting
				if( typeof autoplay !== 'boolean' ) {
					autoplay = el.hasAttribute( 'data-autoplay' ) || !!closestParent( el, '.slide-background' );
				}

				if( autoplay && typeof el.play === 'function' ) {

					if( el.readyState > 1 ) {
						startEmbeddedMedia( { target: el } );
					}
					else {
						el.removeEventListener( 'loadeddata', startEmbeddedMedia ); // remove first to avoid dupes
						el.addEventListener( 'loadeddata', startEmbeddedMedia );
					}

				}
			} );

			// Normal iframes
			toArray( element.querySelectorAll( 'iframe[src]' ) ).forEach( function( el ) {
				if( closestParent( el, '.fragment' ) && !closestParent( el, '.fragment.visible' ) ) {
					return;
				}

				startEmbeddedIframe( { target: el } );
			} );

			// Lazy loading iframes
			toArray( element.querySelectorAll( 'iframe[data-src]' ) ).forEach( function( el ) {
				if( closestParent( el, '.fragment' ) && !closestParent( el, '.fragment.visible' ) ) {
					return;
				}

				if( el.getAttribute( 'src' ) !== el.getAttribute( 'data-src' ) ) {
					el.removeEventListener( 'load', startEmbeddedIframe ); // remove first to avoid dupes
					el.addEventListener( 'load', startEmbeddedIframe );
					el.setAttribute( 'src', el.getAttribute( 'data-src' ) );
				}
			} );

		}

	}

	/**
	 * Starts playing an embedded video/audio element after
	 * it has finished loading.
	 *
	 * @param {object} event
	 */
	function startEmbeddedMedia( event ) {

		var isAttachedToDOM = !!closestParent( event.target, 'html' ),
			isVisible  		= !!closestParent( event.target, '.present' );

		if( isAttachedToDOM && isVisible ) {
			event.target.currentTime = 0;
			event.target.play();
		}

		event.target.removeEventListener( 'loadeddata', startEmbeddedMedia );

	}

	/**
	 * "Starts" the content of an embedded iframe using the
	 * postMessage API.
	 *
	 * @param {object} event
	 */
	function startEmbeddedIframe( event ) {

		var iframe = event.target;

		if( iframe && iframe.contentWindow ) {

			var isAttachedToDOM = !!closestParent( event.target, 'html' ),
				isVisible  		= !!closestParent( event.target, '.present' );

			if( isAttachedToDOM && isVisible ) {

				// Prefer an explicit global autoplay setting
				var autoplay = config.autoPlayMedia;

				// If no global setting is available, fall back on the element's
				// own autoplay setting
				if( typeof autoplay !== 'boolean' ) {
					autoplay = iframe.hasAttribute( 'data-autoplay' ) || !!closestParent( iframe, '.slide-background' );
				}

				// YouTube postMessage API
				if( /youtube\.com\/embed\//.test( iframe.getAttribute( 'src' ) ) && autoplay ) {
					iframe.contentWindow.postMessage( '{"event":"command","func":"playVideo","args":""}', '*' );
				}
				// Vimeo postMessage API
				else if( /player\.vimeo\.com\//.test( iframe.getAttribute( 'src' ) ) && autoplay ) {
					iframe.contentWindow.postMessage( '{"method":"play"}', '*' );
				}
				// Generic postMessage API
				else {
					iframe.contentWindow.postMessage( 'slide:start', '*' );
				}

			}

		}

	}

	/**
	 * Stop playback of any embedded content inside of
	 * the targeted slide.
	 *
	 * @param {HTMLElement} element
	 */
	function stopEmbeddedContent( element, options ) {

		options = extend( {
			// Defaults
			unloadIframes: true
		}, options || {} );

		if( element && element.parentNode ) {
			// HTML5 media elements
			toArray( element.querySelectorAll( 'video, audio' ) ).forEach( function( el ) {
				if( !el.hasAttribute( 'data-ignore' ) && typeof el.pause === 'function' ) {
					el.setAttribute('data-paused-by-reveal', '');
					el.pause();
				}
			} );

			// Generic postMessage API for non-lazy loaded iframes
			toArray( element.querySelectorAll( 'iframe' ) ).forEach( function( el ) {
				if( el.contentWindow ) el.contentWindow.postMessage( 'slide:stop', '*' );
				el.removeEventListener( 'load', startEmbeddedIframe );
			});

			// YouTube postMessage API
			toArray( element.querySelectorAll( 'iframe[src*="youtube.com/embed/"]' ) ).forEach( function( el ) {
				if( !el.hasAttribute( 'data-ignore' ) && el.contentWindow && typeof el.contentWindow.postMessage === 'function' ) {
					el.contentWindow.postMessage( '{"event":"command","func":"pauseVideo","args":""}', '*' );
				}
			});

			// Vimeo postMessage API
			toArray( element.querySelectorAll( 'iframe[src*="player.vimeo.com/"]' ) ).forEach( function( el ) {
				if( !el.hasAttribute( 'data-ignore' ) && el.contentWindow && typeof el.contentWindow.postMessage === 'function' ) {
					el.contentWindow.postMessage( '{"method":"pause"}', '*' );
				}
			});

			if( options.unloadIframes === true ) {
				// Unload lazy-loaded iframes
				toArray( element.querySelectorAll( 'iframe[data-src]' ) ).forEach( function( el ) {
					// Only removing the src doesn't actually unload the frame
					// in all browsers (Firefox) so we set it to blank first
					el.setAttribute( 'src', 'about:blank' );
					el.removeAttribute( 'src' );
				} );
			}
		}

	}

	/**
	 * Returns the number of past slides. This can be used as a global
	 * flattened index for slides.
	 *
	 * @return {number} Past slide count
	 */
	function getSlidePastCount() {

		var horizontalSlides = toArray( dom.wrapper.querySelectorAll( HORIZONTAL_SLIDES_SELECTOR ) );

		// The number of past slides
		var pastCount = 0;

		// Step through all slides and count the past ones
		mainLoop: for( var i = 0; i < horizontalSlides.length; i++ ) {

			var horizontalSlide = horizontalSlides[i];
			var verticalSlides = toArray( horizontalSlide.querySelectorAll( 'section' ) );

			for( var j = 0; j < verticalSlides.length; j++ ) {

				// Stop as soon as we arrive at the present
				if( verticalSlides[j].classList.contains( 'present' ) ) {
					break mainLoop;
				}

				pastCount++;

			}

			// Stop as soon as we arrive at the present
			if( horizontalSlide.classList.contains( 'present' ) ) {
				break;
			}

			// Don't count the wrapping section for vertical slides
			if( horizontalSlide.classList.contains( 'stack' ) === false ) {
				pastCount++;
			}

		}

		return pastCount;

	}

	/**
	 * Returns a value ranging from 0-1 that represents
	 * how far into the presentation we have navigated.
	 *
	 * @return {number}
	 */
	function getProgress() {

		// The number of past and total slides
		var totalCount = getTotalSlides();
		var pastCount = getSlidePastCount();

		if( currentSlide ) {

			var allFragments = currentSlide.querySelectorAll( '.fragment' );

			// If there are fragments in the current slide those should be
			// accounted for in the progress.
			if( allFragments.length > 0 ) {
				var visibleFragments = currentSlide.querySelectorAll( '.fragment.visible' );

				// This value represents how big a portion of the slide progress
				// that is made up by its fragments (0-1)
				var fragmentWeight = 0.9;

				// Add fragment progress to the past slide count
				pastCount += ( visibleFragments.length / allFragments.length ) * fragmentWeight;
			}

		}

		return pastCount / ( totalCount - 1 );

	}

	/**
	 * Checks if this presentation is running inside of the
	 * speaker notes window.
	 *
	 * @return {boolean}
	 */
	function isSpeakerNotes() {

		return !!window.location.search.match( /receiver/gi );

	}

	/**
	 * Reads the current URL (hash) and navigates accordingly.
	 */
	function readURL() {

		var hash = window.location.hash;

		// Attempt to parse the hash as either an index or name
		var bits = hash.slice( 2 ).split( '/' ),
			name = hash.replace( /#|\//gi, '' );

		// If the first bit is invalid and there is a name we can
		// assume that this is a named link
		if( isNaN( parseInt( bits[0], 10 ) ) && name.length ) {
			var element;

			// Ensure the named link is a valid HTML ID attribute
			if( /^[a-zA-Z][\w:.-]*$/.test( name ) ) {
				// Find the slide with the specified ID
				element = document.getElementById( name );
			}

			if( element ) {
				// Find the position of the named slide and navigate to it
				var indices = Reveal.getIndices( element );
				slide( indices.h, indices.v );
			}
			// If the slide doesn't exist, navigate to the current slide
			else {
				slide( indexh || 0, indexv || 0 );
			}
		}
		else {
			// Read the index components of the hash
			var h = parseInt( bits[0], 10 ) || 0,
				v = parseInt( bits[1], 10 ) || 0;

			if( h !== indexh || v !== indexv ) {
				slide( h, v );
			}
		}

	}

	/**
	 * Updates the page URL (hash) to reflect the current
	 * state.
	 *
	 * @param {number} delay The time in ms to wait before
	 * writing the hash
	 */
	function writeURL( delay ) {

		if( config.history ) {

			// Make sure there's never more than one timeout running
			clearTimeout( writeURLTimeout );

			// If a delay is specified, timeout this call
			if( typeof delay === 'number' ) {
				writeURLTimeout = setTimeout( writeURL, delay );
			}
			else if( currentSlide ) {
				var url = '/';

				// Attempt to create a named link based on the slide's ID
				var id = currentSlide.getAttribute( 'id' );
				if( id ) {
					id = id.replace( /[^a-zA-Z0-9\-\_\:\.]/g, '' );
				}

				// If the current slide has an ID, use that as a named link
				if( typeof id === 'string' && id.length ) {
					url = '/' + id;
				}
				// Otherwise use the /h/v index
				else {
					if( indexh > 0 || indexv > 0 ) url += indexh;
					if( indexv > 0 ) url += '/' + indexv;
				}

				window.location.hash = url;
			}
		}

	}
	/**
	 * Retrieves the h/v location and fragment of the current,
	 * or specified, slide.
	 *
	 * @param {HTMLElement} [slide] If specified, the returned
	 * index will be for this slide rather than the currently
	 * active one
	 *
	 * @return {{h: number, v: number, f: number}}
	 */
	function getIndices( slide ) {

		// By default, return the current indices
		var h = indexh,
			v = indexv,
			f;

		// If a slide is specified, return the indices of that slide
		if( slide ) {
			var isVertical = isVerticalSlide( slide );
			var slideh = isVertical ? slide.parentNode : slide;

			// Select all horizontal slides
			var horizontalSlides = toArray( dom.wrapper.querySelectorAll( HORIZONTAL_SLIDES_SELECTOR ) );

			// Now that we know which the horizontal slide is, get its index
			h = Math.max( horizontalSlides.indexOf( slideh ), 0 );

			// Assume we're not vertical
			v = undefined;

			// If this is a vertical slide, grab the vertical index
			if( isVertical ) {
				v = Math.max( toArray( slide.parentNode.querySelectorAll( 'section' ) ).indexOf( slide ), 0 );
			}
		}

		if( !slide && currentSlide ) {
			var hasFragments = currentSlide.querySelectorAll( '.fragment' ).length > 0;
			if( hasFragments ) {
				var currentFragment = currentSlide.querySelector( '.current-fragment' );
				if( currentFragment && currentFragment.hasAttribute( 'data-fragment-index' ) ) {
					f = parseInt( currentFragment.getAttribute( 'data-fragment-index' ), 10 );
				}
				else {
					f = currentSlide.querySelectorAll( '.fragment.visible' ).length - 1;
				}
			}
		}

		return { h: h, v: v, f: f };

	}

	/**
	 * Retrieves all slides in this presentation.
	 */
	function getSlides() {

		return toArray( dom.wrapper.querySelectorAll( SLIDES_SELECTOR + ':not(.stack)' ));

	}

	/**
	 * Retrieves the total number of slides in this presentation.
	 *
	 * @return {number}
	 */
	function getTotalSlides() {

		return getSlides().length;

	}

	/**
	 * Returns the slide element matching the specified index.
	 *
	 * @return {HTMLElement}
	 */
	function getSlide( x, y ) {

		var horizontalSlide = dom.wrapper.querySelectorAll( HORIZONTAL_SLIDES_SELECTOR )[ x ];
		var verticalSlides = horizontalSlide && horizontalSlide.querySelectorAll( 'section' );

		if( verticalSlides && verticalSlides.length && typeof y === 'number' ) {
			return verticalSlides ? verticalSlides[ y ] : undefined;
		}

		return horizontalSlide;

	}

	/**
	 * Returns the background element for the given slide.
	 * All slides, even the ones with no background properties
	 * defined, have a background element so as long as the
	 * index is valid an element will be returned.
	 *
	 * @param {number} x Horizontal background index
	 * @param {number} y Vertical background index
	 * @return {(HTMLElement[]|*)}
	 */
	function getSlideBackground( x, y ) {

		var slide = getSlide( x, y );
		if( slide ) {
			return slide.slideBackgroundElement;
		}

		return undefined;

	}

	/**
	 * Retrieves the speaker notes from a slide. Notes can be
	 * defined in two ways:
	 * 1. As a data-notes attribute on the slide <section>
	 * 2. As an <aside class="notes"> inside of the slide
	 *
	 * @param {HTMLElement} [slide=currentSlide]
	 * @return {(string|null)}
	 */
	function getSlideNotes( slide ) {

		// Default to the current slide
		slide = slide || currentSlide;

		// Notes can be specified via the data-notes attribute...
		if( slide.hasAttribute( 'data-notes' ) ) {
			return slide.getAttribute( 'data-notes' );
		}

		// ... or using an <aside class="notes"> element
		var notesElement = slide.querySelector( 'aside.notes' );
		if( notesElement ) {
			return notesElement.innerHTML;
		}

		return null;

	}

	/**
	 * Retrieves the current state of the presentation as
	 * an object. This state can then be restored at any
	 * time.
	 *
	 * @return {{indexh: number, indexv: number, indexf: number, paused: boolean, overview: boolean}}
	 */
	function getState() {

		var indices = getIndices();

		return {
			indexh: indices.h,
			indexv: indices.v,
			indexf: indices.f,
			paused: isPaused(),
			overview: isOverview()
		};

	}

	/**
	 * Restores the presentation to the given state.
	 *
	 * @param {object} state As generated by getState()
	 * @see {@link getState} generates the parameter `state`
	 */
	function setState( state ) {

		if( typeof state === 'object' ) {
			slide( deserialize( state.indexh ), deserialize( state.indexv ), deserialize( state.indexf ) );

			var pausedFlag = deserialize( state.paused ),
				overviewFlag = deserialize( state.overview );

			if( typeof pausedFlag === 'boolean' && pausedFlag !== isPaused() ) {
				togglePause( pausedFlag );
			}

			if( typeof overviewFlag === 'boolean' && overviewFlag !== isOverview() ) {
				toggleOverview( overviewFlag );
			}
		}

	}

	/**
	 * Return a sorted fragments list, ordered by an increasing
	 * "data-fragment-index" attribute.
	 *
	 * Fragments will be revealed in the order that they are returned by
	 * this function, so you can use the index attributes to control the
	 * order of fragment appearance.
	 *
	 * To maintain a sensible default fragment order, fragments are presumed
	 * to be passed in document order. This function adds a "fragment-index"
	 * attribute to each node if such an attribute is not already present,
	 * and sets that attribute to an integer value which is the position of
	 * the fragment within the fragments list.
	 *
	 * @param {object[]|*} fragments
	 * @return {object[]} sorted Sorted array of fragments
	 */
	function sortFragments( fragments ) {

		fragments = toArray( fragments );

		var ordered = [],
			unordered = [],
			sorted = [];

		// Group ordered and unordered elements
		fragments.forEach( function( fragment, i ) {
			if( fragment.hasAttribute( 'data-fragment-index' ) ) {
				var index = parseInt( fragment.getAttribute( 'data-fragment-index' ), 10 );

				if( !ordered[index] ) {
					ordered[index] = [];
				}

				ordered[index].push( fragment );
			}
			else {
				unordered.push( [ fragment ] );
			}
		} );

		// Append fragments without explicit indices in their
		// DOM order
		ordered = ordered.concat( unordered );

		// Manually count the index up per group to ensure there
		// are no gaps
		var index = 0;

		// Push all fragments in their sorted order to an array,
		// this flattens the groups
		ordered.forEach( function( group ) {
			group.forEach( function( fragment ) {
				sorted.push( fragment );
				fragment.setAttribute( 'data-fragment-index', index );
			} );

			index ++;
		} );

		return sorted;

	}

	/**
	 * Navigate to the specified slide fragment.
	 *
	 * @param {?number} index The index of the fragment that
	 * should be shown, -1 means all are invisible
	 * @param {number} offset Integer offset to apply to the
	 * fragment index
	 *
	 * @return {boolean} true if a change was made in any
	 * fragments visibility as part of this call
	 */
	function navigateFragment( index, offset ) {

		if( currentSlide && config.fragments ) {

			var fragments = sortFragments( currentSlide.querySelectorAll( '.fragment' ) );
			if( fragments.length ) {

				// If no index is specified, find the current
				if( typeof index !== 'number' ) {
					var lastVisibleFragment = sortFragments( currentSlide.querySelectorAll( '.fragment.visible' ) ).pop();

					if( lastVisibleFragment ) {
						index = parseInt( lastVisibleFragment.getAttribute( 'data-fragment-index' ) || 0, 10 );
					}
					else {
						index = -1;
					}
				}

				// If an offset is specified, apply it to the index
				if( typeof offset === 'number' ) {
					index += offset;
				}

				var fragmentsShown = [],
					fragmentsHidden = [];

				toArray( fragments ).forEach( function( element, i ) {

					if( element.hasAttribute( 'data-fragment-index' ) ) {
						i = parseInt( element.getAttribute( 'data-fragment-index' ), 10 );
					}

					// Visible fragments
					if( i <= index ) {
						if( !element.classList.contains( 'visible' ) ) fragmentsShown.push( element );
						element.classList.add( 'visible' );
						element.classList.remove( 'current-fragment' );

						// Announce the fragments one by one to the Screen Reader
						dom.statusDiv.textContent = getStatusText( element );

						if( i === index ) {
							element.classList.add( 'current-fragment' );
							startEmbeddedContent( element );
						}
					}
					// Hidden fragments
					else {
						if( element.classList.contains( 'visible' ) ) fragmentsHidden.push( element );
						element.classList.remove( 'visible' );
						element.classList.remove( 'current-fragment' );
					}

				} );

				if( fragmentsHidden.length ) {
					dispatchEvent( 'fragmenthidden', { fragment: fragmentsHidden[0], fragments: fragmentsHidden } );
				}

				if( fragmentsShown.length ) {
					dispatchEvent( 'fragmentshown', { fragment: fragmentsShown[0], fragments: fragmentsShown } );
				}

				updateControls();
				updateProgress();

				return !!( fragmentsShown.length || fragmentsHidden.length );

			}

		}

		return false;

	}

	/**
	 * Navigate to the next slide fragment.
	 *
	 * @return {boolean} true if there was a next fragment,
	 * false otherwise
	 */
	function nextFragment() {

		return navigateFragment( null, 1 );

	}

	/**
	 * Navigate to the previous slide fragment.
	 *
	 * @return {boolean} true if there was a previous fragment,
	 * false otherwise
	 */
	function previousFragment() {

		return navigateFragment( null, -1 );

	}

	/**
	 * Cues a new automated slide if enabled in the config.
	 */
	function cueAutoSlide() {

		cancelAutoSlide();

		if( currentSlide && config.autoSlide !== false ) {

			var fragment = currentSlide.querySelector( '.current-fragment' );

			// When the slide first appears there is no "current" fragment so
			// we look for a data-autoslide timing on the first fragment
			if( !fragment ) fragment = currentSlide.querySelector( '.fragment' );

			var fragmentAutoSlide = fragment ? fragment.getAttribute( 'data-autoslide' ) : null;
			var parentAutoSlide = currentSlide.parentNode ? currentSlide.parentNode.getAttribute( 'data-autoslide' ) : null;
			var slideAutoSlide = currentSlide.getAttribute( 'data-autoslide' );

			// Pick value in the following priority order:
			// 1. Current fragment's data-autoslide
			// 2. Current slide's data-autoslide
			// 3. Parent slide's data-autoslide
			// 4. Global autoSlide setting
			if( fragmentAutoSlide ) {
				autoSlide = parseInt( fragmentAutoSlide, 10 );
			}
			else if( slideAutoSlide ) {
				autoSlide = parseInt( slideAutoSlide, 10 );
			}
			else if( parentAutoSlide ) {
				autoSlide = parseInt( parentAutoSlide, 10 );
			}
			else {
				autoSlide = config.autoSlide;
			}

			// If there are media elements with data-autoplay,
			// automatically set the autoSlide duration to the
			// length of that media. Not applicable if the slide
			// is divided up into fragments.
			// playbackRate is accounted for in the duration.
			if( currentSlide.querySelectorAll( '.fragment' ).length === 0 ) {
				toArray( currentSlide.querySelectorAll( 'video, audio' ) ).forEach( function( el ) {
					if( el.hasAttribute( 'data-autoplay' ) ) {
						if( autoSlide && (el.duration * 1000 / el.playbackRate ) > autoSlide ) {
							autoSlide = ( el.duration * 1000 / el.playbackRate ) + 1000;
						}
					}
				} );
			}

			// Cue the next auto-slide if:
			// - There is an autoSlide value
			// - Auto-sliding isn't paused by the user
			// - The presentation isn't paused
			// - The overview isn't active
			// - The presentation isn't over
			if( autoSlide && !autoSlidePaused && !isPaused() && !isOverview() && ( !Reveal.isLastSlide() || availableFragments().next || config.loop === true ) ) {
				autoSlideTimeout = setTimeout( function() {
					typeof config.autoSlideMethod === 'function' ? config.autoSlideMethod() : navigateNext();
					cueAutoSlide();
				}, autoSlide );
				autoSlideStartTime = Date.now();
			}

			if( autoSlidePlayer ) {
				autoSlidePlayer.setPlaying( autoSlideTimeout !== -1 );
			}

		}

	}

	/**
	 * Cancels any ongoing request to auto-slide.
	 */
	function cancelAutoSlide() {

		clearTimeout( autoSlideTimeout );
		autoSlideTimeout = -1;

	}

	function pauseAutoSlide() {

		if( autoSlide && !autoSlidePaused ) {
			autoSlidePaused = true;
			dispatchEvent( 'autoslidepaused' );
			clearTimeout( autoSlideTimeout );

			if( autoSlidePlayer ) {
				autoSlidePlayer.setPlaying( false );
			}
		}

	}

	function resumeAutoSlide() {

		if( autoSlide && autoSlidePaused ) {
			autoSlidePaused = false;
			dispatchEvent( 'autoslideresumed' );
			cueAutoSlide();
		}

	}

	function navigateLeft() {

		// Reverse for RTL
		if( config.rtl ) {
			if( ( isOverview() || nextFragment() === false ) && availableRoutes().left ) {
				slide( indexh + 1 );
			}
		}
		// Normal navigation
		else if( ( isOverview() || previousFragment() === false ) && availableRoutes().left ) {
			slide( indexh - 1 );
		}

	}

	function navigateRight() {

		hasNavigatedRight = true;

		// Reverse for RTL
		if( config.rtl ) {
			if( ( isOverview() || previousFragment() === false ) && availableRoutes().right ) {
				slide( indexh - 1 );
			}
		}
		// Normal navigation
		else if( ( isOverview() || nextFragment() === false ) && availableRoutes().right ) {
			slide( indexh + 1 );
		}

	}

	function navigateUp() {

		// Prioritize hiding fragments
		if( ( isOverview() || previousFragment() === false ) && availableRoutes().up ) {
			slide( indexh, indexv - 1 );
		}

	}

	function navigateDown() {

		hasNavigatedDown = true;

		// Prioritize revealing fragments
		if( ( isOverview() || nextFragment() === false ) && availableRoutes().down ) {
			slide( indexh, indexv + 1 );
		}

	}

	/**
	 * Navigates backwards, prioritized in the following order:
	 * 1) Previous fragment
	 * 2) Previous vertical slide
	 * 3) Previous horizontal slide
	 */
	function navigatePrev() {

		// Prioritize revealing fragments
		if( previousFragment() === false ) {
			if( availableRoutes().up ) {
				navigateUp();
			}
			else {
				// Fetch the previous horizontal slide, if there is one
				var previousSlide;

				if( config.rtl ) {
					previousSlide = toArray( dom.wrapper.querySelectorAll( HORIZONTAL_SLIDES_SELECTOR + '.future' ) ).pop();
				}
				else {
					previousSlide = toArray( dom.wrapper.querySelectorAll( HORIZONTAL_SLIDES_SELECTOR + '.past' ) ).pop();
				}

				if( previousSlide ) {
					var v = ( previousSlide.querySelectorAll( 'section' ).length - 1 ) || undefined;
					var h = indexh - 1;
					slide( h, v );
				}
			}
		}

	}

	/**
	 * The reverse of #navigatePrev().
	 */
	function navigateNext() {

		hasNavigatedRight = true;
		hasNavigatedDown = true;

		// Prioritize revealing fragments
		if( nextFragment() === false ) {
			if( availableRoutes().down ) {
				navigateDown();
			}
			else if( config.rtl ) {
				navigateLeft();
			}
			else {
				navigateRight();
			}
		}

	}

	/**
	 * Checks if the target element prevents the triggering of
	 * swipe navigation.
	 */
	function isSwipePrevented( target ) {

		while( target && typeof target.hasAttribute === 'function' ) {
			if( target.hasAttribute( 'data-prevent-swipe' ) ) return true;
			target = target.parentNode;
		}

		return false;

	}


	// --------------------------------------------------------------------//
	// ----------------------------- EVENTS -------------------------------//
	// --------------------------------------------------------------------//

	/**
	 * Called by all event handlers that are based on user
	 * input.
	 *
	 * @param {object} [event]
	 */
	function onUserInput( event ) {

		if( config.autoSlideStoppable ) {
			pauseAutoSlide();
		}

	}

	/**
	 * Handler for the document level 'keypress' event.
	 *
	 * @param {object} event
	 */
	function onDocumentKeyPress( event ) {

		// Check if the pressed key is question mark
		if( event.shiftKey && event.charCode === 63 ) {
			toggleHelp();
		}

	}

	/**
	 * Handler for the document level 'keydown' event.
	 *
	 * @param {object} event
	 */
	function onDocumentKeyDown( event ) {

		// If there's a condition specified and it returns false,
		// ignore this event
		if( typeof config.keyboardCondition === 'function' && config.keyboardCondition() === false ) {
			return true;
		}

		// Remember if auto-sliding was paused so we can toggle it
		var autoSlideWasPaused = autoSlidePaused;

		onUserInput( event );

		// Check if there's a focused element that could be using
		// the keyboard
		var activeElementIsCE = document.activeElement && document.activeElement.contentEditable !== 'inherit';
		var activeElementIsInput = document.activeElement && document.activeElement.tagName && /input|textarea/i.test( document.activeElement.tagName );
		var activeElementIsNotes = document.activeElement && document.activeElement.className && /speaker-notes/i.test( document.activeElement.className);

		// Disregard the event if there's a focused element or a
		// keyboard modifier key is present
		if( activeElementIsCE || activeElementIsInput || activeElementIsNotes || (event.shiftKey && event.keyCode !== 32) || event.altKey || event.ctrlKey || event.metaKey ) return;

		// While paused only allow resume keyboard events; 'b', 'v', '.'
		var resumeKeyCodes = [66,86,190,191];
		var key;

		// Custom key bindings for togglePause should be able to resume
		if( typeof config.keyboard === 'object' ) {
			for( key in config.keyboard ) {
				if( config.keyboard[key] === 'togglePause' ) {
					resumeKeyCodes.push( parseInt( key, 10 ) );
				}
			}
		}

		if( isPaused() && resumeKeyCodes.indexOf( event.keyCode ) === -1 ) {
			return false;
		}

		var triggered = false;

		// 1. User defined key bindings
		if( typeof config.keyboard === 'object' ) {

			for( key in config.keyboard ) {

				// Check if this binding matches the pressed key
				if( parseInt( key, 10 ) === event.keyCode ) {

					var value = config.keyboard[ key ];

					// Callback function
					if( typeof value === 'function' ) {
						value.apply( null, [ event ] );
					}
					// String shortcuts to reveal.js API
					else if( typeof value === 'string' && typeof Reveal[ value ] === 'function' ) {
						Reveal[ value ].call();
					}

					triggered = true;

				}

			}

		}

		// 2. System defined key bindings
		if( triggered === false ) {

			// Assume true and try to prove false
			triggered = true;

			switch( event.keyCode ) {
				// p, page up
				case 80: case 33: navigatePrev(); break;
				// n, page down
				case 78: case 34: navigateNext(); break;
				// h, left
				case 72: case 37: navigateLeft(); break;
				// l, right
				case 76: case 39: navigateRight(); break;
				// k, up
				case 75: case 38: navigateUp(); break;
				// j, down
				case 74: case 40: navigateDown(); break;
				// home
				case 36: slide( 0 ); break;
				// end
				case 35: slide( Number.MAX_VALUE ); break;
				// space
				case 32: isOverview() ? deactivateOverview() : event.shiftKey ? navigatePrev() : navigateNext(); break;
				// return
				case 13: isOverview() ? deactivateOverview() : triggered = false; break;
				// two-spot, semicolon, b, v, period, Logitech presenter tools "black screen" button
				case 58: case 59: case 66: case 86: case 190: case 191: togglePause(); break;
				// f
				case 70: enterFullscreen(); break;
				// a
				case 65: if ( config.autoSlideStoppable ) toggleAutoSlide( autoSlideWasPaused ); break;
				default:
					triggered = false;
			}

		}

		// If the input resulted in a triggered action we should prevent
		// the browsers default behavior
		if( triggered ) {
			event.preventDefault && event.preventDefault();
		}
		// ESC or O key
		else if ( ( event.keyCode === 27 || event.keyCode === 79 ) && features.transforms3d ) {
			if( dom.overlay ) {
				closeOverlay();
			}
			else {
				toggleOverview();
			}

			event.preventDefault && event.preventDefault();
		}

		// If auto-sliding is enabled we need to cue up
		// another timeout
		cueAutoSlide();

	}

	/**
	 * Handler for the 'touchstart' event, enables support for
	 * swipe and pinch gestures.
	 *
	 * @param {object} event
	 */
	function onTouchStart( event ) {

		if( isSwipePrevented( event.target ) ) return true;

		touch.startX = event.touches[0].clientX;
		touch.startY = event.touches[0].clientY;
		touch.startCount = event.touches.length;

		// If there's two touches we need to memorize the distance
		// between those two points to detect pinching
		if( event.touches.length === 2 && config.overview ) {
			touch.startSpan = distanceBetween( {
				x: event.touches[1].clientX,
				y: event.touches[1].clientY
			}, {
				x: touch.startX,
				y: touch.startY
			} );
		}

	}

	/**
	 * Handler for the 'touchmove' event.
	 *
	 * @param {object} event
	 */
	function onTouchMove( event ) {

		if( isSwipePrevented( event.target ) ) return true;

		// Each touch should only trigger one action
		if( !touch.captured ) {
			onUserInput( event );

			var currentX = event.touches[0].clientX;
			var currentY = event.touches[0].clientY;

			// If the touch started with two points and still has
			// two active touches; test for the pinch gesture
			if( event.touches.length === 2 && touch.startCount === 2 && config.overview ) {

				// The current distance in pixels between the two touch points
				var currentSpan = distanceBetween( {
					x: event.touches[1].clientX,
					y: event.touches[1].clientY
				}, {
					x: touch.startX,
					y: touch.startY
				} );

				// If the span is larger than the desire amount we've got
				// ourselves a pinch
				if( Math.abs( touch.startSpan - currentSpan ) > touch.threshold ) {
					touch.captured = true;

					if( currentSpan < touch.startSpan ) {
						activateOverview();
					}
					else {
						deactivateOverview();
					}
				}

				event.preventDefault();

			}
			// There was only one touch point, look for a swipe
			else if( event.touches.length === 1 && touch.startCount !== 2 ) {

				var deltaX = currentX - touch.startX,
					deltaY = currentY - touch.startY;

				if( deltaX > touch.threshold && Math.abs( deltaX ) > Math.abs( deltaY ) ) {
					touch.captured = true;
					navigateLeft();
				}
				else if( deltaX < -touch.threshold && Math.abs( deltaX ) > Math.abs( deltaY ) ) {
					touch.captured = true;
					navigateRight();
				}
				else if( deltaY > touch.threshold ) {
					touch.captured = true;
					navigateUp();
				}
				else if( deltaY < -touch.threshold ) {
					touch.captured = true;
					navigateDown();
				}

				// If we're embedded, only block touch events if they have
				// triggered an action
				if( config.embedded ) {
					if( touch.captured || isVerticalSlide( currentSlide ) ) {
						event.preventDefault();
					}
				}
				// Not embedded? Block them all to avoid needless tossing
				// around of the viewport in iOS
				else {
					event.preventDefault();
				}

			}
		}
		// There's a bug with swiping on some Android devices unless
		// the default action is always prevented
		else if( UA.match( /android/gi ) ) {
			event.preventDefault();
		}

	}

	/**
	 * Handler for the 'touchend' event.
	 *
	 * @param {object} event
	 */
	function onTouchEnd( event ) {

		touch.captured = false;

	}

	/**
	 * Convert pointer down to touch start.
	 *
	 * @param {object} event
	 */
	function onPointerDown( event ) {

		if( event.pointerType === event.MSPOINTER_TYPE_TOUCH || event.pointerType === "touch" ) {
			event.touches = [{ clientX: event.clientX, clientY: event.clientY }];
			onTouchStart( event );
		}

	}

	/**
	 * Convert pointer move to touch move.
	 *
	 * @param {object} event
	 */
	function onPointerMove( event ) {

		if( event.pointerType === event.MSPOINTER_TYPE_TOUCH || event.pointerType === "touch" )  {
			event.touches = [{ clientX: event.clientX, clientY: event.clientY }];
			onTouchMove( event );
		}

	}

	/**
	 * Convert pointer up to touch end.
	 *
	 * @param {object} event
	 */
	function onPointerUp( event ) {

		if( event.pointerType === event.MSPOINTER_TYPE_TOUCH || event.pointerType === "touch" )  {
			event.touches = [{ clientX: event.clientX, clientY: event.clientY }];
			onTouchEnd( event );
		}

	}

	/**
	 * Handles mouse wheel scrolling, throttled to avoid skipping
	 * multiple slides.
	 *
	 * @param {object} event
	 */
	function onDocumentMouseScroll( event ) {

		if( Date.now() - lastMouseWheelStep > 600 ) {

			lastMouseWheelStep = Date.now();

			var delta = event.detail || -event.wheelDelta;
			if( delta > 0 ) {
				navigateNext();
			}
			else if( delta < 0 ) {
				navigatePrev();
			}

		}

	}

	/**
	 * Clicking on the progress bar results in a navigation to the
	 * closest approximate horizontal slide using this equation:
	 *
	 * ( clickX / presentationWidth ) * numberOfSlides
	 *
	 * @param {object} event
	 */
	function onProgressClicked( event ) {

		onUserInput( event );

		event.preventDefault();

		var slidesTotal = toArray( dom.wrapper.querySelectorAll( HORIZONTAL_SLIDES_SELECTOR ) ).length;
		var slideIndex = Math.floor( ( event.clientX / dom.wrapper.offsetWidth ) * slidesTotal );

		if( config.rtl ) {
			slideIndex = slidesTotal - slideIndex;
		}

		slide( slideIndex );

	}

	/**
	 * Event handler for navigation control buttons.
	 */
	function onNavigateLeftClicked( event ) { event.preventDefault(); onUserInput(); navigateLeft(); }
	function onNavigateRightClicked( event ) { event.preventDefault(); onUserInput(); navigateRight(); }
	function onNavigateUpClicked( event ) { event.preventDefault(); onUserInput(); navigateUp(); }
	function onNavigateDownClicked( event ) { event.preventDefault(); onUserInput(); navigateDown(); }
	function onNavigatePrevClicked( event ) { event.preventDefault(); onUserInput(); navigatePrev(); }
	function onNavigateNextClicked( event ) { event.preventDefault(); onUserInput(); navigateNext(); }

	/**
	 * Handler for the window level 'hashchange' event.
	 *
	 * @param {object} [event]
	 */
	function onWindowHashChange( event ) {

		readURL();

	}

	/**
	 * Handler for the window level 'resize' event.
	 *
	 * @param {object} [event]
	 */
	function onWindowResize( event ) {

		layout();

	}

	/**
	 * Handle for the window level 'visibilitychange' event.
	 *
	 * @param {object} [event]
	 */
	function onPageVisibilityChange( event ) {

		var isHidden =  document.webkitHidden ||
						document.msHidden ||
						document.hidden;

		// If, after clicking a link or similar and we're coming back,
		// focus the document.body to ensure we can use keyboard shortcuts
		if( isHidden === false && document.activeElement !== document.body ) {
			// Not all elements support .blur() - SVGs among them.
			if( typeof document.activeElement.blur === 'function' ) {
				document.activeElement.blur();
			}
			document.body.focus();
		}

	}

	/**
	 * Invoked when a slide is and we're in the overview.
	 *
	 * @param {object} event
	 */
	function onOverviewSlideClicked( event ) {

		// TODO There's a bug here where the event listeners are not
		// removed after deactivating the overview.
		if( eventsAreBound && isOverview() ) {
			event.preventDefault();

			var element = event.target;

			while( element && !element.nodeName.match( /section/gi ) ) {
				element = element.parentNode;
			}

			if( element && !element.classList.contains( 'disabled' ) ) {

				deactivateOverview();

				if( element.nodeName.match( /section/gi ) ) {
					var h = parseInt( element.getAttribute( 'data-index-h' ), 10 ),
						v = parseInt( element.getAttribute( 'data-index-v' ), 10 );

					slide( h, v );
				}

			}
		}

	}

	/**
	 * Handles clicks on links that are set to preview in the
	 * iframe overlay.
	 *
	 * @param {object} event
	 */
	function onPreviewLinkClicked( event ) {

		if( event.currentTarget && event.currentTarget.hasAttribute( 'href' ) ) {
			var url = event.currentTarget.getAttribute( 'href' );
			if( url ) {
				showPreview( url );
				event.preventDefault();
			}
		}

	}

	/**
	 * Handles click on the auto-sliding controls element.
	 *
	 * @param {object} [event]
	 */
	function onAutoSlidePlayerClick( event ) {

		// Replay
		if( Reveal.isLastSlide() && config.loop === false ) {
			slide( 0, 0 );
			resumeAutoSlide();
		}
		// Resume
		else if( autoSlidePaused ) {
			resumeAutoSlide();
		}
		// Pause
		else {
			pauseAutoSlide();
		}

	}


	// --------------------------------------------------------------------//
	// ------------------------ PLAYBACK COMPONENT ------------------------//
	// --------------------------------------------------------------------//


	/**
	 * Constructor for the playback component, which displays
	 * play/pause/progress controls.
	 *
	 * @param {HTMLElement} container The component will append
	 * itself to this
	 * @param {function} progressCheck A method which will be
	 * called frequently to get the current progress on a range
	 * of 0-1
	 */
	function Playback( container, progressCheck ) {

		// Cosmetics
		this.diameter = 100;
		this.diameter2 = this.diameter/2;
		this.thickness = 6;

		// Flags if we are currently playing
		this.playing = false;

		// Current progress on a 0-1 range
		this.progress = 0;

		// Used to loop the animation smoothly
		this.progressOffset = 1;

		this.container = container;
		this.progressCheck = progressCheck;

		this.canvas = document.createElement( 'canvas' );
		this.canvas.className = 'playback';
		this.canvas.width = this.diameter;
		this.canvas.height = this.diameter;
		this.canvas.style.width = this.diameter2 + 'px';
		this.canvas.style.height = this.diameter2 + 'px';
		this.context = this.canvas.getContext( '2d' );

		this.container.appendChild( this.canvas );

		this.render();

	}

	/**
	 * @param value
	 */
	Playback.prototype.setPlaying = function( value ) {

		var wasPlaying = this.playing;

		this.playing = value;

		// Start repainting if we weren't already
		if( !wasPlaying && this.playing ) {
			this.animate();
		}
		else {
			this.render();
		}

	};

	Playback.prototype.animate = function() {

		var progressBefore = this.progress;

		this.progress = this.progressCheck();

		// When we loop, offset the progress so that it eases
		// smoothly rather than immediately resetting
		if( progressBefore > 0.8 && this.progress < 0.2 ) {
			this.progressOffset = this.progress;
		}

		this.render();

		if( this.playing ) {
			features.requestAnimationFrameMethod.call( window, this.animate.bind( this ) );
		}

	};

	/**
	 * Renders the current progress and playback state.
	 */
	Playback.prototype.render = function() {

		var progress = this.playing ? this.progress : 0,
			radius = ( this.diameter2 ) - this.thickness,
			x = this.diameter2,
			y = this.diameter2,
			iconSize = 28;

		// Ease towards 1
		this.progressOffset += ( 1 - this.progressOffset ) * 0.1;

		var endAngle = ( - Math.PI / 2 ) + ( progress * ( Math.PI * 2 ) );
		var startAngle = ( - Math.PI / 2 ) + ( this.progressOffset * ( Math.PI * 2 ) );

		this.context.save();
		this.context.clearRect( 0, 0, this.diameter, this.diameter );

		// Solid background color
		this.context.beginPath();
		this.context.arc( x, y, radius + 4, 0, Math.PI * 2, false );
		this.context.fillStyle = 'rgba( 0, 0, 0, 0.4 )';
		this.context.fill();

		// Draw progress track
		this.context.beginPath();
		this.context.arc( x, y, radius, 0, Math.PI * 2, false );
		this.context.lineWidth = this.thickness;
		this.context.strokeStyle = 'rgba( 255, 255, 255, 0.2 )';
		this.context.stroke();

		if( this.playing ) {
			// Draw progress on top of track
			this.context.beginPath();
			this.context.arc( x, y, radius, startAngle, endAngle, false );
			this.context.lineWidth = this.thickness;
			this.context.strokeStyle = '#fff';
			this.context.stroke();
		}

		this.context.translate( x - ( iconSize / 2 ), y - ( iconSize / 2 ) );

		// Draw play/pause icons
		if( this.playing ) {
			this.context.fillStyle = '#fff';
			this.context.fillRect( 0, 0, iconSize / 2 - 4, iconSize );
			this.context.fillRect( iconSize / 2 + 4, 0, iconSize / 2 - 4, iconSize );
		}
		else {
			this.context.beginPath();
			this.context.translate( 4, 0 );
			this.context.moveTo( 0, 0 );
			this.context.lineTo( iconSize - 4, iconSize / 2 );
			this.context.lineTo( 0, iconSize );
			this.context.fillStyle = '#fff';
			this.context.fill();
		}

		this.context.restore();

	};

	Playback.prototype.on = function( type, listener ) {
		this.canvas.addEventListener( type, listener, false );
	};

	Playback.prototype.off = function( type, listener ) {
		this.canvas.removeEventListener( type, listener, false );
	};

	Playback.prototype.destroy = function() {

		this.playing = false;

		if( this.canvas.parentNode ) {
			this.container.removeChild( this.canvas );
		}

	};


	// --------------------------------------------------------------------//
	// ------------------------------- API --------------------------------//
	// --------------------------------------------------------------------//


	Reveal = {
		VERSION: VERSION,

		initialize: initialize,
		configure: configure,
		sync: sync,

		// Navigation methods
		slide: slide,
		left: navigateLeft,
		right: navigateRight,
		up: navigateUp,
		down: navigateDown,
		prev: navigatePrev,
		next: navigateNext,

		// Fragment methods
		navigateFragment: navigateFragment,
		prevFragment: previousFragment,
		nextFragment: nextFragment,

		// Deprecated aliases
		navigateTo: slide,
		navigateLeft: navigateLeft,
		navigateRight: navigateRight,
		navigateUp: navigateUp,
		navigateDown: navigateDown,
		navigatePrev: navigatePrev,
		navigateNext: navigateNext,

		// Forces an update in slide layout
		layout: layout,

		// Randomizes the order of slides
		shuffle: shuffle,

		// Returns an object with the available routes as booleans (left/right/top/bottom)
		availableRoutes: availableRoutes,

		// Returns an object with the available fragments as booleans (prev/next)
		availableFragments: availableFragments,

		// Toggles a help overlay with keyboard shortcuts
		toggleHelp: toggleHelp,

		// Toggles the overview mode on/off
		toggleOverview: toggleOverview,

		// Toggles the "black screen" mode on/off
		togglePause: togglePause,

		// Toggles the auto slide mode on/off
		toggleAutoSlide: toggleAutoSlide,

		// State checks
		isOverview: isOverview,
		isPaused: isPaused,
		isAutoSliding: isAutoSliding,
		isSpeakerNotes: isSpeakerNotes,

		// Slide preloading
		loadSlide: loadSlide,
		unloadSlide: unloadSlide,

		// Adds or removes all internal event listeners (such as keyboard)
		addEventListeners: addEventListeners,
		removeEventListeners: removeEventListeners,

		// Facility for persisting and restoring the presentation state
		getState: getState,
		setState: setState,

		// Presentation progress
		getSlidePastCount: getSlidePastCount,

		// Presentation progress on range of 0-1
		getProgress: getProgress,

		// Returns the indices of the current, or specified, slide
		getIndices: getIndices,

		// Returns an Array of all slides
		getSlides: getSlides,

		// Returns the total number of slides
		getTotalSlides: getTotalSlides,

		// Returns the slide element at the specified index
		getSlide: getSlide,

		// Returns the slide background element at the specified index
		getSlideBackground: getSlideBackground,

		// Returns the speaker notes string for a slide, or null
		getSlideNotes: getSlideNotes,

		// Returns the previous slide element, may be null
		getPreviousSlide: function() {
			return previousSlide;
		},

		// Returns the current slide element
		getCurrentSlide: function() {
			return currentSlide;
		},

		// Returns the current scale of the presentation content
		getScale: function() {
			return scale;
		},

		// Returns the current configuration object
		getConfig: function() {
			return config;
		},

		// Helper method, retrieves query string as a key/value hash
		getQueryHash: function() {
			var query = {};

			location.search.replace( /[A-Z0-9]+?=([\w\.%-]*)/gi, function(a) {
				query[ a.split( '=' ).shift() ] = a.split( '=' ).pop();
			} );

			// Basic deserialization
			for( var i in query ) {
				var value = query[ i ];

				query[ i ] = deserialize( unescape( value ) );
			}

			return query;
		},

		// Returns true if we're currently on the first slide
		isFirstSlide: function() {
			return ( indexh === 0 && indexv === 0 );
		},

		// Returns true if we're currently on the last slide
		isLastSlide: function() {
			if( currentSlide ) {
				// Does this slide has next a sibling?
				if( currentSlide.nextElementSibling ) return false;

				// If it's vertical, does its parent have a next sibling?
				if( isVerticalSlide( currentSlide ) && currentSlide.parentNode.nextElementSibling ) return false;

				return true;
			}

			return false;
		},

		// Checks if reveal.js has been loaded and is ready for use
		isReady: function() {
			return loaded;
		},

		// Forward event binding to the reveal DOM element
		addEventListener: function( type, listener, useCapture ) {
			if( 'addEventListener' in window ) {
				( dom.wrapper || document.querySelector( '.reveal' ) ).addEventListener( type, listener, useCapture );
			}
		},
		removeEventListener: function( type, listener, useCapture ) {
			if( 'addEventListener' in window ) {
				( dom.wrapper || document.querySelector( '.reveal' ) ).removeEventListener( type, listener, useCapture );
			}
		},

		// Programatically triggers a keyboard event
		triggerKey: function( keyCode ) {
			onDocumentKeyDown( { keyCode: keyCode } );
		},

		// Registers a new shortcut to include in the help overlay
		registerKeyboardShortcut: function( key, value ) {
			keyboardShortcuts[key] = value;
		}
	};

	return Reveal;

}));
```
