## Supplementary material for "Artistoo: build, share, and explore simulations of cells and tissues in the web browser": dragdealer.js

```
/**
 * Dragdealer.js 0.9.8
 * http://github.com/skidding/dragdealer
 *
 * (c) 2010+ Ovidiu Cherecheș
 * http://skidding.mit-license.org
 */

(function (root, factory) {
  if (typeof define === 'function' && define.amd) {
    // AMD. Register as an anonymous module.
    define(factory);
  } else {
    // Browser globals
    root.Dragdealer = factory();
  }
}(this, function () {

var Dragdealer = function(wrapper, options) {
  /**
   * Drag-based component that works around two basic DOM elements.
   *
   *   - The wrapper: The top-level element with the .dragdealer class. We
   *                  create a Dragdealer instance with the wrapper as the
   *                  first constructor parameter (it can either receive the ID
   *                  of the wrapper, or the element itself.) The wrapper
   *                  establishes the dragging bounds.
   *
   *   - The handle: A child of the wrapper element, div with a required
   *                 .handle class (may be overridden in options). This will be
   *                 the dragged element, constrained by the wrapper's bounds.
   *
   *
   * The handle can be both smaller or bigger than the wrapper.
   *
   *   - When the handle is smaller, Dragdealer will act as a regular slider,
   *     enabling the handle to be dragged from one side of the wrapper to
   *     another.
   *
   *   - When the handle is bigger, Dragdealer will act a mask for a draggable
   *     surface, where the handle is the draggable surface contrained by the
   *     smaller bounds of the wrapper. The drag action in this case is used
   *     to reveal and "discover" partial content at a time.
   *
   *
   * Simple usage:
   *
   *   // JavaScript
   *   new Dragdealer('simple-slider');
   *
   *   <!-- HTML -->
   *   <div id="simple-slider" class="dragdealer">
   *     <div class="handle">drag me</div>
   *   </div>
   *
   *
   * The second parameter of the Dragdealer constructor is an object used for
   * specifying any of the supported options. All of them are optional.
   *
   *   - bool disabled=false: Init Dragdealer in a disabled state. The handle
   *                          will have a .disabled class.
   *
   *   - bool horizontal=true: Enable horizontal dragging.
   *
   *   - bool vertical=false: Enable vertical dragging.
   *
   *   - number x=0: Initial horizontal (left) position. Accepts a float number
   *                 value between 0 and 1. Read below about positioning in
   *                 Dragdealer.
   *
   *   - number y=0: Initial vertical (top) position. Accepts a float number
   *                 value between 0 and 1. Read below about positoning in
   *                 Dragdealer.
   *
   *   - number steps=0: Limit the positioning of the handle within the bounds
   *                     of the wrapper, by defining a virtual grid made out of
   *                     a number of equally-spaced steps. This restricts
   *                     placing the handle anywhere in-between these steps.
   *                     E.g. setting 3 steps to a regular slider will only
   *                     allow you to move it to the left, to the right or
   *                     exactly in the middle.
   *
   *   - bool snap=false: When a number of steps is set, snap the position of
   *                      the handle to its closest step instantly, even when
   *                      dragging.
   *
   *   - number speed=0.1: Speed can be set between 0 and 1, with 1 being the
   *                       fastest. It represents how fast the handle will slide
   *                       to position after you mouse up.
   *
   *   - bool slide=true: Slide handle after releasing it, depending on the
   *                      movement speed before the mouse/touch release. The
   *                      formula for calculating how much will the handle
   *                      slide after releasing it is defined by simply
   *                      extending the movement of the handle in the current
   *                      direction, with the last movement unit times four (a
   *                      movement unit is considered the distance crossed
   *                      since the last animation loop, which is currently
   *                      25ms.) So if you were to drag the handle 50px in the
   *                      blink of an eye, it will slide another 200px in the
   *                      same direction. Steps interfere with this formula, as
   *                      the closest step is calculated before the sliding
   *                      distance.
   *
   *   - bool loose=false: Loosen-up wrapper boundaries when dragging. This
   *                       allows the handle to be *slightly* dragged outside
   *                       the bounds of the wrapper, but slides it back to the
   *                       margins of the wrapper upon release. The formula for
   *                       calculating how much the handle exceeds the wrapper
   *                       bounds is made out of the actual drag distance
   *                       divided by 4. E.g. Pulling a slider outside its
   *                       frame by 100px will only position it 25px outside
   *                       the frame.
   *
   *   - number top=0: Top padding between the wrapper and the handle.
   *
   *   - number bottom=0: Bottom padding between the wrapper and the handle.
   *
   *   - number left=0: Left padding between the wrapper and the handle.
   *
   *   - number right=0: Right padding between the wrapper and the handle.
   *
   *   - fn callback(x, y): Called when releasing handle, with the projected
   *                        x, y position of the handle. Projected value means
   *                        the value the slider will have after finishing a
   *                        sliding animation, caused by either a step
   *                        restriction or drag motion (see steps and slide
   *                        options.) This implies that the actual position of
   *                        the handle at the time this callback is called
   *                        might not yet reflect the x, y values received.
   *
   *   - fn dragStopCallback(x,y): Same as callback(x,y) but only called after
   *                               a drag motion, not after setting the step
   *                               manually.
   *
   *   - fn dragStartCallback(x,y): Same as dragStopCallback(x,y) but called at
   *                                the beginning of a drag motion and with the
   *                                sliders initial x, y values.
   *
   *   - fn animationCallback(x, y): Called every animation loop, as long as
   *                                 the handle is being dragged or in the
   *                                 process of a sliding animation. The x, y
   *                                 positional values received by this
   *                                 callback reflect the exact position of the
   *                                 handle DOM element, which includes
   *                                 exceeding values (even negative values)
   *                                 when the loose option is set true.
   *
   *   - string handleClass='handle': Custom class of handle element.
   *
   *   - bool css3=true: Use css3 transform in modern browsers instead of
   *                     absolute positioning.
   *
   *   - fn customRequestAnimationFrame: Provide custom requestAnimationFrame
   *                                     function (used in tests).
   *   - fn customCancelAnimationFrame: Provide custom cancelAnimationFrame
   *                                    function (used in tests).
   *
   * Dragdealer also has a few methods to interact with, post-initialization.
   *
   *   - disable: Disable dragging of a Dragdealer instance. Just as with the
   *              disabled option, the handle will receive a .disabled class
   *
   *   - enable: Enable dragging of a Dragdealer instance. The .disabled class
   *             of the handle will be removed.
   *
   *   - reflow: Recalculate the wrapper bounds of a Dragdealer instance, used
   *             when the wrapper is responsive and its parent container
   *             changed its size, or after changing the size of the wrapper
   *             directly.
   *
   *   - getValue: Get the value of a Dragdealer instance programatically. The
   *               value is returned as an [x, y] tuple and is the equivalent
   *               of the (projected) value returned by the regular callback,
   *               not animationCallback.
   *
   *   - getStep: Same as getValue, but the value returned is in step
   *              increments (see steps option)
   *
   *   - setValue(x, y, snap=false): Set the value of a Dragdealer instance
   *                                 programatically. The 3rd parameter allows
   *                                 to snap the handle directly to the desired
   *                                 value, without any sliding transition.
   *
   *   - setStep(x, y, snap=false): Same as setValue, but the value is received
   *                                in step increments (see steps option)
   *
   *
   * Positioning in Dragdealer:
   *
   *   Besides the top, bottom, left and right paddings, which represent a
   *   number of pixels, Dragdealer uses a [0, 1]-based positioning. Both
   *   horizontal and vertical positions are represented by ratios between 0
   *   and 1. This allows the Dragdealer wrapper to have a responsive size and
   *   not revolve around a specific number of pixels. This is how the x, y
   *   options are set, what the callback args contain and what values the
   *   setValue method expects. Once picked up, the ratios can be scaled and
   *   mapped to match any real-life system of coordinates or dimensions.
   */
  this.options = this.applyDefaults(options || {});
  this.bindMethods();
  this.wrapper = this.getWrapperElement(wrapper);
  if (!this.wrapper) {
    return;
  }
  this.handle = this.getHandleElement(this.wrapper, this.options.handleClass);
  if (!this.handle) {
    return;
  }
  this.init();
  this.bindEventListeners();
};


Dragdealer.prototype = {
  defaults: {
    disabled: false,
    horizontal: true,
    vertical: false,
    slide: true,
    steps: 0,
    snap: false,
    loose: false,
    speed: 0.1,
    xPrecision: 0,
    yPrecision: 0,
    handleClass: 'handle',
    css3: true,
    activeClass: 'active',
    tapping: true
  },
  init: function() {
    if (this.options.css3) {
      triggerWebkitHardwareAcceleration(this.handle);
    }
    this.value = {
      prev: [-1, -1],
      current: [this.options.x || 0, this.options.y || 0],
      target: [this.options.x || 0, this.options.y || 0]
    };
    this.offset = {
      wrapper: [0, 0],
      mouse: [0, 0],
      prev: [-999999, -999999],
      current: [0, 0],
      target: [0, 0]
    };
    this.change = [0, 0];
    this.stepRatios = this.calculateStepRatios();

    this.activity = false;
    this.dragging = false;
    this.tapping = false;

    this.reflow();
    if (this.options.disabled) {
      this.disable();
    }
  },
  applyDefaults: function(options) {
    for (var k in this.defaults) {
      if (!options.hasOwnProperty(k)) {
        options[k] = this.defaults[k];
      }
    }
    return options;
  },
  getWrapperElement: function(wrapper) {
    if (typeof(wrapper) == 'string') {
      return document.getElementById(wrapper);
    } else {
      return wrapper;
    }
  },
  getHandleElement: function(wrapper, handleClass) {
    var childElements,
        handleClassMatcher,
        i;
    if (wrapper.getElementsByClassName) {
      childElements = wrapper.getElementsByClassName(handleClass);
      if (childElements.length > 0) {
        return childElements[0];
      }
    } else {
      handleClassMatcher = new RegExp('(^|\\s)' + handleClass + '(\\s|$)');
      childElements = wrapper.getElementsByTagName('*');
      for (i = 0; i < childElements.length; i++) {
        if (handleClassMatcher.test(childElements[i].className)) {
          return childElements[i];
        }
      }
    }
  },
  calculateStepRatios: function() {
    var stepRatios = [];
    if (this.options.steps >= 1) {
      for (var i = 0; i <= this.options.steps - 1; i++) {
        if (this.options.steps > 1) {
          stepRatios[i] = i / (this.options.steps - 1);
        } else {
          // A single step will always have a 0 value
          stepRatios[i] = 0;
        }
      }
    }
    return stepRatios;
  },
  setWrapperOffset: function() {
    this.offset.wrapper = Position.get(this.wrapper);
  },
  calculateBounds: function() {
    // Apply top/bottom/left and right padding options to wrapper extremities
    // when calculating its bounds
    var bounds = {
      top: this.options.top || 0,
      bottom: -(this.options.bottom || 0) + this.wrapper.offsetHeight,
      left: this.options.left || 0,
      right: -(this.options.right || 0) + this.wrapper.offsetWidth
    };
    // The available width and height represents the horizontal and vertical
    // space the handle has for moving. It is determined by the width and
    // height of the wrapper, minus the width and height of the handle
    bounds.availWidth = (bounds.right - bounds.left) - this.handle.offsetWidth;
    bounds.availHeight = (bounds.bottom - bounds.top) - this.handle.offsetHeight;
    return bounds;
  },
  calculateValuePrecision: function() {
    // The sliding transition works by dividing itself until it reaches a min
    // value step; because Dragdealer works with [0-1] values, we need this
    // "min value step" to represent a pixel when applied to the real handle
    // position within the DOM. The xPrecision/yPrecision options can be
    // specified to increase the granularity when we're controlling larger
    // objects from one of the callbacks
    var xPrecision = this.options.xPrecision || Math.abs(this.bounds.availWidth),
        yPrecision = this.options.yPrecision || Math.abs(this.bounds.availHeight);
    return [
      xPrecision ? 1 / xPrecision : 0,
      yPrecision ? 1 / yPrecision : 0
    ];
  },
  bindMethods: function() {
    if (typeof(this.options.customRequestAnimationFrame) === 'function') {
      this.requestAnimationFrame = bind(this.options.customRequestAnimationFrame, window);
    } else {
      this.requestAnimationFrame = bind(requestAnimationFrame, window);
    }
    if (typeof(this.options.customCancelAnimationFrame) === 'function') {
      this.cancelAnimationFrame = bind(this.options.customCancelAnimationFrame, window);
    } else {
      this.cancelAnimationFrame = bind(cancelAnimationFrame, window);
    }
    this.animateWithRequestAnimationFrame = bind(this.animateWithRequestAnimationFrame, this);
    this.animate = bind(this.animate, this);
    this.onHandleMouseDown = bind(this.onHandleMouseDown, this);
    this.onHandleTouchStart = bind(this.onHandleTouchStart, this);
    this.onDocumentMouseMove = bind(this.onDocumentMouseMove, this);
    this.onWrapperTouchMove = bind(this.onWrapperTouchMove, this);
    this.onWrapperMouseDown = bind(this.onWrapperMouseDown, this);
    this.onWrapperTouchStart = bind(this.onWrapperTouchStart, this);
    this.onDocumentMouseUp = bind(this.onDocumentMouseUp, this);
    this.onDocumentTouchEnd = bind(this.onDocumentTouchEnd, this);
    this.onHandleClick = bind(this.onHandleClick, this);
    this.onWindowResize = bind(this.onWindowResize, this);
  },
  bindEventListeners: function() {
    // Start dragging
    addEventListener(this.handle, 'mousedown', this.onHandleMouseDown);
    addEventListener(this.handle, 'touchstart', this.onHandleTouchStart);
    // While dragging
    addEventListener(document, 'mousemove', this.onDocumentMouseMove);
    addEventListener(this.wrapper, 'touchmove', this.onWrapperTouchMove);
    // Start tapping
    addEventListener(this.wrapper, 'mousedown', this.onWrapperMouseDown);
    addEventListener(this.wrapper, 'touchstart', this.onWrapperTouchStart);
    // Stop dragging/tapping
    addEventListener(document, 'mouseup', this.onDocumentMouseUp);
    addEventListener(document, 'touchend', this.onDocumentTouchEnd);

    addEventListener(this.handle, 'click', this.onHandleClick);
    addEventListener(window, 'resize', this.onWindowResize);

    this.animate(false, true);
    this.interval = this.requestAnimationFrame(this.animateWithRequestAnimationFrame);

  },
  unbindEventListeners: function() {
    removeEventListener(this.handle, 'mousedown', this.onHandleMouseDown);
    removeEventListener(this.handle, 'touchstart', this.onHandleTouchStart);
    removeEventListener(document, 'mousemove', this.onDocumentMouseMove);
    removeEventListener(this.wrapper, 'touchmove', this.onWrapperTouchMove);
    removeEventListener(this.wrapper, 'mousedown', this.onWrapperMouseDown);
    removeEventListener(this.wrapper, 'touchstart', this.onWrapperTouchStart);
    removeEventListener(document, 'mouseup', this.onDocumentMouseUp);
    removeEventListener(document, 'touchend', this.onDocumentTouchEnd);
    removeEventListener(this.handle, 'click', this.onHandleClick);
    removeEventListener(window, 'resize', this.onWindowResize);
    this.cancelAnimationFrame(this.interval);
  },
  onHandleMouseDown: function(e) {
    Cursor.refresh(e);
    preventEventDefaults(e);
    stopEventPropagation(e);
    this.activity = false;
    this.startDrag();
  },
  onHandleTouchStart: function(e) {
    Cursor.refresh(e);
    // Unlike in the `mousedown` event handler, we don't prevent defaults here,
    // because this would disable the dragging altogether. Instead, we prevent
    // it in the `touchmove` handler. Read more about touch events
    // https://developer.mozilla.org/en-US/docs/Web/Guide/Events/Touch_events#Handling_clicks
    stopEventPropagation(e);
    this.activity = false;
    this.startDrag();
  },
  onDocumentMouseMove: function(e) {
    Cursor.refresh(e);
    if (this.dragging) {
      this.activity = true;
      preventEventDefaults(e);
    }
  },
  onWrapperTouchMove: function(e) {
    Cursor.refresh(e);
    // Dragging on a disabled axis (horizontal or vertical) shouldn't prevent
    // defaults on touch devices. !this.activity denotes this is the first move
    // inside a drag action; you can drag in any direction after this point if
    // the dragging wasn't stopped
    if (!this.activity && this.draggingOnDisabledAxis()) {
      if (this.dragging) {
        this.stopDrag();
      }
      return;
    }
    // Read comment in `onHandleTouchStart` above, to understand why we're
    // preventing defaults here and not there
    preventEventDefaults(e);
    this.activity = true;
  },
  onWrapperMouseDown: function(e) {
    Cursor.refresh(e);
    preventEventDefaults(e);
    this.startTap();
  },
  onWrapperTouchStart: function(e) {
    Cursor.refresh(e);
    preventEventDefaults(e);
    this.startTap();
  },
  onDocumentMouseUp: function(e) {
    this.stopDrag();
    this.stopTap();
  },
  onDocumentTouchEnd: function(e) {
    this.stopDrag();
    this.stopTap();
  },
  onHandleClick: function(e) {
    // We keep track if any dragging activity has been made between the
    // mouse/touch down and up events; based on this we allow or cancel a click
    // event from inside the handle. i.e. Click events shouldn't be triggered
    // when dragging, but should be allowed when clicking still
    if (this.activity) {
      preventEventDefaults(e);
      stopEventPropagation(e);
    }
  },
  onWindowResize: function(e) {
    this.reflow();
  },
  enable: function() {
    this.disabled = false;
    this.handle.className = this.handle.className.replace(/\s?disabled/g, '');
  },
  disable: function() {
    this.disabled = true;
    this.handle.className += ' disabled';
  },
  reflow: function() {
    this.setWrapperOffset();
    this.bounds = this.calculateBounds();
    this.valuePrecision = this.calculateValuePrecision();
    this.updateOffsetFromValue();
  },
  getStep: function() {
    return [
      this.getStepNumber(this.value.target[0]),
      this.getStepNumber(this.value.target[1])
    ];
  },
  getValue: function() {
    return this.value.target;
  },
  setStep: function(x, y, snap) {
    this.setValue(
      this.options.steps && x > 1 ? (x - 1) / (this.options.steps - 1) : 0,
      this.options.steps && y > 1 ? (y - 1) / (this.options.steps - 1) : 0,
      snap
    );
  },
  setValue: function(x, y, snap) {
    this.setTargetValue([x, y || 0]);
    if (snap) {
      this.groupCopy(this.value.current, this.value.target);
      // Since the current value will be equal to the target one instantly, the
      // animate function won't get to run so we need to update the positions
      // and call the callbacks manually
      this.updateOffsetFromValue();
      this.callAnimationCallback();
    }
  },
  startTap: function() {
    if (this.disabled || !this.options.tapping) {
      return;
    }

    this.tapping = true;
    this.setWrapperOffset();

    this.setTargetValueByOffset([
      Cursor.x - this.offset.wrapper[0] - (this.handle.offsetWidth / 2),
      Cursor.y - this.offset.wrapper[1] - (this.handle.offsetHeight / 2)
    ]);
  },
  stopTap: function() {
    if (this.disabled || !this.tapping) {
      return;
    }
    this.tapping = false;

    this.setTargetValue(this.value.current);
  },
  startDrag: function() {
    if (this.disabled) {
      return;
    }
    this.dragging = true;
    this.setWrapperOffset();

    this.offset.mouse = [
      Cursor.x - Position.get(this.handle)[0],
      Cursor.y - Position.get(this.handle)[1]
    ];
    if (!this.wrapper.className.match(this.options.activeClass)) {
      this.wrapper.className += ' ' + this.options.activeClass;
    }
    this.callDragStartCallback();
  },
  stopDrag: function() {
    if (this.disabled || !this.dragging) {
      return;
    }
    this.dragging = false;

    var target = this.groupClone(this.value.current);
    if (this.options.slide) {
      var ratioChange = this.change;
      target[0] += ratioChange[0] * 4;
      target[1] += ratioChange[1] * 4;
    }
    this.setTargetValue(target);
    this.wrapper.className = this.wrapper.className.replace(' ' + this.options.activeClass, '');
    this.callDragStopCallback();
  },
  callAnimationCallback: function() {
    var value = this.value.current;
    if (this.options.snap && this.options.steps > 1) {
      value = this.getClosestSteps(value);
    }
    if (!this.groupCompare(value, this.value.prev)) {
      if (typeof(this.options.animationCallback) == 'function') {
        this.options.animationCallback.call(this, value[0], value[1]);
      }
      this.groupCopy(this.value.prev, value);
    }
  },
  callTargetCallback: function() {
    if (typeof(this.options.callback) == 'function') {
      this.options.callback.call(this, this.value.target[0], this.value.target[1]);
    }
  },
  callDragStartCallback: function() {
    if (typeof(this.options.dragStartCallback) == 'function') {
      this.options.dragStartCallback.call(this, this.value.target[0], this.value.target[1]);
    }
  },
  callDragStopCallback: function() {
    if (typeof(this.options.dragStopCallback) == 'function') {
      this.options.dragStopCallback.call(this, this.value.target[0], this.value.target[1]);
    }
  },
  animateWithRequestAnimationFrame: function (time) {
    if (time) {
      // using requestAnimationFrame
      this.timeOffset = this.timeStamp ? time - this.timeStamp : 0;
      this.timeStamp = time;
    } else {
      // using setTimeout(callback, 25) polyfill
      this.timeOffset = 25;
    }
    this.animate();
    this.interval = this.requestAnimationFrame(this.animateWithRequestAnimationFrame);
  },
  animate: function(direct, first) {
    if (direct && !this.dragging) {
      return;
    }
    if (this.dragging) {
      var prevTarget = this.groupClone(this.value.target);

      var offset = [
        Cursor.x - this.offset.wrapper[0] - this.offset.mouse[0],
        Cursor.y - this.offset.wrapper[1] - this.offset.mouse[1]
      ];
      this.setTargetValueByOffset(offset, this.options.loose);

      this.change = [
        this.value.target[0] - prevTarget[0],
        this.value.target[1] - prevTarget[1]
      ];
    }
    if (this.dragging || first) {
      this.groupCopy(this.value.current, this.value.target);
    }
    if (this.dragging || this.glide() || first) {
      this.updateOffsetFromValue();
      this.callAnimationCallback();
    }
  },
  glide: function() {
    var diff = [
      this.value.target[0] - this.value.current[0],
      this.value.target[1] - this.value.current[1]
    ];
    if (!diff[0] && !diff[1]) {
      return false;
    }
    if (Math.abs(diff[0]) > this.valuePrecision[0] ||
        Math.abs(diff[1]) > this.valuePrecision[1]) {
      this.value.current[0] += diff[0] * Math.min(this.options.speed * this.timeOffset / 25, 1);
      this.value.current[1] += diff[1] * Math.min(this.options.speed * this.timeOffset / 25, 1);
    } else {
      this.groupCopy(this.value.current, this.value.target);
    }
    return true;
  },
  updateOffsetFromValue: function() {
    if (!this.options.snap) {
      this.offset.current = this.getOffsetsByRatios(this.value.current);
    } else {
      this.offset.current = this.getOffsetsByRatios(
        this.getClosestSteps(this.value.current)
      );
    }
    if (!this.groupCompare(this.offset.current, this.offset.prev)) {
      this.renderHandlePosition();
      this.groupCopy(this.offset.prev, this.offset.current);
    }
  },
  renderHandlePosition: function() {

    var transform = '';
    if (this.options.css3 && StylePrefix.transform) {
      if (this.options.horizontal) {
        transform += 'translateX(' + this.offset.current[0] + 'px)';
      }
      if (this.options.vertical) {
        transform += ' translateY(' + this.offset.current[1] + 'px)';
      }
      this.handle.style[StylePrefix.transform] = transform;
      return;
    }

    if (this.options.horizontal) {
      this.handle.style.left = this.offset.current[0] + 'px';
    }
    if (this.options.vertical) {
      this.handle.style.top = this.offset.current[1] + 'px';
    }
  },
  setTargetValue: function(value, loose) {
    var target = loose ? this.getLooseValue(value) : this.getProperValue(value);

    this.groupCopy(this.value.target, target);
    this.offset.target = this.getOffsetsByRatios(target);

    this.callTargetCallback();
  },
  setTargetValueByOffset: function(offset, loose) {
    var value = this.getRatiosByOffsets(offset);
    var target = loose ? this.getLooseValue(value) : this.getProperValue(value);

    this.groupCopy(this.value.target, target);
    this.offset.target = this.getOffsetsByRatios(target);
  },
  getLooseValue: function(value) {
    var proper = this.getProperValue(value);
    return [
      proper[0] + ((value[0] - proper[0]) / 4),
      proper[1] + ((value[1] - proper[1]) / 4)
    ];
  },
  getProperValue: function(value) {
    var proper = this.groupClone(value);

    proper[0] = Math.max(proper[0], 0);
    proper[1] = Math.max(proper[1], 0);
    proper[0] = Math.min(proper[0], 1);
    proper[1] = Math.min(proper[1], 1);

    if ((!this.dragging && !this.tapping) || this.options.snap) {
      if (this.options.steps > 1) {
        proper = this.getClosestSteps(proper);
      }
    }
    return proper;
  },
  getRatiosByOffsets: function(group) {
    return [
      this.getRatioByOffset(group[0], this.bounds.availWidth, this.bounds.left),
      this.getRatioByOffset(group[1], this.bounds.availHeight, this.bounds.top)
    ];
  },
  getRatioByOffset: function(offset, range, padding) {
    return range ? (offset - padding) / range : 0;
  },
  getOffsetsByRatios: function(group) {
    return [
      this.getOffsetByRatio(group[0], this.bounds.availWidth, this.bounds.left),
      this.getOffsetByRatio(group[1], this.bounds.availHeight, this.bounds.top)
    ];
  },
  getOffsetByRatio: function(ratio, range, padding) {
    return Math.round(ratio * range) + padding;
  },
  getStepNumber: function(value) {
    // Translate a [0-1] value into a number from 1 to N steps (set using the
    // "steps" option)
    return this.getClosestStep(value) * (this.options.steps - 1) + 1;
  },
  getClosestSteps: function(group) {
    return [
      this.getClosestStep(group[0]),
      this.getClosestStep(group[1])
    ];
  },
  getClosestStep: function(value) {
    var k = 0;
    var min = 1;
    for (var i = 0; i <= this.options.steps - 1; i++) {
      if (Math.abs(this.stepRatios[i] - value) < min) {
        min = Math.abs(this.stepRatios[i] - value);
        k = i;
      }
    }
    return this.stepRatios[k];
  },
  groupCompare: function(a, b) {
    return a[0] == b[0] && a[1] == b[1];
  },
  groupCopy: function(a, b) {
    a[0] = b[0];
    a[1] = b[1];
  },
  groupClone: function(a) {
    return [a[0], a[1]];
  },
  draggingOnDisabledAxis: function() {
    return (!this.options.horizontal && Cursor.xDiff > Cursor.yDiff) ||
           (!this.options.vertical && Cursor.yDiff > Cursor.xDiff);
  }
};


var bind = function(fn, context) {
  /**
   * CoffeeScript-like function to bind the scope of a method to an instance,
   * the context of that method, regardless from where it is called
   */
  return function() {
    return fn.apply(context, arguments);
  };
};

// Cross-browser vanilla JS event handling

var addEventListener = function(element, type, callback) {
  if (element.addEventListener) {
    element.addEventListener(type, callback, false);
  } else if (element.attachEvent) {
    element.attachEvent('on' + type, callback);
  }
};

var removeEventListener = function(element, type, callback) {
  if (element.removeEventListener) {
    element.removeEventListener(type, callback, false);
  } else if (element.detachEvent) {
    element.detachEvent('on' + type, callback);
  }
};

var preventEventDefaults = function(e) {
  if (!e) {
    e = window.event;
  }
  if (e.preventDefault) {
    e.preventDefault();
  }
  e.returnValue = false;
};

var stopEventPropagation = function(e) {
  if (!e) {
    e = window.event;
  }
  if (e.stopPropagation) {
    e.stopPropagation();
  }
  e.cancelBubble = true;
};


var Cursor = {
  /**
   * Abstraction for making the combined mouse or touch position available at
   * any time.
   *
   * It picks up the "move" events as an independent component and simply makes
   * the latest x and y mouse/touch position of the user available at any time,
   * which is requested with Cursor.x and Cursor.y respectively.
   *
   * It can receive both mouse and touch events consecutively, extracting the
   * relevant meta data from each type of event.
   *
   * Cursor.refresh(e) is called to update the global x and y values, with a
   * genuine MouseEvent or a TouchEvent from an event listener, e.g.
   * mousedown/up or touchstart/end
   */
  x: 0,
  y: 0,
  xDiff: 0,
  yDiff: 0,
  refresh: function(e) {
    if (!e) {
      e = window.event;
    }
    if (e.type == 'mousemove') {
      this.set(e);
    } else if (e.touches) {
      this.set(e.touches[0]);
    }
  },
  set: function(e) {
    var lastX = this.x,
        lastY = this.y;
    if (e.clientX || e.clientY) {
      this.x = e.clientX;
      this.y = e.clientY;
    } else if (e.pageX || e.pageY) {
      this.x = e.pageX - document.body.scrollLeft - document.documentElement.scrollLeft;
      this.y = e.pageY - document.body.scrollTop - document.documentElement.scrollTop;
    }
    this.xDiff = Math.abs(this.x - lastX);
    this.yDiff = Math.abs(this.y - lastY);
  }
};


var Position = {
  /**
   * Helper for extracting position of a DOM element, relative to the viewport
   *
   * The get(obj) method accepts a DOM element as the only parameter, and
   * returns the position under a (x, y) tuple, as an array with two elements.
   */
  get: function(obj) {
    // Dragdealer relies on getBoundingClientRect to calculate element offsets,
    // but we want to be sure we don't throw any unhandled exceptions and break
    // other code from the page if running from in very old browser that doesn't
    // support this method
    var rect = {left: 0, top: 0};
    if (obj.getBoundingClientRect !== undefined) {
      rect = obj.getBoundingClientRect();
    }
    return [rect.left, rect.top];
  }
};


var StylePrefix = {
  transform: getPrefixedStylePropName('transform'),
  perspective: getPrefixedStylePropName('perspective'),
  backfaceVisibility: getPrefixedStylePropName('backfaceVisibility')
};

function getPrefixedStylePropName(propName) {
  var domPrefixes = 'Webkit Moz ms O'.split(' '),
      elStyle = document.documentElement.style;
  if (elStyle[propName] !== undefined) return propName; // Is supported unprefixed
  propName = propName.charAt(0).toUpperCase() + propName.substr(1);
  for (var i = 0; i < domPrefixes.length; i++) {
    if (elStyle[domPrefixes[i] + propName] !== undefined) {
      return domPrefixes[i] + propName; // Is supported with prefix
    }
  }
};

function triggerWebkitHardwareAcceleration(element) {
  if (StylePrefix.backfaceVisibility && StylePrefix.perspective) {
    element.style[StylePrefix.perspective] = '1000px';
    element.style[StylePrefix.backfaceVisibility] = 'hidden';
  }
};

var vendors = ['webkit', 'moz'];
var requestAnimationFrame = window.requestAnimationFrame;
var cancelAnimationFrame = window.cancelAnimationFrame;

for (var x = 0; x < vendors.length && !requestAnimationFrame; ++x) {
  requestAnimationFrame = window[vendors[x] + 'RequestAnimationFrame'];
  cancelAnimationFrame = window[vendors[x] + 'CancelAnimationFrame'] ||
                         window[vendors[x] + 'CancelRequestAnimationFrame'];
}

if (!requestAnimationFrame) {
  requestAnimationFrame = function (callback) {
    return setTimeout(callback, 25);
  };
  cancelAnimationFrame = clearTimeout;
}

return Dragdealer;

}));
```
