## Supplementary material for "Artistoo: build, share, and explore simulations of cells and tissues in the web browser": teaching-exercise.pdf

### Making a Cellular Potts Model of Collective Migration

Inge Wortel Johannes Textor

May 1st, 2020

In this exercise, you are going to use the Cellular Potts modeling framework to create cells *in silico*. You will first learn how to use the model's many parameters to create cells of realistic shapes and motility patterns. Then you will go a step further and model several migrating cells that interact with each other.

#### Learning Goals

- Understand how the different parameters of a Cellular Potts Model (CPM) interact with each other, and how this influences cell behavior.
- Realize that it can be difficult to tune the parameters of a CPM.
- Apply this knowledge to create different modes of cell migration in the CPM, and explain how this helps understand those migration modes.
- Apply the CPM to investigate the dynamics of collective cell migration *in silico*.

Please go to computational-immunology.org/cpm/collective.html. This is an implementation of a special version of the Cellular Potts Model in which cells can migrate [1]. Note that *this web page does not work properly in Internet Explorer*. It does work in either Firefox, Chrome, or Safari – so we recommend using any of those browsers for this exercise.

#### Notes on the CPM

Remember that in a CPM, copy attempts have a higher chance of succeeding if they lower the total energy $H$ of the system. In other words, $\Delta H = H_{\text{after copy}} - H_{\text{before copy}}$ should be negative to guarantee that the copy attempt will work. Copy attempts with positive $\Delta H$ can still succeed, but do so with a lower probability that depends on the temperature of the system. In general, the formula for $\Delta H$ of a CPM looks something like this:

$$\Delta H = \Delta H_{\text{adhesion}} + \Delta H_{\text{Volume}} + \Delta H_{...} + ... \tag{1}$$

Thus, we build up $\Delta H$ from all the different energetic factors we want the cell to consider – and we can always add stuff to make the model more complex. But what parameters do we need to tune to get realistic cells? In the following exercises, you will get a feel for how you can model cell behavior with a CPM by tuning the parameters that control the energy $\Delta H$.

#### Exercises

##### A very basic CPM

In this exercise, we will first examine a very basic CPM in which cells do not (yet) migrate. Make sure that the field is empty (hit refresh or "remove all cells"), and that the parameters have the following values:

| $\text{Adhesion}_{\text{cell-matrix}}$ | $\text{Adhesion}_{\text{cell-cell}}$ | Volume | $\lambda_{\text{Volume}}$ | Perimeter | $\lambda_{\text{P}}$ | $\text{Max}_{\text{Act}}$ | $\lambda_{\text{Act}}$ | T | Framerate |
|---|---|---|---|---|---|---|---|---|---|
| 20 | 0 | 500 | 50 | 340 | 0 | 0 | 0 | 20 | 1 |

We will now investigate how the basic CPM parameters – controlling adhesion, cell volume, and cell perimeter (circumference) – influence behavior (this means you can ignore the $\lambda_{\text{Act}}$ and $\text{Max}_{\text{Act}}$ parameters for now). This exercise is meant mostly to give you an idea of what the CPM parameters do, and the questions are to guide your thinking – so you don't have to write everything down. Try to spend no longer than 30-40 minutes on this exercise before continuing to the next.

1. Make sure all the parameters are set as in the table above, click "seed cell" and then "start". What do you see? What kind of motion does this cell have?

2. Now set the $\text{Adhesion}_{\text{cell-matrix}}$ to 0. What happens to the cell? Why do you think that happens? (Hint: look back to the description of the CPM and adhesion energy in the lecture...) Also try a negative value for $\text{Adhesion}_{\text{cell-matrix}}$. What is the meaning of positive or negative adhesion values here?

3. *(Optional)* Instead of setting the $\text{Adhesion}_{\text{cell-matrix}}$ back to 20, try setting the $\text{Adhesion}_{\text{cell-cell}}$ to -20 while having the Adhesion $_{\text{cell-matrix}}$ still at 0. Does that have the effect that you expected? Why/why not do you think that is? Hint: try drawing a grid like you saw in the lecture for a copy attempt you are interested in. Do the two adhesion energies change in the same way for that copy attempt?

4. Return to the parameters in the table above. With these parameters, the cell is given an ideal volume (500 pixels), and a "level of importance" of this volume for the energy ($\lambda_{\text{Volume}}$). Try making the cell bigger or smaller (what parameter should you change?). How can you make the volume unimportant for the energy – and what happens then? What happens when you make $\lambda_{\text{Volume}}$ really large (say, 1000)?

5. So far, we have considered $\Delta H$ with only terms for adhesion energy and cell volume. We will now investigate the effect of the cell perimeter (circumference). The cell already has a target perimeter (340), but this is currently not taken into account in the calculation of $\Delta H$. For that, we need to make $\lambda_{\text{P}}$ non-zero. Try setting it to 2. What happens to the cell? Try making the cell "membrane" more or less ruffled. How would you do that?

6. Set the perimeter to 340 and $\lambda_{\text{P}}$ to 2. Now change the $\text{Adhesion}_{\text{cell-matrix}}$ to 0 again. Does this have the same effect as it did in question 2? Why do you think that is?

7. *(Optional)* Play around with the volume and perimeter parameters for a while (using the adhesion parameters from the table, or try your own). How can you change the cell? Can you change the parameters independently of each other? And what happens if you change the temperature?

#### Cell migration: the Act model

We will now investigate the Act model, an extension of the CPM that allows the cells to migrate [1]. This model adds an extra term to the system energy, so that:

$$\Delta H = \Delta H_{\text{adhesion}} + \Delta H_{\text{Volume}} + \Delta H_{\text{Perimeter}} + \Delta H_{\text{Act}} \tag{2}$$

In this model, pixels that were recently added to the cell remember their recent "protrusive activity". This makes them more likely to protrude again. This positive feedback is controlled by the energy term $\Delta H_{\text{Act}}$, which is negative (favourable!) when a recently active pixel tries to copy itself into a less active pixel. The Act model has two extra parameters: $\lambda_{\text{Act}}$, which controls how important the positive feedback is relative to the other $\Delta H$ energies, and $\text{Max}_{\text{Act}}$, which determines how long pixels "remember" that they were active. In this exercise, we will see what happens when we vary those two parameters of the model. In particular, we will see that we can reproduce two very different "modes" of migration: amoeboid and keratocyte-like (see lecture).

Before starting this exercise, please refresh the page and ensure that the parameters of the CPM are set as follows:

| $\text{Adhesion}_{\text{cell-matrix}}$ | $\text{Adhesion}_{\text{cell-cell}}$ | Volume | $\lambda_{\text{Volume}}$ | Perimeter | $\lambda_{\text{P}}$ | $\text{Max}_{\text{Act}}$ | $\lambda_{\text{Act}}$ | T | Framerate |
|---|---|---|---|---|---|---|---|---|---|
| 20 | 0 | 500 | 50 | 340 | 2 | 20 | 0 | 20 | 1 |

1. Seed a cell and click "start". You should now see colored pixels at the border of the cell, which indicate the "activity" that pixels remember (because we have set $\text{Max}_{\text{Act}}$ to 20). Other than the color of the pixels, does the cell behave in a different way than with $\text{Max}_{\text{Act}}$ = 0? Why/why not?

2. Set $\lambda_{\text{Act}}$ to 100. Would you describe this movement as random or persistent?

3. *(Optional)* What happens when you set $\lambda_{\text{P}}$ to 0 now? Why do you think that is? (Reset it to 2 before going to the next question)

4. What happens when you increase $\lambda_{\text{Act}}$ further? (Try steps of 100).

5. *(Optional)* If you increase $\lambda_{\text{Act}}$ to very high values (eg 1000), the cell is prone to breaking in pieces. Can you fix that by altering some other CPM parameter again? (Note: you may have to increase $\lambda_{\text{Act}}$ further when you have done this... Does that make sense to you?)

6. Reset $\lambda_{\text{Act}}$ to 0, change $\text{Max}_{\text{Act}}$ to 80, and repeat questions 1,2, and 4 above. What do you see?

7. *(Optional)* If you have time, play around with different combinations of $\lambda_{\text{Act}}$ and $\text{Max}_{\text{Act}}$. Can you get a clue of what they are doing – beyond the mathematical description given above?

8. *(Optional)* Try halving or doubling the cell's target volume. That won't work. What do you need to change to get the same behavior as before? What does that mean for your model (in other words: to what extent are your choices of parameters important for the behaviour you see? How worried should you be about getting parameters "wrong" and drawing the wrong conclusions?)?

9. The nice thing about computer models is that you know exactly which "rules" you put in, and therefore that those are the only rules that the cells will follow. However, the behavior that follows from those "rules" is not always obvious (that's why we run the simulation in the first place!). In fact, the most interesting computer models can sometimes produce behavior that you did not explicitly put in there. We call this "emergent" behavior. Can you think of an example of emergent behavior in the Act model? What does that tell you about migration in a living cell?

#### Migration in a multicellular system

In this last exercise, we will investigate what happens when there are many cells. Before starting, please refresh the page to clear the grid, and then ensure that the parameters have the following values:

| $\text{Adhesion}_{\text{cell-matrix}}$ | $\text{Adhesion}_{\text{cell-cell}}$ | Volume | $\lambda_{\text{Volume}}$ | Perimeter | $\lambda_{\text{P}}$ | $\text{Max}_{\text{Act}}$ | $\lambda_{\text{Act}}$ | T | Framerate |
|---|---|---|---|---|---|---|---|---|---|
| 20 | 0 | 200 | 50 | 180 | 2 | 20 | 200 | 20 | 5 |

Note that the framerate is not actually a parameter of the model, but it specifies how often the program draws the updated grid (eg framerate of 5 means visualize only 1 in every 5 "frames" of the movie). Setting it to 5 will speed up the animation.

1. To seed many cells at once, click "+100 cells" (this will take a while...). What do you see?

2. Now, refresh the page, reset the parameters, increase $\text{Max}_{\text{Act}}$ to 80, and wait for a while to let the cells adjust. What happens? How is this different from what you saw with $\text{Max}_{\text{Act}}$ = 20?

3. Try other values of $\text{Max}_{\text{Act}}$ in between 20 and 80. What happens as you gradually increase $\text{Max}_{\text{Act}}$?

4. *(Optional)* What happens when you set $\text{Adhesion}_{\text{cell-cell}}$ to a negative value?

5. To what extent are your conclusions dependent on the density of cells on your grid? How could you test that?
