## Supplementary material for "Artistoo: build, share, and explore simulations of cells and tissues in the web browser": artistoo.js

```
var CPM = (function (exports) {
	'use strict';

	/*
	  https://github.com/banksean wrapped Makoto Matsumoto and Takuji Nishimura's code in a namespace
	  so it's better encapsulated. Now you can have multiple random number generators
	  and they won't stomp all over eachother's state.

	  If you want to use this as a substitute for Math.random(), use the random()
	  method like so:

	  var m = new MersenneTwister();
	  var randomNumber = m.random();

	  You can also call the other genrand_{foo}() methods on the instance.

	  If you want to use a specific seed in order to get a repeatable random
	  sequence, pass an integer into the constructor:

	  var m = new MersenneTwister(123);

	  and that will always produce the same random sequence.

	  Sean McCullough
	*/

	/*
	   A C-program for MT19937, with initialization improved 2002/1/26.
	   Coded by Takuji Nishimura and Makoto Matsumoto.

	   Before using, initialize the state by using init_seed(seed)
	   or init_by_array(init_key, key_length).

	   Copyright (C) 1997 - 2002, Makoto Matsumoto and Takuji Nishimura,
	   All rights reserved.

	   Redistribution and use in source and binary forms, with or without
	   modification, are permitted provided that the following conditions
	   are met:

	     1. Redistributions of source code must retain the above copyright
	        notice, this list of conditions and the following disclaimer.

	     2. Redistributions in binary form must reproduce the above copyright
	        notice, this list of conditions and the following disclaimer in the
	        documentation and/or other materials provided with the distribution.

	     3. The names of its contributors may not be used to endorse or promote
	        products derived from this software without specific prior written
	        permission.

	   THIS SOFTWARE IS PROVIDED BY THE COPYRIGHT HOLDERS AND CONTRIBUTORS
	   "AS IS" AND ANY EXPRESS OR IMPLIED WARRANTIES, INCLUDING, BUT NOT
	   LIMITED TO, THE IMPLIED WARRANTIES OF MERCHANTABILITY AND FITNESS FOR
	   A PARTICULAR PURPOSE ARE DISCLAIMED.  IN NO EVENT SHALL THE COPYRIGHT OWNER OR
	   CONTRIBUTORS BE LIABLE FOR ANY DIRECT, INDIRECT, INCIDENTAL, SPECIAL,
	   EXEMPLARY, OR CONSEQUENTIAL DAMAGES (INCLUDING, BUT NOT LIMITED TO,
	   PROCUREMENT OF SUBSTITUTE GOODS OR SERVICES; LOSS OF USE, DATA, OR
	   PROFITS; OR BUSINESS INTERRUPTION) HOWEVER CAUSED AND ON ANY THEORY OF
	   LIABILITY, WHETHER IN CONTRACT, STRICT LIABILITY, OR TORT (INCLUDING
	   NEGLIGENCE OR OTHERWISE) ARISING IN ANY WAY OUT OF THE USE OF THIS
	   SOFTWARE, EVEN IF ADVISED OF THE POSSIBILITY OF SUCH DAMAGE.


	   Any feedback is very welcome.
	   http://www.math.sci.hiroshima-u.ac.jp/~m-mat/MT/emt.html
	   email: m-mat @ math.sci.hiroshima-u.ac.jp (remove space)
	*/

	var MersenneTwister = function(seed) {
		if (seed == undefined) {
			seed = new Date().getTime();
		}

		/* Period parameters */
		this.N = 624;
		this.M = 397;
		this.MATRIX_A = 0x9908b0df;   /* constant vector a */
		this.UPPER_MASK = 0x80000000; /* most significant w-r bits */
		this.LOWER_MASK = 0x7fffffff; /* least significant r bits */

		this.mt = new Array(this.N); /* the array for the state vector */
		this.mti=this.N+1; /* mti==N+1 means mt[N] is not initialized */

		if (seed.constructor == Array) {
			this.init_by_array(seed, seed.length);
		}
		else {
			this.init_seed(seed);
		}
	};

	/* initializes mt[N] with a seed */
	/* origin name init_genrand */
	MersenneTwister.prototype.init_seed = function(s) {
		this.mt[0] = s >>> 0;
		for (this.mti=1; this.mti<this.N; this.mti++) {
			var s = this.mt[this.mti-1] ^ (this.mt[this.mti-1] >>> 30);
			this.mt[this.mti] = (((((s & 0xffff0000) >>> 16) * 1812433253) << 16) + (s & 0x0000ffff) * 1812433253)
			+ this.mti;
			/* See Knuth TAOCP Vol2. 3rd Ed. P.106 for multiplier. */
			/* In the previous versions, MSBs of the seed affect   */
			/* only MSBs of the array mt[].                        */
			/* 2002/01/09 modified by Makoto Matsumoto             */
			this.mt[this.mti] >>>= 0;
			/* for >32 bit machines */
		}
	};

	/* initialize by an array with array-length */
	/* init_key is the array for initializing keys */
	/* key_length is its length */
	/* slight change for C++, 2004/2/26 */
	MersenneTwister.prototype.init_by_array = function(init_key, key_length) {
		var i, j, k;
		this.init_seed(19650218);
		i=1; j=0;
		k = (this.N>key_length ? this.N : key_length);
		for (; k; k--) {
			var s = this.mt[i-1] ^ (this.mt[i-1] >>> 30);
			this.mt[i] = (this.mt[i] ^ (((((s & 0xffff0000) >>> 16) * 1664525) << 16) + ((s & 0x0000ffff) * 1664525)))
			+ init_key[j] + j; /* non linear */
			this.mt[i] >>>= 0; /* for WORDSIZE > 32 machines */
			i++; j++;
			if (i>=this.N) { this.mt[0] = this.mt[this.N-1]; i=1; }
			if (j>=key_length) j=0;
		}
		for (k=this.N-1; k; k--) {
			var s = this.mt[i-1] ^ (this.mt[i-1] >>> 30);
			this.mt[i] = (this.mt[i] ^ (((((s & 0xffff0000) >>> 16) * 1566083941) << 16) + (s & 0x0000ffff) * 1566083941))
			- i; /* non linear */
			this.mt[i] >>>= 0; /* for WORDSIZE > 32 machines */
			i++;
			if (i>=this.N) { this.mt[0] = this.mt[this.N-1]; i=1; }
		}

		this.mt[0] = 0x80000000; /* MSB is 1; assuring non-zero initial array */
	};

	/* generates a random number on [0,0xffffffff]-interval */
	/* origin name genrand_int32 */
	MersenneTwister.prototype.random_int = function() {
		var y;
		var mag01 = new Array(0x0, this.MATRIX_A);
		/* mag01[x] = x * MATRIX_A  for x=0,1 */

		if (this.mti >= this.N) { /* generate N words at one time */
			var kk;

			if (this.mti == this.N+1)  /* if init_seed() has not been called, */
				this.init_seed(5489);  /* a default initial seed is used */

			for (kk=0;kk<this.N-this.M;kk++) {
				y = (this.mt[kk]&this.UPPER_MASK)|(this.mt[kk+1]&this.LOWER_MASK);
				this.mt[kk] = this.mt[kk+this.M] ^ (y >>> 1) ^ mag01[y & 0x1];
			}
			for (;kk<this.N-1;kk++) {
				y = (this.mt[kk]&this.UPPER_MASK)|(this.mt[kk+1]&this.LOWER_MASK);
				this.mt[kk] = this.mt[kk+(this.M-this.N)] ^ (y >>> 1) ^ mag01[y & 0x1];
			}
			y = (this.mt[this.N-1]&this.UPPER_MASK)|(this.mt[0]&this.LOWER_MASK);
			this.mt[this.N-1] = this.mt[this.M-1] ^ (y >>> 1) ^ mag01[y & 0x1];

			this.mti = 0;
		}

		y = this.mt[this.mti++];

		/* Tempering */
		y ^= (y >>> 11);
		y ^= (y << 7) & 0x9d2c5680;
		y ^= (y << 15) & 0xefc60000;
		y ^= (y >>> 18);

		return y >>> 0;
	};

	/* generates a random number on [0,0x7fffffff]-interval */
	/* origin name genrand_int31 */
	MersenneTwister.prototype.random_int31 = function() {
		return (this.random_int()>>>1);
	};

	/* generates a random number on [0,1]-real-interval */
	/* origin name genrand_real1 */
	MersenneTwister.prototype.random_incl = function() {
		return this.random_int()*(1.0/4294967295.0);
		/* divided by 2^32-1 */
	};

	/* generates a random number on [0,1)-real-interval */
	MersenneTwister.prototype.random = function() {
		return this.random_int()*(1.0/4294967296.0);
		/* divided by 2^32 */
	};

	/* generates a random number on (0,1)-real-interval */
	/* origin name genrand_real3 */
	MersenneTwister.prototype.random_excl = function() {
		return (this.random_int() + 0.5)*(1.0/4294967296.0);
		/* divided by 2^32 */
	};

	/* generates a random number on [0,1) with 53-bit resolution*/
	/* origin name genrand_res53 */
	MersenneTwister.prototype.random_long = function() {
		var a=this.random_int()>>>5, b=this.random_int()>>>6;
		return (a*67108864.0+b)*(1.0/9007199254740992.0);
	};

	/* These real versions are due to Isaku Wada, 2002/01/09 added */

	var mersenneTwister = MersenneTwister;

	/** This base class defines a general grid and provides grid methods that do
	 * not depend on the coordinate system used. This class is never used on its
	 * own, as it does not yet contain methods for neighborhood computation etc
	 * (which depend on the coordinate system).
	 * Coordinate system-dependent methods are implemented in extensions of the
	 * Grid class, for example Grid2D and Grid3D. */
	class Grid {

		/** The grid constructor.
		 * @param {GridSize} field_size array of field size in each dimension.
		 * E.g. [100,200] for a grid that is 100 pixels wide and 200 pixels high.
		 * Entries must be positive integer numbers.
		 * @param {boolean[]} [torus=[true,true,...]] - should the borders of the
		 * grid be linked, so that a cell moving out on the left reappears on the
		 * right? Should be an array specifying whether the torus holds in each
		 * dimension; eg [true,false] for a torus in x but not y dimension.
		 * Warning: setting the torus to false can give artifacts if not done
		 * properly, see {@link Grid#torus}.*/
		constructor( field_size, torus ){
		
			torus = torus || [];
		
			/** field_size array of field size in each dimension. E.g. [100,200]
			 * for a grid that is 100 pixels wide and 200 pixels high. Entries must
			 * be positive integer numbers.
			 * @type {GridSize}*/
			this.extents = field_size;
			
			/** Number of dimensions of the grid.
			 * @type {number}*/
			this.ndim = this.extents.length;
			
			
			if( torus.length === 0 ){
				for( let d = 0; d < this.ndim; d++ ){
					torus.push( true );
				}
			} else if ( torus.length !== this.ndim ){
				throw( "Torus should be specified for each dimension, or not " +
					"at all!" )
			}
			/** Should the borders of the grid be linked, so that a cell moving
			 * out on the left reappears on the right? Warning: setting to false
			 * can give artifacts if done incorrectly. If torus is set to false,
			 * artifacts arise because
			 * cells at the border have fewer neighbors. Cells will then stick to
			 * the grid borders where they have fewer neighbors to get adhesion and/or
			 * perimeter penalties from. You will need to specify how to handle the
			 * borders explicitly; see the examples/ folder for details on how to
			 * do this.
			 * Torus can be specified for
			 * each dimension separately.
			 * @type {boolean[]}*/
			this.torus = torus;

			/** Array with values for each pixel stored at the position of its
			 * 	{@link IndexCoordinate}. E.g. the value of pixel with coordinate
			 * 	i is stored as this._pixels[i]. This should be implemented in
			 * 	the grid subclass; see e.g. {@link Grid2D#_pixelArray}.
			 * 	Note that this array is accessed indirectly via the
			 * {@link _pixels} set- and get methods.
			 * 	@private
			 * 	*/
			this._pixelArray = undefined;
			this.datatype = undefined;
			
			/* These are used for rapid conversion between array and index
			coordinates, but not documented as they should not be used from outside.
			*/
			/** @ignore */
			this.X_BITS = 1+Math.floor( Math.log2( this.extents[0] - 1 ) );
			/** @ignore */
			this.Y_BITS = 1+Math.floor( Math.log2( this.extents[1] - 1 ) );
			/** @ignore */
			this.Y_MASK = (1 << this.Y_BITS)-1;
			
			/** Array coordinates to the middle pixel on the grid.
			 * @type {ArrayCoordinate}
			 * */
			this.midpoint = this.extents.map( i => Math.round((i-1)/2) );
		}

		/**
		 * Return the array this._pixelArray, which should be set in the grid
		 * subclass.
		 * @returns {Uint16Array|Float32Array}
		 * @private
		 */
		get _pixels(){
			if ( this._pixelArray !== undefined ){
				return this._pixelArray
			}
			throw("A private array this._pixelArray needs to be generated in every " +
				" Grid subclass! See its documentation for details.")
		}

		/**
		 * Set or update the private this._pixelArray.
		 * @param {Uint16Array|Float32Array} pixels - array of pixels to set.
		 * @private
		 */
		set _pixels( pixels ){
			//noinspection JSValidateTypes
			this._pixelArray = pixels;
		}

		/** Method to check if a given {@link ArrayCoordinate} falls within the
		 * bounds of this grid. Returns an error if this is not the case.
		 * @param {ArrayCoordinate} p - the coordinate to check.
		 */
		checkOnGrid( p ){
			for( let d = 0; d < p.length; d++ ){
				if( p[d] < 0 || p[d] >= this.extents[d] ){
					throw("You are trying to access a coordinate that does not seem" +
						"to lie on the grid! I am expecting values between 0 and " +
						"the grid dimension specified in field_size. (If this is your" +
						"own grid implementation and this assumption is not valid," +
						"please overwrite the liesOnGrid() function in your own" +
						"grid class).")
				}
			}
		}

		/** Method for conversion from an {@link ArrayCoordinate} to an
		 * {@link IndexCoordinate}.
		 * This method should be implemented in the subclass, see
		 * {@link Grid2D#p2i} for an example.
		 * @abstract
		 * @param {ArrayCoordinate} p - the coordinate of the pixel to convert
		 * @return {IndexCoordinate} the converted coordinate.
		 */
		//eslint-disable-next-line no-unused-vars
		p2i ( p ){
			throw( "A p2i method should be implemented in every Grid subclass!")
		}

		/** Method for conversion from an {@link IndexCoordinate} to an
		 * {@link ArrayCoordinate}.
		 * This method should be implemented in the subclass, see
		 * {@link Grid2D#i2p} for an example.
		 * @abstract
		 * @param {IndexCoordinate} i - the coordinate of the pixel to convert
		 * @return {ArrayCoordinate} the converted coordinate.
		 */
		//eslint-disable-next-line no-unused-vars
		i2p ( i ){
			throw( "An i2p method should be implemented in every Grid subclass!")
		}

		/** Method returning the (Moore) neighborhood of a pixel based on its
		 * {@link IndexCoordinate}.
		 * This method should be implemented in the subclass, see
		 * {@link Grid2D#neighi} for an example.
		 * @abstract
		 * @param {IndexCoordinate} i - the coordinate of the pixel to get neighbors
		 * for.
		 * @param {boolean[]} torus are borders of the grid linked so that a cell
		 * leaving on the right re-enters the grid on the left?
		 * @return {IndexCoordinate[]} an array of neighbors.
		 */
		//eslint-disable-next-line no-unused-vars
		neighi ( i, torus = this.torus ){
			throw( "A neighi method should be implemented in every Grid subclass!")
		}

		/** The neigh method returns the neighborhood of a pixel p. This function
		 * uses array coordinates as input and output arguments, but internally
		 * calls a method 'neighi' which computes neighborhoods using index-
		 * coordinates. Since neighborhoods depend on the coordinate system, the
		 * 'neighi' method is defined in the extension class for that specific
		 * coordinate system.
		 * @param {ArrayCoordinate} p array coordinates of a specific pixel
		 * @param {boolean[]} torus are borders of the grid linked so that a cell
		 * leaving on the right re-enters the grid on the left?
		 * @return {ArrayCoordinate[]} an array of neighbors of pixel p, where each
		 * element contains the array coordinates of the neighbor in question.
		 * */
		neigh(p, torus = this.torus){
			let g = this;
			return g.neighi( this.p2i(p), torus ).map( function(i){
				return g.i2p(i)
			} )
		}

		/** Check if a value is valid on this type of grid.
		 * This function forbids trying to set forbidden (negative/float) values
		 * on an integer grid, which could cause bugs if the setpix(i) methods try
		 * to set such a value unnoticed.
		 * @private
		 * @param {number} t - the value that would be stored on the grid.
		 * @param {number} [tol=1e-6] - numeric tolerance for comparing a number
		 * with its rounded version, to check if it is integer (e.g. setting
		 * a value 1.5 on an integer grid would throw an error, but setting
		 * 1.000000000001 would not if the tolerance is 1e-6.
		 * @return {void} - return without problem or throw an error when an
		 * incorrect value is set.
		 * */
		_isValidValue ( t, tol = 1e-6 ) {
			if( this.datatype === "Uint16" ){
				if( t < 0 || Math.abs( t - Math.round(t) ) > tol  ){
					throw( "You cannot set a negative or floating point number to a Uint16 grid!" )
				}
			}
		}

		/** The setpix method changes the value of a pixel p on the grid to t.
		 * @param {ArrayCoordinate} p array coordinates of the pixel to change the
		 * value of
		 * @param {number} t the value to assign to this pixel. This can
		 * be integers or floating point numbers, depending on the grid subclass
		 * used (see eg Grid2D). */
		setpix( p, t ){
			this._isValidValue(t);
			this._pixels[this.p2i(p)] = t;
		}

		/** The setpixi method changes the value of a pixel i on the grid to t.
		 * @param {IndexCoordinate} i index coordinates of the pixel to change the
		 * value of
		 * @param {number} t the value to assign to this pixel. This can be integers
		 * or floating point numbers, depending on the grid subclass used
		 * (see eg Grid2D).
		 * */
		setpixi( i, t ){
			this._isValidValue(t);
			this._pixels[i] = t;
		}

		/** The pixt method finds the current value of a pixel p on the grid.
		 * @param {ArrayCoordinate} p array coordinates of the pixel to find the
		 * value of
		 * @return {number} t the value of p on the grid. This can be integers or
		 * floating point numbers, depending on the grid subclass used (see eg
		 * Grid2D).
		 */
		pixt( p ){
			return this._pixels[this.p2i(p)]
		}

		/** The pixti method finds the current value of a pixel i on the grid.
		 * @param {IndexCoordinate} i index coordinates of the pixel to find the
		 * value of
		 * @return {number} t the value of i on the grid. This can be integers or
		 * floating point numbers, depending on the grid subclass used (see eg
		 * Grid2D).
		*/
		pixti( i ){
			return this._pixels[i]
		}

		/** A pixel on the grid.
		 * @typedef {Object[]} Pixel
		 * @property {ArrayCoordinate} Pixel[0] - pixel coordinate
		 * @property {number} Pixel[1] - pixel value
		 * */
		
		/* /** A pixel on the grid.
		 * @typedef {Object[]} iPixel
		 * @property {IndexCoordinate} Pixel[0] - pixel coordinate
		 * @property {number} Pixel[1] - pixel value
		 * */

		/** This iterator returns locations and values of all non-zero pixels.
		 * This method isn't actually called because the subclasses implement
		 * it themselves due to efficiency reasons. It serves as a template to
		 * document the functionality.
		 * @abstract
		 * @return {Pixel} for each pixel, return an array [p,v] where p are
		 * the pixel's array coordinates on the grid, and v its value.*/
		//eslint-disable-next-line require-yield
		* pixels() {
			//noinspection JSValidateTypes
			throw("Iterator 'pixels' not implemented!")

			/*
			// example code:
			for( let i of this.pixelsi() ){
				if( this._pixels[i] > 0 ){
					yield [this.i2p(i),this._pixels[i]]
				}
			}*/
		}

		/** This iterator returns locations all pixels including background.
		 * This method isn't actually called because the subclasses implement
		 * it themselves due to efficiency reasons. It serves as a template to
		 * document the functionality.
		 * @abstract
		 * @return {IndexCoordinate} for each pixel, because this method should be
		 * implemented in a grid subclass.
		 * */
		//eslint-disable-next-line require-yield
		* pixelsi() {
			throw("Iterator 'pixelsi' not implemented!")
			//yield undefined
		}

		/** This method pre-allocates an array of the correct datatype to make
		 * a copy of the current pixel values. Values are not actually copied yet.
		 * @return {Uint16Array|Float32Array} an array with an element for each
		 * pixel. The datatype is determined by the datatype of this._pixels
		 * (implemented in the subclass), which can be either Uint16Array or
		 * Float32Array. */
		pixelsBuffer() {

			/** For storing a copy of all pixel values; eg for synchronous updating
			 * of some sort.
			 * @type {Uint16Array|Float32Array}*/
			this._pixelsbuffer = new Uint16Array(this._pixels.length);
		
			if( this._pixels instanceof Uint16Array ){
				this._pixelsbuffer = new Uint16Array(this._pixels.length);
			} else if( this._pixels instanceof Float32Array ){
				this._pixelsbuffer = new Float32Array(this._pixels.length);
			} else {
				throw("unsupported datatype: " + (typeof this._pixels))
			}
		}

		/** Template method to compute the gradient at location i on the grid
		 * (location given in index coordinates).
		 * This method throws an error, which is overwritten when a subclass
		 * implements a gradienti method.
		 * @param {IndexCoordinate} i index coordinate of a pixel to compute the
		 * gradient at.
		 * @return {number[]} the gradient
		 * @see https://en.wikipedia.org/wiki/Gradient*/
		//eslint-disable-next-line no-unused-vars
		gradienti( i ){
			throw("method 'gradienti' not implemented! ")
		}

		/** Method to compute the gradient at location p on the grid (location
		 * given in array coordinates). It internally calls the gradienti
		 * method using index coordinates, which should be implemented in the grid
		 * subclass.
		 * @param {ArrayCoordinate} p array coordinates of a pixel p to compute
		 * the gradient at
		 * @return {number} the gradient at position p.
		 * @see https://en.wikipedia.org/wiki/Gradient*/
		gradient( p ){
			//noinspection JSValidateTypes
			return this.gradienti( this.p2i( p ) )
		}

		/** Method to compute the laplacian at location p on the grid
		 * (location given in array coordinates). It internally calls the laplaciani
		 * method that does the same but uses index coordinates.
		 * @param {ArrayCoordinate} p array coordinates of a pixel p to compute the
		 * laplacian at
		 * @return {number} the laplacian at position p.
		 * @see https://en.wikipedia.org/wiki/Laplace_operator#Coordinate_expressions */
		laplacian( p ){
			return this.laplaciani( this.p2i( p ) )
		}

		/**
		 * A method to compute the Neumann neighborhood should be implemented in the
		 * Grid subclass if the laplacian (see below) is used.
		 * This mock function ensures that an error is thrown when there is no
		 * method called neighNeumanni in the grid subclass.
		 * @abstract
		 * @see https://en.wikipedia.org/wiki/Von_Neumann_neighborhood
		 * @param {IndexCoordinate} i - location of the pixel to get neighbors of.
		 * @param {boolean[]} [torus=[true,true]] - does the grid have linked
		 * borders? Defaults to the setting on this grid, see {@link torus}
		 * @return {IndexCoordinate[]} - an array of coordinates for all the neighbors of i.
		 */
		//eslint-disable-next-line no-unused-vars,require-yield
		* neighNeumanni ( i, torus = this.torus ){
			throw( "Trying to call the method neighNeumanni, but you haven't " +
				"implemented this method in the Grid subclass you are using!")
		}

		/** Method to compute the laplacian at location i on the grid (location
		 * given in index coordinates). It internally uses the neighNeumanni method
		 * to compute a Neumann neighborhood, which should be implemented in the
		 * grid subclass. It then uses the finite difference method (see link) with
		 * h = 1.
		 * @param {IndexCoordinate} i index coordinates of a pixel to compute the
		 * laplacian at
		 * @return {number} the laplacian at position p.
		 * @see https://en.wikipedia.org/wiki/Laplace_operator#Coordinate_expressions
		 * @see https://en.wikipedia.org/wiki/Discrete_Laplace_operator#Finite_differences
		 * */
		laplaciani( i ){
			let L = 0, n = 0;

			// For now: forbid computing a laplacian on an integer grid as it makes
			// no sense and could happen by accident if you forget to specify the
			// datatype.
			// If this is too strict, we can set an option to overrule this error.
			// This way you still get to see it if you try this by accident.
			if( this.datatype === "Uint16" ){
				throw("Diffusion/laplacian methods do not work on a Uint16 grid! " +
					"Consider setting datatype='Float32'.")
			}

			//noinspection JSUnresolvedFunction
			for( let x of this.neighNeumanni(i) ){
				L += this.pixti( x ); n ++;
			} 
			return L - n * this.pixti( i )
		}

		/** Perform a diffusion step on the grid, updating the values of all pixels
		 * according to Fick's second law of diffusion.
		 * @param {number} D diffusion coefficient
		 * @see https://en.wikipedia.org/wiki/Diffusion#Fick's_law_and_equations
		 * @see https://en.wikipedia.org/wiki/Discrete_Laplace_operator#Mesh_Laplacians
		 * */
		diffusion( D ){
			// For synchronous updating of the grid: compute updated values in a copy
			// of the current pixels
			if( ! this._pixelsbuffer ) this.pixelsBuffer();
			for( let i of this.pixelsi() ){
				// Diffusion: new value is current value + change.
				// the change is given by the diffusion coefficient D times the laplacian.
				this._pixelsbuffer[i] = this.pixti( i ) + D * this.laplaciani( i );
			}
			// swap the copy and the original
			[this._pixelsbuffer, this._pixels] = [this._pixels, this._pixelsbuffer];
		}

		
		/** Function that updates a gridpoint depending on its current value and
		 * that of its neighbors.
		 * @typedef {function} updatePixelFunction
		 * @param {ArrayCoordinate} p - pixel to update
		 * @param {ArrayCoordinate[]} neighbors - coordinates of neighbors of p
		 * @return {number} value - the updated value, based on the current value
		 * of p and its neighbors.
		 */

		/** Apply a function to all pixel values on the grid. It acts on
		 * this._pixels, which is implemented in the grid subclass.
		 * @param {updatePixelFunction} f - the function to apply to each pixel. */
		applyLocally( f ){
			if( ! this._pixelsbuffer ) this.pixelsBuffer();
			for( let i of this.pixelsi() ){
				let p = this.i2p(i);
				this._pixelsbuffer[i] = f( p, this.neigh(p) ); 
			}
			[this._pixelsbuffer, this._pixels] = [this._pixels, this._pixelsbuffer];
		}
		
		/** Multiply all pixel values on the grid with a constant factor r.
		 * This method acts on this._pixels, which is implemented in the grid
		 * subclass.
		 * @param {number} r the multiplication factor. */
		multiplyBy( r ){
			for( let i of this.pixelsi() ){
				this._pixels[i] *= r;
			}
		}

	}

	/** A class containing (mostly static) utility functions for dealing with 2D
	 *  grids. Extends the Grid class but implements functions specific to the 2D
	 *  grid (such as neighborhoods).
	 *
	 *  @example <caption>Diffusion on a 2D chemokine grid</caption>
	 *  let CPM = require("path/to/build")
	 *
	 *  // Make a grid with a chemokine, add some chemokine at the middle pixel
	 * let chemoGrid = new CPM.Grid2D( [100,100], [true,true], "Float32" )
	 * chemoGrid.setpix( [50,50], 100 )
	 *
	 * // Measure chemokine at different spots before and after diffusion
	 * let p1 = [50,50], p2 = [50,51]
	 * console.log( "p1: " + chemoGrid.pixt( p1 ) + ", p2: " + chemoGrid.pixt(p2) )
	 * chemoGrid.diffusion( 0.001 )
	 * console.log( "p1: " + chemoGrid.pixt( p1 ) + ", p2: " + chemoGrid.pixt(p2) )
	 * chemoGrid.multiplyBy( 0.9 )	 // decay of the chemokine
	 * console.log( "p1: " + chemoGrid.pixt( p1 ) + ", p2: " + chemoGrid.pixt(p2) )
	 *
	 * @example <caption>Neighborhoods on a 2D grid</caption>
	 *
	 * let CPM = require("path/to/build")
	 *
	 * // Make a grid with a torus, and find the neighborhood of the upper left pixel [0,0]
	 * let grid = new CPM.Grid2D( [100,100], [true,true] )
	 * console.log( grid.neigh( [0,0] ) ) // returns coordinates of 8 neighbors
	 *
	 * // Now try a grid without torus; the corner now has fewer neighbors.
	 * let grid2 = new CPM.Grid2D( [100,100], [false,false] )
	 * console.log( grid2.neigh( [0,0] ) ) // returns only 3 neighbors
	 *
	 * // Or try a Neumann neighborhood using the iterator
	 * for( let i of grid.neighNeumanni( 0 ) ){
	 * 	console.log( grid.i2p(i).join(" ") )
	 * }
	 */

	class Grid2D extends Grid {

		/** Constructor of the Grid2D object.
		 * @param {GridSize} extents - the size of the grid in each dimension
		 * @param {boolean[]} [torus=[true,true]] - should the borders of the grid
		 * be linked, so that a cell moving out on the left reappears on the right?
		 * Should be an array specifying whether the torus holds in each dimension;
		 * eg [true,false] for a torus in x but not y dimension.
		 * Warning: setting the torus to false can give artifacts if not done
		 * properly, see {@link Grid#torus}.
		 * @param {string} [datatype="Uint16"] - What datatype are the values
		 * associated with each pixel on the grid? Choose from "Uint16" or
		 * "Float32". */
		constructor( extents, torus=[true,true], datatype="Uint16" ){
			super( extents, torus );
			
			/** @ignore */
			this.X_STEP = 1 << this.Y_BITS; // for neighborhoods based on pixel index
			/** @ignore */
			this.Y_MASK = this.X_STEP-1;
			// Check that the grid size is not too big to store pixel ID in 32-bit
			// number, and allow fast conversion of coordinates to unique
			// ID numbers.
			if( this.X_BITS + this.Y_BITS > 32 ){
				throw("Field size too large -- field cannot be represented as 32-bit number")
			}

			this.datatype = datatype;

			// Attributes per pixel: CellId of the current pixel.
			if( datatype === "Uint16" ){
				/** Array with values for each pixel stored at the position of its
				 * {@link IndexCoordinate}. E.g. the value of pixel with coordinate
				 * i is stored as this._pixelArray[i].
				 * 	Note that this array is accessed indirectly via the
				 * {@link _pixels} set- and get methods.
				 * @private
				 * @type {Uint16Array|Float32Array} */
				this._pixelArray = new Uint16Array(this.p2i(this.extents));
			} else if( datatype === "Float32" ){
				this._pixelArray = new Float32Array(this.p2i(this.extents));
			} else {
				throw("unsupported datatype: " + datatype)
			}
		}

		/** This iterator returns locations and values of all pixels.
		 * Whereas the {@link pixels} generator yields only non-background pixels
		 * and specifies both their {@link ArrayCoordinate} and value, this
		 * generator yields all pixels by {@link IndexCoordinate} and does not
		 * report value.
		 *
		 * @return {IndexCoordinate} for each pixel on the grid (including
		 * background pixels).
		 *
		 * @example
		 * let CPM = require( "path/to/build" )
		 * // make a grid and set some values
		 * let grid = new CPM.Grid2D( [100,100], [true,true] )
		 * grid.setpixi( 0, 1 )
		 * grid.setpixi( 1, 5 )
		 *
		 * // iterator
		 * for( let i of grid.pixelsi() ){
		 * 	console.log( i )
		 * }
		 */
		* pixelsi() {
			let ii = 0, c = 0;
			for( let i = 0 ; i < this.extents[0] ; i ++ ){
				for( let j = 0 ; j < this.extents[1] ; j ++ ){
					yield ii;
					ii ++;
				}
				c += this.X_STEP;
				ii = c;
			}
		}
		
		/** This iterator returns locations and values of all non-zero pixels.
		 * @return {Pixel} for each pixel, return an array [p,v] where p are
		 * the pixel's array coordinates on the grid, and v its value.
		 *
		 * @example
		 * let CPM = require( "path/to/build" )
		 * // make a grid and set some values
		 * let grid = new CPM.Grid2D( [100,100], [true,true] )
		 * grid.setpix( [0,0], 1 )
		 * grid.setpix( [0,1], 5 )
		 *
		 * // iterator
		 * for( let p of grid.pixels() ){
		 * 	console.log( p )
		 * }
		 */
		* pixels() {
			let ii = 0, c = 0;
			// Loop over coordinates [i,j] on the grid
			// For each pixel with cellId != 0 (so non-background pixels),
			// return an array with in the first element the pixel 
			// coordinates p = [i,j], and in the second element the cellId of this pixel.
			for( let i = 0 ; i < this.extents[0] ; i ++ ){
				for( let j = 0 ; j < this.extents[1] ; j ++ ){

					//noinspection JSUnresolvedVariable
					let pixels = this._pixels;
					if( pixels[ii] > 0 ){ //check non-background
						yield [[i,j], pixels[ii]];
					}
					ii ++;
				}
				c += this.X_STEP;
				ii = c;
			}
		}

		/**	Simple method for neighborhood computation. This method is not
		 * actually used in the framework, except to test the less intuitive
		 * (but faster) neighi method that works directly on index coordinates
		 * rather than first converting to array positions (which is expensive
		 * as  neighborhoods are computed very often in the CPM).
		 * @ignore
		 * @param {IndexCoordinate} i - location of the pixel to get neighbors of.
		 * @return {IndexCoordinate[]} - an array of coordinates for all the
		 * neighbors of i.
		*/
		neighiSimple( i ){
			let p = this.i2p(i);
			let xx = [];
			for( let d = 0 ; d <= 1 ; d ++ ){
				if( p[d] === 0 ){
					if( this.torus[d] ){
						xx[d] = [p[d],this.extents[d]-1,p[d]+1];
					} else {
						xx[d] = [p[d],p[d]+1];
					}
				} else if( p[d] === this.extents[d]-1 ){
					if( this.torus[d] ){
						xx[d] = [p[d],p[d]-1,0];
					} else {
						xx[d] = [p[d],p[d]-1];
					}
				} else {
					xx[d] = [p[d],p[d]-1,p[d]+1];
				}
			}

			let r = [], first=true;
			for( let x of xx[0] ){
				for( let y of xx[1] ){
					if( first ){
						first = false; 
					} else {
						r.push( this.p2i( [x,y] ) );
					}
				}
			}
			return r
		}
		
		/**	Return array of {@link IndexCoordinate} of von Neumann-4 neighbor
		 * pixels of the pixel at coordinate i. This function takes the 2D
		 * Neumann-4 neighborhood, excluding the pixel itself.
		 * @see https://en.wikipedia.org/wiki/Von_Neumann_neighborhood
		 * @param {IndexCoordinate} i - location of the pixel to get neighbors of.
		 * @param {boolean[]} [torus=[true,true]] - does the grid have linked
		 * borders? Defaults to the setting on this grid, see {@link torus}
		 * @return {IndexCoordinate[]} - an array of coordinates for all the
		 * neighbors of i.
		*/
		* neighNeumanni( i, torus = this.torus ){
			// normal computation of neighbor indices (top left-middle-right, 
			// left, right, bottom left-middle-right)
			let t = i-1, l = i-this.X_STEP, r = i+this.X_STEP, b = i+1;
			
			// if pixel is part of one of the borders, adjust the 
			// indices accordingly
			// if torus is false, return NaN for all neighbors that cross
			// the border.

			// left border
			if( i < this.extents[1] ){
				if( torus[0] ){
					l += this.extents[0] * this.X_STEP;
					yield l;
				} 
			} else {
				yield l;
			}
			// right border
			if( i >= this.X_STEP*( this.extents[0] - 1 ) ){
				if( torus[0] ){
					r -= this.extents[0] * this.X_STEP;
					yield r;
				}
			} else {
				yield r;
			}
			// top border
			if( i % this.X_STEP === 0 ){
				if( torus[1] ){
					t += this.extents[1];
					yield t;
				} 
			} else {
				yield t;
			}
			// bottom border
			if( (i+1-this.extents[1]) % this.X_STEP === 0 ){
				if( torus[1] ){
					b -= this.extents[1];
					yield b;
				} 
			} else {
				yield b;
			}
		}
		/**	Return array of {@link IndexCoordinate} of Moore-8 neighbor pixels of
		 * the pixel at coordinate i. This function takes the 2D Moore-8
		 * neighborhood, excluding the pixel itself.
		 * @see https://en.wikipedia.org/wiki/Moore_neighborhood
		 * @param {IndexCoordinate} i - location of the pixel to get neighbors of.
		 * @param {boolean[]} [torus] - does the grid have linked borders? Defaults
		 * to the setting on this grid, see {@link torus}
		 * @return {IndexCoordinate[]} - an array of coordinates for all the
		 * neighbors of i.
		*/
		neighi( i, torus = this.torus ){	
			// normal computation of neighbor indices (top left-middle-right, 
			// left, right, bottom left-middle-right)
			let tl, tm, tr, l, r, bl, bm, br;
			
			tl = i-1-this.X_STEP; tm = i-1; tr = i-1+this.X_STEP;
			l = i-this.X_STEP; r = i+this.X_STEP;
			bl = i+1-this.X_STEP; bm = i+1; br = i+1+this.X_STEP;
			
			// if pixel is part of one of the borders, adjust the 
			// indices accordingly
			let add = NaN; // if torus is false, return NaN for all neighbors that cross
			// the border.
			// 
			// left border
			if( i < this.extents[1] ){
				if( torus[0] ){
					add = this.extents[0] * this.X_STEP;
				}
				tl += add; l += add; bl += add; 	
			}
			
			// right border
			if( i >= this.X_STEP*( this.extents[0] - 1 ) ){
				if( torus[0] ){
					add = -this.extents[0] * this.X_STEP;
				}
				tr += add; r += add; br += add;
			}

			add = NaN;
			// top border
			if( i % this.X_STEP === 0 ){
				if( torus[1] ){
					add = this.extents[1];
				}
				tl += add; tm += add; tr += add;	
			}
			
			// bottom border
			if( (i+1-this.extents[1]) % this.X_STEP === 0 ){
				if( torus[1] ){
					add = -this.extents[1];
				}
				bl += add; bm += add; br += add;
			}
			if( !(torus[0]&&torus[1]) ){
				return [ tl, l, bl, tm, bm, tr, r, br ].filter( isFinite )
			} else {
				return [ tl, l, bl, tm, bm, tr, r, br ]
			}
		}
		
		/** Method for conversion from an {@link ArrayCoordinate} to an
		 * {@link IndexCoordinate}.
		 *
		 * See also {@link Grid2D#i2p} for the backward conversion.
		 *
		 * @param {ArrayCoordinate} p - the coordinate of the pixel to convert
		 * @return {IndexCoordinate} the converted coordinate.
		 *
		 * @example
		 * let grid = new CPM.Grid2D( [100,100], [true,true] )
		 * let p = grid.i2p( 5 )
		 * console.log( p )
		 * console.log( grid.p2i( p ))
		 */
		p2i ( p ){
			// using bitwise operators for speed.
			return ( p[0] << this.Y_BITS ) + p[1]
		}

		/** Method for conversion from an {@link IndexCoordinate} to an
		 * {@link ArrayCoordinate}.
		 * See also {@link Grid2D#p2i} for the backward conversion.
		 *
		 * @param {IndexCoordinate} i - the coordinate of the pixel to convert
		 * @return {ArrayCoordinate} the converted coordinate.
		 *
		 * @example
		 * let grid = new CPM.Grid2D( [100,100], [true,true] )
		 * let p = grid.i2p( 5 )
		 * console.log( p )
		 * console.log( grid.p2i( p ))
		 */
		i2p ( i ){
			// using bitwise operators for speed.
			return [i >> this.Y_BITS, i & this.Y_MASK]
		}
		
		/** Method to compute the gradient at location i on the grid (location
		 * given as an {@link IndexCoordinate}).
		 * @param {IndexCoordinate} i - index coordinate of a pixel to compute the
		 * gradient at
		 * @return {number[]} the gradient at position i.
		 * @see https://en.wikipedia.org/wiki/Gradient */
		gradienti( i ){
			let t = i-1, b = i+1, l = i-this.X_STEP, r = i+this.X_STEP,
				torus = this.torus;
			//noinspection JSUnresolvedVariable
			const pixels = this._pixels;
			
			let dx;
			if( i < this.extents[1] ){ // left border
				if( torus[0] ){
					l += this.extents[0] * this.X_STEP;
					dx = ((pixels[r]-pixels[i])+
						(pixels[i]-pixels[l]))/2;
				} else {
					dx = pixels[r]-pixels[i];
				}
			} else { 
				if( i >= this.X_STEP*( this.extents[0] - 1 ) ){ // right border
					if( torus[0] ){
						r -= this.extents[0] * this.X_STEP;
						dx = ((pixels[r]-pixels[i])+
							(pixels[i]-pixels[l]))/2;
					} else {
						dx = pixels[i]-pixels[l];
					}
				} else {
					dx = ((pixels[r]-pixels[i])+
						(pixels[i]-pixels[l]))/2;
				}
			}

			let dy;
			if( i % this.X_STEP === 0 ){ // top border
				if( torus[1] ){
					t += this.extents[1];
					dy = ((pixels[b]-pixels[i])+
						(pixels[i]-pixels[t]))/2;
				}	else {
					dy = pixels[b]-pixels[i];
				}
			} else { 
				if( (i+1-this.extents[1]) % this.X_STEP === 0 ){ // bottom border
					if( torus[1] ){
						b -= this.extents[1];
						dy = ((pixels[b]-pixels[i])+
							(pixels[i]-pixels[t]))/2;
					} else {
						dy = pixels[i]-pixels[t];
					}
				} else {
					dy = ((pixels[b]-pixels[i])+
						(pixels[i]-pixels[t]))/2;
				}
			}
			return [dx, dy]
		}
	}

	/** A class containing (mostly static) utility functions for dealing with 3D
	 * 	grids. Extends the Grid class but implements functions specific to the 3D
	 * 	grid (such as neighborhoods).
	 *
	 * @example <caption>Neighborhoods on a 3D grid</caption>
	 *
	 * let CPM = require("path/to/build")
	 *
	 * // Make a grid with a torus, and find the neighborhood of the upper left pixel [0,0,0]
	 * let grid = new CPM.Grid3D( [100,100,100], [true,true,true] )
	 * console.log( grid.neigh( [0,0,0] ) ) // returns coordinates of 26 neighbors (9+8+9)
	 *
	 * // Now try a grid without torus; the corner now has fewer neighbors.
	 * let grid2 = new CPM.Grid3D( [100,100,100], [false,false,false] )
	 * console.log( grid2.neigh( [0,0,0] ) ) // returns only 7 neighbors
	 *
	 * // Or try a torus in one dimension
	 * let grid2 = new CPM.Grid3D( [100,100,100], [true,false,false] )
	 * console.log( grid2.neigh( [0,0,0] ) )
	 */
	class Grid3D extends Grid {

		/** Constructor of the Grid3D object.
		 * @param {GridSize} extents - the size of the grid in each dimension
		 * @param {boolean[]} [torus = [true,true,true]] - should the borders of
		 * the grid be linked, so that a cell moving out on the left reappears on
		 * the right?
		 * Warning: setting the torus to false can give artifacts if not done
		 * properly, see {@link Grid#torus}.*/
		constructor( extents, torus = [true,true,true] ){
			super( extents, torus );
			// Check that the grid size is not too big to store pixel ID in 32-bit number,
			// and allow fast conversion of coordinates to unique ID numbers.
			/** @ignore */
			this.Z_BITS = 1+Math.floor( Math.log2( this.extents[2] - 1 ) );
			if( this.X_BITS + this.Y_BITS + this.Z_BITS > 32 ){
				throw("Field size too large -- field cannot be represented as 32-bit number")
			}
			/** @ignore */
			this.Z_MASK = (1 << this.Z_BITS)-1;
			/** @ignore */
			this.Z_STEP = 1;
			/** @ignore */
			this.Y_STEP = 1 << (this.Z_BITS);
			/** @ignore */
			this.X_STEP = 1 << (this.Z_BITS +this.Y_BITS);
			/** Array with values for each pixel stored at the position of its
			 * {@link IndexCoordinate}. E.g. the value of pixel with coordinate i
			 * is stored as this._pixelArray[i].
			 * 	Note that this array is accessed indirectly via the
			 * {@link _pixels} set- and get methods.
			 * @private
			 * @type {Uint16Array} */
			this._pixelArray = new Uint16Array(this.p2i(extents));
			this.datatype = "Uint16";
		}
		/** Method for conversion from an {@link ArrayCoordinate} to an
		 * {@link IndexCoordinate}.
		 *
		 * See also {@link Grid3D#i2p} for the backward conversion.
		 *
		 * @param {ArrayCoordinate} p - the coordinate of the pixel to convert
		 * @return {IndexCoordinate} the converted coordinate.
		 *
		 * @example
		 * let grid = new CPM.Grid3D( [100,100,100], [true,true,true] )
		 * let p = grid.i2p( 5 )
		 * console.log( p )
		 * console.log( grid.p2i( p ))
		 */
		p2i( p ){
			return ( p[0] << ( this.Z_BITS + this.Y_BITS ) ) + 
				( p[1] << this.Z_BITS ) + 
				p[2]
		}
		/** Method for conversion from an {@link IndexCoordinate} to an
		 * {@link ArrayCoordinate}.
		 *
		 * See also {@link Grid3D#p2i} for the backward conversion.
		 *
		 * @param {IndexCoordinate} i - the coordinate of the pixel to convert
		 * @return {ArrayCoordinate} the converted coordinate.
		 *
		 * @example
		 * let grid = new CPM.Grid3D( [100,100,100], [true,true,true] )
		 * let p = grid.i2p( 5 )
		 * console.log( p )
		 * console.log( grid.p2i( p ))
		 */
		i2p( i ){
			return [i >> (this.Y_BITS + this.Z_BITS), 
				( i >> this.Z_BITS ) & this.Y_MASK, i & this.Z_MASK ]
		}
		
		/** This iterator returns locations and values of all non-zero pixels.
		 * Whereas the {@link pixels} generator yields only non-background pixels
		 * and specifies both their {@link ArrayCoordinate} and value, this
		 * generator yields all pixels by {@link IndexCoordinate} and does not
		 * report value.
		 *
		 * @return {IndexCoordinate} for each pixel, return its
		 * {@link IndexCoordinate}.
		 *
		 *
		 * @example
		 * let CPM = require( "path/to/build" )
		 * // make a grid and set some values
		 * let grid = new CPM.Grid3D( [100,100,100], [true,true,true] )
		 * grid.setpixi( 0, 1 )
		 * grid.setpixi( 1, 5 )
		 *
		 * // iterator
		 * for( let i of grid.pixelsi() ){
		 * 	console.log( i )
		 * }
		 */
		* pixelsi() {
			let ii = 0, c = 0;
			for( let i = 0 ; i < this.extents[0] ; i ++ ){
				let d = 0;
				for( let j = 0 ; j < this.extents[1] ; j ++ ){
					for( let k = 0 ; k < this.extents[2] ; k ++ ){
						yield ii;
						ii++;
					}
					d += this.Y_STEP;
					ii = c + d;
				}
				c += this.X_STEP;
				ii = c;
			}
		}

		/** This iterator returns locations and values of all non-zero pixels.
		 * @return {Pixel} for each pixel, return an array [p,v] where p are
		 * the pixel's array coordinates on the grid, and v its value.
		 *
		 * @example
		 * let CPM = require( "path/to/build" )
		 * // make a grid and set some values
		 * let grid = new CPM.Grid3D( [100,100,100], [true,true,true] )
		 * grid.setpix( [0,0,0], 1 )
		 * grid.setpix( [0,0,1], 5 )
		 *
		 * // iterator
		 * for( let p of grid.pixels() ){
		 * 	console.log( p )
		 * }
		 */
		* pixels() {
			let ii = 0, c = 0;
			for( let i = 0 ; i < this.extents[0] ; i ++ ){
				let d = 0;
				for( let j = 0 ; j < this.extents[1] ; j ++ ){
					for( let k = 0 ; k < this.extents[2] ; k ++ ){
						//noinspection JSUnresolvedVariable
						let pixels = this._pixels;
						if( pixels[ii] > 0 ){
							yield [[i,j,k], pixels[ii]];
						}
						ii++;
					}
					d += this.Y_STEP;
					ii = c + d;
				}
				c += this.X_STEP;
				ii = c;
			}
		}

		/**	Return array of {@link IndexCoordinate} of the Moore neighbor pixels
		 * of the pixel at coordinate i. This function takes the 3D equivalent of
		 * the 2D Moore-8 neighborhood, excluding the pixel itself.
		 * @see https://en.wikipedia.org/wiki/Moore_neighborhood
		 * @param {IndexCoordinate} i - location of the pixel to get neighbors of.
		 * @param {boolean[]} [torus=[true,true,true]] - does the grid have linked
		 * borders? Defaults to the setting on this grid, see {@link torus}
		 * @return {IndexCoordinate[]} - an array of coordinates for all the
		 * neighbors of i.
		*/
		neighi( i, torus = this.torus ){
			let p = this.i2p(i);

			let xx = [];
			for( let d = 0 ; d <= 2 ; d ++ ){
				if( p[d] === 0 ){
					if( torus[d] ){
						xx[d] = [p[d],this.extents[d]-1,p[d]+1];
					} else {
						xx[d] = [p[d],p[d]+1];
					}
				} else if( p[d] === this.extents[d]-1 ){
					if( torus[d] ){
						xx[d] = [p[d],p[d]-1,0];
					} else {
						xx[d] = [p[d],p[d]-1];
					}
				} else {
					xx[d] = [p[d],p[d]-1,p[d]+1];
				}
			}
			let r = [], first=true;
			for( let x of xx[0] ){
				for( let y of xx[1] ){
					for( let z of xx[2] ){
						if( first ){
							first = false; 
						} else {
							r.push( this.p2i( [x,y,z] ) );
						}
					}
				}
			}
			return r
		}
	}

	/** Base class for grid-based models. This class is not used by itself; see
	{@link CPM} for a Cellular Potts Model and {@link CA} for a Cellular Automaton. 
	*/
	class GridBasedModel {

		/** The constructor of a GridBasedModel automatically attaches a grid of 
		class {@link Grid2D} or {@link Grid3D}, depending on the grid dimensions
		given in the 'extents' parameter. Configuration options for the model can
		be supplied in 'conf'.
		@param {GridSize} extents - the size of the grid of the model.
		@param {object} conf - configuration options. See below for its elements,
		but subclasses can have more.
		@param {boolean[]} [conf.torus=[true,true,...]] - should the grid have linked borders?
		@param {boolean} [conf.hexGrid=false] - should the grid be hexagonal? Grids
		 are square by default.
		@param {number} [seed] - seed for the random number generator. If left unspecified,
		a random number from the Math.random() generator is used to make one.  */
		constructor( extents, conf ){
			let seed = conf.seed || Math.floor(Math.random()*Number.MAX_SAFE_INTEGER);
			
			/** Attach a random number generation with a seed.
			@type {MersenneTwister}*/
			this.mt = new mersenneTwister( seed );
			if( !("torus" in conf) ){
				let torus = [];
				for( let d = 0; d < extents.length; d++ ){
					torus.push( true );
				}
				conf["torus"] = torus;
			}

			// Attributes based on input parameters
			/** Dimensionality of the grid 
			@type {number}*/
			this.ndim = extents.length; // grid dimensions (2 or 3)
			if( this.ndim !== 2 && this.ndim !== 3 ){
				throw("only 2D and 3D models are implemented!")
			}
			
			/** Input parameter settings; see the constructor of subclasses documentation.
			@type {object}*/
			this.conf = conf; // input parameter settings; see documentation.

			// Some functions/attributes depend on ndim:
			//this.hexGrid = conf.hexGrid || false

			if( this.ndim === 2 ){
				this.grid=new Grid2D(extents,conf.torus);
				//if( this.hexGrid ){
				//	/** The grid.
				//	@type {Grid2D|Grid3D|HexGrid2D}*/
				//	this.grid = new HexGrid2D(extents,conf.torus)
				//} else {
				//	this.grid = new Grid2D(extents,conf.torus)
				//}

			} else {
				/*if( this.hexGrid ){
					throw( "There is no 3D hexagonal grid!" )
				}*/
				this.grid = new Grid3D(extents,conf.torus);
			}
			// Pull up some things from the grid object so we don't have to access it
			// from the outside
			/** Midpoint of the grid.
			@type {ArrayCoordinate}*/
			this.midpoint = this.grid.midpoint;
			/** Size of the grid in object format.
			@type {object}*/
			this.field_size = this.grid.field_size;
			/** The {@link Grid2D#pixels} or {@link Grid3D#pixels} iterator function of the underlying grid.
			@type {function}*/
			this.pixels = this.grid.pixels.bind(this.grid);
			/** The {@link Grid#pixti} iterator function of the underlying grid.
			@type {function}*/
			this.pixti = this.grid.pixti.bind(this.grid);
			/** The {@link Grid2D#neighi} or {@link Grid3D#neighi} iterator function of the underlying grid.
			@type {function}*/
			this.neighi = this.grid.neighi.bind(this.grid);
			/** Size of the grid.
			@type {GridSize}*/
			this.extents = this.grid.extents;


			/** This tracks the volumes of all non-background cells on the grid.
			cellvolumes will be added with key = {@link CellId}, value = volume.
			@type{number[]}*/
			this.cellvolume = [];
			/** Tracks the elapsed time in MCS
			@type {number}*/
			this.time = 0;
			/** Objects of class {@link Stat} that have been computed on this model.
			@type {Stat}*/
			this.stats = [];
			/** Cached values of these stats. Object with stat name as key and its cached
			value as value. The cache must be cleared when the grid changes!
			@type {object} */
			this.stat_values = {};
		}

		/** Get the {@link CellKind} of the cell with {@link CellId} t. For this model, they
		are just the same.
		@param {CellId} t - id of the cell to get kind of.
		@return {CellKind} the cellkind. */
		cellKind( t ){
			return t 
		}

		/** Iterator for all the {@link CellId}s that are currently on the grid. 
		@return {CellId}*/
		* cellIDs() {
			yield* Object.keys( this.cellvolume );
		}

		/** Get neighbourhood of position p, using neighborhood functions of the underlying
		grid class.
		@param {ArrayCoordinate} p - coordinate of a pixel to get the neighborhood of.
		@param {boolean[]} [torus=[true,true,...]]  Does the grid have linked borders? If left unspecified,
		this is determined by this.conf.torus.*/
		neigh(p, torus=this.conf.torus){
			let g = this.grid;
			return g.neighi( g.p2i(p), torus ).map( function(i){ return g.i2p(i) } )
		}

		/** Get {@link CellId} of the pixel at coordinates p. 
		@param {ArrayCoordinate} p - pixel to get cellid of.
		@return {CellId} ID of the cell p belongs to.*/
		pixt( p ){
			return this.grid.pixti( this.grid.p2i(p) )
		}

		/** Change the pixel at position i into {@link CellId} t. 
			This standard implementation also keeps track of cell volumes
			for all nonzero cell IDs. Subclasses may want to do more, 
			such as also keeping track of perimeters or even centroids.
			In that case, this method needs to be overridden. 
			
			See also {@link setpix} for a method working with {@link ArrayCoordinate}s.
			
			@param {IndexCoordinate} i - coordinate of pixel to change.
			@param {CellId} t - cellid to change this pixel into.
			*/
		setpixi ( i, t ){		
			const t_old = this.grid.pixti(i);
			if( t_old > 0 ){
				// also update volume of the old cell
				this.cellvolume[t_old] --;
				// if this was the last pixel belonging to this cell, 
				// remove the cell altogether.
				if( this.cellvolume[t_old] == 0 ){
					delete this.cellvolume[t_old];
				}
			}
			// update volume of the new cell and cellid of the pixel.
			this.grid.setpixi( i, t );
			if( t > 0 ){
				if( !this.cellvolume[t] ){
					this.cellvolume[t] = 1;
				} else {
					this.cellvolume[t] ++;
				}
			}
		}

		/** Change the pixel at position p into {@link CellId} t. 
			This just calls the {@link setpixi} method internally.
			
			@param {ArrayCoordinate} p - coordinate of pixel to change.
			@param {CellId} t - cellid to change this pixel into.
			*/
		setpix ( p, t ){
			this.setpixi( this.grid.p2i(p), t );
		}

		/* ------------- MATH HELPER FUNCTIONS --------------- */
		/** Get a random number from the seeded number generator.
		@return {number} a random number between 0 and 1, uniformly sampled.*/
		random (){
			return this.mt.random()
		}

		/** Get a random integer number between incl_min and incl_max, uniformly sampled.
		@param {number} incl_min - lower end of the sampling range.
		@param {number} incl_max - upper end of the sampling range. 
		@return {number} the randomly sampled integer.*/
		ran (incl_min, incl_max) {
			return Math.floor(this.random() * (1.0 + incl_max - incl_min)) + incl_min
		}

		/** Compute a statistic on this model. If necessary, this produces an object
		of the right {@link Stat} subclass and runs the compute method. Stats are 
		cached because many stats use each other; this prevents that 'expensive' stats are
		computed twice. 
		@param {Stat} s - the stat to compute.
		@return {anything} - the value of the computed stat. This is often a {@link CellObject}
		or a {@link CellArrayObject}. 
		
		@example
		* let CPM = require( "path/to/dir")
		* let C = new CPM.CPM( [200,200], {T:20, torus:[false,false]} )
		* let gm = new CPM.GridManipulator( C )
		* gm.seedCell( 1 )
		* gm.seedCell( 1 )
		* C.getStat( Centroids )
		*/
		getStat( s ){
			/* Instantiate stats class if it doesn't exist yet and bind to this model */
			if( !(s.name in this.stats) ){
				let t = new s();
				this.stats[s.name] = t;
				t.model = this;
				
			}
			/* Cache stat value if it hasn't been done yet */
			if( !(s.name in this.stat_values) ){
				this.stat_values[s.name] = this.stats[s.name].compute();
			}
			/* Return cached value */
			return this.stat_values[s.name]
		}
		
		/** Update the grid in one timestep. This method is model-dependent and 
		must be implemented in the subclass.
		@abstract */
		timeStep (){
			throw("implemented in subclasses")
		}
	}

	// pass in RNG

	/** This class implements a data structure with constant-time insertion, deletion, and random
	    sampling. That's crucial for the CPM metropolis algorithm, which repeatedly needs to sample
	    pixels at cell borders. Elements in this set must be unique.*/
	class DiceSet{

		/** An object of class MersenneTwister. 
		@see https://www.npmjs.com/package/mersenne-twister
		@typedef {object} MersenneTwister
		*/

		/** The constructor of class DiceSet takes a MersenneTwister object as input, to allow
		seeding of the random number generator used for random sampling.
		@param {MersenneTwister} mt MersenneTwister object used for random numbers.*/
		constructor( mt ) {

			/** Object or hash map used to check in constant time whether a pixel is at the
			cell border. Keys are the actual values stored in the DiceSet, numbers are their
			location in the elements arrray.
			Currently (Mar 6, 2019), it seems that vanilla objects perform BETTER than ES6 maps,
			at least in nodejs. This is weird given that in vanilla objects, all keys are 
			converted to strings, which does not happen for Maps.
			@type {object}
			*/
			this.indices = {}; //new Map() // {}
			//this.indices = {}

			/** Use an array for constant time random sampling of pixels at the border of cells.
			@type {number[]} */
			this.elements = [];

			/** The number of elements currently present in the DiceSet. 
			@type {number}
			*/
			this.length = 0;

			/** @ignore */
			this.mt = mt;
		}

		/** Unique identifier of some element. This can be a number (integer) or a string,
		but it must uniquely identify one element in a set.
		@typedef {number|string} uniqueID*/

		/** Insert a new element. It is added as an index in the indices, and pushed
		to the end of the elements array.
		@param {uniqueID} v The element to add.
		*/
		insert( v ){
			if( this.indices[v] ){
				return
			}
			// Add element to both the hash map and the array.
			//this.indices.set( v, this.length )
			this.indices[v] = this.length;
		
			this.elements.push( v );
			this.length ++; 
		}

		/** Remove element v.
		@param {uniqueID} v The element to remove. 
		*/
		remove( v ){
			// Check whether element is present before it can be removed.
			if( !this.indices[v] ){
				return
			}
			/* The hash map gives the index in the array of the value to be removed.
			The value is removed directly from the hash map, but from the array we
			initially remove the last element, which we then substitute for the 
			element that should be removed.*/
			//const i = this.indices.get(v)
			const i = this.indices[v];

			//this.indices.delete(v)
			delete this.indices[v];

			const e = this.elements.pop();
			this.length --;
			if( e == v ){
				return
			}
			this.elements[i] = e;

			//this.indices.set(e,i)
			this.indices[e] = i;
		}
		/** Check if the DiceSet already contains element v. 
		@param {uniqueID} v The element to check presence of. 
		@return {boolean} true or false depending on whether the element is present or not.
		*/
		contains( v ){
			//return this.indices.has(v)
			return (v in this.indices)
		}
		
		/** Sample a random element from v.
		@return {uniqueID} the element sampled.
		*/
		sample(){
			return this.elements[Math.floor(this.mt.random()*this.length)]
		}
	}

	/** This base class defines a general CPM constraint and provides methods that do not 
	depend on the specific constraint used. This class is never used on its own, 
	as it does not yet contain the actual definition of a constraint (such as a deltaH method).

	In general, we distinguish between two types of constraint:

	- a {@link HardConstraint} is a hard rule that *must* be fulfilled in order for a copy
	attempt to succeed;
	- a {@link SoftConstraint} is an energy term in the Hamiltonian that can make a copy 
	attempt either more or less energetically favourable, but does not by itself determine
	whether a copy attempt will succeed. An unfavourable outcome may be outbalanced by 
	favourable energies from other terms, and even a copy attempt net unfavourable 
	energy (deltaH > 0) may succeed with a success chance P = exp(-DeltaH/T). 

	See the subclasses {@link SoftConstraint} and {@link HardConstraint} for details. Each
	implemented constraint is in turn a subclass of one of these two.
	*/
	class Constraint {

		/** This method is actually implemented in the subclass.
		@abstract
		*/
		get CONSTRAINT_TYPE() {
			throw("You need to implement the 'CONSTRAINT_TYPE' getter for this constraint!")
		}
		
		/** Get the parameters of this constraint from the conf object. 
		@return {object} conf - configuration settings for this constraint, containing the
		relevant parameters.
		*/
		get parameters(){
			return this.conf
		}
		/** The constructor of a constraint takes a configuration object.
		This method is usually overwritten by the actual constraint so that the entries
		of this object can be documented.
		@param {object} conf - configuration settings for this constraint, containing the
		relevant parameters.
		@abstract*/
		constructor( conf ){
			/** Configuration object for this constraint.
			@type {object}*/
			this.conf = conf;
		}
		/** This function attaches the relevant CPM to this constraint, so that information
		about this cpm can be requested from the constraint. 
		@todo Check why some constraints overwrite this? Because that disables the automatic
		usage of a confChecker() when it is implemented. 
		@param {CPM} C - the CPM to attach to this constraint.*/
		set CPM(C){
			/** CPM on which this constraint acts.
			@type {CPM}*/
			this.C = C;
			if( typeof this.confChecker === "function" ){
				this.confChecker();
			}
		}
		/** The optional confChecker method should verify that all the required conf parameters
		are actually present in the conf object and have the right format. It is implemented in
		the subclass that specifies the actual constraint.
		@abstract */
		confChecker( ){
		}

	}

	/** Extension of class {@link Constraint} used for a soft constraint. See description in
	 {@link Constraint} for details. This class is not used on its own but serves as a base
	 class for a soft constraint. */
	class SoftConstraint extends Constraint {

		/** Let the CPM know that this is a soft constraint, so return 'soft'. 
		@return {string} "soft"*/
		get CONSTRAINT_TYPE() {
			return "soft"
		}
		
		/** Soft constraints must have a deltaH method to compute the Hamiltonian. This method
		must be implemented in any SoftConstraint subclass before it works.
		@abstract
		 @param {IndexCoordinate} src_i - coordinate of the source pixel that tries to copy.
		 @param {IndexCoordinate} tgt_i - coordinate of the target pixel the source is trying
		 to copy into.
		 @param {CellId} src_type - cellid of the source pixel.
		 @param {CellId} tgt_type - cellid of the target pixel. 
		 @return {number} the change in Hamiltonian for this copy attempt and this constraint.*/ 
		// eslint-disable-next-line no-unused-vars
		deltaH( src_i, tgt_i, src_type, tgt_type ){
			throw("You need to implement the 'deltaH' method for this constraint!")
		}
	}

	/** Class for checking if parameters are present in a conf object and are of the expected
	structure and type. This is not meant for usage from outside the package, but you can
	use it when you are building your own constraints and want to specify the required parameters.

	@example
	* // Add this above your constraint class
	* import ParameterChecker from "./ParameterChecker.js"
	* 
	* // from inside the confChecker() function of your constraint:
	* let checker = new ParameterChecker( this.conf, this.C )
	* 
	* // Most parameters have a standard structure and value type;
	* // you can check them like this:
	* checker.confCheckParameter( "MY_PARAMETER", "KindMatrix", "Number" ) // see Adhesion
	* checker.confCheckParameter( "MY_PARAMETER_2", "KindArray", "NonNegative" )
	* 
	* // Or you can just check their presence and do a custom check:
	* checker.confCheckPresenceOf( "MY_WEIRD_PARAMETER" )
	* if( !myCondition ){
	* 	throw( "Some error because MY_WEIRD_PARAMETER does not fulfill myCondition!" )
	* }
	*/
	class ParameterChecker {

		/** The constructor of the ParameterChecker takes a configuration object.
		@param {object} conf - configuration settings as supplied to a constraint, containing the
		relevant parameters.
		@param {CPM} C - the attached CPM.
		*/
		constructor( conf, C ){
		
			/** The configuration object to check parameters in 
			@type {object}*/
			this.conf = conf;
			
			/** The attached CPM. This is used to check if parameter array lengths match
			the number of cellkinds
			@type {CPM}*/
			this.C = C;
		}
		
		/* ========= CHECKING PARAMETER PRESENCE ======== */

		/** Method to check if a parameter of a given name is present in the conf object 
		supplied to the constraint, and if it is defined. Throws an error if this is not the case.
		@param {string} name - the name of the parameter, which should be present as a key
		in the object.
		
		@example
		* let conf = { MY_PARAMETER : "something" }
		* // This works, because  MY_PARAMETER is present in conf
		* let checker = new ParameterChecker( conf, myCPM )
		* checker.confCheckPresenceOf( "MY_PARAMETER" )
		* 
		* // There will be an error if MY_PARAMETER is absent/undefined:
		* conf = {}
		* checker = new ParameterChecker( conf, myCPM )
		* checker.confCheckPresenceOf( "MY_PARAMETER" )
		* conf = { MY_PARAMETER : "undefined" }
		* checker = new ParameterChecker( conf, myCPM )
		* checker.confCheckPresenceOf( "MY_PARAMETER" )
		*/
		confCheckPresenceOf( name ){
			if( !this.conf.hasOwnProperty( name ) ){
				throw( "Cannot find parameter " + name + " in the conf object!" )
			}
			if( this.conf[name] == "undefined" ){
				throw( "Parameter " + name + " is undefined!" )
			}
		}
		
		/* ========= CHECKING PARAMETER STRUCTURE ======== */
		
		/** Helper function. Some parameters need to be specified for each {@link CellKind}, 
		so to check those we first need to know the number of cellkinds on the CPM.
		This number is retrieved from the CPM or added to it if it isn't there yet.
		@param {number} n_default - a number of cellkinds (including background), which is used
		to set the number of cellkinds in the CPM if it isn't there yet.
		@return {number} the number of non-background cellkinds as cached in the CPM.
		@private
		*/
		confCheckCellKinds( n_default ){
			if( !this.C ){
				throw("confCheck method called before addition to CPM!")
			}
			if( !("n_cell_kinds" in this.C) ){
				this.C.n_cell_kinds = n_default - 1;
			}
			return this.C.n_cell_kinds
		}
		
		/** Parameter structure for parameters that should come as a single value.
		This value can be of type string, boolean, or number.
		@example
		* let ACT_MEAN = "arithmetic" // Is a SingleValue parameter
		@typedef {number|string|boolean} SingleValue
		*/
		
		/** Check if a parameter consists of a single value (rather than an object or array),
		which can be a string, number, or boolean. 
		@param {SingleValue} p - the parameter to check, which must be a single
		string/number/boolean value.
		@param {string} name - the name of this parameter in the conf object, used for the
		error message if p is not a single value.
		
		@example
		* let checker = new ParameterChecker( conf, C )
		* let p1 = true, p2 = 1, p3 = "hi", p4 = [1,2]
		* // Nothing happens for parameters of type SingleValue:
		* checker.confCheckStructureSingle( p1, "MY_PARAMETER" )
		* checker.confCheckStructureSingle( p2, "MY_PARAMETER" )
		* checker.confCheckStructureSingle( p3, "MY_PARAMETER" )
		*
		* // This will throw an error because p4 is an array.
		* checker.confCheckStructureSingle( p4, "MY_PARAMETER" )
		*/
		confCheckStructureSingle( p, name ){
		
			// single values are of type string, number, or boolean. If that is the case,
			// just return.
			let type = typeof p;
			if( type == "string" || type == "number" || type == "boolean" ){
				return
			} else {
				throw( "Parameter " + name + " should be a single value!" )
			}

		}
		
		/** Parameter structure for parameters that should come in an array with an element
		for each {@link CellKind} including background.
		@example
		* let V = [0,5] // Is a KindArray parameter
		@typedef {Array} KindArray
		*/
		
		/** Check if a parameter has a {@link KindArray} structure.
		@param {KindArray} p - the parameter to check
		@param {string} name - the name of this parameter in the conf object, used for the
		error message if p is not a {@link KindArray}.
		
		@example
		* // C is a CPM which has 2 cellkinds including background:
		* let checker = new ParameterChecker( conf, C )
		* let p1 = [1,1], p2 = ["hi","hi"], p3 = "hi", p4 = [1,2,3]
		* // Nothing happens when parameters are indeed arrays of length 2
		* // (regardless of what type of array)
		* checker.confCheckStructureKindArray( p1, "MY_PARAMETER" )
		* checker.confCheckStructureKindArray( p2, "MY_PARAMETER" )
		*
		* // You'll get an error when p is no array, or when 
		* // its length doesn't match the number of cellkinds:
		* checker.confCheckStructureSingle( p3, "MY_PARAMETER" )
		* checker.confCheckStructureSingle( p4, "MY_PARAMETER" )
		*/
		confCheckStructureKindArray( p, name ){
			if( !(p instanceof Array) ){
				throw( "Parameter " + name + " should be an array!" )
			}	
			
			// Check if the array has an element for each cellkind incl. background
			let n_cell_kinds = this.confCheckCellKinds( p.length );
			if( this.conf[name].length != n_cell_kinds + 1 ){
				throw( "Parameter " + name + 
				" should be an array with an element for each cellkind including background!" )
			}
		}
		
		/** Parameter structure for parameters that specify interactions between two cells 
		with each a {@link CellKind}. Should be an array of arrays ("matrix") 
		where each array has an element for each cellkind including background. 
		Thus, M[n][m] specifies the parameter for an interaction between a cell of 
		cellkind n and a cell of cellkind m.
		@example
		* let J = [[0,20],[20,10]] // is a KindMatrix parameter.
		@typedef {Array} KindMatrix
		*/
		
		/** Checker if a parameter has a {@link KindMatrix} structure.
		If this is not the case, the method throws an error.
		@param {KindMatrix} p - the parameter to check
		@param {string} name - the name of this parameter in the conf object, used for the
		error message if p is not a {@link KindMatrix}.
		
		@example
		* // C is a CPM which has 2 cellkinds including background:
		* let checker = new ParameterChecker( conf, C )
		* let p1 = [[1,1],[1,1]], p2 = [["a","a"],["a","a"]] 
		* // Nothing happens when parameters are indeed arrays of length 2
		* // with sub-arrays of length 2 (regardless of what is in the elements)
		* checker.confCheckStructureKindArray( p1, "MY_PARAMETER" ) //OK
		* checker.confCheckStructureKindArray( p2, "MY_PARAMETER" ) //OK
		*
		* // You'll get an error when p is no array, or when 
		* // its length doesn't match the number of cellkinds:
		* let p3 = 1, p4 = [1,2,3], p5 = [[1,2],[1,2],[1,2]]
		* checker.confCheckStructureSingle( p3, "MY_PARAMETER" ) //error
		* checker.confCheckStructureSingle( p4, "MY_PARAMETER" ) //error
		* checker.confCheckStructureSingle( p5, "MY_PARAMETER" ) //error
		*/
		confCheckStructureKindMatrix( p, name ){

			let err1 = false, err2 = false;
			let n_cell_kinds;

			// err1: Check if the main array is an array and has the right size
			if( !(p instanceof Array) ){
				err1 = true;
			} else {
				n_cell_kinds = this.confCheckCellKinds( p.length );
				if( !( p.length == n_cell_kinds + 1 ) ){
					err1 = true;
				}		
			}
			if( err1 ){
				throw( "Parameter " + name + 
				" must be an array with a sub-array for each cellkind including background!" )
			}
			
			// Check if subarrays have the right size
			for( let e of p ){
				if( !(e instanceof Array) ){
					err2 = true;
				} else {
					if( !( e.length == n_cell_kinds + 1 ) ){
						err2 = true;
					}		
				}
				if( err2 ){
					throw( "Sub-arrays of " + name + 
					" must have an element for each cellkind including background!" )
				}
			}
		}
		
		/** Method for checking if the parameter has the right structure. Throws an error
		message if the parameter does not have this structure.
		
		It internally uses one of the following functions, depending on the structure argument:
		{@link confCheckStructureSingle}, {@link confCheckStructureKindArray}, or
		{@link confCheckStructureKindMatrix}.
		
		@param {SingleValue|KindArray|KindMatrix} p - the parameter to check the structure of
		@param {string} name - the name of this parameter in the conf object, used for the
		error message.
		@param {string} structure - parameter structure, which must be one of 
		"{@link SingleValue}", "{@link KindArray}", or "{@link KindMatrix}".
		
		@example
		* // My own configuration object
		* let conf = {
		*	P1 : true,
		* 	P2 : [0,2],
		* 	P3 : [-1,2,4],
		* 	P4 : [[1,2],[1,2]]
		* }
		* // C is a CPM which has 2 cellkinds including background:
		* let checker = new ParameterChecker( conf, C )
		* // These checks work out fine:
		* checker.confCheckStructure( conf["P1"],"P1","SingleValue")
		* checker.confCheckStructure( conf["P2"],"P2","KindArray")
		* checker.confCheckStructure( conf["P4"],"P4","KindMatrix")
		* 
		* // These checks throw an error:
		* checker.confCheckStructure( conf["P1"],"P1","KindArray") // not an array
		* checker.confCheckStructure( conf["P2"],"P3","KindArray") // too long
		*/
		confCheckStructure( p, name, structure ){
			if( structure == "SingleValue" ){
				this.confCheckStructureSingle( p, name );
			} else if ( structure == "KindArray" ){
				this.confCheckStructureKindArray( p, name );
			} else if ( structure == "KindMatrix" ){
				this.confCheckStructureKindMatrix( p, name );
			} else {
				throw("Unknown structure " + structure + ", please choose 'SingleValue', 'KindArray', or 'KindMatrix'.")
			}
		}
		
		/* ========= CHECKING VALUE TYPE ======== */
		
		/** Check if a value is of type 'number'.
		@param {number} v - value to check.
		@return {boolean} is v a number?
		@example
		this.isNumber( -1 ) // true
		this.isNumber( 0.5 ) // true
		this.isNumber( true ) // false
		this.isNumber( [1,2 ] ) // false
		this.isNumber( "hello world" ) // false
		@private
		*/
		isNumber( v ){
			return ( typeof v === "number" )
		}
		/** Check if a value is of type 'number' and non-negative.
		@param {number} v - value to check.
		@return {boolean} is v a non-negative number?
		@example
		this.isNonNegative( -1 ) // false
		this.isNonNegative( 0.5 ) // true
		this.isNumber( true ) // false
		this.isNumber( [1,2 ] ) // false
		this.isNumber( "hello world" ) // false
		@private
		*/
		isNonNegative( v ){
			if( !( typeof v === "number" ) || v < 0 ){
				return false
			}
			return true
		}
		/** Check if a value is of type 'number' and between 0 and 1.
		@param {number} v - value to check.
		@return {boolean} is v a number between 0 and 1?
		@example
		this.isProbability( -1 ) // false
		this.isProbability( 0.5 ) // true
		this.isProbability( true ) // false
		this.isProbability( [1,2 ] ) // false
		this.isProbability( "hello world" ) // false
		@private
		*/
		isProbability( v ){
			if( !( typeof v === "number" ) || v < 0 || v > 1 ){
				return false
			}
			return true
		}
		/** Check if a value is of type 'string' and has one of a set of
		predefined values.
		@param {number} v - value to check.
		@param {string[]} [values=[]] - possible values. If left empty,
		any string is considered OK.
		@return {boolean} is v a string and does it match one of the predefined values?
		@example
		this.isString( true ) // false
		this.isString( ["a","b"] ) // false
		this.isString( "hello world" ) // true
		@private
		*/
		isString( v, values = [] ){
			if( !( typeof v === "string" ) ){
				return false
			}
			let found = false;
			for( let val of values ){
				if( val == v ){
					found = true;
				}
			}
			return found
		}
		/** Check if a value is of type 'boolean'.
		@param {number} v - value to check.
		@return {boolean} is v a boolean?
		@example
		this.isBoolean( true ) // true
		this.isBoolean( [true,false] ) // false
		this.isBoolean( "hello world" ) // true
		@private
		*/
		isBoolean( v ){
			return( typeof v === "boolean" )
		}
		
		/** Check if a value is a coordinate in the dimensions of the current grid.
		@param {number} v - value to check.
		@return {boolean} is v a coordinate of the right dimensions?
		@public
		*/
		isCoordinate( v ){
			if( !( v instanceof Array ) ){
				return false
			}
			if( !v.length == this.C.extents.length ){
				return false
			}
			return true
		}
		
		/** Check if the elements of a given parameter are of the right type.
		It throws an error if this is not the case.
		@param {anything} p - parameter to check.
		@param {string} name - parameter name used for any error messages.
		@param {string} structure - parameter structure, which must be one of 
		"{@link SingleValue}", "{@link KindArray}", or "{@link KindMatrix}".
		@param {string} valuetype - type of value, which must be one of 
		"Number", "NonNegative", "Probability", "String", or "Boolean". 
		@param {string[]} values - predefined specific options for the value. 
		If left empty, this is ignored.
		@private
		*/
		confCheckValueType( p, name, structure, valuetype, values = [] ){
		
			// Determine which checkfunction will be used.
			let valuechecker;
			if( valuetype == "Number" ){
				valuechecker = this.isNumber;
			} else if ( valuetype == "NonNegative" ){
				valuechecker = this.isNonNegative;
			} else if ( valuetype == "Probability" ){
				valuechecker = this.isProbability;
			} else if ( valuetype == "String" ){
				valuechecker = this.isString;
			} else if ( valuetype == "Boolean" ){
				valuechecker = this.isBoolean;
			} else {
				throw( "Unsupported valuetype in check for parameter " + name +
					", please choose from: 'Number','NonNegative', 'Probability', 'String', 'Boolean'.")
			}
			
			let message =  "Parameter " + name + " : incorrect type. Expecting values of type " + valuetype + "."; 
			
			// structure determines how the checker is applied.
			if( structure == "SingleValue" ){
				if( p == "undefined" || !valuechecker( p, values ) ){ throw(message) }
			} else if ( structure == "KindArray" ){
				for( let i of p ){
					if( i == "undefined" || !valuechecker( i, values ) ){ throw(message) }
				}
			} else if ( structure == "KindMatrix" ){
				for( let i of p ){
					for( let j of i ){
						if( j == "undefined" || !valuechecker( j, values ) ){ throw(message) }
					}
				}
			} else {
				throw("Unknown structure " + structure + ", please choose 'SingleValue', 'KindArray', or 'KindMatrix'.")
			}
		}
		
		/** Check if a parameter exists and is defined, has the right structure, and if all 
		its elements are of the correct type. Throw an error if any of these do not hold.
		@param {string} name - parameter name used for any error messages.
		@param {string} structure - parameter structure, which must be one of 
		"{@link SingleValue}", "{@link KindArray}", or "{@link KindMatrix}".
		@param {string} valuetype - type of value, which must be one of 
		"Number", "NonNegative", "Probability", "String", or "Boolean". 
		@param {string[]} [values =[]] - predefined specific options for the value. 
		If left empty, this is ignored.
		
		@example
		* // from inside the confChecker() function of your constraint:
		* let checker = new ParameterChecker( this.conf, this.C )
		* 
		* checker.confCheckParameter( "MY_PARAMETER", "KindMatrix", "Number" ) // see Adhesion
		* checker.confCheckParameter( "MY_PARAMETER_2", "KindArray", "NonNegative" )
		*/
		confCheckParameter( name, structure, valuetype, values = [] ){
			this.confCheckPresenceOf( name );
			let p = this.conf[name];
			this.confCheckStructure( p, name, structure );
			this.confCheckValueType( p, name, structure, valuetype, values );
		}
		
		
	}

	/** 
	 * Implements the adhesion constraint of Potts models. 
	 * Each pair of neighboring pixels [n,m] gets a positive energy penalty deltaH if n and m
	 * do not belong to the same {@link CellId}.
	 *
	 * @example
	 * // Build a CPM and add the constraint
	 * let CPM = require( "path/to/build" )
	 * let C = new CPM.CPM( [200,200], { T : 20 } )
	 * C.add( new CPM.Adhesion( { J : [[0,20],[20,10]] } ) )
	 * 
	 * // Or add automatically by entering the parameters in the CPM
	 * let C2 = new CPM.CPM( [200,200], {
	 * 	T : 20,
	 * 	J : [[0,20],[20,10]]
	 * })
	 */
	class Adhesion extends SoftConstraint {

		/** The constructor of Adhesion requires a conf object with a single parameter J.
		@param {object} conf - parameter object for this constraint
		@param {CellKindInteractionMatrix} conf.J - J[n][m] gives the adhesion energy between a pixel of
		{@link CellKind} n and a pixel of {@link CellKind} m. J[n][n] is only non-zero
		when the pixels in question are of the same {@link CellKind}, but a different 
		{@link CellId}. Energies are given as non-negative numbers.
		*/
		constructor( conf ){
			super( conf );		
		}

		/** This method checks that all required parameters are present in the object supplied to
		the constructor, and that they are of the right format. It throws an error when this
		is not the case.*/
		confChecker(){
			let checker = new ParameterChecker( this.conf, this.C );
			checker.confCheckParameter( "J", "KindMatrix", "Number" );
		}

		/**  Get adhesion between two cells t1,t2 from "conf". 
		@param {CellId} t1 - cellid of the first cell.
		@param {CellId} t2 - cellid of the second cell.
		@return {number} adhesion between a pixel of t1 and one of t2.
		@private
		*/
		J( t1, t2 ){
			return this.conf["J"][this.C.cellKind(t1)][this.C.cellKind(t2)]
		}
		/**  Returns the Hamiltonian around a pixel i with cellid tp by checking all its
		neighbors that belong to a different cellid.
		@param {IndexCoordinate} i - coordinate of the pixel to evaluate hamiltonian at.
		@param {CellId} tp - cellid of this pixel.
		@return {number} sum over all neighbors of adhesion energies (only non-zero for 
		neighbors belonging to a different cellid).	
		@private
		 */
		H( i, tp ){
			let r = 0, tn;
			/* eslint-disable */
			const N = this.C.grid.neighi( i );
			for( let j = 0 ; j < N.length ; j ++ ){
				tn = this.C.pixti( N[j] );
				if( tn != tp ) r += this.J( tn, tp );
			}
			return r
		}
		/** Method to compute the Hamiltonian for this constraint. 
		 @param {IndexCoordinate} sourcei - coordinate of the source pixel that tries to copy.
		 @param {IndexCoordinate} targeti - coordinate of the target pixel the source is trying
		 to copy into.
		 @param {CellId} src_type - cellid of the source pixel.
		 @param {CellId} tgt_type - cellid of the target pixel. 
		 @return {number} the change in Hamiltonian for this copy attempt and this constraint.*/ 
		deltaH( sourcei, targeti, src_type, tgt_type ){
			return this.H( targeti, src_type ) - this.H( targeti, tgt_type )
		}
	}

	/** 
	 * Implements the volume constraint of Potts models. 
	 * 
	 * This constraint is typically used together with {@link Adhesion}.
	 * 
	 * See {@link VolumeConstraint#constructor} for the required parameters.
	 *
	 * @example
	 * // Build a CPM and add the constraint
	 * let CPM = require( "path/to/build" )
	 * let C = new CPM.CPM( [200,200], {
	 * 	T : 20,
	 * 	J : [[0,20],[20,10]]
	 * })
	 * C.add( new CPM.VolumeConstraint( {
	 * 	V : [0,500],
	 * 	LAMBDA_V : [0,5] 	
	 * } ) )
	 * 
	 * // Or add automatically by entering the parameters in the CPM
	 * let C2 = new CPM.CPM( [200,200], {
	 * 	T : 20,
	 * 	J : [[0,20],[20,10]],
	 * 	V : [0,500],
	 * 	LAMBDA_V : [0,5]
	 * })
	 */
	class VolumeConstraint extends SoftConstraint {


		/** The constructor of the VolumeConstraint requires a conf object with parameters.
		@param {object} conf - parameter object for this constraint
		@param {PerKindNonNegative} conf.LAMBDA_V - strength of the constraint per cellkind.
		@param {PerKindNonNegative} conf.V - Target volume per cellkind.
		*/
		constructor( conf ){
			super( conf );
		}
		
		/** This method checks that all required parameters are present in the object supplied to
		the constructor, and that they are of the right format. It throws an error when this
		is not the case.*/
		confChecker(){
			let checker = new ParameterChecker( this.conf, this.C );
			checker.confCheckParameter( "LAMBDA_V", "KindArray", "NonNegative" );
			checker.confCheckParameter( "V", "KindArray", "NonNegative" );
		}

		/** Method to compute the Hamiltonian for this constraint. 
		 @param {IndexCoordinate} sourcei - coordinate of the source pixel that tries to copy.
		 @param {IndexCoordinate} targeti - coordinate of the target pixel the source is trying
		 to copy into.
		 @param {CellId} src_type - cellid of the source pixel.
		 @param {CellId} tgt_type - cellid of the target pixel. 
		 @return {number} the change in Hamiltonian for this copy attempt and this constraint.*/ 
		deltaH( sourcei, targeti, src_type, tgt_type ){
			// volume gain of src cell
			let deltaH = this.volconstraint( 1, src_type ) - 
				this.volconstraint( 0, src_type );
			// volume loss of tgt cell
			deltaH += this.volconstraint( -1, tgt_type ) - 
				this.volconstraint( 0, tgt_type );
			return deltaH
		}
		/* ======= VOLUME ======= */

		/** The volume constraint term of the Hamiltonian for the cell with id t.
		@param {number} vgain - Use vgain=0 for energy of current volume, vgain=1 
			for energy if cell gains a pixel, and vgain = -1 for energy if cell loses a pixel.
		@param {CellId} t - the cellid of the cell whose volume energy we are computing.
		@return {number} the volume energy of this cell.
		*/
		volconstraint ( vgain, t ){
			const k = this.C.cellKind(t), l = this.conf["LAMBDA_V"][k];
			// the background "cell" has no volume constraint.
			if( t == 0 || l == 0 ) return 0
			const vdiff = this.conf["V"][k] - (this.C.getVolume(t) + vgain);
			return l*vdiff*vdiff
		}
	}

	/** This class implements the activity constraint of Potts models published in:
	 *
	 *	Niculescu I, Textor J, de Boer RJ (2015) 
	 *	Crawling and Gliding: A Computational Model for Shape-Driven Cell Migration. 
	 *	PLoS Comput Biol 11(10): e1004280. 
	 * 
	 * Pixels recently added to a cell get an "activity", which then declines with every MCS.
	 * Copy attempts from more active into less active pixels have a higher success rate,
	 * which puts a positive feedback on protrusive activity and leads to cell migration.
	 * 
	 * This constraint is generally used together with {@link Adhesion}, {@link VolumeConstraint},
	 * and {@link PerimeterConstraint}.
	 * 
	 * @see https://doi.org/10.1371/journal.pcbi.1004280
	 *
	 * @example
	 * // Build a CPM and add the constraint
	 * let CPM = require( "path/to/build" )
	 * let C = new CPM.CPM( [200,200], {
	 * 	T : 20,
	 * 	J : [[0,20],[20,10]],
	 * 	V : [0,500],
	 * 	LAMBDA_V : [0,5],
	 * 	P : [0,260],
	 * 	LAMBDA_P : [0,2] 	
	 * })
	 * C.add( new CPM.ActivityConstraint( {
	 * 	LAMBDA_ACT : [0,500],
	 * 	MAX_ACT : [0,30],
	 * 	ACT_MEAN : "geometric"
	 * } ) )
	 * 
	 * // Or add automatically by entering the parameters in the CPM
	 * let C2 = new CPM.CPM( [200,200], {
	 * 	T : 20,
	 * 	J : [[0,20],[20,10]],
	 * 	V : [0,500],
	 * 	LAMBDA_V : [0,5],
	 * 	P : [0,260],
	 * 	LAMBDA_P : [0,2],
	 * 	LAMBDA_ACT : [0,500],
	 * 	MAX_ACT : [0,30],
	 * 	ACT_MEAN : "geometric"	
	 * })
	 */
	class ActivityConstraint extends SoftConstraint {

		/** The constructor of the ActivityConstraint requires a conf object with parameters.
		@param {object} conf - parameter object for this constraint
		@param {string} [conf.ACT_MEAN="geometric"] - should local mean activity be measured with an
		"arithmetic" or a "geometric" mean?
		@param {PerKindNonNegative} conf.LAMBDA_ACT - strength of the activityconstraint per cellkind.
		@param {PerKindNonNegative} conf.MAX_ACT - how long do pixels remember their activity? Given per cellkind.
		*/
		constructor( conf ){
			super( conf );

			/** Activity of all cellpixels with a non-zero activity is stored in this object,
			with the {@link IndexCoordinate} of each pixel as key and its current activity as
			value. When the activity reaches 0, the pixel is removed from the object until it
			is added again. 
			@type {object}*/
			this.cellpixelsact = {};
			
			/** Wrapper: select function to compute activities based on ACT_MEAN in conf.
			Default is to use the {@link activityAtGeom} for a geometric mean.
			@type {function}*/
			this.activityAt = this.activityAtGeom;
			if( this.conf.ACT_MEAN == "arithmetic" ){
				this.activityAt = this.activityAtArith;
			} 
			
		}
		
		/** This method checks that all required parameters are present in the object supplied to
		the constructor, and that they are of the right format. It throws an error when this
		is not the case.*/
		confChecker(){
			let checker = new ParameterChecker( this.conf, this.C );
			checker.confCheckParameter( "ACT_MEAN", "SingleValue", "String", [ "geometric", "arithmetic" ] );
			checker.confCheckParameter( "LAMBDA_ACT", "KindArray", "NonNegative" );
			checker.confCheckParameter( "MAX_ACT", "KindArray", "NonNegative" );
		}
		
		/* ======= ACT MODEL ======= */

		/* Act model : compute local activity values within cell around pixel i.
		 * Depending on settings in conf, this is an arithmetic (activityAtArith)
		 * or geometric (activityAtGeom) mean of the activities of the neighbors
		 * of pixel i.
		 */
		/** Method to compute the Hamiltonian for this constraint. 
		 @param {IndexCoordinate} sourcei - coordinate of the source pixel that tries to copy.
		 @param {IndexCoordinate} targeti - coordinate of the target pixel the source is trying
		 to copy into.
		 @param {CellId} src_type - cellid of the source pixel.
		 @param {CellId} tgt_type - cellid of the target pixel. 
		 @return {number} the change in Hamiltonian for this copy attempt and this constraint.*/ 
		deltaH ( sourcei, targeti, src_type, tgt_type ){

			let deltaH = 0, maxact, lambdaact;
			const src_kind = this.C.cellKind( src_type );
			const tgt_kind = this.C.cellKind( tgt_type );

			// use parameters for the source cell, unless that is the background.
			// In that case, use parameters of the target cell.
			if( src_type != 0 ){
				maxact = this.conf["MAX_ACT"][src_kind];
				lambdaact = this.conf["LAMBDA_ACT"][src_kind];
			} else {
				// special case: punishment for a copy attempt from background into
				// an active cell. This effectively means that the active cell retracts,
				// which is different from one cell pushing into another (active) cell.
				maxact = this.conf["MAX_ACT"][tgt_kind];
				lambdaact = this.conf["LAMBDA_ACT"][tgt_kind];
			}
			if( !maxact || !lambdaact ){
				return 0
			}

			// compute the Hamiltonian. The activityAt method is a wrapper for either activityAtArith
			// or activityAtGeom, depending on conf (see constructor).	
			deltaH += lambdaact*(this.activityAt( targeti ) - this.activityAt( sourcei ))/maxact;
			return deltaH
		}

		/** Activity mean computation methods for arithmetic mean. It computes the mean activity
			of a pixel and all its neighbors belonging to the same cell.
			
			This method is generally called indirectly via {@link activityAt}, which is set
			based on the value of ACT_MEAN in the configuration object given to the constructor.
			
			@param {IndexCoordinate} i - pixel to evaluate local activity at.
			@return {number} the arithmetic mean of activities in this part of the cell.
			@private
		*/
		activityAtArith( i ){
			const t = this.C.pixti( i );
			
			// no activity for background/stroma
			if( t <= 0 ){ return 0 }
			
			// neighborhood pixels
			const N = this.C.neighi(i);
			
			// r activity summed, nN number of neighbors
			// we start with the current pixel. 
			let r = this.pxact(i), nN = 1;
			
			// loop over neighbor pixels
			for( let j = 0 ; j < N.length ; j ++ ){ 
				const tn = this.C.pixti( N[j] ); 
				
				// a neighbor only contributes if it belongs to the same cell
				if( tn == t ){
					r += this.pxact( N[j] );
					nN ++; 
				}
			}

			// average is summed r divided by num neighbors.
			return r/nN
		}
		/** Activity mean computation methods for geometric mean. It computes the mean activity
			of a pixel and all its neighbors belonging to the same cell.
			
			This method is generally called indirectly via {@link activityAt}, which is set
			based on the value of ACT_MEAN in the configuration object given to the constructor.
			
			@param {IndexCoordinate} i - pixel to evaluate local activity at.
			@return {number} the geometric mean of activities in this part of the cell.
			@private
		*/
		activityAtGeom ( i ){
			const t = this.C.pixti( i );

			// no activity for background/stroma
			if( t <= 0 ){ return 0 }
			
			//neighborhood pixels
			const N = this.C.neighi( i );
			
			// r activity product, nN number of neighbors.
			// we start with the current pixel.
			let nN = 1, r = this.pxact( i );

			// loop over neighbor pixels
			for( let j = 0 ; j < N.length ; j ++ ){ 
				const tn = this.C.pixti( N[j] ); 

				// a neighbor only contributes if it belongs to the same cell.
				// if it does and has activity 0, the product will also be zero so
				// we can already return.
				if( tn == t ){
					if( this.pxact( N[j] ) == 0 ) return 0
					r *= this.pxact( N[j] );
					nN ++; 
				}
			}
			
			// Geometric mean computation. 
			return Math.pow(r,1/nN)
		}


		/** Current activity (under the Act model) of the pixel at position i. 
		@param {IndexCoordinate} i the position of the pixel to evaluate the activity of.
		@return {number} the current activity of this pixel, which is >= 0.*/
		pxact ( i ){
			// If the pixel is not in the cellpixelsact object, it has activity 0.
			// Otherwise, its activity is stored in the object.
			return this.cellpixelsact[i] || 0
		}
		
		/** The postSetpixListener of the ActivityConstraint ensures that pixels are 
		given their maximal activity when they are freshly added to a CPM.
		@listens {CPM#setpixi} because when a new pixel is set (which is determined in the CPM),
		its activity must change so that this class knows about the update.
		@param {IndexCoordinate} i - the coordinate of the pixel that is changed.
		@param {CellId} t_old - the cellid of this pixel before the copy
		@param {CellId} t - the cellid of this pixel after the copy.
		*/
		/* eslint-disable no-unused-vars*/
		postSetpixListener( i, t_old, t ){
			// After setting a pixel, it gets the MAX_ACT value of its cellkind.
			const k = this.C.cellKind( t );
			this.cellpixelsact[i] = this.conf["MAX_ACT"][k];
		}
		
		/** The postMCSListener of the ActivityConstraint ensures that pixel activities
		decline with one after every MCS.
		@listens {CPM#timeStep} because when the CPM has finished an MCS, the activities must go down.
		*/
		postMCSListener(){
			// iterate over cellpixelsage and decrease all activities by one.
			for( let key in this.cellpixelsact ){
				// activities that reach zero no longer need to be stored.
				if( --this.cellpixelsact[ key ] <= 0 ){
					delete this.cellpixelsact[ key ];
				}
			}
		}


	}

	/** 
	 * Implements the perimeter constraint of Potts models. 
	 * A cell's "perimeter" is the number over all its borderpixels of the number of 
	 * neighbors that do not belong to the cell itself. 
	 * 
	 * This constraint is typically used together with {@link Adhesion} and {@VolumeConstraint}.
	 * 
	 * See {@link PerimeterConstraint#constructor} for the required parameters.
	 *
	 * @example
	 * // Build a CPM and add the constraint
	 * let CPM = require( "path/to/build" )
	 * let C = new CPM.CPM( [200,200], {
	 * 	T : 20,
	 * 	J : [[0,20],[20,10]],
	 * 	V : [0,500],
	 * 	LAMBDA_V : [0,5]
	 * })
	 * C.add( new CPM.PerimeterConstraint( {
	 * 	P : [0,260],
	 * 	LAMBDA_P : [0,2] 	
	 * } ) )
	 * 
	 * // Or add automatically by entering the parameters in the CPM
	 * let C2 = new CPM.CPM( [200,200], {
	 * 	T : 20,
	 * 	J : [[0,20],[20,10]],
	 * 	V : [0,500],
	 * 	LAMBDA_V : [0,5],
	 * 	P : [0,260],
	 * 	LAMBDA_P : [0,2]
	 * })
	 */
	class PerimeterConstraint extends SoftConstraint {

		/** The constructor of the PerimeterConstraint requires a conf object with
		 * parameters.
		 * @param {object} conf - parameter object for this constraint
		 * @param {PerKindNonNegative} conf.LAMBDA_P - strength of the perimeter
		 * 	constraint per cellkind.
		 * @param {PerKindNonNegative} conf.P - Target perimeter per cellkind.
		*/
		constructor( conf ){
			super( conf );
			
			/** The perimeter size of each pixel is tracked.
			@type {CellObject}*/
			this.cellperimeters = {};
		}

		/** Set the CPM attached to this constraint.
		@param {CPM} C - the CPM to attach.*/
		set CPM(C){
			/** The CPM this constraint acts on.
			@type {CPM}*/
			this.C = C;

			this.confChecker();

			// if C already has cells, initialize perimeters
			if( C.cellvolume.length !== 0 ){
				this.initializePerimeters();
			}
		}
		
		/** This method checks that all required parameters are present in the
		 * object supplied to the constructor, and that they are of the right
		 * format. It throws an error when this is not the case.*/
		confChecker(){
			let checker = new ParameterChecker( this.conf, this.C );
			checker.confCheckParameter( "LAMBDA_P", "KindArray", "NonNegative" );
			checker.confCheckParameter( "P", "KindArray", "NonNegative" );
		}

		/** This method initializes the this.cellperimeters object when the
		 * constraint is added to a non-empty CPM. */
		initializePerimeters(){

			for( let bp of this.C.cellBorderPixels() ){
				const p = bp[0];
				let cid = this.C.pixt(p);
				if( !(cid in this.cellperimeters) ){
					this.cellperimeters[cid] = 0;
				}
				const i = this.C.grid.p2i( p );
				this.cellperimeters[cid] += this.C.perimeterNeighbours[i];
			}

		}

		/** The postSetpixListener of the PerimeterConstraint ensures that cell
		 * perimeters are updated after each copy in the CPM.
		 * @listens {CPM#setpixi} because when a new pixel is set (which is
		 * 	determined in the CPM),	some of the cell perimeters will change.
		 * @param {IndexCoordinate} i - the coordinate of the pixel that is changed.
		 * @param {CellId} t_old - the cellid of this pixel before the copy
		 * @param {CellId} t_new - the cellid of this pixel after the copy.
		*/
		/* eslint-disable no-unused-vars*/
		postSetpixListener( i, t_old, t_new ){
			if( t_old === t_new ){ return }
			
			// Neighborhood of the pixel that changes
			const Ni = this.C.neighi( i );
			
			// Keep track of perimeter before and after copy
			let n_new = 0, n_old = 0;
			
			// Loop over the neighborhood. 
			for( let i = 0 ; i < Ni.length ; i ++  ){
				const nt = this.C.pixti(Ni[i]);
				
				// neighbors are added to the perimeter if they are
				// of a different cellID than the current pixel
				if( nt !== t_new ){
					n_new ++; 
				}
				if( nt !== t_old ){
					n_old ++;
				}
				
				// if the neighbor is non-background, the perimeter
				// of the cell it belongs to may also have to be updated.
				if( nt !== 0 ){
				
					// if it was of t_old, its perimeter goes up because the
					// current pixel will no longer be t_old. This means it will
					// have a different type and start counting as perimeter.
					if( nt === t_old ){
						this.cellperimeters[nt] ++;
					}
					// opposite if it is t_new.
					if( nt === t_new ){
						this.cellperimeters[nt] --;
					}
				}
			}
			
			// update perimeters of the old and new type if they are non-background
			if( t_old !== 0 ){
				this.cellperimeters[t_old] -= n_old;
			}
			if( t_new !== 0 ){
				if( !(t_new in this.cellperimeters) ){
					this.cellperimeters[t_new] = 0;
				}
				this.cellperimeters[t_new] += n_new;
			}
		}
		
		/** Method to compute the Hamiltonian for this constraint.
		 * @param {IndexCoordinate} sourcei - coordinate of the source pixel that
		 * 	tries to copy.
		 * @param {IndexCoordinate} targeti - coordinate of the target pixel the
		 * 	source is trying to copy into.
		 * @param {CellId} src_type - cellid of the source pixel.
		 * @param {CellId} tgt_type - cellid of the target pixel.
		 * @return {number} the change in Hamiltonian for this copy attempt and
		 * this constraint.*/
		deltaH( sourcei, targeti, src_type, tgt_type ){
			if( src_type === tgt_type ){
				return 0
			}
			const ts = this.C.cellKind(src_type);
			const ls = this.conf["LAMBDA_P"][ts];
			const tt = this.C.cellKind(tgt_type);
			const lt = this.conf["LAMBDA_P"][tt];
			if( !(ls>0) && !(lt>0) ){
				return 0
			}
			const Ni = this.C.neighi( targeti );
			let pchange = {};
			pchange[src_type] = 0; pchange[tgt_type] = 0;
			for( let i = 0 ; i < Ni.length ; i ++  ){
				const nt = this.C.pixti(Ni[i]);
				if( nt !== src_type ){
					pchange[src_type]++; 
				}
				if( nt !== tgt_type ){
					pchange[tgt_type]--;
				}
				if( nt === tgt_type ){
					pchange[nt] ++;
				}
				if( nt === src_type ){
					pchange[nt] --;
				}
			}
			let r = 0.0;
			if( ls > 0 ){
				const pt = this.conf["P"][ts],
					ps = this.cellperimeters[src_type];
				const hnew = (ps+pchange[src_type])-pt,
					hold = ps-pt;
				r += ls*((hnew*hnew)-(hold*hold));
			}
			if( lt > 0 ){
				const pt = this.conf["P"][tt],
					ps = this.cellperimeters[tgt_type];
				const hnew = (ps+pchange[tgt_type])-pt,
					hold = ps-pt;
				r += lt*((hnew*hnew)-(hold*hold));
			}
			// eslint-disable-next-line
			//console.log( r )
			return r
		}
	}

	/** Extension of class {@link Constraint} used for a hard constraint. See description in
	 {@link Constraint} for details. This class is not used on its own but serves as a base
	 class for a hard constraint. */
	class HardConstraint extends Constraint {

		/** Let the CPM know that this is a soft constraint, so return 'soft'.
		@return {string} "hard"*/
		get CONSTRAINT_TYPE() {
			return "hard"
		}
		/*constructor( conf ){
			this.conf = conf
		}*/
		/*set CPM(C){
			this.C = C
		}*/
		
		/** Hard constraints must have a 'fulfilled' method to compute whether the copy attempt fulfills the rule.
		This method must be implemented in the subclass.
		 @param {IndexCoordinate} src_i - coordinate of the source pixel that tries to copy.
		 @param {IndexCoordinate} tgt_i - coordinate of the target pixel the source is trying
		 to copy into.
		 @param {CellId} src_type - cellid of the source pixel.
		 @param {CellId} tgt_type - cellid of the target pixel. 
		 @return {boolean} whether the copy attempt satisfies the constraint.
		 @abstract
		 */ 
		// eslint-disable-next-line no-unused-vars
		fulfilled( src_i, tgt_i, src_type, tgt_type ){
			throw("You need to implement the 'fulfilled' method for this constraint!")
		}
	}

	/** 
	 * This constraint allows a "barrier" celltype from and into which copy attempts are forbidden.
	 * @example
	 * // Build a CPM and add the constraint
	 * let CPM = require( "path/to/build" )
	 * let C = new CPM.CPM( [200,200], {
	 * 	T : 20,
	 * 	J : [[0,0,20],[0,0,5],[20,5,10]],
	 * 	V : [0,0,500],
	 * 	LAMBDA_V : [0,0,5],
	 * })
	 * C.add( new CPM.BarrierConstraint( {
	 * 	IS_BARRIER : [false,true,false]
	 * } ) )
	 * 
	 * // OR: add automatically by entering the parameters in the CPM
	 * C = new CPM.CPM( [200,200], {
	 * 	T : 20,
	 * 	J : [[0,0,20],[0,0,5],[20,5,10]],
	 * 	V : [0,0,500],
	 * 	LAMBDA_V : [0,0,5],
	 * 	IS_BARRIER : [false,true,false]
	 * })
	 * 
	 * // Make a horizontal line barrier (cellkind 1 )
	 * let cid = C.makeNewCellID( 1 )
	 * for( let x = 0; x < 200; x++ ){
	 * 	C.setpix( [x,95], cid )
	 * }
	 * // Seed a cell (cellkind2)
	 * let gm = new CPM.GridManipulator( C )
	 * gm.seedCell(2)
	 */
	class BarrierConstraint extends HardConstraint {

		/** The constructor of the BarrierConstraint requires a conf object with a single parameter.
		@param {object} conf - parameter object for this constraint.
		@param {PerKindBoolean} conf.IS_BARRIER - specify for each cellkind if it should be 
	 	considered as a barrier. If so, all copy attempts into and from it are forbidden.
		*/
		constructor( conf ){
			super(conf);
		}

		/** This method checks that all required parameters are present in the object supplied to
		the constructor, and that they are of the right format. It throws an error when this
		is not the case.*/
		confChecker(){
			let checker = new ParameterChecker( this.conf, this.C );
			checker.confCheckParameter( "IS_BARRIER", "KindArray", "Boolean" );
		}

		/** Method for hard constraints to compute whether the copy attempt fulfills the rule.
		 @param {IndexCoordinate} src_i - coordinate of the source pixel that tries to copy.
		 @param {IndexCoordinate} tgt_i - coordinate of the target pixel the source is trying
		 to copy into.
		 @param {CellId} src_type - cellid of the source pixel.
		 @param {CellId} tgt_type - cellid of the target pixel. 
		 @return {boolean} whether the copy attempt satisfies the constraint.*/ 
		fulfilled( src_i, tgt_i, src_type, tgt_type ){
		
			// Fulfilled = false when either src or tgt pixel is of the barrier cellkind	
			if( this.conf["IS_BARRIER"][this.C.cellKind( src_type ) ] ){
				return false
			}

			if( this.conf["IS_BARRIER"][this.C.cellKind( tgt_type ) ] ){
				return false
			}

			return true
		}
	}

	/** This class enables automatic addition of Hamiltonian terms to a CPM
	 * through their parameter names.
	 *
	 * For each parameter name, we specify one Hamiltonian term. If the parameter
	 * is present, then a new instance of this term is initialized with the CPM's
	 * configuration as parameter and added to the CPM. 
	@type {object}
	@property {Constraint} J - An {@link Adhesion} constraint is added when there is a parameter J.
	@property {Constraint} LAMBDA_V - A {@link VolumeConstraint} is added when there is a parameter LAMBDA_V.
	@property {Constraint} LAMBDA_P - A {@link PerimeterConstraint} is added when there is a parameter LAMBDA_P.
	@property {Constraint} LAMBDA_ACT - An {@link ActivityConstraint} is added when there is a parameter LAMBDA_ACT.
	@property {Constraint} IS_BARRIER - A {@link BarrierConstraint} is added when there is a parameter IS_BARRIER.
	*/
	let AutoAdderConfig = {
		J : Adhesion,
		LAMBDA_V : VolumeConstraint,
		LAMBDA_ACT : ActivityConstraint,
		LAMBDA_P : PerimeterConstraint,
		IS_BARRIER : BarrierConstraint
	};

	/** The core CPM class. Can be used for two- or three-dimensional simulations.
	*/
	class CPM extends GridBasedModel {

		/** The constructor of class CA.
		 * @param {GridSize} field_size - the size of the grid of the model.
		 * @param {object} conf - configuration options; see below. In addition,
		 * the conf object can have parameters to constraints added to the CPM.
		 * See the different {@link Constraint} subclasses for options. For some
		 * constraints, adding its parameter to the CPM conf object automatically
		 * adds the constraint; see {@link AutoAdderConfig} to see for which
		 * constraints this is supported.
		 * @param {boolean[]} [conf.torus=[true,true,...]] - should the grid have
		 * linked borders?
		 * @param {number} [conf.T] - the temperature of this CPM. At higher
		 * temperatures, unfavourable copy attempts are more likely to be accepted.
		 * @param {number} [conf.seed] - seed for the random number generator. If
		 * left unspecified, a random number from the Math.random() generator is
		 * used to make one.
		 * */
		constructor( field_size, conf ){
			super( field_size, conf );

			// ---------- CPM specific stuff here
			
			/** Number of non-background cells currently on the grid.
			@type{number}*/
			this.nr_cells = 0;
			/** Highest cell ID previously assigned. 
			@type{number}*/
			this.last_cell_id = 0;
			/** track border pixels for speed 
			@type {DiceSet}*/
			this.borderpixels = new DiceSet( this.mt );
			/** Private property used by {@link updateborderneari} to track borders. 
			@private
			@type {Uint16Array} */
			this._neighbours = new Uint16Array(this.grid.p2i(field_size));

			//  ---------- Attributes per cell:
			/** Store the {@CellKind} of each cell on the grid. 
			@example
			this.t2k[1] // cellkind of cell with cellId 1
			@type {CellObject}
			*/
			this.t2k = [];	// cell type ("kind"). Example: this.t2k[1] is the cellKind of cell 1.
			this.t2k[0] = 0;	// Background cell; there is just one cell of this type.

			//  ---------- CPM constraints
			/** Array of objects of (@link SoftConstraint) subclasses attached to the CPM.
			These are used to determine {@link deltaH}.
			@type {Array}*/
			this.soft_constraints = [];
			/** Object showing which constraints are where in {@link soft_constraints}. Used
			by the {@link getConstraint} method to find an attached constraint by name.
			@type {object}*/
			this.soft_constraints_indices = {};
			/** Array of objects of (@link HardConstraint) subclasses attached to the CPM.
			These are used to determine which copy attempts are allowed in a {@link timeStep}.
			@type {Array}*/
			this.hard_constraints = [];
			/** Object showing which constraints are where in {@link soft_constraints}. Used
			by the {@link getConstraint} method to find an attached constraint by name.
			@type {object}*/
			this.hard_constraints_indices = {};
			/** Array of functions that need to be executed after every {@link setpixi} event.
			These functions are often implemented in subclasses of {@link Constraint} that
			need to track some specific property on the grid. 
			@type {function[]}*/
			this.post_setpix_listeners = [];
			/** Array of functions that need to be executed after every {@link timeStep} event.
			These functions are often implemented in subclasses of {@link Constraint} that
			need to track some specific property on the grid. 
			@type {function[]}*/
			this.post_mcs_listeners = [];
			
			/* Automatically add constraints by their parameters in conf. This only works
			for some constraints specified in AutoAdderConfig. */
			for( let x of Object.keys( conf ) ){
				if( x in AutoAdderConfig ){
					this.add( new AutoAdderConfig[x]( conf ) );
				}
			}
		}

		/* This is no different from the GridBasedModel function and can go. 
		neigh(p, torus=this.conf.torus){
			let g = this.grid
			return g.neighi( g.p2i(p), torus ).map( function(i){ return g.i2p(i) } )
		}*/

		/** Iterator returning non-background pixels on the grid.
		@return {Pixel} for each pixel, return an array [p,v] where p are
			the pixel's array coordinates on the grid, and v its value.*/
		* cellPixels() {
			for( let p of this.grid.pixels() ){
				if( p[1] !== 0 ){
					yield p;
				}
			}
		}

		/** Iterator returning non-background border pixels on the grid.
		See {@link cellBorderPixelIndices} for a version returning pixels
		by their {@link IndexCoordinate} instead of {@link ArrayCoordinate}.
		
		@return {Pixel} for each pixel, return an array [p,v] where p are
			the pixel's array coordinates on the grid, and v its value.*/
		* cellBorderPixels() {
			for( let i of this.borderpixels.elements ){
				const t = this.grid.pixti(i);
				if( t !== 0 ){
					yield [this.grid.i2p(i),t];
				}
			}
		}

		/** Iterator returning non-background border pixels on the grid.
		See {@link cellBorderPixels} for a version returning pixels
		by their {@link ArrayCoordinate} instead of {@link IndexCoordinate}.
		
		@return {iPixel} for each pixel, return an array [p,v] where p are
			the pixel's array coordinates on the grid, and v its value.*/
		* cellBorderPixelIndices() {
			for( let i of this.borderpixels.elements ){
				const t = this.grid.pixti(i);
				if( t !== 0 ){
					yield [i,t];
				}
			}
		}

		/** Add a constraint to the CPM, ensuring that its {@link SoftConstraint#deltaH} or
		{@link HardConstraint#fulfilled} methods are called appropriately during a copy attempt.
		Any postSetpixListeners and postMCSListeners are also executed at the appropriate times.
		@param {Constraint} t - the constraint object to add.
		*/
		add( t ){
			let tName = t.constructor.name, i;
			if( t.CONSTRAINT_TYPE ){
				switch( t.CONSTRAINT_TYPE ){
				
				case "soft": 
					// Add constraint to the array of soft constraints
					i = this.soft_constraints.push( t );
					
					// Write this index to an array in the 
					// this.soft_constraints_indices object, for lookup later. 
					if( !this.soft_constraints_indices.hasOwnProperty(tName) ){
						this.soft_constraints_indices[tName] = [];
					}
					this.soft_constraints_indices[tName].push( i-1 );
					break
					
				case "hard": 
					// Add constraint to the array of soft constraints
					i = this.hard_constraints.push( t );
					
					// Write this index to an array in the 
					// this.soft_constraints_indices object, for lookup later. 
					if( !this.hard_constraints_indices.hasOwnProperty(tName) ){
						this.hard_constraints_indices[tName] = [];
					}
					this.hard_constraints_indices[tName].push( i-1 );
					break
				}
			}
			if( typeof t["postSetpixListener"] === "function" ){
				this.post_setpix_listeners.push( t.postSetpixListener.bind(t) );
			}
			if( typeof t["postMCSListener"] === "function" ){
				this.post_mcs_listeners.push( t.postMCSListener.bind(t) );
			}
			t.CPM = this;
			if( typeof t["postAdd"] === "function" ){
				t.postAdd();
			}
		}
		
		/** Get a {@link Constraint} object linked to this CPM by the name of its class.
		By default, the first constraint found of this class is returned. It is possible
		that there are multiple constraints of the same type on the CPM; in that case,
		supply its number (by order in which the constraints of this type were added) to 
		get a specific one. 
		
		This function can be useful if you need to access information in the constraint object,
		such as the cell directions in a {@PersistenceConstraint}, from outside. You can use
		this for stuff like drawing.
		
		@param {string} constraintname - name of the constraint class you are looking for.
		@param {number} [num = 0] - if multiple constraints of this class are present, 
		return the num-th one added to the CPM. 
		*/
		getConstraint( constraintname, num ){
		
			if( !num ){
				num = 0;
			}
			let i;
			
			if( this.hard_constraints_indices.hasOwnProperty( constraintname ) ){
				i = this.hard_constraints_indices[constraintname][num];
				return this.hard_constraints[i]
			} else if ( this.soft_constraints_indices.hasOwnProperty( constraintname ) ){
				i = this.soft_constraints_indices[constraintname][num];
				return this.soft_constraints[i]
			} else {
				throw("No constraint of name " + " exists in this CPM!")
			}	
		
		}

		/** Get {@link CellId} of the pixel at coordinates p. 
		@param {ArrayCoordinate} p - pixel to get cellid of.
		@return {CellId} ID of the cell p belongs to.*/
		pixt( p ){
			return this.grid.pixti( this.grid.p2i(p) )
		}

		/** Get volume of the cell with {@link CellId} t 
		@param {CellId} t - id of the cell to get volume of.
		@return {number} the cell's current volume. */ 
		getVolume( t ){
			return this.cellvolume[t]
		}

		/** Get the {@link CellKind} of the cell with {@link CellId} t. 
		Overwrites {@link GridBasedModel#cellKind} because in a CPM, the two are not the same.
		@param {CellId} t - id of the cell to get kind of.
		@return {CellKind} the cellkind. */
		cellKind( t ){
			return this.t2k[ t ]
		}

		/** Assign the cell with {@link CellId} t to {@link CellKind} k.
		@param {CellId} t - id of the cell to assing
		@param {CellKind} k - cellkind to give it. 
		*/
		setCellKind( t, k ){
			this.t2k[ t ] = k;
		}
		
		
		/* ------------- COMPUTING THE HAMILTONIAN --------------- */

		/** returns total change in hamiltonian for all registered soft constraints together.
		 @param {IndexCoordinate} sourcei - coordinate of the source pixel that tries to copy.
		 @param {IndexCoordinate} targeti - coordinate of the target pixel the source is trying
		 to copy into.
		 @param {CellId} src_type - cellid of the source pixel.
		 @param {CellId} tgt_type - cellid of the target pixel. 
		 @return {number} the change in Hamiltonian for this copy attempt.
		*/
		deltaH ( sourcei, targeti, src_type, tgt_type ){
			let r = 0.0;
			for( let t of this.soft_constraints ){
				r += t.deltaH( sourcei, targeti, src_type, tgt_type );
			}
			return r
		}
		/* ------------- COPY ATTEMPTS --------------- */

		/** Simulate one Monte Carlo Step. We now just use {@link timeStep} for consistency
		with other {@link GridBasedModel}s, but we have kept this method for compatibility
		with earlier version. Internally, it just calls {@link timeStep}.
		*/
		monteCarloStep () {
			this.timeStep();
		}
		
		/** A time step in the CPM is a Monte Carlo step. This performs a 
		  	 number of copy attempts depending on grid size:
		  	 
			1) Randomly sample one of the border pixels for the copy attempt.
			2) Compute the change in Hamiltonian for the suggested copy attempt.
			3) With a probability depending on this change, decline or accept the 
			   copy attempt and update the grid accordingly. 

			@todo TODO it is quite confusing that the "borderpixels" array also
			contains border pixels of the background.
		*/
		timeStep (){
			let delta_t = 0.0;
			// this loop tracks the number of copy attempts until one MCS is completed.
			while( delta_t < 1.0 ){
				// This is the expected time (in MCS) you would expect it to take to
				// randomly draw another border pixel.
				delta_t += 1./(this.borderpixels.length);

				// sample a random pixel that borders at least 1 cell of another type,
				// and pick a random neighbour of tha pixel
				const tgt_i = this.borderpixels.sample();
				const Ni = this.grid.neighi( tgt_i );
				const src_i = Ni[this.ran(0,Ni.length-1)];
			
				const src_type = this.grid.pixti( src_i );
				const tgt_type = this.grid.pixti( tgt_i );

				// only compute the Hamiltonian if source and target belong to a different cell,
				// and do not allow a copy attempt into the stroma. Only continue if the copy attempt
				// would result in a viable cell.
				if( tgt_type != src_type ){
					let ok = true;
					for( let h of this.hard_constraints ){
						if( !h.fulfilled( src_i, tgt_i, src_type, tgt_type ) ){
							ok = false; break
						}
					}
					if( ok ){
						const hamiltonian = this.deltaH( src_i, tgt_i, src_type, tgt_type );
						// probabilistic success of copy attempt 
						if( this.docopy( hamiltonian ) ){
							this.setpixi( tgt_i, src_type );
						}
					}
				} 
			}
			this.time++; // update time with one MCS.
			/** Cached values of these stats. Object with stat name as key and its cached
			value as value. The cache must be cleared when the grid changes!
			@type {object} */
			this.stat_values = {}; // invalidate stat value cache
			for( let l of this.post_mcs_listeners ){
				l();
			}
		}	

		/** Determine whether copy attempt will succeed depending on deltaH (stochastic). 
		@param {number} deltaH - energy change associated with the potential copy.
		@return {boolean} whether the copy attempt succeeds.
		*/
		docopy ( deltaH ){
			if( deltaH < 0 ) return true
			return this.random() < Math.exp( -deltaH / this.conf.T )
		}
		
		/** Change the pixel at position i into {@link CellId} t. 
		This method overrides {@link GridBasedModel#setpixi} because we want to
		add postSetpixListeners for all the constraints, to keep track of relevant information.
		
		See also {@link setpix} for a method working with {@link ArrayCoordinate}s.
		
		@param {IndexCoordinate} i - coordinate of pixel to change.
		@param {CellId} t - cellid to change this pixel into.
		*/
		setpixi ( i, t ){		
			const t_old = this.grid.pixti(i);
			if( t_old > 0 ){
				// also update volume of the old cell
				// (unless it is background/stroma)
				this.cellvolume[t_old] --;
				
				// if this was the last pixel belonging to this cell, 
				// remove the cell altogether.
				if( this.cellvolume[t_old] == 0 ){
					delete this.cellvolume[t_old];
					delete this.t2k[t_old];
					this.nr_cells--;
				}
			}
			// update volume of the new cell and cellid of the pixel.
			this.grid.setpixi(i,t);
			if( t > 0 ){
				this.cellvolume[t] ++;
			}
			this.updateborderneari( i, t_old, t );
			//this.stat_values = {} // invalidate stat value cache
			for( let l of this.post_setpix_listeners ){
				l( i, t_old, t );
			}
		}

		get perimeterNeighbours(){
			return this._neighbours
		}

		/** Update border elements ({@link borderpixels}) after a successful copy attempt. 
		@listens {setpixi} because borders change when a copy succeeds.
		@param {IndexCoordinate} i - coordinate of pixel that has changed.
		@param {CellId} t_old - id of the cell the pixel belonged to before the copy.
		@param {CellId} t_new - id of the cell the pixel has changed into.
		*/
		updateborderneari ( i, t_old, t_new ){
			if( t_old == t_new ) return
			const Ni = this.grid.neighi( i );
			const wasborder = this._neighbours[i] > 0; 
			this._neighbours[i] = 0;
			for( let ni of Ni  ){
				const nt = this.grid.pixti(ni);
				if( nt != t_new ){
					this._neighbours[i] ++; 
				}
				if( nt == t_old ){
					if( this._neighbours[ni] ++ == 0 ){
						this.borderpixels.insert( ni );
					}
				}
				if( nt == t_new ){
					if( --this._neighbours[ni] == 0 ){
						this.borderpixels.remove( ni );
					}
				}
			}

			if( !wasborder && this._neighbours[i] > 0 ){
				this.borderpixels.insert( i );
			}
			if( wasborder &&  this._neighbours[i] == 0 ){
				this.borderpixels.remove( i );
			}
		}

		/* ------------- MANIPULATING CELLS ON THE GRID --------------- */

		/** Initiate a new {@link CellId} for a cell of {@link CellKind} "kind", and create elements
		   for this cell in the relevant arrays (cellvolume, t2k).
		   @param {CellKind} kind - cellkind of the cell that has to be made.
		   @return {CellId} of the new cell.*/
		makeNewCellID ( kind ){
			const newid = ++ this.last_cell_id;
			this.cellvolume[newid] = 0;
			this.setCellKind( newid, kind );
			return newid
		}

	}

	/** This class encapsulates a lower-resolution grid and makes it
	   visible as a higher-resolution grid. Only exact subsampling by
	   a constant factor per dimension is supported. 
	   
	   This class is useful when combining information of grids of
	   different sizes. This is often the case for chemotaxis, where
	   we let diffusion occur on a lower resolution grid to speed things up.
	   This class then allows you to obtain chemokine information from the 
	   low resolution chemokine grid using coordinates from the linked,
	   higher resolution model grid.
	   
	   @example <caption>Linear interpolation on a low resolution chemokine grid</caption>
	   * let CPM = require( "path/to/build" )
	   * 
	   * // Define a grid with float values for chemokine values, and set the middle pixel
	   * let chemogrid = new CPM.Grid2D( [50,50], [true,true], "Float32" )
	   * chemogrid.setpix( [99,99], 100 )
	   * 
	   * // Make a coarse grid at 5x as high resolution, which is then 500x500 pixels.
	   * let coarsegrid = new CPM.CoarseGrid( chemogrid, 5 )
	   * 
	   * // Use interpolation. Pixels close to the midpoint won't have the exact same
	   * // value of either 100 or 0, but something inbetween.
	   * let p1 = [250,250], p2 = [250,251]
	   * console.log( "p1 : " + coarsegrid.pixt(p1) + ", p2 : " + coarsegrid.pixt(p2) )
	   * // p1 : 100, p2 : 80 
	   * 
	   * // Or draw it to see this. Compare these two:
	   * let Cim1 = new CPM.Canvas( coarsegrid )
	   * Cim1.drawField()
	   * let Cim2 = new CPM.Canvas( chemogrid, {zoom:5} )
	   * Cim2.drawField()
	*/
	class CoarseGrid extends Grid2D {
		/** The constructor of class CoarseGrid takes a low resolution grid as input
		and a factor 'upscale', which is how much bigger the dimensions of the high
		resolution grid are (must be a constant factor). 
		@param {Grid2D} grid the grid to scale up; currently only supports the {@link Grid2D} class.
		@param {number} upscale The (integer) factor to magnify the original grid with. */
		constructor( grid, upscale = 3 ){
		
			let extents = new Array( grid.extents.length );
			for( let i = 0 ; i < grid.extents.length ; i++ ){
				extents[i] = upscale * grid.extents[i];
			}
			super( extents, grid.torus, "Float32" );
		
			/** Size of the new grid in all dimensions.
			@type {GridSize} with a non-negative integer number for each dimension. */
			this.extents = extents;
			/** The original, low-resolution grid. 
			@type {Grid2D}*/
			this.grid = grid;
			
			/** The upscale factor (a positive integer number).
			@private
			@type {number} */
			this.upscale = upscale;
		}

		/** The pixt method takes as input a coordinate on the bigger grid, and maps it
		to the corresponding value on the resized small grid via bilinear interpolation.
		This prevents artefacts from the lower resolution of the second grid: the 
		[upscale x upscale] pixels that map to the same pixel in the low resolution grid
		do not get the same value.
		@param {ArrayCoordinate} p array coordinates on the high resolution grid.
		@return {number} interpolated value from the low resolution grid at this position. */
		pixt( p ){
		
			// 2D bilinear interpolation. Find the 4 positions on the original, low resolution grid
			// that are closest to the requested position p: x-coordinate l,r (left/right) 
			// and y-coordinate t,b (top/bottom)
		
			let positions = this.positions(p); // [t,r,b,l,h,v]
			let t = positions[0], r = positions[1], b = positions[2], l = positions[3],
				h = positions[4], v = positions[5];

			// Get the values on those 4 positions
			let f_lt = this.grid.pixt([l,t]);
			let f_rt = this.grid.pixt([r,t]);
			let f_lb = this.grid.pixt([l,b]);
			let f_rb = this.grid.pixt([r,b]);

			// Average these weighted by their distance to the current pixel.
			let f_x_b = f_lb * (1-h) + f_rb * h; 
			let f_x_t = f_lt * (1-h) + f_rt * h;

			return f_x_t*(1-v) + f_x_b * v
		}
		
		/** This method takes as input a coordinate on the bigger grid, and 'adds' additional
		value to it by adding the proper amount to the corresponding positions on the low
		resolution grid.
		@param {ArrayCoordinate} p array coordinates on the high resolution grid.
		@param {number} value - value that should be added to this position.
		*/
		addValue( p, value ){
			
			// 2D bilinear interpolation, the other way around.
			// Find the 4 positions on the original, low res grid that are closest to the
			// requested position p
			
			let positions = this.positions(p); 
			let t = positions[0], r = positions[1], b = positions[2], l = positions[3],
				h = positions[4], v = positions[5];
				
			
			let v_lt = value * (1-h) * (1-v);
			let v_lb = value * (1-h) * v;
			let v_rt = value * h * (1-v);
			let v_rb = value * h * v;
			
			
			this.grid.setpix( [l,t], this.grid.pixt([l,t]) + v_lt );
			this.grid.setpix( [l,b], this.grid.pixt([l,b]) + v_lb );
			this.grid.setpix( [r,t], this.grid.pixt([r,t]) + v_rt );
			this.grid.setpix( [r,b], this.grid.pixt([r,b]) + v_rb );
			
		}
		/** @private 
		@ignore */
		positions( p ){
			// Find the 4 positions on the original, low resolution grid
			// that are closest to the requested position p: x-coordinate l,r (left/right) 
			// and y-coordinate t,b (top/bottom)
			let l = ~~(p[0] / this.upscale); // ~~ is a fast alternative for Math.floor
			let r = l+1;
			
			let t = ~~(p[1] / this.upscale);
			let b = t+1;
			
			// Find the horizontal/vertical distances of these positions to p
			let h = (p[0]%this.upscale)/this.upscale;
			let v = (p[1]%this.upscale)/this.upscale;
			
			// Correct grid boundaries depending on torus
			if( r > this.grid.extents[0] ){
				if( this.grid.torus[0] ){
					r = 0;
				} else {
					r = this.grid.extents[0];
					h = 0.5;
				}
			}
			
			if( b > this.grid.extents[1] ){
				if( this.grid.torus[1] ){
					b = 0;
				} else {
					b = this.grid.extents[1];
					v = 0.5;
				}
				
			}
			
			return [t,r,b,l,h,v]
		}

		/*gradient( p ){
			let ps = new Array( p.length )
			for( let i = 0 ; i < p.length ; i ++ ){
				ps[i] = ~~(p[i]/this.upscale)
			}
			return this.grid.gradient( ps )
		}*/
	}

	/** Base class for a statistic that can be computed on a GridBasedModel. 
	This class by itself is not usable; see its subclasses for stats that are 
	currently supported. */
	class Stat {

		/** The constructor of class Stat takes a 'conf' object as argument.
		However, Stats should not really be configurable in the sense that they should always
		provide an expected output. The 'conf' object is mainly intended
		to provide an option to configure logging / debugging output. That
		is not implemented yet.	
		@param {object} conf configuration options for the Stat, which should change nothing
		about the return value produced by the compute() method but may be used for logging
		and debugging options.*/
		constructor( conf ){
			/** Configuration object for the stat, which should not change its value but
			may be used for logging and debugging options.
			@type {object}*/
			this.conf = conf || {};
		}
		
		/** Every stat is linked to a specific model.
		@param {GridBasedModel} M the model to compute the stat on.*/
		set model( M ){
		
			/** The model to compute the stat on.
			@type {GridBasedModel} */
			this.M = M;
		}
		
		/** The compute method of the base Stat class throws an error, 
		enforcing that you have to implement this method when you build a new 
		stat class extending this base class. 
		@abstract */
		compute(){
			throw("compute method not implemented for subclass of Stat")
		}
	}

	/** This Stat creates an object with the cellpixels of each cell on the grid. 
		Keys are the {@link CellId} of all cells on the grid, corresponding values are arrays
		containing the pixels belonging to that cell. Each element of that array contains
		the {@link ArrayCoordinate} for that pixel.
		
		@example
		* let CPM = require( "path/to/build" )
		*
		* // Make a CPM, seed a cell, and get the PixelsByCell
		* let C = new CPM.CPM( [100,100], { 
		* 	T:20,
		* 	J:[[0,20],[20,10]],
		* 	V:[0,200],
		* 	LAMBDA_V:[0,2]
		* } )
		* let gm = new CPM.GridManipulator( C )
		* gm.seedCell(1)
		* gm.seedCell(1)
		* for( let t = 0; t < 100; t++ ){ C.timeStep() }
		* C.getStat( CPM.PixelsByCell )
	*/
	class PixelsByCell extends Stat {

		/** The compute method of PixelsByCell creates an object with cellpixels of each
		cell on the grid.
		@return {CellArrayObject} object with for each cell on the grid
		an array of pixels (specified by {@link ArrayCoordinate}) belonging to that cell.
		*/
		compute(){
			// initialize the object
			let cellpixels = { };
			// The this.M.pixels() iterator returns coordinates and cellid for all 
			// non-background pixels on the grid. See the appropriate Grid class for
			// its implementation.
			for( let [p,i] of this.M.pixels() ){
				if( !cellpixels[i] ){
					cellpixels[i] = [p];
				} else {
					cellpixels[i].push( p );
				}
			}
			return cellpixels
		}
	}

	/**
	 * The ActivityMultiBackground constraint implements the activity constraint of Potts models,
	 but allows users to specify locations on the grid where LAMBDA_ACT is different. 
	 See {@link ActivityConstraint} for the normal version of this constraint.
	 See {@link ActivityMultiBackground#constructor} for an explanation of the parameters.
	 */
	class ActivityMultiBackground extends ActivityConstraint {

		/** Creates an instance of the ActivityMultiBackground constraint 
		* @param {object} conf - Configuration object with the parameters.
		* ACT_MEAN is a single string determining whether the activity mean should be computed
		* using a "geometric" or "arithmetic" mean. 
		*/
		/** The constructor of the ActivityConstraint requires a conf object with parameters.
		@param {object} conf - parameter object for this constraint
		@param {string} [conf.ACT_MEAN="geometric"] - should local mean activity be measured with an
		"arithmetic" or a "geometric" mean?
		@param {PerKindArray} conf.LAMBDA_ACT_MBG - strength of the activityconstraint per cellkind and per background.
		@param {PerKindNonNegative} conf.MAX_ACT - how long do pixels remember their activity? Given per cellkind.
		@param {Array} conf.BACKGROUND_VOXELS - an array where each element represents a different background type.
		This is again an array of {@ArrayCoordinate}s of the pixels belonging to that backgroundtype. These pixels
		will have the LAMBDA_ACT_MBG value of that backgroundtype, instead of the standard value.
		*/
		constructor( conf ){
			super( conf );

			/** Activity of all cellpixels with a non-zero activity is stored in this object,
			with the {@link IndexCoordinate} of each pixel as key and its current activity as
			value. When the activity reaches 0, the pixel is removed from the object until it
			is added again. 
			@type {object}*/
			this.cellpixelsact = {}; // activity of cellpixels with a non-zero activity
			
			/** Wrapper: select function to compute activities based on ACT_MEAN in conf.
			Default is to use the {@link activityAtGeom} for a geometric mean.
			@type {function}*/
			this.activityAt = this.activityAtGeom;
			if( this.conf.ACT_MEAN == "arithmetic" ){
				this.activityAt = this.activityAtArith;
			} 
			
			/** Store which pixels belong to which background type 
			@type {Array}*/
			this.bgvoxels = [];
			
			/** Track if this.bgvoxels has been set.
			@type {boolean}*/
			this.setup = false;
		}
		
		/** This method checks that all required parameters are present in the object supplied to
		the constructor, and that they are of the right format. It throws an error when this
		is not the case.*/
		confChecker(){
			let checker = new ParameterChecker( this.conf, this.C );
			checker.confCheckParameter( "ACT_MEAN", "SingleValue", "String", [ "geometric", "arithmetic" ] );
			checker.confCheckPresenceOf( "LAMBDA_ACT_MBG" );
			checker.confCheckParameter( "MAX_ACT", "KindArray", "NonNegative" );
			
			// Custom checks
			checker.confCheckStructureKindArray( this.conf["LAMBDA_ACT_MBG"], "LAMBDA_ACT_MBG" );
			for( let e of this.conf["LAMBDA_ACT_MBG"] ){
				for( let i of e ){
					if( !checker.isNonNegative(i) ){
						throw("Elements of LAMBDA_ACT_MBG must be non-negative numbers!")
					}
				}
			}
			checker.confCheckPresenceOf( "BACKGROUND_VOXELS" );
			let bgvox = this.conf["BACKGROUND_VOXELS"];
			// Background voxels must be an array of arrays
			if( !(bgvox instanceof Array) ){
				throw( "Parameter BACKGROUND_VOXELS should be an array of at least two arrays!" )
			} else if ( bgvox.length < 2 ){
				throw( "Parameter BACKGROUND_VOXELS should be an array of at least two arrays!" )
			}
			// Elements of the initial array must be arrays.
			for( let e of bgvox ){
				if( !(e instanceof Array) ){
					throw( "Parameter BACKGROUND_VOXELS should be an array of at least two arrays!" )
				}
				
				// Entries of this array must be pixel coordinates, which are arrays of length C.extents.length
				for( let ee of e ){
					let isCoordinate = true;
					if( !(ee instanceof Array) ){
						isCoordinate = false;
					} else if ( ee.length != this.C.extents.length ){
						isCoordinate = false;
					}
					if( !isCoordinate ){
						throw( "Parameter BACKGROUND_VOXELS: subarray elements should be ArrayCoordinates; arrays of length " + this.C.extents.length + "!" )
					}
				}
			}
		}
		
		/** Get the background voxels from input argument or the conf object and store them in a correct format
		in this.bgvoxels. This only has to be done once, but can be called from outside to
		change the background voxels during a simulation (eg in a HTML page).
		 */	
		setBackgroundVoxels( voxels ){
		
			voxels = voxels || this.conf["BACKGROUND_VOXELS"];
		
			// reset if any exist already
			this.bgvoxels = [];
			for( let bgkind = 0; bgkind < voxels.length; bgkind++ ){
				this.bgvoxels.push({});
				for( let v of voxels[bgkind] ){
					this.bgvoxels[bgkind][ this.C.grid.p2i(v) ] = true;
				}
			}
			this.setup = true;

		}
		
		/* ======= ACT MODEL ======= */

		/* Act model : compute local activity values within cell around pixel i.
		 * Depending on settings in conf, this is an arithmetic (activityAtArith)
		 * or geometric (activityAtGeom) mean of the activities of the neighbors
		 * of pixel i.
		 */
		/** Method to compute the Hamiltonian for this constraint. 
		 @param {IndexCoordinate} sourcei - coordinate of the source pixel that tries to copy.
		 @param {IndexCoordinate} targeti - coordinate of the target pixel the source is trying
		 to copy into.
		 @param {CellId} src_type - cellid of the source pixel.
		 @param {CellId} tgt_type - cellid of the target pixel. 
		 @return {number} the change in Hamiltonian for this copy attempt and this constraint.*/ 
		deltaH ( sourcei, targeti, src_type, tgt_type ){
		
			if( ! this.setup ){
				this.setBackgroundVoxels();
			}

			let deltaH = 0, maxact, lambdaact;
			const src_kind = this.C.cellKind( src_type );
			const tgt_kind = this.C.cellKind( tgt_type );
			let bgindex1 = 0, bgindex2 = 0;
			
			for( let bgkind = 0; bgkind < this.bgvoxels.length; bgkind++ ){
				if( sourcei in this.bgvoxels[bgkind] ){
					bgindex1 = bgkind;
				}
				if( targeti in this.bgvoxels[bgkind] ){
					bgindex2 = bgkind;
				}
			}
			

			// use parameters for the source cell, unless that is the background.
			// In that case, use parameters of the target cell.
			if( src_type != 0 ){
				maxact = this.conf["MAX_ACT"][src_kind];
				lambdaact = this.conf["LAMBDA_ACT_MBG"][src_kind][bgindex1];
			} else {
				// special case: punishment for a copy attempt from background into
				// an active cell. This effectively means that the active cell retracts,
				// which is different from one cell pushing into another (active) cell.
				maxact = this.conf["MAX_ACT"][tgt_kind];
				lambdaact = this.conf["LAMBDA_ACT_MBG"][tgt_kind][bgindex2];
			}
			if( !maxact || !lambdaact ){
				return 0
			}

			// compute the Hamiltonian. The activityAt method is a wrapper for either activityAtArith
			// or activityAtGeom, depending on conf (see constructor).	
			deltaH += lambdaact*(this.activityAt( targeti ) - this.activityAt( sourcei ))/maxact;
			return deltaH
		}


	}

	/**
	 * Class for taking a CPM grid and displaying it in either browser or with
	 *  nodejs.
	 * Note: when using this class from outside the module, you don't need to import
	 *  it separately but can access it from CPM.Canvas. */
	class Canvas {
		/** The Canvas constructor accepts a CPM object C or a Grid2D object.
		@param {GridBasedModel|Grid2D|CoarseGrid} C - the object to draw, which must
		 be an object of class {@link GridBasedModel} (or its subclasses {@link CPM}
		 and {@link CA}), or a 2D grid ({@link Grid2D} or {@link CoarseGrid}).
		 Drawing of other grids is currently not supported.
		@param {object} [options = {}] - Configuration settings
		@param {number} [options.zoom = 1]- positive number specifying the zoom
		 level to draw with.
		@param {number[]} [options.wrap = [0,0,0]] - if nonzero: 'wrap' the grid to
		 these dimensions; eg a pixel with x coordinate 201 and wrap[0] = 200 is
		 displayed at x = 1.
		@param {string} [options.parentElement = document.body] - the element on
		 the html page where the canvas will be appended.

		@example <caption>A CPM with Canvas</caption>
		* let CPM = require( "path/to/build" )
		*
		* // Create a CPM, corresponding Canvas and GridManipulator
		* // (Use CPM. prefix from outside the module)
		* let C = new CPM.CPM( [200,200], {
		* 	T : 20,
		* 	J : [[0,20][20,10]],
		* 	V:[0,500],
		* 	LAMBDA_V:[0,5]
		* } )
		* let Cim = new CPM.Canvas( C, {zoom:2} )
		* let gm = new CPM.GridManipulator( C )
		*
		* // Seed a cell at [x=100,y=100] and run 100 MCS.
		* gm.seedCellAt( 1, [100,100] )
		* for( let t = 0; t < 100; t++ ){
		* 	C.timeStep()
		* }
		*
		* // Draw the cell and save an image
		* Cim.drawCells( 1, "FF0000" )			// draw cells of CellKind 1 in red
		* Cim.writePNG( "my-cell-t100.png" )
		*/
		constructor( C, options ){
			if( C instanceof GridBasedModel ){
				/**
				 * The underlying model that is drawn on the canvas.
				 * @type {GridBasedModel|CPM|CA}
				 */
				this.C = C;
				/**
				 * The underlying grid that is drawn on the canvas.
				 * @type {Grid2D|CoarseGrid}
				 */
				this.grid = this.C.grid;

				/** Grid size in each dimension, taken from the CPM or grid object
				 * to draw.
				 * @type {GridSize} each element is the grid size in that dimension
				 * in pixels */
				this.extents = C.extents;
			} else if( C instanceof Grid2D  ||  C instanceof CoarseGrid ){

				this.grid = C;
				this.extents = C.extents;
			}
			/** Zoom level to draw the canvas with, set to options.zoom or its
			 * default value 1.
			 * @type {number}*/
			this.zoom = (options && options.zoom) || 1;
			/** if nonzero: 'wrap' the grid to these dimensions; eg a pixel with x
			 * coordinate 201 and wrap[0] = 200 is displayed at x = 1.
			 * @type {number[]} */
			this.wrap = (options && options.wrap) || [0,0,0];

			/** Width of the canvas in pixels (in its unzoomed state)
			 * @type {number}*/
			this.width = this.wrap[0];
			/** Height of the canvas in pixels (in its unzoomed state)
			 * @type {number}*/
			this.height = this.wrap[1];

			if( this.width === 0 || this.extents[0] < this.width ){
				this.width = this.extents[0];
			}
			if( this.height === 0 || this.extents[1] < this.height ){
				this.height = this.extents[1];
			}

			if( typeof document !== "undefined" ){
				/** @ignore */
				this.el = document.createElement("canvas");
				this.el.width = this.width*this.zoom;
				this.el.height = this.height*this.zoom;//extents[1]*this.zoom
				let parent_element = (options && options.parentElement) || document.body;
				parent_element.appendChild( this.el );
			} else {
				const {createCanvas} = require("canvas");
				/** @ignore */
				this.el = createCanvas( this.width*this.zoom,
					this.height*this.zoom );
				/** @ignore */
				this.fs = require("fs");
			}

			/** @ignore */
			this.ctx = this.el.getContext("2d");
			this.ctx.lineWidth = .2;
			this.ctx.lineCap="butt";
		}

		/** Give the canvas element an ID supplied as argument. Useful for building
		 * an HTML page where you want to get this canvas by its ID.
		 * @param {string} idString - the name to give the canvas element.
		 * */
		setCanvasId( idString ){
			this.el.id = idString;
		}


		/* Several internal helper functions (used by drawing functions below) : */

		/** @private
		 * @ignore*/
		pxf( p ){
			this.ctx.fillRect( this.zoom*p[0], this.zoom*p[1], this.zoom, this.zoom );
		}

		/** @private
		 * @ignore */
		pxfi( p, alpha=1 ){
			const dy = this.zoom*this.width;
			const off = (this.zoom*p[1]*dy + this.zoom*p[0])*4;
			for( let i = 0 ; i < this.zoom*4 ; i += 4 ){
				for( let j = 0 ; j < this.zoom*dy*4 ; j += dy*4 ){
					this.px[i+j+off] = this.col_r;
					this.px[i+j+off + 1] = this.col_g;
					this.px[i+j+off + 2] = this.col_b;
					this.px[i+j+off + 3] = alpha*255;
				}
			}
		}

		/** @private
		 * @ignore */
		pxfir( p ){
			const dy = this.zoom*this.width;
			const off = (p[1]*dy + p[0])*4;
			this.px[off] = this.col_r;
			this.px[off + 1] = this.col_g;
			this.px[off + 2] = this.col_b;
			this.px[off + 3] = 255;
		}

		/** @private
		 * @ignore*/
		getImageData(){
			/** @ignore */
			this.image_data = this.ctx.getImageData(0, 0, this.width*this.zoom, this.height*this.zoom);
			/** @ignore */
			this.px = this.image_data.data;
		}

		/** @private
		 * @ignore*/
		putImageData(){
			this.ctx.putImageData(this.image_data, 0, 0);
		}

		/** @private
		 * @ignore*/
		pxfnozoom( p ){
			this.ctx.fillRect( this.zoom*p[0], this.zoom*p[1], 1, 1 );
		}

		/** draw a line left (l), right (r), down (d), or up (u) of pixel p
		 * @private
		 * @ignore */
		pxdrawl( p ){
			for( let i = this.zoom*p[1] ; i < this.zoom*(p[1]+1) ; i ++ ){
				this.pxfir( [this.zoom*p[0],i] );
			}
		}

		/** @private
		 * @ignore */
		pxdrawr( p ){
			for( let i = this.zoom*p[1] ; i < this.zoom*(p[1]+1) ; i ++ ){
				this.pxfir( [this.zoom*(p[0]+1),i] );
			}
		}
		/** @private
		 * @ignore */
		pxdrawd( p ){
			for( let i = this.zoom*p[0] ; i < this.zoom*(p[0]+1) ; i ++ ){
				this.pxfir( [i,this.zoom*(p[1]+1)] );
			}
		}
		/** @private
		 * @ignore */
		pxdrawu( p ){
			for( let i = this.zoom*p[0] ; i < this.zoom*(p[0]+1) ; i ++ ){
				this.pxfir( [i,this.zoom*p[1]] );
			}
		}

		/** For easier color naming
		 * @private
		 * @ignore */
		col( hex ){
			this.ctx.fillStyle="#"+hex;
			/** @ignore */
			this.col_r = parseInt( hex.substr(0,2), 16 );
			/** @ignore */
			this.col_g = parseInt( hex.substr(2,2), 16 );
			/** @ignore */
			this.col_b = parseInt( hex.substr(4,2), 16 );
		}

		/** Hex code string for a color.
		 * @typedef {string} HexColor*/

		/** Color the whole grid in color [col], or in black if no argument is given.
		 * @param {HexColor} [col = "000000"] -hex code for the color to use, defaults to black.
		 */
		clear( col ){
			col = col || "000000";
			this.ctx.fillStyle="#"+col;
			this.ctx.fillRect( 0,0, this.el.width, this.el.height );
		}

		/** Rendering context of canvas.
		 * @see https://developer.mozilla.org/en-US/docs/Web/API/CanvasRenderingContext2D
		 * @typedef {object} CanvasRenderingContext2D
		 * */

		/** Return the current drawing context.
		 * @return {CanvasRenderingContext2D} current drawing context on the canvas.
		 * */
		context(){
			return this.ctx
		}
		/** @private
		 * @ignore */
		p2pdraw( p ){
			for( let dim = 0; dim < p.length; dim++ ){
				if( this.wrap[dim] !== 0 ){
					p[dim] = p[dim] % this.wrap[dim];
				}
			}
			return p
		}

		/* DRAWING FUNCTIONS ---------------------- */

		/** Use to color a grid according to its values. High values are colored in
		 * a brighter color.
		 * @param {Grid2D|CoarseGrid} [cc] - the grid to draw values for. If left
		 * unspecified, the grid that was originally supplied to the Canvas
		 * constructor is used.
		 * @param {HexColor} [col = "0000FF"] - the color to draw the chemokine in.
		 * */
		drawField( cc, col = "0000FF" ){
			if( !cc ){
				cc = this.grid;
			}
			this.col(col);
			let maxval = 0;
			for( let i = 0 ; i < cc.extents[0] ; i ++ ){
				for( let j = 0 ; j < cc.extents[1] ; j ++ ){
					let p = Math.log(.1+cc.pixt([i,j]));
					if( maxval < p ){
						maxval = p;
					}
				}
			}
			this.getImageData();
			//this.col_g = 0
			//this.col_b = 0
			for( let i = 0 ; i < cc.extents[0] ; i ++ ){
				for( let j = 0 ; j < cc.extents[1] ; j ++ ){
					//let colval = 255*(Math.log(.1+cc.pixt( [i,j] ))/maxval)
					let alpha = (Math.log(.1+cc.pixt( [i,j] ))/maxval);
					//this.col_r = colval
					//this.col_g = colval
					this.pxfi([i,j], alpha);
				}
			}
			this.putImageData();
			this.ctx.globalAlpha = 1;
		}
		/** Use to color a grid according to its values. High values are colored in
		 * a brighter color.
		 * @param {Grid2D|CoarseGrid} [cc] - the grid to draw values for. If left
		 * unspecified, the grid that was originally supplied to the Canvas
		 * constructor is used.
		 * @param {number} [nsteps = 10] - the number of contour lines to draw.
		 * Contour lines are evenly spaced between the min and max log10 of the
		 * chemokine.
		 * @param {HexColor} [col = "FFFF00"] - the color to draw contours with.
		 * */
		drawFieldContour( cc, nsteps = 10, col = "FFFF00" ){
			if( !cc ){
				cc = this.grid;
			}
			this.col(col);
			let maxval = 0;
			let minval = Math.log(0.1);
			for( let i = 0 ; i < cc.extents[0] ; i ++ ){
				for( let j = 0 ; j < cc.extents[1] ; j ++ ){
					let p = Math.log(.1+cc.pixt([i,j]));
					if( maxval < p ){
						maxval = p;
					}
					if( minval > p ){
						minval = p;
					}
				}
			}


			this.getImageData();
			//this.col_g = 0
			//this.col_b = 0
			//this.col_r = 255

			let step = (maxval-minval)/nsteps;
			for( let v = minval; v < maxval; v+= step ){

				for( let i = 0 ; i < cc.extents[0] ; i ++ ){
					for( let j = 0 ; j < cc.extents[1] ; j ++ ){

						let pixelval = Math.log( .1 + cc.pixt( [i,j] ) );
						if( Math.abs( v - pixelval ) < 0.05*maxval ){
							let below = false, above = false;
							for( let n of this.grid.neighNeumanni( this.grid.p2i( [i,j] ) ) ){

								let nval = Math.log(0.1 + cc.pixt(this.grid.i2p(n)) );
								if( nval < v ){
									below = true;
								}
								if( nval >= v ){
									above = true;
								}
								if( above && below ){
									//this.col_r = 150*((v-minval)/(maxval-minval)) + 105
									let alpha = 0.7*((v-minval)/(maxval-minval)) + 0.3;
									this.pxfi( [i,j], alpha );
									break
								}
							}
						}


					}
				}

			}


			this.putImageData();
		}


		/** @desc Method for drawing the cell borders for a given cellkind in the
		 * color specified in "col" (hex format). This function draws a line around
		 * the cell (rather than coloring the outer pixels). If [kind] is negative,
		 * simply draw all borders.
		 *
		 * See {@link drawOnCellBorders} to color the outer pixels of the cell.
		 *
		 * @param {CellKind} kind - Integer specifying the cellkind to color.
		 * Should be a positive integer as 0 is reserved for the background.
		 * @param {HexColor}  [col = "000000"] - hex code for the color to use,
		 * defaults to black.
	   */
		drawCellBorders( kind, col ){

			let isCPM = ( this.C instanceof CPM ), C = this.C;
			let getBorderPixels = function*(){
				for( let p of C.cellBorderPixels() ){
					yield p;
				}
			};
			if( !isCPM ){
				// in a non-cpm, simply draw borders of all pixels
				getBorderPixels = function*(){
					for( let p of C.grid.pixels() ){
						yield p;
					}
				};
			}


			col = col || "000000";
			let pc, pu, pd, pl, pr, pdraw;
			this.col( col );
			this.getImageData();
			// cst contains indices of pixels at the border of cells
			for( let x of getBorderPixels() ){

				let pKind;
				if( isCPM ){
					pKind = this.C.cellKind( x[1] );
				} else {
					pKind = x[1];
				}

				let p = x[0];
				if( kind < 0 || pKind === kind ){
					pdraw = this.p2pdraw( p );

					pc = this.C.pixt( [p[0],p[1]] );
					pr = this.C.pixt( [p[0]+1,p[1]] );
					pl = this.C.pixt( [p[0]-1,p[1]] );
					pd = this.C.pixt( [p[0],p[1]+1] );
					pu = this.C.pixt( [p[0],p[1]-1] );

					if( pc !== pl  ){
						this.pxdrawl( pdraw );
					}
					if( pc !== pr ){
						this.pxdrawr( pdraw );
					}
					if( pc !== pd ){
						this.pxdrawd( pdraw );
					}
					if( pc !== pu ){
						this.pxdrawu( pdraw );
					}
				}

			}
			this.putImageData();
		}

		/** Use to show activity values of the act model using a color gradient, for
		 * cells in the grid of cellkind "kind". The constraint holding the activity
		 * values can be supplied as an argument. Otherwise, the current CPM is
		 * searched for the first registered activity constraint and that is then
		 * used.
		 *
		 * @param {CellKind} kind - Integer specifying the cellkind to color.
		 * If negative, draw values for all cellkinds.
		 * @param {ActivityConstraint|ActivityMultiBackground} [A] - the constraint
		 * object to use, which must be of class {@link ActivityConstraint} or
		 * {@link ActivityMultiBackground} If left unspecified, this is the first
		 * instance of an ActivityConstraint or ActivityMultiBackground object found
		 * in the soft_constraints of the attached CPM.
		 * @param {Function} [col] - a function that returns a color for a number
		 * in [0,1] as an array of red/green/blue values, for example, [255,0,0]
		 * would be the color red. If unspecified, a green-to-red heatmap is used.
		 * */
		drawActivityValues( kind, A, col ){
			if( !( this.C instanceof CPM) ){
				throw("You cannot use the drawActivityValues method on a non-CPM model!")
			}
			if( !A ){
				for( let c of this.C.soft_constraints ){
					if( c instanceof ActivityConstraint || c instanceof ActivityMultiBackground ){
						A = c; break
					}
				}
			}
			if( !A ){
				throw("Cannot find activity values to draw!")
			}
			if( !col ){
				col = function(a){
					let r = [0,0,0];
					if( a > 0.5 ){
						r[0] = 255;
						r[1] = (2-2*a)*255;
					} else {
						r[0] = (2*a)*255;
						r[1] = 255;
					}
					return r
				};
			}
			// cst contains the pixel ids of all non-background/non-stroma cells in
			// the grid. 
			let ii, sigma, a, k;
			// loop over all pixels belonging to non-background, non-stroma
			this.col("FF0000");
			this.getImageData();
			this.col_b = 0;
			//this.col_g = 0
			for( let x of this.C.cellPixels() ){
				ii = x[0];
				sigma = x[1];
				k = this.C.cellKind(sigma);

				// For all pixels that belong to the current kind, compute
				// color based on activity values, convert to hex, and draw.
				if( ( kind < 0 && A.conf["MAX_ACT"][k] > 0 ) || k === kind ){
					a = A.pxact( this.C.grid.p2i( ii ) )/A.conf["MAX_ACT"][k];
					if( a > 0 ){
						if( a > 0.5 ){
							this.col_r = 255;
							this.col_g = (2-2*a)*255;
						} else {
							this.col_r = (2*a)*255;
							this.col_g = 255;
						}
						let r = col( a );
						this.col_r = r[0];
						this.col_g = r[1];
						this.col_b = r[2];
						this.pxfi( ii );
					}
				}
			}
			this.putImageData();
		}

		/** Color outer pixel of all cells of kind [kind] in col [col].
		 * See {@link drawCellBorders} to actually draw around the cell rather than
		 * coloring the outer pixels. If you're using this model on a CA,
		 * {@link CellKind} is not defined and the parameter "kind" is instead
		 * interpreted as {@link CellId}.
		 *
		 * @param {CellKind} kind - Integer specifying the cellkind to color.
		 * Should be a positive integer as 0 is reserved for the background.
		 * @param {HexColor|function} col - Optional: hex code for the color to use.
		 * If left unspecified, it gets the default value of black ("000000").
		 * col can also be a function that returns a hex value for a cell id. */
		drawOnCellBorders( kind, col ){
			col = col || "000000";

			let isCPM = ( this.C instanceof CPM ), C = this.C;
			let getBorderPixels = function*(){
				for( let p of C.cellBorderPixels() ){
					yield p;
				}
			};
			if( !isCPM ){
				// in a non-cpm, simply draw borders of all pixels
				getBorderPixels = this.C.pixels;
			}

			this.getImageData();
			this.col( col );
			for( let p of getBorderPixels() ){

				let pKind;
				if( isCPM ){
					pKind = this.C.cellKind( p[1] );
				} else {
					pKind = p[1];
				}

				if( kind < 0 || pKind === kind ){
					if( typeof col == "function" ){
						this.col( col(p[1]) );
					}
					this.pxfi( p[0] );
				}
			}
			this.putImageData();
		}

		/**
		 * Draw all cells of cellid "id" in color col (hex). Note that this function
		 * also works for CA.
		 *
		 * @param {CellId} id - id of the cell to color.
		 * @param {HexColor} col - Optional: hex code for the color to use.
		 * If left unspecified, it gets the default value of black ("000000").
		 *
		 * */
		drawCellsOfId( id, col ){
			if( !col ){
				col = "000000";
			}
			if( typeof col == "string" ){
				this.col(col);
			}


			// Use the pixels() iterator to get the id of all non-background pixels.
			this.getImageData();

			for( let x of this.C.pixels() ){
				if( x[1] === id ){

					this.pxfi( x[0] );

				}
			}

			this.putImageData();
		}

		/** Draw all cells of cellkind "kind" in color col (hex). This method is
		 * meant for models of class {@link CPM}, where the {@link CellKind} is
		 * defined. If you apply this method on a {@link CA} model, this method
		 * will internally call {@link drawCellsOfId} by just supplying the
		 * "kind" parameter as {@link CellId}.
		 *
		 * @param {CellKind} kind - Integer specifying the cellkind to color.
		 * Should be a positive integer as 0 is reserved for the background.
		 * @param {HexColor|function} col - Optional: hex code for the color to use.
		 * If left unspecified, it gets the default value of black ("000000").
		 * col can also be a function that returns a hex value for a cell id, but
		 * this is only supported for CPMs.
		 * */
		drawCells( kind, col ){
			if( !( this.C instanceof CPM ) ){
				if( typeof col != "string" ){
					throw("If you use the drawCells method on a CA, you cannot " +
						"specify the color as function! Please specify a single string.")
				}
				this.drawCellsOfId( kind, col );
			} else {
				if (!col) {
					col = "000000";
				}
				if (typeof col == "string") {
					this.col(col);
				}
				// Object cst contains pixel index of all pixels belonging to non-background,
				// non-stroma cells.

				let cellpixelsbyid = this.C.getStat(PixelsByCell);

				/*for( let x of this.C.pixels() ){
					if( kind < 0 || this.C.cellKind(x[1]) == kind ){
						if( !cellpixelsbyid[x[1]] ){
							cellpixelsbyid[x[1]] = []
						}
						cellpixelsbyid[x[1]].push( x[0] )
					}
				}*/

				this.getImageData();
				for (let cid of Object.keys(cellpixelsbyid)) {
					if (kind < 0 || this.C.cellKind(cid) === kind) {
						if (typeof col == "function") {
							this.col(col(cid));
						}
						for (let cp of cellpixelsbyid[cid]) {
							this.pxfi(cp);
						}
					}
				}
				this.putImageData();
			}
		}

		/** General drawing function to draw all pixels in a supplied set in a given
		 * color.
		 * @param {ArrayCoordinate[]} pixelarray - an array of
		 * {@link ArrayCoordinate}s of pixels to color.
		 * @param {HexColor|function} col - Optional: hex code for the color to use.
		 * If left unspecified, it gets the default value of black ("000000").
		 * col can also be a function that returns a hex value for a cell id.
		 * */
		drawPixelSet( pixelarray, col ){
			if( ! col ){
				col = "000000";
			}
			if( typeof col == "string" ){
				this.col(col);
			}
			this.getImageData();
			for( let p of pixelarray ){
				this.pxfi( p );
			}
			this.putImageData();
		}

		/** Draw grid to the png file "fname".
		 *
		 * @param {string} fname Path to the file to write. Any parent folders in
		 * this path must already exist.*/
		writePNG( fname ){

			try {
				this.fs.writeFileSync(fname, this.el.toBuffer());
			}
			catch (err) {
				if (err.code === "ENOENT") {
					let message = "Canvas.writePNG: cannot write to file " + fname +
						", are you sure the directory exists?";
					throw(message)
				}
			}

		}
	}

	/** Extension of the {@link GridBasedModel} class suitable for
	a Cellular Automaton (CA). Currently only supports synchronous CAs.

	@example <caption>Conway's Game of Life </caption>
	*	let CPM = require( "path/to/build" )
	*	let C = new CPM.CA( [200,200], {
	*		"UPDATE_RULE": 	function(p,N){
	*			let nalive = 0
	*			for( let pn of N ){
	*				nalive += (this.pixt(pn)==1)
	*			}	
	*			if( this.pixt(p) == 1 ){
	*				if( nalive == 2 || nalive == 3 ){
	*					return 1
	*				}
	*			} else {
	*				if( nalive == 3 ) return 1
	*			}
	*			return 0
	*		}
	*	})
	*	let initialpixels = [ [100,100], [101,100], [102,100], [102,101], [101,102] ]
	*	for( p of initialpixels ){
	*		C.setpix( p, 1 )
	* 	}
	*	// Run it.
	*	for( let t = 0; t < 10; t++ ){ C.timeStep() }

	@todo Include asynchronous updating scheme?
	*/
	class CA extends GridBasedModel {

		/** The constructor of class CA.
		@param {GridSize} extents - the size of the grid of the model.
		@param {object} conf - configuration options. 
		@param {boolean} [conf.torus=[true,true,...]] - should the grid have linked borders?
		@param {number} [seed] - seed for the random number generator. If left unspecified,
		a random number from the Math.random() generator is used to make one.
		@param {updatePixelFunction} conf.UPDATE_RULE - the update rule of the CA. 
		*/
		constructor( extents, conf ){
			super( extents, conf );
			/** Bind the supplied updaterule to the object.
			@type {updatePixelFunction}*/
			this.updateRule = conf["UPDATE_RULE"].bind(this);
		}

		/** A timestep in a CA just applies the update rule and clears any cached stats after
		doing so. */
		timeStep(){
			this.grid.applyLocally( this.updateRule );
			
			/** Cached values of these stats. Object with stat name as key and its cached
			value as value. The cache must be cleared when the grid changes!
			@type {object} */
			this.stat_values = {};
		}
	}

	/**	This Stat creates a {@link CellArrayObject} with the border cellpixels of each cell on the grid. 
		Keys are the {@link CellId} of cells on the grid, corresponding values are arrays
		containing the pixels belonging to that cell. Coordinates are stored as {@link ArrayCoordinate}.
		
		@example
		* let CPM = require( "path/to/build" )
		*
		* // Make a CPM, seed a cell, and get the BorderPixelsByCell
		* let C = new CPM.CPM( [100,100], { 
		* 	T:20,
		* 	J:[[0,20],[20,10]],
		* 	V:[0,200],
		* 	LAMBDA_V:[0,2]
		* } )
		* C.setpix( [50,50], 1 )
		* C.getStat( CPM.BorderPixelsByCell )
	*/
	class BorderPixelsByCell extends Stat {

		/** The set model function of BorderPixelsByCell requires an object of type CPM.
		@param {CPM} M The CPM to compute cellborderpixels of.*/
		set model( M ){
			if( M instanceof CPM ){
				/** The CPM to compute borderpixels for.
				@type {CPM} */
				this.M = M;
			} else {
				throw( "The stat BorderPixelsByCell is only implemented for CPMs, where cellborderpixels are stored!" )
			}
			
		}

		/** The compute method of BorderPixelsByCell creates an object with the borderpixels of
		each cell on the grid.
		@returns {CellArrayObject} An object with a key for each cell on the grid, and as
		corresponding value an array with all the borderpixels of that 
		cell. Each pixel is stored by its {@link ArrayCoordinate}.*/
		compute(){
			// initialize the object
			let cellborderpixels = { };
			
			// The this.M.cellBorderPixels() iterator returns coordinates and cellid for all 
			// non-background pixels on the grid. See the appropriate Grid class for
			// its implementation.
			for( let [p,i] of this.M.cellBorderPixels() ){
				if( !cellborderpixels[i] ){
					cellborderpixels[i] = [p];
				} else {
					cellborderpixels[i].push( p );
				}
			}
			return cellborderpixels
		}
	}

	/**	This Stat computes the centroid of a cell when grid has a torus. 

		!!! Assumption: cell pixels never extend for more than half the size of the grid.
		If this assumption does not hold, centroids may be computed wrongly.
		
		See also {@link Centroids} for a version without torus correction.
			
		@example
		* let CPM = require( "path/to/build" )
		*
		* // Make a CPM, seed two cells, run a little, and get their centroids
		* let C = new CPM.CPM( [100,100], { 
		* 	T:20,
		* 	torus:[true,true],
		* 	J:[[0,20],[20,10]],
		* 	V:[0,200],
		* 	LAMBDA_V:[0,2]
		* } )
		* let gm = new CPM.GridManipulator( C )
		* gm.seedCell(1)
		* gm.seedCell(1)
		* for( let t = 0; t < 100; t++ ){ C.timeStep() }
		*
		* C.getStat( CPM.CentroidsWithTorusCorrection ) 
	*/

	class CentroidsWithTorusCorrection extends Stat {

		/** The set model method of class CentroidsWithTorusCorrection.
		@param {GridBasedModel} M - the model to compute centroids on. */
		set model( M ){
		
			/** The model to compute centroids on. 
			@type {GridBasedModel}*/
			this.M = M;
			// Half the grid dimensions; if pixels with the same cellid are further apart,
			// we assume they are on the border of the grid and that we need to correct
			// their positions to compute the centroid.
			/** @ignore */
			this.halfsize = new Array( this.M.ndim).fill(0);
			for( let i = 0 ; i < this.M.ndim ; i ++ ){
				this.halfsize[i] = this.M.extents[i]/2;
			}
		}
		
		/** @ignore */
		constructor( conf ){
			super(conf);
		}
			
		/** This method computes the centroid of a specific cell with id = <cellid>. 
		The cellpixels object is given as an argument so that it only has to be requested
		once for all cells together.
		@param {CellId} cellid ID number of the cell to get centroid of. 
		@param {CellArrayObject} cellpixels object produced by {@link PixelsByCell}, 
		where keys are the cellids
		of all non-background cells on the grid, and the corresponding value is an array
		of the pixels belonging to that cell specified by their {@link ArrayCoordinate}.
		@return {ArrayCoordinate} the centroid of the current cell.
		*/
		computeCentroidOfCell( cellid, cellpixels  ){
		
			//let cellpixels = this.M.getStat( PixelsByCell ) 
		
			const pixels = cellpixels[ cellid ];
			
			// cvec will contain the x, y, (z) coordinate of the centroid.
			// Loop over the dimensions to compute each element separately.
			let cvec = new Array(this.M.ndim).fill(0);
			for( let dim = 0 ; dim < this.M.ndim ; dim ++ ){
				
				let mi = 0.;
				const hsi = this.halfsize[dim], si = this.M.extents[dim];
				
				// Loop over the pixels;
				// compute mean position per dimension with online algorithm
				for( let j = 0 ; j < pixels.length ; j ++ ){
					
					// Check distance of current pixel to the accumulated mean in this dim.
					// Check if this distance is greater than half the grid size in this
					// dimension; if so, this indicates that the cell has moved to the
					// other end of the grid because of the torus. Note that this only
					// holds AFTER the first pixel (so for j > 0), when we actually have
					// an idea of where the cell is.
					let dx = pixels[j][dim] - mi;
					if( this.M.grid.torus[dim] && j > 0 ){
						// If distance is greater than half the grid size, correct the
						// coordinate.
						if( dx > hsi ){
							dx -= si;
						} else if( dx < -hsi ){
							dx += si;
						}
					}
					// Update the mean with the appropriate weight. 
					mi += dx/(j+1);
				}
				
				// Correct the final position so that it falls in the current grid.
				// (Because of the torus, it can happen to get a centroid at eg x = -1. )
				if( mi < 0 ){
					mi += si;
				} else if( mi > si ){
					mi -= si;
				}
				
				// Set the mean position in the cvec vector.
				cvec[dim] = mi;
			}
			return cvec
			
		}
			
		/** This method computes the centroids of all cells on the grid. 
		@return {CellObject} with an {@link ArrayCoordinate} of the centroid for each cell
		 on the grid (see {@link computeCentroidOfCell}). */
		compute(){
			// Get object with arrays of pixels for each cell on the grid, and get
			// the array for the current cell.
			let cellpixels = this.M.getStat( PixelsByCell ); 
			
			// Create an object for the centroids. Add the centroid array for each cell.
			let centroids = {};
			for( let cid of this.M.cellIDs() ){
				centroids[cid] = this.computeCentroidOfCell( cid, cellpixels );
			}
			
			return centroids
			
		}
	}

	/**	This Stat computes the centroid of a cell. When the cell resides on a torus, the
		centroid may be well outside the cell, and other stats may be preferable (e.g.
		{@link CentroidsWithTorusCorrection}).
		
		@example
		* let CPM = require( "path/to/build" )
		*
		* // Make a CPM, seed two cells, run a little, and get their centroids
		* let C = new CPM.CPM( [100,100], { 
		* 	T:20,
		*	torus:[false,false],
		* 	J:[[0,20],[20,10]],
		* 	V:[0,200],
		* 	LAMBDA_V:[0,2]
		* } )
		* let gm = new CPM.GridManipulator( C )
		* gm.seedCell(1)
		* gm.seedCell(1)
		* for( let t = 0; t < 100; t++ ){ C.timeStep() }
		*
		* C.getStat( CPM.Centroids ) 
	*/
	class Centroids extends Stat {

		/** The set model method of class CentroidsWithTorusCorrection.
		@param {GridBasedModel} M - the model to compute centroids on. */
		set model( M ){
		
			/** The model to compute centroids on. 
			@type {GridBasedModel}*/
			this.M = M;
			
			/* Check if the grid has a torus; if so, warn that this method may not be
			appropriate. */
			let torus = false;
			for( let d = 0; d < this.M.ndim; d++ ){
				if( this.M.grid.torus[d] ){
					torus = true;
					break
				}
			}
			
			if(torus){
				// eslint-disable-next-line no-console
				console.warn( "Your model grid has a torus, and the 'Centroids' stat is not compatible with torus! Consider using 'CentroidsWithTorusCorrection' instead." );
			}
			
			// Half the grid dimensions; if pixels with the same cellid are further apart,
			// we assume they are on the border of the grid and that we need to correct
			// their positions to compute the centroid.
			/** @ignore */
			this.halfsize = new Array( this.M.ndim).fill(0);
			for( let i = 0 ; i < this.M.ndim ; i ++ ){
				this.halfsize[i] = this.M.extents[i]/2;
			}
		}
		/** @ignore */
		constructor( conf ){
			super(conf);
		}
		/** This method computes the centroid of a specific cell. 
			@param {CellId} cellid the unique cell id of the cell to get centroid of.
			@param {CellArrayObject} cellpixels object produced by {@link PixelsByCell}, 
			with keys for each cellid
			and as corresponding value the pixel coordinates of their pixels.
			@returns {ArrayCoordinate} coordinate of the centroid.
		*/
		computeCentroidOfCell( cellid, cellpixels  ){
		
			//let cellpixels = this.M.getStat( PixelsByCell ) 
		
			const pixels = cellpixels[ cellid ];
			
			// cvec will contain the x, y, (z) coordinate of the centroid.
			// Loop over the dimensions to compute each element separately.
			let cvec = new Array(this.M.ndim).fill(0);
			for( let dim = 0 ; dim < this.M.ndim ; dim ++ ){
				
				let mi = 0.;
				// Loop over the pixels;
				// compute mean position per dimension with online algorithm
				for( let j = 0 ; j < pixels.length ; j ++ ){
					// Check distance of current pixel to the accumulated mean in this dim.
					// Check if this distance is greater than half the grid size in this
					// dimension; if so, this indicates that the cell has moved to the
					// other end of the grid because of the torus. Note that this only
					// holds AFTER the first pixel (so for j > 0), when we actually have
					// an idea of where the cell is.
					let dx = pixels[j][dim] - mi;
					// Update the mean with the appropriate weight. 
					mi += dx/(j+1);
				}			
				// Set the mean position in the cvec vector.
				cvec[dim] = mi;
			}
			return cvec
			
		}
			
		/** Compute centroids for all cells on the grid. 
		@return {CellObject} with an {@link ArrayCoordinate} of the centroid for each cell
		 on the grid (see {@link computeCentroidOfCell}). */
		compute(){
			// Get object with arrays of pixels for each cell on the grid, and get
			// the array for the current cell.
			let cellpixels = this.M.getStat( PixelsByCell ); 
			
			// Create an object for the centroids. Add the centroid array for each cell.
			let centroids = {};
			for( let cid of this.M.cellIDs() ){
				centroids[cid] = this.computeCentroidOfCell( cid, cellpixels );
			}
			return centroids
		}
	}

	/** This stat computes a list of all cell ids of the cells that border to "cell" and 
		belong to a different cellid, also giving the interface length for each contact. 
		@experimental 
		@example
		* let CPM = require("path/to/build")
		* 
		* // Set up a CPM and manipulator
		* let C = new CPM.CPM( [300,300], {
		* 	T:20, 
		* 	torus:[false,false],
		* 	J:[[0,20],[20,10]], 
		* 	V:[0,200], 
		* 	LAMBDA_V:[0,2]
		* })
		* let gm = new CPM.GridManipulator( C )
		* 
		* // Seed a cells, run a little, then divide it
		* gm.seedCell(1)
		* for( let t = 0; t < 50; t++ ){
		* 	C.timeStep()
		* }
		* gm.divideCell(1)
		* 
		* // Get neighborlist
		* console.log(  C.getStat( CPM.CellNeighborList ) )	
		*/	
	class CellNeighborList extends Stat {


		/** The set model function of CellNeighborList requires an object of type CPM.
		@param {CPM} M The CPM to compute bordering cells for.*/
		set model( M ){
			if( M instanceof CPM ){
				/** The CPM to compute borderpixels for.
				@type {CPM} */
				this.M = M;
			} else {
				throw( "The stat CellNeighborList is only implemented for CPMs, where cellborderpixels are stored!" )
			}
			
		}

		/** The getNeighborsOfCell method of CellNeighborList computes a list of all pixels
			that border to "cell" and belong to a different cellid.
			@param {CellId} cellid the unique cell id of the cell to get neighbors from.
			@param {CellArrayObject} cellborderpixels object produced by {@link BorderPixelsByCell}, with keys for each cellid
			and as corresponding value the border pixel indices of their pixels.
			@returns {CellObject} a dictionairy with keys = neighbor cell ids, and 
			values = number of neighbor cellpixels at the border.
		*/
		getNeighborsOfCell( cellid, cellborderpixels ){
					
			let neigh_cell_amountborder = { };
			let cbp = cellborderpixels[cellid];
			
			//loop over border pixels of cell
			for ( let cellpix = 0; cellpix < cbp.length; cellpix++ ) {

				//get neighbouring pixels of borderpixel of cell
				let neighbours_of_borderpixel_cell = this.M.neigh( cbp[cellpix] );

				//don't add a pixel in cell more than twice
				//loop over neighbouring pixels and store the parent cell if it is different from
				//cell, add or increment the key corresponding to the neighbor in the dictionairy
				for ( let neighborpix of neighbours_of_borderpixel_cell ) {
					
					let neighbor_id = this.M.pixt( neighborpix );

					if (neighbor_id != cellid) {
						neigh_cell_amountborder[neighbor_id] = neigh_cell_amountborder[neighbor_id]+1 || 1;
					}
				}
			}
			return neigh_cell_amountborder

		
		}
		
		/** The compute method of CellNeighborList computes for each cell on the grid 
			a list of all pixels at its border that belong to a different cellid.
			@returns {CellObject} a dictionairy with keys = cell ids, and 
			values = an object produced by {@link getNeighborsOfCell} (which has keys for each
			neighboring cellid and values the number of contacting pixels for that cell).
		*/
		compute(){
			
			let cellborderpixels = this.M.getStat( PixelsByCell );
			let neighborlist = {};
			
			// the this.M.cellIDs() iterator returns non-background cellids on the grid.
			for( let i of this.M.cellIDs() ){
				neighborlist[i] = this.getNeighborsOfCell( i, cellborderpixels );
			}
			
			return neighborlist

		}
	}

	/** This Stat creates an object with the connected components of each cell on the grid. 
		Keys are the {@link CellId} of all cells on the grid, corresponding values are objects
		where each element is a connected component. Each element of that array contains 
		the {@link ArrayCoordinate} for that pixel.
		
		@example
		* let CPM = require( "path/to/build" )
		*
		* // Make a CPM, seed a cell, and get the ConnectedComponentsByCell
		* let C = new CPM.CPM( [100,100], { 
		* 	T:20,
		* 	J:[[0,20],[20,10]],
		* 	V:[0,200],
		* 	LAMBDA_V:[0,2]
		* } )
		* let gm = new CPM.GridManipulator( C )
		* gm.seedCell(1)
		* gm.seedCell(1)
		* for( let t = 0; t < 100; t++ ){ C.timeStep() }
		* C.getStat( CPM.ConnectedComponentsByCell )
	*/
	class ConnectedComponentsByCell extends Stat {

		/** This method computes the connected components of a specific cell. 
			@param {CellId} cellid the unique cell id of the cell to get connected components of.
			@returns {object} object of cell connected components. These components in turn consist of the pixels 
		(specified by {@link ArrayCoordinate}) belonging to that cell.
		*/
		connectedComponentsOfCell( cellid ){
		
			const cbp = this.M.getStat( PixelsByCell );
			const cbpi = cbp[cellid];
			let M = this.M;
		
		
			let visited = {}, k=0, pixels = [];
			let labelComponent = function(seed, k){
				let q = [seed];
				visited[q[0]] = 1;
				pixels[k] = [];
				while( q.length > 0 ){
					let e = q.pop();
					pixels[k].push( M.grid.i2p(e) );
					let ne = M.grid.neighi( e );
					for( let i = 0 ; i < ne.length ; i ++ ){
						if( M.pixti( ne[i] ) == cellid &&
							!(ne[i] in visited) ){
							q.push(ne[i]);
							visited[ne[i]]=1;
						}
					}
				}
			};
			for( let i = 0 ; i < cbpi.length ; i ++ ){
				let pi = this.M.grid.p2i( cbpi[i] );
				if( !(pi in visited) ){
					labelComponent( pi, k );
					k++;
				}
			}
			return pixels
		}

		/** The compute method of ConnectedComponentsByCell creates an object with 
		connected components of the border of each cell on the grid.
		@return {CellObject} object with for each cell on the grid
		an object of components. These components in turn consist of the pixels 
		(specified by {@link ArrayCoordinate}) belonging to that cell.
		*/
		compute(){
			// initialize the object
			let components = { };
			// The this.M.pixels() iterator returns coordinates and cellid for all 
			// non-background pixels on the grid. See the appropriate Grid class for
			// its implementation.
			for( let ci of this.M.cellIDs() ){
				components[ci] = this.connectedComponentsOfCell( ci );
			}
			return components
		}
	}

	/** This Stat computes the 'connectedness' of cells on the grid. 
		Keys are the {@link CellId} of all cells on the grid, corresponding values the
		connectedness of the corresponding cell. 
		
		@example
		* let CPM = require( "path/to/build" )
		*
		* // Make a CPM, seed a cell, and get the Connectedness
		* let C = new CPM.CPM( [100,100], { 
		* 	T:20,
		* 	J:[[0,20],[20,10]],
		* 	V:[0,200],
		* 	LAMBDA_V:[0,2]
		* } )
		* let gm = new CPM.GridManipulator( C )
		* gm.seedCell(1)
		* for( let t = 0; t < 100; t++ ){ C.timeStep() }
		* C.getStat( CPM.Connectedness )
	*/
	class Connectedness extends Stat {

		/** This method computes the connectedness of a specific cell. 
		@return {number} the connectedness value of this cell, a number between 0 and 1.
		*/
		connectednessOfCell( cellid ){
		
			let ccbc = this.M.getStat( ConnectedComponentsByCell );
			const v = ccbc[cellid];
		
			//let s = {}, r = {}, i, j
			let s = 0, r = 0;
			
			for( let comp in Object.keys( v ) ){
				let volume = v[comp].length;
				s += volume;
			}
			for( let comp in Object.keys( v ) ){
				let volume = v[comp].length;
				r += (volume/s)*(volume/s);
			}
			
			return r

		}

		/** The compute method of Connectedness creates an object with 
		connectedness of each cell on the grid.
		@return {CellObject} object with for each cell on the grid
		a connectedness value. 
		*/
		compute(){
			// initialize the object
			let connectedness = { };
			// The this.M.pixels() iterator returns coordinates and cellid for all 
			// non-background pixels on the grid. See the appropriate Grid class for
			// its implementation.
			for( let ci of this.M.cellIDs() ){
				connectedness[ci] = this.connectednessOfCell( ci );
			}
			return connectedness
		}
	}

	/** This class contains methods that should be executed once per Monte Carlo Step.
	 Examples are cell division, cell death etc.
	 
	 Methods are written for CPMs, but some of the methods may also apply to other
		models of class ({@link GridBasedModel}, e.g. the cell seeding methods) 
		or even a general grid ({@link Grid}, e.g. the {@link makePlane} and {@link changeKind} 
		methods).
		
	 @example
	 // Build CPM and attach a gridmanipulator 
	 let C = new CPM.CPM( [100,100], {T:20, J:[[0,20],[20,10]]} )
	 let gm = new CPM.GridManipulator( C )
	 */
	class GridManipulator {
		/** Constructor of class GridManipulator.
		@param {CPM|GridBasedModel|Grid} C - the model whose grid you wish to manipulate.
		Methods are written for CPMs, but some of the methods may also apply to other
		models of class ({@link GridBasedModel}, e.g. the cell seeding methods) 
		or even a general grid ({@link Grid}, e.g. the {@link makePlane} and {@link changeKind} 
		methods).
		*/
		constructor( C ){
			/** The model whose grid we are manipulating.
			@type {CPM|GridBasedModel|Grid}*/
			this.C = C;
		}
		
		/** @experimental
		 */
		killCell( cellid ){
			let cp = this.C.getStat( PixelsByCell );
			let cpi = cp[cellid];
			
			for( let p of cpi ){
				this.C.setpixi( this.C.grid.p2i(p), 0 );
			}
			
			// update stats
			if( "PixelsByCell" in this.C.stat_values ){
				delete this.C.stat_values["PixelsByCell"][cellid];
			}
		}
		
		/** Seed a new cell at a random position. Return 0 if failed, ID of new cell otherwise.
		 * Try a specified number of times, then give up if grid is too full. 
		 * The first cell will always be seeded at the midpoint of the grid. 
		 
		 See also {@link seedCellAt} to seed a cell on a predefined position.
		 
		 @param {CellKind} kind - what kind of cell should be seeded? This determines the CPM
		 parameters that will be used for that cell.
		 @param {number} [max_attempts = 10000] - number of tries allowed. The method will
		 attempt to seed a cell at a random position, but this will fail if the position is 
		 already occupied. After max_attempts fails, it will not try again. This can happen
		 if the grid is very full.
		 @return {CellId} - the {@link CellId} of the newly seeded cell, or 0 if the seeding
		 has failed.
		 
		 @example
		 * // Build CPM and attach a gridmanipulator
		 * let C = new CPM.CPM( [100,100], {T:20, J:[[0,20],[20,10]]} )
		 * let gm = new CPM.GridManipulator( C )
		 *
		 * // Seed some cells
		 * gm.seedCell( 1 )
		 * gm.seedCell( 1 )
		 *
		 * // Check which pixels are nonzero. One is always the grid midpoint.
		 * for( let p of C.pixels() ){
		 * 	console.log( p )
		 * }
		 */
		seedCell( kind, max_attempts = 10000 ){
			let p = this.C.midpoint;
			while( this.C.pixt( p ) != 0 && max_attempts-- > 0 ){
				for( let i = 0 ; i < p.length ; i ++ ){
					p[i] = this.C.ran(0,this.C.extents[i]-1);
				}
			}
			if( this.C.pixt(p) != 0 ){
				return 0 // failed
			}
			const newid = this.C.makeNewCellID( kind );
			this.C.setpix( p, newid );
			return newid
		}
		/**  Seed a new cell of celltype "kind" onto position "p".
			This succeeds regardless of whether there is already a cell there.
			
			See also {@link seedCell} to seed a cell on a random position.
			
			@param {CellKind} kind - what kind of cell should be seeded? This determines the CPM
			parameters that will be used for that cell.
			@param {ArrayCoordinate} p - position to seed the cell at. 
			@return {CellId} - the {@link CellId} of the newly seeded cell, or 0 if the seeding
			has failed.
			
		 @example
		 * // Build CPM and attach a gridmanipulator
		 * let C = new CPM.CPM( [100,100], {T:20, J:[[0,20],[20,10]]} )
		 * let gm = new CPM.GridManipulator( C )
		 *
		 * // Seed some cells
		 * gm.seedCellAt( 1, [2,4] )
		 * gm.seedCellAt( 1, [9,3] )
		 *
		 * // Check which pixels are nonzero. These should be the positions defined above.
		 * for( let p of C.pixels() ){
		 * 	console.log( p )
		 * }
		 */
		seedCellAt( kind, p ){
			const newid = this.C.makeNewCellID( kind );
			this.C.grid.checkOnGrid(p);
			this.C.setpix( p, newid );
			return newid
		}
		
		/**  Seed "n" cells of celltype "kind" at random points lying within a circle 
			surrounding "center" with radius "radius". 
			
			See also {@link seedCell} to seed a cell on a random position in the entire grid,
			and {@link seedCellAt} to seed a cell at a specific position.
			
			@param {CellKind} kind - what kind of cell should be seeded? This determines the CPM
			parameters that will be used for that cell.
			@param {number} n - the number of cells to seed (must be integer).
			@param {ArrayCoordinate} center - position on the grid where the center of the
			circle should be.
			@param {number} radius - the radius of the circle to seed cells in.
			@param {number} max_attempts - the maximum number of attempts to seed a cell.
			Seeding can fail if the randomly chosen position is outside the circle, or if 
			there is already a cell there. After max_attempts the method will stop trying
			and throw an error.
			
		 @example
		 * // Build CPM and attach a gridmanipulator
		 * let C = new CPM.CPM( [100,100], {T:20, J:[[0,20],[20,10]]} )
		 * let gm = new CPM.GridManipulator( C )
		 *
		 * // Seed cells within a circle
		 * gm.seedCellsInCircle( 1, 10, [50,30], 20 )
		 *
		 * // Check which pixels are nonzero. These should be within the circle.
		 * for( let p of C.pixels() ){
		 * 	console.log( p )
		 * }
		 */
		seedCellsInCircle( kind, n, center, radius, max_attempts = 10000 ){
			if( !max_attempts ){
				max_attempts = 10*n;
			}
			let C = this.C;
			while( n > 0 ){
				if( --max_attempts == 0 ){
					throw("too many attempts to seed cells!")
				}
				let p = center.map( function(i){ return C.ran(Math.ceil(i-radius),Math.floor(i+radius)) } );
				let d = 0;
				for( let i = 0 ; i < p.length ; i ++ ){
					d += (p[i]-center[i])*(p[i]-center[i]);
				}
				if( d < radius*radius && this.C.pixt(p) == 0 ){
					this.seedCellAt( kind, p );
					n--;
				}
			}
		}
		/** Helper method to set an entire plane or line of pixels to a certain CellId at once.
		The method takes an existing array of coordinates (which can be empty) and adds the pixels
		of the specified plane to it. See {@link changeKind} for a method that sets such a 
		pixel set to a new value.
		
		The plane is specified by fixing one coordinate (x,y,or z) to a fixed value, and
		letting the others range from their min value 0 to their max value.
		
		@param {ArrayCoordinate[]} voxels - Existing array of pixels; this can be empty [].
		@param {number} coord - the dimension to fix the coordinate of: 0 = x, 1 = y, 2 = z.
		@param {number} coordvalue - the value of the coordinate in the fixed dimension; location
		of the plane.
		@return {ArrayCoordinate[]} the updated array of pixels. 
		
		@example
		* let C = new CPM.CPM( [10,10], {T:20, J:[[0,20],[20,10]]} )
		* let gm = new CPM.GridManipulator( C )
		* let myline = gm.makePlane( [], 0, 2 )
		* gm.changeKind( myline, 1 )
		*/
		makePlane ( voxels, coord, coordvalue ){
			let x,y,z;
			let minc = [0,0,0];
			let maxc = [0,0,0];
			for( let dim = 0; dim < this.C.ndim; dim++ ){
				maxc[dim] = this.C.extents[dim]-1;
			}
			minc[coord] = coordvalue;
			maxc[coord] = coordvalue;

			// For every coordinate x,y,z, loop over all possible values from min to max.
			// one of these loops will have only one iteration because min = max = coordvalue.
			for( x = minc[0]; x <= maxc[0]; x++ ){
				for( y = minc[1]; y<=maxc[1]; y++ ){
					for( z = minc[2]; z<=maxc[2]; z++ ){
						if( this.C.ndim == 3 ){
							voxels.push( [x,y,z] );	
						} else {
							//console.log(x,y)
							voxels.push( [x,y] );
						}
					}
				}
			}

			return voxels
		}
		/** Helper method that converts all pixels in a given array to a specific cellkind:
		   changes the pixels defined by voxels (array of coordinates p) into
		   the given cellkind. 
		
		@param {ArrayCoordinate[]} voxels - Array of pixels to change.
		@param {CellKind} cellkind - cellkind to change these pixels into.
		   
		@example
		* let C = new CPM.CPM( [10,10], {T:20, J:[[0,20],[20,10]]} )
		* let gm = new CPM.GridManipulator( C )
		* let myline = gm.makePlane( [], 0, 2 )
		* gm.changeKind( myline, 1 )
		*/
		changeKind ( voxels, cellkind ){
			
			let newid = this.C.makeNewCellID( cellkind );
			for( let p of voxels ){
				this.C.setpix( p, newid );
			}
			
		}

		/** Let cell "id" divide by splitting it along a line perpendicular to
		 * its major axis. 
		 
		 @param {CellId} id - the id of the cell that needs to divide.
		 @return {CellId} the id of the newly generated daughter cell.
		   
			@example
			* let C = new CPM.CPM( [10,10], {
			* 	T:20, 
			*	J:[[0,20],[20,10]], 
			*	V:[0,200], 
			*	LAMBDA_V:[0,2] 
			* })
			* let gm = new CPM.GridManipulator( C )
			*
			* // Seed a single cell
			* gm.seedCell( 1 )
			* 
			* // Perform some Monte Carlo Steps before dividing the cell
			* for( let t = 0; t < 100; t++ ){
			* 	C.timeStep()
			* }
			* gm.divideCell( 1 )
			* 
			* // Check which pixels belong to which cell. Should be roughly half half.
			* C.getStat( PixelsByCell )
		 */
		divideCell( id ){
			let C = this.C;
			let torus = C.conf.torus.indexOf(true) >= 0;
			if( C.ndim != 2 || torus ){
				throw("The divideCell methods is only implemented for 2D non-torus lattices yet!")
			}
			let cp = C.getStat( PixelsByCell )[id], com = C.getStat( Centroids )[id];
			let bxx = 0, bxy = 0, byy=0, cx, cy, x2, y2, side, T, D, x0, y0, x1, y1, L2;

			// Loop over the pixels belonging to this cell
			for( let j = 0 ; j < cp.length ; j ++ ){
				cx = cp[j][0] - com[0]; // x position rel to centroid
				cy = cp[j][1] - com[1]; // y position rel to centroid

				// sum of squared distances:
				bxx += cx*cx;
				bxy += cx*cy;
				byy += cy*cy;
			}

			// This code computes a "dividing line", which is perpendicular to the longest
			// axis of the cell.
			if( bxy == 0 ){
				x0 = 0;
				y0 = 0;
				x1 = 1;
				y1 = 0;
			} else {
				T = bxx + byy;
				D = bxx*byy - bxy*bxy;
				//L1 = T/2 + Math.sqrt(T*T/4 - D)
				L2 = T/2 - Math.sqrt(T*T/4 - D);
				x0 = 0;
				y0 = 0;
				x1 = L2 - byy;
				y1 = bxy;
			}

			// create a new ID for the second cell
			let nid = C.makeNewCellID( C.cellKind( id ) );

			// Loop over the pixels belonging to this cell
			//let sidea = 0, sideb = 0
			//let pix_id = []
			//let pix_nid = []
			//let sidea = 0, sideb=0

			for( let j = 0 ; j < cp.length ; j ++ ){
				// coordinates of current cell relative to center of mass
				x2 = cp[j][0]-com[0];
				y2 = cp[j][1]-com[1];

				// Depending on which side of the dividing line this pixel is,
				// set it to the new type
				side = (x1 - x0)*(y2 - y0) - (x2 - x0)*(y1 - y0);
				if( side > 0 ){
					//sidea++
					C.setpix( cp[j], nid ); 
					// console.log( cp[j] + " " + C.cellKind( id ) )
					//pix_nid.push( cp[j] )
				}
			}
			//console.log( "3 " + C.cellKind( id ) )
			//cp[id] = pix_id
			//cp[nid] = pix_nid
			C.stat_values = {}; // remove cached stats or this will crash!!!
			return nid
		}
	}

	/**
	 * This is a constraint in which each cell has a preferred direction of migration. 
	 * This direction is only dependent on the cell, not on the specific pixel of a cell.
	 * 
	 * This constraint works with torus as long as the field size is large enough. 
	 */
	class PersistenceConstraint extends SoftConstraint {

		/** The constructor of the PersistenceConstraint requires a conf object with parameters.
		@param {object} conf - parameter object for this constraint
		@param {PerKindNonNegative} conf.LAMBDA_DIR - strength of the persistenceconstraint per cellkind.
		@param {PerKindNonNegative} [conf.DELTA_T = [10,10,...]] - the number of MCS over which the current direction is 
		determined. Eg if DELTA_T = 10 for a cellkind, then its current direction is given by
		the vector from its centroid 10 MCS ago to its centroid now.
		@param {PerKindProb} conf.PERSIST - persistence per cellkind. If this is 1, its new
		target direction is exactly equal to its current direction. If it is lower than 1, 
		angular noise is added accordingly. 
		*/
		constructor( conf ){
			super( conf );
			
			/** Cache centroids over the previous conf.DELTA_T MCS to determine directions.
			@type {CellObject}
			*/
			this.cellcentroidlists = {};
			/** Target direction of movement of each cell.
			@type {CellObject}
			*/
			this.celldirections = {};
		}
		
		/** Set the CPM attached to this constraint.
		@param {CPM} C - the CPM to attach.*/
		set CPM(C){
		
			/** @ignore */
			this.halfsize = new Array(C.ndim).fill(0);
			
			/** The CPM this constraint acts on.
			@type {CPM}*/
			this.C = C;
			for( let i = 0 ; i < C.ndim ; i ++ ){
				this.halfsize[i] = C.extents[i]/2;
			}
			this.confChecker();
		}
		
		/** This method checks that all required parameters are present in the object supplied to
		the constructor, and that they are of the right format. It throws an error when this
		is not the case.*/
		confChecker(){
			let checker = new ParameterChecker( this.conf, this.C );
			checker.confCheckParameter( "LAMBDA_DIR", "KindArray", "NonNegative" );
			checker.confCheckParameter( "PERSIST", "KindArray", "Probability" );
		}
		
		/** Method to compute the Hamiltonian for this constraint. 
		 @param {IndexCoordinate} sourcei - coordinate of the source pixel that tries to copy.
		 @param {IndexCoordinate} targeti - coordinate of the target pixel the source is trying
		 to copy into.
		 @param {CellId} src_type - cellid of the source pixel.
		 @param {CellId} tgt_type - cellid of the target pixel. This argument is not used by this
		 method, but is supplied for consistency with other SoftConstraints. The CPM will always
		 call this method supplying the tgt_type as fourth argument.
		 @return {number} the change in Hamiltonian for this copy attempt and this constraint.*/ 
		/* eslint-disable no-unused-vars*/
		deltaH ( sourcei, targeti, src_type, tgt_type ) {
			if( src_type == 0 || !(src_type in this.celldirections) ) return 0
			let b = this.celldirections[src_type];
			let p1 = this.C.grid.i2p(sourcei), p2 = this.C.grid.i2p(targeti);
			let a = [];
			for( let i = 0 ; i < p1.length ; i ++ ){
				a[i] = p2[i]-p1[i];
				// Correct for torus if necessary
				if( this.C.grid.torus[i] ){
					if( a[i] > this.halfsize[i] ){
						a[i] -= this.C.extents[i];
					} else if( a[i] < -this.halfsize[i] ){
						a[i] += this.C.extents[i];
					}
				}
			}
			let dp = 0;
			for( let i = 0 ; i < a.length ; i ++ ){
				dp += a[i]*b[i];
			}
			return - dp
		}
		
		/** Normalize a vector a by its length.
		@param {number[]} a - vector to normalize.
		@return {number[]} normalized version of this vector.
		@private*/
		normalize( a ){
			let norm = 0;
			for( let i = 0 ; i < a.length ; i ++ ){
				norm += a[i]*a[i];
			}
			norm = Math.sqrt(norm);
			for( let i = 0 ; i < a.length ; i ++ ){
				a[i] /= norm;
			}
		}
		/** this function samples a random number from a normal distribution
		@param {number} [mu=0] - mean of the normal distribution.
		@param {number} [sigma=1] - SD of the normal distribution.
		@return {number} the random number generated.
		@private
		*/
		sampleNorm (mu=0, sigma=1) {
			let u1 = this.C.random();
			let u2 = this.C.random();
			let z0 = Math.sqrt(-2.0 * Math.log(u1)) * Math.cos(Math.PI*2 * u2);
			return z0 * sigma + mu
		}
		/** This function samples a random direction vector with length 1
		@param {number} [n=3] - number of dimensions of the space to make the vector in.
		@return {number[]} - a normalized direction vector.
		*/
		randDir (n=3) {
			let dir = [];
			while(n-- > 0){
				dir.push(this.sampleNorm());
			}
			this.normalize(dir);
			return dir
		}
		/** Change the target direction of a cell to a given vector.
			@param {CellId} t - the cellid of the cell to change the direction of.
			@param {number[]} dx - the new direction this cell should get.
		*/
		setDirection( t, dx ){
			this.celldirections[t] = dx;
		}
		
		/** After each MCS, update the target direction of each cell based on its actual
		direction over the last {conf.DELTA_T[cellkind]} steps, and some angular noise
		depending on {conf.PERSIST[cellkind]}.
		@listens {CPM#timeStep} because when the CPM has finished an MCS, cells have new 
		centroids and their direction must be updated. 
		*/
		postMCSListener(){
			let centroids;
			if( this.C.conf.torus ){
				centroids = this.C.getStat( CentroidsWithTorusCorrection );
			} else {
				centroids = this.C.getStat( Centroids );
			}
			for( let t of this.C.cellIDs() ){
				const k = this.C.cellKind(t);
				let ld = this.conf["LAMBDA_DIR"][k];
				let dt = this.conf["DELTA_T"] && this.conf["DELTA_T"][k] ? 
					this.conf["DELTA_T"][k] : 10;
				if( ld == 0 ){
					delete this.cellcentroidlists[t];
					delete this.celldirections[t];
					continue
				}
				if( !(t in this.cellcentroidlists ) ){
					this.cellcentroidlists[t] = [];
					this.celldirections[t] = this.randDir(this.C.ndim);
				}

				let ci = centroids[t];
				this.cellcentroidlists[t].unshift(ci);
				if( this.cellcentroidlists[t].length >= dt ){
					// note, dt could change during execution
					let l;
					while( this.cellcentroidlists[t].length >= dt ){
						l = this.cellcentroidlists[t].pop();
					}
					let dx = [];
					for( let j = 0 ; j < l.length ; j ++ ){
						dx[j] = ci[j] - l[j];
						
						// torus correction; do only if CPM actually has a torus in this dimension.
						if( this.C.grid.torus[j] ){
							if( dx[j] > this.halfsize[j] ){
								dx[j] -= this.C.extents[j];
							} else if( dx[j] < -this.halfsize[j] ){
								dx[j] += this.C.extents[j];
							}					
						}
					}
					// apply angular diffusion to target direction if needed
					let per = this.conf["PERSIST"][k];
					if( per < 1 ){
						this.normalize(dx);
						this.normalize(this.celldirections[t]);
						for( let j = 0 ; j < dx.length ; j ++ ){
							dx[j] = (1-per)*dx[j] + per*this.celldirections[t][j];
						}
						this.normalize(dx);
						for( let j = 0 ; j < dx.length ; j ++ ){
							dx[j] *= ld;
						}
						this.celldirections[t] = dx;
					}
				}
			}
		}
	}

	/** Implements a global bias direction of motion.
		This constraint computes the *unnormalized* dot product 
		between copy attempt vector and target direction vector.

		Supply the target direction vector in normalized form, or 
		use the length of the vector to influence the strength 
		of this bias.

		Works for torus grids, if they are "big enough".
		
		 * @example
		 * // Build a CPM and add the constraint
		 * let CPM = require( "path/to/build" )
		 * let C = new CPM.CPM( [200,200], { T : 20 } )
		 * C.add( new CPM.PreferredDirectionConstraint( {
		 * 	LAMBDA_DIR : [0,50], 
		 * 	DIR : [[0,0],[1,1]]
		 * } ) )
		*/
	class PreferredDirectionConstraint extends SoftConstraint {

		
		/** The constructor of the PreferredDirectionConstraint requires a conf object with parameters.
		@param {object} conf - parameter object for this constraint
		@param {PerKindNonNegative} conf.LAMBDA_DIR - strength of the constraint per cellkind.
		@param {ArrayCoordinate[]} conf.DIR 'vector' with the preferred direction. This is
		an array with the {@link ArrayCoordinate}s of the start and endpoints of this vector. 
		*/
		constructor(conf){
			super(conf);
		}
		
		/** This method checks that all required parameters are present in the object supplied to
		the constructor, and that they are of the right format. It throws an error when this
		is not the case.*/
		confChecker(){
			let checker = new ParameterChecker( this.conf, this.C );
			checker.confCheckParameter( "LAMBDA_DIR", "KindArray", "NonNegative" );
			
			// Custom check for the attractionpoint
			checker.confCheckPresenceOf( "DIR" );
			let pt = this.conf["DIR"];
			if( !( pt instanceof Array ) ){
				throw( "DIR must be an array with the start and end coordinate of the preferred direction vector!" )
			}
			for( let p of pt ){
			
				if( !checker.isCoordinate(p) ){
					throw("DIR elements must be coordinate arrays with the same dimensions as the grid!")
				}
			}
		}
		
		/** Method to compute the Hamiltonian for this constraint. 
		 @param {IndexCoordinate} src_i - coordinate of the source pixel that tries to copy.
		 @param {IndexCoordinate} tgt_i - coordinate of the target pixel the source is trying
		 to copy into.
		 @param {CellId} src_type - cellid of the source pixel.
		 @param {CellId} tgt_type - cellid of the target pixel. This argument is not actually
		 used but is given for consistency with other soft constraints; the CPM always calls
		 this method with four arguments.
		 @return {number} the change in Hamiltonian for this copy attempt and this constraint.*/ 
		/* eslint-disable no-unused-vars*/
		deltaH( src_i, tgt_i, src_type, tgt_type ){
			let l = this.conf["LAMBDA_DIR"][this.C.cellKind( src_type )];
			if( !l ){
				return 0
			}
			let torus = this.C.conf.torus;
			let dir = this.conf["DIR"][this.C.cellKind( src_type )];
			let p1 = this.C.grid.i2p( src_i ), p2 = this.C.grid.i2p( tgt_i );
			// To bias a copy attempt p1 -> p2 in the direction of vector 'dir'.
			let r = 0.;
			// loops over the coordinates x,y,(z)
			for( let i = 0; i < p1.length ; i++ ){
				let si = this.C.extents[i];
				// direction of the copy attempt on this coordinate is from p1 to p2
				let dx = p2[i] - p1[i];
				if( torus[i] ){
					// If distance is greater than half the grid size, correct the
					// coordinate.
					if( dx > si/2 ){
						dx -= si;
					} else if( dx < -si/2 ){
						dx += si;
					}
				}
				// direction of the gradient
				r += dx * dir[i]; 
			}
			return - r * l
		}
	}

	/** 
	 * This class implements a constraint for cells moving up a chemotactic gradient.
	 * It checks the chemokine gradient in the direction of the attempted copy, and 
	 * rewards copy attempts that go up the gradient. This effect is stronger when the
	 * gradient is steep. Copy attempts going to a lower chemokine value are punished.
	 * 
	 * The grid with the chemokine must be supplied, see the {@link constructor}.
	 *
	 * @example
	 * // Build a chemotaxis field
	 * let CPM = require( "path/to/build" )
	 * let chemogrid = new CPM.Grid2D( [200,200], [true,true], "Float32" )
	 * chemogrid.setpix( [100,100], 100 )
	 * 
	 * // Build a CPM with the constraint
	 * let C = new CPM.CPM( [200,200], {
	 * 	T : 20,
	 * 	J : [[0,20],[20,10]],
	 * 	V : [0,500],
	 * 	LAMBDA_V : [0,5]	
	 * })
	 * C.add( new CPM.ChemotaxisConstraint( {
	 * 	LAMBDA_CH : [0,5000],
	 * 	CH_FIELD : chemogrid
	 * } ) )
	 */
	class ChemotaxisConstraint extends SoftConstraint {

		/** Set the CPM attached to this constraint.
		@param {CPM} C - the CPM to attach.*/
		set CPM(C){
			/** The CPM this constraint acts on.
			@type {CPM}*/
			this.C = C;
			
			this.confChecker();
			this.checkField();
		}
		
		/** @todo add checks for dimensions, better check for type.*/
		checkField(){
			if( !( this.field instanceof CoarseGrid || this.field instanceof Grid2D ) ){
				throw( "CH_FIELD must be a CoarseGrid or a Grid2D!" )
			}
		}
		
		/** This method checks that all required parameters are present in the object supplied to
		the constructor, and that they are of the right format. It throws an error when this
		is not the case.*/
		confChecker(){
			let checker = new ParameterChecker( this.conf, this.C );
			checker.confCheckParameter( "LAMBDA_CH", "KindArray", "NonNegative" );

			// Custom check for the chemotactic field
			checker.confCheckPresenceOf( "CH_FIELD" );
		}

		/** The constructor of the ChemotaxisConstraint requires a conf object with a parameter
		and a chemotactic field.
		@todo what kinds of grids are allowed for the chemotactic field? Do we need to check
		somewhere that its properties "match" that of the CPM? (That is, the same resolution and
		torus properties)?
		@param {object} conf - parameter object for this constraint
		@param {PerKindNonNegative} conf.LAMBDA_CH - chemotactic sensitivity per cellkind.
		@param {CoarseGrid|Grid2D} conf.CH_FIELD - the chemotactic field where the chemokine lives.
		*/
		constructor( conf ){
			super( conf );
			
			/** @todo is this ever used? */
			this.conf = conf;
			/** The field where the chemokine lives.
			@type {CoarseGrid|Grid2D}*/
			this.field = conf.CH_FIELD;
			if( this.field instanceof CoarseGrid ){
				this.deltaH = this.deltaHCoarse;
			}
		}

		/** Method to compute the Hamiltonian if the chemotactic field is a {@link CoarseGrid}.
			This method is used instead of the regular {@link deltaH} whenever this is true.
		 @param {IndexCoordinate} sourcei - coordinate of the source pixel that tries to copy.
		 @param {IndexCoordinate} targeti - coordinate of the target pixel the source is trying
		 to copy into.
		 @param {CellId} src_type - cellid of the source pixel.
		 @param {CellId} tgt_type - cellid of the target pixel. This argument is not actually
		 used by this method, but is supplied for compatibility; the CPM will always call the
		 deltaH method with all 4 arguments.
		 @return {number} the change in Hamiltonian for this copy attempt and this constraint.
		 @private */
		/* eslint-disable no-unused-vars*/
		deltaHCoarse( sourcei, targeti, src_type, tgt_type ){
			let sp = this.C.grid.i2p( sourcei ), tp = this.C.grid.i2p( targeti );
			let delta = this.field.pixt( tp ) - this.field.pixt( sp );
			let lambdachem = this.conf["LAMBDA_CH"][this.C.cellKind(src_type)];
			return -delta*lambdachem
		}

		/** Method to compute the Hamiltonian for this constraint. 
		 @param {IndexCoordinate} sourcei - coordinate of the source pixel that tries to copy.
		 @param {IndexCoordinate} targeti - coordinate of the target pixel the source is trying
		 to copy into.
		 @param {CellId} src_type - cellid of the source pixel.
		 @param {CellId} tgt_type - cellid of the target pixel. This argument is not actually
		 used by this method, but is supplied for compatibility; the CPM will always call the
		 deltaH method with all 4 arguments.
		 @return {number} the change in Hamiltonian for this copy attempt and this constraint.*/
		/* eslint-disable no-unused-vars*/
		deltaH( sourcei, targeti, src_type, tgt_type  ){
			let delta = this.field.pixt( targeti ) - this.field.pixt( sourcei );
			let lambdachem = this.conf["LAMBDA_CH"][this.C.cellKind(src_type)];
			return -delta*lambdachem
		}
	}

	/**Implements bias of motion in the direction of a supplied "attraction point".
	 * This constraint computes the cosine of the angle alpha between the direction
	 * of the copy attempt (from source to target pixel), and the direction from the
	 * source to the attraction point. This cosine is 1 if these directions are
	 * aligned, 0 if they are perpendicular, and 1 if they are opposite.
	 * We take the negative (so that deltaH is negative for a copy attempt in the
	 * right direction), and modify the strength of this bias using the lambda
	 * parameter. The constraint only acts on copy attempts *from* the cell that
	 * is responding to the field; it does not take into account the target pixel
	 * (except for its location to determine the direction of the copy attempt).
	 *
	 * The current implementation works for torus grids as long as the grid size in
	 * each dimension is larger than a few pixels.
	 *
	 * Automatic adding of this constraint via the conf object is currently not
	 * supported, so you will have to add this constraint using the
	 * {@link CPM#add} method.
	 *
	 * @example
	 * // Build a CPM and add the constraint
	 * let CPM = require( "path/to/build" )
	 * let C = new CPM.CPM( [200,200], { T : 20 } )
	 * C.add( new CPM.AttractionPointConstraint( {
	 * 	LAMBDA_ATTRACTIONPOINT : [0,100],
	 * 	ATTRACTIONPOINT: [100,100],
	 * } ) )
	 *
	 * // We can even add a second one at a different location
	 * C.add( new CPM.AttractionPointConstraint( {
	 * 	LAMBDA_ATTRACTIONPOINT : [0,100],
	 * 	ATTRACTIONPOINT: [50,50],
	 * } ) )
	 */
	class AttractionPointConstraint extends SoftConstraint {

		/** The constructor of an AttractionPointConstraint requires a conf object
		 * with two parameters.
		 * @param {object} conf - parameter object for this constraint
		 * @param {PerKindNonNegative} conf.LAMBDA_ATTRACTIONPOINT - strength of
		 * the constraint per cellkind.
		 * @param {ArrayCoordinate} conf.ATTRACTIONPOINT coordinate of the
		 * attraction point.
		*/
		constructor( conf ){
			super( conf );		
		}
		
		/** This method checks that all required parameters are present in the
		 * bject supplied to the constructor, and that they are of the right format.
		 * It throws an error when this is not the case.*/
		confChecker(){
			let checker = new ParameterChecker( this.conf, this.C );
			checker.confCheckParameter( "LAMBDA_ATTRACTIONPOINT",
				"KindArray", "NonNegative" );
			
			// Custom check for the attractionpoint
			checker.confCheckPresenceOf( "ATTRACTIONPOINT" );
			let pt = this.conf["ATTRACTIONPOINT"];
			if( !checker.isCoordinate(pt) ){
				throw( "ATTRACTIONPOINT must be a coordinate array with the same " +
					"dimensions as the grid!" )
			}
		}

		/** Method to compute the Hamiltonian for this constraint.
		 * @param {IndexCoordinate} src_i - coordinate of the source pixel that
		 * tries to copy.
		 * @param {IndexCoordinate} tgt_i - coordinate of the target pixel the
		 * source is trying to copy into.
		 * @param {CellId} src_type - cellid of the source pixel.
		 * @param {CellId} tgt_type - cellid of the target pixel. This argument is
		 * not actually used but is given for consistency with other soft
		 * constraints; the CPM always calls this method with four arguments.
		 * @return {number} the change in Hamiltonian for this copy attempt and
		 * this constraint. */
		/* eslint-disable no-unused-vars*/
		deltaH( src_i, tgt_i, src_type, tgt_type ){

			// deltaH is only non-zero when the source pixel belongs to a cell with
			// an attraction point, so it does not act on copy attempts where the
			// background would invade the cell.
			let l = this.conf["LAMBDA_ATTRACTIONPOINT"][this.C.cellKind( src_type )];
			if( !l ){
				return 0
			}

			// To assess whether the copy attempt lies in the direction of the
			// attraction point, we must take into account whether the grid has
			// wrapped boundaries (torus; see below).
			let torus = this.C.conf.torus;

			// tgt is the attraction point; p1 is the source location and p2 is
			// the location of the target pixel.
			let tgt = this.conf["ATTRACTIONPOINT"][this.C.cellKind( src_type )];
			let p1 = this.C.grid.i2p( src_i ), p2 = this.C.grid.i2p( tgt_i );

			// To bias a copy attempt p1 -> p2 in the direction of vector 'dir'.
			// r will contain the dot product of the copy attempt vector and the
			// vector pointing from the source pixel to the attraction point.
			// The copy attempt vector always has length one, but the vector to the
			// attraction point has a variable length that will be stored in ldir
			// (actually, we store the squared length).
			let r = 0., ldir = 0.;

			// loops over the coordinates x,y,(z)
			for( let i = 0; i < p1.length ; i++ ){

				// compute the distance between the target and the current position
				// in this dimension, and add it in squared form to the total.
				let dir_i = tgt[i] - p1[i];
				ldir += dir_i * dir_i;

				// similarly, the distance between the source and target pixel in this
				// dimension (direction of the copy attempt is from p1 to p2)
				let dx = p2[i] - p1[i];

				// we may have to correct for torus if a copy attempt crosses the
				// boundary.
				let si = this.C.extents[i];
				if( torus[i] ){
					// If distance is greater than half the grid size, correct the
					// coordinate.
					if( dx > si/2 ){
						dx -= si;
					} else if( dx < -si/2 ){
						dx += si;
					}
				}

				// direction of the gradient; add contribution of the current
				// dimension to the dot product.
				r += dx * dir_i; 
			}

			// divide dot product by squared length of directional vector to obtain
			// cosine of the angle between the copy attempt direction and the
			// direction to the attraction point. This cosine is 1 if they are
			// perfectly aligned, 0 if they are perpendicular, and negative
			// if the directions are opposite. Since we want to reward copy attempts
			// in the right direction, deltaH is the negative of this (and
			// multiplied by the lambda weight factor).
			return - r * l / Math.sqrt( ldir )
		}
	}

	/** This constraint enforces that cells stay 'connected' throughout any copy attempts.
	Copy attempts that break the cell into two parts are therefore forbidden. To speed things
	up, this constraint only checks if the borderpixels of the cells stay connected.
	@experimental
	*/
	class ConnectivityConstraint extends HardConstraint {

		/** The constructor of the ConnectivityConstraint requires a conf object with one parameter.
		@param {object} conf - parameter object for this constraint.
		@param {PerKindBoolean} conf.CONNECTED - should the cellkind be connected or not?
		*/
		constructor( conf ){
			super(conf);
			
			/** Object tracking the borderpixels of each cell. This is kept up to date after
			every copy attempt.
			@type {CellObject}*/
			this.borderpixelsbycell = {};
		}
		
		/** The set CPM method attaches the CPM to the constraint. */
		set CPM(C){
			/** CPM on which this constraint acts.
			@type {CPM}*/
			this.C = C;
			
			/** Private property used by {@link updateBorderPixels} to track borders. 
			@private
			@type {Uint16Array} */
			this._neighbours = new Uint16Array(this.C.grid.p2i(this.C.extents));
		}
		
		/** This method checks that all required parameters are present in the object supplied to
		the constructor, and that they are of the right format. It throws an error when this
		is not the case.*/
		confChecker(){
			let checker = new ParameterChecker( this.conf, this.C );
			checker.confCheckParameter( "CONNECTED", "KindArray", "Boolean" );
		}
		
		/** Update the borderpixels when pixel i changes from t_old into t_new.
		@param {IndexCoordinate} i - the pixel to change
		@param {CellId} t_old - the cell the pixel belonged to previously
		@param {CellId} t_new - the cell the pixel belongs to now. */
		updateBorderPixels( i, t_old, t_new  ){
			if( t_old == t_new ) return
			if( !(t_new in this.borderpixelsbycell) ){
				this.borderpixelsbycell[t_new] = {};
			}
			const Ni = this.C.grid.neighi( i );
			const wasborder = this._neighbours[i] > 0;
			// current neighbors of pixel i, set counter to zero and loop over nbh.
			this._neighbours[i] = 0;
			for( let ni of Ni  ){
				// type of the neighbor.
				const nt = this.C.grid.pixti(ni);
				
				// If type is not the t_new of pixel i, nbi++ because the neighbor belongs
				// to a different cell. 
				if( nt != t_new ){
					this._neighbours[i] ++; 
				}
				
				// If neighbor type is t_old, the border of t_old may have to be adjusted. 
				// It gets an extra neighbor because the current pixel becomes t_new.
				if( nt == t_old ){
					// If this wasn't a borderpixel of t_old, it now becomes one because
					// it has a neighbor belonging to t_new
					if( this._neighbours[ni] ++ == 0 ){
						if( !(t_old in this.borderpixelsbycell) ){
							this.borderpixelsbycell[t_old] = {};
						}
						this.borderpixelsbycell[t_old][ni] = true;
					}

				}
				// If type is t_new, the neighbor may no longer be a borderpixel
				if( nt == t_new ){
					if( --this._neighbours[ni] == 0 && ( ni in this.borderpixelsbycell[t_new] ) ){
						delete this.borderpixelsbycell[t_new][ni];
					}
				}
			}
			// Case 1: 
			// If current pixel is a borderpixel, add it to those of the current cell.
			if( this._neighbours[i] > 0 ){
				this.borderpixelsbycell[t_new][i]=true;
			}
			
			// Case 2:
			// Current pixel was a borderpixel. Remove from the old cell. 
			if( wasborder ){
				// It was a borderpixel from the old cell, but no longer belongs to that cell.
				if( i in this.borderpixelsbycell[t_old] ){ 
					delete this.borderpixelsbycell[t_old][i];
				}
			}
			
		}
		
		/** Get the connected components of the borderpixels of the current cell.
		@param {CellId} cellid - cell to check the connected components of.
		@return {object} an array with an element for every connected component, which is in
		turn an array of the {@link ArrayCoordinate}s of the pixels belonging to that component.  */
		connectedComponentsOfCellBorder( cellid ){
		
			/* Note that to get number of connected components, we only need to look at cellborderpixels. */
			if( !( cellid in this.borderpixelsbycell ) ){
				return []
			}
			
			//let cbpi = Object.keys( this.borderpixelsbycell[cellid] ), cbpobject = this.borderpixelsbycell[cellid]
			return this.connectedComponentsOf( this.borderpixelsbycell[cellid] )
				
			
		}
		
		
		/** Get the connected components of a set of pixels.
		@param {object} pixelobject - an object with as keys the {@link IndexCoordinate}s of the pixels to check.
		@return {object} an array with an element for every connected component, which is in
		turn an array of the {@link ArrayCoordinate}s of the pixels belonging to that component.  */
		connectedComponentsOf( pixelobject ){
		
			let cbpi = Object.keys( pixelobject );
			
			let visited = {}, k=0, pixels = [], C = this.C;
			let labelComponent = function(seed, k){
				let q = [seed];
				let cellid = C.pixti(q);
				visited[q[0]] = 1;
				pixels[k] = [];
				while( q.length > 0 ){
					let e = q.pop();
					pixels[k].push(C.grid.i2p(e) );
					let ne = C.grid.neighi( e );
					for( let i = 0 ; i < ne.length ; i ++ ){
						if( C.pixti( ne[i] ) == cellid &&
							!(ne[i] in visited) && (ne[i] in pixelobject) ){
							q.push(ne[i]);
							visited[ne[i]]=1;
						}
					}
				}
			};
			for( let i = 0 ; i < cbpi.length ; i ++ ){
				let pi = cbpi[i];
				if( !(pi in visited) ){
					labelComponent( pi, k );
					k++;
				}
			}
			return pixels
		}

		/** The postSetpixListener of the ConnectivityConstraint updates the internally
		tracked borderpixels after every copy.
		@param {IndexCoordinate} i - the pixel to change
		@param {CellId} t_old - the cell the pixel belonged to previously
		@param {CellId} t - the cell the pixel belongs to now.	
		*/
		postSetpixListener(  i, t_old, t ){
			this.updateBorderPixels( i, t_old, t );
		}	
		
		/** To speed things up: first check if a pixel change disrupts the local connectivity
		in its direct neighborhood. If local connectivity is not disrupted, we don't have to
		check global connectivity at all. This currently only works in 2D, so it returns 
		false for 3D (ensuring that connectivity is checked globally).
		@param {IndexCoordinate} tgt_i - the pixel to change
		@param {CellId} tgt_type - the cell the pixel belonged to before the copy attempt.
		@return {boolean} does the local neighborhood remain connected if this pixel changes?
		*/
		localConnected( tgt_i, tgt_type ){
		
			if( this.C.extents.length != 2 ){
				return false
			}
		
			let neighbors = 0;
			for( let i of this.C.grid.neighNeumanni(tgt_i) ){
				if( this.C.pixti(i) != tgt_type ){
					neighbors++;
				}
			}
			
			if( neighbors >= 2 ){
				return false
			}
			return true
			
		}
		
		/** This method checks if the connectivity still holds after pixel tgt_i is changed from
		tgt_type to src_type. 
		@param {IndexCoordinate} tgt_i - the pixel to change
		@param {CellId} src_type - the new cell for this pixel.
		@param {CellId} tgt_type - the cell the pixel belonged to previously. 	
		*/
		checkConnected( tgt_i, src_type, tgt_type ){
		
			// If local connectivity is preserved, global connectivity holds too.
			if( this.localConnected( tgt_i, tgt_type ) ){
				return true
			}
		
			// Otherwise, check connected components of the cell border. Before the copy attempt:
			let comp1 = this.connectedComponentsOfCellBorder( tgt_type );
			let length_before = comp1.length;
		
			// Update the borderpixels as if the change occurs
			this.updateBorderPixels( tgt_i, tgt_type, src_type );
			let comp = this.connectedComponentsOfCellBorder( tgt_type );
			let length_after = comp.length;
			
			// The src pixels copies its type, so the cell src_type gains a pixel. This
			// pixel is by definition connected because the copy happens from a neighbor.
			// So we only have to check if tgt_type remains connected
			let connected = true;
			if( length_after > length_before ){
				connected = false;
			}
			
			// Change borderpixels back because the copy attempt hasn't actually gone through yet.
			this.updateBorderPixels( tgt_i, src_type, tgt_type );
			
			return connected
			
		}

		/** Method for hard constraints to compute whether the copy attempt fulfills the rule.
		 @param {IndexCoordinate} src_i - coordinate of the source pixel that tries to copy.
		 @param {IndexCoordinate} tgt_i - coordinate of the target pixel the source is trying
		 to copy into.
		 @param {CellId} src_type - cellid of the source pixel.
		 @param {CellId} tgt_type - cellid of the target pixel. 
		 @return {boolean} whether the copy attempt satisfies the constraint.*/ 
		fulfilled( src_i, tgt_i, src_type, tgt_type ){
			// connectedness of src cell cannot change if it was connected in the first place.
			
			// connectedness of tgt cell
			if( tgt_type != 0 && this.conf["CONNECTED"][this.C.cellKind(tgt_type)] ){
				return this.checkConnected( tgt_i, src_type, tgt_type )
			}
			
			return true
		}
	}

	/** This constraint encourages that cells stay 'connected' throughout any copy attempts.
	In contrast to the hard version of the {@link ConnectivityConstraint}, this version does
	not completely forbid copy attempts that break the cell connectivity, but punishes them
	through a positive term in the Hamiltonian. 
	@experimental
	*/
	class SoftConnectivityConstraint extends SoftConstraint {

		/** The constructor of the ConnectivityConstraint requires a conf object with one parameter.
		@param {object} conf - parameter object for this constraint.
		@param {PerKindBoolean} conf.LAMBDA_CONNECTIVITY - should the cellkind be connected or not?
		*/
		constructor( conf ){
			super(conf);
			
			/** Object tracking the borderpixels of each cell. This is kept up to date after
			every copy attempt.
			@type {CellObject}*/
			this.borderpixelsbycell = {};
			
			this.components = [];
		}
		
		/** The set CPM method attaches the CPM to the constraint. */
		set CPM(C){
			/** CPM on which this constraint acts.
			@type {CPM}*/
			this.C = C;
			
			/** Private property used by {@link updateBorderPixels} to track borders. 
			@private
			@type {Uint16Array} */
			this._neighbours = new Uint16Array(this.C.grid.p2i(this.C.extents));
		}
		
		/** This method checks that all required parameters are present in the object supplied to
		the constructor, and that they are of the right format. It throws an error when this
		is not the case.*/
		confChecker(){
			let checker = new ParameterChecker( this.conf, this.C );
			checker.confCheckParameter( "LAMBDA_CONNECTIVITY", "KindArray", "NonNegative" );
		}
		
		/** Update the borderpixels when pixel i changes from t_old into t_new.
		@param {IndexCoordinate} i - the pixel to change
		@param {CellId} t_old - the cell the pixel belonged to previously
		@param {CellId} t_new - the cell the pixel belongs to now. */
		updateBorderPixels( i, t_old, t_new  ){
			if( t_old == t_new ) return
			if( !(t_new in this.borderpixelsbycell) ){
				this.borderpixelsbycell[t_new] = {};
			}
			const Ni = this.C.grid.neighi( i );
			const wasborder = this._neighbours[i] > 0;
			// current neighbors of pixel i, set counter to zero and loop over nbh.
			this._neighbours[i] = 0;
			for( let ni of Ni  ){
				// type of the neighbor.
				const nt = this.C.grid.pixti(ni);
				
				// If type is not the t_new of pixel i, nbi++ because the neighbor belongs
				// to a different cell. 
				if( nt != t_new ){
					this._neighbours[i] ++; 
				}
				
				// If neighbor type is t_old, the border of t_old may have to be adjusted. 
				// It gets an extra neighbor because the current pixel becomes t_new.
				if( nt == t_old ){
					// If this wasn't a borderpixel of t_old, it now becomes one because
					// it has a neighbor belonging to t_new
					if( this._neighbours[ni] ++ == 0 ){
						if( !(t_old in this.borderpixelsbycell) ){
							this.borderpixelsbycell[t_old] = {};
						}
						this.borderpixelsbycell[t_old][ni] = true;
					}

				}
				// If type is t_new, the neighbor may no longer be a borderpixel
				if( nt == t_new ){
					if( --this._neighbours[ni] == 0 && ( ni in this.borderpixelsbycell[t_new] ) ){
						delete this.borderpixelsbycell[t_new][ni];
					}
				}
			}
			// Case 1: 
			// If current pixel is a borderpixel, add it to those of the current cell.
			if( this._neighbours[i] > 0 ){
				this.borderpixelsbycell[t_new][i]=true;
			}
			
			// Case 2:
			// Current pixel was a borderpixel. Remove from the old cell. 
			if( wasborder ){
				// It was a borderpixel from the old cell, but no longer belongs to that cell.
				if( i in this.borderpixelsbycell[t_old] ){ 
					delete this.borderpixelsbycell[t_old][i];
				}
			}
			
		}
		
		/** Get the connected components of the borderpixels of the current cell.
		@param {CellId} cellid - cell to check the connected components of.
		@return {object} an array with an element for every connected component, which is in
		turn an array of the {@link ArrayCoordinate}s of the pixels belonging to that component.  */
		connectedComponentsOfCellBorder( cellid ){
		
			/* Note that to get number of connected components, we only need to look at cellborderpixels. */
			if( !( cellid in this.borderpixelsbycell ) ){
				return []
			}
			
			//let cbpi = Object.keys( this.borderpixelsbycell[cellid] ), cbpobject = this.borderpixelsbycell[cellid]
			return this.connectedComponentsOf( this.borderpixelsbycell[cellid] )
		}
		
		/** Get the connected components of a set of pixels.
		@param {object} pixelobject - an object with as keys the {@link IndexCoordinate}s of the pixels to check.
		@return {object} an array with an element for every connected component, which is in
		turn an array of the {@link ArrayCoordinate}s of the pixels belonging to that component.  */
		connectedComponentsOf( pixelobject ){
		
			let cbpi = Object.keys( pixelobject );
			
			let visited = {}, k=0, pixels = [], C = this.C;
			let labelComponent = function(seed, k){
				let q = [seed];
				let cellid = C.pixti(q);
				visited[q[0]] = 1;
				pixels[k] = [];
				while( q.length > 0 ){
					let e = q.pop();
					pixels[k].push(C.grid.i2p(e) );
					let ne = C.grid.neighi( e );
					for( let i = 0 ; i < ne.length ; i ++ ){
						if( C.pixti( ne[i] ) == cellid &&
							!(ne[i] in visited) && (ne[i] in pixelobject) ){
							q.push(ne[i]);
							visited[ne[i]]=1;
						}
					}
				}
			};
			for( let i = 0 ; i < cbpi.length ; i ++ ){
				let pi = parseInt( cbpi[i] );
				if( !(pi in visited) ){
					labelComponent( pi, k );
					k++;
				}
			}
			return pixels
		}
			
		/** The postSetpixListener of the ConnectivityConstraint updates the internally
		tracked borderpixels after every copy.
		@param {IndexCoordinate} i - the pixel to change
		@param {CellId} t_old - the cell the pixel belonged to previously
		@param {CellId} t - the cell the pixel belongs to now.	
		*/
		postSetpixListener(  i, t_old, t ){
			this.updateBorderPixels( i, t_old, t );
		}	
		
		
		/** To speed things up: first check if a pixel change disrupts the local connectivity
		in its direct neighborhood. If local connectivity is not disrupted, we don't have to
		check global connectivity at all. This currently only works in 2D, so it returns 
		false for 3D (ensuring that connectivity is checked globally).
		@param {IndexCoordinate} tgt_i - the pixel to change
		@param {CellId} tgt_type - the cell the pixel belonged to before the copy attempt.
		@return {boolean} does the local neighborhood remain connected if this pixel changes?
		*/
		localConnected( tgt_i, tgt_type ){
		
			// Get neighbors of the target pixel
			let nbh = this.C.grid.neighi( tgt_i );
			
			// object storing the neighbors of tgt_type
			let nbhobj = {};
			
			for( let n of nbh ){
			
				// add it and its neighbors to the neighborhood object
				if( this.C.pixti(n) == tgt_type ){
					nbhobj[n] = true;
				}
			}
			
			// Get connected components.
			let conn = this.connectedComponentsOf( nbhobj );
			this.components = conn;
			//console.log(conn.length)
			
			let connected = ( conn.length == 1 );
			//console.log(connected)
			return connected
			
		}
		
		/** Compute the 'connectivity' of a cell; a number between 0 and 1. If the cell
		is completely connected, this returns 1. A cell split into many parts gets a 
		connectivity approaching zero. It also matters how the cell is split: splitting
		the cell in two near-equal parts results in a lower connectivity than separating
		one pixel from the rest of the cell.
		@param {Array} components - an array of arrays (one array per connected component, 
		in which each entry is the {@link ArrayCoordinate} of a pixel belonging to that component).
		@param {CellId} cellid - the cell these components belong to.
		@return {number} connectivity of this cell.*/
		connectivity( components, cellid ){
			if( components.length <= 1 ){
				return 1
			} else {
				
				let Vtot = Object.keys( this.borderpixelsbycell[cellid] ).length;
				let Ci = 0;
				for( let c of components ){
					let Vc = c.length;
					Ci += (Vc/Vtot)*(Vc/Vtot);
				}
				//console.log( Ci )
				return Ci
				
			}
		}
		
		/** This method checks the difference in connectivity when pixel tgt_i is changed from
		tgt_type to src_type. 
		@param {IndexCoordinate} tgt_i - the pixel to change
		@param {CellId} src_type - the new cell for this pixel.
		@param {CellId} tgt_type - the cell the pixel belonged to previously. 	
		@return {number} conndiff - the difference: connectivity_after - connectivity_before.
		*/
		checkConnected( tgt_i, src_type, tgt_type ){
		
			//return this.localConnected( tgt_i, tgt_type )
			
		
			
			if( this.localConnected( tgt_i, tgt_type ) ){
				return 0
			} 
			
			/*else {
				let conn_new = this.connectivity( this.components, tgt_type )
				
				// current components
				let nbh = this.C.grid.neighi( tgt_i )
				let nbhobj = {}
				nbhobj[tgt_i] = true
			
				for( let n of nbh ){
					if( this.C.pixti(n) == tgt_type ){
						nbhobj[n] = true
					}
				}
				
				this.components = this.connectedComponentsOf( nbhobj )
				let conn_old = this.connectivity( this.components, tgt_type )
				
				let conndiff = conn_old - conn_new
				return conndiff
				
				
				
			}*/
			
			
		
			let comp1 = this.connectedComponentsOfCellBorder( tgt_type );
			let conn1 = Math.pow( (1-this.connectivity( comp1, tgt_type )),2 );
		
			// Update the borderpixels as if the change occurs
			this.updateBorderPixels( tgt_i, tgt_type, src_type );
			let comp = this.connectedComponentsOfCellBorder( tgt_type );
			let conn2 = Math.pow((1-this.connectivity( comp, tgt_type )),2);
			
			
			let conndiff = conn2 - conn1;
			/*if( conn2 > conn1 ){
				conndiff = -conndiff
			} */
			
			// Change borderpixels back because the copy attempt hasn't actually gone through yet.
			this.updateBorderPixels( tgt_i, src_type, tgt_type );
			
			return conndiff
			
		}

		/** Method for hard constraints to compute whether the copy attempt fulfills the rule.
		 @param {IndexCoordinate} src_i - coordinate of the source pixel that tries to copy.
		 @param {IndexCoordinate} tgt_i - coordinate of the target pixel the source is trying
		 to copy into.
		 @param {CellId} src_type - cellid of the source pixel.
		 @param {CellId} tgt_type - cellid of the target pixel. 
		 @return {boolean} whether the copy attempt satisfies the constraint.*/ 
		deltaH( src_i, tgt_i, src_type, tgt_type ){
			// connectedness of src cell cannot change if it was connected in the first place.
			
			let lambda = this.conf["LAMBDA_CONNECTIVITY"][this.C.cellKind(tgt_type)];
			
			// connectedness of tgt cell
			if( tgt_type != 0 && lambda > 0 ){
				return lambda*this.checkConnected( tgt_i, src_type, tgt_type )
			}
			
			return 0
		}
	}

	/** Version of the {@link ConnectivityConstraint} which only checks local
	 * connectivity.
	 * @experimental
	 * */
	class LocalConnectivityConstraint extends HardConstraint {

		/** The constructor of the LocalConnectivityConstraint requires a conf
		 * object with one parameter.
		 * @param {object} conf - parameter object for this constraint.
		 * @param {PerKindBoolean} conf.CONNECTED - should the cellkind be connected
		 * or not?
		 * */
		constructor( conf ){
			super(conf);
		}
		

		/** This method checks that all required parameters are present in the
		 * object supplied to the constructor, and that they are of the right
		 * format. It throws an error when this is not the case.
		 * */
		confChecker(){
			let checker = new ParameterChecker( this.conf, this.C );
			checker.confCheckParameter( "CONNECTED", "KindArray", "Boolean" );
		}
		
		/** Get the connected components of a set of pixels.
		 * @param {object} pixelObject - an object with as keys the
		 * {@link IndexCoordinate}s of the pixels to check.
		 * @return {object} an array with an element for every connected component,
		 * which is in turn an array of the {@link ArrayCoordinate}s of the pixels
		 * belonging to that component.
		 * */
		connectedComponentsOf( pixelObject ){
		
			let cbpi = Object.keys( pixelObject );
			
			let visited = {}, k=0, pixels = [], C = this.C;
			let labelComponent = function(seed, k){
				let q = [seed];
				let cellId = C.grid.pixti(q);
				visited[q[0]] = 1;
				pixels[k] = [];
				while( q.length > 0 ){
					let e = q.pop();
					pixels[k].push(C.grid.i2p(e) );
					let ne = C.grid.neighi( e );
					for( let i = 0 ; i < ne.length ; i ++ ){
						if( C.grid.pixti( ne[i] ) === cellId &&
							!(ne[i] in visited) && (ne[i] in pixelObject) ){
							q.push(ne[i]);
							visited[ne[i]]=1;
						}
					}
				}
			};
			for( let i = 0 ; i < cbpi.length ; i ++ ){
				let pi = parseInt( cbpi[i] );
				if( !(pi in visited) ){
					labelComponent( pi, k );
					k++;
				}
			}
			return pixels
		}

		
		/** This method checks if the connectivity still holds after pixel tgt_i is
		 * changed from tgt_type to src_type.
		 * @param {IndexCoordinate} tgt_i - the pixel to change.
		 * @param {CellId} src_type - the new cell for this pixel.
		 * @param {CellId} tgt_type - the cell the pixel belonged to previously.
		 * */
		checkConnected( tgt_i, src_type, tgt_type ){
		
			// Get neighbors of the target pixel
			let nbh = this.C.grid.neighi( tgt_i );
			
			// object storing the neighbors of tgt_type
			let nbhObj = {};
			
			for( let n of nbh ){
			
				// add it and its neighbors to the neighborhood object
				if( this.C.grid.pixti(n) === tgt_type ){
					nbhObj[n] = true;
				}
				
				/*for( let i of this.C.grid.neighi(n) ){
				
					if( ( this.C.grid.pixti(i) == tgt_type ) && !( i == tgt_i ) ){
						nbhObj[i] = true
					}
				}*/
			}
			
			// Get connected components.
			let conn = this.connectedComponentsOf( nbhObj );
			
			return ( conn.length === 1 )
			
		}

		/** Method for hard constraints to compute whether the copy attempt fulfills
		 * the rule.
		 * @param {IndexCoordinate} src_i - coordinate of the source pixel that
		 * tries to copy.
		 * @param {IndexCoordinate} tgt_i - coordinate of the target pixel the
		 * source is trying to copy into.
		 * @param {CellId} src_type - cellId of the source pixel.
		 * @param {CellId} tgt_type - cellId of the target pixel.
		 * @return {boolean} whether the copy attempt satisfies the constraint.
		 * */
		fulfilled( src_i, tgt_i, src_type, tgt_type ){
			// connectedness of src cell cannot change if it was connected in the first place.
			
			// connectedness of tgt cell
			if( tgt_type !== 0 && this.conf["CONNECTED"][this.C.cellKind(tgt_type)] ){
				return this.checkConnected( tgt_i, src_type, tgt_type )
			}
			
			return true
		}
	}

	/** Soft version of the {@link ConnectivityConstraint} which only checks local connectivity.
	@experimental
	*/
	class SoftLocalConnectivityConstraint extends SoftConstraint {

		/** The constructor of the SoftLocalConnectivityConstraint requires a conf object with one or two parameters.
		@param {object} conf - parameter object for this constraint.
		@param {PerKindBoolean} conf.LAMBDA_CONNECTIVITY - strength of the penalty for breaking connectivity.
		#param {string} conf.NBH_TYPE - should a Neumann (default) or Moore neighborhood be used to determine
		whether the cell locally stays connected? The default is Neumann since the Moore neighborhood tends to
		give artefacts. Also, LAMBDA should be much higher if the Moore neighborhood is used. 
		*/
		constructor( conf ){
			super(conf);
			
			/** Should a Neumann or Moore neighborhood be used for determining connectivity?
			See {@link SoftLocalConnectivityConstraint#constructor} for details.
			@type {string}*/
			this.nbhtype = "Neumann";
		}
		
		/** The set CPM method attaches the CPM to the constraint. It checks whether the
		CPM is 2D or 3D, because this constraint is currently only tested in 2D. */
		set CPM(C){
			/** CPM on which this constraint acts.
			@type {CPM}*/
			this.C = C;
			
			if( this.C.ndim != 2 ){
				throw("You are trying to add a SoftLocalConnectivityConstraint to a 3D CPM, but this constraint is currently only supported in 2D!")
			}
			
			this.confChecker();
		}
		
		/** This method checks that all required parameters are present in the object supplied to
		the constructor, and that they are of the right format. It throws an error when this
		is not the case.*/
		confChecker(){
			let checker = new ParameterChecker( this.conf, this.C );
			checker.confCheckParameter( "LAMBDA_CONNECTIVITY", "KindArray", "NonNegative" );

			//
			if( "NBH_TYPE" in this.conf ){
				let v = this.conf["NBH_TYPE"];
				let values = [ "Neumann", "Moore" ];
				let found = false;
				for( let val of values ){
					if( val == v ){
						found = true;
						this.nbhtype = val;
					}
				}
				if( !found ){
					throw( "In the SoftLocalConnectivityConstraint, NBH_TYPE must be either 'Neumann' or 'Moore'")
				}
			}

		}
		
		/** Get the connected components of a set of pixels.
		@param {object} pixelobject - an object with as keys the {@link IndexCoordinate}s of the pixels to check.
		@return {object} an array with an element for every connected component, which is in
		turn an array of the {@link ArrayCoordinate}s of the pixels belonging to that component.  */
		connectedComponentsOf( pixelobject ){
		
			let cbpi = Object.keys( pixelobject );
			
			let visited = {}, k=0, pixels = [], C = this.C, nbhtype = this.nbhtype;
			let labelComponent = function(seed, k){
				let q = [seed];
				let cellid = C.pixti(q);
				visited[q[0]] = 1;
				pixels[k] = [];
				while( q.length > 0 ){
					let e = q.pop();
					pixels[k].push(C.grid.i2p(e) );
					
					if( nbhtype == "Neumann" ){
						for( let i of C.grid.neighNeumanni( e ) ){
							if( C.pixti( i ) == cellid &&
								!(i in visited) && (i in pixelobject) ){
								q.push(i);
								visited[i]=1;
							}
						}
					} else {
						let ne = C.grid.neighi(e);
						for( let j = 0; j < ne.length; j++ ){
						
							let i = ne[j];
							if( C.pixti( i ) == cellid &&
								!(i in visited) && (i in pixelobject) ){
								q.push(i);
								visited[i]=1;
							}
						}
					
					}
					
					
					
					//let ne = C.grid.neighi( e )
					
				}
			};
			for( let i = 0 ; i < cbpi.length ; i ++ ){
				let pi = parseInt( cbpi[i] );
				if( !(pi in visited) ){
					labelComponent( pi, k );
					k++;
				}
			}
			return pixels
		}

		/** This method checks if the connectivity still holds after pixel tgt_i is changed from
		tgt_type to src_type. 
		@param {IndexCoordinate} tgt_i - the pixel to change
		@param {CellId} src_type - the new cell for this pixel.
		@param {CellId} tgt_type - the cell the pixel belonged to previously. 	
		@return {number} 1 if connectivity is broken, 0 if the connectivity remains. 
		*/
		checkConnected( tgt_i, src_type, tgt_type ){
		
			// Get neighbors of the target pixel
			let nbh = this.C.grid.neighi( tgt_i );
			
			// object storing the neighbors of tgt_type
			let nbhobj = {};
			
			for( let n of nbh ){
			
				// add it and its neighbors to the neighborhood object
				if( this.C.pixti(n) == tgt_type ){
					nbhobj[n] = true;
				}
			}
			
			// Get connected components.
			let conn = this.connectedComponentsOf( nbhobj );
			//console.log(conn.length)
			
			let disconnected = 0;
			if( conn.length > 1 ){
				disconnected = 1;
			}
			return disconnected
			
		}
		
		/** Method for hard constraints to compute whether the copy attempt fulfills the rule.
		 @param {IndexCoordinate} src_i - coordinate of the source pixel that tries to copy.
		 @param {IndexCoordinate} tgt_i - coordinate of the target pixel the source is trying
		 to copy into.
		 @param {CellId} src_type - cellid of the source pixel.
		 @param {CellId} tgt_type - cellid of the target pixel. 
		 @return {boolean} whether the copy attempt satisfies the constraint.*/ 
		deltaH( src_i, tgt_i, src_type, tgt_type ){
			// connectedness of src cell cannot change if it was connected in the first place.
			
			let lambda = this.conf["LAMBDA_CONNECTIVITY"][this.C.cellKind(tgt_type)];
			
			// connectedness of tgt cell. Only check when the lambda is non-zero.
			if( tgt_type != 0 && lambda > 0 ){
				return lambda*this.checkConnected( tgt_i, src_type, tgt_type )
			}
			
			return 0
		}

		

	}

	/** 
	 * This constraint forbids that cells exceed or fall below a certain size range. 
	 * @example
	 * // Build a CPM and add the constraint
	 * let CPM = require( "path/to/build" )
	 * let C = new CPM.CPM( [200,200], {T : 4})
	 * C.add( new CPM.HardVolumeRangeConstraint( {
	 * 	LAMBDA_VRANGE_MIN : [0,1],
	 * 	LAMBDA_VRANGE_MAX : [0,2]
	 * } ) )
	 */
	class HardVolumeRangeConstraint extends HardConstraint {

		/** The constructor of the HardVolumeRangeConstraint requires a conf object with two parameters.
		@param {object} conf - parameter object for this constraint.
		@param {PerKindNonNegative} conf.LAMBDA_VRANGE_MIN - minimum volume of each cellkind.
		@param {PerKindNonNegative} conf.LAMBDA_VRANGE_MAX - maximum volume of each cellkind.
		*/
		constructor( conf ){
			super(conf);
		}
		
		/** This method checks that all required parameters are present in the object supplied to
		the constructor, and that they are of the right format. It throws an error when this
		is not the case.*/
		confChecker(){
			let checker = new ParameterChecker( this.conf, this.C );
			checker.confCheckParameter( "LAMBDA_VRANGE_MAX", "KindArray", "NonNegative" );
			checker.confCheckParameter( "LAMBDA_VRANGE_MIN", "KindArray", "NonNegative" );
		}

		/** Method for hard constraints to compute whether the copy attempt fulfills the rule.
		 @param {IndexCoordinate} src_i - coordinate of the source pixel that tries to copy.
		 @param {IndexCoordinate} tgt_i - coordinate of the target pixel the source is trying
		 to copy into.
		 @param {CellId} src_type - cellid of the source pixel.
		 @param {CellId} tgt_type - cellid of the target pixel. 
		 @return {boolean} whether the copy attempt satisfies the constraint.*/ 
		fulfilled( src_i, tgt_i, src_type, tgt_type ){
			// volume gain of src cell
			if( src_type != 0 && this.C.getVolume(src_type) + 1 > 
				this.conf["LAMBDA_VRANGE_MAX"][this.C.cellKind(src_type)] ){
				return false
			}
			// volume loss of tgt cell
			if( tgt_type != 0 && this.C.getVolume(tgt_type) - 1 < 
				this.conf["LAMBDA_VRANGE_MIN"][this.C.cellKind(tgt_type)] ){
				return false
			}
			return true
		}
	}

	/** 
	 * This constraint allows a set of "barrier" background pixels, into 
	 * which copy attempts are forbidden.
	 * @example
	 * // Build a CPM and add the constraint
	 * let CPM = require( "path/to/build" )
	 * let C = new CPM.CPM( [200,200], {
	 * 	T : 20,
	 * 	J : [[0,20],[20,10]],
	 * 	V : [0,500],
	 * 	LAMBDA_V : [0,5],
	 * })
	 * 
	 * // Build a barrier and add the border constraint
	 * let border = []
	 * let channelwidth = 10
	 * for( let x = 0; x < C.extents[0]; x++ ){
	 * 	let ymin = Math.floor( C.extents[1]/2 )
	 *  let ymax = ymin + channelwidth
	 *  border.push( [x,ymin] )
	 *  border.push( [x,ymax] )
	 * }
	 * 
	 * C.add( new CPM.BorderConstraint( {
	 * 	BARRIER_VOXELS : border
	 * } ) )
	 * 
	 * // Seed a cell
	 * let gm = new CPM.GridManipulator( C )
	 * gm.seedCell(1)
	 */
	class BorderConstraint extends HardConstraint {

		/** Creates an instance of the ActivityMultiBackground constraint 
		* @param {object} conf - Configuration object with the parameters.
		* ACT_MEAN is a single string determining whether the activity mean should be computed
		* using a "geometric" or "arithmetic" mean. 
		*/
		/** The constructor of the ActivityConstraint requires a conf object with parameters.
		@param {object} conf - parameter object for this constraint
		@param {string} [conf.ACT_MEAN="geometric"] - should local mean activity be measured with an
		"arithmetic" or a "geometric" mean?
		@param {PerKindArray} conf.LAMBDA_ACT_MBG - strength of the activityconstraint per cellkind and per background.
		@param {PerKindNonNegative} conf.MAX_ACT - how long do pixels remember their activity? Given per cellkind.
		@param {Array} conf.BACKGROUND_VOXELS - an array where each element represents a different background type.
		This is again an array of {@ArrayCoordinate}s of the pixels belonging to that backgroundtype. These pixels
		will have the LAMBDA_ACT_MBG value of that backgroundtype, instead of the standard value.
		*/
		constructor( conf ){
			super( conf );
		
			/** Store which pixels are barrier pixels. Each entry has key the {@IndexCoordinate} of
			the pixel, and value equal to true.
			@type {object}*/
			this.barriervoxels = {};
			
			/** Track if this.barriervoxels has been set.
			@type {boolean}*/
			this.setup = false;
		}
		
		
		
		/** This method checks that all required parameters are present in the object supplied to
		the constructor, and that they are of the right format. It throws an error when this
		is not the case.*/
		confChecker(){
			let checker = new ParameterChecker( this.conf, this.C );

			checker.confCheckPresenceOf( "BARRIER_VOXELS" );
			let barriervox = this.conf["BARRIER_VOXELS"];
			// Barrier voxels must be an array of arrays
			if( !(barriervox instanceof Array) ){
				throw( "Parameter BARRIER_VOXELS should be an array!" )
			} 
			// Elements of the initial array must be arrays.
			for( let e of barriervox ){
				
				let isCoordinate = true;
				if( !(e instanceof Array)){
					isCoordinate = false;
				} else if( e.length != this.C.extents.length ){
					isCoordinate = false;
				}
				if( !isCoordinate ){
					throw( "Parameter BARRIER_VOXELS: elements should be ArrayCoordinates; arrays of length " + this.C.extents.length + "!" )
					
				}
				
			}
		}
		
		/** Get the background voxels from input argument or the conf object and store them in a correct format
		in this.barriervoxels. This only has to be done once, but can be called from outside to
		change the background voxels during a simulation (eg in a HTML page).
		@param {ArrayCoordinate[]} voxels - the pixels that should act as barrier.
		 */	
		setBarrierVoxels( voxels ){
		
			voxels = voxels || this.conf["BARRIER_VOXELS"];
		
			// reset if any exist already
			this.barriervoxels = {};
			for( let v of voxels ){
				this.barriervoxels[ this.C.grid.p2i(v) ] = true;
			}
			this.setup = true;

		}
		
		/** Method for hard constraints to compute whether the copy attempt fulfills the rule.
		 @param {IndexCoordinate} src_i - coordinate of the source pixel that tries to copy.
		 @param {IndexCoordinate} tgt_i - coordinate of the target pixel the source is trying
		 to copy into.
		 @param {CellId} src_type - cellid of the source pixel.
		 @param {CellId} tgt_type - cellid of the target pixel. 
		 @return {boolean} whether the copy attempt satisfies the constraint.*/ 
		// eslint-disable-next-line no-unused-vars
		fulfilled( src_i, tgt_i, src_type, tgt_type ){
		
			if( !this.setup ){
				this.setBarrierVoxels();
			}
		
			// If the target pixel is a barrier pixel, forbid the copy attempt.
			if( tgt_i in this.barriervoxels ){
				return false
			}
			
			// Otherwise accept it.
			return true
		}


	}

	/** 
	This class provides some boilerplate code for creating simulations easily.
	It comes with defaults for seeding cells, drawing, logging of statistics, saving output
	images, and running the simulation. Each of these default methods can be overwritten
	by the user while keeping the other default methods intact. See the {@link constructor}
	for details on how to configure a simulation.
	@see ../examples
	*/
	class Simulation {
		/** The constructor of class Simulation takes two arguments.
			@param {object} config - overall configuration settings. This is an object
			with multiple entries, see below.
			@param {GridSize} config.field_size - size of the CPM to build.
			@param {object} config.conf - configuration settings for the CPM;
			see its {@link CPM#constructor} for details.
			@param {object} simsettings - configuration settings for the simulation 
			itself and for controlling the outputs. See the parameters below for details.
			@param {number[]} simsettings.NRCELLS - array with number of cells to seed for
				every non-background {@link CellKind}.
			@param {number} simsettings.BURNIN - number of MCS to run before the actual
				simulation starts (let cells get their target volume before starting).
			@param {number} simsettings.RUNTIME - number of MCS the simulation should run.
				Only necessary if you plan to use the {@link run} method.
			@param {number} [ simsettings.IMGFRAMERATE = 1 ]- draw the grid every [x] MCS.
			@param {number} [ simsettings.LOGRATE = 1 ] - log stats every [x] MCS.
			@param {object} [ simsettings.LOGSTATS = {browser:false,node:true} ] - 
				whether stats should be logged in browser and node.
			@param {boolean} [ simsettings.SAVEIMG = false ] - should images be saved? (node only).
			@param {string} [ simsettings.SAVEPATH ] - where should images be saved? You only have
				to give this argument when SAVEIMG = true. 
			@param {string} [ simsettings.EXPNAME = "myexp" ] - string used to construct the
				filename of any saved image. 
			@param {HexColor} [ simsettings.CANVASCOLOR = "FFFFFF" ] - color to draw the background in; defaults to white.
			@param {HexColor[]} [ simsettings.CELLCOLOR ] - color to draw each non-background 
				{@link CellKind} in. If left unspecified, the {@link Canvas} will use black.
			@param {boolean[]} [simsettings.ACTCOLOR ] - should activities of the {@link ActivityConstraint}
				be drawn for each {@link CellKind}? If left unspecified, these are not drawn.
			@param {boolean[]} [simsettings.SHOWBORDERS = false] - should borders of each {@link CellKind}
				be drawn? Defaults to false.
			@param {HexColor[]} [simsettings.BORDERCOL = "000000"] - color to draw cellborders of
				each {@link CellKind} in. Defaults to black. 
			*/
		constructor( config, custommethods ){
		
			// ========= configuration and custom methods
			
			custommethods = custommethods || {};
		
			// overwrite default method if methods are supplied in custommethods
			// these can be initializeGrid(), drawCanvas(), logStats(),
			// postMCSListener().
			for( let m of Object.keys( custommethods ) ){
			
				/** Any function suplied in the custommethods argument to
				the {@link constructor} is bound to the object. */
				this[m] = custommethods[m];
			}
			
			/** Configuration of the simulation environment 
			@type {object}*/
			this.conf = config.simsettings;
			
			// ========= controlling outputs
			
			/** Draw the canvas every [rate] MCS.
			@type {number}*/
			this.imgrate = this.conf["IMGFRAMERATE"] || 1;
			
			/** Log stats every [rate] MCS.
			@type {number}*/
			this.lograte = this.conf["LOGRATE"] || 1;
			
			/** See if code is run in browser or via node, which will be used
				below to determine what the output should be.
				@type {string}*/
			this.mode = "node";
			if( typeof window !== "undefined" && typeof window.document !== "undefined" ){
				
				this.mode = "browser";
			} 
			
			/** Log stats or not.
			@type {boolean}*/
			this.logstats = false;
			
			/** Log stats or not, specified for both browser and node mode.
			@type {object} */
			this.logstats2 = this.conf["STATSOUT"] || { browser: false, node: true };
			
			
			this.logstats = this.logstats2[this.mode];
			
			/** Saving images or not.
			@type {boolean}*/
			this.saveimg = this.conf["SAVEIMG"] || false;
			/** Where to save images.
			@type {string}*/
			this.savepath = this.conf["SAVEPATH"] || "undefined";
			
			if( this.saveimg && this.savepath === "undefined" ){
				throw( "You need to specify the SAVEPATH option in the configuration object of your simulation!")
			}
			
			// ========= tracking simulation progress
			
			/** Track the time of the simulation. 
			@type {number}*/
			this.time = 0;
			
			/** Should the simulation be running? Change this to pause;
			see the {@link toggleRunning} method.
			@private
			@type {boolean}*/
			this.running = true;
			
			// ========= Attached objects
			
			/** Make CPM object based on configuration settings and attach it.
			@type {CPM} */
			this.C = new CPM( config.field_size, config.conf );
					
			/** See if objects of class {@link Canvas} and {@link GridManipulator} already 
			exist. These are added automatically when required. This will set
			their values in helpClasses to 'true', so they don't have to be added again.
			@type {object}*/
			this.helpClasses = { gm: false, canvas: false };
			
			
			// ========= Begin.
			// Initialize the grid and run the burnin.
			this.initializeGrid();
			this.runBurnin();
			
		}

		/** Adds a {@link GridManipulator} object when required. */
		addGridManipulator(){
			/** Attached {@link GridManipulator} object.
			@type {GridManipulator}*/
			this.gm = new GridManipulator( this.C );
			this.helpClasses[ "gm" ] = true;
		}
		
		/** Adds a {@link Canvas} object when required. */
		addCanvas(){
			//let zoom = this.conf.zoom || 2
			/** Attached {@link Canvas} object.
			@type {Canvas}*/
			this.Cim = new Canvas( this.C, this.conf );
			this.helpClasses[ "canvas" ] = true;
		}
		
		/** Method to initialize the Grid should be implemented in each simulation. 
		The default method checks in the simsettings.NRCELLS array how many cells to
		seed for each {@CellKind}, and does this (at random positions). 
		
		Often you'll want to do other things here. In that case you can use the 
		custommethods argument of the {@link constructor} to overwrite this with your
		own initializeGrid method.
		*/
		initializeGrid(){
		
			// add the initializer if not already there
			if( !this.helpClasses["gm"] ){ this.addGridManipulator(); }
		
			let nrcells = this.conf["NRCELLS"], cellkind, i;
			
			// Seed the right number of cells for each cellkind
			for( cellkind = 0; cellkind < nrcells.length; cellkind ++ ){
				
				for( i = 0; i < nrcells[cellkind]; i++ ){
					// first cell always at the midpoint. Any other cells
					// randomly.				
					
					this.gm.seedCell( cellkind+1 );
					
				}
			}


		}
		
		/** Run the brunin period as defined by simsettings.BURNIN : run this number
		of MCS before the {@link time} of this simulation object starts ticking, and 
		before we start drawing etc. 
		*/	
		runBurnin(){
			// Simulate the burnin phase
			let burnin = this.conf["BURNIN"] || 0;
			for( let i = 0; i < burnin; i++ ){
				this.C.monteCarloStep();
			}
		}

		
		/** Method to draw the canvas.
		The default method draws the canvas, cells, cellborders, and activityvalues
		as specified in the simsettings object (see the {@link constructor} for details).
		
		This will be enough for most scenarios, but if you want to draw more complicated stuff,
		you can use the custommethods argument of the {@link constructor} to overwrite 
		this with your own drawCanvas method.
		*/
		drawCanvas(){
		
			// Add the canvas if required
			if( !this.helpClasses["canvas"] ){ this.addCanvas(); }
		
			// Clear canvas and draw stroma border
			this.Cim.clear( this.conf["CANVASCOLOR"] || "FFFFFF" );


			// Call the drawBelow method for if it is defined. 
			this.drawBelow();

			// Draw each cellkind appropriately
			let cellcolor=( this.conf["CELLCOLOR"] || [] ), actcolor=this.conf["ACTCOLOR"], 
				nrcells=this.conf["NRCELLS"], cellkind, cellborders = this.conf["SHOWBORDERS"];
			for( cellkind = 0; cellkind < nrcells.length; cellkind ++ ){
			
				// draw the cells of each kind in the right color
				if( cellcolor[ cellkind ] !== -1 ){
					this.Cim.drawCells( cellkind+1, cellcolor[cellkind] );
				}
				
				// Draw borders if required
				if(  this.conf.hasOwnProperty("SHOWBORDERS") && cellborders[ cellkind  ] ){
					let bordercol = "000000";
					if( this.conf.hasOwnProperty("BORDERCOL") ){
						bordercol = this.conf["BORDERCOL"][cellkind] || "000000";
					}
					this.Cim.drawCellBorders( cellkind+1, bordercol );
				}
				
				// if there is an activity constraint, draw activity values depending on color.
				if( this.C.conf["LAMBDA_ACT"] !== undefined && this.C.conf["LAMBDA_ACT"][ cellkind + 1 ] > 0 ){ //this.constraints.hasOwnProperty( "ActivityConstraint" ) ){
					let colorAct;
					if( typeof actcolor !== "undefined" ){
						colorAct = actcolor[ cellkind ] || false;
					} else {
						colorAct = false;
					}
					if( ( colorAct ) ){
						this.Cim.drawActivityValues( cellkind + 1 );//, this.constraints["ActivityConstraint"] )
					}			
				}

			}
			
			// Call the drawOnTop() method for if it is defined. 
			this.drawOnTop();
			
		}
		
		/** Methods drawBelow and {@link drawOnTop} allow you to draw extra stuff below and
		on top of the output from {@link drawCanvas}, respectively. You can use them if you
		wish to visualize additional properties but don't want to remove the standard visualization.
		They are called at the beginning and end of {@link drawCanvas}, so they do not work
		if you overwrite this method. 
		*/
		drawBelow(){
		
		}
		
		/** Methods drawBelow and {@link drawOnTop} allow you to draw extra stuff below and
		on top of the output from {@link drawCanvas}, respectively. You can use them if you
		wish to visualize additional properties but don't want to remove the standard visualization.
		They are called at the beginning and end of {@link drawCanvas}, so they do not work
		if you overwrite this method. 
		*/
		drawOnTop(){
		
		}
		
		
		/** Method to log statistics.
		The default method logs time, {@link CellId}, {@link CellKind}, and the 
		{@ArrayCoordinate} of the cell's centroid to the console.
		
		If you want to compute other stats (see subclasses of {@link Stat} for options)
		you can use the custommethods argument of the {@link constructor} to overwrite 
		this with your own drawCanvas method.
		*/
		logStats(){
			
			// compute centroids for all cells
			let allcentroids; 
			let torus = false;
			for( let d = 0; d < this.C.grid.ndim; d++ ){
				if( this.C.grid.torus[d] ){
					torus = true;
				}
			}
			if( torus ){
				allcentroids = this.C.getStat( CentroidsWithTorusCorrection );
			} else {
				allcentroids = this.C.getStat( Centroids );
			} 
			
			for( let cid of this.C.cellIDs() ){
			
				let thecentroid = allcentroids[cid];
				
				// eslint-disable-next-line no-console
				console.log( this.time + "\t" + cid + "\t" + 
					this.C.cellKind(cid) + "\t" + thecentroid.join("\t") );
				
			}

		}
		
		/** Listener for something that needs to be done after every monte carlo step.
		This method is empty but can be overwritten via the custommethods 
		argument of the {@link constructor}.*/
		postMCSListener(){
		
		}
		
		/** This automatically creates all outputs (images and logged stats) at the correct
		rates. See the {@link constructor} documentation for options on how to control these
		outputs. */
		createOutputs(){
			// Draw the canvas every IMGFRAMERATE steps
			if( this.imgrate > 0 && this.time % this.imgrate == 0 ){
				
				if( this.mode == "browser" ){
					this.drawCanvas();
				}
				
				// Save the image if required and if we're in node (not possible in browser)
				if( this.mode == "node" && this.saveimg ){
					this.drawCanvas();
					let outpath = this.conf["SAVEPATH"], expname = this.conf["EXPNAME"] || "mysim";
					this.Cim.writePNG( outpath +"/" + expname + "-t"+this.time+".png" );
				}
			}
			
			// Log stats every LOGRATE steps
			if( this.logstats && this.time % this.lograte == 0 ){
				this.logStats();
			}
		}
		
		/** Run a montecarlostep, produce outputs if required, run any {@link postMCSListener},
		and update the time. */
		step(){
			if( this.running ){
				this.C.monteCarloStep();
				this.postMCSListener();
				this.createOutputs();
				this.time++;
			}
		}
		
		/** Use this to pause or restart the simulation from an HTML page. */
		toggleRunning(){
			this.running = !this.running;
		}
		
		/** Run the entire simulation. This function is meant for nodejs, as you'll
		want to perform individual {@link step}s in a requestAnimationFrame for an 
		animation in a HTML page. */
		run(){
			while( this.time < this.conf["RUNTIME"] ){
			
				this.step();
				
			}
		}
		
	}

	// NOTE : This file is now auto-generated by app/automatic-index.bash when you compile the build using 'make'.

	exports.Canvas = Canvas;
	exports.CA = CA;
	exports.CPM = CPM;
	exports.GridBasedModel = GridBasedModel;
	exports.Stat = Stat;
	exports.PixelsByCell = PixelsByCell;
	exports.BorderPixelsByCell = BorderPixelsByCell;
	exports.CentroidsWithTorusCorrection = CentroidsWithTorusCorrection;
	exports.Centroids = Centroids;
	exports.CellNeighborList = CellNeighborList;
	exports.ConnectedComponentsByCell = ConnectedComponentsByCell;
	exports.Connectedness = Connectedness;
	exports.Grid = Grid;
	exports.Grid2D = Grid2D;
	exports.Grid3D = Grid3D;
	exports.GridManipulator = GridManipulator;
	exports.CoarseGrid = CoarseGrid;
	exports.SoftConstraint = SoftConstraint;
	exports.Adhesion = Adhesion;
	exports.VolumeConstraint = VolumeConstraint;
	exports.PerimeterConstraint = PerimeterConstraint;
	exports.ActivityConstraint = ActivityConstraint;
	exports.ActivityMultiBackground = ActivityMultiBackground;
	exports.PersistenceConstraint = PersistenceConstraint;
	exports.PreferredDirectionConstraint = PreferredDirectionConstraint;
	exports.ChemotaxisConstraint = ChemotaxisConstraint;
	exports.AttractionPointConstraint = AttractionPointConstraint;
	exports.ConnectivityConstraint = ConnectivityConstraint;
	exports.SoftConnectivityConstraint = SoftConnectivityConstraint;
	exports.LocalConnectivityConstraint = LocalConnectivityConstraint;
	exports.SoftLocalConnectivityConstraint = SoftLocalConnectivityConstraint;
	exports.HardConstraint = HardConstraint;
	exports.HardVolumeRangeConstraint = HardVolumeRangeConstraint;
	exports.BarrierConstraint = BarrierConstraint;
	exports.BorderConstraint = BorderConstraint;
	exports.Simulation = Simulation;

	return exports;

}({}));
```
