## Supplementary material for "Artistoo: build, share, and explore simulations of cells and tissues in the web browser": run-ActModel.js

```
let fs = require('fs')
let dir = "output"

if (!fs.existsSync(dir)){
    fs.mkdirSync(dir);
}

let CPM = require("../build/artistoo-cjs.js")


/*	----------------------------------
	CONFIGURATION SETTINGS
	----------------------------------
*/
let config = {

	// Grid settings
	ndim : 2,
	field_size : [200,200],
	
	// CPM parameters and configuration
	conf : {
		// Basic CPM parameters
		torus : [true,true],						// Should the grid have linked borders?
		T : 20,								// CPM temperature
		
		// Constraint parameters. 
		// Mostly these have the format of an array in which each element specifies the
		// parameter value for one of the cellkinds on the grid.
		// First value is always cellkind 0 (the background) and is often not used.
				
		// Adhesion parameters:
		J: [[0,140], [140,0]],
		
		// VolumeConstraint parameters
		LAMBDA_V: [0,50],					// VolumeConstraint importance per cellkind
		V: [0,2500],							// Target volume of each cellkind
		
		// PerimeterConstraint parameters
		LAMBDA_P: [0,5],						// PerimeterConstraint importance per cellkind
		P : [0,650],						// Target perimeter of each cellkind
		
		// ActivityConstraint parameters
		LAMBDA_ACT : [0,80],				// ActivityConstraint importance per cellkind
		MAX_ACT : [0,200],					// Activity memory duration per cellkind
		ACT_MEAN : "geometric"				// Is neighborhood activity computed as a
											// "geometric" or "arithmetic" mean?

	},
	
	// Simulation setup and configuration
	simsettings : {
	
		// Cells on the grid
		NRCELLS : [1],						// Number of cells to seed for all
											// non-background cellkinds.
		// Runtime etc
		BURNIN : 0,
		RUNTIME : 15001,
		RUNTIME_BROWSER : "Inf",
		
		// Visualization
		CANVASCOLOR : "eaecef",
		CELLCOLOR : ["000000"],
		ACTCOLOR : [true],					// Should pixel activity values be displayed?
		SHOWBORDERS : [false],				// Should cellborders be displayed?
		zoom : 1,							// zoom in on canvas with this factor.
		
		// Output images
		SAVEIMG : true,					// Should a png image of the grid be saved
											// during the simulation?
		IMGFRAMERATE : 250,					// If so, do this every <IMGFRAMERATE> MCS.
		SAVEPATH : dir,	// ... And save the image in this folder.
		EXPNAME : "ActModel",					// Used for the filename of output images.
		
		// Output stats etc
		STATSOUT : { browser: false, node: true }, // Should stats be computed?
		LOGRATE : 10							// Output stats every <LOGRATE> MCS.

	}
}
/*	---------------------------------- */

let custommethods = {
	 	initializeGrid : initializeGrid,
	 	buildChannel : buildChannel,
	 	drawBelow : drawBelow
	 }

let sim = new CPM.Simulation( config, custommethods )

function drawBelow(){
	this.Cim.drawPixelSet( this.channelvoxels, "AAAAAA" ) 
}

function initializeGrid(){
	
		// add the initializer if not already there
		if( !this.helpClasses["gm"] ){ this.addGridManipulator() }
	
		let nrcells = this.conf["NRCELLS"], cellkind, i
		this.buildChannel()
		
		// Seed the right number of cells for each cellkind
		for( cellkind = 0; cellkind < nrcells.length; cellkind ++ ){
			
			for( i = 0; i < nrcells[cellkind]; i++ ){
				// first cell always at the midpoint. Any other cells
				// randomly.				
				if( i == 0 ){
					this.gm.seedCellAt( cellkind+1, this.C.midpoint )
				} else {
					this.gm.seedCell( cellkind+1 )
				}
			}
		}
}
	
function buildChannel(){
		
	
		this.channelvoxels = []
		
		//obstacle 1 at 50,100 w radius 10
		let center = [50,100], radius = 10
		for( let xx = center[0]-radius; xx <= center[0]+radius; xx++ ){
			for( let yy = center[1]-radius; yy <= center[1]+radius; yy++){
				let dx = Math.abs( xx-center[0] ), dy = Math.abs( yy-center[1] )
				if( Math.sqrt( dx*dx + dy*dy ) < radius ){
					this.channelvoxels.push( [xx,yy] )
				}	
			}
		}
		
		// obstacle 2 at 150,100 w radius 10
		center = [150,100]
		for( let xx = center[0]-radius; xx <= center[0]+radius; xx++ ){
			for( let yy = center[1]-radius; yy <= center[1]+radius; yy++){
				let dx = Math.abs( xx-center[0] ), dy = Math.abs( yy-center[1] )
				if( Math.sqrt( dx*dx + dy*dy ) < radius ){
					this.channelvoxels.push( [xx,yy] )
				}	
			}
		}
		
		
		this.C.add( new CPM.BorderConstraint({
			BARRIER_VOXELS : this.channelvoxels
		}) )
		
}


// No custom methods.


sim.run()
```
