## Supplementary material for "Artistoo: build, share, and explore simulations of cells and tissues in the web browser": run-CellDivision.js

```
let fs = require('fs')
let dir = "output"

if (!fs.existsSync(dir)){
    fs.mkdirSync(dir);
}

let CPM = require("../build/artistoo-cjs.js")


/*	----------------------------------
	CONFIGURATION SETTINGS
	----------------------------------
*/
let config = {

	// Grid settings
	ndim : 2,
	field_size : [500,500],
	
	// CPM parameters and configuration
	conf : {
		// Basic CPM parameters
		torus : [false,false],				// Should the grid have linked borders?
		T : 3,								// CPM temperature
		
		// Constraint parameters. 
		// Mostly these have the format of an array in which each element specifies the
		// parameter value for one of the cellkinds on the grid.
		// First value is always cellkind 0 (the background) and is often not used.

		
		// Adhesion parameters:
		J : [ 
		 	[0,0],
			[0,-4]
		],
		
		// VolumeConstraint parameters
		LAMBDA_V : [0,1],				// VolumeConstraint importance per cellkind
		V : [0,500],					// Target volume of each cellkind
		
		LAMBDA_P : [0,1],
		P : [0,240]
		
	},
	
	// Simulation setup and configuration: this controls stuff like grid initialization,
	// runtime, and what the output should look like.
	simsettings : {
	
		// Cells on the grid
		NRCELLS : [1],						// Number of cells to seed for all
											// non-background cellkinds.
		// Runtime etc
		BURNIN : 0,
		RUNTIME : 40001,
		RUNTIME_BROWSER : "Inf",
		
		// Visualization
		CANVASCOLOR : "EEEEEE",
		CELLCOLOR : ["000000"],
		SHOWBORDERS : [true],				// Should cellborders be displayed?
		BORDERCOL : ["FFFFFF"],				// color of the cell borders
		zoom : 1,							// zoom in on canvas with this factor.
		
		// Output images
		SAVEIMG : true,						// Should a png image of the grid be saved
											// during the simulation?
		IMGFRAMERATE : 1000,					// If so, do this every <IMGFRAMERATE> MCS.
		SAVEPATH : dir,	// ... And save the image in this folder.
		EXPNAME : "CellDivision",					// Used for the filename of output images.
		
		// Output stats etc
		STATSOUT : { browser: false, node: true }, // Should stats be computed?
		LOGRATE : 100							// Output stats every <LOGRATE> MCS.

	}
}
/*	---------------------------------- */


/* 	The following functions are defined below and will be added to
	the simulation object. If Custom-methods above is set to false,
	this object is ignored and not used in the html/node files. */
let custommethods = {
	postMCSListener : postMCSListener,
	initializeGrid : initializeGrid,
	logStats : logStats
}
let sim = new CPM.Simulation( config, custommethods )

function logStats(){
	
		console.log( this.time + "\t" + ( this.C.t2k.length - 1) )

}


/* The following custom methods will be added to the simulation object*/
function postMCSListener(){
	
		// add the initializer if not already there
		if( !this.helpClasses["gm"] ){ this.addGridManipulator() }
	
		//prolif rate 
		let prate = 5e-5
		let interval = 10
		if( this.time % interval == 0 ){
			for( let i of this.C.cellIDs() ){
				if( this.C.getVolume(i) > this.C.conf.V[1]*0.95 && this.C.random() < prate*interval ){
					this.gm.divideCell(i)
					this.C.stats=[]
				}
			}
		}
		
		
		
}
	
function initializeGrid(){
	
		// add the GridManipulator if not already there and if you need it
		if( !this.helpClasses["gm"] ){ this.addGridManipulator() }
	
		this.gm.seedCellsInCircle( 1, 20, this.C.midpoint, 35 )

}


sim.run()
```
