## Supplementary material for "Artistoo: build, share, and explore simulations of cells and tissues in the web browser": run-CellSorting-varN.js

```
let nCells = parseInt( process.argv[2] )

let CPM = require("../build/artistoo-cjs.js")

const cellVolume = 200

// Compute gridSize and seeding radius from nCells
const totalPixels = nCells*cellVolume*2 // two cell types
const gridSize = Math.ceil( Math.sqrt( totalPixels*1.5 ) )
const seedRadius = Math.ceil( 0.8* Math.sqrt( totalPixels/ Math.PI ) )


/*	----------------------------------
	CONFIGURATION SETTINGS
	----------------------------------
*/
let config = {

	// Grid settings
	ndim : 2,
	field_size : [gridSize,gridSize],
	
	// CPM parameters and configuration
	conf : {
		// Basic CPM parameters
		torus : [true,true],						// Should the grid have linked borders?
		seed : 1,							// Seed for random number generation.
		T : 15, //15,								// CPM temperature

		// Constraint parameters. 
		// Mostly these have the format of an array in which each element specifies the
		// parameter value for one of the cellkinds on the grid.
		// First value is always cellkind 0 (the background) and is often not used.
		
		// Adhesion parameters:
		J : [ [NaN, 12, 6], [12, 6, 16], [6, 16, 6] ],
		
		// VolumeConstraint parameters
			// VolumeConstraint importance per cellkind
			// Target volume of each cellkind
		LAMBDA_V : [0,1,1], //2 2
		V : [0,cellVolume,cellVolume]		//25 25

		
	},
	
	// Simulation setup and configuration
	simsettings : {
	
		// Cells on the grid
		NRCELLS : [1,1],					// Number of cells to seed for all
											// non-background cellkinds.
	
		// Runtime etc
		BURNIN : 500,
		RUNTIME : 2000,
		RUNTIME_BROWSER : 20000,
		
		// Visualization
		CANVASCOLOR : "eaecef",
		CELLCOLOR : ["000000","FF0000"],
		ACTCOLOR : [true,false],			// Should pixel activity values be displayed?
		SHOWBORDERS : [true,true],				// Should cellborders be displayed?
		zoom : 1,							// zoom in on canvas with this factor.
		
		// Output images
		SAVEIMG : false,					// Should a png image of the grid be saved
											// during the simulation?
		IMGFRAMERATE : 100,					// If so, do this every <IMGFRAMERATE> MCS.
		SAVEPATH : "output/img/CellSorting-varN-"+nCells,				// ... And save the image in this folder.
		EXPNAME : "CellSorting-varN-"+nCells,					// Used for the filename of output images.
		
		// Output stats etc
		STATSOUT : { browser: false, node: false }, // Should stats be computed?
		LOGRATE : 10							// Output stats every <LOGRATE> MCS.

	}
}
/*	---------------------------------- */


/* 	The following functions are defined below and will be added to
	the simulation object.*/
let custommethods = {
	initializeGrid : initializeGrid
}
let sim = new CPM.Simulation( config, custommethods )


/* The following custom methods will be added to the simulation object
below. */
function initializeGrid(){
	
		// add the GridManipulator if not already there and if you need it
		if( !this.helpClasses["gm"] ){ this.addGridManipulator() }
	
		this.gm.seedCellsInCircle( 1, nCells, this.C.midpoint, seedRadius )
		this.gm.seedCellsInCircle( 2, nCells, this.C.midpoint, seedRadius )

}


sim.run()
```
