## Supplementary material for "Artistoo: build, share, and explore simulations of cells and tissues in the web browser": run-GameOfLife.js

```
let fs = require('fs')
let dir = "output"

if (!fs.existsSync(dir)){
    fs.mkdirSync(dir);
}

let CPM = require("../build/artistoo-cjs.js")


/* */

"use strict"


// Configuration
let conf = {
	gsize : [50,50],		// dimensions of the grid to build
	zoom : 3,				// zoom for displaying the grid
	torus: [true,true],			// Should grid boundaries be connected?
	runtime : 500
}

let C, Cim, cid=0, meter, t = 0


// Setup the grid and needed objects
function setup(){
		C = new CPM.CA( [conf.gsize[0],conf.gsize[1]], {
		"UPDATE_RULE": 	function(p,N){
			let nalive = 0
			for( let pn of N ){
				nalive += (this.pixt(pn)==1)
			}	
			if( this.pixt(p) == 1 ){
				if( nalive == 2 || nalive == 3 ){
					return 1
				}
			} else {
				if( nalive == 3 ) return 1
			}
			return 0
		}
	})
	Cim = new CPM.Canvas( C, {zoom:conf.zoom} )
}

// Place something on the grid
function initializeGrid(){
	for( let x = 0 ; x < conf.gsize[0] ; x ++ ){
		for( let y = 0 ; y < conf.gsize[1] ; y ++ ){
			if( C.random() < 0.5 ){
				C.setpix( [x,y], 1 )
			}
		}
	}
}

// Produce output, like drawing on the canvas and logging stats
function output(){
	// Clear the canvas (in the backgroundcolor white), and redraw:
	Cim.clear( "FFFFFF" )
	// The cell in red
	Cim.drawCells( 1, "AA0000" )
	
	
	if( t % 20 == 0 ){
		let outpath = dir, expname = "GameOfLife"
		Cim.writePNG( outpath +"/" + expname + "-t"+t+".png" )
	}
}

// Run everything needed for a single step (output and computation),
// and update the current time
function step(){
	C.timeStep()
	output()
	t++
}

// Starts up the simulation
function initialize(){
	setup()
	initializeGrid()
	run()
}
// all steps
function run(){
	 while( t < conf.runtime){
			step()
	 }
}
initialize()
```
