## Supplementary material for "Artistoo: build, share, and explore simulations of cells and tissues in the web browser": ActModel.html


### Cell migration

Cellular Potts Model of a migrating cell. Adjust the max_act_ and
λ_act_ parameters using the sliders below.

|  |  |  |  |
| --- | --- | --- | --- |
| max_act_ | 0 |  | 100 |
| λ_act_ | 0 |  | 1000 |

  
