## Supplementary material for "Artistoo: build, share, and explore simulations of cells and tissues in the web browser": CellSorting.html


### Cell sorting

Cellular Potts Model of cells sorting through differential adhesion.
Adjust the different adhesion parameters using the sliders below.

|  |  |  |  |
| --- | --- | --- | --- |
| J_bg,1_ | 0 |  | 100 |
| J_bg,2_ | 0 |  | 100 |
| J_1,1_ | 0 |  | 100 |
| J_2,2_ | 0 |  | 100 |
| J_1,2_ | 0 |  | 100 |

  
