## Supplementary material for "Artistoo: build, share, and explore simulations of cells and tissues in the web browser": CollectiveMigration.html

### CollectiveMigration

start
stop
seed cell
+10 cells
+100 cells
remove cell
remove all cells
  

|  |  |  |
| --- | --- | --- |
| Adhesion_cell-matrix_ |  | Adhesion_cell-cell_ |
| Volume |  | λ_Volume_ |
| Perimeter |  | λ_P_ |
| Max_Act_ |  | λ_Act_ |
| T |  | Framerate |
