## Supplementary material for "Artistoo: build, share, and explore simulations of cells and tissues in the web browser": sliders.js

```
function initializeSliders(){
	const allRanges = document.querySelectorAll(".range-wrap");
	allRanges.forEach(wrap => {
	  const range = wrap.querySelector(".range");
	  const bubble = wrap.querySelector(".bubble");

	  range.addEventListener("input", () => {
	  	/*if( range.id == "mact" ){
	  		sim.C.getConstraint("ActivityConstraint").conf.MAX_ACT[1]=range.value
	  	} else if ( range.id == "lact" ){
	  		sim.C.getConstraint("ActivityConstraint").conf.LAMBDA_ACT[1]=range.value
	  	}*/
		setBubble(range, bubble);
	  });
	  setBubble(range, bubble);
	});
	

}

function setBubble(range, bubble) {
  const val = range.value;
  const min = range.min ? range.min : 0;
  const max = range.max ? range.max : 100;
  const newVal = Number(((val - min) * 100) / (max - min));
  bubble.innerHTML = val;

  // Sorta magic numbers based on size of the native UI thumb
  bubble.style.left = `calc(${newVal}% + (${8 - newVal * 0.15}px))`;
}
```
