## Supplementary material for "Artistoo: build, share, and explore simulations of cells and tissues in the web browser": Makefile

```
all : supplement.pdf

latex : 
	mkdir -p $@


supplement.pdf : latex/supplement.pdf
	cp $< $@
	
latex/supplement.pdf : latex/supplement.tex
	cd latex && pdflatex supplement.tex
	
latex/supplement.tex : supplement.tex | latex
	cp $< $@
```
