## Supplementary material for "Artistoo: build, share, and explore simulations of cells and tissues in the web browser": supplement.pdf

May 1, 2020

### Overview of materials

This directory contains the following supplementary materials:

- `build/` folder: the Artistoo source code required for the interactive HTML pages and node scripts;
- `code/` folder: node scripts of Artistoo simulations used to assess performance, and xml files of the corresponding Morpheus simulations;
- `interactive-html/` folder: examples of interactive simulations;
- `applications/` folder: example applications

See the corresponding sections below for details.

### Code

The `code/` folder contains the code used to assess Artistoo performance. Before you can run these scripts, you must ensure that nodejs is installed on your computer, and run `npm install` in the base folder to install the required node modules. (Note that while no software installation is required for browser simulations, the console simulations rely on nodejs and require users to install node and its packages).

### Interactive web pages

The `interactive-html/` folder contains the several interactive web pages.

`ActModel.html` contains the simulation shown in panel B of the main figure. `CellMigration.html`, `CellDivision.html`, `GameOfLife.html`, and `CellSorting.html` are interactive versions of the simulations for which performance was tested in panel C. `CollectiveMigration.html` is the interactive simulation we use for education, see also the section *Example applications*.

Please note that the interactive web pages load files from `./build/` and from `./interactive-html/src/`. An isolated HTML file copied to some other folder will therefore not work; please copy the entire directory or update the paths in the HTML file.

### Example applications

#### Teaching

We frequently use Artistoo for teaching workshops on the CPM. The file `applications/teaching-exercise.pdf` contains an exercise we use for beginning CPM users, which readers are free to use in their own education. The file refers to an online simulation, but this simulation can also be found under `interactive-html/CollectiveMigration.html`.

### Slides with interactive simulation

The revealjs framework allows users to build slidesets in HTML. These slidesets can also contain interactive Artistoo simulations. See `applications/slides- example/index.html`: open it in your web browser to see the slides. Note that the simulations here use the artistoo build in the `build/` folder, so if you move the presentation elsewhere you will also have to change the links in the HTML files in `applications/slides-example/simulations/`.

### Poster website

Interactive HTML simulations made with Artistoo can also be shared on a poster, by making a small web page and sharing this via a QR code on the poster. An example of such a website can be found at:
`https://computational-immunology.org/inge/poster-cpmjs/`

### Interactive parameter tuning

CPM parameter tuning can be easier in an interactive web page where effects of changes are visible immediately (as are artefacts such as cell breaking). An example of such a page is the `interactive-html/CollectiveMigration.html` page we also use for teaching.
