## Supplementary material for "Artistoo: build, share, and explore simulations of cells and tissues in the web browser": supplement.tex

\documentclass{article}

\usepackage{url}
\usepackage{a4wide}
%\usepackage{hyperref}

\title{Supplementary Materials}

\author{Inge Wortel and Johannes Textor}


\begin{document}
\maketitle

\section*{Overview of materials}

This directory contains the following supplementary materials:

\begin{itemize}
	\item \texttt{build/} folder: the Artistoo source code required for the interactive HTML pages
	and node scripts;
	\item \texttt{code/} folder: node scripts of Artistoo simulations used to assess
	performance, and xml files of the corresponding Morpheus simulations;
	\item \texttt{interactive-html/} folder: examples of interactive simulations;
	\item \texttt{applications/} folder: example applications
\end{itemize}

See the corresponding sections below for details.


\section*{Code}

The \texttt{code/} folder contains the code used to assess Artistoo performance. Before you can run
these scripts, you must ensure that nodejs is installed on your computer, and 
run \texttt{npm install} in the base folder to install the required node modules. 
(Note that while no software installation is required for browser simulations, the 
console simulations rely on nodejs and require users to install node and its packages).


\section*{Interactive web pages}

The \texttt{interactive-html/} folder contains the several interactive web pages. 

\texttt{ActModel.html} contains
the simulation shown in panel B of the main figure. \texttt{CellMigration.html},
\texttt{CellDivision.html}, \texttt{GameOfLife.html}, and \texttt{CellSorting.html} are
interactive versions of the simulations for which performance was tested in panel C.
\texttt{CollectiveMigration.html} is the interactive simulation we use for education,
see also the section \textit{Example applications}.

Please note that the interactive web pages load files from \texttt{./build/} and from
\texttt{./interactive-html/src/}. An isolated HTML file copied to some other folder
will therefore not work; please copy the entire directory or update the paths in the
HTML file.


\section*{Example applications}

\subsection*{Teaching}
We frequently use Artistoo for teaching workshops on the CPM. The file \newline
\texttt{applications/teaching-exercise.pdf} contains an exercise we use for beginning
CPM users, which readers are free to use in their own education. The file refers to an
online simulation, but this simulation can also be found under 
\texttt{interactive-html/CollectiveMigration.html}.


\subsection*{Slides with interactive simulation}
The revealjs framework allows users to build slidesets in HTML. These slidesets can also
contain interactive Artistoo simulations. See \texttt{applications/slides- example/index.html}:
open it in your web browser to see the slides. Note that the simulations here use the
artistoo build in the \texttt{build/} folder, so if you move the presentation elsewhere
you will also have to change the links in the HTML files in \texttt{applications/slides-example/simulations/}.


\subsection*{Poster website}
Interactive HTML simulations made with Artistoo can also be shared on a poster,
by making a small web page and sharing this via a QR code on the poster.
An example of such a website can be found at: \newline 
\url{https://computational-immunology.org/inge/poster-cpmjs/}


\subsection*{Interactive parameter tuning}
CPM parameter tuning can be easier in an interactive web page where effects of changes
are visible immediately (as are artefacts such as cell breaking). An example of such 
a page is the \texttt{interactive-html/CollectiveMigration.html} page we also use for
teaching. 


\end{document}
