## Supplementary figures and images for "Artistoo: build, share, and explore simulations of cells and tissues in the web browser"

### nijmegen.png

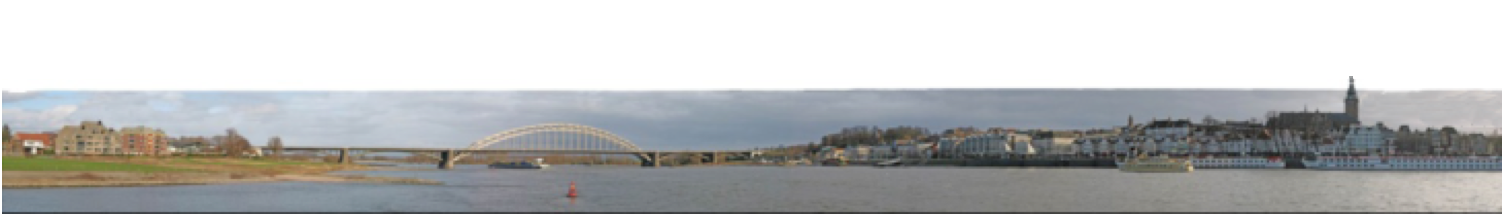

### Radboudumc_ENGELS_700px_RGB.jpg

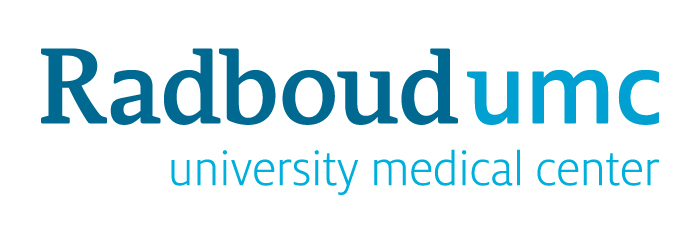

### Radboudumc_ENGELS_700px_RGB.png

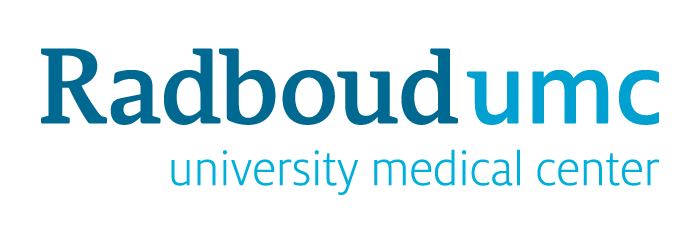

### ru-rumc.jpg

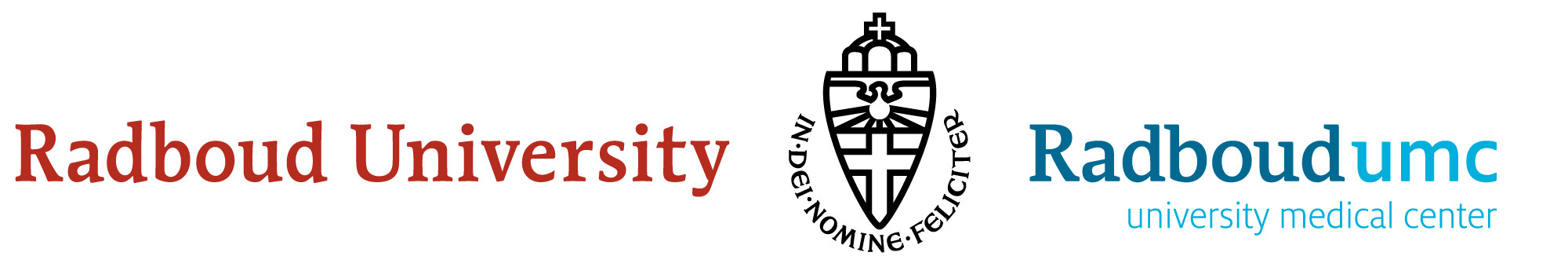
